# Reference genome for the WHO reference strain for *Mycobacterium bovis* BCG Danish, the present tuberculosis vaccine

**DOI:** 10.1101/513408

**Authors:** Katlyn Borgers, Jheng-Yang Ou, Po-Xing Zheng, Petra Tiels, Annelies Van Hecke, Evelyn Plets, Gitte Michielsen, Nele Festjens, Nico Callewaert, Yao-Cheng Lin

## Abstract

*Mycobacterium bovis* bacillus Calmette-Guérin (*M. bovis* BCG) is the only vaccine available against tuberculosis (TB). This study reports on an integrated genome analysis workflow for BCG, resulting in the completely assembled genome sequence of BCG Danish 1331 (07/270), one of the WHO reference strains for BCG vaccines. We demonstrate how this analysis workflow enables the resolution of genome duplications and of the genome of engineered derivatives of this vaccine strain.

---

The BCG live attenuated TB vaccine is one of the oldest and most widely used vaccines in human medicine. Each year, BCG vaccines are administered to over 100 million newborns (i.e. 75% of all newborns on the planet). The original BCG strain was developed in 1921 at the Pasteur Institute, through attenuation of the bovine TB pathogen *M. bovis*, by 231 serial passages on potato slices soaked in glycerol-ox bile over a time-span of 13 years^1^. This BCG Pasteur strain was subsequently distributed to laboratories around the world and different laboratories maintained their own daughter strains by passaging. Over the years, different substrains arose with different protective efficacy^2, 3^. The establishment of a frozen seed-lot system in 1956 and the WHO recommendation of 1966 that vaccines should not be prepared from cultures that had undergone >12 passages starting from a defined freeze-dried seed lot, halted the accumulation of additional genetic changes^1^. In an effort to further standardize the vaccine production and to prevent severe adverse reactions related to BCG vaccination, three substrains, i.e. Danish 1331, Tokyo 172-1 and Russian BCG-1 were established as the WHO reference strains in 2009 and 2010^4^. Of these, the BCG Danish 1331 strain is the most frequently used one, and it also serves as a basis of most current “next-generation” engineering efforts to improve the BCG vaccine or to use it as a “carrier” for antigens of other pathogens^5, 6^. Complete genome elucidation of BCG strains is challenging by the occurrence of large genome segment duplications and a high GC content. Therefore, no fully assembled reference genome is yet available for BCG Danish, only incomplete ones^7, 8^, which hinders further standardization efforts.

By combining second (Illumina) and third (PacBio) generation sequencing technologies and an integrated bioinformatics workflow we have for the first time fully assembled the BCG Danish 1331 (07/270) strain genome sequence. Ambiguous regions were locally reassembled and/or experimentally verified. The single circular chromosome is 4,411,814 bp in length and encodes 4,084 genes, including 4,004 genes encoding for proteins, 5 genes for rRNA, 45 genes for tRNA and 30 pseudogenes (Fig. 1a). Compared to the reference genome sequence of BCG Pasteur 1173P2, 42 SNPs were identified and a selected subset was validated (Suppl. Table 1 and 5). Genetic features determinative for BCG Danish, as described by Abdallah *et al*.^8^, were identified, including the region of difference (RD) Denmark/Glaxo and the DU2 type III, that was completely resolved in the assembly (Fig. 1a-b). Additionally, a 1 bp deletion in Mb3865 and a 465 bp insertion in PE_PGRS54 compared to BCG Pasteur were found. The organization of 2 repeats (A and B) in PE_PGRS54 has been reported to differ between the BCG strains^9^. We report a A-A-B-B-B-B organization for BCG Danish in contrast to BCG Tokyo (A-A-B-B-B) and BCG Pasteur (A-B-B-B-B). Previously, two separate genetic populations for BCG Danish 1331 have been described, which differ in the SenX3-RegX3 region (having 2 or 3 repeats of 77 bp)^10^. For BCG Danish 1331 07/270 we document only 3 repeats of 77 bp (Suppl. Fig. 2). Two features described by Abdallah *et al*.^8^ to be determinative for BCG Danish were not identified, namely the rearrangement of the *fadD26-pssA* gene region and a 894 bp del in Mb0096c-Mb0098c. In addition, a 399 bp instead of 118 bp insertion was detected in *leuA*, giving 12 direct repeats of 57 bp, as in the Pasteur strain (previously denoted as S-RD13^11^). These differences are likely due to inherent repeat structures in these regions, which cannot be resolved by short sequencing reads (as used in Abdallah *et al*.^8^), but require long sequencing reads, as generated by PacBio SMRT sequencing in this study.

**Figure 1.**
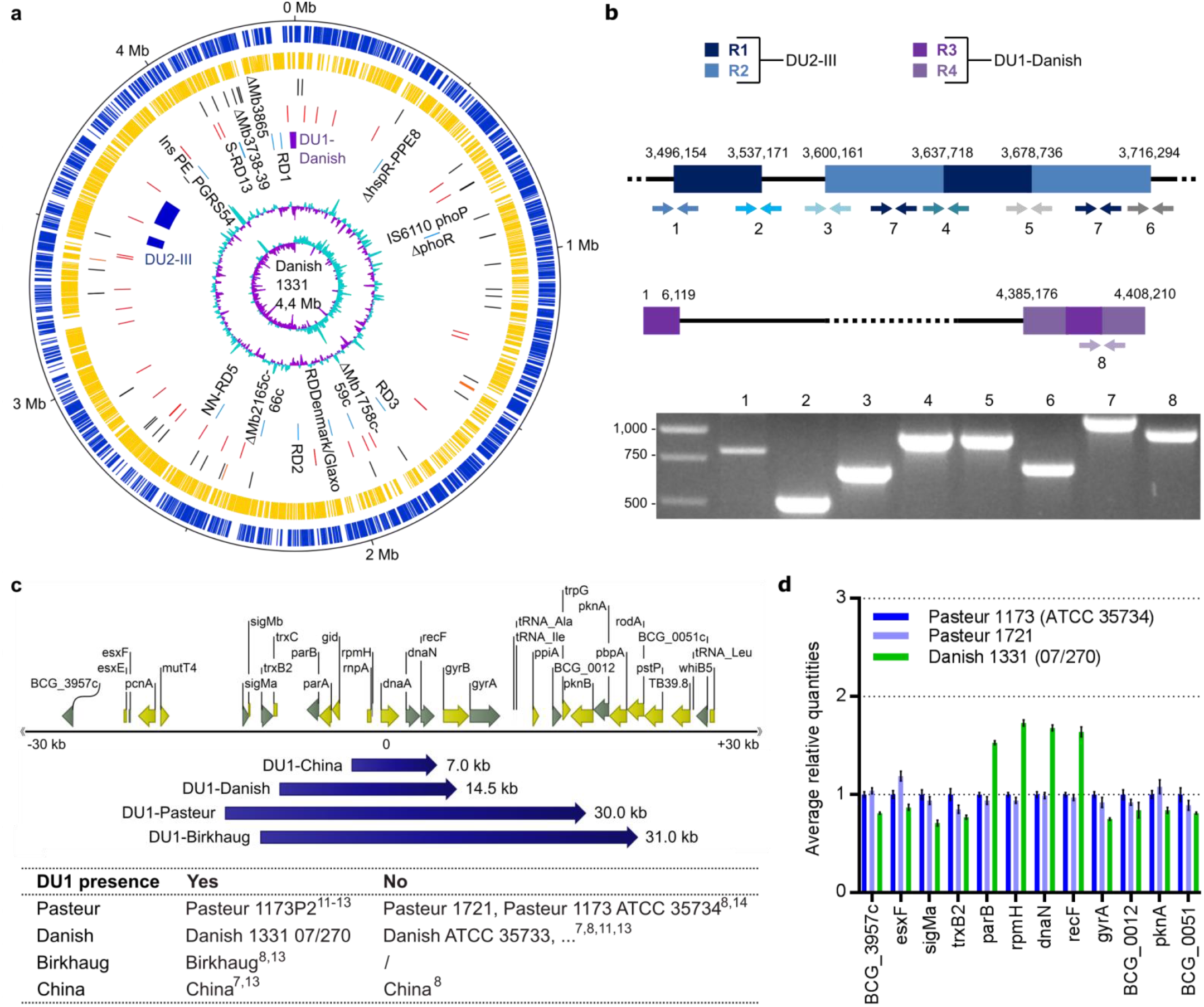
Organization of the BCG Danish 1331 (07/270) genome, focusing on the DU1 and DU2. **a)** Circular representation of the BCG Danish chromosome. The scale is shown in megabases on the outer black circle. Moving inward, the next two circles show forward (dark blue) and reverse (yellow) strand CDS. The next circle shows tRNA (black) and rRNA (orange) genes, followed by 42 SNPs (red) detected between BCG Danish and Pasteur. The subsequent circle shows DU2-III (dark blue), DU1-Danish (purple) and RD (light blue, names of RD in black) that are typical for BCG Danish. The two inner circles represent G+C content and GC skew. **b)** Organization of the two tandem duplications in BCG Danish and confirmation by PCR. The DU2 is made up by two repeats (R1 and R2), as well as the DU1-Danish (R3 and R4). Used primer pairs (1-8) to validate their organization are indicated. **c)** Visual representation of the *oriC* with position and size of DU1-China, -Danish, -Pasteur and -Birkhaug. The table indicates which substrains have the DU1. **d)** Copy-number analysis of genes (indicated in grey in **subfigure c)** in and surrounding the DU1 region for Pasteur 1173 (ATCC 35734), Pasteur 1721 and Danish 1331 (07/270). The represented data are averages (± SD) of four technical replicates.

Two large tandem chromosomal duplications characterize the BCG strains; the DU2 and DU1 (Fig. 2, **Suppl. Table 6**). While four different forms of the DU2 exist, the DU1 is supposed to be exclusively present in BCG Pasteur^11–13^; it spans the chromosomal origin of replication or *oriC* (*dnaA-dnaN* region) and encodes key components of the replication initiation and cell division machinery. Surprisingly, we detected a DU1-like duplication of 14,577 bp in BCG Danish (Fig. 1). To adapt an unambiguous terminology, we considered all duplications spanning the *oriC* as DU1, while specifying the strain in which the duplication was found. Investigation of other publicly available data for BCG Danish did not show presence of a DU1 (Fig. 1c, Suppl. Fig. 1), indicating that only the Danish 1331 substrain deposited as the WHO reference at the National Institute for Biological Standards and Control (NIBSC) contains this duplication. Additional inconsistencies in DU1 presence/absence were detected by reanalyzing publicly available data (Fig. 1c, Suppl. Fig. 1). In contrast to the literature, we detected BCG Pasteur substrains with a DU1 (data ref^13^) and without a DU1 (data ref^8, 14^). Similarly, experimental analysis of our in-house Pasteur strains (1721, 1173 ATCC 35734) showed absence of a DU1 (Fig. 1d). Additionally, a DU1-China was detected (data ref^7, 13^), but not in the data of Abdallah *et al.* 2015^8^, which could be explained by the use of two different substrains of BCG that are both named BCG China^8^. DU1-Birkhaug was consistently detected in all reported sequencing data of that BCG strain. The genealogy of BCG strains is thus further complicated by the genomic instability of the *oriC* during *in vitro* cultivation (Fig. 2, **Suppl. Table 6**). A DU1-like duplication has also been identified in a ‘non-vaccine’ strain; in a clinical isolate (3281), identified as BCG, a 7-kb region that covered six genes and crossed the *oriC* was repeated three times^15^, further indicating that this region is prone to duplication. Together, these data underline the importance of the genomic characterization of BCG strains used as vaccines, including their dynamic duplications.

**Figure 2.**
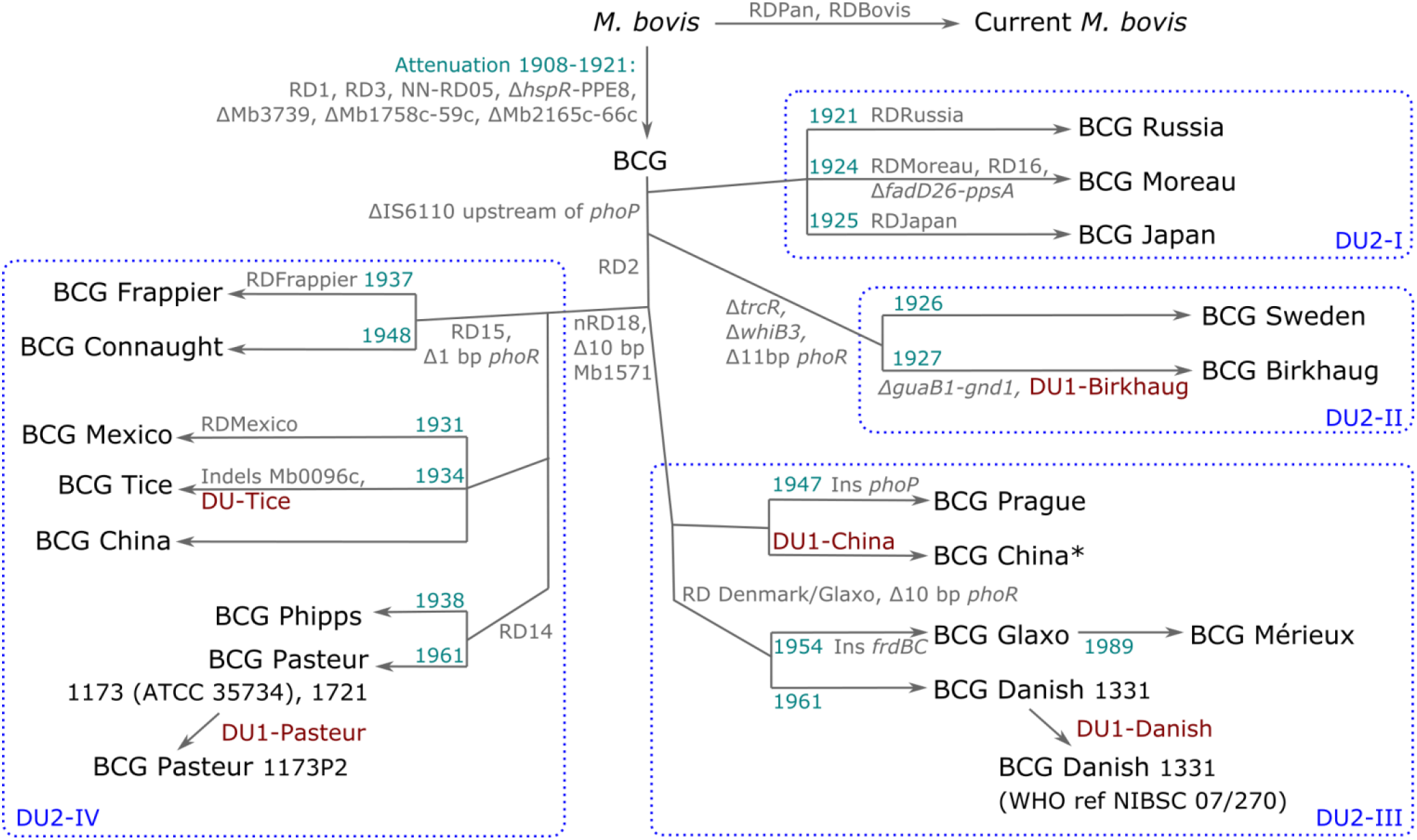
Refined genealogy of BCG vaccine strains. The year when the strain was obtained per geographical location is indicated where possible (indigo). The scheme shows regions of difference (RD), insertions (Ins), deletions (‘Δ’), indels, tandem duplications (DU), which differentiate the different BCG strains (**Suppl. Table 6**). The blue dashed squares indicate the different DU2-forms, which classify the BCG strains into four major lineages. When the DU1 is not found in all substrains of a certain strain, this is indicated on the scheme. According to the literature, two different substrains of BCG are named BCG China or Beijing^8^. Therefore, the scheme contains two ‘BCG China’ strains: BCG China^8^ and BCG China*^7, 13^. Adapted from refs ^8, 11, 13, 16, 17^ Concerning ref ^8^, only the RD and deleted genes that could be verified on the assembled genomes are included.

To demonstrate how this genome analysis methodology contributes to full characterization of improved BCG-derived engineered vaccines, we applied it to a knock-out mutant (KO) for the *sapM* secreted acid phosphatase, located in the analytically challenging long duplication region DU2^11^. Our BCG genome analysis workflow unequivocally demonstrated that the KO engineering had inadvertently out-recombined one of the copies of this DU2 and had given rise to a single SNP (Suppl. Fig. 4, Suppl. Table 1 and 5). Such unexpected genomic alterations are likely common in engineered live attenuated TB vaccines, but have so far gone unnoticed due to lack of a complete reference genome and or suitable genome analysis methodology.

The implementation of both short (Illumina) and long (PacBio) sequencing reads in one genome analysis methodology allows the straightforward generation of completely assembled genomes of BCG strains. These include the decomposition of the analytically challenging long duplications regions DU1 and DU2, wherefore one formerly needed additional experimental methods (e.g. gap closure via PCR^15^).

The availability of the complete reference genome for BCG Danish 1331 as well as the associated genome analysis workflow, now permits full genomic characterization of (engineered) TB vaccine strains, which should contribute to more consistent manufacturing of this highly cost-effective vaccine that protects the world’s newborns from disseminated TB.

## METHODS

### Mycobacterial strains, gDNA and reference genomes

The strains used include the *M. bovis* BCG Danish 1331 sub-strain (1^st^ WHO Reference Reagent, 07/270, National Institute for Biological Standards and Control (NIBSC), Hertfordshire), the BCG Pasteur 1173 strain (ATCC®35734™, ATCC, Manassas), the streptomycin-resistant BCG Pasteur 1721 strain^18^ (*RpsL*: K43R; a gift of Dr. P. Sander, Institute for Medical Microbiology, Zürich). From the Danish 1331 strain a *ΔsapM* (KO) strain was constructed (detailed procedure of the strain construction can be found in **Suppl. Methods**). Strains were grown in Middlebrook 7H9 broth (Difco) supplemented with 0.05% Tween-80 and Middlebrook OADC (Becton Dickinson). Preparation of gDNA from mycobacterial strains was performed as previously described^19^. As reference genomes, *M. tb* H37Rv (NC_000962.3), *M. bovis* AF2122_97 (NC_002945.4) and BCG Pasteur 1173P2 (NC_008769.1) were used.

### Whole genome sequencing of BCG Danish 1331 WT and *sapM* KO strain

For PacBio SMRT sequencing, the gDNA was sheared (large hydropore, Megaruptor device, Diagenode), used for a PacBio SMRT library prep (SMRTbell Temp Prep Kit 1.0, Pacific Biosciences) and sequenced on an PacBio RSII instrument (DNA/Polymerase Binding Kit P6 v2, Pacific Biosciences). 1 SMRT-cell was run for the KO sample (229x coverage) and 2 SMRT-cells were run for the WT sample (140x and 95x coverage). For Illumina sequencing, libraries were prepared with Nextera DNA Library Preparation kit and sequenced on an Illumina MiSeq instrument (MiSeq Reagent Kit v2 Nano, PE250, 500 Mb), with an average of 55-56x coverage per genome.

### Genome assembly and analysis (Suppl. Fig. 3)

Illumina reads were quality-filtered and trimmed (Trimmomatic-0.36) after which the paired-end reads were merged with overlapping sequence (BBmerged v36.69). PacBio reads were corrected using the trimmed Illumina reads (Lordec v0.6). The unmerged and merged Illumina reads were assembled (SPAdes v3.9.0) into a draft assembly, which was scaffolded using the corrected PacBio reads (SSPACE-LongRead v3.0). Gaps in the scaffold were closed (GapFiller v1.10) and finally the assembly was improved using the trimmed Illumina reads (Pilon v1.20).

The exact sequence of the DU1 region was based on a second round of local *de novo* assembly (SPAdes v3.9.0) using soft-clipped Illumina reads surrounding the draft DU1 region where the Illumina read coverage is more than two times higher than the background coverage. The DU2 repeat was resolved by comparing SPAdes assembly with the assembly from HINGE (v201705), where the R1 and R2 regions have been separated. The junction sequences of DU1 and DU2 were further confirmed by aligning uniquely mapped PacBio reads, or by PCR and Sanger sequencing.

Annotation was done by combining an automatic gene prediction program with heuristic models (GeneMark.hmm) and the existing Pasteur and *M. tb* reference gene models (GMAP and TBLASTN) along with UniProt database (BLASTP). Non-coding RNA were predicted (tRNAScan-SE and Infernal). The assigned annotations were manually checked (Artemis and CLC Main Workbench 8, e.g. correct start codon), by comparative analysis with the 3 reference genomes for *M. tb, M. bovis* and Pasteur, as listed above. Inconsistencies in the annotation and/or assembly were analyzed in detail and/or verified by PCR and Sanger Sequencing.

A probabilistic variant analysis was performed by mapping the BBmerged Illumina reads to the Pasteur reference genome (BWA-MEM) and calling variants by GATK UnifiedGenotyper (Count ≥ 10 & Variant Probability > 0.9), whereafter variant annotations and functional effect prediction were carried out with SnpEff and SnpSift. The orthologous relationships between *M. tb*, BCG Pasteur and BCG Danish WT and *sapM* KO were investigated: the proteins of strains (BCG Danish WT and *sapM* KO, BCG Pasteur 1173P2, H37Rv) were searched using all-against-all with BLASTP, after which the result was analyzed by TribeMCL and i-ADHoRe 3.0 based on the genome synteny information (**Suppl. Table 7**).

To validate the detection of the DU1, the DU1 duplication region was reanalyzed with published genome data from ref ^7, 8, 13, 14^. Illumina short sequencing reads or probes on tiling array were mapped to the *M. tb* reference strain (BWA-MEM) after which the DU1 duplications were detected (cn.mops).

In Suppl. Fig. 3 a graphical overview of the performed genome analysis pipeline is given and in **Supplementary Methods** a citation list was incorporated for the used bioinformatics tools and databases.

### PCR analysis, gel electrophoresis and Sanger Sequencing

PCR (GoTaq®Green, Promega) was performed on gDNA using primers listed in Suppl. Table 2. PCR products were run on a 1.2% agarose gel, stained with Midori Green and visualized under ultraviolet light. To confirm the SNP variants, regions of interest were amplified (Phusion High-Fidelity DNA Polymerase, NEB) from gDNA with primers listed in Suppl. Table 3. The resulting PCR products were purified (AMPure XP beads) and Sanger sequenced with (a) nested primer(s) (Suppl. Table 3).

### Copy number profiling via qPCR

Real-time quantitative PCR was done on a LightCycler 480 (Roche Diagnostics) using the SensiFast SYBR-NoRox kit (Bioline) in quadruplicate for each gDNA sample using primers listed in Suppl. Table 4. Determination of the average relative quantities was performed using the qbasePLUS software (Biogazelle). All results were normalized using the reference genes 16S rRNA, *nuoG* and *mptpB.*

## Supporting information

Supplemental Table 7 Worksheet 1

Supplemental Table 7 Worksheet 2

## ACKNOWLEDGEMENTS

We thank Dr. Peter Sander (Institute for Medical Microbiology, Faculty of Medicine, University of Zurich) for providing us with the BCG strain 1721. We thank the DRESDEN-concept Genome Center (MPI-CBG, Dresden) for the PacBio library prep and sequencing services. We thank Insight Genomics Inc. (Tainan, Taiwan) for the Illumina library prep and sequencing services. We thank the VIB Genetics Service Facility, (http://www.vibgeneticservicefacility.be, Antwerp) for the Sanger sequencing services. Research was funded through a PhD fellowship to K.B. from the Flanders Innovation & Entrepreneurship agency (VLAIO), an ERC Consolidator grant ‘GlycoTarget’ to N.C., VIB and UGhent institutional funding to N.C., and Academia Sinica institutional funding to Y.-C.L.

The authors declare no competing interests.

## AUTHOR CONTRIBUTIONS

K.B. designed and performed the experiments, analyzed the data and co-wrote the manuscript. J.-Y.O performed the bioinformatics analysis. P.-X.Z. assisted in the data analysis. G.M. assisted in performing the experiments. E.P. performed the RT-qPCR analysis. P.T. and A.H. constructed pGoal17SapM700 and transformed BCG Danish 1331 with this plasmid. N.F. assisted in experimental interpretation and carefully revised the manuscript. N.C. initiated the project, assisted in experimental design and interpretation and co-wrote the manuscript. Y.-C.L. performed the bioinformatics analysis, assisted in experimental design and interpretation and co-wrote the manuscript.

## DATA AVAILABILITY STATEMENT

The raw sequencing data (raw Illumina and PacBio reads, and PacBio base modification files) generated by this study for the BCG Danish 1331 WT and *sapM* KO strain, the complete genome assemblies and annotation have been deposited in GenBank with the primary accession codes BioProject PRJNA494982. The data (other than the next-generation sequencing data) that support the findings of this study are available on request from the corresponding author N.C.

## SUPPLEMENTARY DATA

Supplementary Methods

Supplementary Figure 1. DU1 duplication detection in BCG strains

Supplementary Figure 2. Analysis of the SenX3-RegX3 region in BCG strains

Supplementary Figure 3. Genome analysis pipeline

Supplementary Figure 4. Generation and characterization of BCG Danish 1331 *sapM* KO

Supplementary Table 1. Summary table of SNPs detected in *M. bovis* Danish 1331 WT and *sapM* KO compared to the Pasteur reference 1173P2 (NC_008769.1)

Supplementary Table 2. PCR primer pairs for confirmation of the genome assembly

Supplementary Table 3. PCR primer pairs for confirmation of the SNP variants

Supplementary Table 4. qPCR primer pairs for copy number profiling

Supplementary Table 5*. SNPs detected in *M. bovis* Danish 1331 WT and *sapM* KO compared to the Pasteur reference 1173P2 (NC_008769.1)

Supplementary Table 6*. Distribution of regions of difference, deletions and tandem duplications (DU1 and DU2) in the different BCG strains compared to *M. bovis*

Supplementary Table 7*. Ortholog of mycobacterial genes between *M. tb* H37Rv, *M. bovis* BCG Pasteur, M. bovis BCG Danish WT and *sapM* KO.

* Supplementary Table 5 to 7 are provided as Excel files.

## Supplementary Methods

### Generation of *M. bovis* BCG Danish 1331 *sapM* KO

A *sapM* KO construct was made with the p2NIL and pGOAL17 vectors^20^. Hereto, a 5’ and 3’ sequence part of *sapM* of 700 bp was amplified by PCR from gDNA of *M. bovis* BCG (5’ Fw primer: ctgcagggctggtgggtttgctcgtcg, 5’ Rv primer: attaccctgttatccctacggcgaacgcctgggccatc, 3’ Fw primer: tagggataacagggtaattagccgccgtcgctattctgtg, 3’ Rv primer aagcttctcgtcgtcggactcggccg). In the 5’ Rv primer a stop codon was inserted to ensure that a truncated *sapM* is formed. The 5’ and 3’ part were fused by performing a PCR with the 5’ Fw primer and 3’ Rv primer. The resulting fragment was cut with PstI and HindIII and ligated in the p2NIL vector cut with the same restriction enzymes. This p2NILSapM700 vector was then cloned into the PacI site of the pGOAL17 vector to create the pGOAL17SapM700 vector (Suppl. Fig. 4a).

For the electroporation, *M. bovis* BCG Danish 1331 was grown to mid-log phase (OD_600_ 0.4-0.8) in 7H9-ADS-Tw medium (7H9 + 50 g/L bovine serum albumin fraction V, 20 g/L dextrose, 8.5 g/L NaCl, 0.05% Tween-80). 1.5% glycine was added into the culture the day before electroporation. On the day of electroporation, cells were harvested in 50 ml conical tubes at room temperature at 3700 rpm for 10 minutes. The cells were washed twice with 50 ml 0.05% Tween-80 (pre-warmed at 37°C), after which the cells were resuspended in 1 ml of 0.05% Tween-80. The UV-irradiated plasmid (100 mJ/cm^2^, to stimulate homologous recombination) was added to 200 μl of bacterial cells after which the electroporation was performed (GenePulser apparatus (Bio-Rad) set at 2500 mV, resistance 800 ohms, capacitance 25 μF). The cells were diluted with 1 ml of 7H9-ADS-Tw (pre-warmed at 37°C) after which 4 ml medium was added and the culture was placed overnight at 37°C. The culture was plated out on 7H10 with 50 μg/ml kanamycin and 50 μg/ml X-gal. The one blue colony (presence of *lacZ*) that was formed, was tested with colony PCR for the integration of the plasmid and grown in liquid 7H9 medium without kanamycin to stimulate a second homologous recombination, resulting in knocking out the *sapM* gene. To select for the clones that have lost the plasmid, the culture was plated on 7H10 + 2% sucrose + 50 μg/ml X-gal (presence of *sacB* which inhibits growth on sucrose medium). Several white colonies were tested for the absence of *sapM*, by means of PCR. One clone was selected for further work, which showed absence of SapM expression and thus lost both *sapM* loci (Suppl. Fig. 4b). This was confirmed by PCR, Southern Blot, copy number analysis via qPCR, qPCR-RT analysis, SapM ELISA and phosphatase assay (Suppl. Fig. 4c-i). The SapM ELISA and phosphatase assay were performed as described in ref^14^.

### Southern blot

To verify if deletion of the *sapM* gene had occurred, genomic DNA of the strains was digested with PvuII. The digested samples were blotted to an Amersham Hybond-N+ membrane (GE Healthcare) by the neutral denaturing procedure (see manufacturer’s instructions). We hybridized the membranes with a DIG-labelled *sapM* probe, created by PCR amplifying a region overlapping with the 5’ end of *sapM* (primers GGCTGGTGGGTTTGCTCGTCG and TGCCAGACCCACTTGTGGGACA) using a DIG-labeled synthetic dNTP mix (Roche Life Sciences). The membrane was incubated with an anti-DIG-AP antibody (1:10,000) (Roche Life Sciences), washed twice with washing buffer (0.1 M of maleic acid, 0.15 M of NaCl and 0.3% Tween-20, pH 7.5) and developed with the Amersham CDP-Star substrate in detection buffer (1:100 dilution) (GE Healthcare). The luminescent signal was measured by exposure to an X-ray film. The expected Southern Blot band for the WT was 2206 bp and 1598 bp for the *sapM* KO (Suppl. Fig. 4c).

### RT-qPCR analysis

*M. bovis* BCG cultures (grown in standard 7H9 medium until an OD_600_ of 0.8 – 1.0) were centrifuged and the pellets were washed once with sterile water containing 0.5% Tween-80. The pellet was then resuspended in 500 μl of RLT buffer (RNeasy Mini Kit, Qiagen; supplemented with β-ME). The cells were disrupted with glass beads in a Retsch MM2000 bead beater at 4°C in screw-cap tubes (pre-baked at 150°C). After centrifugation (2 min, 13,000 rpm, 4°C), the supernatant was transferred to a fresh eppendorf tube. To recover the lysate trapped in between the beads, 800 μl of chloroform was added to the beads and centrifuged, after which the upper phase was transferred to the same eppendorf tube as before. Then, 1 volume of Acid Phenol/Chloroform (Ambion) was added, incubated for 2 minutes and centrifuged (5 min, 13,000 rpm, 4°C). The upper aqueous phase was transferred to a fresh eppendorf tube and this last step was repeated once. An equal volume of 70% ethanol was added and the sample was transferred to an RNeasy spin column (RNeasy Mini Kit, Qiagen). The kit manufacturer’s instructions were followed to purify the RNA. After elution in RNase-free water (30 μl), an extra DNase digestion was performed with DNaseI (10 U of enzyme, 50 μl reaction volume). Then an extra clean-up step was performed with the RNeasy Mini Kit (Qiagen). Finally, the RNA concentration was determined on a Nanodrop instrument.

cDNA was prepared from 1 μg of DNase-treated RNA using the iScript Synthesis Kit (BioRad) and a control reaction lacking reverse transcriptase was included for each sample. The RT-PCR program was as follows: 10 min at 25°C, 30 min at 42°C, 5 min at 85°C.

Real time quantitative PCR was done on a LightCycler 480 (Roche Diagnostics) using the SensiFast SYBR-NoRox kit (BioLine), in triplicate for each cDNA sample. All gene expression values were normalized using the geometric mean of the *gap* and *pgk* rRNA. Determination of amplification efficiencies and conversion of raw Cq values to normalized relative quantities (NRQ) was performed using the qbasePLUS software. Statistical analysis of the NRQs was done with the Prism6.04 software package using an unpaired t-test, we corrected for multiple comparisons using the Holm-Sidak method (alpha = 0.05).

Primers used in the experiment: *gap* (TGGGAGTTAACGACGACAAG and ACTCATCGTCGAGCACTTTG), *pgk* (GAAACCAGCAAGAACGATGA and AACAGGGTTGCGATGTCATA), *upp* (CGGGTCGCGGCTAAC and GGGCAGCGAGTCCAGATA), *sapM* (TGCGGCCCGGAACTTACAACGAGA and CAAGCGGATGGGTACGAGGTCAGC) and BCG_3376 (AAGTTCTTCAACGGCAATCC and GTGCTGATGATCTCGTCGAT).

### Citation list for the used bioinformatics tools and database

*Artemis* – Carver, T., Harris, S. R., Berriman, M., Parkhill, J. & McQuillan, J. A. Artemis: an integrated platform for visualization and analysis of high-throughput sequence-based experimental data. Bioinformatics 28, 464-469 (2012).

*BBmerge* – Bushnell, B., Rood, J. & Singer, E. BBMerge – Accurate paired shotgun read merging via overlap. PLoS One 12, e0185056 (2017).

*BLAST* – Altschul, S. F., Gish, W., Miller, W., Myers, E. W. & Lipman, D. J. Basic local alignment search tool. J. Mol. Biol. 215, 403-410 (1990).

*BWA-MEM* – Li, H. Aligning sequence reads, clone sequences and assembly contigs with BWA-MEM. ArXiv E-Prints (2013).

*CLC Main Workbench 8* – Qiagen. CLC Main Workbench version 8. Available at: https://www.qiagenbioinformatics.com/.

*cn.mops* – Klambauer, G. et al. cn.MOPS: mixture of Poissons for discovering copy number variations in next-generation sequencing data with a low false discovery rate. Nucleic Acids Res. 40, e69 (2012).

*GapFiller* – Nadalin, F., Vezzi, F. & Policriti, A. GapFiller: a de novo assembly approach to fill the gap within paired reads. BMC Bioinformatics 13 Suppl 14, S8 (2012).

*GATK* – McKenna, A. et al. The Genome Analysis Toolkit: a MapReduce framework for analyzing next-generation DNA sequencing data. Genome Res 20, 1297-303 (2010).

*GeneMark.hmm* – Besemer, J. GeneMarkS: a self-training method for prediction of gene starts in microbial genomes. Implications for finding sequence motifs in regulatory regions. Nucleic Acids Res. 29, 2607-2618 (2001).

*GMAP* – Wu, T. D. & Nacu, S. Fast and SNP-tolerant detection of complex variants and splicing in short reads. Bioinformatics 26, 873-81 (2010).

*HINGE* – Kamath, G. M., Shomorony, I., Xia, F., Courtade, T. A. & Tse, D. N. HINGE: long-read assembly achieves optimal repeat resolution. Genome Res 27, 747-756 (2017).

*i-ADHoRe 3.0* – Proost, S. et al. i-ADHoRe 3.0–fast and sensitive detection of genomic homology in extremely large data sets. Nucleic Acids Res. 40, e11 (2012).

*Infernal* – Nawrocki, E. P. & Eddy, S. R. Infernal 1.1: 100-fold faster RNA homology searches. Bioinformatics 29, 2933-2935 (2013).

*Lordec* – Salmela, L. & Rivals, E. LoRDEC: accurate and efficient long read error correction. Bioinformatics 30, 3506-14 (2014).

*Pilon* – Walker, B. J. et al. Pilon: an integrated tool for comprehensive microbial variant detection and genome assembly improvement. PLoS One 9, e112963 (2014).

*SnpEff and SnpSift* – Cingolani, P. et al. A program for annotating and predicting the effects of single nucleotide polymorphisms, SnpEff. Fly (Austin) 6, 80-92 (2012).

*SPAdes* – Bankevich, A. et al. SPAdes: a new genome assembly algorithm and its applications to single-cell sequencing. J Comput Biol 19, 455-77 (2012).

*SSPACE-LongRead v3.0* – Boetzer, M. & Pirovano, W. SSPACE-LongRead: scaffolding bacterial draft genomes using long read sequence information. BMC Bioinformatics 15, 211 (2014).

*TribeMCL* – van Dongen, S. & Abreu-Goodger, C. Using MCL to extract clusters from networks. Methods Mol Biol 804, 281-95 (2012).

*Trimmomatic* – Bolger, A. M., Lohse, M. & Usadel, B. Trimmomatic: a flexible trimmer for Illumina sequence data. Bioinformatics 30, 2114-20 (2014).

*tRNAscan-SE* – Lowe, T. M. & Eddy, S. R. tRNAscan-SE: a program for improved detection of transfer RNA genes in genomic sequence. Nucleic Acids Res 25, 955-64 (1997).

*UniProt* – The UniProt, C. UniProt: the universal protein knowledgebase. Nucleic Acids Res 45, D158-D169 (2017).

**Supplementary Figure 1.**
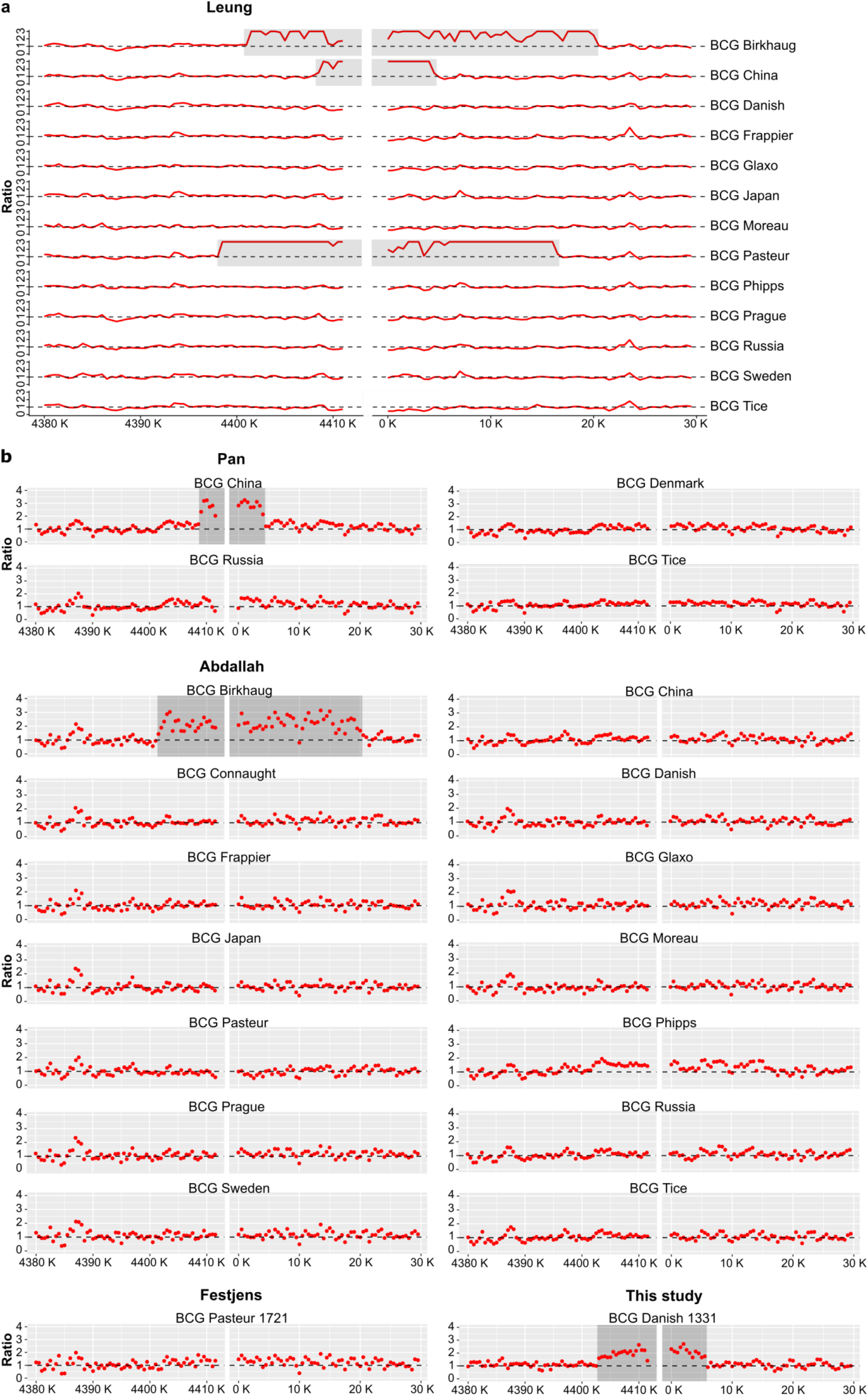
DU1 duplication detection in BCG strains. Tiling array data **(a)** from Leung *et al.* 2008^13^ and Illumina sequencing data **(b)** for BCG Danish 1331 (this study) as well as published genome data from Pan *et al.* 2011^7^, Abdallah *et al.* 2015^8^ and Festjens *et al.* 2018^14^ were reanalyzed for the presence of a DU1 in the region of the *oriC.* These references were chosen as they contain BCG Danish genome sequencing data. The graphs in (a) depict the ratio of the reference (M. *tb* H37Rv) probe intensity (Cy5) divided by the test (BCG strain) probe intensity as originally presented in Leung *et al.* 2008^13^. The graphs in (b) depict the ratio of mean whole genome read coverage divided by the mean read coverage in 500 bp window size. Detection of a DU1-like duplication in BCG Pasteur 1173P2^13^, Birkhaug^8, 13^, Danish 1331 07/270 (this study) and BCG China^7, 13^, indicated in grey. No detection of DU1-duplication for other BCG Pasteur^8, 14^, Danish^7, 8^ and China^8^ strains.

**Supplementary Figure 2.**
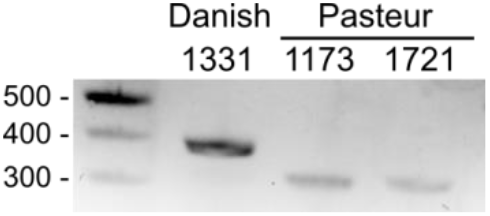
Analysis of the SenX3-RegX3 region in BCG strains. PCR was performed on gDNA of BCG Danish 1331, Pasteur 1173 and 1721 using primer pair 9. This primer set was originally designed by Bedwell *et al.* 2001^10^ to identify BCG substrains by multiplex-PCR. Dependent on the number of repeats of 77 bp in the SenX3-RegX3 region a different amplicon is formed; a 353 bp amplicon for 3 repeats and a 276 bp amplicon for 2 repeats.

**Supplementary Figure 3.**
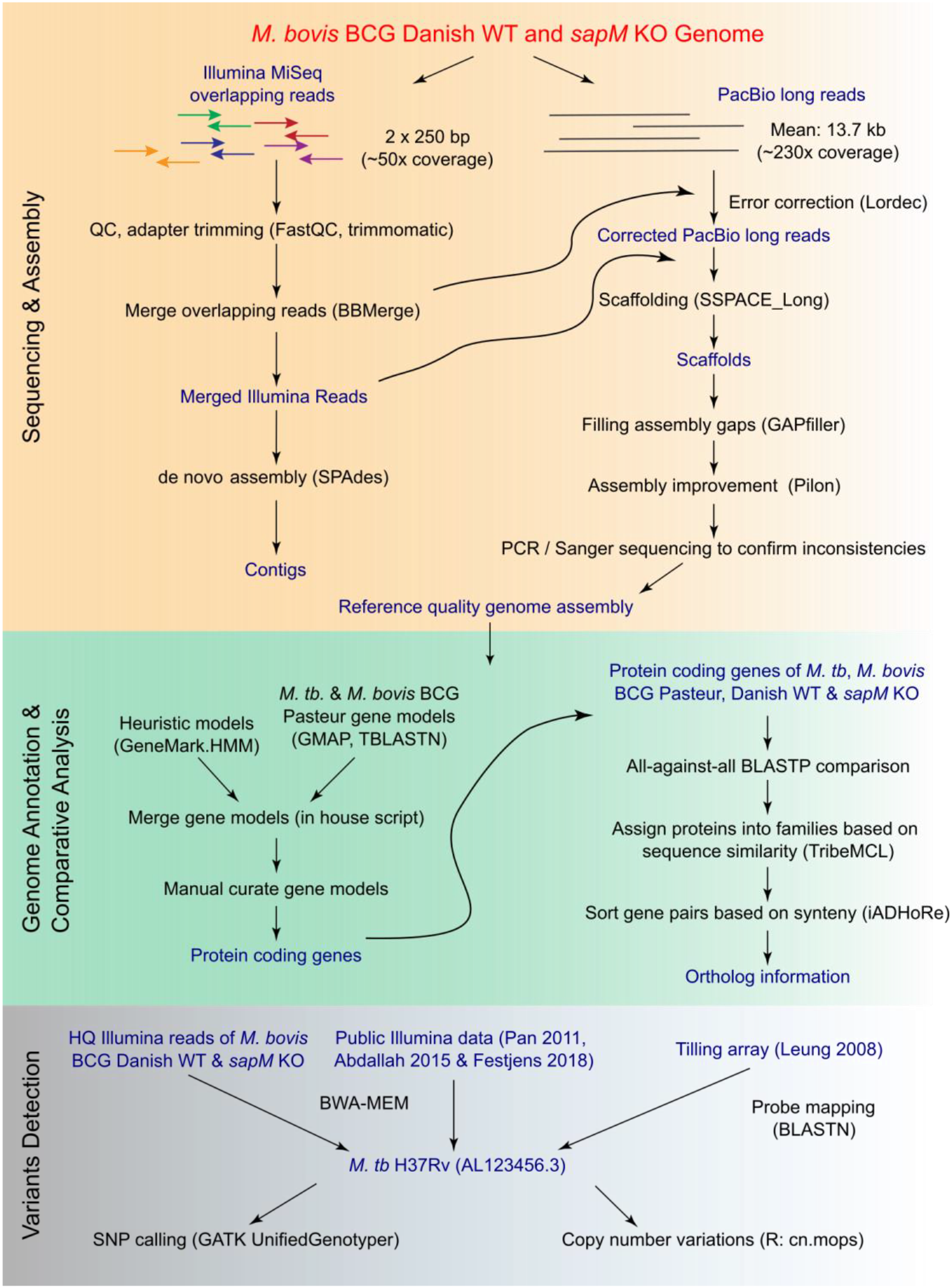
Genome analysis pipeline.

**Supplementary Figure 4.**
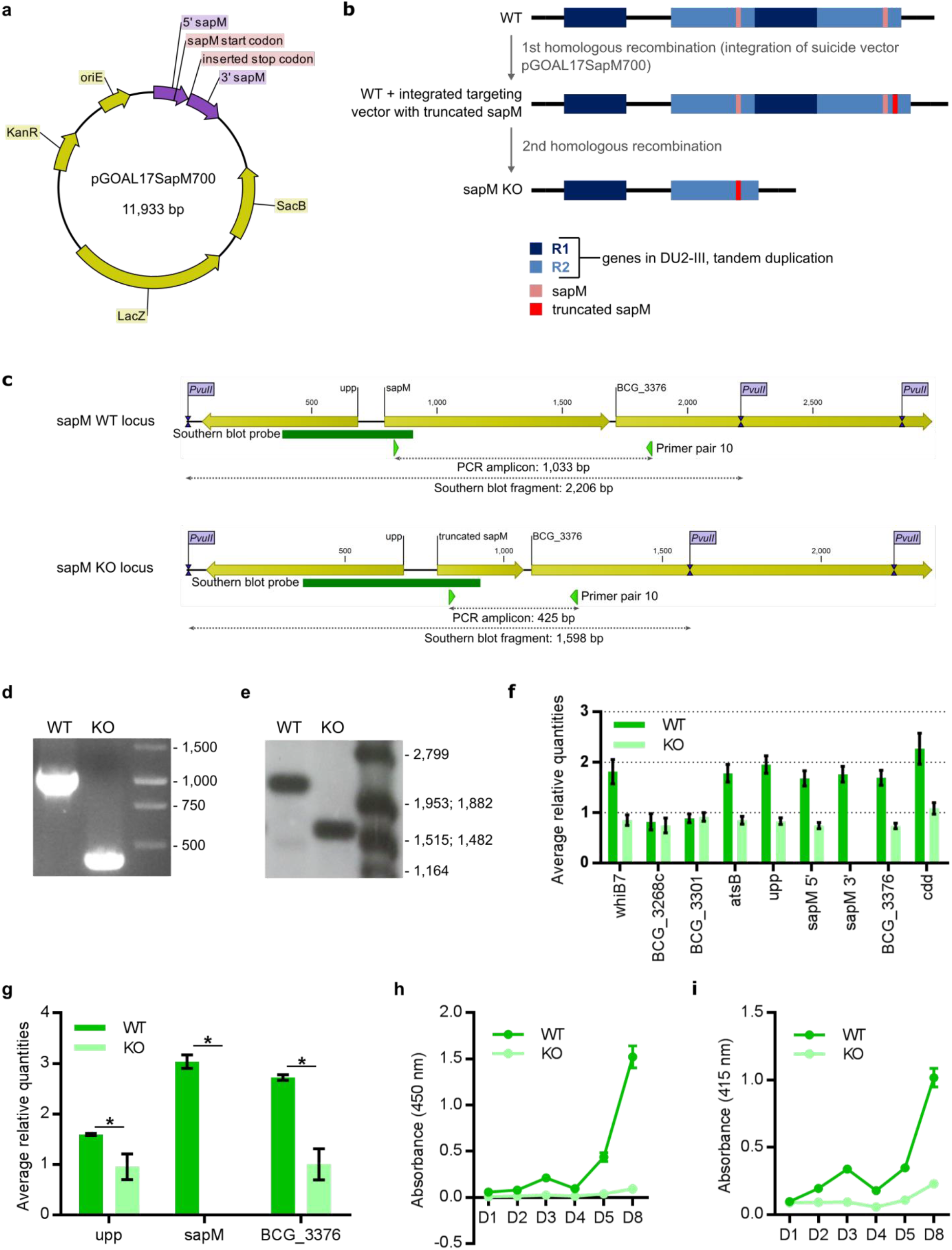
Generation and characterization of BCG Danish 1331 *sapM* KO. **a)** Plasmid map of suicide vector pGOAL17SapM700. **b)** Hypothesis for the sequence of genomic rearrangements during production of BCG Danish 1331 *sapM* KO from BCG Danish 1331 WT, which contains two *sapM* loci, due to the presence of the *sapM* locus in the DU2 duplicated genomic region. **c)** Representation of the *sapM* WT and KO locus with indication of theoretical PCR amplicons **(d)** and Southern blot fragments (PvuII digest and hybridization with Southern blot probe) (**e**). **d)** PCR analysis of the *sapM* locus. WT and *sapM* KO gDNA were amplified with primer pair 10. **e)** Southern blot for the 5’ part of *sapM* of *M. bovis* BCG WT and *sapM* KO after *PvuII* digest. **f)** Copy-number analysis of *sapM* and its surrounding genes via qPCR on gDNA. Next to the truncation of *sapM*, the DU2 has been lost in the *sapM* KO, since all duplicated genes in the DU2 are reduced to one copy in the *sapM* KO. The represented data are averages (± SD) of four technical replicates. **g)** RT-PCR analysis of the *sapM* locus. RNA was prepared of cultures of biological triplicates of the *sapM* KO and parental strain. RT-PCR on the cDNA was performed using primer sets directed against *sapM* and the directly up- and downstream genes (*upp* and BCG_3376). The data presented here are averages (± SEM) of three biological replicates. (*: p < 0.05). **h-i)** Analysis of SapM protein quantity and enzymatic activity. Strains were subcultured from a mid-log culture to an OD_600_ of 0.1, from which medium samples were collected for 8 days. ELISA using an anti-SapM polyclonal antibody **(h)** and *in vitro* phosphatase assay using pNPP as a substrate to check phosphatase activity **(i)**. The represented data are averages (± SD) of three biological replicates.

**Supplementary Table 1:** Summary table of SNPs detected in *M. bovis* Danish 1331 WT and *sapM* KO compared to the Pasteur reference 1173P2 (NC_008679.1).

| Variants | Type | WT | KO | non-shared |
| --- | --- | --- | --- | --- |
| SNP's | intergenic_region | 9 | 9 | 0 |
|  | missense_variant | 23 | 24 | 1 |
|  | stop_gained | 1 | 1 | 0 |
|  | synonymous_variant | 9 | 9 | 0 |
|  | **total** | **42** | **43** | **1** |
| genes affected by SNP's | missense_variants | 18 | 19 | 1 |
|  | stop_gained | 1 | 1 | 0 |
|  | **total** | **19** | **20** | **1** |

**Supplementary Table 2.**
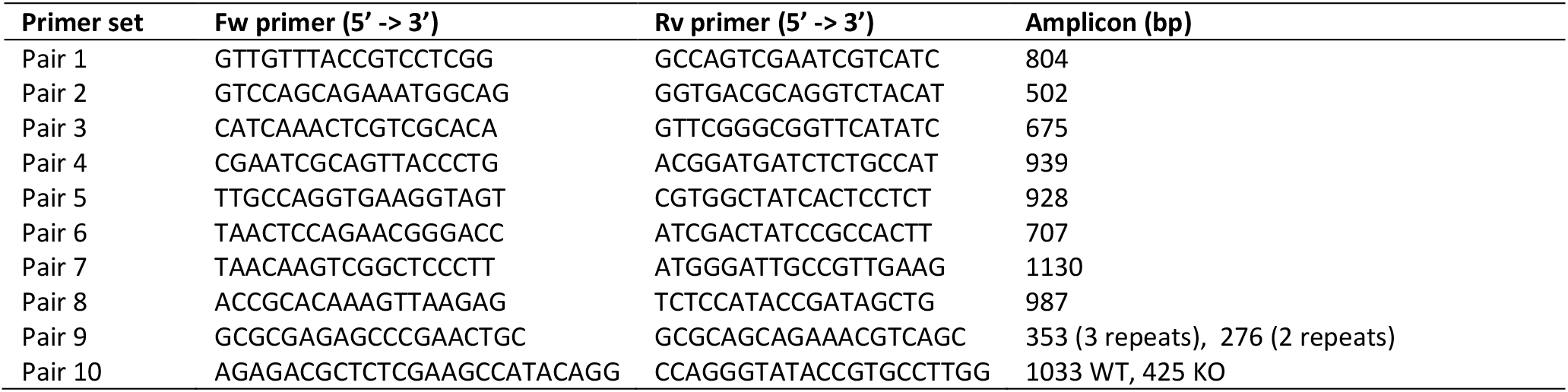
PCR primer pairs for confirmation of the genome assembly (pair 1-8), analysis of the SenX3-RegX3 region (pair 9) and validation of the KO engineering (pair 10)

**Supplementary Table 3.**
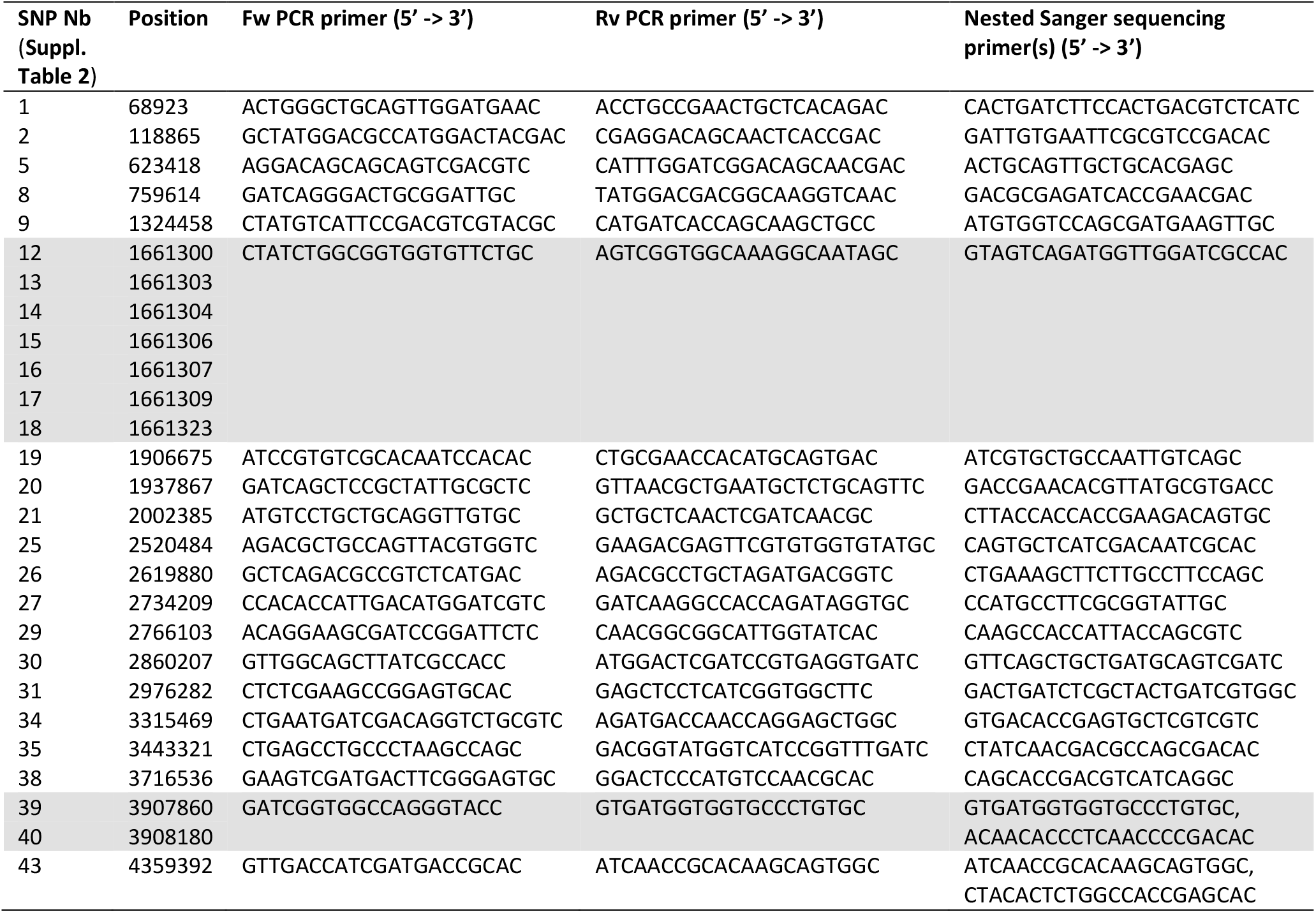
PCR primer pairs for confirmation of the SNP variants.

**Supplementary Table 4.**
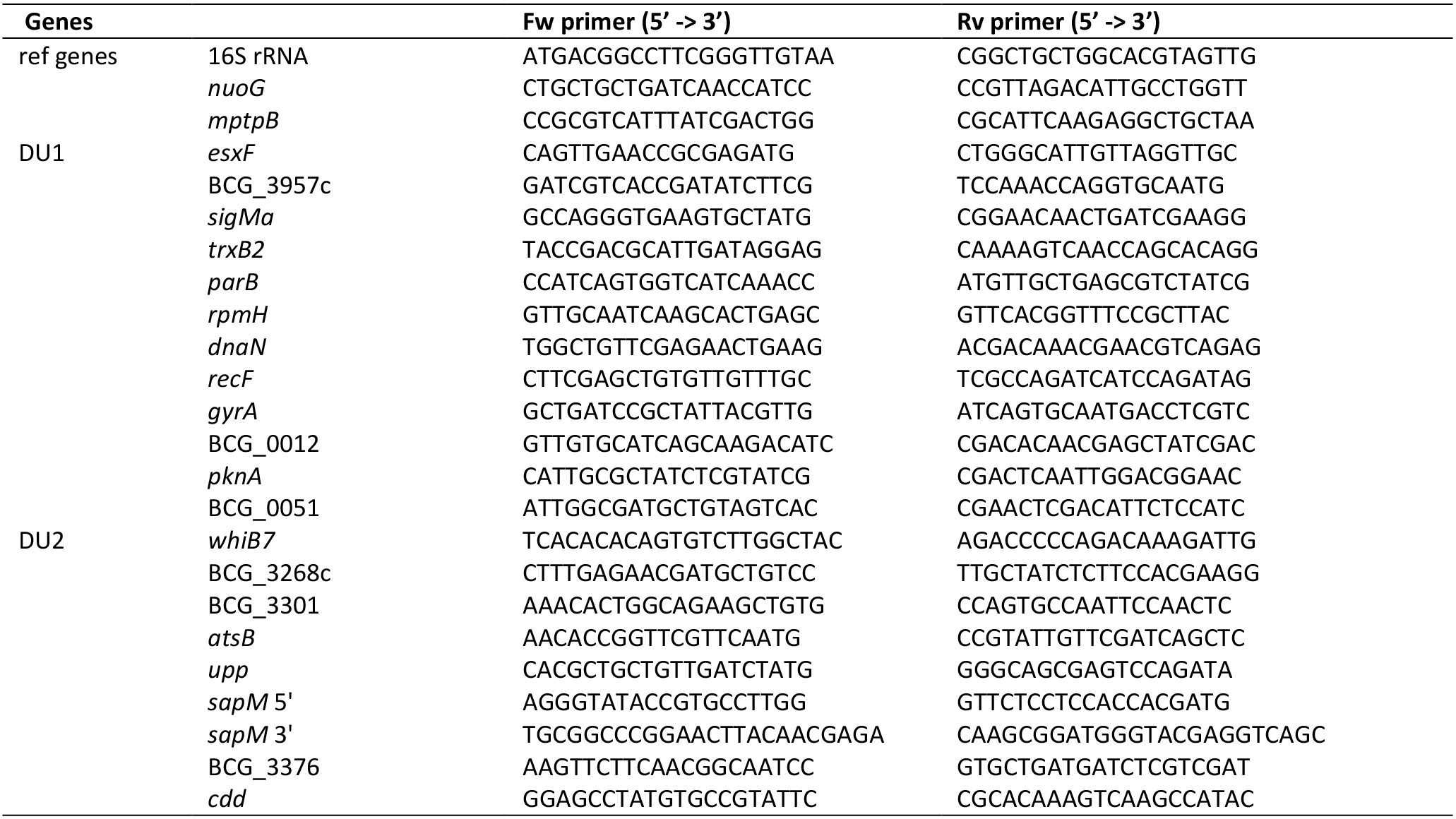
qPCR primer pairs for copy number profiling.

