## Supplemental Table 5 for "Reference genome for the WHO reference strain for *Mycobacterium bovis* BCG Danish, the present tuberculosis vaccine"

**Supplementary Excel 1: SNPs detected in *M. bovis* Danish 1331 WT and *sapM* KO compared to the Pasteur reference 1173P2 (NC_008679.1).** SNP validation via PCR and Sanger sequencing for all genes containing missense or stop gained SNPs.(*) PCR and/or Sanger sequencing was unsuccessful, all these regions are GC-rich or contain repeats.

| **Nb** | **Position in ref** | **Pasteur ref** | **Danish 1331** | | **SNP effect** | **SNP effect on gene** | | | | | **PCR + Sanger sequencing performed** |
|---|---|---|---|---|---|---|---|---|---|---|---|
| | | | **WT** | **KO** | | **Effect** | **GeneName** | **GeneID** | **Start gene** | **End gene** | |
| 1 | 68923 | A | G | G | missense variant | p.Leu186Pro | BCG_0067c | BCG_0067c | 68799 | 69479 | yes |
| 2 | 118865 | T | C | C | missense variant | p.Cys35Arg | BCG_0113 | BCG_0113 | 118763 | 119221 | yes |
| 3 | 190828 | C | T | T | synonymous variant | p.Ala21Ala | BCG_0167 | BCG_0167 | 190766 | 191248 | no |
| 4 | 370270 | C | T | T | intergenic region | . | . | . | . | . | no |
| 5 | 623418 | C | T | T | stop gained | p.Gln323* | galE2 | BCG_0544 | 622452 | 623582 | yes |
| 6 | 705623 | G | A | A | synonymous variant | p.His392His | PE_PGRS7 | BCG_0623c | 702887 | 706798 | no |
| 7 | 705626 | G | C | C | synonymous variant | p.Ala391Ala | PE_PGRS7 | BCG_0623c | 702887 | 706798 | no |
| 8 | 759614 | T | C | C | missense variant | p.Met184Val | echA3 | BCG_0679c | 759468 | 760163 | yes |
| 9 | 1324458 | T | C | C | missense variant | p.Leu106Pro | narJ | BCG_1225 | 1324142 | 1324747 | yes |
| 10 | 1344671 | C | G | G | intergenic region | . | . | . | . | . | no |
| 11 | 1344672 | G | C | C | intergenic region | . | . | . | . | . | no |
| x | 1661298 | C | T | T | missense variant | p.Ala651Gly | PE_PGRS28 | BCG_1513c | 1661026 | 1663248 | no (*) |
| 12 | 1661300 | A | C | C | missense variant | p.Val650Gly | PE_PGRS28 | BCG_1513c | 1661026 | 1663248 | no (*) |
| 13 | 1661303 | A | T | T | missense variant | p.Ile649Asn | PE_PGRS28 | BCG_1513c | 1661026 | 1663248 | no (*) |
| 14 | 1661304 | T | C | C | missense variant | p.Ile649Val | PE_PGRS28 | BCG_1513c | 1661026 | 1663248 | no (*) |
| 15 | 1661306 | A | C | C | missense variant | p.Ile648Ser | PE_PGRS28 | BCG_1513c | 1661026 | 1663248 | no (*) |
| 16 | 1661307 | T | C | C | missense variant | p.Ile648Val | PE_PGRS28 | BCG_1513c | 1661026 | 1663248 | no (*) |
| 17 | 1661309 | T | C | C | missense variant | p.Asp647Gly | PE_PGRS28 | BCG_1513c | 1661026 | 1663248 | no (*) |
| 18 | 1661323 | G | C | C | synonymous variant | p.Gly642Gly | PE_PGRS28 | BCG_1513c | 1661026 | 1663248 | no (*) |
| 19 | 1906675 | G | A | A | missense variant | p.Asp119Asn | BCG_1714 | BCG_1714 | 1906321 | 1907025 | yes |
| 20 | 1937867 | G | T | T | missense variant | p.Ala2Asp | PPE22 | BCG_1743c | 1936714 | 1937871 | yes |
| 21 | 2002385 | G | C | C | missense variant | p.Arg129Gly | wag22b | BCG_1799c | 2000361 | 2002769 | no (*) |
| 22 | 2143356 | T | G | G | intergenic region | . | . | . | . | . | no |
| 23 | 2143357 | A | G | G | intergenic region | . | . | . | . | . | no |
| 24 | 2442228 | G | A | A | synonymous variant | p.Gly23Gly | BCG_2215c | BCG_2215c | 2441877 | 2442296 | no |
| 25 | 2520484 | C | T | T | missense variant | p.Arg260His | BCG_2284c | BCG_2284c | 2520096 | 2521262 | yes |
| 26 | 2619880 | T | C | C | missense variant | p.Gln79Arg | hrcA | BCG_2388c | 2619084 | 2620115 | yes |
| 27 | 2734209 | G | A | A | missense variant | p.Met18Ile | pepN | BCG_2487 | 2734156 | 2736741 | yes |
| 28 | 2764157 | T | A | A | synonymous variant | p.Thr177Thr | BCG_2507c | BCG_2507c | 2761274 | 2764687 | no |
| 29 | 2766103 | G | C | C | missense variant | p.Ala804Gly | PE_PGRS43b | BCG_2509c | 2765061 | 2768513 | no (*) |
| 30 | 2860207 | A | G | G | missense variant | p.Cys68Arg | BCG_2594c | BCG_2594c | 2859341 | 2860408 | yes |
| 31 | 2976282 | T | C | C | missense variant | p.Val46Ala | sigB | BCG_2723 | 2976146 | 2977117 | yes |
| 32 | 3179819 | T | C | C | synonymous variant | p.Leu133Leu | ffh | BCG_2937c | 3178640 | 3180217 | no |
| 33 | 3237056 | G | T | T | intergenic region | . | . | . | . | . | no |
| 34 | 3315469 | A | G | G | missense variant | p.Ile50Thr | ilvH | BCG_3024c | 3315111 | 3315617 | yes |
| 35 | 3443321 | G | A | A | missense variant | p.Ser14Leu | PPE49a | BCG_3147c | 3442513 | 3443361 | yes |
| 36 | 3452980 | G | A | A | intergenic region | . | . | . | . | . | no |
| 37 | 3606542 | C | T | T | synonymous variant | p.Asp208Asp | BCG_3301 | BCG_3301 | 3605919 | 3607103 | no |
| 38 | 3716536 | A | G | G | missense variant | p.Val198Ala | icd1 | BCG_3409c | 3715899 | 3717128 | yes |
| 39 | 3907860 | A | G | G | missense variant | p.Asn599Asp | PE_PGRS53 | BCG_3571 | 3906066 | 3910184 | no (*) |
| 40 | 3908180 | C | T | T | synonymous variant | p.Gly705Gly | PE_PGRS53 | BCG_3571 | 3906066 | 3910184 | no (*) |
| 41 | 4061576 | C | G | G | intergenic region | . | . | . | . | . | no |
| 42 | 4080517 | T | A | A | intergenic region | . | . | . | . | . | no |
| 43 | 4359392 | A | A | C | missense variant | p.Asn737Thr | BCG_3966 | BCG_3966 | 4357183 | 4359591 | yes |
