## Supplemental Table 6 for "Reference genome for the WHO reference strain for *Mycobacterium bovis* BCG Danish, the present tuberculosis vaccine"

**Supplementary Excel 2. Distribution of regions of difference, deletions and tandem duplications (DU1 and DU2) in the different BCG strains compared to *M. bovis* AF2122_97 (NC_002945.4).** Adapted from Abdallah *et al.* , 2015; only the RD and deleted genes that could be verified on the assembled genomes upon manual inspection are included. The regions of difference, deletions and DU are grouped according to their occurrence in the different strains (instead of alphabetically) to allow easy discrimination of the different lineages; which is further promoted by the used color coding. In green 'normal' situation, in pink 'deletion', in blue 'insertion'. ND = no (sufficient) data, del = deletion, ins = insertion, indel = insertion-deletion. According to literature two different groups of BCG exist, that are both named BCG China or BCG Beijing, BCG China corresponds with data from Abdallah *et al.* 2015 and BCG China* corresponds with data from Leung *et al.* 2008 and Pan *et al.* 2011.

| **Region of difference** | ***M. bovis*** | **Russia** | **Moreau** | **Japan (Tokyo)** | **Sweden** | **Birkhaug** | **Prague** | **China*** | **Glaxo** | **Danish (Copenhagen)** | **Frappier** | **Connaught** | **Mexico** | **Tice** | **China** | **Phipps** | **Pasteur** |
|---|---|---|---|---|---|---|---|---|---|---|---|---|---|---|---|---|---|
| RDPan (Rv3887c-Rv3889c) | Deleted | Present | Present | Present | Present | Present | Present | Present | Present | Present | Present | Present | Present | Present | Present | Present | Present |
| RD1 | Mb3901 to Mb3909c | Deleted | Deleted | Deleted | Deleted | Deleted | Deleted | Deleted | Deleted | Deleted | Deleted | Deleted | Deleted | Deleted | Deleted | Deleted | Deleted |
| RD3 | Mb1599 to Mb1612c | Deleted | Deleted | Deleted | Deleted | Deleted | Deleted | Deleted | Deleted | Deleted | Deleted | Deleted | Deleted | Deleted | Deleted | Deleted | Deleted |
| NN-RD5 | Mb2377c | Deletion | Deletion | Deletion | Deletion | Deletion | Deletion | Deletion | Deletion | Deletion | Deletion | Deletion | Deletion | Deletion | Deletion | Deletion | Deletion |
| *hspR* -PPE8 | Mb0361-62c | 102 bp del | 102 bp del | 102 bp del | 102 bp del | 102 bp del | 102 bp del | 102 bp del | 102 bp del | 102 bp del | 102 bp del | 102 bp del | 102 bp del | 102 bp del | 102 bp del | 102 bp del | 102 bp del |
| Mb1758c-59c | Mb1758c-59c | 114 bp del | 114 bp del | 114 bp del | 114 bp del | 114 bp del | 114 bp del | 114 bp del | 114 bp del | 114 bp del | 114 bp del | 114 bp del | 114 bp del | 114 bp del | 114 bp del | 114 bp del | 114 bp del |
| Mb2165c-66c | Mb2165c-66c | 116 bp del | 116 bp del | 116 bp del | 116 bp del | 116 bp del | 116 bp del | 116 bp del | 116 bp del | 116 bp del | 116 bp del | 116 bp del | 116 bp del | 116 bp del | 116 bp del | 116 bp del | 116 bp del |
| *dnaQ*- Mb3739 | Mb3738c-39 | 59 bp del | 59 bp del | 59 bp del | 59 bp del | 59 bp del | 59 bp del | 59 bp del | 59 bp del | 59 bp del | 59 bp del | 59 bp del | 59 bp del | 59 bp del | 59 bp del | 59 bp del | 59 bp del |
| IS6110 upstream of *PhoP* | Mb0780 | Present | Present | Present | Deleted | Deleted | Deleted | Deleted | Deleted | Deleted | Deleted | Deleted | Deleted | Deleted | Deleted | Deleted | Deleted |
| RDRussia | Mb3723c to Mb3724 | Deleted | Present | Present | Present | Present | Present | Present | Present | Present | Present | Present | Present | Present | Present | Present | Present |
| RDMoreau (Rv3887c) | Deleted (<RDPan) | Present | 876 bp del | Present | Present | Present | Present | Present | Present | Present | Present | Present | Present | Present | Present | Present | Present |
| RD16 | Mb3433 to Mb3439c | Present | Deleted | Present | Present | Present | Present | Present | Present | Present | Present | Present | Present | Present | Present | Present | Present |
| *fadD26-ppsA* | Mb2955 to Mb2956 | Present | 976 bp del | Present | Present | Present | Present | Present | Present | Present | Present | Present | Present | Present | Present | Present | Present |
| RDJapan | Mb3439c | Present | Deleted (<RD16) | 22 bp del | Present | Present | Present | Present | Present | Present | Present | Present | Present | Present | Present | Present | Present |
| *trcR* | Mb1062c | Present | Present | Present | 244 bp del | 244 bp del | Present | Present | Present | Present | Present | Present | Present | Present | Present | Present | Present |
| *whiB3* | Mb3450 | Present | Present | Present | 111 bp del | 111 bp del | Present | Present | Present | Present | Present | Present | Present | Present | Present | Present | Present |
| *guaB1-gnd1* | Mb1874c-75c | Present | Present | Present | Present | 118 bp del | Present | Present | Present | Present | Present | Present | Present | Present | Present | Present | Present |
| RD2 | Mb2000 to Mb2010 | Present | Present | Present | Present | Present | Deleted | Deleted | Deleted | Deleted | Deleted | Deleted | Deleted | Deleted | Deleted | Deleted | Deleted |
| nRD18 | Mb1221 to Mb1223 | Present | Present | Present | Present | Present | Present | Present | Present | Present | Deleted | Deleted | Deleted | Deleted | Deleted | Deleted | Deleted |
| Mb1571 | Mb1571 | Present | Present | Present | Present | Present | Present | Present | Present | Present | 10 bp del | 10 bp del | 10 bp del | 10 bp del | 10 bp del | 10 bp del | 10 bp del |
| RD15 (Brosch 2007), RD08 (Behr 1999) | Mb0317 to Mb0320 | Present | Present | Present | Present | Present | Present | Present | Present | Present | Deleted | Deleted | Present | Present | Present | Present | Present |
| RDFrappier | Mb3525c to Mb3527c | Present | Present | Present | Present | Present | Present | Present | Present | Present | Deleted | Present | Present | Present | Present | Present | Present |
| RDMexico | Mb3890 to Mb3892c | Present | Present | Present | Present | Present | Present | Present | Present | Present | Present | Present | Deleted | Present | Present | Present | Present |
| Mb0096c | Mb0096c | Present | Present | Present | Present | Present | Present | Present | Present | Present | Present | Present | Present | 43 bp indel | Present | Present | Present |
| RD14 | Mb1795 to Mb1802c | Present | Present | Present | Present | Present | Present | Present | Present | Present | Present | Present | Present | Present | Present | Deleted | Deleted |
| RDDenmark/Glaxo | Mb1839 to Mb1840 | Present | Present | Present | Present | Present | Present | Present | Deleted | Deleted | Present | Present | Present | Present | Present | Present | Present |
| *frdBC* | Mb1579 | Present | Present | Present | Present | Present | Present | Present | 5 bp ins | Present | Present | Present | Present | Present | Present | Present | Present |
| *PhoR* | Mb0781 | Present | Present | Present | 11 bp del | 11 bp del | Present | ND | 10 bp del | 10 bp del | 1 bp del | 1 bp del | Present | Present | Present | Present | Present |
| *PhoP* | Mb0780 | Present | Present | Present | Present | Present | Ins G | ND | Present | Present | Present | Present | Present | Present | Present | Present | Present |
| DU1-like duplication | - | - | - | - | - | DU1-Birkhaug | - | - | - | DU1-Danish in certain substrains, e.g. NIBSC 07/270 | - | - | - | - | DU1-China | - | DU1-Pasteur in certain substrains, e.g. Pasteur 1173P2 |
| other type of DU than DU1 or DU2 | - | - | - | - | - | - | - | - | - | - | - | - | - | DU-Tice | - | - | - |
| type of DU2 | - | I | | | II | | III | | | | IV | | | | | | |
