## Supplemental Table 7 Worksheet 1 for "Reference genome for the WHO reference strain for *Mycobacterium bovis* BCG Danish, the present tuberculosis vaccine"

**Supplementary Excel 3. Ortholog of mycobacterial genes between *M. tb* H37Rv, *M. bovis* BCG Pasteur, *M. bovis* BCG Danish WT and *sapM* KO.** In worksheet 1 all CDS for the different strains are depicted, as well as two Venn diagrams illustrating the ortholog between the different strains. In worksheet 2 only the duplicated CDS are incorporated.

| *M. tuberculosis* H37Rv (AL123456.3, 4032 CDS) | | | | | *M. bovis* BCG Pasteur 1173P2 (AM408590.1, 3988 CDS) | | | | | *M. bovis* BCG Danish 1331 (NC….. (in process), 4034 CDS) | | | | | *M. bovis* BCG Danish 1331 *sapM* KO (NC…. (in process),3959 CDS) | | | | |
|---|---|---|---|---|---|---|---|---|---|---|---|---|---|---|---|---|---|---|---|
| start M. tb | end M. tb | CDS name/l | locus_tag M. | length M. tb | start Pasteur | end Pasteur | CDS name/l | locus_tag Pa | length Paste | start Danish | end Danish | CDS name/locus_ | locus_tag Dani | length Danis | start Danish | end Danish s | CDS name/locus_tag Dar | locus_tag Danish sapM K | length Danish sapM KO |
| 1 | 1524 | dnaA | Rv0001 | 1524 | 1 | 1524 | dnaA_1 | BCG_0001 | 1524 | 1 | 1524 | dnaA_1 | BCGDan_0001 | 1524 | 1 | 1524 | dnaA_1 | BCGDan_sapMKO_0001 | 1524 |
| 2052 | 3260 | dnaN | Rv0002 | 1209 | 2052 | 3260 | dnaN_1 | BCG_0002 | 1209 | 2052 | 3260 | dnaN_1 | BCGDan_0002 | 1209 | 2127 | 3260 | dnaN_1 | BCGDan_sapMKO_0002 | 1134 |
| 3280 | 4437 | recF | Rv0003 | 1158 | 3280 | 4437 | recF_1 | BCG_0003 | 1158 | 3280 | 4437 | recF_1 | BCGDan_0003 | 1158 | 3451 | 4437 | recF_1 | BCGDan_sapMKO_0003 | 987 |
| 4434 | 4997 | Rv0004 | Rv0004 | 564 | 4434 | 4997 | BCG_0004 | BCG_0004 | 564 | 4434 | 4997 | BCGDan_0004 | BCGDan_0004 | 564 | 4434 | 4997 | BCGDan_sapMKO_0004 | BCGDan_sapMKO_0004 | 564 |
| 5240 | 7267 | gyrB | Rv0005 | 2028 | 5123 | 7267 | gyrB_1 | BCG_0005 | 2145 | 5123 | 7267 | gyrB_1 | BCGDan_0005 | 2145 | 5123 | 7267 | gyrB_1 | BCGDan_sapMKO_0005 | 2145 |
| 7302 | 9818 | gyrA | Rv0006 | 2517 | 7302 | 9818 | gyrA_1 | BCG_0006 | 2517 | 7302 | 9818 | gyrA | BCGDan_0006 | 2517 | 7302 | 9818 | gyrA | BCGDan_sapMKO_0006 | 2517 |
| 9914 | 10828 | Rv0007 | Rv0007 | 915 | 9914 | 10828 | BCG_0007 | BCG_0007 | 915 | 9914 | 10828 | BCGDan_0007 | BCGDan_0007 | 915 | 9914 | 10828 | BCGDan_sapMKO_0007 | BCGDan_sapMKO_0007 | 915 |
| 11874 | 12311 | Rv0008c | Rv0008c | 438 | 11874 | 12311 | BCG_0008c | BCG_0008c | 438 | 11874 | 12311 | BCGDan_0010 | BCGDan_0010 | 438 | 11874 | 12311 | BCGDan_sapMKO_0010 | BCGDan_sapMKO_0010 | 438 |
| 12468 | 13016 | ppiA | Rv0009 | 549 | 12468 | 13016 | ppiA_1 | BCG_0009 | 549 | 12468 | 13016 | ppiA_2 | BCGDan_0011 | 549 | 12468 | 13016 | ppiA_2 | BCGDan_sapMKO_0011 | 549 |
| 13133 | 13558 | Rv0010c | Rv0010c | 426 | 13222 | 13557 | BCG_0010c | BCG_0010c | 336 | 13222 | 13557 | BCGDan_0012 | BCGDan_0012 | 336 | 13222 | 13557 | BCGDan_sapMKO_0012 | BCGDan_sapMKO_0012 | 336 |
| 13714 | 13995 | Rv0011c | Rv0011c | 282 | 13713 | 13994 | BCG_0011c | BCG_0011c | 282 | 13713 | 13997 | BCGDan_0013 | BCGDan_0013 | 285 | 13713 | 13997 | BCGDan_sapMKO_0013 | BCGDan_sapMKO_0013 | 285 |
| 14089 | 14877 | Rv0012 | Rv0012 | 789 | 14088 | 14876 | BCG_0012 | BCG_0012 | 789 | 14088 | 14876 | BCGDan_0014 | BCGDan_0014 | 789 | 14178 | 14876 | BCGDan_sapMKO_0014 | BCGDan_sapMKO_0014 | 699 |
| 14914 | 15612 | trpG | Rv0013 | 699 | 14913 | 15611 | trpG_1 | BCG_0013 | 699 | 14913 | 15611 | trpG | BCGDan_0015 | 699 | 14913 | 15611 | trpG | BCGDan_sapMKO_0015 | 699 |
| 15590 | 17470 | pknB | Rv0014c | 1881 | 15589 | 16734 | 'pknB | BCG_0014c | 1146 | 15589 | 17469 | pknB | BCGDan_0016 | 1881 | 15589 | 17469 | pknB | BCGDan_sapMKO_0016 | 1881 |
| 17467 | 18762 | pknA | Rv0015c | 1296 | 47133 | 48428 | pknA | BCG_0045c | 1296 | 17466 | 18761 | pknA | BCGDan_0017 | 1296 | 17466 | 18743 | pknA | BCGDan_sapMKO_0017 | 1278 |
| 18759 | 20234 | pbpA | Rv0016c | 1476 | 48425 | 49900 | pbpA | BCG_0046c | 1476 | 18758 | 20233 | pbpA | BCGDan_0018 | 1476 | 18758 | 20233 | pbpA | BCGDan_sapMKO_0018 | 1476 |
| 20231 | 21640 | rodA | Rv0017c | 1410 | 49897 | 51306 | rodA | BCG_0047c | 1410 | 20230 | 21639 | rodA | BCGDan_0019 | 1410 | 20230 | 21639 | rodA | BCGDan_sapMKO_0019 | 1410 |
| 21637 | 23181 | pstP | Rv0018c | 1545 | 51303 | 52847 | pstP | BCG_0048c | 1545 | 21636 | 23180 | pstP | BCGDan_0020 | 1545 | 21636 | 23171 | pstP | BCGDan_sapMKO_0020 | 1536 |
| 23270 | 23737 | fhaB | Rv0019c | 468 | 52936 | 53403 | BCG_0049c | BCG_0049c | 468 | 23269 | 23736 | BCGDan_0021 | BCGDan_0021 | 468 | 23269 | 23736 | BCGDan_sapMKO_0021 | BCGDan_sapMKO_0021 | 468 |
| 23861 | 25444 | fhaA | Rv0020c | 1584 | 53527 | 55092 | TB39.8 | BCG_0050c | 1566 | 23860 | 25425 | TB39.8 | BCGDan_0022 | 1566 | 23860 | 25425 | TB39.8 | BCGDan_sapMKO_0022 | 1566 |
| 25913 | 26881 | Rv0021c | Rv0021c | 969 | 55561 | 56529 | BCG_0051c | BCG_0051c | 969 | 25894 | 26862 | BCGDan_0024 | BCGDan_0024 | 969 | 25894 | 26835 | BCGDan_sapMKO_0024 | BCGDan_sapMKO_0024 | 942 |
| 27023 | 27442 | whiB5 | Rv0022c | 420 | 56671 | 57090 | whiB5 | BCG_0052c | 420 | 27004 | 27423 | whiB5 | BCGDan_0025 | 420 | 27004 | 27423 | whiB5 | BCGDan_sapMKO_0025 | 420 |
| 27595 | 28365 | Rv0023 | Rv0023 | 771 | 57243 | 58013 | BCG_0053 | BCG_0053 | 771 | 27576 | 28346 | BCGDan_0026 | BCGDan_0026 | 771 | 27576 | 28346 | BCGDan_sapMKO_0026 | BCGDan_sapMKO_0026 | 771 |
| 28362 | 29207 | Rv0024 | Rv0024 | 846 | 58010 | 58843 | BCG_0054 | BCG_0054 | 834 | 28343 | 29176 | BCGDan_0027 | BCGDan_0027 | 834 | 28343 | 29176 | BCGDan_sapMKO_0027 | BCGDan_sapMKO_0027 | 834 |
| NA | NA | NA | NA | NA | 58585 | 58854 | BCG_0055 | BCG_0055 | 270 | 28918 | 29187 | BCGDan_0028 | BCGDan_0028 | 270 | 28918 | 29184 | BCGDan_sapMKO_0028 | BCGDan_sapMKO_0028 | 267 |
| 29245 | 29607 | Rv0025 | Rv0025 | 363 | 58892 | 59254 | BCG_0056 | BCG_0056 | 363 | 29225 | 29587 | BCGDan_0029 | BCGDan_0029 | 363 | 29225 | 29587 | BCGDan_sapMKO_0029 | BCGDan_sapMKO_0029 | 363 |
| 29722 | 31068 | Rv0026 | Rv0026 | 1347 | 59369 | 60802 | BCG_0057 | BCG_0057 | 1434 | 29702 | 31135 | BCGDan_0030 | BCGDan_0030 | 1434 | 29720 | 31135 | BCGDan_sapMKO_0030 | BCGDan_sapMKO_0030 | 1416 |
| 31189 | 31506 | Rv0027 | Rv0027 | 318 | 60840 | 61157 | BCG_0058 | BCG_0058 | 318 | 31173 | 31490 | BCGDan_0031 | BCGDan_0031 | 318 | 31173 | 31490 | BCGDan_sapMKO_0031 | BCGDan_sapMKO_0031 | 318 |
| 31514 | 31819 | Rv0028 | Rv0028 | 306 | 61165 | 61470 | BCG_0059 | BCG_0059 | 306 | 31498 | 31803 | BCGDan_0032 | BCGDan_0032 | 306 | 31498 | 31803 | BCGDan_sapMKO_0032 | BCGDan_sapMKO_0032 | 306 |
| 32057 | 33154 | Rv0029 | Rv0029 | 1098 | 61708 | 62805 | BCG_0060 | BCG_0060 | 1098 | 32041 | 33138 | BCGDan_0033 | BCGDan_0033 | 1098 | 32041 | 33138 | BCGDan_sapMKO_0033 | BCGDan_sapMKO_0033 | 1098 |
| 33224 | 33553 | Rv0030 | Rv0030 | 330 | 62875 | 63204 | BCG_0061 | BCG_0061 | 330 | 33208 | 33537 | BCGDan_0034 | BCGDan_0034 | 330 | 33208 | 33537 | BCGDan_sapMKO_0034 | BCGDan_sapMKO_0034 | 330 |
| 33582 | 33794 | Rv0031 | Rv0031 | 213 | 63233 | 63445 | BCG_0062 | BCG_0062 | 213 | 33566 | 33778 | BCGDan_0035 | BCGDan_0035 | 213 | 33566 | 33778 | BCGDan_sapMKO_0035 | BCGDan_sapMKO_0035 | 213 |
| 34295 | 36610 | bioF2 | Rv0032 | 2316 | 63946 | 66261 | bioF2 | BCG_0063 | 2316 | 34279 | 36594 | bioF2 | BCGDan_0036 | 2316 | 34279 | 36594 | bioF2 | BCGDan_sapMKO_0036 | 2316 |
| 36607 | 36870 | acpA | Rv0033 | 264 | 66258 | 66521 | acpA | BCG_0064 | 264 | 36591 | 36854 | acpA | BCGDan_0037 | 264 | 36591 | 36854 | acpA | BCGDan_sapMKO_0037 | 264 |
| 36867 | 37262 | Rv0034 | Rv0034 | 396 | 66518 | 66913 | BCG_0065 | BCG_0065 | 396 | 36851 | 37246 | BCGDan_0038 | BCGDan_0038 | 396 | 36851 | 37246 | BCGDan_sapMKO_0038 | BCGDan_sapMKO_0038 | 396 |
| 37259 | 38947 | fadD34 | Rv0035 | 1689 | 66910 | 68598 | fadD34 | BCG_0066 | 1689 | 37243 | 38931 | fadD34 | BCGDan_0039 | 1689 | 37243 | 38931 | fadD34 | BCGDan_sapMKO_0039 | 1689 |
| 39056 | 39829 | Rv0036c | Rv0036c | 774 | 68799 | 69479 | BCG_0067c | BCG_0067c | 681 | 39132 | 39812 | BCGDan_0040 | BCGDan_0040 | 681 | 39132 | 39818 | BCGDan_sapMKO_0040 | BCGDan_sapMKO_0040 | 687 |
| 39877 | 41202 | Rv0037c | Rv0037c | 1326 | 69527 | 70852 | BCG_0068c | BCG_0068c | 1326 | 39860 | 41185 | BCGDan_0041 | BCGDan_0041 | 1326 | 39860 | 41146 | BCGDan_sapMKO_0041 | BCGDan_sapMKO_0041 | 1287 |
| 41304 | 41912 | Rv0038 | Rv0038 | 609 | 70954 | 71562 | BCG_0069 | BCG_0069 | 609 | 41287 | 41895 | BCGDan_0042 | BCGDan_0042 | 609 | 41263 | 41895 | BCGDan_sapMKO_0042 | BCGDan_sapMKO_0042 | 633 |
| 42004 | 42351 | Rv0039c | Rv0039c | 348 | 71654 | 72001 | BCG_0070c | BCG_0070c | 348 | 41987 | 42334 | BCGDan_0043 | BCGDan_0043 | 348 | 41987 | 42334 | BCGDan_sapMKO_0043 | BCGDan_sapMKO_0043 | 348 |
| 42433 | 43365 | mtc28 | Rv0040c | 933 | 72083 | 73015 | mtc28 | BCG_0071c | 933 | 42416 | 43348 | mtc28 | BCGDan_0044 | 933 | 42416 | 43348 | mtc28 | BCGDan_sapMKO_0044 | 933 |
| 43562 | 46471 | leuS | Rv0041 | 2910 | 73212 | 76121 | leuS | BCG_0072 | 2910 | 43545 | 46454 | leuS | BCGDan_0045 | 2910 | 43578 | 46454 | leuS | BCGDan_sapMKO_0045 | 2877 |
| 46581 | 47207 | Rv0042c | Rv0042c | 627 | 76231 | 76857 | BCG_0073c | BCG_0073c | 627 | 46564 | 47190 | BCGDan_0046 | BCGDan_0046 | 627 | 46564 | 47190 | BCGDan_sapMKO_0046 | BCGDan_sapMKO_0046 | 627 |
| 47366 | 48100 | Rv0043c | Rv0043c | 735 | 77016 | 77750 | BCG_0074c | BCG_0074c | 735 | 47349 | 48083 | BCGDan_0047 | BCGDan_0047 | 735 | 47349 | 48083 | BCGDan_sapMKO_0047 | BCGDan_sapMKO_0047 | 735 |
| 48233 | 49027 | Rv0044c | Rv0044c | 795 | 77883 | 78677 | BCG_0075c | BCG_0075c | 795 | 48216 | 49010 | BCGDan_0048 | BCGDan_0048 | 795 | 48216 | 49010 | BCGDan_sapMKO_0048 | BCGDan_sapMKO_0048 | 795 |
| 49043 | 49939 | Rv0045c | Rv0045c | 897 | 78693 | 79589 | BCG_0076c | BCG_0076c | 897 | 49026 | 49922 | BCGDan_0049 | BCGDan_0049 | 897 | 49026 | 49922 | BCGDan_sapMKO_0049 | BCGDan_sapMKO_0049 | 897 |
| 50021 | 51124 | ino1 | Rv0046c | 1104 | 79671 | 80774 | ino1 | BCG_0077c | 1104 | 50004 | 51107 | ino1 | BCGDan_0050 | 1104 | 50004 | 51107 | ino1 | BCGDan_sapMKO_0050 | 1104 |
| 51185 | 51727 | Rv0047c | Rv0047c | 543 | 80835 | 81377 | BCG_0078c | BCG_0078c | 543 | 51168 | 51710 | BCGDan_0051 | BCGDan_0051 | 543 | 51168 | 51710 | BCGDan_sapMKO_0051 | BCGDan_sapMKO_0051 | 543 |
| 51828 | 52697 | Rv0048c | Rv0048c | 870 | 81478 | 82347 | BCG_0079c | BCG_0079c | 870 | 51811 | 52680 | BCGDan_0052 | BCGDan_0052 | 870 | 51811 | 52680 | BCGDan_sapMKO_0052 | BCGDan_sapMKO_0052 | 870 |
| 52831 | 53244 | Rv0049 | Rv0049 | 414 | 82481 | 82894 | BCG_0080 | BCG_0080 | 414 | 52814 | 53227 | BCGDan_0053 | BCGDan_0053 | 414 | 52793 | 53227 | BCGDan_sapMKO_0053 | BCGDan_sapMKO_0053 | 435 |
| 53663 | 55699 | ponA1 | Rv0050 | 2037 | 83313 | 85358 | ponA1 | BCG_0081 | 2046 | 53646 | 55691 | ponA1 | BCGDan_0054 | 2046 | 53337 | 55691 | ponA1 | BCGDan_sapMKO_0054 | 2355 |
| 55696 | 57378 | Rv0051 | Rv0051 | 1683 | 85355 | 87037 | BCG_0082 | BCG_0082 | 1683 | 55688 | 57370 | BCGDan_0055 | BCGDan_0055 | 1683 | 55688 | 57370 | BCGDan_sapMKO_0055 | BCGDan_sapMKO_0055 | 1683 |
| 57410 | 57973 | Rv0052 | Rv0052 | 564 | 87069 | 87632 | BCG_0083 | BCG_0083 | 564 | 57402 | 57965 | BCGDan_0056 | BCGDan_0056 | 564 | 57327 | 57965 | BCGDan_sapMKO_0056 | BCGDan_sapMKO_0056 | 639 |
| 58192 | 58482 | rpsF | Rv0053 | 291 | 87851 | 88141 | rpsF | BCG_0084 | 291 | 58184 | 58474 | rpsF | BCGDan_0057 | 291 | 58184 | 58474 | rpsF | BCGDan_sapMKO_0057 | 291 |
| 58586 | 59080 | ssb | Rv0054 | 495 | 88245 | 88739 | ssb | BCG_0085 | 495 | 58578 | 59072 | ssb | BCGDan_0058 | 495 | 58578 | 59072 | ssb | BCGDan_sapMKO_0058 | 495 |
| 59122 | 59376 | rpsR1 | Rv0055 | 255 | 88781 | 89035 | rpsR1 | BCG_0086 | 255 | 59114 | 59368 | rpsR1 | BCGDan_0059 | 255 | 59114 | 59368 | rpsR1 | BCGDan_sapMKO_0059 | 255 |
| 59409 | 59867 | rplI | Rv0056 | 459 | 89068 | 89526 | rplI | BCG_0087 | 459 | 59401 | 59859 | rplI | BCGDan_0060 | 459 | 59401 | 59859 | rplI | BCGDan_sapMKO_0060 | 459 |
| 59896 | 60417 | Rv0057 | Rv0057 | 522 | 89555 | 90076 | BCG_0088 | BCG_0088 | 522 | 59888 | 60409 | BCGDan_0061 | BCGDan_0061 | 522 | 59888 | 60409 | BCGDan_sapMKO_0061 | BCGDan_sapMKO_0061 | 522 |
| 60396 | 63020 | dnaB | Rv0058 | 2625 | 90055 | 92679 | dnaB | BCG_0089 | 2625 | 60388 | 63012 | dnaB | BCGDan_0062 | 2625 | 60418 | 63012 | dnaB | BCGDan_sapMKO_0062 | 2595 |
| 63200 | 63892 | Rv0059 | Rv0059 | 693 | 92859 | 93551 | BCG_0090 | BCG_0090 | 693 | 63192 | 63884 | BCGDan_0063 | BCGDan_0063 | 693 | 63192 | 63884 | BCGDan_sapMKO_0063 | BCGDan_sapMKO_0063 | 693 |
| 63909 | 64967 | Rv0060 | Rv0060 | 1059 | 93568 | 94626 | BCG_0091 | BCG_0091 | 1059 | 63901 | 64959 | BCGDan_0064 | BCGDan_0064 | 1059 | 63901 | 64959 | BCGDan_sapMKO_0064 | BCGDan_sapMKO_0064 | 1059 |
| NA | NA | NA | NA | NA | 94650 | 95075 | BCG_0092 | BCG_0092 | 426 | 64983 | 65408 | BCGDan_0065 | BCGDan_0065 | 426 | 64983 | 65408 | BCGDan_sapMKO_0065 | BCGDan_sapMKO_0065 | 426 |
| 65552 | 66694 | celA1 | Rv0062 | 1143 | 95211 | 96353 | celA1 | BCG_0093 | 1143 | 65544 | 66686 | celA1 | BCGDan_0066 | 1143 | 65544 | 66686 | celA1 | BCGDan_sapMKO_0066 | 1143 |
| 66923 | 68362 | Rv0063 | Rv0063 | 1440 | 96582 | 98021 | BCG_0094 | BCG_0094 | 1440 | 66915 | 68354 | BCGDan_0067 | BCGDan_0067 | 1440 | 66915 | 68354 | BCGDan_sapMKO_0067 | BCGDan_sapMKO_0067 | 1440 |
| 68620 | 71559 | Rv0064 | Rv0064 | 2940 | 98279 | 101218 | BCG_0095 | BCG_0095 | 2940 | 68612 | 71551 | BCGDan_0068 | BCGDan_0068 | 2940 | 68612 | 71551 | BCGDan_sapMKO_0068 | BCGDan_sapMKO_0068 | 2940 |
| 71589 | 71828 | vapB1 | Rv0064A | 240 | 101285 | 101524 | BCG_0095A | BCG_0095A | 240 | 71618 | 71857 | BCGDan_0069 | BCGDan_0069 | 240 | 71618 | 71857 | BCGDan_sapMKO_0069 | BCGDan_sapMKO_0069 | 240 |
| 71821 | 72222 | vapC1 | Rv0065 | 402 | 101517 | 101918 | BCG_0096 | BCG_0096 | 402 | 71850 | 72251 | BCGDan_0070 | BCGDan_0070 | 402 | 71850 | 72251 | BCGDan_sapMKO_0070 | BCGDan_sapMKO_0070 | 402 |
| 72274 | 74511 | icd2 | Rv0066c | 2238 | 101970 | 104207 | icd2 | BCG_0097c | 2238 | 72303 | 74540 | icd2 | BCGDan_0071 | 2238 | 72303 | 74540 | icd2 | BCGDan_sapMKO_0071 | 2238 |
| 74629 | 75198 | Rv0067c | Rv0067c | 570 | 104325 | 104894 | BCG_0098c | BCG_0098c | 570 | 74658 | 75227 | BCGDan_0072 | BCGDan_0072 | 570 | 74658 | 75206 | BCGDan_sapMKO_0072 | BCGDan_sapMKO_0072 | 549 |
| 75301 | 76212 | Rv0068 | Rv0068 | 912 | 104997 | 105908 | BCG_0099 | BCG_0099 | 912 | 75330 | 76241 | BCGDan_0073 | BCGDan_0073 | 912 | 75330 | 76241 | BCGDan_sapMKO_0073 | BCGDan_sapMKO_0073 | 912 |
| 76237 | 77622 | sdaA | Rv0069c | 1386 | 105933 | 107318 | sdaA | BCG_0100c | 1386 | 76266 | 77651 | sdaA | BCGDan_0074 | 1386 | 76266 | 77651 | sdaA | BCGDan_sapMKO_0074 | 1386 |
| 77619 | 78896 | glyA2 | Rv0070c | 1278 | 107315 | 108592 | glyA2 | BCG_0101c | 1278 | 77648 | 78925 | glyA2 | BCGDan_0075 | 1278 | 77648 | 78925 | glyA2 | BCGDan_sapMKO_0075 | 1278 |
| 79486 | 80193 | Rv0071 | Rv0071 | 708 | 109182 | 109934 | BCG_0102 | BCG_0102 | 753 | 79515 | 80267 | BCGDan_0076 | BCGDan_0076 | 753 | 79515 | 80267 | BCGDan_sapMKO_0076 | BCGDan_sapMKO_0076 | 753 |
| 80624 | 81673 | Rv0072 | Rv0072 | 1050 | 110365 | 111414 | BCG_0103 | BCG_0103 | 1050 | 80698 | 81747 | BCGDan_0077 | BCGDan_0077 | 1050 | 80698 | 81747 | BCGDan_sapMKO_0077 | BCGDan_sapMKO_0077 | 1050 |
| 81676 | 82668 | Rv0073 | Rv0073 | 993 | 111417 | 112409 | BCG_0104 | BCG_0104 | 993 | 81750 | 82742 | BCGDan_0078 | BCGDan_0078 | 993 | 81750 | 82742 | BCGDan_sapMKO_0078 | BCGDan_sapMKO_0078 | 993 |
| 82748 | 83983 | Rv0074 | Rv0074 | 1236 | 112489 | 113724 | BCG_0105 | BCG_0105 | 1236 | 82822 | 84057 | BCGDan_0079 | BCGDan_0079 | 1236 | 82888 | 84057 | BCGDan_sapMKO_0079 | BCGDan_sapMKO_0079 | 1170 |
| 83996 | 85168 | Rv0075 | Rv0075 | 1173 | 113737 | 114909 | BCG_0106 | BCG_0106 | 1173 | 84070 | 85242 | BCGDan_0080 | BCGDan_0080 | 1173 | 84070 | 85242 | BCGDan_sapMKO_0080 | BCGDan_sapMKO_0080 | 1173 |
| 85183 | 85572 | Rv0076c | Rv0076c | 390 | 114924 | 115313 | BCG_0107c | BCG_0107c | 390 | 85257 | 85646 | BCGDan_0081 | BCGDan_0081 | 390 | 85257 | 85646 | BCGDan_sapMKO_0081 | BCGDan_sapMKO_0081 | 390 |
| 85636 | 86466 | Rv0077c | Rv0077c | 831 | 115377 | 116207 | BCG_0108c | BCG_0108c | 831 | 85710 | 86540 | BCGDan_0082 | BCGDan_0082 | 831 | 85710 | 86564 | BCGDan_sapMKO_0082 | BCGDan_sapMKO_0082 | 855 |
| 86528 | 87133 | Rv0078 | Rv0078 | 606 | 116269 | 116874 | BCG_0109 | BCG_0109 | 606 | 86602 | 87207 | BCGDan_0083 | BCGDan_0083 | 606 | 86602 | 87207 | BCGDan_sapMKO_0083 | BCGDan_sapMKO_0083 | 606 |
| 87208 | 87801 | Rv0078A | Rv0078A | 594 | 116949 | 117542 | BCG_0110c | BCG_0110c | 594 | 87282 | 87875 | BCGDan_0084 | BCGDan_0084 | 594 | 87282 | 87875 | BCGDan_sapMKO_0084 | BCGDan_sapMKO_0084 | 594 |

**Legend**

| | |
|---|---|
| | DU1-Danish |
| | DU2-III in BCG Danish |
| | DU1-Pasteur |
| | DU2-IV in BCG Pasteur |
| | *M. tb* gene is split in BCG |
| | truncated *sapM* |
| NA | not applicable |

**Venn diagrams showing the shared and unique CDS**

among *M. tb* H37Rv, BCG Pasteur and BCG Danish genome

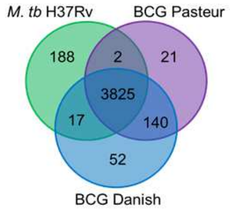


among Danish WT and *sapM* KO genome

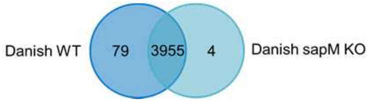


**Remark** Annotation of the *sapM* KO was done using an automatic gene prediction pipeline. The assigned annotations were not manually curated. Resulting in small differences in annotation with its parental strain BCG Danish 1331 (07/270) (the annotation for this strain was manually curated) causing incomplete mapping of the derived ortholog information for the KO. 3 genetic differences were noted between the parental strain and the *sapM* KO, namely the truncated *sapM* gene and the deleted DU2, and a missense SNP in BCG3966 or BCGDan4053.

| | | | | | | | | | | | | | | | | | | | |
|---|---|---|---|---|---|---|---|---|---|---|---|---|---|---|---|---|---|---|---|
| 87798 | 88004 | Rv0078B | Rv0078B | 207 | 117539 | 117751 | BCG_0111c | BCG_0111c | 213 | 87872 | 88078 | BCGDan_0085 | BCGDan_0085 | 207 | 87872 | 88078 | BCGDan_sapMKO_0085 | BCGDan_sapMKO_0085 | 207 |
| 88204 | 89025 | Rv0079 | Rv0079 | 822 | 117945 | 118766 | BCG_0112 | BCG_0112 | 822 | 88278 | 89099 | BCGDan_0086 | BCGDan_0086 | 822 | 88278 | 89099 | BCGDan_sapMKO_0086 | BCGDan_sapMKO_0086 | 822 |
| 89022 | 89480 | Rv0080 | Rv0080 | 459 | 118763 | 119221 | BCG_0113 | BCG_0113 | 459 | 89096 | 89554 | BCGDan_0087 | BCGDan_0087 | 459 | 89096 | 89554 | BCGDan_sapMKO_0087 | BCGDan_sapMKO_0087 | 459 |
| 89575 | 89919 | Rv0081 | Rv0081 | 345 | 119316 | 119660 | BCG_0114 | BCG_0114 | 345 | 89649 | 89993 | BCGDan_0088 | BCGDan_0088 | 345 | 89649 | 89993 | BCGDan_sapMKO_0088 | BCGDan_sapMKO_0088 | 345 |
| 89924 | 90403 | Rv0082 | Rv0082 | 480 | 119665 | 120144 | BCG_0115 | BCG_0115 | 480 | 89998 | 90477 | BCGDan_0089 | BCGDan_0089 | 480 | 89998 | 90477 | BCGDan_sapMKO_0089 | BCGDan_sapMKO_0089 | 480 |
| 90400 | 92322 | Rv0083 | Rv0083 | 1923 | 120141 | 122069 | BCG_0116 | BCG_0116 | 1929 | 90474 | 92402 | BCGDan_0090 | BCGDan_0090 | 1929 | 90474 | 92402 | BCGDan_sapMKO_0090 | BCGDan_sapMKO_0090 | 1929 |
| 92328 | 93278 | hycD | Rv0084 | 951 | 122069 | 123019 | hycD | BCG_0117 | 951 | 92402 | 93352 | hycD | BCGDan_0091 | 951 | 92402 | 93352 | hycD | BCGDan_sapMKO_0091 | 951 |
| 93289 | 93951 | hycP | Rv0085 | 663 | 123030 | 123692 | hycP | BCG_0118 | 663 | 93363 | 94025 | hycP | BCGDan_0092 | 663 | 93363 | 94025 | hycP | BCGDan_sapMKO_0092 | 663 |
| 93951 | 95417 | hycQ | Rv0086 | 1467 | 123692 | 125158 | hycQ | BCG_0119 | 1467 | 94025 | 95491 | hycQ | BCGDan_0093 | 1467 | 94025 | 95491 | hycQ | BCGDan_sapMKO_0093 | 1467 |
| 95414 | 96892 | hycE | Rv0087 | 1479 | 125155 | 126633 | hycE | BCG_0120 | 1479 | 95488 | 96966 | hycE | BCGDan_0094 | 1479 | 95491 | 96966 | hycE | BCGDan_sapMKO_0094 | 1476 |
| 96927 | 97601 | Rv0088 | Rv0088 | 675 | 126668 | 127342 | BCG_0121 | BCG_0121 | 675 | 97001 | 97675 | BCGDan_0095 | BCGDan_0095 | 675 | 97097 | 97675 | BCGDan_sapMKO_0095 | BCGDan_sapMKO_0095 | 579 |
| 97758 | 98351 | Rv0089 | Rv0089 | 594 | 127499 | 128092 | BCG_0122 | BCG_0122 | 594 | 97832 | 98425 | BCGDan_0096 | BCGDan_0096 | 594 | 97832 | 98425 | BCGDan_sapMKO_0096 | BCGDan_sapMKO_0096 | 594 |
| 98480 | 99250 | Rv0090 | Rv0090 | 771 | 128221 | 128991 | BCG_0123 | BCG_0123 | 771 | 98554 | 99324 | BCGDan_0097 | BCGDan_0097 | 771 | 98554 | 99324 | BCGDan_sapMKO_0097 | BCGDan_sapMKO_0097 | 771 |
| 99684 | 100451 | mtn | Rv0091 | 768 | 129425 | 130192 | mtn | BCG_0124 | 768 | 99758 | 100525 | mtn | BCGDan_0098 | 768 | 99758 | 100525 | mtn | BCGDan_sapMKO_0098 | 768 |
| 100583 | 102868 | ctpA | Rv0092 | 2286 | 130324 | 132609 | ctpA | BCG_0125 | 2286 | 100657 | 102942 | ctpA | BCGDan_0099 | 2286 | 100750 | 102942 | ctpA | BCGDan_sapMKO_0099 | 2193 |
| 102815 | 103663 | Rv0093c | Rv0093c | 849 | 132556 | 133404 | BCG_0126c | BCG_0126c | 849 | 102889 | 103737 | BCGDan_0100 | BCGDan_0100 | 849 | 102889 | 103737 | BCGDan_sapMKO_0100 | BCGDan_sapMKO_0100 | 849 |
| 103710 | 104663 | Rv0094c | Rv0094c | 954 | 133451 | 134404 | BCG_0127c | BCG_0127c | 954 | 103784 | 104737 | BCGDan_0101 | BCGDan_0101 | 954 | 103784 | 104743 | BCGDan_sapMKO_0101 | BCGDan_sapMKO_0101 | 960 |
| 104805 | 105215 | Rv0095c | Rv0095c | 411 | 134175 | 134957 | BCG_0128c | BCG_0128c | 783 | 104508 | 105290 | BCGDan_0102 | BCGDan_0102 | 783 | 104508 | 105290 | BCGDan_sapMKO_0102 | BCGDan_sapMKO_0102 | 783 |
| 105324 | 106715 | PPE1 | Rv0096 | 1392 | 135066 | 136457 | PPE1 | BCG_0129 | 1392 | 105399 | 106790 | PPE1 | BCGDan_0103 | 1392 | 105417 | 106790 | PPE1 | BCGDan_sapMKO_0103 | 1374 |
| 106734 | 107603 | Rv0097 | Rv0097 | 870 | 136476 | 137345 | BCG_0130 | BCG_0130 | 870 | 106809 | 107678 | BCGDan_0104 | BCGDan_0104 | 870 | 106809 | 107678 | BCGDan_sapMKO_0104 | BCGDan_sapMKO_0104 | 870 |
| 107600 | 108151 | fcoT | Rv0098 | 552 | 137342 | 137893 | BCG_0131 | BCG_0131 | 552 | 107675 | 108226 | BCGDan_0105 | BCGDan_0105 | 552 | 107675 | 108226 | BCGDan_sapMKO_0105 | BCGDan_sapMKO_0105 | 552 |
| 108156 | 109778 | fadD10 | Rv0099 | 1623 | 137898 | 139520 | fadD10 | BCG_0132 | 1623 | 108231 | 109853 | fadD10 | BCGDan_0106 | 1623 | 108276 | 109853 | fadD10 | BCGDan_sapMKO_0106 | 1578 |
| 109783 | 110019 | Rv0100 | Rv0100 | 237 | 139525 | 139761 | BCG_0133 | BCG_0133 | 237 | 109858 | 110094 | BCGDan_0107 | BCGDan_0107 | 237 | 109846 | 110094 | BCGDan_sapMKO_0107 | BCGDan_sapMKO_0107 | 249 |
| 110001 | 117539 | nrp | Rv0101 | 7539 | 139743 | 147281 | nrp | BCG_0134 | 7539 | 110076 | 117614 | nrp | BCGDan_0108 | 7539 | 110124 | 117614 | nrp | BCGDan_sapMKO_0108 | 7491 |
| 117714 | 119699 | Rv0102 | Rv0102 | 1986 | 147456 | 149441 | BCG_0135 | BCG_0135 | 1986 | 117789 | 119774 | BCGDan_0109 | BCGDan_0109 | 1986 | 117852 | 119774 | BCGDan_sapMKO_0109 | BCGDan_sapMKO_0109 | 1923 |
| 119915 | 122173 | ctpB | Rv0103c | 2259 | 149657 | 151915 | ctpB | BCG_0136c | 2259 | 119990 | 122248 | ctpB | BCGDan_0110 | 2259 | 119990 | 122233 | ctpB | BCGDan_sapMKO_0110 | 2244 |
| 122317 | 123831 | Rv0104 | Rv0104 | 1515 | 152059 | 153573 | BCG_0137 | BCG_0137 | 1515 | 122392 | 123906 | BCGDan_0111 | BCGDan_0111 | 1515 | 122419 | 123906 | BCGDan_sapMKO_0111 | BCGDan_sapMKO_0111 | 1488 |
| 123980 | 124264 | rpmB1 | Rv0105c | 285 | 153722 | 154006 | rpmB1 | BCG_0138c | 285 | 124055 | 124339 | rpmB1 | BCGDan_0112 | 285 | 124055 | 124339 | rpmB1 | BCGDan_sapMKO_0112 | 285 |
| 124374 | 125570 | Rv0106 | Rv0106 | 1197 | 154116 | 155312 | BCG_0139 | BCG_0139 | 1197 | 124449 | 125645 | BCGDan_0113 | BCGDan_0113 | 1197 | 124449 | 125645 | BCGDan_sapMKO_0113 | BCGDan_sapMKO_0113 | 1197 |
| 125643 | 130541 | ctpI | Rv0107c | 4899 | 155407 | 160284 | ctpI | BCG_0140c | 4878 | 125740 | 130617 | ctpI | BCGDan_0114 | 4878 | 125740 | 130617 | ctpI | BCGDan_sapMKO_0114 | 4878 |
| 130895 | 131104 | Rv0108c | Rv0108c | 210 | 160638 | 160847 | BCG_0141c | BCG_0141c | 210 | 130971 | 131180 | BCGDan_0115 | BCGDan_0115 | 210 | 130971 | 131213 | BCGDan_sapMKO_0115 | BCGDan_sapMKO_0115 | 243 |
| 131382 | 132872 | PE_PGRS1 | Rv0109 | 1491 | 161126 | 162616 | PE_PGRS1 | BCG_0142 | 1491 | 131459 | 132949 | PE_PGRS1 | BCGDan_0116 | 1491 | 131459 | 132949 | PE_PGRS1 | BCGDan_sapMKO_0116 | 1491 |
| 133020 | 133769 | Rv0110 | Rv0110 | 750 | 162764 | 163513 | BCG_0143 | BCG_0143 | 750 | 133097 | 133846 | BCGDan_0117 | BCGDan_0117 | 750 | 133208 | 133846 | BCGDan_sapMKO_0117 | BCGDan_sapMKO_0117 | 639 |
| 133950 | 136007 | Rv0111 | Rv0111 | 2058 | 163694 | 165751 | BCG_0144 | BCG_0144 | 2058 | 134060 | 136084 | BCGDan_0118 | BCGDan_0118 | 2025 | 134060 | 136084 | BCGDan_sapMKO_0118 | BCGDan_sapMKO_0118 | 2025 |
| 136289 | 137245 | gca | Rv0112 | 957 | 166033 | 166989 | gca | BCG_0145 | 957 | 136366 | 137322 | gca | BCGDan_0119 | 957 | 136366 | 137322 | gca | BCGDan_sapMKO_0119 | 957 |
| 137319 | 137909 | gmhA | Rv0113 | 591 | 167063 | 167653 | gmhA | BCG_0146 | 591 | 137396 | 137986 | gmhA | BCGDan_0120 | 591 | 137396 | 137986 | gmhA | BCGDan_sapMKO_0120 | 591 |
| 137941 | 138513 | gmhB | Rv0114 | 573 | 167685 | 168257 | BCG_0147 | BCG_0147 | 573 | 138018 | 138590 | BCGDan_0121 | BCGDan_0121 | 573 | 138087 | 138590 | BCGDan_sapMKO_0121 | BCGDan_sapMKO_0121 | 504 |
| 138513 | 139673 | hddA | Rv0115 | 1161 | 168257 | 169324 | BCG_0149 | BCG_0149 | 1068 | 138590 | 139657 | BCGDan_0122 | BCGDan_0122 | 1068 | 138590 | 139657 | BCGDan_sapMKO_0122 | BCGDan_sapMKO_0122 | 1068 |
| 140267 | 141022 | ldtA | Rv0116c | 756 | 170012 | 170767 | BCG_0150c | BCG_0150c | 756 | 140345 | 141100 | BCGDan_0123 | BCGDan_0123 | 756 | 140345 | 141109 | BCGDan_sapMKO_0123 | BCGDan_sapMKO_0123 | 765 |
| 141200 | 142144 | oxyS | Rv0117 | 945 | 170945 | 171889 | oxyS | BCG_0151 | 945 | 141278 | 142222 | oxyS | BCGDan_0124 | 945 | 141272 | 142222 | oxyS | BCGDan_sapMKO_0124 | 951 |
| 142128 | 143876 | oxcA | Rv0118c | 1749 | 171873 | 173621 | oxcA | BCG_0152c | 1749 | 142206 | 143954 | oxcA | BCGDan_0125 | 1749 | 142206 | 143924 | oxcA | BCGDan_sapMKO_0125 | 1719 |
| 144049 | 145626 | fadD7 | Rv0119 | 1578 | 173794 | 175371 | fadD7 | BCG_0153 | 1578 | 144127 | 145704 | fadD7 | BCGDan_0126 | 1578 | 144076 | 145704 | fadD7 | BCGDan_sapMKO_0126 | 1629 |
| 145627 | 147771 | fusA2 | Rv0120c | 2145 | 175372 | 177516 | fusA2b | BCG_0154c | 2145 | 145705 | 147849 | fusA2b | BCGDan_0127 | 2145 | 145705 | 147849 | fusA2b | BCGDan_sapMKO_0127 | 2145 |
| 147908 | 148342 | Rv0121c | Rv0121c | 435 | 177652 | 178086 | BCG_0155c | BCG_0155c | 435 | 147985 | 148419 | BCGDan_0128 | BCGDan_0128 | 435 | 147985 | 148419 | BCGDan_sapMKO_0128 | BCGDan_sapMKO_0128 | 435 |
| 148491 | 148859 | Rv0122 | Rv0122 | 369 | 178235 | 178603 | BCG_0156 | BCG_0156 | 369 | 148568 | 148936 | BCGDan_0129 | BCGDan_0129 | 369 | 148568 | 148936 | BCGDan_sapMKO_0129 | BCGDan_sapMKO_0129 | 369 |
| 148856 | 149224 | Rv0123 | Rv0123 | 369 | 178600 | 178968 | BCG_0157 | BCG_0157 | 369 | 148933 | 149301 | BCGDan_0130 | BCGDan_0130 | 369 | 148933 | 149301 | BCGDan_sapMKO_0130 | BCGDan_sapMKO_0130 | 369 |
| 149533 | 150996 | PE_PGRS2 | Rv0124 | 1464 | 179277 | 180893 | PE_PGRS2 | BCG_0158 | 1617 | 149610 | 151226 | PE_PGRS2 | BCGDan_0131 | 1617 | 149610 | 151226 | PE_PGRS2 | BCGDan_sapMKO_0131 | 1617 |
| 151148 | 152215 | pepA | Rv0125 | 1068 | 181045 | 182112 | pepA | BCG_0159 | 1068 | 151378 | 152445 | pepA | BCGDan_0132 | 1068 | 151378 | 152445 | pepA | BCGDan_sapMKO_0132 | 1068 |
| 152324 | 154129 | treS | Rv0126 | 1806 | 182221 | 184026 | treS | BCG_0160 | 1806 | 152554 | 154359 | treS | BCGDan_0133 | 1806 | 152554 | 154359 | treS | BCGDan_sapMKO_0133 | 1806 |
| 154232 | 155599 | mak | Rv0127 | 1368 | 184129 | 185496 | BCG_0161 | BCG_0161 | 1368 | 154462 | 155829 | BCGDan_0134 | BCGDan_0134 | 1368 | 154462 | 155829 | BCGDan_sapMKO_0134 | BCGDan_sapMKO_0134 | 1368 |
| 155667 | 156446 | Rv0128 | Rv0128 | 780 | 185564 | 186343 | BCG_0162 | BCG_0162 | 780 | 155897 | 156676 | BCGDan_0135 | BCGDan_0135 | 780 | 155897 | 156676 | BCGDan_sapMKO_0135 | BCGDan_sapMKO_0135 | 780 |
| 156578 | 157600 | fbpC | Rv0129c | 1023 | 186475 | 187497 | fbpC | BCG_0163c | 1023 | 156808 | 157830 | fbpC | BCGDan_0136 | 1023 | 156808 | 157830 | fbpC | BCGDan_sapMKO_0136 | 1023 |
| 157847 | 158302 | htdZ | Rv0130 | 456 | 187744 | 188199 | BCG_0164 | BCG_0164 | 456 | 158077 | 158532 | BCGDan_0137 | BCGDan_0137 | 456 | 158077 | 158532 | BCGDan_sapMKO_0137 | BCGDan_sapMKO_0137 | 456 |
| 158315 | 159658 | fadE1 | Rv0131c | 1344 | 188212 | 189555 | fadE1 | BCG_0165c | 1344 | 158545 | 159888 | fadE1 | BCGDan_0138 | 1344 | 158545 | 159888 | fadE1 | BCGDan_sapMKO_0138 | 1344 |
| 159700 | 160782 | fgd2 | Rv0132c | 1083 | 189597 | 190679 | fgd2 | BCG_0166c | 1083 | 159930 | 161012 | fgd2 | BCGDan_0139 | 1083 | 159930 | 161012 | fgd2 | BCGDan_sapMKO_0139 | 1083 |
| 160869 | 161474 | Rv0133 | Rv0133 | 606 | 190766 | 191248 | BCG_0167 | BCG_0167 | 483 | 161099 | 161581 | BCGDan_0140 | BCGDan_0140 | 483 | 161099 | 161581 | BCGDan_sapMKO_0140 | BCGDan_sapMKO_0140 | 483 |
| 161771 | 162673 | ephF | Rv0134 | 903 | 191668 | 192051 | BCG_0168 | BCG_0168 | 384 | 162001 | 162384 | BCGDan_0141 | BCGDan_0141 | 384 | 162001 | 162384 | BCGDan_sapMKO_0141 | BCGDan_sapMKO_0141 | 384 |
| NA | NA | NA | NA | NA | 192051 | 192158 | BCG_0169 | BCG_0169 | 108 | 162384 | 162491 | BCGDan_0142 | BCGDan_0142 | 108 | 162384 | 162491 | BCGDan_sapMKO_0142 | BCGDan_sapMKO_0142 | 108 |
| NA | NA | NA | NA | NA | 192160 | 192570 | BCG_0170 | BCG_0170 | 411 | 162493 | 162903 | BCGDan_0143 | BCGDan_0143 | 411 | 162493 | 162900 | BCGDan_sapMKO_0143 | BCGDan_sapMKO_0143 | 408 |
| 162644 | 163249 | Rv0135c | Rv0135c | 606 | 192541 | 193146 | BCG_0171c | BCG_0171c | 606 | 162874 | 163479 | BCGDan_0144 | BCGDan_0144 | 606 | 162874 | 163479 | BCGDan_sapMKO_0144 | BCGDan_sapMKO_0144 | 606 |
| 163366 | 164691 | cyp138 | Rv0136 | 1326 | 193263 | 194588 | cyp138 | BCG_0172 | 1326 | 163596 | 164921 | cyp138 | BCGDan_0145 | 1326 | 163596 | 164921 | cyp138 | BCGDan_sapMKO_0145 | 1326 |
| 164712 | 165260 | msrA | Rv0137c | 549 | 194609 | 195157 | msrA | BCG_0173c | 549 | 164942 | 165490 | msrA | BCGDan_0146 | 549 | 164942 | 165490 | msrA | BCGDan_sapMKO_0146 | 549 |
| 165323 | 165826 | Rv0138 | Rv0138 | 504 | 195220 | 195723 | BCG_0174 | BCG_0174 | 504 | 165553 | 166056 | BCGDan_0147 | BCGDan_0147 | 504 | 165553 | 166056 | BCGDan_sapMKO_0147 | BCGDan_sapMKO_0147 | 504 |
| 165827 | 166849 | Rv0139 | Rv0139 | 1023 | 195724 | 196746 | BCG_0175 | BCG_0175 | 1023 | 166057 | 167079 | BCGDan_0148 | BCGDan_0148 | 1023 | 166057 | 167079 | BCGDan_sapMKO_0148 | BCGDan_sapMKO_0148 | 1023 |
| 166910 | 167290 | Rv0140 | Rv0140 | 381 | 196807 | 197187 | BCG_0176 | BCG_0176 | 381 | 167140 | 167520 | BCGDan_0149 | BCGDan_0149 | 381 | 167140 | 167520 | BCGDan_sapMKO_0149 | BCGDan_sapMKO_0149 | 381 |
| 167271 | 167681 | Rv0141c | Rv0141c | 411 | 197168 | 197578 | BCG_0177c | BCG_0177c | 411 | 167501 | 167911 | BCGDan_0150 | BCGDan_0150 | 411 | 167501 | 167911 | BCGDan_sapMKO_0150 | BCGDan_sapMKO_0150 | 411 |
| 167711 | 168637 | Rv0142 | Rv0142 | 927 | 197608 | 198534 | BCG_0178 | BCG_0178 | 927 | 167941 | 168867 | BCGDan_0151 | BCGDan_0151 | 927 | 167956 | 168867 | BCGDan_sapMKO_0151 | BCGDan_sapMKO_0151 | 912 |
| 168704 | 170182 | Rv0143c | Rv0143c | 1479 | 198601 | 200079 | BCG_0179c | BCG_0179c | 1479 | 168934 | 170412 | BCGDan_0152 | BCGDan_0152 | 1479 | 168934 | 170412 | BCGDan_sapMKO_0152 | BCGDan_sapMKO_0152 | 1479 |
| 170284 | 171126 | Rv0144 | Rv0144 | 843 | 200181 | 201023 | BCG_0180 | BCG_0180 | 843 | 170514 | 171356 | BCGDan_0153 | BCGDan_0153 | 843 | 170514 | 171356 | BCGDan_sapMKO_0153 | BCGDan_sapMKO_0153 | 843 |
| 171215 | 172168 | Rv0145 | Rv0145 | 954 | 201112 | 202065 | BCG_0181 | BCG_0181 | 954 | 171445 | 172398 | BCGDan_0154 | BCGDan_0154 | 954 | 171445 | 172398 | BCGDan_sapMKO_0154 | BCGDan_sapMKO_0154 | 954 |
| 172211 | 173143 | Rv0146 | Rv0146 | 933 | 202108 | 203040 | BCG_0182 | BCG_0182 | 933 | 172441 | 173373 | BCGDan_0155 | BCGDan_0155 | 933 | 172432 | 173373 | BCGDan_sapMKO_0155 | BCGDan_sapMKO_0155 | 942 |
| 173238 | 174758 | Rv0147 | Rv0147 | 1521 | 203135 | 204655 | BCG_0183 | BCG_0183 | 1521 | 173468 | 174988 | BCGDan_0156 | BCGDan_0156 | 1521 | 173468 | 174988 | BCGDan_sapMKO_0156 | BCGDan_sapMKO_0156 | 1521 |
| 174833 | 175693 | Rv0148 | Rv0148 | 861 | 204730 | 205590 | BCG_0184 | BCG_0184 | 861 | 175063 | 175923 | BCGDan_0157 | BCGDan_0157 | 861 | 175063 | 175923 | BCGDan_sapMKO_0157 | BCGDan_sapMKO_0157 | 861 |
| 175700 | 176668 | Rv0149 | Rv0149 | 969 | 205597 | 206565 | BCG_0185 | BCG_0185 | 969 | 175930 | 176898 | BCGDan_0158 | BCGDan_0158 | 969 | 175972 | 176898 | BCGDan_sapMKO_0158 | BCGDan_sapMKO_0158 | 927 |
| 176665 | 176952 | Rv0150c | Rv0150c | 288 | 206562 | 206849 | BCG_0186c | BCG_0186c | 288 | 176895 | 177182 | BCGDan_0159 | BCGDan_0159 | 288 | 176895 | 177218 | BCGDan_sapMKO_0159 | BCGDan_sapMKO_0159 | 324 |
| 177543 | 179309 | PE1 | Rv0151c | 1767 | 207440 | 209206 | PE1 | BCG_0187c | 1767 | 177773 | 179539 | PE1 | BCGDan_0160 | 1767 | 177773 | 179455 | PE1 | BCGDan_sapMKO_0160 | 1683 |
| 179319 | 180896 | PE2 | Rv0152c | 1578 | 209216 | 210793 | PE2 | BCG_0188c | 1578 | 179549 | 181126 | PE2 | BCGDan_0161 | 1578 | 179549 | 181021 | PE2 | BCGDan_sapMKO_0161 | 1473 |
| 181155 | 181985 | ptbB | Rv0153c | 831 | 211052 | 211882 | ptbB | BCG_0189c | 831 | 181385 | 182215 | ptbB | BCGDan_0162 | 831 | 181385 | 182215 | ptbB | BCGDan_sapMKO_0162 | 831 |
| 181987 | 183198 | fadE2 | Rv0154c | 1212 | 211884 | 213095 | fadE2 | BCG_0190c | 1212 | 182217 | 183428 | fadE2 | BCGDan_0163 | 1212 | 182217 | 183440 | fadE2 | BCGDan_sapMKO_0163 | 1224 |
| 183622 | 184722 | pntAa | Rv0155 | 1101 | 213519 | 214619 | pntAa | BCG_0191 | 1101 | 183852 | 184952 | pntAa | BCGDan_0164 | 1101 | 183852 | 184952 | pntAa | BCGDan_sapMKO_0164 | 1101 |
| 184723 | 185055 | pntAb | Rv0156 | 333 | 214620 | 214952 | pntAb | BCG_0192 | 333 | 184953 | 185285 | pntAb | BCGDan_0165 | 333 | 184953 | 185285 | pntAb | BCGDan_sapMKO_0165 | 333 |
| 185052 | 186479 | pntB | Rv0157 | 1428 | 214949 | 216376 | pntB | BCG_0193 | 1428 | 185282 | 186709 | pntB | BCGDan_0166 | 1428 | 185282 | 186709 | pntB | BCGDan_sapMKO_0166 | 1428 |
| 186785 | 187429 | Rv0158 | Rv0158 | 645 | 216682 | 217326 | BCG_0194 | BCG_0194 | 645 | 187015 | 187659 | BCGDan_0167 | BCGDan_0167 | 645 | 187015 | 187659 | BCGDan_sapMKO_0167 | BCGDan_sapMKO_0167 | 645 |
| 187433 | 188839 | PE3 | Rv0159c | 1407 | 217330 | 218736 | PE3 | BCG_0195c | 1407 | 187663 | 189069 | PE3 | BCGDan_0168 | 1407 | 187663 | 189069 | PE3 | BCGDan_sapMKO_0168 | 1407 |
| 188931 | 190439 | PE4 | Rv0160c | 1509 | 218828 | 220336 | PE4 | BCG_0196c | 1509 | 189161 | 190669 | PE4 | BCGDan_0169 | 1509 | 189161 | 190669 | PE4 | BCGDan_sapMKO_0169 | 1509 |
| 190607 | 191956 | Rv0161 | Rv0161 | 1350 | 220504 | 221853 | BCG_0197 | BCG_0197 | 1350 | 190837 | 192186 | BCGDan_0170 | BCGDan_0170 | 1350 | 190813 | 192186 | BCGDan_sapMKO_0170 | BCGDan_sapMKO_0170 | 1374 |

| | | | | | | | | | | | | | | | | | | | |
|---|---|---|---|---|---|---|---|---|---|---|---|---|---|---|---|---|---|---|---|
| 191984 | 193135 | adhE1 | Rv0162c | 1152 | 221881 | 223032 | adhE1 | BCG_0198c | 1152 | 192214 | 193365 | adhE1 | BCGDan_0171 | 1152 | 192214 | 193344 | adhE1 | BCGDan_sapMKO_0171 | 1131 |
| 193117 | 193572 | Rv0163 | Rv0163 | 456 | 223014 | 223469 | BCG_0199 | BCG_0199 | 456 | 193347 | 193802 | BCGDan_0172 | BCGDan_0172 | 456 | 193347 | 193802 | BCGDan_sapMKO_0172 | BCGDan_sapMKO_0172 | 456 |
| 193626 | 194111 | TB18.5 | Rv0164 | 486 | 223523 | 224008 | TB18.5 | BCG_0200 | 486 | 193856 | 194341 | TB18.5 | BCGDan_0173 | 486 | 193901 | 194341 | TB18.5 | BCGDan_sapMKO_0173 | 441 |
| 194144 | 194815 | mce1R | Rv0165c | 672 | 224022 | 224837 | BCG_0201c | BCG_0201c | 816 | 194355 | 195170 | BCGDan_0174 | BCGDan_0174 | 816 | 194355 | 195170 | BCGDan_sapMKO_0174 | BCGDan_sapMKO_0174 | 816 |
| NA | NA | NA | NA | NA | 224041 | 224220 | BCG_0202c | BCG_0202c | 180 | 194374 | 194553 | BCGDan_0175 | BCGDan_0175 | 180 | 194377 | 194553 | BCGDan_sapMKO_0175 | BCGDan_sapMKO_0175 | 177 |
| 194993 | 196657 | fadD5 | Rv0166 | 1665 | 224892 | 226556 | fadD5 | BCG_0203 | 1665 | 195225 | 196889 | fadD5 | BCGDan_0176 | 1665 | 195333 | 196889 | fadD5 | BCGDan_sapMKO_0176 | 1557 |
| 196861 | 197658 | yrbE1A | Rv0167 | 798 | 226760 | 227557 | yrbE1A | BCG_0204 | 798 | 197093 | 197890 | yrbE1A | BCGDan_0177 | 798 | 197093 | 197890 | yrbE1A | BCGDan_sapMKO_0177 | 798 |
| 197660 | 198529 | yrbE1B | Rv0168 | 870 | 227559 | 228428 | yrbE1B | BCG_0205 | 870 | 197892 | 198761 | yrbE1B | BCGDan_0178 | 870 | 197892 | 198761 | yrbE1B | BCGDan_sapMKO_0178 | 870 |
| 198534 | 199898 | mce1A | Rv0169 | 1365 | 228433 | 229797 | mce1A | BCG_0206 | 1365 | 198766 | 200130 | mce1A | BCGDan_0179 | 1365 | 198766 | 200130 | mce1A | BCGDan_sapMKO_0179 | 1365 |
| 199895 | 200935 | mce1B | Rv0170 | 1041 | 229794 | 230834 | mce1B | BCG_0207 | 1041 | 200127 | 201167 | mce1B | BCGDan_0180 | 1041 | 200127 | 201167 | mce1B | BCGDan_sapMKO_0180 | 1041 |
| 200932 | 202479 | mce1C | Rv0171 | 1548 | 230831 | 232378 | mce1C | BCG_0208 | 1548 | 201164 | 202711 | mce1C | BCGDan_0181 | 1548 | 201206 | 202711 | mce1C | BCGDan_sapMKO_0181 | 1506 |
| 202476 | 204068 | mce1D | Rv0172 | 1593 | 232375 | 233967 | mce1D | BCG_0209 | 1593 | 202708 | 204300 | mce1D | BCGDan_0182 | 1593 | 202708 | 204300 | mce1D | BCGDan_sapMKO_0182 | 1593 |
| 204065 | 205237 | lprK | Rv0173 | 1173 | 233964 | 235136 | lprK | BCG_0210 | 1173 | 204297 | 205469 | lprK | BCGDan_0183 | 1173 | 204300 | 205469 | lprK | BCGDan_sapMKO_0183 | 1170 |
| 205231 | 206778 | mce1F | Rv0174 | 1548 | 235130 | 236677 | mce1F | BCG_0211 | 1548 | 205463 | 207010 | mce1F | BCGDan_0184 | 1548 | 205463 | 207010 | mce1F | BCGDan_sapMKO_0184 | 1548 |
| 206814 | 207455 | Rv0175 | Rv0175 | 642 | 236713 | 237354 | BCG_0212 | BCG_0212 | 642 | 207046 | 207687 | BCGDan_0185 | BCGDan_0185 | 642 | 206980 | 207687 | BCGDan_sapMKO_0185 | BCGDan_sapMKO_0185 | 708 |
| 207452 | 208420 | Rv0176 | Rv0176 | 969 | 237351 | 238319 | BCG_0213 | BCG_0213 | 969 | 207684 | 208652 | BCGDan_0186 | BCGDan_0186 | 969 | 207684 | 208652 | BCGDan_sapMKO_0186 | BCGDan_sapMKO_0186 | 969 |
| 208417 | 208971 | Rv0177 | Rv0177 | 555 | 238316 | 238870 | BCG_0214 | BCG_0214 | 555 | 208649 | 209203 | BCGDan_0187 | BCGDan_0187 | 555 | 208649 | 209203 | BCGDan_sapMKO_0187 | BCGDan_sapMKO_0187 | 555 |
| 208938 | 209672 | Rv0178 | Rv0178 | 735 | 238837 | 239571 | BCG_0215 | BCG_0215 | 735 | 209170 | 209904 | BCGDan_0188 | BCGDan_0188 | 735 | 209170 | 209904 | BCGDan_sapMKO_0188 | BCGDan_sapMKO_0188 | 735 |
| 209703 | 210812 | lprO | Rv0179c | 1110 | 239602 | 240711 | lprO | BCG_0216c | 1110 | 209935 | 211044 | lprO | BCGDan_0189 | 1110 | 209935 | 211044 | lprO | BCGDan_sapMKO_0189 | 1110 |
| 210892 | 212250 | Rv0180c | Rv0180c | 1359 | 240791 | 242149 | BCG_0217c | BCG_0217c | 1359 | 211124 | 212482 | BCGDan_0190 | BCGDan_0190 | 1359 | 211124 | 212482 | BCGDan_sapMKO_0190 | BCGDan_sapMKO_0190 | 1359 |
| 212277 | 213011 | Rv0181c | Rv0181c | 735 | 242176 | 242910 | BCG_0218c | BCG_0218c | 735 | 212509 | 213243 | BCGDan_0191 | BCGDan_0191 | 735 | 212509 | 213243 | BCGDan_sapMKO_0191 | BCGDan_sapMKO_0191 | 735 |
| 213028 | 214140 | sigG | Rv0182c | 1113 | 242927 | 244039 | sigG | BCG_0219c | 1113 | 213260 | 214372 | sigG | BCGDan_0192 | 1113 | 213260 | 214258 | sigG | BCGDan_sapMKO_0192 | 999 |
| 214088 | 214927 | Rv0183 | Rv0183 | 840 | 243987 | 244826 | BCG_0220 | BCG_0220 | 840 | 214320 | 215159 | BCGDan_0193 | BCGDan_0193 | 840 | 214320 | 215159 | BCGDan_sapMKO_0193 | BCGDan_sapMKO_0193 | 840 |
| 214969 | 215718 | Rv0184 | Rv0184 | 750 | 244868 | 245617 | BCG_0221 | BCG_0221 | 750 | 215201 | 215950 | BCGDan_0194 | BCGDan_0194 | 750 | 215201 | 215950 | BCGDan_sapMKO_0194 | BCGDan_sapMKO_0194 | 750 |
| 215715 | 216224 | Rv0185 | Rv0185 | 510 | 245614 | 246123 | BCG_0222 | BCG_0222 | 510 | 215947 | 216456 | BCGDan_0195 | BCGDan_0195 | 510 | 215989 | 216456 | BCGDan_sapMKO_0195 | BCGDan_sapMKO_0195 | 468 |
| 216269 | 218344 | bglS | Rv0186 | 2076 | 246168 | 248243 | bglS | BCG_0223 | 2076 | 216501 | 218576 | bglS | BCGDan_0196 | 2076 | 216501 | 218576 | bglS | BCGDan_sapMKO_0196 | 2076 |
| 218705 | 219367 | Rv0187 | Rv0187 | 663 | 248604 | 249266 | BCG_0224 | BCG_0224 | 663 | 218937 | 219599 | BCGDan_0197 | BCGDan_0197 | 663 | 218943 | 219599 | BCGDan_sapMKO_0197 | BCGDan_sapMKO_0197 | 657 |
| 219486 | 219917 | Rv0188 | Rv0188 | 432 | 249385 | 249816 | BCG_0225 | BCG_0225 | 432 | 219718 | 220149 | BCGDan_0198 | BCGDan_0198 | 432 | 219757 | 220149 | BCGDan_sapMKO_0198 | BCGDan_sapMKO_0198 | 393 |
| 219996 | 221723 | ilvD | Rv0189c | 1728 | 249895 | 251622 | ilvD | BCG_0226c | 1728 | 220228 | 221955 | ilvD | BCGDan_0199 | 1728 | 220228 | 221955 | ilvD | BCGDan_sapMKO_0199 | 1728 |
| 221871 | 222161 | Rv0190 | Rv0190 | 291 | 251770 | 252060 | BCG_0227 | BCG_0227 | 291 | 222103 | 222393 | BCGDan_0200 | BCGDan_0200 | 291 | 222103 | 222393 | BCGDan_sapMKO_0200 | BCGDan_sapMKO_0200 | 291 |
| 222289 | 223530 | Rv0191 | Rv0191 | 1242 | 252188 | 253429 | BCG_0228 | BCG_0228 | 1242 | 222521 | 223762 | BCGDan_0201 | BCGDan_0201 | 1242 | 222593 | 223762 | BCGDan_sapMKO_0201 | BCGDan_sapMKO_0201 | 1170 |
| 223564 | 224664 | Rv0192 | Rv0192 | 1101 | 253506 | 254564 | BCG_0229 | BCG_0229 | 1059 | 223839 | 224897 | BCGDan_0202 | BCGDan_0202 | 1059 | 223902 | 224897 | BCGDan_sapMKO_0202 | BCGDan_sapMKO_0202 | 996 |
| 224724 | 226571 | Rv0193c | Rv0193c | 1848 | 254624 | 256471 | BCG_0230c | BCG_0230c | 1848 | 224957 | 226804 | BCGDan_0203 | BCGDan_0203 | 1848 | 224957 | 226804 | BCGDan_sapMKO_0203 | BCGDan_sapMKO_0203 | 1848 |
| 226878 | 230462 | Rv0194 | Rv0194 | 3585 | 256778 | 260362 | BCG_0231 | BCG_0231 | 3585 | 227111 | 230695 | BCGDan_0204 | BCGDan_0204 | 3585 | 227111 | 230695 | BCGDan_sapMKO_0204 | BCGDan_sapMKO_0204 | 3585 |
| 230899 | 231534 | Rv0195 | Rv0195 | 636 | 260800 | 261435 | BCG_0232 | BCG_0232 | 636 | 231133 | 231768 | BCGDan_0205 | BCGDan_0205 | 636 | 231070 | 231768 | BCGDan_sapMKO_0205 | BCGDan_sapMKO_0205 | 699 |
| 231647 | 232231 | Rv0196 | Rv0196 | 585 | 261548 | 262132 | BCG_0233 | BCG_0233 | 585 | 231881 | 232465 | BCGDan_0206 | BCGDan_0206 | 585 | 231902 | 232465 | BCGDan_sapMKO_0206 | BCGDan_sapMKO_0206 | 564 |
| 232231 | 234519 | Rv0197 | Rv0197 | 2289 | 262132 | 264378 | BCG_0234 | BCG_0234 | 2247 | 232465 | 234711 | BCGDan_0207 | BCGDan_0207 | 2247 | 232465 | 234711 | BCGDan_sapMKO_0207 | BCGDan_sapMKO_0207 | 2247 |
| 234516 | 236507 | zmp1 | Rv0198c | 1992 | 264419 | 266410 | BCG_0235c | BCG_0235c | 1992 | 234752 | 236743 | BCGDan_0208 | BCGDan_0208 | 1992 | 234752 | 236743 | BCGDan_sapMKO_0208 | BCGDan_sapMKO_0208 | 1992 |
| 236550 | 237209 | Rv0199 | Rv0199 | 660 | 266453 | 267112 | BCG_0236 | BCG_0236 | 660 | 236786 | 237445 | BCGDan_0209 | BCGDan_0209 | 660 | 236786 | 237445 | BCGDan_sapMKO_0209 | BCGDan_sapMKO_0209 | 660 |
| 237206 | 237895 | Rv0200 | Rv0200 | 690 | 267109 | 267798 | BCG_0237 | BCG_0237 | 690 | 237442 | 238131 | BCGDan_0210 | BCGDan_0210 | 690 | 237442 | 238131 | BCGDan_sapMKO_0210 | BCGDan_sapMKO_0210 | 690 |
| 237892 | 238395 | Rv0201c | Rv0201c | 504 | 267795 | 268298 | BCG_0238c | BCG_0238c | 504 | 238128 | 238631 | BCGDan_0211 | BCGDan_0211 | 504 | 238128 | 238631 | BCGDan_sapMKO_0211 | BCGDan_sapMKO_0211 | 504 |
| 238392 | 241292 | mmpL11 | Rv0202c | 2901 | 268295 | 271195 | mmpL11 | BCG_0239c | 2901 | 238628 | 241528 | mmpL11 | BCGDan_0212 | 2901 | 238628 | 241528 | mmpL11 | BCGDan_sapMKO_0212 | 2901 |
| 241514 | 241924 | Rv0203 | Rv0203 | 411 | 271417 | 271827 | BCG_0240 | BCG_0240 | 411 | 241750 | 242160 | BCGDan_0213 | BCGDan_0213 | 411 | 241750 | 242160 | BCGDan_sapMKO_0213 | BCGDan_sapMKO_0213 | 411 |
| 241976 | 243214 | Rv0204c | Rv0204c | 1239 | 271879 | 273162 | BCG_0241c | BCG_0241c | 1284 | 242212 | 243495 | BCGDan_0214 | BCGDan_0214 | 1284 | 242212 | 243495 | BCGDan_sapMKO_0214 | BCGDan_sapMKO_0214 | 1284 |
| 243384 | 244487 | Rv0205 | Rv0205 | 1104 | 273287 | 274390 | BCG_0242 | BCG_0242 | 1104 | 243620 | 244723 | BCGDan_0215 | BCGDan_0215 | 1104 | 243620 | 244723 | BCGDan_sapMKO_0215 | BCGDan_sapMKO_0215 | 1104 |
| 244484 | 247318 | mmpL3 | Rv0206c | 2835 | 274387 | 277221 | mmpL3 | BCG_0243c | 2835 | 244720 | 247554 | mmpL3 | BCGDan_0216 | 2835 | 244720 | 247530 | mmpL3 | BCGDan_sapMKO_0216 | 2811 |
| 247384 | 248112 | Rv0207c | Rv0207c | 729 | 277287 | 278015 | BCG_0244c | BCG_0244c | 729 | 247620 | 248348 | BCGDan_0217 | BCGDan_0217 | 729 | 247620 | 248348 | BCGDan_sapMKO_0217 | BCGDan_sapMKO_0217 | 729 |
| 248115 | 248906 | Rv0208c | Rv0208c | 792 | 278018 | 278809 | BCG_0245c | BCG_0245c | 792 | 248351 | 249142 | BCGDan_0218 | BCGDan_0218 | 792 | 248351 | 249142 | BCGDan_sapMKO_0218 | BCGDan_sapMKO_0218 | 792 |
| 249038 | 250123 | Rv0209 | Rv0209 | 1086 | 278941 | 280026 | BCG_0246 | BCG_0246 | 1086 | 249274 | 250359 | BCGDan_0219 | BCGDan_0219 | 1086 | 249274 | 250359 | BCGDan_sapMKO_0219 | BCGDan_sapMKO_0219 | 1086 |
| 250120 | 251598 | Rv0210 | Rv0210 | 1479 | 280023 | 281501 | BCG_0247 | BCG_0247 | 1479 | 250356 | 251834 | BCGDan_0220 | BCGDan_0220 | 1479 | 250356 | 251834 | BCGDan_sapMKO_0220 | BCGDan_sapMKO_0220 | 1479 |
| 251782 | 253602 | pckA | Rv0211 | 1821 | 281685 | 283505 | pckA | BCG_0248 | 1821 | 252018 | 253838 | pckA | BCGDan_0221 | 1821 | 252018 | 253838 | pckA | BCGDan_sapMKO_0221 | 1821 |
| 253669 | 254640 | nadR | Rv0212c | 972 | 283572 | 284543 | nadR | BCG_0249c | 972 | 253905 | 254876 | nadR | BCGDan_0222 | 972 | 253905 | 254864 | nadR | BCGDan_sapMKO_0222 | 960 |
| 254637 | 255950 | Rv0213c | Rv0213c | 1314 | 284540 | 285853 | BCG_0250c | BCG_0250c | 1314 | 254873 | 256186 | BCGDan_0223 | BCGDan_0223 | 1314 | 254873 | 256210 | BCGDan_sapMKO_0223 | BCGDan_sapMKO_0223 | 1338 |
| 256064 | 257677 | fadD4 | Rv0214 | 1614 | 285967 | 287580 | fadD4 | BCG_0251 | 1614 | 256300 | 257913 | fadD4 | BCGDan_0224 | 1614 | 256396 | 257913 | fadD4 | BCGDan_sapMKO_0224 | 1518 |
| 257783 | 258856 | fadE3 | Rv0215c | 1074 | 287622 | 288788 | fadE3 | BCG_0252c | 1167 | 257955 | 259121 | fadE3 | BCGDan_0225 | 1167 | 257955 | 259121 | fadE3 | BCGDan_sapMKO_0225 | 1167 |
| 258913 | 259926 | Rv0216 | Rv0216 | 1014 | 288845 | 289858 | BCG_0253 | BCG_0253 | 1014 | 259178 | 260191 | BCGDan_0226 | BCGDan_0226 | 1014 | 259178 | 260191 | BCGDan_sapMKO_0226 | BCGDan_sapMKO_0226 | 1014 |
| 259923 | 260831 | lipW | Rv0217c | 909 | 289855 | 290763 | lipW | BCG_0254c | 909 | 260188 | 261096 | lipW | BCGDan_0227 | 909 | 260188 | 261096 | lipW | BCGDan_sapMKO_0227 | 909 |
| 260924 | 262252 | Rv0218 | Rv0218 | 1329 | 290856 | 292184 | BCG_0255 | BCG_0255 | 1329 | 261189 | 262517 | BCGDan_0228 | BCGDan_0228 | 1329 | 261189 | 262517 | BCGDan_sapMKO_0228 | BCGDan_sapMKO_0228 | 1329 |
| 262254 | 262802 | Rv0219 | Rv0219 | 549 | 292186 | 292734 | BCG_0256 | BCG_0256 | 549 | 262519 | 263067 | BCGDan_0229 | BCGDan_0229 | 549 | 262519 | 263067 | BCGDan_sapMKO_0229 | BCGDan_sapMKO_0229 | 549 |
| 262812 | 264023 | lipC | Rv0220 | 1212 | 292744 | 293985 | lipC | BCG_0257 | 1242 | 263077 | 264318 | lipC | BCGDan_0230 | 1242 | 263077 | 264318 | lipC | BCGDan_sapMKO_0230 | 1242 |
| 264067 | 265476 | Rv0221 | Rv0221 | 1410 | 293999 | 294772 | BCG_0258 | BCG_0258 | 774 | 264332 | 265105 | BCGDan_0231 | BCGDan_0231 | 774 | 264332 | 265105 | BCGDan_sapMKO_0231 | BCGDan_sapMKO_0231 | 774 |
| 266301 | 267764 | Rv0223c | Rv0223c | 1464 | 294638 | 295447 | BCG_0259c | BCG_0259c | 810 | 264971 | 265780 | BCGDan_0232 | BCGDan_0232 | 810 | 264971 | 265780 | BCGDan_sapMKO_0232 | BCGDan_sapMKO_0232 | 810 |
| NA | NA | NA | NA | NA | 295444 | 295794 | BCG_0260c | BCG_0260c | 351 | 265777 | 266127 | BCGDan_0233 | BCGDan_0233 | 351 | 265777 | 266127 | BCGDan_sapMKO_0233 | BCGDan_sapMKO_0233 | 351 |
| 267863 | 268627 | Rv0224c | Rv0224c | 765 | 295893 | 296657 | BCG_0261c | BCG_0261c | 765 | 266226 | 266990 | BCGDan_0234 | BCGDan_0234 | 765 | 266226 | 266999 | BCGDan_sapMKO_0234 | BCGDan_sapMKO_0234 | 774 |
| 268663 | 269817 | Rv0225 | Rv0225 | 1155 | 296693 | 297847 | BCG_0262 | BCG_0262 | 1155 | 267026 | 268180 | BCGDan_0235 | BCGDan_0235 | 1155 | 267026 | 268180 | BCGDan_sapMKO_0235 | BCGDan_sapMKO_0235 | 1155 |
| 269834 | 271564 | Rv0226c | Rv0226c | 1731 | 297864 | 299594 | BCG_0263c | BCG_0263c | 1731 | 268197 | 269927 | BCGDan_0236 | BCGDan_0236 | 1731 | 268197 | 269927 | BCGDan_sapMKO_0236 | BCGDan_sapMKO_0236 | 1731 |
| 271574 | 272839 | Rv0227c | Rv0227c | 1266 | 299604 | 300869 | BCG_0264c | BCG_0264c | 1266 | 269937 | 271202 | BCGDan_0237 | BCGDan_0237 | 1266 | 269937 | 271202 | BCGDan_sapMKO_0237 | BCGDan_sapMKO_0237 | 1266 |
| 273055 | 274278 | Rv0228 | Rv0228 | 1224 | 301085 | 302308 | BCG_0265 | BCG_0265 | 1224 | 271418 | 272641 | BCGDan_0238 | BCGDan_0238 | 1224 | 271418 | 272641 | BCGDan_sapMKO_0238 | BCGDan_sapMKO_0238 | 1224 |
| 274306 | 274986 | Rv0229c | Rv0229c | 681 | 302336 | 303016 | BCG_0266c | BCG_0266c | 681 | 272669 | 273349 | BCGDan_0239 | BCGDan_0239 | 681 | 272669 | 273349 | BCGDan_sapMKO_0239 | BCGDan_sapMKO_0239 | 681 |
| 274983 | 275963 | php | Rv0230c | 981 | 303013 | 303993 | php | BCG_0267c | 981 | 273346 | 274326 | php | BCGDan_0240 | 981 | 273346 | 274326 | php | BCGDan_sapMKO_0240 | 981 |
| 276058 | 277764 | fadE4 | Rv0231 | 1707 | 304088 | 305794 | fadE4 | BCG_0268 | 1707 | 274421 | 276127 | fadE4 | BCGDan_0241 | 1707 | 274421 | 276127 | fadE4 | BCGDan_sapMKO_0241 | 1707 |
| 277899 | 278588 | Rv0232 | Rv0232 | 690 | 305929 | 306618 | BCG_0269 | BCG_0269 | 690 | 276262 | 276951 | BCGDan_0242 | BCGDan_0242 | 690 | 276262 | 276951 | BCGDan_sapMKO_0242 | BCGDan_sapMKO_0242 | 690 |
| 278585 | 279529 | nrdB | Rv0233 | 945 | 306615 | 307559 | nrdB | BCG_0270 | 945 | 276948 | 277892 | nrdB | BCGDan_0243 | 945 | 276948 | 277892 | nrdB | BCGDan_sapMKO_0243 | 945 |
| 279605 | 281140 | gabD1 | Rv0234c | 1536 | 307635 | 309170 | gabD1 | BCG_0271c | 1536 | 277968 | 279503 | gabD1 | BCGDan_0244 | 1536 | 277968 | 279341 | gabD1 | BCGDan_sapMKO_0244 | 1374 |
| 281166 | 282614 | Rv0235c | Rv0235c | 1449 | 309196 | 310644 | BCG_0272c | BCG_0272c | 1449 | 279529 | 280977 | BCGDan_0245 | BCGDan_0245 | 1449 | 279529 | 280977 | BCGDan_sapMKO_0245 | BCGDan_sapMKO_0245 | 1449 |
| 282649 | 286851 | aftD | Rv0236c | 4203 | 310679 | 314881 | BCG_0273c | BCG_0273c | 4203 | 281012 | 285214 | BCGDan_0246 | BCGDan_0246 | 4203 | 281012 | 285214 | BCGDan_sapMKO_0246 | BCGDan_sapMKO_0246 | 4203 |
| 286898 | 287071 | Rv0236A | Rv0236A | 174 | 314928 | 315101 | BCG_0274c | BCG_0274c | 174 | 285261 | 285434 | BCGDan_0247 | BCGDan_0247 | 174 | 285261 | 285434 | BCGDan_sapMKO_0247 | BCGDan_sapMKO_0247 | 174 |
| 287186 | 288352 | lpqI | Rv0237 | 1167 | 315216 | 316382 | lpqI | BCG_0275 | 1167 | 285549 | 286715 | lpqI | BCGDan_0248 | 1167 | 285549 | 286715 | lpqI | BCGDan_sapMKO_0248 | 1167 |
| 288428 | 289042 | Rv0238 | Rv0238 | 615 | 316458 | 317072 | BCG_0276 | BCG_0276 | 615 | 286791 | 287405 | BCGDan_0249 | BCGDan_0249 | 615 | 286839 | 287405 | BCGDan_sapMKO_0249 | BCGDan_sapMKO_0249 | 567 |
| 289104 | 289337 | vapB24 | Rv0239 | 234 | 317134 | 317367 | BCG_0277 | BCG_0277 | 234 | 287467 | 287700 | BCGDan_0250 | BCGDan_0250 | 234 | 287467 | 287700 | BCGDan_sapMKO_0250 | BCGDan_sapMKO_0250 | 234 |
| 289345 | 289782 | vapC24 | Rv0240 | 438 | 317375 | 317812 | BCG_0278 | BCG_0278 | 438 | 287708 | 288145 | BCGDan_0251 | BCGDan_0251 | 438 | 287708 | 288145 | BCGDan_sapMKO_0251 | BCGDan_sapMKO_0251 | 438 |
| 289812 | 290654 | htdX | Rv0241c | 843 | 317842 | 318684 | BCG_0279c | BCG_0279c | 843 | 288175 | 289017 | BCGDan_0252 | BCGDan_0252 | 843 | 288175 | 289017 | BCGDan_sapMKO_0252 | BCGDan_sapMKO_0252 | 843 |
| 290665 | 292029 | fabG4 | Rv0242c | 1365 | 318695 | 320059 | fabG4 | BCG_0280c | 1365 | 289028 | 290392 | fabG4 | BCGDan_0253 | 1365 | 289028 | 290392 | fabG4 | BCGDan_sapMKO_0253 | 1365 |
| 292171 | 293493 | fadA2 | Rv0243 | 1323 | 320201 | 321523 | fadA2 | BCG_0281 | 1323 | 290534 | 291856 | fadA2 | BCGDan_0254 | 1323 | 290534 | 291856 | fadA2 | BCGDan_sapMKO_0254 | 1323 |
| 293798 | 295633 | fadE5 | Rv0244c | 1836 | 321829 | 323664 | fadE5 | BCG_0282c | 1836 | 292162 | 293997 | fadE5 | BCGDan_0255 | 1836 | 292162 | 293997 | fadE5 | BCGDan_sapMKO_0255 | 1836 |
| 296005 | 296493 | Rv0245 | Rv0245 | 489 | 324036 | 324524 | BCG_0283 | BCG_0283 | 489 | 294369 | 294857 | BCGDan_0256 | BCGDan_0256 | 489 | 294369 | 294857 | BCGDan_sapMKO_0256 | BCGDan_sapMKO_0256 | 489 |

| | | | | | | | | | | | | | | | | | | | |
|---|---|---|---|---|---|---|---|---|---|---|---|---|---|---|---|---|---|---|---|
| 296809 | 298119 | Rv0246 | Rv0246 | 1311 | 324840 | 326150 | BCG_0284 | BCG_0284 | 1311 | 295173 | 296483 | BCGDan_0257 | BCGDan_0257 | 1311 | 295194 | 296483 | BCGDan_sapMKO_0257 | BCGDan_sapMKO_0257 | 1290 |
| 298116 | 298862 | Rv0247c | Rv0247c | 747 | 326147 | 326893 | BCG_0285c | BCG_0285c | 747 | 296480 | 297226 | BCGDan_0258 | BCGDan_0258 | 747 | 296480 | 297208 | BCGDan_sapMKO_0258 | BCGDan_sapMKO_0258 | 729 |
| 298863 | 300803 | Rv0248c | Rv0248c | 1941 | 326894 | 328834 | BCG_0286c | BCG_0286c | 1941 | 297227 | 299167 | BCGDan_0259 | BCGDan_0259 | 1941 | 297227 | 299167 | BCGDan_sapMKO_0259 | BCGDan_sapMKO_0259 | 1941 |
| 300834 | 301655 | Rv0249c | Rv0249c | 822 | 328865 | 329686 | BCG_0287c | BCG_0287c | 822 | 299198 | 300019 | BCGDan_0260 | BCGDan_0260 | 822 | 299198 | 300019 | BCGDan_sapMKO_0260 | BCGDan_sapMKO_0260 | 822 |
| 301735 | 302028 | Rv0250c | Rv0250c | 294 | 329766 | 330059 | BCG_0288c | BCG_0288c | 294 | 300099 | 300392 | BCGDan_0261 | BCGDan_0261 | 294 | 300099 | 300392 | BCGDan_sapMKO_0261 | BCGDan_sapMKO_0261 | 294 |
| 302173 | 302652 | hsp | Rv0251c | 480 | 330204 | 330683 | hsp | BCG_0289c | 480 | 300537 | 301016 | hsp | BCGDan_0262 | 480 | 300537 | 301016 | hsp | BCGDan_sapMKO_0262 | 480 |
| 302866 | 305427 | nirB | Rv0252 | 2562 | 330897 | 333458 | nirB | BCG_0290 | 2562 | 301230 | 303791 | nirB | BCGDan_0263 | 2562 | 301281 | 303791 | nirB | BCGDan_sapMKO_0263 | 2511 |
| 305453 | 305809 | nirD | Rv0253 | 357 | 333484 | 333840 | nirD | BCG_0291 | 357 | 303817 | 304173 | nirD | BCGDan_0264 | 357 | 303817 | 304173 | nirD | BCGDan_sapMKO_0264 | 357 |
| 305825 | 306349 | cobU | Rv0254c | 525 | 333856 | 334380 | cobU | BCG_0292c | 525 | 304189 | 304713 | cobU | BCGDan_0265 | 525 | 304189 | 304713 | cobU | BCGDan_sapMKO_0265 | 525 |
| 306374 | 307858 | cobQ1 | Rv0255c | 1485 | 334405 | 335889 | cobQ1 | BCG_0293c | 1485 | 304738 | 306222 | cobQ1 | BCGDan_0266 | 1485 | 304738 | 306222 | cobQ1 | BCGDan_sapMKO_0266 | 1485 |
| 307877 | 309547 | PPE2 | Rv0256c | 1671 | 335908 | 337578 | PPE2 | BCG_0294c | 1671 | 306241 | 307911 | PPE2 | BCGDan_0267 | 1671 | 306241 | 307911 | PPE2 | BCGDan_sapMKO_0267 | 1671 |
| 309699 | 310073 | Rv0257 | Rv0257 | 375 | 337886 | 338104 | BCG_0295 | BCG_0295 | 219 | 308219 | 308437 | BCGDan_0268 | BCGDan_0268 | 219 | 308219 | 308437 | BCGDan_sapMKO_0268 | BCGDan_sapMKO_0268 | 219 |
| 310294 | 310749 | Rv0258c | Rv0258c | 456 | 338325 | 338780 | BCG_0296c | BCG_0296c | 456 | 308658 | 309113 | BCGDan_0269 | BCGDan_0269 | 456 | 308658 | 309062 | BCGDan_sapMKO_0269 | BCGDan_sapMKO_0269 | 405 |
| 310774 | 311517 | Rv0259c | Rv0259c | 744 | 338805 | 339548 | BCG_0297c | BCG_0297c | 744 | 309138 | 309881 | BCGDan_0270 | BCGDan_0270 | 744 | 309138 | 309881 | BCGDan_sapMKO_0270 | BCGDan_sapMKO_0270 | 744 |
| 311514 | 312659 | Rv0260c | Rv0260c | 1146 | 339545 | 340690 | BCG_0298c | BCG_0298c | 1146 | 309878 | 311023 | BCGDan_0271 | BCGDan_0271 | 1146 | 309878 | 311023 | BCGDan_sapMKO_0271 | BCGDan_sapMKO_0271 | 1146 |
| 312759 | 314168 | narK3 | Rv0261c | 1410 | 340790 | 342199 | narK3 | BCG_0299c | 1410 | 311123 | 312532 | narK3 | BCGDan_0272 | 1410 | 311123 | 312532 | narK3 | BCGDan_sapMKO_0272 | 1410 |
| 314309 | 314854 | aac | Rv0262c | 546 | 342340 | 342885 | aac | BCG_0300c | 546 | 312673 | 313218 | aac | BCGDan_0273 | 546 | 312673 | 313218 | aac | BCGDan_sapMKO_0273 | 546 |
| 314864 | 315766 | Rv0263c | Rv0263c | 903 | 342895 | 343797 | BCG_0301c | BCG_0301c | 903 | 313228 | 314130 | BCGDan_0274 | BCGDan_0274 | 903 | 313228 | 314130 | BCGDan_sapMKO_0274 | BCGDan_sapMKO_0274 | 903 |
| 315783 | 316415 | Rv0264c | Rv0264c | 633 | 343814 | 344446 | BCG_0302c | BCG_0302c | 633 | 314147 | 314779 | BCGDan_0275 | BCGDan_0275 | 633 | 314147 | 314800 | BCGDan_sapMKO_0275 | BCGDan_sapMKO_0275 | 654 |
| 316511 | 317503 | Rv0265c | Rv0265c | 993 | 344542 | 345534 | BCG_0303c | BCG_0303c | 993 | 314875 | 315867 | BCGDan_0276 | BCGDan_0276 | 993 | 314875 | 315837 | BCGDan_sapMKO_0276 | BCGDan_sapMKO_0276 | 963 |
| 317525 | 321154 | oplA | Rv0266c | 3630 | 345556 | 349185 | oplA | BCG_0304c | 3630 | 315889 | 319518 | oplA | BCGDan_0277 | 3630 | 315889 | 319518 | oplA | BCGDan_sapMKO_0277 | 3630 |
| 321331 | 322722 | narU | Rv0267 | 1392 | 349362 | 350753 | narU | BCG_0305 | 1392 | 319695 | 321086 | narU | BCGDan_0278 | 1392 | 319695 | 321086 | narU | BCGDan_sapMKO_0278 | 1392 |
| 322764 | 323273 | Rv0268c | Rv0268c | 510 | 350795 | 351304 | BCG_0306c | BCG_0306c | 510 | 321128 | 321637 | BCGDan_0279 | BCGDan_0279 | 510 | 321128 | 321637 | BCGDan_sapMKO_0279 | BCGDan_sapMKO_0279 | 510 |
| 323338 | 324531 | Rv0269c | Rv0269c | 1194 | 351369 | 352562 | BCG_0307c | BCG_0307c | 1194 | 321702 | 322895 | BCGDan_0280 | BCGDan_0280 | 1194 | 321702 | 322895 | BCGDan_sapMKO_0280 | BCGDan_sapMKO_0280 | 1194 |
| 324567 | 326249 | fadD2 | Rv0270 | 1683 | 352598 | 354280 | fadD2 | BCG_0308 | 1683 | 322931 | 324613 | fadD2 | BCGDan_0281 | 1683 | 322931 | 324613 | fadD2 | BCGDan_sapMKO_0281 | 1683 |
| 326266 | 328461 | fadE6 | Rv0271c | 2196 | 354297 | 356492 | fadE6 | BCG_0309c | 2196 | 324630 | 326825 | fadE6 | BCGDan_0282 | 2196 | 324630 | 326825 | fadE6 | BCGDan_sapMKO_0282 | 2196 |
| 328575 | 329708 | Rv0272c | Rv0272c | 1134 | 356606 | 357739 | BCG_0310c | BCG_0310c | 1134 | 326939 | 328072 | BCGDan_0283 | BCGDan_0283 | 1134 | 326939 | 327955 | BCGDan_sapMKO_0283 | BCGDan_sapMKO_0283 | 1017 |
| 329705 | 330325 | Rv0273c | Rv0273c | 621 | 357736 | 358356 | BCG_0311c | BCG_0311c | 621 | 328069 | 328689 | BCGDan_0284 | BCGDan_0284 | 621 | 328069 | 328689 | BCGDan_sapMKO_0284 | BCGDan_sapMKO_0284 | 621 |
| 330422 | 331003 | Rv0274 | Rv0274 | 582 | 358453 | 359034 | BCG_0312 | BCG_0312 | 582 | 328786 | 329367 | BCGDan_0285 | BCGDan_0285 | 582 | 328786 | 329367 | BCGDan_sapMKO_0285 | BCGDan_sapMKO_0285 | 582 |
| 330933 | 331658 | Rv0275c | Rv0275c | 726 | 358964 | 359689 | BCG_0313c | BCG_0313c | 726 | 329297 | 330022 | BCGDan_0286 | BCGDan_0286 | 726 | 329297 | 330022 | BCGDan_sapMKO_0286 | BCGDan_sapMKO_0286 | 726 |
| 331748 | 332668 | Rv0276 | Rv0276 | 921 | 359779 | 360699 | BCG_0314 | BCG_0314 | 921 | 330112 | 331032 | BCGDan_0287 | BCGDan_0287 | 921 | 330112 | 331032 | BCGDan_sapMKO_0287 | BCGDan_sapMKO_0287 | 921 |
| 332708 | 333136 | vapC25 | Rv0277c | 429 | 360739 | 361167 | BCG_0315c | BCG_0315c | 429 | 331072 | 331500 | BCGDan_0288 | BCGDan_0288 | 429 | 331072 | 331500 | BCGDan_sapMKO_0288 | BCGDan_sapMKO_0288 | 429 |
| 333160 | 333417 | vapB25 | Rv0277A | 258 | 361191 | 361448 | BCG_0315A | BCG_0315A | 258 | 331524 | 331781 | BCGDan_0289 | BCGDan_0289 | 258 | 331524 | 331781 | BCGDan_sapMKO_0289 | BCGDan_sapMKO_0289 | 258 |
| NA | NA | NA | NA | NA | 361468 | 361710 | PE_PGRS3b | BCG_0316c | 243 | 331801 | 332043 | PE_PGRS3b | BCGDan_0290 | 243 | NA | NA | NA | NA | NA |
| NA | NA | NA | NA | NA | 361533 | 364283 | PE_PGRS3a | BCG_0317c | 2751 | 331866 | 334616 | PE_PGRS3a | BCGDan_0291 | 2751 | 331866 | 334616 | PE_PGRS3a | BCGDan_sapMKO_0290 | 2751 |
| 333437 | 336310 | PE_PGRS3 | Rv0278c | 2874 | 364547 | 367246 | PE_PGRS3 | BCG_0318c | 2700 | 334880 | 337579 | PE_PGRS3 | BCGDan_0292 | 2700 | 334880 | 337579 | PE_PGRS3 | BCGDan_sapMKO_0291 | 2700 |
| 336560 | 339073 | PE_PGRS4 | Rv0279c | 2514 | 367496 | 370000 | PE_PGRS4 | BCG_0319c | 2505 | 337829 | 340333 | PE_PGRS4 | BCGDan_0293 | 2505 | 337829 | 340333 | PE_PGRS4 | BCGDan_sapMKO_0292 | 2505 |
| 339364 | 340974 | PPE3 | Rv0280 | 1611 | 370291 | 371901 | PPE3 | BCG_0320 | 1611 | 340624 | 342234 | PPE3 | BCGDan_0294 | 1611 | 340624 | 342234 | PPE3 | BCGDan_sapMKO_0293 | 1611 |
| 340998 | 341906 | Rv0281 | Rv0281 | 909 | 371925 | 372833 | BCG_0321 | BCG_0321 | 909 | 342258 | 343166 | BCGDan_0295 | BCGDan_0295 | 909 | 342258 | 343166 | BCGDan_sapMKO_0294 | BCGDan_sapMKO_0294 | 909 |
| 342130 | 344025 | eccA3 | Rv0282 | 1896 | 373058 | 374953 | BCG_0322 | BCG_0322 | 1896 | 343390 | 345285 | BCGDan_0296 | BCGDan_0296 | 1896 | 343423 | 345285 | BCGDan_sapMKO_0295 | BCGDan_sapMKO_0295 | 1863 |
| 344022 | 345638 | eccB3 | Rv0283 | 1617 | 374950 | 376566 | BCG_0323 | BCG_0323 | 1617 | 345282 | 346898 | BCGDan_0297 | BCGDan_0297 | 1617 | 345282 | 346898 | BCGDan_sapMKO_0296 | BCGDan_sapMKO_0296 | 1617 |
| 345635 | 349627 | eccC3 | Rv0284 | 3993 | 376563 | 380555 | BCG_0324 | BCG_0324 | 3993 | 346895 | 350887 | BCGDan_0298 | BCGDan_0298 | 3993 | 346895 | 350887 | BCGDan_sapMKO_0297 | BCGDan_sapMKO_0297 | 3993 |
| 349624 | 349932 | PE5 | Rv0285 | 309 | 380552 | 380860 | PE5 | BCG_0325 | 309 | 350884 | 351192 | PE5 | BCGDan_0299 | 309 | 350884 | 351192 | PE5 | BCGDan_sapMKO_0298 | 309 |
| 349935 | 351476 | PPE4 | Rv0286 | 1542 | 380863 | 382404 | PPE4 | BCG_0326 | 1542 | 351195 | 352736 | PPE4 | BCGDan_0300 | 1542 | 351195 | 352736 | PPE4 | BCGDan_sapMKO_0299 | 1542 |
| 351525 | 351818 | esxG | Rv0287 | 294 | 382453 | 382746 | esxG | BCG_0327 | 294 | 352785 | 353078 | esxG | BCGDan_0301 | 294 | 352785 | 353078 | esxG | BCGDan_sapMKO_0300 | 294 |
| 351848 | 352138 | esxH | Rv0288 | 291 | 382776 | 383066 | esxH | BCG_0328 | 291 | 353108 | 353398 | esxH | BCGDan_0302 | 291 | 353108 | 353398 | esxH | BCGDan_sapMKO_0301 | 291 |
| 352149 | 353036 | espG3 | Rv0289 | 888 | 383077 | 383964 | BCG_0329 | BCG_0329 | 888 | 353409 | 354296 | BCGDan_0303 | BCGDan_0303 | 888 | 353409 | 354296 | BCGDan_sapMKO_0302 | BCGDan_sapMKO_0302 | 888 |
| 353083 | 354501 | eccD3 | Rv0290 | 1419 | 384011 | 385429 | BCG_0330 | BCG_0330 | 1419 | 354343 | 355761 | BCGDan_0304 | BCGDan_0304 | 1419 | 354343 | 355761 | BCGDan_sapMKO_0303 | BCGDan_sapMKO_0303 | 1419 |
| 354498 | 355883 | mycP3 | Rv0291 | 1386 | 385426 | 386811 | BCG_0331 | BCG_0331 | 1386 | 355758 | 357143 | BCGDan_0305 | BCGDan_0305 | 1386 | 355758 | 357143 | BCGDan_sapMKO_0304 | BCGDan_sapMKO_0304 | 1386 |
| 355880 | 356875 | eccE3 | Rv0292 | 996 | 386808 | 387803 | BCG_0332 | BCG_0332 | 996 | 357140 | 358135 | BCGDan_0306 | BCGDan_0306 | 996 | 357140 | 358135 | BCGDan_sapMKO_0305 | BCGDan_sapMKO_0305 | 996 |
| 356862 | 358064 | Rv0293c | Rv0293c | 1203 | 387790 | 388992 | BCG_0333c | BCG_0333c | 1203 | 358122 | 359324 | BCGDan_0307 | BCGDan_0307 | 1203 | 358122 | 359333 | BCGDan_sapMKO_0306 | BCGDan_sapMKO_0306 | 1212 |
| 358171 | 358956 | tam | Rv0294 | 786 | 389099 | 389884 | tam | BCG_0334 | 786 | 359431 | 360216 | tam | BCGDan_0308 | 786 | 359431 | 360216 | tam | BCGDan_sapMKO_0307 | 786 |
| 358945 | 359748 | Rv0295c | Rv0295c | 804 | 389873 | 390676 | BCG_0335c | BCG_0335c | 804 | 360205 | 361008 | BCGDan_0309 | BCGDan_0309 | 804 | 360205 | 361008 | BCGDan_sapMKO_0308 | BCGDan_sapMKO_0308 | 804 |
| 359758 | 361155 | Rv0296c | Rv0296c | 1398 | 390686 | 392083 | BCG_0336c | BCG_0336c | 1398 | 361018 | 362415 | BCGDan_0310 | BCGDan_0310 | 1398 | 361018 | 362442 | BCGDan_sapMKO_0309 | BCGDan_sapMKO_0309 | 1425 |
| 361334 | 363109 | PE_PGRS5 | Rv0297 | 1776 | 392262 | 394082 | PE_PGRS5 | BCG_0337 | 1821 | 362594 | 364414 | PE_PGRS5 | BCGDan_0311 | 1821 | 362594 | 364414 | PE_PGRS5 | BCGDan_sapMKO_0310 | 1821 |
| 363252 | 363479 | Rv0298 | Rv0298 | 228 | 394225 | 394452 | BCG_0338 | BCG_0338 | 228 | 364557 | 364784 | BCGDan_0312 | BCGDan_0312 | 228 | 364557 | 364784 | BCGDan_sapMKO_0311 | BCGDan_sapMKO_0311 | 228 |
| 363476 | 363778 | Rv0299 | Rv0299 | 303 | 394449 | 394751 | BCG_0339 | BCG_0339 | 303 | 364781 | 365083 | BCGDan_0313 | BCGDan_0313 | 303 | 364781 | 365083 | BCGDan_sapMKO_0312 | BCGDan_sapMKO_0312 | 303 |
| 363826 | 364047 | vapB2 | Rv0300 | 222 | 394799 | 395020 | BCG_0340 | BCG_0340 | 222 | 365131 | 365352 | BCGDan_0314 | BCGDan_0314 | 222 | 365131 | 365352 | BCGDan_sapMKO_0313 | BCGDan_sapMKO_0313 | 222 |
| 364044 | 364469 | vapC2 | Rv0301 | 426 | 395017 | 395442 | BCG_0341 | BCG_0341 | 426 | 365349 | 365774 | BCGDan_0315 | BCGDan_0315 | 426 | 365343 | 365774 | BCGDan_sapMKO_0314 | BCGDan_sapMKO_0314 | 432 |
| 364605 | 365237 | Rv0302 | Rv0302 | 633 | 395579 | 396211 | BCG_0342 | BCG_0342 | 633 | 365911 | 366543 | BCGDan_0316 | BCGDan_0316 | 633 | 365989 | 366543 | BCGDan_sapMKO_0315 | BCGDan_sapMKO_0315 | 555 |
| 365234 | 366142 | Rv0303 | Rv0303 | 909 | 396208 | 397116 | BCG_0343 | BCG_0343 | 909 | 366540 | 367448 | BCGDan_0317 | BCGDan_0317 | 909 | 366540 | 367448 | BCGDan_sapMKO_0316 | BCGDan_sapMKO_0316 | 909 |
| 366150 | 372764 | PPE5 | Rv0304c | 6615 | 397124 | 400567 | PPE5 | BCG_0344c | 3444 | 367456 | 370899 | PPE5 | BCGDan_0318 | 3444 | 367456 | 370899 | PPE5 | BCGDan_sapMKO_0317 | 3444 |
| 372820 | 375711 | PPE6 | Rv0305c | 2892 | 400711 | 406638 | PPE6 | BCG_0345c | 5928 | 371043 | 376970 | PPE6 | BCGDan_0319 | 5928 | 371043 | 376970 | PPE6 | BCGDan_sapMKO_0318 | 5928 |
| 375914 | 376585 | Rv0306 | Rv0306 | 672 | 406841 | 407512 | BCG_0346 | BCG_0346 | 672 | 377173 | 377844 | BCGDan_0320 | BCGDan_0320 | 672 | 377200 | 377844 | BCGDan_sapMKO_0319 | BCGDan_sapMKO_0319 | 645 |
| 376573 | 377055 | Rv0307c | Rv0307c | 483 | 407500 | 407982 | BCG_0347c | BCG_0347c | 483 | 377832 | 378314 | BCGDan_0321 | BCGDan_0321 | 483 | 377832 | 378314 | BCGDan_sapMKO_0320 | BCGDan_sapMKO_0320 | 483 |
| 377113 | 377829 | Rv0308 | Rv0308 | 717 | 408040 | 408756 | BCG_0348 | BCG_0348 | 717 | 378372 | 379088 | BCGDan_0322 | BCGDan_0322 | 717 | 378372 | 379088 | BCGDan_sapMKO_0321 | BCGDan_sapMKO_0321 | 717 |
| 377931 | 378587 | Rv0309 | Rv0309 | 657 | 408858 | 409514 | BCG_0349 | BCG_0349 | 657 | 379190 | 379846 | BCGDan_0323 | BCGDan_0323 | 657 | 379259 | 379846 | BCGDan_sapMKO_0322 | BCGDan_sapMKO_0322 | 588 |
| 378657 | 379148 | Rv0310c | Rv0310c | 492 | 409584 | 410075 | BCG_0350c | BCG_0350c | 492 | 379916 | 380407 | BCGDan_0324 | BCGDan_0324 | 492 | 379916 | 380407 | BCGDan_sapMKO_0323 | BCGDan_sapMKO_0323 | 492 |
| 379172 | 380401 | Rv0311 | Rv0311 | 1230 | 410099 | 411328 | BCG_0351 | BCG_0351 | 1230 | 380431 | 381660 | BCGDan_0325 | BCGDan_0325 | 1230 | 380431 | 381660 | BCGDan_sapMKO_0324 | BCGDan_sapMKO_0324 | 1230 |
| 380556 | 382418 | Rv0312 | Rv0312 | 1863 | 411483 | 413345 | BCG_0352 | BCG_0352 | 1863 | 381815 | 383677 | BCGDan_0326 | BCGDan_0326 | 1863 | 381815 | 383677 | BCGDan_sapMKO_0325 | BCGDan_sapMKO_0325 | 1863 |
| 382490 | 382876 | Rv0313 | Rv0313 | 387 | 413417 | 413803 | BCG_0353 | BCG_0353 | 387 | 383749 | 384135 | BCGDan_0327 | BCGDan_0327 | 387 | 383749 | 384135 | BCGDan_sapMKO_0326 | BCGDan_sapMKO_0326 | 387 |
| 382879 | 383541 | Rv0314c | Rv0314c | 663 | 413806 | 414468 | BCG_0354c | BCG_0354c | 663 | 384138 | 384800 | BCGDan_0328 | BCGDan_0328 | 663 | 384138 | 384773 | BCGDan_sapMKO_0327 | BCGDan_sapMKO_0327 | 636 |
| 383602 | 384486 | Rv0315 | Rv0315 | 885 | 414529 | 415413 | BCG_0355 | BCG_0355 | 885 | 384861 | 385745 | BCGDan_0329 | BCGDan_0329 | 885 | 384891 | 385745 | BCGDan_sapMKO_0328 | BCGDan_sapMKO_0328 | 855 |
| 384535 | 385149 | Rv0316 | Rv0316 | 615 | 415462 | 416076 | BCG_0356 | BCG_0356 | 615 | 385794 | 386408 | BCGDan_0330 | BCGDan_0330 | 615 | 385767 | 386408 | BCGDan_sapMKO_0329 | BCGDan_sapMKO_0329 | 642 |
| 385173 | 385943 | glpQ2 | Rv0317c | 771 | 416100 | 416870 | glpQ2 | BCG_0357c | 771 | 386432 | 387202 | glpQ2 | BCGDan_0331 | 771 | 386432 | 387133 | glpQ2 | BCGDan_sapMKO_0330 | 702 |
| 386305 | 387099 | Rv0318c | Rv0318c | 795 | 417232 | 418026 | BCG_0358c | BCG_0358c | 795 | 387564 | 388358 | BCGDan_0333 | BCGDan_0333 | 795 | 387564 | 388340 | BCGDan_sapMKO_0332 | BCGDan_sapMKO_0332 | 777 |
| 387148 | 387816 | pcp | Rv0319 | 669 | 418075 | 418743 | pcp | BCG_0359 | 669 | 388407 | 389075 | pcp | BCGDan_0334 | 669 | 388407 | 389075 | pcp | BCGDan_sapMKO_0333 | 669 |
| 387888 | 388550 | Rv0320 | Rv0320 | 663 | 418815 | 419477 | BCG_0360 | BCG_0360 | 663 | 389147 | 389809 | BCGDan_0335 | BCGDan_0335 | 663 | 389222 | 389809 | BCGDan_sapMKO_0334 | BCGDan_sapMKO_0334 | 588 |
| 388582 | 389154 | dcd | Rv0321 | 573 | 419509 | 420081 | dcd | BCG_0361 | 573 | 389841 | 390413 | dcd | BCGDan_0336 | 573 | 389841 | 390413 | dcd | BCGDan_sapMKO_0335 | 573 |
| 389260 | 390591 | udgA | Rv0322 | 1332 | 420187 | 421518 | udgA | BCG_0362 | 1332 | 390519 | 391850 | udgA | BCGDan_0337 | 1332 | 390519 | 391850 | udgA | BCGDan_sapMKO_0336 | 1332 |
| 390580 | 391251 | Rv0323c | Rv0323c | 672 | 421507 | 422178 | BCG_0363c | BCG_0363c | 672 | 391839 | 392510 | BCGDan_0338 | BCGDan_0338 | 672 | 391839 | 392474 | BCGDan_sapMKO_0337 | BCGDan_sapMKO_0337 | 636 |
| 391352 | 392032 | Rv0324 | Rv0324 | 681 | 422279 | 422959 | BCG_0364 | BCG_0364 | 681 | 392611 | 393291 | BCGDan_0339 | BCGDan_0339 | 681 | 392611 | 393291 | BCGDan_sapMKO_0338 | BCGDan_sapMKO_0338 | 681 |
| 392039 | 392263 | Rv0325 | Rv0325 | 225 | 422966 | 423655 | BCG_0365 | BCG_0365 | 690 | 393298 | 393987 | BCGDan_0340 | BCGDan_0340 | 690 | 393325 | 393987 | BCGDan_sapMKO_0339 | BCGDan_sapMKO_0339 | 663 |
| 392696 | 394045 | cyp135A1 | Rv0327c | 1350 | 423623 | 424972 | cyp135A1 | BCG_0366c | 1350 | 393955 | 395304 | cyp135A1 | BCGDan_0341 | 1350 | 393955 | 395304 | cyp135A1 | BCGDan_sapMKO_0340 | 1350 |
| 394111 | 394713 | Rv0328 | Rv0328 | 603 | 425038 | 425640 | BCG_0367 | BCG_0367 | 603 | 395370 | 395972 | BCGDan_0342 | BCGDan_0342 | 603 | 395370 | 395972 | BCGDan_sapMKO_0341 | BCGDan_sapMKO_0341 | 603 |
| 394694 | 395320 | Rv0329c | Rv0329c | 627 | 425621 | 426247 | BCG_0368c | BCG_0368c | 627 | 395953 | 396579 | BCGDan_0343 | BCGDan_0343 | 627 | 395953 | 396579 | BCGDan_sapMKO_0342 | BCGDan_sapMKO_0342 | 627 |

| | | | | | | | | | | | | | | | | | | | |
|---|---|---|---|---|---|---|---|---|---|---|---|---|---|---|---|---|---|---|---|
| 395347 | 396087 | Rv0330c | Rv0330c | 741 | 426274 | 427014 | BCG_0369c | BCG_0369c | 741 | 396606 | 397346 | BCGDan_0344 | BCGDan_0344 | 741 | 396606 | 397346 | BCGDan_sapMKO_0343 | BCGDan_sapMKO_0343 | 741 |
| 396201 | 397367 | Rv0331 | Rv0331 | 1167 | 427128 | 428294 | BCG_0370 | BCG_0370 | 1167 | 397460 | 398626 | BCGDan_0345 | BCGDan_0345 | 1167 | 397460 | 398626 | BCGDan_sapMKO_0344 | BCGDan_sapMKO_0344 | 1167 |
| 397442 | 398227 | Rv0332 | Rv0332 | 786 | 428369 | 429154 | BCG_0371 | BCG_0371 | 786 | 398701 | 399486 | BCGDan_0346 | BCGDan_0346 | 786 | 398731 | 399486 | BCGDan_sapMKO_0345 | BCGDan_sapMKO_0345 | 756 |
| 398254 | 398628 | Rv0333 | Rv0333 | 375 | 429181 | 429555 | BCG_0372 | BCG_0372 | 375 | 399513 | 399887 | BCGDan_0347 | BCGDan_0347 | 375 | 399462 | 399887 | BCGDan_sapMKO_0346 | BCGDan_sapMKO_0346 | 426 |
| 398658 | 399524 | rmlA | Rv0334 | 867 | 429585 | 430451 | rmlA | BCG_0373 | 867 | 399917 | 400783 | rmlA | BCGDan_0348 | 867 | 399917 | 400783 | rmlA | BCGDan_sapMKO_0347 | 867 |
| 399535 | 400050 | PE6 | Rv0335c | 516 | 430462 | 430977 | PE6 | BCG_0374c | 516 | 400794 | 401309 | PE6 | BCGDan_0349 | 516 | 400794 | 401309 | PE6 | BCGDan_sapMKO_0348 | 516 |
| 400192 | 401703 | Rv0336 | Rv0336 | 1512 | 431119 | 432630 | BCG_0375 | BCG_0375 | 1512 | 401451 | 402962 | BCGDan_0350 | BCGDan_0350 | 1512 | 401427 | 402962 | BCGDan_sapMKO_0349 | BCGDan_sapMKO_0349 | 1536 |
| 401873 | 403162 | aspC | Rv0337c | 1290 | 432800 | 434089 | aspC | BCG_0376c | 1290 | 403132 | 404421 | aspC | BCGDan_0351 | 1290 | 403132 | 404421 | aspC | BCGDan_sapMKO_0350 | 1290 |
| 403193 | 405841 | Rv0338c | Rv0338c | 2649 | 434120 | 436768 | BCG_0377c | BCG_0377c | 2649 | 404452 | 407100 | BCGDan_0352 | BCGDan_0352 | 2649 | 404452 | 407100 | BCGDan_sapMKO_0351 | BCGDan_sapMKO_0351 | 2649 |
| 405950 | 408448 | Rv0339c | Rv0339c | 2499 | 436877 | 439375 | BCG_0378c | BCG_0378c | 2499 | 407209 | 409707 | BCGDan_0353 | BCGDan_0353 | 2499 | 407209 | 409659 | BCGDan_sapMKO_0352 | BCGDan_sapMKO_0352 | 2451 |
| 408634 | 409173 | Rv0340 | Rv0340 | 540 | 439561 | 440100 | BCG_0379 | BCG_0379 | 540 | 409893 | 410432 | BCGDan_0354 | BCGDan_0354 | 540 | 409893 | 410432 | BCGDan_sapMKO_0353 | BCGDan_sapMKO_0353 | 540 |
| 409362 | 410801 | iniB | Rv0341 | 1440 | 440289 | 441728 | iniB | BCG_0380 | 1440 | 410621 | 412060 | iniB | BCGDan_0355 | 1440 | 410615 | 412060 | iniB | BCGDan_sapMKO_0354 | 1446 |
| 410838 | 412760 | iniA | Rv0342 | 1923 | 441765 | 443687 | iniA | BCG_0381 | 1923 | 412097 | 414019 | iniA | BCGDan_0356 | 1923 | 412166 | 414019 | iniA | BCGDan_sapMKO_0355 | 1854 |
| 412757 | 414238 | iniC | Rv0343 | 1482 | 443684 | 445165 | iniC | BCG_0382 | 1482 | 414016 | 415497 | iniC | BCGDan_0357 | 1482 | 414016 | 415497 | iniC | BCGDan_sapMKO_0356 | 1482 |
| 414381 | 414941 | lpqJ | Rv0344c | 561 | 445308 | 445868 | lpqJ | BCG_0383c | 561 | 415640 | 416200 | lpqJ | BCGDan_0358 | 561 | 415640 | 416200 | lpqJ | BCGDan_sapMKO_0357 | 561 |
| 415050 | 415460 | Rv0345 | Rv0345 | 411 | 445977 | 446387 | BCG_0384 | BCG_0384 | 411 | 416309 | 416719 | BCGDan_0359 | BCGDan_0359 | 411 | 416303 | 416719 | BCGDan_sapMKO_0358 | BCGDan_sapMKO_0358 | 417 |
| 415502 | 416965 | ansP2 | Rv0346c | 1464 | 446429 | 447892 | ansP2 | BCG_0385c | 1464 | 416761 | 418224 | ansP2 | BCGDan_0360 | 1464 | 416761 | 418224 | ansP2 | BCGDan_sapMKO_0359 | 1464 |
| 417304 | 418290 | Rv0347 | Rv0347 | 987 | 448231 | 449217 | BCG_0386 | BCG_0386 | 987 | 418563 | 419549 | BCGDan_0361 | BCGDan_0361 | 987 | 418563 | 419549 | BCGDan_sapMKO_0360 | BCGDan_sapMKO_0360 | 987 |
| 418293 | 418946 | Rv0348 | Rv0348 | 654 | 449220 | 449873 | BCG_0387 | BCG_0387 | 654 | 419552 | 420205 | BCGDan_0362 | BCGDan_0362 | 654 | 419552 | 420205 | BCGDan_sapMKO_0361 | BCGDan_sapMKO_0361 | 654 |
| 418949 | 419608 | Rv0349 | Rv0349 | 660 | 449876 | 450535 | BCG_0388 | BCG_0388 | 660 | 420208 | 420867 | BCGDan_0363 | BCGDan_0363 | 660 | 420208 | 420867 | BCGDan_sapMKO_0362 | BCGDan_sapMKO_0362 | 660 |
| 419835 | 421712 | dnaK | Rv0350 | 1878 | 450762 | 452639 | dnaK | BCG_0389 | 1878 | 421094 | 422971 | dnaK | BCGDan_0364 | 1878 | 421094 | 422971 | dnaK | BCGDan_sapMKO_0363 | 1878 |
| 421709 | 422416 | grpE | Rv0351 | 708 | 452636 | 453343 | grpE | BCG_0390 | 708 | 422968 | 423675 | grpE | BCGDan_0365 | 708 | 422968 | 423675 | grpE | BCGDan_sapMKO_0364 | 708 |
| 422452 | 423639 | dnaJ1 | Rv0352 | 1188 | 453379 | 454566 | dnaJ1 | BCG_0391 | 1188 | 423711 | 424898 | dnaJ1 | BCGDan_0366 | 1188 | 423711 | 424898 | dnaJ1 | BCGDan_sapMKO_0365 | 1188 |
| 423639 | 424019 | hspR | Rv0353 | 381 | 454566 | 454946 | hspR | BCG_0392 | 381 | 424898 | 425278 | hspR | BCGDan_0367 | 381 | 424898 | 425278 | hspR | BCGDan_sapMKO_0366 | 381 |
| 424269 | 424694 | PPE7 | Rv0354c | 426 | 454969 | 465492 | PPE8 | BCG_0393c | 10524 | 425301 | 435824 | PPE8 | BCGDan_0368 | 10524 | 425301 | 435824 | PPE8 | BCGDan_sapMKO_0367 | 10524 |
| 434830 | 435474 | Rv0356c | Rv0356c | 645 | 465643 | 466287 | BCG_0394c | BCG_0394c | 645 | 435975 | 436619 | BCGDan_0369 | BCGDan_0369 | 645 | 435975 | 436604 | BCGDan_sapMKO_0368 | BCGDan_sapMKO_0368 | 630 |
| 435471 | 436769 | purA | Rv0357c | 1299 | 466284 | 467582 | purA | BCG_0395c | 1299 | 436616 | 437914 | purA | BCGDan_0370 | 1299 | 436616 | 437914 | purA | BCGDan_sapMKO_0369 | 1299 |
| 436860 | 437507 | Rv0358 | Rv0358 | 648 | 467673 | 468320 | BCG_0396 | BCG_0396 | 648 | 438005 | 438652 | BCGDan_0371 | BCGDan_0371 | 648 | 438005 | 438652 | BCGDan_sapMKO_0370 | BCGDan_sapMKO_0370 | 648 |
| 437518 | 438297 | Rv0359 | Rv0359 | 780 | 468331 | 469110 | BCG_0397 | BCG_0397 | 780 | 438663 | 439442 | BCGDan_0372 | BCGDan_0372 | 780 | 438663 | 439442 | BCGDan_sapMKO_0371 | BCGDan_sapMKO_0371 | 780 |
| 438302 | 438739 | Rv0360c | Rv0360c | 438 | 469115 | 469552 | BCG_0398c | BCG_0398c | 438 | 439447 | 439884 | BCGDan_0373 | BCGDan_0373 | 438 | 439447 | 439860 | BCGDan_sapMKO_0372 | BCGDan_sapMKO_0372 | 414 |
| 438822 | 439649 | Rv0361 | Rv0361 | 828 | 469635 | 470462 | BCG_0399 | BCG_0399 | 828 | 439967 | 440794 | BCGDan_0374 | BCGDan_0374 | 828 | 439967 | 440794 | BCGDan_sapMKO_0373 | BCGDan_sapMKO_0373 | 828 |
| 439871 | 441253 | mgtE | Rv0362 | 1383 | 470684 | 472066 | mgtE | BCG_0400 | 1383 | 441016 | 442398 | mgtE | BCGDan_0375 | 1383 | 441016 | 442398 | mgtE | BCGDan_sapMKO_0374 | 1383 |
| 441265 | 442299 | fba | Rv0363c | 1035 | 472078 | 473112 | fba | BCG_0401c | 1035 | 442410 | 443444 | fba | BCGDan_0376 | 1035 | 442410 | 443444 | fba | BCGDan_sapMKO_0375 | 1035 |
| 442395 | 443078 | Rv0364 | Rv0364 | 684 | 473208 | 473891 | BCG_0402 | BCG_0402 | 684 | 443540 | 444223 | BCGDan_0377 | BCGDan_0377 | 684 | 443540 | 444223 | BCGDan_sapMKO_0376 | BCGDan_sapMKO_0376 | 684 |
| 443067 | 444197 | Rv0365c | Rv0365c | 1131 | 473880 | 475010 | BCG_0403c | BCG_0403c | 1131 | 444212 | 445342 | BCGDan_0378 | BCGDan_0378 | 1131 | 444212 | 445342 | BCGDan_sapMKO_0377 | BCGDan_sapMKO_0377 | 1131 |
| 444222 | 444815 | Rv0366c | Rv0366c | 594 | 475035 | 475628 | BCG_0404c | BCG_0404c | 594 | 445367 | 445960 | BCGDan_0379 | BCGDan_0379 | 594 | 445367 | 445960 | BCGDan_sapMKO_0378 | BCGDan_sapMKO_0378 | 594 |
| 444844 | 445233 | Rv0367c | Rv0367c | 390 | 475657 | 476046 | BCG_0405c | BCG_0405c | 390 | 445989 | 446378 | BCGDan_0380 | BCGDan_0380 | 390 | 445989 | 446378 | BCGDan_sapMKO_0379 | BCGDan_sapMKO_0379 | 390 |
| 445314 | 446525 | Rv0368c | Rv0368c | 1212 | 476127 | 477338 | BCG_0406c | BCG_0406c | 1212 | 446459 | 447670 | BCGDan_0381 | BCGDan_0381 | 1212 | 446459 | 447679 | BCGDan_sapMKO_0380 | BCGDan_sapMKO_0380 | 1221 |
| 446531 | 447046 | Rv0369c | Rv0369c | 516 | 477344 | 477859 | BCG_0407c | BCG_0407c | 516 | 447676 | 448191 | BCGDan_0382 | BCGDan_0382 | 516 | 447676 | 448278 | BCGDan_sapMKO_0381 | BCGDan_sapMKO_0381 | 603 |
| 447147 | 448043 | Rv0370c | Rv0370c | 897 | 477960 | 478856 | BCG_0408c | BCG_0408c | 897 | 448292 | 449188 | BCGDan_0383 | BCGDan_0383 | 897 | 448292 | 449188 | BCGDan_sapMKO_0382 | BCGDan_sapMKO_0382 | 897 |
| 448040 | 448633 | Rv0371c | Rv0371c | 594 | 478853 | 479446 | BCG_0409c | BCG_0409c | 594 | 449185 | 449778 | BCGDan_0384 | BCGDan_0384 | 594 | 449185 | 449778 | BCGDan_sapMKO_0383 | BCGDan_sapMKO_0383 | 594 |
| 448630 | 449385 | Rv0372c | Rv0372c | 756 | 479443 | 480198 | BCG_0410c | BCG_0410c | 756 | 449775 | 450530 | BCGDan_0385 | BCGDan_0385 | 756 | 449775 | 450530 | BCGDan_sapMKO_0384 | BCGDan_sapMKO_0384 | 756 |
| 449404 | 451803 | Rv0373c | Rv0373c | 2400 | 480217 | 482616 | BCG_0411c | BCG_0411c | 2400 | 450549 | 452948 | BCGDan_0386 | BCGDan_0386 | 2400 | 450549 | 452948 | BCGDan_sapMKO_0385 | BCGDan_sapMKO_0385 | 2400 |
| 451800 | 452279 | Rv0374c | Rv0374c | 480 | 482613 | 483092 | BCG_0412c | BCG_0412c | 480 | 452945 | 453424 | BCGDan_0387 | BCGDan_0387 | 480 | 452945 | 453424 | BCGDan_sapMKO_0386 | BCGDan_sapMKO_0386 | 480 |
| 452294 | 453154 | Rv0375c | Rv0375c | 861 | 483107 | 483967 | BCG_0413c | BCG_0413c | 861 | 453439 | 454299 | BCGDan_0388 | BCGDan_0388 | 861 | 453439 | 454341 | BCGDan_sapMKO_0387 | BCGDan_sapMKO_0387 | 903 |
| 453230 | 454372 | Rv0376c | Rv0376c | 1143 | 484043 | 485185 | BCG_0414c | BCG_0414c | 1143 | 454375 | 455517 | BCGDan_0389 | BCGDan_0389 | 1143 | 454375 | 455541 | BCGDan_sapMKO_0388 | BCGDan_sapMKO_0388 | 1167 |
| 454421 | 455386 | Rv0377 | Rv0377 | 966 | 485234 | 486199 | BCG_0415 | BCG_0415 | 966 | 455566 | 456531 | BCGDan_0390 | BCGDan_0390 | 966 | 455566 | 456531 | BCGDan_sapMKO_0389 | BCGDan_sapMKO_0389 | 966 |
| 455637 | 455858 | Rv0378 | Rv0378 | 222 | 486450 | 486671 | BCG_0416 | BCG_0416 | 222 | 456776 | 457003 | BCGDan_0391 | BCGDan_0391 | 228 | 456776 | 457003 | BCGDan_sapMKO_0390 | BCGDan_sapMKO_0390 | 228 |
| 455977 | 456192 | secE2 | Rv0379 | 216 | 486790 | 487005 | secE2 | BCG_0417 | 216 | 457122 | 457337 | secE2 | BCGDan_0392 | 216 | 457122 | 457337 | secE2 | BCGDan_sapMKO_0391 | 216 |
| 456268 | 456819 | Rv0380c | Rv0380c | 552 | 487081 | 487632 | BCG_0418c | BCG_0418c | 552 | 457413 | 457964 | BCGDan_0393 | BCGDan_0393 | 552 | 457413 | 458012 | BCGDan_sapMKO_0392 | BCGDan_sapMKO_0392 | 600 |
| 456915 | 457823 | Rv0381c | Rv0381c | 909 | 487728 | 488636 | BCG_0419c | BCG_0419c | 909 | 458060 | 458968 | BCGDan_0394 | BCGDan_0394 | 909 | 458060 | 458968 | BCGDan_sapMKO_0393 | BCGDan_sapMKO_0393 | 909 |
| 457841 | 458380 | pyrE | Rv0382c | 540 | 488654 | 489193 | pyrE | BCG_0420c | 540 | 458986 | 459525 | pyrE | BCGDan_0395 | 540 | 458986 | 459525 | pyrE | BCGDan_sapMKO_0394 | 540 |
| 458461 | 459315 | Rv0383c | Rv0383c | 855 | 489274 | 490128 | BCG_0421c | BCG_0421c | 855 | 459606 | 460460 | BCGDan_0396 | BCGDan_0396 | 855 | 459606 | 460460 | BCGDan_sapMKO_0395 | BCGDan_sapMKO_0395 | 855 |
| 459456 | 462002 | clpB | Rv0384c | 2547 | 490269 | 492815 | clpB | BCG_0422c | 2547 | 460601 | 463147 | clpB | BCGDan_0397 | 2547 | 460601 | 463234 | clpB | BCGDan_sapMKO_0396 | 2634 |
| 462135 | 463307 | Rv0385 | Rv0385 | 1173 | 492948 | 494120 | BCG_0423 | BCG_0423 | 1173 | 463280 | 464452 | BCGDan_0398 | BCGDan_0398 | 1173 | 463280 | 464452 | BCGDan_sapMKO_0397 | BCGDan_sapMKO_0397 | 1173 |
| 463411 | 466668 | Rv0386 | Rv0386 | 3258 | 494224 | 497481 | BCG_0424 | BCG_0424 | 3258 | 464556 | 467813 | BCGDan_0399 | BCGDan_0399 | 3258 | 464556 | 467813 | BCGDan_sapMKO_0398 | BCGDan_sapMKO_0398 | 3258 |
| 466672 | 467406 | Rv0387c | Rv0387c | 735 | 497485 | 498816 | PPE9 | BCG_0425c | 1332 | 467817 | 469148 | PPE9 | BCGDan_0400 | 1332 | 467817 | 469148 | PPE9 | BCGDan_sapMKO_0399 | 1332 |
| 468335 | 469594 | purT | Rv0389 | 1260 | 499150 | 500409 | purT | BCG_0426 | 1260 | 469482 | 470741 | purT | BCGDan_0401 | 1260 | 469566 | 470741 | purT | BCGDan_sapMKO_0400 | 1176 |
| 469591 | 470013 | Rv0390 | Rv0390 | 423 | 500406 | 500828 | BCG_0427 | BCG_0427 | 423 | 470738 | 471160 | BCGDan_0402 | BCGDan_0402 | 423 | 470738 | 471160 | BCGDan_sapMKO_0401 | BCGDan_sapMKO_0401 | 423 |
| 470010 | 471230 | metZ | Rv0391 | 1221 | 500825 | 502045 | metZ | BCG_0428 | 1221 | 471157 | 472377 | metZ | BCGDan_0403 | 1221 | 471157 | 472377 | metZ | BCGDan_sapMKO_0402 | 1221 |
| 471227 | 472639 | ndhA | Rv0392c | 1413 | 502042 | 503454 | ndhA | BCG_0429c | 1413 | 472374 | 473786 | ndhA | BCGDan_0404 | 1413 | 472374 | 473786 | ndhA | BCGDan_sapMKO_0403 | 1413 |
| 472781 | 474106 | Rv0393 | Rv0393 | 1326 | 503596 | 504921 | BCG_0430 | BCG_0430 | 1326 | 473928 | 475253 | BCGDan_0405 | BCGDan_0405 | 1326 | 473928 | 475253 | BCGDan_sapMKO_0404 | BCGDan_sapMKO_0404 | 1326 |
| 474122 | 474841 | Rv0394c | Rv0394c | 720 | 504937 | 505656 | BCG_0431c | BCG_0431c | 720 | 475269 | 475988 | BCGDan_0406 | BCGDan_0406 | 720 | 475269 | 475988 | BCGDan_sapMKO_0405 | BCGDan_sapMKO_0405 | 720 |
| 474940 | 475344 | Rv0395 | Rv0395 | 405 | 505755 | 506159 | BCG_0432 | BCG_0432 | 405 | 476087 | 476491 | BCGDan_0407 | BCGDan_0407 | 405 | 476051 | 476491 | BCGDan_sapMKO_0406 | BCGDan_sapMKO_0406 | 441 |
| 475350 | 475742 | Rv0396 | Rv0396 | 393 | 506165 | 506557 | BCG_0433 | BCG_0433 | 393 | 476497 | 476889 | BCGDan_0408 | BCGDan_0408 | 393 | 476497 | 476889 | BCGDan_sapMKO_0407 | BCGDan_sapMKO_0407 | 393 |
| 475816 | 476184 | Rv0397 | Rv0397 | 369 | 506631 | 506999 | BCG_0434 | BCG_0434 | 369 | 476963 | 477331 | BCGDan_0409 | BCGDan_0409 | 369 | 476963 | 477331 | BCGDan_sapMKO_0408 | BCGDan_sapMKO_0408 | 369 |
| 476679 | 477320 | Rv0398c | Rv0398c | 642 | 507494 | 508135 | BCG_0435c | BCG_0435c | 642 | 477826 | 478467 | BCGDan_0410 | BCGDan_0410 | 642 | 477826 | 478467 | BCGDan_sapMKO_0409 | BCGDan_sapMKO_0409 | 642 |
| 477327 | 478556 | lpqK | Rv0399c | 1230 | 508142 | 509371 | lpqK | BCG_0436c | 1230 | 478474 | 479703 | lpqK | BCGDan_0411 | 1230 | 478474 | 479703 | lpqK | BCGDan_sapMKO_0410 | 1230 |
| 478566 | 479753 | fadE7 | Rv0400c | 1188 | 509381 | 510568 | fadE7 | BCG_0437c | 1188 | 479713 | 480900 | fadE7 | BCGDan_0412 | 1188 | 479713 | 480900 | fadE7 | BCGDan_sapMKO_0411 | 1188 |
| 479789 | 480160 | Rv0401 | Rv0401 | 372 | 510604 | 510975 | BCG_0438 | BCG_0438 | 372 | 480936 | 481307 | BCGDan_0413 | BCGDan_0413 | 372 | 480936 | 481307 | BCGDan_sapMKO_0412 | BCGDan_sapMKO_0412 | 372 |
| 480355 | 483231 | mmpL1 | Rv0402c | 2877 | 511170 | 512273 | mmpL1b | BCG_0439c | 1104 | 481502 | 482608 | mmpL1b | BCGDan_0414 | 1107 | 481502 | 482617 | mmpL1b | BCGDan_sapMKO_0413 | 1116 |
| NA | NA | NA | NA | NA | 512270 | 514045 | mmpL1a | BCG_0440c | 1776 | 482602 | 484377 | mmpL1a | BCGDan_0415 | 1776 | 482602 | 484377 | mmpL1a | BCGDan_sapMKO_0414 | 1776 |
| 483228 | 483656 | mmpS1 | Rv0403c | 429 | 514042 | 514470 | mmpS1 | BCG_0441c | 429 | 484374 | 484802 | mmpS1 | BCGDan_0416 | 429 | 484374 | 484802 | mmpS1 | BCGDan_sapMKO_0415 | 429 |
| 483977 | 485734 | fadD30 | Rv0404 | 1758 | 514791 | 516548 | fadD30 | BCG_0442 | 1758 | 485123 | 486880 | fadD30 | BCGDan_0417 | 1758 | 485123 | 486880 | fadD30 | BCGDan_sapMKO_0416 | 1758 |
| 485731 | 489939 | pks6 | Rv0405 | 4209 | 516545 | 517927 | pks6a | BCG_0443 | 1383 | 486877 | 488259 | pks6a | BCGDan_0418 | 1383 | 486877 | 488259 | pks6a | BCGDan_sapMKO_0417 | 1383 |
| NA | NA | NA | NA | NA | 517914 | 520754 | pks6b | BCG_0444 | 2841 | 488246 | 491086 | pks6b | BCGDan_0419 | 2841 | 488393 | 491086 | pks6b | BCGDan_sapMKO_0418 | 2694 |
| 489887 | 490705 | Rv0406c | Rv0406c | 819 | 520702 | 521520 | BCG_0445c | BCG_0445c | 819 | 491034 | 491852 | BCGDan_0420 | BCGDan_0420 | 819 | 491034 | 491852 | BCGDan_sapMKO_0419 | BCGDan_sapMKO_0419 | 819 |
| 490783 | 491793 | fgd1 | Rv0407 | 1011 | 521598 | 522608 | fgd1 | BCG_0446 | 1011 | 491930 | 492940 | fgd1 | BCGDan_0421 | 1011 | 491930 | 492940 | fgd1 | BCGDan_sapMKO_0420 | 1011 |
| 491786 | 493858 | pta | Rv0408 | 2073 | 522601 | 524673 | pta | BCG_0447 | 2073 | 492933 | 495005 | pta | BCGDan_0422 | 2073 | 492933 | 495005 | pta | BCGDan_sapMKO_0421 | 2073 |
| 493851 | 495008 | ackA | Rv0409 | 1158 | 524666 | 525823 | ackA | BCG_0448 | 1158 | 494998 | 496155 | ackA | BCGDan_0423 | 1158 | 494998 | 496155 | ackA | BCGDan_sapMKO_0422 | 1158 |
| 495062 | 497314 | pknG | Rv0410c | 2253 | 525877 | 528129 | pknG | BCG_0449c | 2253 | 496209 | 498461 | pknG | BCGDan_0424 | 2253 | 496209 | 498518 | pknG | BCGDan_sapMKO_0423 | 2310 |
| 497314 | 498300 | glnH | Rv0411c | 987 | 528129 | 529115 | glnH | BCG_0450c | 987 | 498461 | 499447 | glnH | BCGDan_0425 | 987 | 498461 | 499447 | glnH | BCGDan_sapMKO_0424 | 987 |
| 498300 | 499619 | Rv0412c | Rv0412c | 1320 | 529115 | 530434 | BCG_0451c | BCG_0451c | 1320 | 499447 | 500766 | BCGDan_0426 | BCGDan_0426 | 1320 | 499447 | 500766 | BCGDan_sapMKO_0425 | BCGDan_sapMKO_0425 | 1320 |
| 499713 | 500366 | mutT3 | Rv0413 | 654 | 530528 | 531181 | mutT3 | BCG_0452 | 654 | 500860 | 501513 | mutT3 | BCGDan_0427 | 654 | 500893 | 501513 | mutT3 | BCGDan_sapMKO_0426 | 621 |
| 500350 | 501018 | thiE | Rv0414c | 669 | 531165 | 531833 | thiE | BCG_0453c | 669 | 501497 | 502165 | thiE | BCGDan_0428 | 669 | 501497 | 502165 | thiE | BCGDan_sapMKO_0427 | 669 |
| 501148 | 502170 | thiO | Rv0415 | 1023 | 531963 | 532985 | thiO | BCG_0454 | 1023 | 502295 | 503317 | thiO | BCGDan_0429 | 1023 | 502295 | 503317 | thiO | BCGDan_sapMKO_0428 | 1023 |

| | | | | | | | | | | | | | | | | | | | |
|---|---|---|---|---|---|---|---|---|---|---|---|---|---|---|---|---|---|---|---|
| 502167 | 502373 | thiS | Rv0416 | 207 | 532982 | 533188 | thiS | BCG_0455 | 207 | 503314 | 503520 | thiS | BCGDan_0430 | 207 | 503314 | 503520 | thiS | BCGDan_sapMKO_0429 | 207 |
| 502366 | 503124 | thiG | Rv0417 | 759 | 533181 | 533939 | thiG | BCG_0456 | 759 | 503513 | 504271 | thiG | BCGDan_0431 | 759 | 503513 | 504271 | thiG | BCGDan_sapMKO_0430 | 759 |
| 503496 | 504998 | lpqL | Rv0418 | 1503 | 534311 | 535813 | lpqL | BCG_0457 | 1503 | 504643 | 506145 | lpqL | BCGDan_0432 | 1503 | 504643 | 506145 | lpqL | BCGDan_sapMKO_0431 | 1503 |
| 505086 | 506582 | lpqM | Rv0419 | 1497 | 535901 | 537397 | lpqM | BCG_0458 | 1497 | 506233 | 507729 | lpqM | BCGDan_0433 | 1497 | 506287 | 507729 | lpqM | BCGDan_sapMKO_0432 | 1443 |
| 506561 | 506971 | Rv0420c | Rv0420c | 411 | 537376 | 537786 | BCG_0459c | BCG_0459c | 411 | 507708 | 508118 | BCGDan_0434 | BCGDan_0434 | 411 | 507708 | 508118 | BCGDan_sapMKO_0433 | BCGDan_sapMKO_0433 | 411 |
| 507132 | 507761 | Rv0421c | Rv0421c | 630 | 537947 | 538576 | BCG_0460c | BCG_0460c | 630 | 508279 | 508908 | BCGDan_0435 | BCGDan_0435 | 630 | 508279 | 508908 | BCGDan_sapMKO_0434 | BCGDan_sapMKO_0434 | 630 |
| 507758 | 508555 | thiD | Rv0422c | 798 | 538573 | 539370 | thiD | BCG_0461c | 798 | 508905 | 509702 | thiD | BCGDan_0436 | 798 | 508905 | 509732 | thiD | BCGDan_sapMKO_0435 | 828 |
| 508582 | 510225 | thiC | Rv0423c | 1644 | 539397 | 541040 | thiC | BCG_0462c | 1644 | 509729 | 511372 | thiC | BCGDan_0437 | 1644 | 509729 | 511372 | thiC | BCGDan_sapMKO_0436 | 1644 |
| 510377 | 510652 | Rv0424c | Rv0424c | 276 | 541192 | 541467 | BCG_0463c | BCG_0463c | 276 | 511524 | 511799 | BCGDan_0438 | BCGDan_0438 | 276 | 511524 | 511802 | BCGDan_sapMKO_0437 | BCGDan_sapMKO_0437 | 279 |
| 510702 | 515321 | ctpH | Rv0425c | 4620 | 541517 | 546136 | ctpH | BCG_0464c | 4620 | 511849 | 516468 | ctpH | BCGDan_0439 | 4620 | 511849 | 516468 | ctpH | BCGDan_sapMKO_0438 | 4620 |
| 515373 | 515816 | Rv0426c | Rv0426c | 444 | 546188 | 546631 | BCG_0465c | BCG_0465c | 444 | 516520 | 516963 | BCGDan_0440 | BCGDan_0440 | 444 | 516520 | 516921 | BCGDan_sapMKO_0439 | BCGDan_sapMKO_0439 | 402 |
| 516017 | 516892 | xthA | Rv0427c | 876 | 546832 | 547707 | xthA | BCG_0466c | 876 | 517164 | 518039 | xthA | BCGDan_0441 | 876 | 517164 | 518039 | xthA | BCGDan_sapMKO_0440 | 876 |
| 516895 | 517803 | Rv0428c | Rv0428c | 909 | 547710 | 548618 | BCG_0467c | BCG_0467c | 909 | 518042 | 518950 | BCGDan_0442 | BCGDan_0442 | 909 | 518042 | 518914 | BCGDan_sapMKO_0441 | BCGDan_sapMKO_0441 | 873 |
| 517803 | 518396 | def | Rv0429c | 594 | 548618 | 549211 | def | BCG_0468c | 594 | 518950 | 519543 | def | BCGDan_0443 | 594 | 518950 | 519543 | def | BCGDan_sapMKO_0442 | 594 |
| 518733 | 519041 | Rv0430 | Rv0430 | 309 | 549548 | 549856 | BCG_0469 | BCG_0469 | 309 | 519880 | 520188 | BCGDan_0444 | BCGDan_0444 | 309 | 519880 | 520188 | BCGDan_sapMKO_0443 | BCGDan_sapMKO_0443 | 309 |
| 519073 | 519567 | Rv0431 | Rv0431 | 495 | 549888 | 550382 | BCG_0470 | BCG_0470 | 495 | 520220 | 520714 | BCGDan_0445 | BCGDan_0445 | 495 | 520286 | 520714 | BCGDan_sapMKO_0444 | BCGDan_sapMKO_0444 | 429 |
| 519600 | 520322 | sodC | Rv0432 | 723 | 550415 | 551137 | sodC | BCG_0471 | 723 | 520747 | 521469 | sodC | BCGDan_0446 | 723 | 520747 | 521469 | sodC | BCGDan_sapMKO_0445 | 723 |
| 520324 | 521454 | Rv0433 | Rv0433 | 1131 | 551139 | 552269 | BCG_0472 | BCG_0472 | 1131 | 521471 | 522601 | BCGDan_0447 | BCGDan_0447 | 1131 | 521534 | 522601 | BCGDan_sapMKO_0446 | BCGDan_sapMKO_0446 | 1068 |
| 521514 | 522167 | Rv0434 | Rv0434 | 654 | 552329 | 552982 | BCG_0473 | BCG_0473 | 654 | 522661 | 523314 | BCGDan_0448 | BCGDan_0448 | 654 | 522661 | 523314 | BCGDan_sapMKO_0447 | BCGDan_sapMKO_0447 | 654 |
| 522347 | 524533 | Rv0435c | Rv0435c | 2187 | 553162 | 555348 | BCG_0474c | BCG_0474c | 2187 | 523494 | 525680 | BCGDan_0449 | BCGDan_0449 | 2187 | 523494 | 525680 | BCGDan_sapMKO_0448 | BCGDan_sapMKO_0448 | 2187 |
| 524530 | 525390 | pssA | Rv0436c | 861 | 555345 | 556205 | pssA | BCG_0475c | 861 | 525677 | 526537 | pssA | BCGDan_0450 | 861 | 525677 | 526543 | pssA | BCGDan_sapMKO_0449 | 867 |
| 525387 | 526082 | psd | Rv0437c | 696 | 556202 | 556897 | psd | BCG_0476c | 696 | 526534 | 527229 | psd | BCGDan_0451 | 696 | 526534 | 527229 | psd | BCGDan_sapMKO_0450 | 696 |
| 526143 | 527360 | moeA2 | Rv0438c | 1218 | 556958 | 558175 | moeA2 | BCG_0477c | 1218 | 527290 | 528507 | moeA2 | BCGDan_0452 | 1218 | 527290 | 528468 | moeA2 | BCGDan_sapMKO_0451 | 1179 |
| 527379 | 528314 | Rv0439c | Rv0439c | 936 | 558194 | 559129 | BCG_0478c | BCG_0478c | 936 | 528526 | 529461 | BCGDan_0453 | BCGDan_0453 | 936 | 528526 | 529461 | BCGDan_sapMKO_0452 | BCGDan_sapMKO_0452 | 936 |
| 528608 | 530230 | groEL2 | Rv0440 | 1623 | 559423 | 561045 | groEL2 | BCG_0479 | 1623 | 529755 | 531377 | groEL2 | BCGDan_0454 | 1623 | 529755 | 531377 | groEL2 | BCGDan_sapMKO_0453 | 1623 |
| 530296 | 530724 | Rv0441c | Rv0441c | 429 | 561111 | 561539 | BCG_0480c | BCG_0480c | 429 | 531443 | 531871 | BCGDan_0455 | BCGDan_0455 | 429 | 531443 | 531871 | BCGDan_sapMKO_0454 | BCGDan_sapMKO_0454 | 429 |
| 530751 | 532214 | PPE10 | Rv0442c | 1464 | 561566 | 563005 | PPE10 | BCG_0481c | 1440 | 531898 | 533337 | PPE10 | BCGDan_0456 | 1440 | 531898 | 533337 | PPE10 | BCGDan_sapMKO_0455 | 1440 |
| 532396 | 532911 | Rv0443 | Rv0443 | 516 | 563211 | 563726 | BCG_0482 | BCG_0482 | 516 | 533543 | 534058 | BCGDan_0457 | BCGDan_0457 | 516 | 533543 | 534058 | BCGDan_sapMKO_0456 | BCGDan_sapMKO_0456 | 516 |
| 533091 | 533789 | rskA | Rv0444c | 699 | 563906 | 564604 | BCG_0483c | BCG_0483c | 699 | 534238 | 534936 | BCGDan_0458 | BCGDan_0458 | 699 | 534238 | 534936 | BCGDan_sapMKO_0457 | BCGDan_sapMKO_0457 | 699 |
| 533833 | 534396 | sigK | Rv0445c | 564 | 564648 | 565211 | sigK | BCG_0484c | 564 | 534980 | 535543 | sigK | BCGDan_0459 | 564 | 534980 | 535501 | sigK | BCGDan_sapMKO_0458 | 522 |
| 534445 | 535215 | Rv0446c | Rv0446c | 771 | 565260 | 566030 | BCG_0485c | BCG_0485c | 771 | 535592 | 536362 | BCGDan_0460 | BCGDan_0460 | 771 | 535592 | 536374 | BCGDan_sapMKO_0459 | BCGDan_sapMKO_0459 | 783 |
| 535224 | 536507 | ufaA1 | Rv0447c | 1284 | 566039 | 567322 | ufaA1 | BCG_0486c | 1284 | 536371 | 537654 | ufaA1 | BCGDan_0461 | 1284 | 536371 | 537654 | ufaA1 | BCGDan_sapMKO_0460 | 1284 |
| 536504 | 537169 | Rv0448c | Rv0448c | 666 | 567319 | 567984 | BCG_0487c | BCG_0487c | 666 | 537651 | 538316 | BCGDan_0462 | BCGDan_0462 | 666 | 537651 | 538382 | BCGDan_sapMKO_0461 | BCGDan_sapMKO_0461 | 732 |
| 537229 | 538548 | Rv0449c | Rv0449c | 1320 | 568044 | 569363 | BCG_0488c | BCG_0488c | 1320 | 538376 | 539695 | BCGDan_0463 | BCGDan_0463 | 1320 | 538376 | 539671 | BCGDan_sapMKO_0462 | BCGDan_sapMKO_0462 | 1296 |
| 538588 | 541491 | mmpL4 | Rv0450c | 2904 | 569403 | 572306 | mmpL4 | BCG_0489c | 2904 | 539735 | 542638 | mmpL4 | BCGDan_0464 | 2904 | 539735 | 542638 | mmpL4 | BCGDan_sapMKO_0463 | 2904 |
| 541488 | 541910 | mmpS4 | Rv0451c | 423 | 572303 | 572725 | mmpS4 | BCG_0490c | 423 | 542635 | 543051 | mmpS4 | BCGDan_0465 | 417 | 542635 | 543051 | mmpS4 | BCGDan_sapMKO_0464 | 417 |
| 542142 | 542852 | Rv0452 | Rv0452 | 711 | 572957 | 573667 | BCG_0491 | BCG_0491 | 711 | 543289 | 543999 | BCGDan_0466 | BCGDan_0466 | 711 | 543289 | 543999 | BCGDan_sapMKO_0465 | BCGDan_sapMKO_0465 | 711 |
| 543174 | 544730 | PPE11 | Rv0453 | 1557 | 573989 | 575545 | PPE11 | BCG_0492 | 1557 | 544321 | 545877 | PPE11 | BCGDan_0467 | 1557 | 544342 | 545877 | PPE11 | BCGDan_sapMKO_0466 | 1536 |
| 544835 | 545185 | Rv0454 | Rv0454 | 351 | 575650 | 576000 | BCG_0493 | BCG_0493 | 351 | 545982 | 546332 | BCGDan_0468 | BCGDan_0468 | 351 | 545982 | 546332 | BCGDan_sapMKO_0467 | BCGDan_sapMKO_0467 | 351 |
| 545375 | 545821 | Rv0455c | Rv0455c | 447 | 576190 | 576636 | BCG_0494c | BCG_0494c | 447 | 546522 | 546968 | BCGDan_0469 | BCGDan_0469 | 447 | 546522 | 546968 | BCGDan_sapMKO_0468 | BCGDan_sapMKO_0468 | 447 |
| 545889 | 546803 | echA2 | Rv0456c | 915 | 576704 | 577618 | echA2 | BCG_0495c | 915 | 547036 | 547950 | echA2 | BCGDan_0470 | 915 | 547036 | 547950 | echA2 | BCGDan_sapMKO_0469 | 915 |
| 547076 | 547357 | mazF1 | Rv0456A | 282 | 577891 | 578172 | BCG_0496c | BCG_0496c | 282 | 548223 | 548504 | BCGDan_0471 | BCGDan_0471 | 282 | 548223 | 548504 | BCGDan_sapMKO_0470 | BCGDan_sapMKO_0470 | 282 |
| 547586 | 549607 | Rv0457c | Rv0457c | 2022 | 578401 | 580431 | BCG_0497c | BCG_0497c | 2031 | 548733 | 550763 | BCGDan_0472 | BCGDan_0472 | 2031 | 548733 | 550754 | BCGDan_sapMKO_0471 | BCGDan_sapMKO_0471 | 2022 |
| 549675 | 551198 | Rv0458 | Rv0458 | 1524 | 580490 | 582013 | BCG_0498 | BCG_0498 | 1524 | 550822 | 552345 | BCGDan_0473 | BCGDan_0473 | 1524 | 550822 | 552345 | BCGDan_sapMKO_0472 | BCGDan_sapMKO_0472 | 1524 |
| 551198 | 551689 | Rv0459 | Rv0459 | 492 | 582013 | 582504 | BCG_0499 | BCG_0499 | 492 | 552345 | 552836 | BCGDan_0474 | BCGDan_0474 | 492 | 552345 | 552836 | BCGDan_sapMKO_0473 | BCGDan_sapMKO_0473 | 492 |
| 551749 | 551988 | Rv0460 | Rv0460 | 240 | 582564 | 582803 | BCG_0500 | BCG_0500 | 240 | 552896 | 553135 | BCGDan_0475 | BCGDan_0475 | 240 | 552857 | 553135 | BCGDan_sapMKO_0474 | BCGDan_sapMKO_0474 | 279 |
| 552026 | 552550 | Rv0461 | Rv0461 | 525 | 582841 | 583365 | BCG_0501 | BCG_0501 | 525 | 553173 | 553697 | BCGDan_0476 | BCGDan_0476 | 525 | 553128 | 553697 | BCGDan_sapMKO_0475 | BCGDan_sapMKO_0475 | 570 |
| 552614 | 554008 | lpdC | Rv0462 | 1395 | 583429 | 584823 | lpd | BCG_0502 | 1395 | 553761 | 555155 | lpd | BCGDan_0477 | 1395 | 553761 | 555155 | lpd | BCGDan_sapMKO_0476 | 1395 |
| 554016 | 554309 | Rv0463 | Rv0463 | 294 | 584831 | 585124 | BCG_0503 | BCG_0503 | 294 | 555163 | 555456 | BCGDan_0478 | BCGDan_0478 | 294 | 555163 | 555456 | BCGDan_sapMKO_0477 | BCGDan_sapMKO_0477 | 294 |
| 554313 | 554885 | Rv0464c | Rv0464c | 573 | 585128 | 585700 | BCG_0504c | BCG_0504c | 573 | 555460 | 556032 | BCGDan_0479 | BCGDan_0479 | 573 | 555460 | 556032 | BCGDan_sapMKO_0478 | BCGDan_sapMKO_0478 | 573 |
| 554882 | 556306 | Rv0465c | Rv0465c | 1425 | 585697 | 587121 | BCG_0505c | BCG_0505c | 1425 | 556029 | 557453 | BCGDan_0480 | BCGDan_0480 | 1425 | 556029 | 557453 | BCGDan_sapMKO_0479 | BCGDan_sapMKO_0479 | 1425 |
| 556458 | 557252 | Rv0466 | Rv0466 | 795 | 587273 | 588067 | BCG_0506 | BCG_0506 | 795 | 557605 | 558399 | BCGDan_0481 | BCGDan_0481 | 795 | 557632 | 558399 | BCGDan_sapMKO_0480 | BCGDan_sapMKO_0480 | 768 |
| 557527 | 558813 | icl1 | Rv0467 | 1287 | 588342 | 589628 | icl | BCG_0507 | 1287 | 558674 | 559960 | icl | BCGDan_0482 | 1287 | 558674 | 559960 | icl | BCGDan_sapMKO_0481 | 1287 |
| 558895 | 559755 | fadB2 | Rv0468 | 861 | 589710 | 590570 | fadB2 | BCG_0508 | 861 | 560042 | 560902 | fadB2 | BCGDan_0483 | 861 | 560087 | 560902 | fadB2 | BCGDan_sapMKO_0482 | 816 |
| 559888 | 560748 | umaA | Rv0469 | 861 | 590703 | 591563 | umaA1 | BCG_0509 | 861 | 561035 | 561895 | umaA1 | BCGDan_0484 | 861 | 561035 | 561895 | umaA1 | BCGDan_sapMKO_0483 | 861 |
| 560848 | 561711 | pcaA | Rv0470c | 864 | 591663 | 592526 | pcaA | BCG_0510c | 864 | 561995 | 562858 | pcaA | BCGDan_0485 | 864 | 561995 | 562858 | pcaA | BCGDan_sapMKO_0484 | 864 |
| 561854 | 562294 | Rv0470A | Rv0470A | 441 | 592669 | 593109 | BCG_0511c | BCG_0511c | 441 | 563001 | 563441 | BCGDan_0486 | BCGDan_0486 | 441 | 563001 | 563441 | BCGDan_sapMKO_0485 | BCGDan_sapMKO_0485 | 441 |
| 562225 | 562713 | Rv0471c | Rv0471c | 489 | 593040 | 593528 | BCG_0512c | BCG_0512c | 489 | 563372 | 563860 | BCGDan_0487 | BCGDan_0487 | 489 | 563372 | 563860 | BCGDan_sapMKO_0486 | BCGDan_sapMKO_0486 | 489 |
| 562723 | 563427 | Rv0472c | Rv0472c | 705 | 593538 | 594242 | BCG_0513c | BCG_0513c | 705 | 563870 | 564574 | BCGDan_0488 | BCGDan_0488 | 705 | 563870 | 564574 | BCGDan_sapMKO_0487 | BCGDan_sapMKO_0487 | 705 |
| 563564 | 564934 | Rv0473 | Rv0473 | 1371 | 594379 | 595749 | BCG_0514 | BCG_0514 | 1371 | 564711 | 566081 | BCGDan_0489 | BCGDan_0489 | 1371 | 564687 | 566081 | BCGDan_sapMKO_0488 | BCGDan_sapMKO_0488 | 1395 |
| 565021 | 565443 | Rv0474 | Rv0474 | 423 | 595836 | 596258 | BCG_0515 | BCG_0515 | 423 | 566168 | 566590 | BCGDan_0490 | BCGDan_0490 | 423 | 566168 | 566590 | BCGDan_sapMKO_0489 | BCGDan_sapMKO_0489 | 423 |
| 565797 | 566396 | hbhA | Rv0475 | 600 | 596612 | 597211 | hbhA | BCG_0516 | 600 | 566944 | 567543 | hbhA | BCGDan_0491 | 600 | 566944 | 567543 | hbhA | BCGDan_sapMKO_0490 | 600 |
| 566508 | 566771 | Rv0476 | Rv0476 | 264 | 597323 | 597586 | BCG_0517 | BCG_0517 | 264 | 567655 | 567918 | BCGDan_0492 | BCGDan_0492 | 264 | 567631 | 567918 | BCGDan_sapMKO_0491 | BCGDan_sapMKO_0491 | 288 |
| 566776 | 567222 | Rv0477 | Rv0477 | 447 | 597591 | 598037 | BCG_0518 | BCG_0518 | 447 | 567923 | 568369 | BCGDan_0493 | BCGDan_0493 | 447 | 567923 | 568369 | BCGDan_sapMKO_0492 | BCGDan_sapMKO_0492 | 447 |
| 567222 | 567896 | deoC | Rv0478 | 675 | 598037 | 598711 | deoC | BCG_0519 | 675 | 568369 | 569043 | deoC | BCGDan_0494 | 675 | 568369 | 569043 | deoC | BCGDan_sapMKO_0493 | 675 |
| 567921 | 568967 | Rv0479c | Rv0479c | 1047 | 598736 | 599782 | BCG_0520c | BCG_0520c | 1047 | 569068 | 570114 | BCGDan_0495 | BCGDan_0495 | 1047 | 569068 | 570114 | BCGDan_sapMKO_0494 | BCGDan_sapMKO_0494 | 1047 |
| 568964 | 569806 | Rv0480c | Rv0480c | 843 | 599779 | 600714 | BCG_0521c | BCG_0521c | 936 | 570111 | 571046 | BCGDan_0496 | BCGDan_0496 | 936 | 570111 | 571046 | BCGDan_sapMKO_0495 | BCGDan_sapMKO_0495 | 936 |
| 569988 | 570512 | Rv0481c | Rv0481c | 525 | 600747 | 601271 | BCG_0522c | BCG_0522c | 525 | 571079 | 571603 | BCGDan_0497 | BCGDan_0497 | 525 | 571079 | 571585 | BCGDan_sapMKO_0496 | BCGDan_sapMKO_0496 | 507 |
| 570539 | 571648 | murB | Rv0482 | 1110 | 601298 | 602407 | murB | BCG_0523 | 1110 | 571630 | 572739 | murB | BCGDan_0498 | 1110 | 571630 | 572739 | murB | BCGDan_sapMKO_0497 | 1110 |
| 571710 | 573065 | lprQ | Rv0483 | 1356 | 602469 | 603824 | lprQ | BCG_0524 | 1356 | 572801 | 574156 | lprQ | BCGDan_0499 | 1356 | 572918 | 574156 | lprQ | BCGDan_sapMKO_0498 | 1239 |
| 573046 | 573801 | Rv0484c | Rv0484c | 756 | 603805 | 604560 | BCG_0525c | BCG_0525c | 756 | 574137 | 574892 | BCGDan_0500 | BCGDan_0500 | 756 | 574137 | 574892 | BCGDan_sapMKO_0499 | BCGDan_sapMKO_0499 | 756 |
| 573984 | 575300 | Rv0485 | Rv0485 | 1317 | 604743 | 606059 | BCG_0526 | BCG_0526 | 1317 | 575075 | 576391 | BCGDan_0501 | BCGDan_0501 | 1317 | 575075 | 576391 | BCGDan_sapMKO_0500 | BCGDan_sapMKO_0500 | 1317 |
| 575348 | 576790 | mshA | Rv0486 | 1443 | 606107 | 607549 | BCG_0527 | BCG_0527 | 1443 | 576439 | 577881 | BCGDan_0502 | BCGDan_0502 | 1443 | 576439 | 577881 | BCGDan_sapMKO_0501 | BCGDan_sapMKO_0501 | 1443 |
| 576787 | 577338 | Rv0487 | Rv0487 | 552 | 607546 | 608097 | BCG_0528 | BCG_0528 | 552 | 577878 | 578429 | BCGDan_0503 | BCGDan_0503 | 552 | 577899 | 578429 | BCGDan_sapMKO_0502 | BCGDan_sapMKO_0502 | 531 |
| 577664 | 578269 | Rv0488 | Rv0488 | 606 | 608539 | 609144 | BCG_0529 | BCG_0529 | 606 | 578813 | 579418 | BCGDan_0504 | BCGDan_0504 | 606 | 578896 | 579534 | BCGDan_sapMKO_0503 | BCGDan_sapMKO_0503 | 639 |
| 578426 | 579175 | gpm1 | Rv0489 | 750 | 609301 | 610050 | gpm1 | BCG_0530 | 750 | 579575 | 580324 | gpm1 | BCGDan_0505 | 750 | 579691 | 580440 | gpm1 | BCGDan_sapMKO_0504 | 750 |
| 579349 | 580581 | senX3 | Rv0490 | 1233 | 610224 | 611456 | senX3 | BCG_0531 | 1233 | 580498 | 581730 | senX3 | BCGDan_0506 | 1233 | 580614 | 581846 | senX3 | BCGDan_sapMKO_0505 | 1233 |
| 580809 | 581492 | regX3 | Rv0491 | 684 | 611607 | 612290 | regX3 | BCG_0532 | 684 | 581958 | 582641 | regX3 | BCGDan_0507 | 684 | 582065 | 582757 | regX3 | BCGDan_sapMKO_0506 | 693 |
| 581489 | 583378 | Rv0492c | Rv0492c | 1890 | 612287 | 614176 | BCG_0533c | BCG_0533c | 1890 | 582638 | 584527 | BCGDan_0508 | BCGDan_0508 | 1890 | 582754 | 584643 | BCGDan_sapMKO_0507 | BCGDan_sapMKO_0507 | 1890 |
| 583375 | 583704 | Rv0492A | Rv0492A | 330 | 614173 | 614502 | BCG_0534c | BCG_0534c | 330 | 584524 | 584853 | BCGDan_0509 | BCGDan_0509 | 330 | 584640 | 584969 | BCGDan_sapMKO_0508 | BCGDan_sapMKO_0508 | 330 |
| 583701 | 584690 | Rv0493c | Rv0493c | 990 | 614499 | 615488 | BCG_0535c | BCG_0535c | 990 | 584850 | 585839 | BCGDan_0510 | BCGDan_0510 | 990 | 584966 | 585916 | BCGDan_sapMKO_0509 | BCGDan_sapMKO_0509 | 951 |
| 584695 | 585423 | Rv0494 | Rv0494 | 729 | 615493 | 616221 | BCG_0536 | BCG_0536 | 729 | 585844 | 586572 | BCGDan_0511 | BCGDan_0511 | 729 | 585972 | 586688 | BCGDan_sapMKO_0510 | BCGDan_sapMKO_0510 | 717 |
| 585424 | 586314 | Rv0495c | Rv0495c | 891 | 616222 | 617112 | BCG_0537c | BCG_0537c | 891 | 586573 | 587463 | BCGDan_0512 | BCGDan_0512 | 891 | 586689 | 587459 | BCGDan_sapMKO_0511 | BCGDan_sapMKO_0511 | 771 |
| 586394 | 587380 | Rv0496 | Rv0496 | 987 | 617192 | 618178 | BCG_0538 | BCG_0538 | 987 | 587543 | 588529 | BCGDan_0513 | BCGDan_0513 | 987 | 587611 | 588645 | BCGDan_sapMKO_0512 | BCGDan_sapMKO_0512 | 1035 |
| 587377 | 588309 | Rv0497 | Rv0497 | 933 | 618175 | 619107 | BCG_0539 | BCG_0539 | 933 | 588526 | 589458 | BCGDan_0514 | BCGDan_0514 | 933 | 588642 | 589574 | BCGDan_sapMKO_0513 | BCGDan_sapMKO_0513 | 933 |
| 588325 | 589167 | Rv0498 | Rv0498 | 843 | 619123 | 619965 | BCG_0540 | BCG_0540 | 843 | 589474 | 590316 | BCGDan_0515 | BCGDan_0515 | 843 | 589590 | 590432 | BCGDan_sapMKO_0514 | BCGDan_sapMKO_0514 | 843 |

| | | | | | | | | | | | | | | | | | | | |
|---|---|---|---|---|---|---|---|---|---|---|---|---|---|---|---|---|---|---|---|
| 589183 | 590058 | Rv0499 | Rv0499 | 876 | 619981 | 620856 | BCG_0541 | BCG_0541 | 876 | 590332 | 591207 | BCGDan_0516 | BCGDan_0516 | 876 | 590448 | 591323 | BCGDan_sapMKO_0515 | BCGDan_sapMKO_0515 | 876 |
| 590083 | 590970 | proC | Rv0500 | 888 | 620881 | 621768 | proC | BCG_0542 | 888 | 591232 | 592119 | proC | BCGDan_0517 | 888 | 591348 | 592235 | proC | BCGDan_sapMKO_0516 | 888 |
| 591111 | 591347 | Rv0500A | Rv0500A | 237 | 621909 | 622145 | BCG_0543 | BCG_0543 | 237 | 592260 | 592496 | BCGDan_0518 | BCGDan_0518 | 237 | 592376 | 592612 | BCGDan_sapMKO_0517 | BCGDan_sapMKO_0517 | 237 |
| 591654 | 592784 | galE2 | Rv0501 | 1131 | 622452 | 623582 | galE2 | BCG_0544 | 1131 | 592803 | 593771 | galE2 | BCGDan_0519 | 969 | 592919 | 594049 | galE2 | BCGDan_sapMKO_0518 | 1131 |
| 592791 | 593867 | Rv0502 | Rv0502 | 1077 | 623589 | 624665 | BCG_0545 | BCG_0545 | 1077 | 593940 | 595016 | BCGDan_0520 | BCGDan_0520 | 1077 | 594173 | 595132 | BCGDan_sapMKO_0519 | BCGDan_sapMKO_0519 | 960 |
| 593871 | 594779 | cmaA2 | Rv0503c | 909 | 624669 | 625577 | cmaA2 | BCG_0546c | 909 | 595020 | 595928 | cmaA2 | BCGDan_0521 | 909 | 595136 | 596044 | cmaA2 | BCGDan_sapMKO_0520 | 909 |
| 594802 | 595302 | Rv0504c | Rv0504c | 501 | 625600 | 626100 | BCG_0547c | BCG_0547c | 501 | 595951 | 596451 | BCGDan_0522 | BCGDan_0522 | 501 | 596067 | 596567 | BCGDan_sapMKO_0521 | BCGDan_sapMKO_0521 | 501 |
| 595464 | 596585 | serB1 | Rv0505c | 1122 | 626262 | 627383 | serB1 | BCG_0548c | 1122 | 596613 | 597734 | serB1 | BCGDan_0523 | 1122 | 596729 | 597850 | serB1 | BCGDan_sapMKO_0522 | 1122 |
| 596759 | 597202 | mmpS2 | Rv0506 | 444 | 627557 | 628000 | mmpS2 | BCG_0549 | 444 | 597908 | 598351 | mmpS2 | BCGDan_0524 | 444 | 598060 | 598467 | mmpS2 | BCGDan_sapMKO_0523 | 408 |
| 597199 | 600105 | mmpL2 | Rv0507 | 2907 | 627997 | 630903 | mmpL2 | BCG_0550 | 2907 | 598348 | 601254 | mmpL2 | BCGDan_0525 | 2907 | 598464 | 601370 | mmpL2 | BCGDan_sapMKO_0524 | 2907 |
| 600098 | 600391 | Rv0508 | Rv0508 | 294 | 630896 | 631189 | BCG_0551 | BCG_0551 | 294 | 601247 | 601540 | BCGDan_0526 | BCGDan_0526 | 294 | 601363 | 601656 | BCGDan_sapMKO_0525 | BCGDan_sapMKO_0525 | 294 |
| 600441 | 601847 | hemA | Rv0509 | 1407 | 631239 | 632645 | hemA | BCG_0552 | 1407 | 601590 | 602996 | hemA | BCGDan_0527 | 1407 | 601706 | 603112 | hemA | BCGDan_sapMKO_0526 | 1407 |
| 601857 | 602786 | hemC | Rv0510 | 930 | 632655 | 633584 | hemC | BCG_0553 | 930 | 603006 | 603935 | hemC | BCGDan_0528 | 930 | 603122 | 604051 | hemC | BCGDan_sapMKO_0527 | 930 |
| 602819 | 604516 | hemD | Rv0511 | 1698 | 633617 | 635314 | hemD | BCG_0554 | 1698 | 603968 | 605665 | hemD | BCGDan_0529 | 1698 | 604126 | 605781 | hemD | BCGDan_sapMKO_0528 | 1656 |
| 604602 | 605591 | hemB | Rv0512 | 990 | 635400 | 636389 | hemB | BCG_0555 | 990 | 605751 | 606740 | hemB | BCGDan_0530 | 990 | 605924 | 606856 | hemB | BCGDan_sapMKO_0529 | 933 |
| 605604 | 606152 | Rv0513 | Rv0513 | 549 | 636402 | 636950 | BCG_0556 | BCG_0556 | 549 | 606753 | 607301 | BCGDan_0531 | BCGDan_0531 | 549 | 606869 | 607417 | BCGDan_sapMKO_0530 | BCGDan_sapMKO_0530 | 549 |
| 606149 | 606448 | Rv0514 | Rv0514 | 300 | 636947 | 637246 | BCG_0557 | BCG_0557 | 300 | 607298 | 607597 | BCGDan_0532 | BCGDan_0532 | 300 | 607414 | 607713 | BCGDan_sapMKO_0531 | BCGDan_sapMKO_0531 | 300 |
| 606551 | 608062 | Rv0515 | Rv0515 | 1512 | 637349 | 638860 | BCG_0558 | BCG_0558 | 1512 | 607700 | 609211 | BCGDan_0533 | BCGDan_0533 | 1512 | 607792 | 609327 | BCGDan_sapMKO_0532 | BCGDan_sapMKO_0532 | 1536 |
| 608059 | 608535 | Rv0516c | Rv0516c | 477 | 638857 | 639333 | BCG_0559c | BCG_0559c | 477 | 609208 | 609684 | BCGDan_0534 | BCGDan_0534 | 477 | 609324 | 609800 | BCGDan_sapMKO_0533 | BCGDan_sapMKO_0533 | 477 |
| 608746 | 610056 | Rv0517 | Rv0517 | 1311 | 639544 | 640854 | BCG_0560 | BCG_0560 | 1311 | 609895 | 611205 | BCGDan_0535 | BCGDan_0535 | 1311 | 610089 | 611321 | BCGDan_sapMKO_0534 | BCGDan_sapMKO_0534 | 1233 |
| 610188 | 610883 | Rv0518 | Rv0518 | 696 | 640986 | 641681 | BCG_0561 | BCG_0561 | 696 | 611337 | 612032 | BCGDan_0536 | BCGDan_0536 | 696 | 611453 | 612148 | BCGDan_sapMKO_0535 | BCGDan_sapMKO_0535 | 696 |
| 611172 | 612074 | Rv0519c | Rv0519c | 903 | 641970 | 642872 | BCG_0562c | BCG_0562c | 903 | 612321 | 613223 | BCGDan_0537 | BCGDan_0537 | 903 | 612437 | 613339 | BCGDan_sapMKO_0536 | BCGDan_sapMKO_0536 | 903 |
| 612255 | 612605 | Rv0520 | Rv0520 | 351 | 643053 | 643403 | BCG_0563 | BCG_0563 | 351 | 613404 | 613754 | BCGDan_0538 | BCGDan_0538 | 351 | 613520 | 613870 | BCGDan_sapMKO_0537 | BCGDan_sapMKO_0537 | 351 |
| 612598 | 612903 | Rv0521 | Rv0521 | 306 | 643396 | 643701 | BCG_0564 | BCG_0564 | 306 | 613747 | 614052 | BCGDan_0539 | BCGDan_0539 | 306 | 613863 | 614168 | BCGDan_sapMKO_0538 | BCGDan_sapMKO_0538 | 306 |
| 613038 | 614342 | gabP | Rv0522 | 1305 | 643836 | 645140 | gabP | BCG_0565 | 1305 | 614187 | 615491 | gabP | BCGDan_0540 | 1305 | 614303 | 615607 | gabP | BCGDan_sapMKO_0539 | 1305 |
| 614326 | 614721 | Rv0523c | Rv0523c | 396 | 645124 | 645519 | BCG_0566c | BCG_0566c | 396 | 615475 | 615870 | BCGDan_0541 | BCGDan_0541 | 396 | 615591 | 615995 | BCGDan_sapMKO_0540 | BCGDan_sapMKO_0540 | 405 |
| 614835 | 616223 | hemL | Rv0524 | 1389 | 645633 | 647021 | hemL | BCG_0567 | 1389 | 615984 | 617372 | hemL | BCGDan_0542 | 1389 | 616100 | 617488 | hemL | BCGDan_sapMKO_0541 | 1389 |
| 616223 | 616831 | Rv0525 | Rv0525 | 609 | 647021 | 647629 | BCG_0568 | BCG_0568 | 609 | 617372 | 617980 | BCGDan_0543 | BCGDan_0543 | 609 | 617488 | 618096 | BCGDan_sapMKO_0542 | BCGDan_sapMKO_0542 | 609 |
| 616846 | 617496 | Rv0526 | Rv0526 | 651 | 647644 | 648294 | BCG_0569 | BCG_0569 | 651 | 617995 | 618645 | BCGDan_0544 | BCGDan_0544 | 651 | 618150 | 618761 | BCGDan_sapMKO_0543 | BCGDan_sapMKO_0543 | 612 |
| 617493 | 618272 | ccdA | Rv0527 | 780 | 648291 | 649070 | ccdA | BCG_0570 | 780 | 618642 | 619421 | ccdA | BCGDan_0545 | 780 | 618758 | 619537 | ccdA | BCGDan_sapMKO_0544 | 780 |
| 618305 | 619894 | Rv0528 | Rv0528 | 1590 | 649103 | 650692 | BCG_0571 | BCG_0571 | 1590 | 619454 | 621043 | BCGDan_0546 | BCGDan_0546 | 1590 | 619591 | 621159 | BCGDan_sapMKO_0545 | BCGDan_sapMKO_0545 | 1569 |
| 619891 | 620865 | ccsA | Rv0529 | 975 | 650689 | 651663 | ccsA | BCG_0572 | 975 | 621040 | 622014 | ccsA | BCGDan_0547 | 975 | 621156 | 622130 | ccsA | BCGDan_sapMKO_0546 | 975 |
| 620907 | 622124 | Rv0530 | Rv0530 | 1218 | 651705 | 652922 | BCG_0573 | BCG_0573 | 1218 | 622056 | 623273 | BCGDan_0548 | BCGDan_0548 | 1218 | 622172 | 623389 | BCGDan_sapMKO_0547 | BCGDan_sapMKO_0547 | 1218 |
| 622329 | 622646 | Rv0531 | Rv0531 | 318 | 653127 | 653444 | BCG_0574 | BCG_0574 | 318 | 623478 | 623795 | BCGDan_0549 | BCGDan_0549 | 318 | 623594 | 623911 | BCGDan_sapMKO_0548 | BCGDan_sapMKO_0548 | 318 |
| 622793 | 624577 | PE_PGRS6 | Rv0532 | 1785 | 653591 | 655336 | PE_PGRS6a | BCG_0575 | 1746 | 623942 | 625678 | PE_PGRS6a | BCGDan_0550 | 1737 | 624058 | 625794 | PE_PGRS6a | BCGDan_sapMKO_0549 | 1737 |
| NA | NA | NA | NA | NA | 655242 | 655469 | PE_PGRS6b | BCG_0576 | 228 | 625584 | 625811 | PE_PGRS6b | BCGDan_0551 | 228 | NA | NA | NA | NA | NA |
| 624473 | 625480 | fabH | Rv0533c | 1008 | 655365 | 656372 | fabH | BCG_0577c | 1008 | 625707 | 626714 | fabH | BCGDan_0552 | 1008 | 625823 | 626830 | fabH | BCGDan_sapMKO_0550 | 1008 |
| 625562 | 626440 | menA | Rv0534c | 879 | 656454 | 657332 | menA | BCG_0578c | 879 | 626796 | 627674 | menA | BCGDan_0553 | 879 | 626912 | 627790 | menA | BCGDan_sapMKO_0551 | 879 |
| 626457 | 627251 | pnp | Rv0535 | 795 | 657349 | 658143 | pnp | BCG_0579 | 795 | 627691 | 628485 | pnp | BCGDan_0554 | 795 | 627807 | 628601 | pnp | BCGDan_sapMKO_0552 | 795 |
| 627248 | 628288 | galE3 | Rv0536 | 1041 | 658140 | 659180 | galE3 | BCG_0580 | 1041 | 628482 | 629522 | galE3 | BCGDan_0555 | 1041 | 628604 | 629638 | galE3 | BCGDan_sapMKO_0553 | 1035 |
| 628298 | 629731 | Rv0537c | Rv0537c | 1434 | 659190 | 660623 | BCG_0581c | BCG_0581c | 1434 | 629532 | 630965 | BCGDan_0556 | BCGDan_0556 | 1434 | 629648 | 631081 | BCGDan_sapMKO_0554 | BCGDan_sapMKO_0554 | 1434 |
| 630040 | 631686 | Rv0538 | Rv0538 | 1647 | 660932 | 662578 | BCG_0582 | BCG_0582 | 1647 | 631274 | 632920 | BCGDan_0557 | BCGDan_0557 | 1647 | 631390 | 633036 | BCGDan_sapMKO_0555 | BCGDan_sapMKO_0555 | 1647 |
| 631743 | 632375 | Rv0539 | Rv0539 | 633 | 662635 | 663267 | BCG_0583 | BCG_0583 | 633 | 632977 | 633609 | BCGDan_0558 | BCGDan_0558 | 633 | 633069 | 633725 | BCGDan_sapMKO_0556 | BCGDan_sapMKO_0556 | 657 |
| 632372 | 633034 | Rv0540 | Rv0540 | 663 | 663264 | 663926 | BCG_0584 | BCG_0584 | 663 | 633606 | 634268 | BCGDan_0559 | BCGDan_0559 | 663 | 633722 | 634384 | BCGDan_sapMKO_0557 | BCGDan_sapMKO_0557 | 663 |
| 633055 | 634404 | Rv0541c | Rv0541c | 1350 | 663947 | 665296 | BCG_0585c | BCG_0585c | 1350 | 634289 | 635638 | BCGDan_0560 | BCGDan_0560 | 1350 | 634405 | 635754 | BCGDan_sapMKO_0558 | BCGDan_sapMKO_0558 | 1350 |
| 634416 | 635504 | menE | Rv0542c | 1089 | 665308 | 666396 | menE | BCG_0586c | 1089 | 635650 | 636738 | menE | BCGDan_0561 | 1089 | 635766 | 636854 | menE | BCGDan_sapMKO_0559 | 1089 |
| 635573 | 635875 | Rv0543c | Rv0543c | 303 | 666465 | 666767 | BCG_0587c | BCG_0587c | 303 | 636807 | 637109 | BCGDan_0562 | BCGDan_0562 | 303 | 636923 | 637225 | BCGDan_sapMKO_0560 | BCGDan_sapMKO_0560 | 303 |
| 635935 | 636213 | Rv0544c | Rv0544c | 279 | 666827 | 667105 | BCG_0588c | BCG_0588c | 279 | 637169 | 637447 | BCGDan_0563 | BCGDan_0563 | 279 | 637285 | 637563 | BCGDan_sapMKO_0561 | BCGDan_sapMKO_0561 | 279 |
| 636210 | 637463 | pitA | Rv0545c | 1254 | 667102 | 668355 | pitA | BCG_0589c | 1254 | 637444 | 638697 | pitA | BCGDan_0564 | 1254 | 637560 | 638813 | pitA | BCGDan_sapMKO_0562 | 1254 |
| 637583 | 637969 | Rv0546c | Rv0546c | 387 | 668475 | 668861 | BCG_0590c | BCG_0590c | 387 | 638817 | 639203 | BCGDan_0565 | BCGDan_0565 | 387 | 638933 | 639319 | BCGDan_sapMKO_0563 | BCGDan_sapMKO_0563 | 387 |
| 638032 | 638916 | Rv0547c | Rv0547c | 885 | 668924 | 669808 | BCG_0591c | BCG_0591c | 885 | 639266 | 640150 | BCGDan_0566 | BCGDan_0566 | 885 | 639382 | 640266 | BCGDan_sapMKO_0564 | BCGDan_sapMKO_0564 | 885 |
| 639012 | 639956 | menB | Rv0548c | 945 | 669904 | 670848 | menB | BCG_0592c | 945 | 640246 | 641190 | menB | BCGDan_0567 | 945 | 640362 | 641225 | menB | BCGDan_sapMKO_0565 | 864 |
| 640228 | 640641 | vapC3 | Rv0549c | 414 | 671120 | 671257 | BCG_0593c | BCG_0593c | 138 | 641462 | 641599 | BCGDan_0568 | BCGDan_0568 | 138 | 641581 | 641715 | BCGDan_sapMKO_0566 | BCGDan_sapMKO_0566 | 135 |
| NA | NA | NA | NA | NA | 671224 | 671532 | BCG_0594c | BCG_0594c | 309 | 641566 | 641874 | BCGDan_0569 | BCGDan_0569 | 309 | 641682 | 641990 | BCGDan_sapMKO_0567 | BCGDan_sapMKO_0567 | 309 |
| 640638 | 640904 | vapB3 | Rv0550c | 267 | 671529 | 671795 | BCG_0595c | BCG_0595c | 267 | 641871 | 642137 | BCGDan_0570 | BCGDan_0570 | 267 | 641987 | 642253 | BCGDan_sapMKO_0568 | BCGDan_sapMKO_0568 | 267 |
| 641096 | 642811 | fadD8 | Rv0551c | 1716 | 671987 | 673702 | fadD8 | BCG_0596c | 1716 | 642329 | 644044 | fadD8 | BCGDan_0571 | 1716 | 642445 | 644061 | fadD8 | BCGDan_sapMKO_0569 | 1617 |
| 642889 | 644493 | Rv0552 | Rv0552 | 1605 | 673780 | 675384 | BCG_0597 | BCG_0597 | 1605 | 644122 | 645726 | BCGDan_0572 | BCGDan_0572 | 1605 | 644238 | 645842 | BCGDan_sapMKO_0570 | BCGDan_sapMKO_0570 | 1605 |
| 644490 | 645470 | menC | Rv0553 | 981 | 675381 | 676361 | menC | BCG_0598 | 981 | 645723 | 646703 | menC | BCGDan_0573 | 981 | 645839 | 646819 | menC | BCGDan_sapMKO_0571 | 981 |
| 645467 | 646255 | bpoC | Rv0554 | 789 | 676358 | 677146 | bpoC | BCG_0599 | 789 | 646700 | 647488 | bpoC | BCGDan_0574 | 789 | 646816 | 647604 | bpoC | BCGDan_sapMKO_0572 | 789 |
| 646298 | 647962 | menD | Rv0555 | 1665 | 677189 | 678853 | menD | BCG_0600 | 1665 | 647531 | 649195 | menD | BCGDan_0575 | 1665 | 647647 | 649311 | menD | BCGDan_sapMKO_0573 | 1665 |
| 647959 | 648474 | Rv0556 | Rv0556 | 516 | 678850 | 679365 | BCG_0601 | BCG_0601 | 516 | 649192 | 649707 | BCGDan_0576 | BCGDan_0576 | 516 | 649308 | 649823 | BCGDan_sapMKO_0574 | BCGDan_sapMKO_0574 | 516 |
| 648536 | 649672 | mgtA | Rv0557 | 1137 | 679427 | 680563 | pimB | BCG_0602 | 1137 | 649769 | 650905 | pimB | BCGDan_0577 | 1137 | 649894 | 651021 | pimB | BCGDan_sapMKO_0575 | 1128 |
| 649689 | 650393 | menH | Rv0558 | 705 | 680580 | 681284 | menH | BCG_0603 | 705 | 650922 | 651626 | menH | BCGDan_0578 | 705 | 651038 | 651742 | menH | BCGDan_sapMKO_0576 | 705 |
| 650407 | 650745 | Rv0559c | Rv0559c | 339 | 681298 | 681636 | BCG_0604c | BCG_0604c | 339 | 651640 | 651978 | BCGDan_0579 | BCGDan_0579 | 339 | 651756 | 652094 | BCGDan_sapMKO_0577 | BCGDan_sapMKO_0577 | 339 |
| 650779 | 651504 | Rv0560c | Rv0560c | 726 | 681670 | 682395 | BCG_0605c | BCG_0605c | 726 | 652012 | 652737 | BCGDan_0580 | BCGDan_0580 | 726 | 652128 | 652805 | BCGDan_sapMKO_0578 | BCGDan_sapMKO_0578 | 678 |
| 651529 | 652755 | Rv0561c | Rv0561c | 1227 | 682420 | 683646 | BCG_0606c | BCG_0606c | 1227 | 652762 | 653988 | BCGDan_0581 | BCGDan_0581 | 1227 | 652878 | 654212 | BCGDan_sapMKO_0579 | BCGDan_sapMKO_0579 | 1335 |
| 652771 | 653778 | grcC1 | Rv0562 | 1008 | 683662 | 684669 | grcC1 | BCG_0607 | 1008 | 654004 | 655011 | grcC1 | BCGDan_0582 | 1008 | 654138 | 655127 | grcC1 | BCGDan_sapMKO_0580 | 990 |
| 653879 | 654739 | htpX | Rv0563 | 861 | 684770 | 685630 | htpX | BCG_0608 | 861 | 655112 | 655972 | htpX | BCGDan_0583 | 861 | 655228 | 656088 | htpX | BCGDan_sapMKO_0581 | 861 |
| 654924 | 655949 | gpdA1 | Rv0564c | 1026 | 685815 | 686840 | gpdA1 | BCG_0609c | 1026 | 656157 | 657182 | gpdA1 | BCGDan_0584 | 1026 | 656273 | 657358 | gpdA1 | BCGDan_sapMKO_0582 | 1086 |
| 656010 | 657470 | Rv0565c | Rv0565c | 1461 | 686901 | 688361 | BCG_0610c | BCG_0610c | 1461 | 657243 | 658703 | BCGDan_0585 | BCGDan_0585 | 1461 | 657359 | 658819 | BCGDan_sapMKO_0583 | BCGDan_sapMKO_0583 | 1461 |
| 657548 | 658039 | Rv0566c | Rv0566c | 492 | 688439 | 688930 | BCG_0611c | BCG_0611c | 492 | 658781 | 659272 | BCGDan_0586 | BCGDan_0586 | 492 | 658897 | 659388 | BCGDan_sapMKO_0584 | BCGDan_sapMKO_0584 | 492 |
| 658321 | 659340 | Rv0567 | Rv0567 | 1020 | 689212 | 690231 | BCG_0612 | BCG_0612 | 1020 | 659554 | 660573 | BCGDan_0588 | BCGDan_0588 | 1020 | 659667 | 660689 | BCGDan_sapMKO_0586 | BCGDan_sapMKO_0586 | 1023 |
| 659450 | 660868 | cyp135B1 | Rv0568 | 1419 | 690341 | 691759 | cyp135B1 | BCG_0613 | 1419 | 660683 | 662101 | cyp135B1 | BCGDan_0589 | 1419 | 660853 | 662217 | cyp135B1 | BCGDan_sapMKO_0587 | 1365 |
| 661003 | 661269 | Rv0569 | Rv0569 | 267 | 691894 | 692160 | BCG_0614 | BCG_0614 | 267 | 662236 | 662502 | BCGDan_0590 | BCGDan_0590 | 267 | 662352 | 662618 | BCGDan_sapMKO_0588 | BCGDan_sapMKO_0588 | 267 |
| 661295 | 663373 | nrdZ | Rv0570 | 2079 | 692186 | 694318 | nrdZ | BCG_0615 | 2133 | 662528 | 664660 | nrdZ | BCGDan_0591 | 2133 | 662644 | 664776 | nrdZ | BCGDan_sapMKO_0589 | 2133 |
| 663487 | 664818 | Rv0571c | Rv0571c | 1332 | 694377 | 695708 | BCG_0616c | BCG_0616c | 1332 | 664719 | 666050 | BCGDan_0592 | BCGDan_0592 | 1332 | 664835 | 666166 | BCGDan_sapMKO_0590 | BCGDan_sapMKO_0590 | 1332 |
| 665042 | 665383 | Rv0572c | Rv0572c | 342 | 695932 | 696273 | BCG_0617c | BCG_0617c | 342 | 666274 | 666615 | BCGDan_0593 | BCGDan_0593 | 342 | 666390 | 666731 | BCGDan_sapMKO_0591 | BCGDan_sapMKO_0591 | 342 |
| 665851 | 667242 | pncB2 | Rv0573c | 1392 | 696741 | 698132 | BCG_0618c | BCG_0618c | 1392 | 667083 | 668474 | BCGDan_0594 | BCGDan_0594 | 1392 | 667199 | 668590 | BCGDan_sapMKO_0592 | BCGDan_sapMKO_0592 | 1392 |
| 667252 | 668394 | Rv0574c | Rv0574c | 1143 | 698142 | 699284 | BCG_0619c | BCG_0619c | 1143 | 668484 | 669626 | BCGDan_0595 | BCGDan_0595 | 1143 | 668600 | 669742 | BCGDan_sapMKO_0593 | BCGDan_sapMKO_0593 | 1143 |
| 668579 | 669745 | Rv0575c | Rv0575c | 1167 | 699500 | 700636 | BCG_0620c | BCG_0620c | 1137 | 669842 | 670978 | BCGDan_0596 | BCGDan_0596 | 1137 | 669958 | 671094 | BCGDan_sapMKO_0594 | BCGDan_sapMKO_0594 | 1137 |
| 669848 | 671152 | Rv0576 | Rv0576 | 1305 | 700739 | 702043 | BCG_0621 | BCG_0621 | 1305 | 671081 | 672385 | BCGDan_0597 | BCGDan_0597 | 1305 | 671188 | 672501 | BCGDan_sapMKO_0595 | BCGDan_sapMKO_0595 | 1314 |
| 671166 | 671951 | TB27.3 | Rv0577 | 786 | 702057 | 702842 | TB27.3 | BCG_0622 | 786 | 672399 | 673184 | TB27.3 | BCGDan_0598 | 786 | 672509 | 673300 | TB27.3 | BCGDan_sapMKO_0596 | 792 |
| 671996 | 675916 | PE_PGRS7 | Rv0578c | 3921 | 702887 | 706798 | PE_PGRS7 | BCG_0623c | 3912 | 673229 | 677140 | PE_PGRS7 | BCGDan_0599 | 3912 | 673345 | 677256 | PE_PGRS7 | BCGDan_sapMKO_0597 | 3912 |
| 676238 | 676996 | Rv0579 | Rv0579 | 759 | 707120 | 707878 | BCG_0624 | BCG_0624 | 759 | 677462 | 678220 | BCGDan_0600 | BCGDan_0600 | 759 | 677578 | 678336 | BCGDan_sapMKO_0598 | BCGDan_sapMKO_0598 | 759 |
| 677125 | 677616 | Rv0580c | Rv0580c | 492 | 708007 | 708498 | BCG_0625c | BCG_0625c | 492 | 678349 | 678840 | BCGDan_0601 | BCGDan_0601 | 492 | 678465 | 678956 | BCGDan_sapMKO_0599 | BCGDan_sapMKO_0599 | 492 |
| 677710 | 677925 | vapB26 | Rv0581 | 216 | 708592 | 708807 | BCG_0626 | BCG_0626 | 216 | 678934 | 679149 | BCGDan_0602 | BCGDan_0602 | 216 | 679050 | 679265 | BCGDan_sapMKO_0600 | BCGDan_sapMKO_0600 | 216 |

| | | | | | | | | | | | | | | | | | | | |
|---|---|---|---|---|---|---|---|---|---|---|---|---|---|---|---|---|---|---|---|
| 677922 | 678329 | vapC26 | Rv0582 | 408 | 708804 | 709211 | BCG_0627 | BCG_0627 | 408 | 679146 | 679553 | BCGDan_0603 | BCGDan_0603 | 408 | 679262 | 679669 | BCGDan_sapMKO_0601 | BCGDan_sapMKO_0601 | 408 |
| 678389 | 679075 | lpqN | Rv0583c | 687 | 709271 | 709957 | lpqN | BCG_0628c | 687 | 679613 | 680299 | lpqN | BCGDan_0604 | 687 | 679729 | 680415 | lpqN | BCGDan_sapMKO_0602 | 687 |
| 679229 | 681862 | Rv0584 | Rv0584 | 2634 | 710111 | 712744 | BCG_0629 | BCG_0629 | 2634 | 680453 | 683086 | BCGDan_0605 | BCGDan_0605 | 2634 | 680569 | 683202 | BCGDan_sapMKO_0603 | BCGDan_sapMKO_0603 | 2634 |
| 681885 | 684272 | Rv0585c | Rv0585c | 2388 | 712767 | 715154 | BCG_0630c | BCG_0630c | 2388 | 683109 | 685496 | BCGDan_0606 | BCGDan_0606 | 2388 | 683225 | 685612 | BCGDan_sapMKO_0604 | BCGDan_sapMKO_0604 | 2388 |
| 684410 | 685132 | mce2R | Rv0586 | 723 | 715292 | 716014 | BCG_0631 | BCG_0631 | 723 | 685634 | 686356 | BCGDan_0607 | BCGDan_0607 | 723 | 685750 | 686472 | BCGDan_sapMKO_0605 | BCGDan_sapMKO_0605 | 723 |
| 685129 | 685926 | yrbE2A | Rv0587 | 798 | 716011 | 716808 | yrbE2A | BCG_0632 | 798 | 686353 | 687150 | yrbE2A | BCGDan_0608 | 798 | 686469 | 687266 | yrbE2A | BCGDan_sapMKO_0606 | 798 |
| 685928 | 686815 | yrbE2B | Rv0588 | 888 | 716810 | 717697 | yrbE2B | BCG_0633 | 888 | 687152 | 688039 | yrbE2B | BCGDan_0609 | 888 | 687268 | 688155 | yrbE2B | BCGDan_sapMKO_0607 | 888 |
| 686821 | 688035 | mce2A | Rv0589 | 1215 | 717703 | 718917 | mce2A | BCG_0634 | 1215 | 688045 | 689259 | mce2A | BCGDan_0610 | 1215 | 688161 | 689375 | mce2A | BCGDan_sapMKO_0608 | 1215 |
| 688032 | 688859 | mce2B | Rv0590 | 828 | 718914 | 719945 | mce2B | BCG_0635 | 1032 | 689256 | 690287 | mce2B | BCGDan_0611 | 1032 | 689372 | 690403 | mce2B | BCGDan_sapMKO_0609 | 1032 |
| 689059 | 690504 | mce2C | Rv0591 | 1446 | 719942 | 721387 | mce2C | BCG_0636 | 1446 | 690284 | 691729 | mce2C | BCGDan_0612 | 1446 | 690400 | 691845 | mce2C | BCGDan_sapMKO_0610 | 1446 |
| 690501 | 692027 | mce2D | Rv0592 | 1527 | 721384 | 722934 | mce2Da | BCG_0637 | 1551 | 691726 | 693276 | mce2Da | BCGDan_0613 | 1551 | 691842 | 693392 | mce2Da | BCGDan_sapMKO_0611 | 1551 |
| 692024 | 693232 | lprL | Rv0593 | 1209 | 722909 | 724117 | lprL | BCG_0639 | 1209 | 693251 | 694459 | lprL | BCGDan_0614 | 1209 | 693433 | 694575 | lprL | BCGDan_sapMKO_0612 | 1143 |
| 693237 | 694787 | mce2F | Rv0594 | 1551 | 724122 | 725672 | mce2F | BCG_0640 | 1551 | 694464 | 696014 | mce2F | BCGDan_0615 | 1551 | 694535 | 696130 | mce2F | BCGDan_sapMKO_0613 | 1596 |
| 694839 | 695231 | vapC4 | Rv0595c | 393 | 725724 | 726116 | BCG_0641c | BCG_0641c | 393 | 696066 | 696458 | BCGDan_0616 | BCGDan_0616 | 393 | 696182 | 696577 | BCGDan_sapMKO_0614 | BCGDan_sapMKO_0614 | 396 |
| 695228 | 695485 | vapB4 | Rv0596c | 258 | 726113 | 726370 | BCG_0642c | BCG_0642c | 258 | 696455 | 696712 | BCGDan_0617 | BCGDan_0617 | 258 | 696571 | 696828 | BCGDan_sapMKO_0615 | BCGDan_sapMKO_0615 | 258 |
| 695668 | 696903 | Rv0597c | Rv0597c | 1236 | 726553 | 727788 | BCG_0643c | BCG_0643c | 1236 | 696895 | 698130 | BCGDan_0618 | BCGDan_0618 | 1236 | 697011 | 698246 | BCGDan_sapMKO_0616 | BCGDan_sapMKO_0616 | 1236 |
| 697154 | 697567 | vapC27 | Rv0598c | 414 | 728039 | 728452 | BCG_0644c | BCG_0644c | 414 | 698381 | 698794 | BCGDan_0619 | BCGDan_0619 | 414 | 698497 | 698910 | BCGDan_sapMKO_0617 | BCGDan_sapMKO_0617 | 414 |
| 697564 | 697800 | vapB27 | Rv0599c | 237 | 728449 | 728685 | BCG_0645c | BCG_0645c | 237 | 698791 | 699027 | BCGDan_0620 | BCGDan_0620 | 237 | 698907 | 699143 | BCGDan_sapMKO_0618 | BCGDan_sapMKO_0618 | 237 |
| 697904 | 698410 | Rv0600c | Rv0600c | 507 | 728789 | 729295 | BCG_0646c | BCG_0646c | 507 | 699131 | 699637 | BCGDan_0621 | BCGDan_0621 | 507 | 699247 | 699753 | BCGDan_sapMKO_0619 | BCGDan_sapMKO_0619 | 507 |
| 698524 | 698994 | Rv0601c | Rv0601c | 471 | 729409 | 729879 | BCG_0647c | BCG_0647c | 471 | 699751 | 700221 | BCGDan_0622 | BCGDan_0622 | 471 | 699867 | 700397 | BCGDan_sapMKO_0620 | BCGDan_sapMKO_0620 | 531 |
| 699038 | 699799 | tcrA | Rv0602c | 762 | 729923 | 730684 | tcrA | BCG_0648c | 762 | 700265 | 701026 | tcrA | BCGDan_0623 | 762 | 700381 | 701085 | tcrA | BCGDan_sapMKO_0621 | 705 |
| 699856 | 700167 | Rv0603 | Rv0603 | 312 | 730741 | 731052 | BCG_0649 | BCG_0649 | 312 | 701083 | 701394 | BCGDan_0624 | BCGDan_0624 | 312 | 701199 | 701510 | BCGDan_sapMKO_0622 | BCGDan_sapMKO_0622 | 312 |
| 700239 | 701189 | lpqO | Rv0604 | 951 | 731124 | 732074 | lpqO | BCG_0650 | 951 | 701466 | 702416 | lpqO | BCGDan_0625 | 951 | 701606 | 702532 | lpqO | BCGDan_sapMKO_0623 | 927 |
| 701406 | 702014 | Rv0605 | Rv0605 | 609 | 732291 | 732899 | BCG_0651 | BCG_0651 | 609 | 702633 | 703241 | BCGDan_0626 | BCGDan_0626 | 609 | 702773 | 703357 | BCGDan_sapMKO_0624 | BCGDan_sapMKO_0624 | 585 |
| 702016 | 702759 | Rv0606 | Rv0606 | 744 | 732901 | 733644 | BCG_0652 | BCG_0652 | 744 | 703243 | 703986 | BCGDan_0627 | BCGDan_0627 | 744 | 703359 | 704102 | BCGDan_sapMKO_0625 | BCGDan_sapMKO_0625 | 744 |
| 702813 | 703199 | Rv0607 | Rv0607 | 387 | 733698 | 734084 | BCG_0653 | BCG_0653 | 387 | 704040 | 704426 | BCGDan_0628 | BCGDan_0628 | 387 | 704156 | 704542 | BCGDan_sapMKO_0626 | BCGDan_sapMKO_0626 | 387 |
| 703244 | 703489 | vapB28 | Rv0608 | 246 | 734129 | 734374 | BCG_0654 | BCG_0654 | 246 | 704471 | 704716 | BCGDan_0629 | BCGDan_0629 | 246 | 704587 | 704832 | BCGDan_sapMKO_0627 | BCGDan_sapMKO_0627 | 246 |
| 703486 | 703887 | vapC28 | Rv0609 | 402 | 734371 | 734772 | BCG_0655 | BCG_0655 | 402 | 704713 | 705114 | BCGDan_0630 | BCGDan_0630 | 402 | 704829 | 705230 | BCGDan_sapMKO_0628 | BCGDan_sapMKO_0628 | 402 |
| 703830 | 704057 | Rv0609A | Rv0609A | 228 | 734715 | 734942 | BCG_0656 | BCG_0656 | 228 | 705057 | 705284 | BCGDan_0631 | BCGDan_0631 | 228 | 705173 | 705400 | BCGDan_sapMKO_0629 | BCGDan_sapMKO_0629 | 228 |
| 704752 | 705909 | Rv0610c | Rv0610c | 1158 | 735637 | 736794 | BCG_0657c | BCG_0657c | 1158 | 705979 | 707136 | BCGDan_0632 | BCGDan_0632 | 1158 | 706095 | 707252 | BCGDan_sapMKO_0630 | BCGDan_sapMKO_0630 | 1158 |
| 705961 | 706344 | Rv0611c | Rv0611c | 384 | 736846 | 737229 | BCG_0658c | BCG_0658c | 384 | 707188 | 707571 | BCGDan_0633 | BCGDan_0633 | 384 | 707304 | 707687 | BCGDan_sapMKO_0631 | BCGDan_sapMKO_0631 | 384 |
| 706324 | 706929 | Rv0612 | Rv0612 | 606 | 737209 | 737814 | BCG_0659 | BCG_0659 | 606 | 707551 | 708156 | BCGDan_0634 | BCGDan_0634 | 606 | 707667 | 708272 | BCGDan_sapMKO_0632 | BCGDan_sapMKO_0632 | 606 |
| 706948 | 709515 | Rv0613c | Rv0613c | 2568 | 737833 | 740400 | BCG_0660c | BCG_0660c | 2568 | 708175 | 710742 | BCGDan_0635 | BCGDan_0635 | 2568 | 708291 | 710867 | BCGDan_sapMKO_0633 | BCGDan_sapMKO_0633 | 2577 |
| 709356 | 710348 | Rv0614 | Rv0614 | 993 | 740241 | 741233 | BCG_0661 | BCG_0661 | 993 | 710583 | 711575 | BCGDan_0636 | BCGDan_0636 | 993 | 710699 | 711691 | BCGDan_sapMKO_0634 | BCGDan_sapMKO_0634 | 993 |
| 710345 | 710587 | Rv0615 | Rv0615 | 243 | 741230 | 741472 | BCG_0662 | BCG_0662 | 243 | 711572 | 711814 | BCGDan_0637 | BCGDan_0637 | 243 | 711691 | 711930 | BCGDan_sapMKO_0635 | BCGDan_sapMKO_0635 | 240 |
| 710584 | 710850 | Rv0616c | Rv0616c | 267 | 741469 | 741735 | BCG_0663c | BCG_0663c | 267 | 711811 | 712077 | BCGDan_0638 | BCGDan_0638 | 267 | 711927 | 712193 | BCGDan_sapMKO_0636 | BCGDan_sapMKO_0636 | 267 |
| 710782 | 711009 | vapB29 | Rv0616A | 228 | 741667 | 741894 | BCG_0663A | BCG_0663A | 228 | 712009 | 712236 | BCGDan_0639 | BCGDan_0639 | 228 | 712125 | 712352 | BCGDan_sapMKO_0637 | BCGDan_sapMKO_0637 | 228 |
| 711006 | 711407 | vapC29 | Rv0617 | 402 | 741891 | 742292 | BCG_0664 | BCG_0664 | 402 | 712233 | 712634 | BCGDan_0640 | BCGDan_0640 | 402 | 712349 | 712750 | BCGDan_sapMKO_0638 | BCGDan_sapMKO_0638 | 402 |
| 711536 | 712231 | galTa | Rv0618 | 696 | 742421 | 743605 | galT | BCG_0665 | 1185 | 712763 | 713947 | galT | BCGDan_0641 | 1185 | 712879 | 714063 | galT | BCGDan_sapMKO_0639 | 1185 |
| 712716 | 713807 | galK | Rv0620 | 1092 | 743602 | 744693 | galK | BCG_0666 | 1092 | 713944 | 715035 | galK | BCGDan_0642 | 1092 | 714060 | 715151 | galK | BCGDan_sapMKO_0640 | 1092 |
| 714202 | 715266 | Rv0621 | Rv0621 | 1065 | 745088 | 746245 | BCG_0667 | BCG_0667 | 1158 | 715430 | 716587 | BCGDan_0643 | BCGDan_0643 | 1158 | 715651 | 716703 | BCGDan_sapMKO_0641 | BCGDan_sapMKO_0641 | 1053 |
| 715370 | 716317 | Rv0622 | Rv0622 | 948 | 746256 | 747203 | BCG_0668 | BCG_0668 | 948 | 716598 | 717545 | BCGDan_0644 | BCGDan_0644 | 948 | 716726 | 717661 | BCGDan_sapMKO_0642 | BCGDan_sapMKO_0642 | 936 |
| 716410 | 716664 | vapB30 | Rv0623 | 255 | 747296 | 747550 | BCG_0669 | BCG_0669 | 255 | 717638 | 717892 | BCGDan_0645 | BCGDan_0645 | 255 | 717754 | 718008 | BCGDan_sapMKO_0643 | BCGDan_sapMKO_0643 | 255 |
| 716664 | 717059 | vapC30 | Rv0624 | 396 | 747550 | 747945 | BCG_0670 | BCG_0670 | 396 | 717892 | 718287 | BCGDan_0646 | BCGDan_0646 | 396 | 718008 | 718403 | BCGDan_sapMKO_0644 | BCGDan_sapMKO_0644 | 396 |
| 717153 | 717893 | Rv0625c | Rv0625c | 741 | 748039 | 748779 | BCG_0671c | BCG_0671c | 741 | 718381 | 719121 | BCGDan_0647 | BCGDan_0647 | 741 | 718497 | 719237 | BCGDan_sapMKO_0645 | BCGDan_sapMKO_0645 | 741 |
| 718025 | 718285 | vapB5 | Rv0626 | 261 | 748911 | 749171 | BCG_0672 | BCG_0672 | 261 | 719253 | 719513 | BCGDan_0648 | BCGDan_0648 | 261 | 719369 | 719629 | BCGDan_sapMKO_0646 | BCGDan_sapMKO_0646 | 261 |
| 718282 | 718689 | vapC5 | Rv0627 | 408 | 749168 | 749575 | BCG_0673 | BCG_0673 | 408 | 719510 | 719917 | BCGDan_0649 | BCGDan_0649 | 408 | 719626 | 720033 | BCGDan_sapMKO_0647 | BCGDan_sapMKO_0647 | 408 |
| 718761 | 719912 | Rv0628c | Rv0628c | 1152 | 749647 | 750798 | BCG_0674c | BCG_0674c | 1152 | 719989 | 721140 | BCGDan_0650 | BCGDan_0650 | 1152 | 720105 | 721256 | BCGDan_sapMKO_0648 | BCGDan_sapMKO_0648 | 1152 |
| 720005 | 721732 | recD | Rv0629c | 1728 | 750891 | 752618 | recD | BCG_0675c | 1728 | 721233 | 722960 | recD | BCGDan_0651 | 1728 | 721349 | 723076 | recD | BCGDan_sapMKO_0649 | 1728 |
| 721729 | 725013 | recB | Rv0630c | 3285 | 752615 | 753319 | recBb | BCG_0676c | 705 | 722957 | 723661 | recBb | BCGDan_0652 | 705 | 723073 | 723756 | BCGDan_sapMKO_0650 | BCGDan_sapMKO_0650 | 684 |
| NA | NA | NA | NA | NA | 753205 | 755898 | recBa | BCG_0677c | 2694 | 723547 | 726240 | recBa | BCGDan_0653 | 2694 | 723663 | 726356 | recBa | BCGDan_sapMKO_0651 | 2694 |
| 725013 | 728306 | recC | Rv0631c | 3294 | 755898 | 759191 | recC | BCG_0678c | 3294 | 726240 | 729533 | recC | BCGDan_0654 | 3294 | 726356 | 729649 | recC | BCGDan_sapMKO_0652 | 3294 |
| 728583 | 729278 | echA3 | Rv0632c | 696 | 759468 | 760163 | echA3 | BCG_0679c | 696 | 729810 | 730505 | echA3 | BCGDan_0655 | 696 | 729926 | 730621 | echA3 | BCGDan_sapMKO_0653 | 696 |
| 729327 | 730166 | Rv0633c | Rv0633c | 840 | 760212 | 761042 | BCG_0680c | BCG_0680c | 831 | 730554 | 731384 | BCGDan_0656 | BCGDan_0656 | 831 | 730670 | 731500 | BCGDan_sapMKO_0654 | BCGDan_sapMKO_0654 | 831 |
| 730320 | 731033 | Rv0634c | Rv0634c | 714 | 761196 | 761909 | BCG_0681c | BCG_0681c | 714 | 731538 | 732251 | BCGDan_0657 | BCGDan_0657 | 714 | 731654 | 732367 | BCGDan_sapMKO_0655 | BCGDan_sapMKO_0655 | 714 |
| 731113 | 731364 | Rv0634A | Rv0634A | 252 | 761989 | 762240 | BCG_0682 | BCG_0682 | 252 | 732331 | 732582 | BCGDan_0658 | BCGDan_0658 | 252 | 732447 | 732698 | BCGDan_sapMKO_0656 | BCGDan_sapMKO_0656 | 252 |
| 731712 | 731879 | rpmG2 | Rv0634B | 168 | 762588 | 762755 | rpmG2 | BCG_0683 | 168 | 732930 | 733097 | rpmG2 | BCGDan_0661 | 168 | 733046 | 733213 | rpmG2 | BCGDan_sapMKO_0659 | 168 |
| 731930 | 732406 | hadA | Rv0635 | 477 | 762806 | 763282 | BCG_0684 | BCG_0684 | 477 | 733148 | 733624 | BCGDan_0662 | BCGDan_0662 | 477 | 733291 | 733740 | BCGDan_sapMKO_0660 | BCGDan_sapMKO_0660 | 450 |
| 732393 | 732821 | hadB | Rv0636 | 429 | 763269 | 763697 | BCG_0685 | BCG_0685 | 429 | 733611 | 734039 | BCGDan_0663 | BCGDan_0663 | 429 | 733727 | 734155 | BCGDan_sapMKO_0661 | BCGDan_sapMKO_0661 | 429 |
| 732825 | 733325 | hadC | Rv0637 | 501 | 763701 | 764201 | BCG_0686 | BCG_0686 | 501 | 734043 | 734543 | BCGDan_0664 | BCGDan_0664 | 501 | 734159 | 734659 | BCGDan_sapMKO_0662 | BCGDan_sapMKO_0662 | 501 |
| 733737 | 734222 | secE1 | Rv0638 | 486 | 764613 | 765098 | secE1 | BCG_0687 | 486 | 734955 | 735440 | secE1 | BCGDan_0666 | 486 | 735071 | 735556 | secE1 | BCGDan_sapMKO_0664 | 486 |
| 734254 | 734970 | nusG | Rv0639 | 717 | 765130 | 765846 | nusG | BCG_0688 | 717 | 735472 | 736188 | nusG | BCGDan_0667 | 717 | 735588 | 736304 | nusG | BCGDan_sapMKO_0665 | 717 |
| 735022 | 735450 | rplK | Rv0640 | 429 | 765898 | 766326 | rplK | BCG_0689 | 429 | 736240 | 736668 | rplK | BCGDan_0668 | 429 | 736356 | 736784 | rplK | BCGDan_sapMKO_0666 | 429 |
| 735517 | 736224 | rplA | Rv0641 | 708 | 766393 | 767100 | rplA | BCG_0690 | 708 | 736735 | 737442 | rplA | BCGDan_0669 | 708 | 736851 | 737558 | rplA | BCGDan_sapMKO_0667 | 708 |
| 736298 | 737203 | mmaA4 | Rv0642c | 906 | 767174 | 768079 | mmaA4 | BCG_0691c | 906 | 737516 | 738421 | mmaA4 | BCGDan_0670 | 906 | 737632 | 738528 | mmaA4 | BCGDan_sapMKO_0668 | 897 |
| 737268 | 738149 | mmaA3 | Rv0643c | 882 | 768144 | 769025 | mmaA3 | BCG_0692c | 882 | 738486 | 739367 | mmaA3 | BCGDan_0671 | 882 | 738602 | 739483 | mmaA3 | BCGDan_sapMKO_0669 | 882 |
| 738297 | 739160 | mmaA2 | Rv0644c | 864 | 769173 | 770036 | mmaA2 | BCG_0693c | 864 | 739515 | 740378 | mmaA2 | BCGDan_0672 | 864 | 739631 | 740494 | mmaA2 | BCGDan_sapMKO_0670 | 864 |
| 739327 | 740187 | mmaA1 | Rv0645c | 861 | 770203 | 771063 | mmaA1 | BCG_0694c | 861 | 740545 | 741405 | mmaA1 | BCGDan_0673 | 861 | 740661 | 741521 | mmaA1 | BCGDan_sapMKO_0671 | 861 |
| 740234 | 741139 | lipG | Rv0646c | 906 | 771110 | 772015 | lipG | BCG_0695c | 906 | 741452 | 742357 | lipG | BCGDan_0674 | 906 | 741568 | 742473 | lipG | BCGDan_sapMKO_0672 | 906 |
| 741151 | 742617 | Rv0647c | Rv0647c | 1467 | 772027 | 773493 | BCG_0696c | BCG_0696c | 1467 | 742369 | 743835 | BCGDan_0675 | BCGDan_0675 | 1467 | 742485 | 743831 | BCGDan_sapMKO_0673 | BCGDan_sapMKO_0673 | 1347 |
| 742719 | 746366 | Rv0648 | Rv0648 | 3648 | 773579 | 777775 | BCG_0697 | BCG_0697 | 4197 | 743921 | 748117 | BCGDan_0676 | BCGDan_0676 | 4197 | 744037 | 748233 | BCGDan_sapMKO_0674 | BCGDan_sapMKO_0674 | 4197 |
| 746363 | 747037 | fabD2 | Rv0649 | 675 | 777772 | 778446 | fabD2 | BCG_0698 | 675 | 748114 | 748788 | fabD2 | BCGDan_0677 | 675 | 748230 | 748904 | fabD2 | BCGDan_sapMKO_0675 | 675 |
| 747037 | 747945 | Rv0650 | Rv0650 | 909 | 778446 | 779354 | BCG_0699 | BCG_0699 | 909 | 748788 | 749696 | BCGDan_0678 | BCGDan_0678 | 909 | 748904 | 749812 | BCGDan_sapMKO_0676 | BCGDan_sapMKO_0676 | 909 |
| 748276 | 748812 | rplJ | Rv0651 | 537 | 779685 | 780221 | rplJ | BCG_0700 | 537 | 750027 | 750563 | rplJ | BCGDan_0679 | 537 | 750143 | 750679 | rplJ | BCGDan_sapMKO_0677 | 537 |
| 748849 | 749241 | rplL | Rv0652 | 393 | 780258 | 780650 | rplL | BCG_0701 | 393 | 750600 | 750992 | rplL | BCGDan_0680 | 393 | 750716 | 751108 | rplL | BCGDan_sapMKO_0678 | 393 |
| 749234 | 749929 | Rv0653c | Rv0653c | 696 | 780643 | 781338 | BCG_0702c | BCG_0702c | 696 | 750985 | 751680 | BCGDan_0681 | BCGDan_0681 | 696 | 751101 | 751868 | BCGDan_sapMKO_0679 | BCGDan_sapMKO_0679 | 768 |
| 750000 | 751505 | Rv0654 | Rv0654 | 1506 | 781409 | 782914 | BCG_0703 | BCG_0703 | 1506 | 751751 | 753256 | BCGDan_0682 | BCGDan_0682 | 1506 | 751867 | 753372 | BCGDan_sapMKO_0680 | BCGDan_sapMKO_0680 | 1506 |
| 751517 | 752596 | mkl | Rv0655 | 1080 | 782926 | 784005 | mkl | BCG_0704 | 1080 | 753268 | 754347 | mkl | BCGDan_0683 | 1080 | 753453 | 754463 | mkl | BCGDan_sapMKO_0681 | 1011 |
| 752984 | 753367 | vapC6 | Rv0656c | 384 | 784393 | 784776 | BCG_0705c | BCG_0705c | 384 | 754735 | 755118 | BCGDan_0684 | BCGDan_0684 | 384 | 754851 | 755234 | BCGDan_sapMKO_0682 | BCGDan_sapMKO_0682 | 384 |
| 753462 | 753617 | vapB6 | Rv0657c | 156 | 784871 | 785026 | BCG_0706c | BCG_0706c | 156 | 755213 | 755368 | BCGDan_0685 | BCGDan_0685 | 156 | 755329 | 755484 | BCGDan_sapMKO_0683 | BCGDan_sapMKO_0683 | 156 |
| 753693 | 754409 | Rv0658c | Rv0658c | 717 | 785102 | 785818 | BCG_0707c | BCG_0707c | 717 | 755444 | 756160 | BCGDan_0686 | BCGDan_0686 | 717 | 755560 | 756276 | BCGDan_sapMKO_0684 | BCGDan_sapMKO_0684 | 717 |
| 754685 | 754993 | mazF2 | Rv0659c | 309 | 786094 | 786402 | BCG_0708c | BCG_0708c | 309 | 756436 | 756744 | BCGDan_0687 | BCGDan_0687 | 309 | 756552 | 756860 | BCGDan_sapMKO_0685 | BCGDan_sapMKO_0685 | 309 |
| 754980 | 755225 | mazE2 | Rv0660c | 246 | 786389 | 786634 | BCG_0709c | BCG_0709c | 246 | 756731 | 756976 | BCGDan_0688 | BCGDan_0688 | 246 | 756847 | 757098 | BCGDan_sapMKO_0686 | BCGDan_sapMKO_0686 | 252 |
| 755335 | 755772 | vapC7 | Rv0661c | 438 | 786744 | 787181 | BCG_0710c | BCG_0710c | 438 | 757086 | 757523 | BCGDan_0689 | BCGDan_0689 | 438 | 757202 | 757633 | BCGDan_sapMKO_0687 | BCGDan_sapMKO_0687 | 432 |
| 755769 | 756023 | vapB7 | Rv0662c | 255 | 787178 | 787546 | BCG_0711c | BCG_0711c | 369 | 757520 | 757774 | BCGDan_0690 | BCGDan_0690 | 255 | 757636 | 758004 | BCGDan_sapMKO_0688 | BCGDan_sapMKO_0688 | 369 |
| 756137 | 758500 | atsD | Rv0663 | 2364 | 787546 | 789909 | atsD | BCG_0712 | 2364 | 757888 | 760251 | atsD | BCGDan_0691 | 2364 | 758004 | 760367 | atsD | BCGDan_sapMKO_0689 | 2364 |

| | | | | | | | | | | | | | | | | | | | |
|---|---|---|---|---|---|---|---|---|---|---|---|---|---|---|---|---|---|---|---|
| 758532 | 758804 | vapB8 | Rv0664 | 273 | 789941 | 790213 | BCG_0713 | BCG_0713 | 273 | 760283 | 760555 | BCGDan_0692 | BCGDan_0692 | 273 | 760399 | 760671 | BCGDan_sapMKO_0690 | BCGDan_sapMKO_0690 | 273 |
| 758801 | 759139 | vapC8 | Rv0665 | 339 | 790210 | 790548 | BCG_0714 | BCG_0714 | 339 | 760552 | 760890 | BCGDan_0693 | BCGDan_0693 | 339 | 760668 | 761006 | BCGDan_sapMKO_0691 | BCGDan_sapMKO_0691 | 339 |
| 759136 | 759309 | Rv0666 | Rv0666 | 174 | 790545 | 790718 | BCG_0715 | BCG_0715 | 174 | 760887 | 761060 | BCGDan_0694 | BCGDan_0694 | 174 | 761003 | 761176 | BCGDan_sapMKO_0692 | BCGDan_sapMKO_0692 | 174 |
| 759807 | 763325 | rpoB | Rv0667 | 3519 | 791216 | 794734 | rpoB | BCG_0716 | 3519 | 761558 | 765076 | rpoB | BCGDan_0695 | 3519 | 761902 | 765192 | rpoB | BCGDan_sapMKO_0693 | 3291 |
| 763370 | 767320 | rpoC | Rv0668 | 3951 | 794779 | 798729 | rpoC | BCG_0717 | 3951 | 765121 | 769071 | rpoC | BCGDan_0696 | 3951 | 765237 | 769187 | rpoC | BCGDan_sapMKO_0694 | 3951 |
| 767684 | 769597 | Rv0669c | Rv0669c | 1914 | 799093 | 801006 | BCG_0718c | BCG_0718c | 1914 | 769435 | 771348 | BCGDan_0697 | BCGDan_0697 | 1914 | 769551 | 771464 | BCGDan_sapMKO_0695 | BCGDan_sapMKO_0695 | 1914 |
| 769792 | 770550 | end | Rv0670 | 759 | 801201 | 801959 | end | BCG_0719 | 759 | 771543 | 772301 | end | BCGDan_0698 | 759 | 771659 | 772417 | end | BCGDan_sapMKO_0696 | 759 |
| 770582 | 771424 | lpqP | Rv0671 | 843 | 801991 | 802833 | lpqP | BCG_0720 | 843 | 772333 | 773175 | lpqP | BCGDan_0699 | 843 | 772449 | 773291 | lpqP | BCGDan_sapMKO_0697 | 843 |
| 771484 | 773112 | fadE8 | Rv0672 | 1629 | 802893 | 804521 | fadE8 | BCG_0721 | 1629 | 773235 | 774863 | fadE8 | BCGDan_0700 | 1629 | 773351 | 774979 | fadE8 | BCGDan_sapMKO_0698 | 1629 |
| 773123 | 774061 | echA4 | Rv0673 | 939 | 804532 | 805470 | echA4 | BCG_0722 | 939 | 774874 | 775812 | echA4 | BCGDan_0701 | 939 | 774990 | 775928 | echA4 | BCGDan_sapMKO_0699 | 939 |
| 774064 | 774786 | Rv0674 | Rv0674 | 723 | 805473 | 806195 | BCG_0723 | BCG_0723 | 723 | 775815 | 776537 | BCGDan_0702 | BCGDan_0702 | 723 | 775931 | 776653 | BCGDan_sapMKO_0700 | BCGDan_sapMKO_0700 | 723 |
| 774783 | 775574 | echA5 | Rv0675 | 792 | 806192 | 806983 | echA5 | BCG_0724 | 792 | 776534 | 777325 | echA5 | BCGDan_0703 | 792 | 776650 | 777441 | echA5 | BCGDan_sapMKO_0701 | 792 |
| 775586 | 778480 | mmpL5 | Rv0676c | 2895 | 806995 | 809889 | mmpL5 | BCG_0725c | 2895 | 777337 | 780231 | mmpL5 | BCGDan_0704 | 2895 | 777453 | 780269 | mmpL5 | BCGDan_sapMKO_0702 | 2817 |
| 778477 | 778905 | mmpS5 | Rv0677c | 429 | 809886 | 810314 | mmpS5 | BCG_0726c | 429 | 780228 | 780656 | mmpS5 | BCGDan_0705 | 429 | 780344 | 780832 | mmpS5 | BCGDan_sapMKO_0703 | 489 |
| 778990 | 779487 | Rv0678 | Rv0678 | 498 | 810399 | 810896 | BCG_0727 | BCG_0727 | 498 | 780741 | 781238 | BCGDan_0706 | BCGDan_0706 | 498 | 780857 | 781354 | BCGDan_sapMKO_0704 | BCGDan_sapMKO_0704 | 498 |
| 779543 | 780040 | Rv0679c | Rv0679c | 498 | 810952 | 811449 | BCG_0728c | BCG_0728c | 498 | 781294 | 781791 | BCGDan_0707 | BCGDan_0707 | 498 | 781410 | 781907 | BCGDan_sapMKO_0705 | BCGDan_sapMKO_0705 | 498 |
| 780042 | 780416 | Rv0680c | Rv0680c | 375 | 811451 | 811825 | BCG_0729c | BCG_0729c | 375 | 781793 | 782167 | BCGDan_0708 | BCGDan_0708 | 375 | 781909 | 782283 | BCGDan_sapMKO_0706 | BCGDan_sapMKO_0706 | 375 |
| 780721 | 781311 | Rv0681 | Rv0681 | 591 | 812130 | 812720 | BCG_0730 | BCG_0730 | 591 | 782472 | 783062 | BCGDan_0709 | BCGDan_0709 | 591 | 782579 | 783178 | BCGDan_sapMKO_0707 | BCGDan_sapMKO_0707 | 600 |
| 781560 | 781934 | rpsL | Rv0682 | 375 | 812969 | 813343 | rpsL | BCG_0731 | 375 | 783311 | 783685 | rpsL | BCGDan_0710 | 375 | 783427 | 783801 | rpsL | BCGDan_sapMKO_0708 | 375 |
| 781934 | 782404 | rpsG | Rv0683 | 471 | 813343 | 813813 | rpsG | BCG_0732 | 471 | 783685 | 784155 | rpsG | BCGDan_0711 | 471 | 783801 | 784271 | rpsG | BCGDan_sapMKO_0709 | 471 |
| 782485 | 784590 | fusA1 | Rv0684 | 2106 | 813894 | 815999 | fusA1 | BCG_0733 | 2106 | 784236 | 786341 | fusA1 | BCGDan_0712 | 2106 | 784352 | 786457 | fusA1 | BCGDan_sapMKO_0710 | 2106 |
| 784821 | 786011 | tuf | Rv0685 | 1191 | 816230 | 817420 | tuf | BCG_0734 | 1191 | 786572 | 787762 | tuf | BCGDan_0713 | 1191 | 786688 | 787878 | tuf | BCGDan_sapMKO_0711 | 1191 |
| 786149 | 786946 | Rv0686 | Rv0686 | 798 | 817558 | 818355 | BCG_0735 | BCG_0735 | 798 | 787900 | 788697 | BCGDan_0714 | BCGDan_0714 | 798 | 788016 | 788813 | BCGDan_sapMKO_0712 | BCGDan_sapMKO_0712 | 798 |
| 787099 | 787926 | Rv0687 | Rv0687 | 828 | 818508 | 819335 | BCG_0736 | BCG_0736 | 828 | 788850 | 789677 | BCGDan_0715 | BCGDan_0715 | 828 | 788999 | 789793 | BCGDan_sapMKO_0713 | BCGDan_sapMKO_0713 | 795 |
| 787940 | 789160 | Rv0688 | Rv0688 | 1221 | 819349 | 820569 | BCG_0737 | BCG_0737 | 1221 | 789691 | 790911 | BCGDan_0716 | BCGDan_0716 | 1221 | 789807 | 791027 | BCGDan_sapMKO_0714 | BCGDan_sapMKO_0714 | 1221 |
| 789157 | 789411 | Rv0689c | Rv0689c | 255 | 820566 | 820820 | BCG_0738c | BCG_0738c | 255 | 790908 | 791162 | BCGDan_0717 | BCGDan_0717 | 255 | 791024 | 791278 | BCGDan_sapMKO_0715 | BCGDan_sapMKO_0715 | 255 |
| 790024 | 791073 | Rv0690c | Rv0690c | 1050 | 821433 | 822482 | BCG_0739c | BCG_0739c | 1050 | 791775 | 792824 | BCGDan_0718 | BCGDan_0718 | 1050 | 791891 | 792940 | BCGDan_sapMKO_0716 | BCGDan_sapMKO_0716 | 1050 |
| 791070 | 791666 | Rv0691c | Rv0691c | 597 | 822479 | 823075 | BCG_0740c | BCG_0740c | 597 | 792821 | 793417 | BCGDan_0719 | BCGDan_0719 | 597 | 792937 | 793533 | BCGDan_sapMKO_0717 | BCGDan_sapMKO_0717 | 597 |
| 791658 | 791846 | Rv0691A | Rv0691A | 189 | NA | NA | NA | NA | NA | 793409 | 793597 | BCGDan_0720 | BCGDan_0720 | 189 | 793624 | 793713 | BCGDan_sapMKO_0718 | BCGDan_sapMKO_0718 | 90 |
| 791831 | 792160 | Rv0692 | Rv0692 | 330 | 823240 | 823569 | BCG_0741 | BCG_0741 | 330 | 793582 | 793911 | BCGDan_0721 | BCGDan_0721 | 330 | 793716 | 794027 | BCGDan_sapMKO_0719 | BCGDan_sapMKO_0719 | 312 |
| 792157 | 793332 | pqqE | Rv0693 | 1176 | 823566 | 824741 | pqqE | BCG_0742 | 1176 | 793908 | 795083 | pqqE | BCGDan_0722 | 1176 | 794024 | 795199 | pqqE | BCGDan_sapMKO_0720 | 1176 |
| 793335 | 794525 | lldD1 | Rv0694 | 1191 | 824744 | 825934 | lldD1 | BCG_0743 | 1191 | 795086 | 796276 | lldD1 | BCGDan_0723 | 1191 | 795202 | 796392 | lldD1 | BCGDan_sapMKO_0721 | 1191 |
| 794715 | 795470 | Rv0695 | Rv0695 | 756 | 826124 | 826879 | BCG_0744 | BCG_0744 | 756 | 796466 | 797221 | BCGDan_0724 | BCGDan_0724 | 756 | 796672 | 797337 | BCGDan_sapMKO_0722 | BCGDan_sapMKO_0722 | 666 |
| 795519 | 796931 | Rv0696 | Rv0696 | 1413 | 826928 | 828340 | BCG_0745 | BCG_0745 | 1413 | 797270 | 798682 | BCGDan_0725 | BCGDan_0725 | 1413 | 797386 | 798798 | BCGDan_sapMKO_0723 | BCGDan_sapMKO_0723 | 1413 |
| 796933 | 798372 | Rv0697 | Rv0697 | 1440 | 828342 | 829781 | BCG_0746 | BCG_0746 | 1440 | 798684 | 800123 | BCGDan_0726 | BCGDan_0726 | 1440 | 798800 | 800239 | BCGDan_sapMKO_0724 | BCGDan_sapMKO_0724 | 1440 |
| 798833 | 799444 | Rv0698 | Rv0698 | 612 | 830242 | 830571 | BCG_0747 | BCG_0747 | 330 | 800584 | 800913 | BCGDan_0727 | BCGDan_0727 | 330 | 800739 | 801029 | BCGDan_sapMKO_0725 | BCGDan_sapMKO_0725 | 291 |
| 799629 | 799850 | Rv0699 | Rv0699 | 222 | 831037 | 831258 | BCG_0749 | BCG_0749 | 222 | 801379 | 801600 | BCGDan_0728 | BCGDan_0728 | 222 | 801495 | 801716 | BCGDan_sapMKO_0727 | BCGDan_sapMKO_0727 | 222 |
| 800487 | 800792 | rpsJ | Rv0700 | 306 | 831895 | 832200 | rpsJ | BCG_0750 | 306 | 802237 | 802542 | rpsJ | BCGDan_0729 | 306 | 802353 | 802658 | rpsJ | BCGDan_sapMKO_0728 | 306 |
| 800809 | 801462 | rplC | Rv0701 | 654 | 832217 | 832870 | rplC | BCG_0751 | 654 | 802559 | 803212 | rplC | BCGDan_0730 | 654 | 802675 | 803328 | rplC | BCGDan_sapMKO_0729 | 654 |
| 801462 | 802133 | rplD | Rv0702 | 672 | 832870 | 833541 | rplD | BCG_0752 | 672 | 803212 | 803883 | rplD | BCGDan_0731 | 672 | 803328 | 803999 | rplD | BCGDan_sapMKO_0730 | 672 |
| 802133 | 802435 | rplW | Rv0703 | 303 | 833541 | 833843 | rplW | BCG_0753 | 303 | 803883 | 804185 | rplW | BCGDan_0732 | 303 | 803999 | 804301 | rplW | BCGDan_sapMKO_0731 | 303 |
| 802528 | 803370 | rplB | Rv0704 | 843 | 833990 | 834832 | rplB | BCG_0754 | 843 | 804332 | 805174 | rplB | BCGDan_0733 | 843 | 804448 | 805290 | rplB | BCGDan_sapMKO_0732 | 843 |
| 803411 | 803692 | rpsS | Rv0705 | 282 | 834873 | 835154 | rpsS | BCG_0755 | 282 | 805215 | 805496 | rpsS | BCGDan_0734 | 282 | 805331 | 805612 | rpsS | BCGDan_sapMKO_0733 | 282 |
| 803689 | 804282 | rplV | Rv0706 | 594 | 835151 | 835744 | rplV | BCG_0756 | 594 | 805493 | 806086 | rplV | BCGDan_0735 | 594 | 805609 | 806202 | rplV | BCGDan_sapMKO_0734 | 594 |
| 804282 | 805106 | rpsC | Rv0707 | 825 | 835744 | 836568 | rpsC | BCG_0757 | 825 | 806086 | 806910 | rpsC | BCGDan_0736 | 825 | 806202 | 807026 | rpsC | BCGDan_sapMKO_0735 | 825 |
| 805110 | 805526 | rplP | Rv0708 | 417 | 836572 | 836988 | rplP | BCG_0758 | 417 | 806914 | 807330 | rplP | BCGDan_0737 | 417 | 807030 | 807446 | rplP | BCGDan_sapMKO_0736 | 417 |
| 805526 | 805759 | rpmC | Rv0709 | 234 | 836988 | 837221 | rpmC | BCG_0759 | 234 | 807330 | 807563 | rpmC | BCGDan_0738 | 234 | 807446 | 807679 | rpmC | BCGDan_sapMKO_0737 | 234 |
| 805756 | 806166 | rpsQ | Rv0710 | 411 | 837218 | 837628 | rpsQ | BCG_0760 | 411 | 807563 | 807970 | rpsQ | BCGDan_0739 | 408 | 807679 | 808086 | rpsQ | BCGDan_sapMKO_0738 | 408 |
| 806335 | 808698 | atsA | Rv0711 | 2364 | 837797 | 840160 | atsA | BCG_0761 | 2364 | 808139 | 810502 | atsA | BCGDan_0740 | 2364 | 808255 | 810618 | atsA | BCGDan_sapMKO_0739 | 2364 |
| 808746 | 809645 | Rv0712 | Rv0712 | 900 | 840208 | 841107 | BCG_0762 | BCG_0762 | 900 | 810550 | 811449 | BCGDan_0741 | BCGDan_0741 | 900 | 810666 | 811565 | BCGDan_sapMKO_0740 | BCGDan_sapMKO_0740 | 900 |
| 809946 | 810887 | Rv0713 | Rv0713 | 942 | 841408 | 842349 | BCG_0763 | BCG_0763 | 942 | 811750 | 812691 | BCGDan_0742 | BCGDan_0742 | 942 | 811947 | 812807 | BCGDan_sapMKO_0741 | BCGDan_sapMKO_0741 | 861 |
| 811373 | 811741 | rplN | Rv0714 | 369 | 842836 | 843204 | rplN | BCG_0764 | 369 | 813177 | 813545 | rplN | BCGDan_0743 | 369 | 813293 | 813661 | rplN | BCGDan_sapMKO_0742 | 369 |
| 811742 | 812059 | rplX | Rv0715 | 318 | 843205 | 843522 | rplX | BCG_0765 | 318 | 813546 | 813863 | rplX | BCGDan_0744 | 318 | 813662 | 813979 | rplX | BCGDan_sapMKO_0743 | 318 |
| 812059 | 812622 | rplE | Rv0716 | 564 | 843522 | 844085 | rplE | BCG_0766 | 564 | 813863 | 814426 | rplE | BCGDan_0745 | 564 | 813979 | 814542 | rplE | BCGDan_sapMKO_0744 | 564 |
| 812627 | 812812 | rpsN1 | Rv0717 | 186 | 844090 | 844275 | rpsN1 | BCG_0767 | 186 | 814431 | 814616 | rpsN1 | BCGDan_0746 | 186 | 814547 | 814732 | rpsN1 | BCGDan_sapMKO_0745 | 186 |
| 812976 | 813374 | rpsH | Rv0718 | 399 | 844439 | 844837 | rpsH | BCG_0768 | 399 | 814780 | 815178 | rpsH | BCGDan_0747 | 399 | 814896 | 815294 | rpsH | BCGDan_sapMKO_0746 | 399 |
| 813398 | 813937 | rplF | Rv0719 | 540 | 844861 | 845400 | rplF | BCG_0769 | 540 | 815202 | 815741 | rplF | BCGDan_0748 | 540 | 815318 | 815857 | rplF | BCGDan_sapMKO_0747 | 540 |
| 813940 | 814308 | rplR | Rv0720 | 369 | 845403 | 845771 | rplR | BCG_0770 | 369 | 815744 | 816112 | rplR | BCGDan_0749 | 369 | 815860 | 816228 | rplR | BCGDan_sapMKO_0748 | 369 |
| 814328 | 814990 | rpsE | Rv0721 | 663 | 845791 | 846453 | rpsE | BCG_0771 | 663 | 816132 | 816794 | rpsE | BCGDan_0750 | 663 | 816248 | 816910 | rpsE | BCGDan_sapMKO_0749 | 663 |
| 814993 | 815190 | rpmD | Rv0722 | 198 | 846456 | 846653 | rpmD | BCG_0772 | 198 | 816797 | 816994 | rpmD | BCGDan_0751 | 198 | 816913 | 817110 | rpmD | BCGDan_sapMKO_0750 | 198 |
| 815190 | 815630 | rplO | Rv0723 | 441 | 846653 | 847093 | rplO | BCG_0773 | 441 | 816994 | 817434 | rplO | BCGDan_0752 | 441 | 817110 | 817550 | rplO | BCGDan_sapMKO_0751 | 441 |
| 815663 | 817534 | sppA | Rv0724 | 1872 | 847126 | 848997 | sppA | BCG_0774 | 1872 | 817467 | 819338 | sppA | BCGDan_0753 | 1872 | 817652 | 819454 | sppA | BCGDan_sapMKO_0752 | 1803 |
| 817539 | 818444 | Rv0725c | Rv0725c | 906 | 849002 | 849907 | BCG_0775c | BCG_0775c | 906 | 819343 | 820248 | BCGDan_0754 | BCGDan_0754 | 906 | 819459 | 820364 | BCGDan_sapMKO_0753 | BCGDan_sapMKO_0753 | 906 |
| 818537 | 819640 | Rv0726c | Rv0726c | 1104 | 850000 | 851103 | BCG_0776c | BCG_0776c | 1104 | 820341 | 821444 | BCGDan_0755 | BCGDan_0755 | 1104 | 820457 | 821560 | BCGDan_sapMKO_0754 | BCGDan_sapMKO_0754 | 1104 |
| 819843 | 820499 | fucA | Rv0727c | 657 | 851306 | 851962 | fucA | BCG_0777c | 657 | 821647 | 822303 | fucA | BCGDan_0756 | 657 | 821763 | 822419 | fucA | BCGDan_sapMKO_0755 | 657 |
| 820496 | 821476 | serA2 | Rv0728c | 981 | 851959 | 852939 | serA2 | BCG_0778c | 981 | 822300 | 823280 | serA2 | BCGDan_0757 | 981 | 822416 | 823372 | serA2 | BCGDan_sapMKO_0756 | 957 |
| 821507 | 822853 | xylB | Rv0729 | 1347 | 852970 | 854316 | xylB | BCG_0779 | 1347 | 823311 | 824657 | xylB | BCGDan_0758 | 1347 | 823427 | 824773 | xylB | BCGDan_sapMKO_0757 | 1347 |
| 822866 | 823594 | Rv0730 | Rv0730 | 729 | 854329 | 855057 | BCG_0780 | BCG_0780 | 729 | 824670 | 825398 | BCGDan_0759 | BCGDan_0759 | 729 | 824828 | 825514 | BCGDan_sapMKO_0758 | BCGDan_sapMKO_0758 | 687 |
| 823683 | 824639 | Rv0731c | Rv0731c | 957 | 855146 | 856102 | BCG_0781c | BCG_0781c | 957 | 825487 | 826443 | BCGDan_0760 | BCGDan_0760 | 957 | 825603 | 826559 | BCGDan_sapMKO_0759 | BCGDan_sapMKO_0759 | 957 |
| 824800 | 826125 | secY | Rv0732 | 1326 | 856263 | 857588 | secY | BCG_0782 | 1326 | 826604 | 827929 | secY | BCGDan_0761 | 1326 | 826720 | 828045 | secY | BCGDan_sapMKO_0760 | 1326 |
| 826122 | 826667 | adk | Rv0733 | 546 | 857585 | 858130 | adk | BCG_0783 | 546 | 827926 | 828471 | adk | BCGDan_0762 | 546 | 828042 | 828587 | adk | BCGDan_sapMKO_0761 | 546 |
| 826670 | 827470 | mapA | Rv0734 | 801 | 858133 | 858933 | mapA | BCG_0784 | 801 | 828474 | 829274 | mapA | BCGDan_0763 | 801 | 828590 | 829390 | mapA | BCGDan_sapMKO_0762 | 801 |
| 827543 | 828076 | sigL | Rv0735 | 534 | 859006 | 859539 | sigL | BCG_0785 | 534 | 829347 | 829880 | sigL | BCGDan_0764 | 534 | 829463 | 829996 | sigL | BCGDan_sapMKO_0763 | 534 |
| 828140 | 828892 | rslA | Rv0736 | 753 | 859603 | 860355 | BCG_0786 | BCG_0786 | 753 | 829944 | 830696 | BCGDan_0765 | BCGDan_0765 | 753 | 830060 | 830812 | BCGDan_sapMKO_0764 | BCGDan_sapMKO_0764 | 753 |
| 829207 | 829704 | Rv0737 | Rv0737 | 498 | 860670 | 861167 | BCG_0787 | BCG_0787 | 498 | 831011 | 831508 | BCGDan_0766 | BCGDan_0766 | 498 | 831127 | 831624 | BCGDan_sapMKO_0765 | BCGDan_sapMKO_0765 | 498 |
| 830062 | 830610 | Rv0738 | Rv0738 | 549 | 861525 | 862073 | BCG_0788 | BCG_0788 | 549 | 831866 | 832414 | BCGDan_0767 | BCGDan_0767 | 549 | 831994 | 832530 | BCGDan_sapMKO_0766 | BCGDan_sapMKO_0766 | 537 |
| 830855 | 831661 | Rv0739 | Rv0739 | 807 | 862278 | 863126 | BCG_0789 | BCG_0789 | 849 | 832619 | 833467 | BCGDan_0768 | BCGDan_0768 | 849 | 832720 | 833583 | BCGDan_sapMKO_0767 | BCGDan_sapMKO_0767 | 864 |
| 831776 | 832303 | Rv0740 | Rv0740 | 528 | 863241 | 863768 | BCG_0790 | BCG_0790 | 528 | 833582 | 834109 | BCGDan_0769 | BCGDan_0769 | 528 | 833617 | 834225 | BCGDan_sapMKO_0768 | BCGDan_sapMKO_0768 | 609 |
| 832534 | 832848 | Rv0741 | Rv0741 | 315 | 863999 | 864313 | BCG_0791 | BCG_0791 | 315 | 834340 | 834654 | BCGDan_0770 | BCGDan_0770 | 315 | 834456 | 834770 | BCGDan_sapMKO_0769 | BCGDan_sapMKO_0769 | 315 |
| 832981 | 833508 | PE_PGRS8 | Rv0742 | 528 | 864455 | 864973 | PE_PGRS8 | BCG_0792 | 519 | 834796 | 835314 | PE_PGRS8 | BCGDan_0771 | 519 | 834912 | 835430 | PE_PGRS8 | BCGDan_sapMKO_0770 | 519 |
| 833886 | 834443 | Rv0743c | Rv0743c | 558 | 865351 | 865908 | BCG_0793c | BCG_0793c | 558 | 835692 | 836249 | BCGDan_0772 | BCGDan_0772 | 558 | 835808 | 836317 | BCGDan_sapMKO_0771 | BCGDan_sapMKO_0771 | 510 |
| 834440 | 834946 | Rv0744c | Rv0744c | 507 | 865905 | 866411 | BCG_0794c | BCG_0794c | 507 | 836246 | 836752 | BCGDan_0773 | BCGDan_0773 | 507 | 836362 | 836901 | BCGDan_sapMKO_0772 | BCGDan_sapMKO_0772 | 540 |
| 835154 | 835681 | Rv0745 | Rv0745 | 528 | 866619 | 867146 | BCG_0795 | BCG_0795 | 528 | 836960 | 837487 | BCGDan_0774 | BCGDan_0774 | 528 | 837076 | 837603 | BCGDan_sapMKO_0773 | BCGDan_sapMKO_0773 | 528 |
| 835701 | 838052 | PE_PGRS9 | Rv0746 | 2352 | 867166 | 869559 | PE_PGRS9 | BCG_0796 | 2394 | 837507 | 839900 | PE_PGRS9 | BCGDan_0775 | 2394 | 837623 | 840016 | PE_PGRS9 | BCGDan_sapMKO_0774 | 2394 |
| 838451 | 840856 | PE_PGRS10 | Rv0747 | 2406 | 869958 | 872639 | PE_PGRS10 | BCG_0797 | 2682 | 840299 | 842980 | PE_PGRS10 | BCGDan_0776 | 2682 | 840427 | 843096 | PE_PGRS10 | BCGDan_sapMKO_0775 | 2670 |
| 840947 | 841204 | vapB31 | Rv0748 | 258 | 872730 | 872987 | BCG_0798 | BCG_0798 | 258 | 843071 | 843328 | BCGDan_0777 | BCGDan_0777 | 258 | 843187 | 843444 | BCGDan_sapMKO_0776 | BCGDan_sapMKO_0776 | 258 |

| | | | | | | | | | | | | | | | | | | | |
|---|---|---|---|---|---|---|---|---|---|---|---|---|---|---|---|---|---|---|---|
| 841228 | 841656 | vapC31 | Rv0749 | 429 | 873011 | 873439 | BCG_0799 | BCG_0799 | 429 | 843352 | 843780 | BCGDan_0778 | BCGDan_0778 | 429 | 843468 | 843896 | BCGDan_sapMKO_0777 | BCGDan_sapMKO_0777 | 429 |
| 841737 | 841874 | Rv0749A | Rv0749A | 138 | 873520 | 873657 | BCG_0800c | BCG_0800c | 138 | 843861 | 843998 | BCGDan_0779 | BCGDan_0779 | 138 | 843977 | 844114 | BCGDan_sapMKO_0778 | BCGDan_sapMKO_0778 | 138 |
| 842033 | 842278 | Rv0750 | Rv0750 | 246 | 873816 | 874061 | BCG_0801 | BCG_0801 | 246 | 844157 | 844402 | BCGDan_0780 | BCGDan_0780 | 246 | 844285 | 844518 | BCGDan_sapMKO_0779 | BCGDan_sapMKO_0779 | 234 |
| 842347 | 843231 | mmsB | Rv0751c | 885 | 874130 | 875014 | mmsB | BCG_0802c | 885 | 844471 | 845355 | mmsB | BCGDan_0781 | 885 | 844587 | 845471 | mmsB | BCGDan_sapMKO_0780 | 885 |
| 843242 | 844414 | fadE9 | Rv0752c | 1173 | 875025 | 876197 | fadE9 | BCG_0803c | 1173 | 845366 | 846538 | fadE9 | BCGDan_0782 | 1173 | 845482 | 846654 | fadE9 | BCGDan_sapMKO_0781 | 1173 |
| 844421 | 845953 | mmsA | Rv0753c | 1533 | 876204 | 877736 | mmsA | BCG_0804c | 1533 | 846545 | 848077 | mmsA | BCGDan_0783 | 1533 | 846661 | 848193 | mmsA | BCGDan_sapMKO_0782 | 1533 |
| 846159 | 847913 | PE_PGRS11 | Rv0754 | 1755 | 877942 | 879696 | PE_PGRS11 | BCG_0805 | 1755 | 848283 | 850037 | PE_PGRS11 | BCGDan_0784 | 1755 | 848399 | 850153 | PE_PGRS11 | BCGDan_sapMKO_0783 | 1755 |
| 848103 | 850040 | PPE12 | Rv0755c | 1938 | 879886 | 881823 | PPE12 | BCG_0806c | 1938 | 850227 | 852164 | PPE12 | BCGDan_0785 | 1938 | 850343 | 852280 | PPE12 | BCGDan_sapMKO_0784 | 1938 |
| 850342 | 850527 | Rv0755A | Rv0755A | 186 | 882125 | 882310 | BCG_0807c | BCG_0807c | 186 | 852466 | 852651 | BCGDan_0786 | BCGDan_0786 | 186 | 852582 | 852767 | BCGDan_sapMKO_0785 | BCGDan_sapMKO_0785 | 186 |
| 850741 | 851466 | Rv0756c | Rv0756c | 726 | 882524 | 883249 | BCG_0808c | BCG_0808c | 726 | 852865 | 853590 | BCGDan_0788 | BCGDan_0788 | 726 | 852981 | 853706 | BCGDan_sapMKO_0787 | BCGDan_sapMKO_0787 | 726 |
| 851608 | 852351 | phoP | Rv0757 | 744 | 883391 | 884134 | phoP | BCG_0809 | 744 | 853732 | 854475 | phoP | BCGDan_0789 | 744 | 853869 | 854591 | phoP | BCGDan_sapMKO_0788 | 723 |
| 852396 | 853853 | phoR | Rv0758 | 1458 | 884179 | 885636 | phoR | BCG_0810 | 1458 | 854520 | 854804 | BCGDan_0790 | BCGDan_0790 | 285 | 854636 | 854920 | BCGDan_sapMKO_0789 | BCGDan_sapMKO_0789 | 285 |
| NA | NA | NA | NA | NA | NA | NA | NA | NA | NA | 854792 | 855967 | BCGDan_0791 | BCGDan_0791 | 1176 | 854908 | 856083 | BCGDan_sapMKO_0790 | BCGDan_sapMKO_0790 | 1176 |
| 853825 | 854157 | Rv0759c | Rv0759c | 333 | 885608 | 885940 | BCG_0811c | BCG_0811c | 333 | 855939 | 856271 | BCGDan_0792 | BCGDan_0792 | 333 | 856055 | 856387 | BCGDan_sapMKO_0791 | BCGDan_sapMKO_0791 | 333 |
| 854267 | 854686 | Rv0760c | Rv0760c | 420 | 886049 | 886468 | BCG_0812c | BCG_0812c | 420 | 856380 | 856799 | BCGDan_0793 | BCGDan_0793 | 420 | 856496 | 856915 | BCGDan_sapMKO_0792 | BCGDan_sapMKO_0792 | 420 |
| 854699 | 855826 | adhB | Rv0761c | 1128 | 886481 | 887608 | adhB | BCG_0813c | 1128 | 856812 | 857939 | adhB | BCGDan_0794 | 1128 | 856928 | 858055 | adhB | BCGDan_sapMKO_0793 | 1128 |
| 855925 | 856470 | Rv0762c | Rv0762c | 546 | 887707 | 888252 | BCG_0814c | BCG_0814c | 546 | 858038 | 858583 | BCGDan_0795 | BCGDan_0795 | 546 | 858154 | 858699 | BCGDan_sapMKO_0794 | BCGDan_sapMKO_0794 | 546 |
| 856473 | 856679 | Rv0763c | Rv0763c | 207 | 888255 | 888461 | BCG_0815c | BCG_0815c | 207 | 858586 | 858792 | BCGDan_0796 | BCGDan_0796 | 207 | 858702 | 858908 | BCGDan_sapMKO_0795 | BCGDan_sapMKO_0795 | 207 |
| 856682 | 858037 | cyp51 | Rv0764c | 1356 | 888464 | 889819 | cyp51 | BCG_0816c | 1356 | 858795 | 860150 | cyp51 | BCGDan_0797 | 1356 | 858911 | 860266 | cyp51 | BCGDan_sapMKO_0796 | 1356 |
| 858037 | 858864 | Rv0765c | Rv0765c | 828 | 889819 | 890646 | BCG_0817c | BCG_0817c | 828 | 860150 | 860977 | BCGDan_0798 | BCGDan_0798 | 828 | 860266 | 861093 | BCGDan_sapMKO_0797 | BCGDan_sapMKO_0797 | 828 |
| 858864 | 860072 | cyp123 | Rv0766c | 1209 | 890228 | 891853 | cyp123 | BCG_0818c | 1626 | 860559 | 862184 | cyp123 | BCGDan_0799 | 1626 | 860675 | 862300 | cyp123 | BCGDan_sapMKO_0798 | 1626 |
| 860069 | 860710 | Rv0767c | Rv0767c | 642 | 891850 | 892491 | BCG_0819c | BCG_0819c | 642 | 862181 | 862822 | BCGDan_0800 | BCGDan_0800 | 642 | 862297 | 862938 | BCGDan_sapMKO_0799 | BCGDan_sapMKO_0799 | 642 |
| 860912 | 862381 | aldA | Rv0768 | 1470 | 892693 | 894162 | aldA | BCG_0820 | 1470 | 863024 | 864493 | aldA | BCGDan_0801 | 1470 | 863140 | 864609 | aldA | BCGDan_sapMKO_0800 | 1470 |
| 862412 | 863158 | Rv0769 | Rv0769 | 747 | 894193 | 894939 | BCG_0821 | BCG_0821 | 747 | 864524 | 865270 | BCGDan_0802 | BCGDan_0802 | 747 | 864640 | 865386 | BCGDan_sapMKO_0801 | BCGDan_sapMKO_0801 | 747 |
| 863256 | 864143 | Rv0770 | Rv0770 | 888 | 895037 | 895924 | BCG_0822 | BCG_0822 | 888 | 865368 | 866255 | BCGDan_0803 | BCGDan_0803 | 888 | 865547 | 866371 | BCGDan_sapMKO_0802 | BCGDan_sapMKO_0802 | 825 |
| 864140 | 864574 | Rv0771 | Rv0771 | 435 | 895921 | 896355 | BCG_0823 | BCG_0823 | 435 | 866252 | 866686 | BCGDan_0804 | BCGDan_0804 | 435 | 866371 | 866802 | BCGDan_sapMKO_0803 | BCGDan_sapMKO_0803 | 432 |
| 864586 | 865854 | purD | Rv0772 | 1269 | 896367 | 897635 | purD | BCG_0824 | 1269 | 866698 | 867966 | purD | BCGDan_0805 | 1269 | 866814 | 868082 | purD | BCGDan_sapMKO_0804 | 1269 |
| 865851 | 867389 | ggtA | Rv0773c | 1539 | 897632 | 899170 | ggtA | BCG_0825c | 1539 | 867963 | 869501 | ggtA | BCGDan_0806 | 1539 | 868079 | 869617 | ggtA | BCGDan_sapMKO_0805 | 1539 |
| 867440 | 868351 | Rv0774c | Rv0774c | 912 | 899221 | 900132 | BCG_0826c | BCG_0826c | 912 | 869552 | 870463 | BCGDan_0807 | BCGDan_0807 | 912 | 869668 | 870579 | BCGDan_sapMKO_0806 | BCGDan_sapMKO_0806 | 912 |
| 868407 | 869030 | Rv0775 | Rv0775 | 624 | 900188 | 900811 | BCG_0827 | BCG_0827 | 624 | 870519 | 871142 | BCGDan_0808 | BCGDan_0808 | 624 | 870635 | 871258 | BCGDan_sapMKO_0807 | BCGDan_sapMKO_0807 | 624 |
| 868984 | 869763 | Rv0776c | Rv0776c | 780 | 900765 | 901544 | BCG_0828c | BCG_0828c | 780 | 871096 | 871875 | BCGDan_0809 | BCGDan_0809 | 780 | 871212 | 871991 | BCGDan_sapMKO_0808 | BCGDan_sapMKO_0808 | 780 |
| 870008 | 871426 | purB | Rv0777 | 1419 | 901789 | 903207 | purB | BCG_0829 | 1419 | 872120 | 873538 | purB | BCGDan_0810 | 1419 | 872236 | 873654 | purB | BCGDan_sapMKO_0809 | 1419 |
| 871431 | 872675 | cyp126 | Rv0778 | 1245 | 903212 | 904456 | cyp126 | BCG_0830 | 1245 | 873543 | 874787 | cyp126 | BCGDan_0811 | 1245 | 873659 | 874903 | cyp126 | BCGDan_sapMKO_0810 | 1245 |
| 872672 | 873292 | Rv0779c | Rv0779c | 621 | 904453 | 905073 | BCG_0831c | BCG_0831c | 621 | 874784 | 875404 | BCGDan_0812 | BCGDan_0812 | 621 | 874900 | 875520 | BCGDan_sapMKO_0811 | BCGDan_sapMKO_0811 | 621 |
| 873343 | 874236 | purC | Rv0780 | 894 | 905124 | 906017 | purC | BCG_0832 | 894 | 875455 | 876348 | purC | BCGDan_0813 | 894 | 875571 | 876464 | purC | BCGDan_sapMKO_0812 | 894 |
| 874233 | 874943 | ptrBa | Rv0781 | 711 | 906014 | 908173 | ptrB | BCG_0833 | 2160 | 876345 | 878504 | ptrB | BCGDan_0814 | 2160 | 876461 | 878620 | ptrB | BCGDan_sapMKO_0813 | 2160 |
| 876818 | 878440 | emrB | Rv0783c | 1623 | 908601 | 910223 | emrB | BCG_0834c | 1623 | 878932 | 880554 | emrB | BCGDan_0815 | 1623 | 879048 | 880670 | emrB | BCGDan_sapMKO_0814 | 1623 |
| 878638 | 879324 | Rv0784 | Rv0784 | 687 | 910421 | 911107 | BCG_0835 | BCG_0835 | 687 | 880752 | 881438 | BCGDan_0816 | BCGDan_0816 | 687 | 880868 | 881554 | BCGDan_sapMKO_0815 | BCGDan_sapMKO_0815 | 687 |
| 879340 | 881040 | Rv0785 | Rv0785 | 1701 | 911123 | 911698 | BCG_0836 | BCG_0836 | 576 | 881454 | 882029 | BCGDan_0817 | BCGDan_0817 | 576 | 881591 | 882145 | BCGDan_sapMKO_0816 | BCGDan_sapMKO_0816 | 555 |
| NA | NA | NA | NA | NA | 911716 | 912822 | BCG_0837 | BCG_0837 | 1107 | 882047 | 883153 | BCGDan_0818 | BCGDan_0818 | 1107 | 882109 | 883269 | BCGDan_sapMKO_0817 | BCGDan_sapMKO_0817 | 1161 |
| 881075 | 881464 | Rv0786c | Rv0786c | 390 | 912857 | 913246 | BCG_0838c | BCG_0838c | 390 | 883188 | 883577 | BCGDan_0819 | BCGDan_0819 | 390 | 883304 | 883693 | BCGDan_sapMKO_0818 | BCGDan_sapMKO_0818 | 390 |
| 881459 | 882418 | Rv0787 | Rv0787 | 960 | 913241 | 914200 | BCG_0839 | BCG_0839 | 960 | 883680 | 884531 | BCGDan_0820 | BCGDan_0820 | 852 | 883796 | 884647 | BCGDan_sapMKO_0819 | BCGDan_sapMKO_0819 | 852 |
| 882524 | 882763 | Rv0787A | Rv0787A | 240 | 914306 | 914545 | BCG_0840 | BCG_0840 | 240 | 884637 | 884876 | BCGDan_0821 | BCGDan_0821 | 240 | 884753 | 884992 | BCGDan_sapMKO_0820 | BCGDan_sapMKO_0820 | 240 |
| 882760 | 883434 | purQ | Rv0788 | 675 | 914542 | 915216 | purQ | BCG_0841 | 675 | 884873 | 885547 | purQ | BCGDan_0822 | 675 | 884989 | 885663 | purQ | BCGDan_sapMKO_0821 | 675 |
| 883451 | 884050 | Rv0789c | Rv0789c | 600 | 915233 | 915832 | BCG_0842c | BCG_0842c | 600 | 885564 | 886163 | BCGDan_0823 | BCGDan_0823 | 600 | 885680 | 886279 | BCGDan_sapMKO_0822 | BCGDan_sapMKO_0822 | 600 |
| 884072 | 884800 | Rv0790c | Rv0790c | 729 | 915854 | 916582 | BCG_0843c | BCG_0843c | 729 | 886185 | 886913 | BCGDan_0824 | BCGDan_0824 | 729 | 886301 | 887029 | BCGDan_sapMKO_0823 | BCGDan_sapMKO_0823 | 729 |
| 884797 | 885840 | Rv0791c | Rv0791c | 1044 | 916579 | 917622 | BCG_0844c | BCG_0844c | 1044 | 886910 | 887953 | BCGDan_0825 | BCGDan_0825 | 1044 | 887026 | 888069 | BCGDan_sapMKO_0824 | BCGDan_sapMKO_0824 | 1044 |
| 885837 | 886646 | Rv0792c | Rv0792c | 810 | 917619 | 918428 | BCG_0845c | BCG_0845c | 810 | 887950 | 888759 | BCGDan_0826 | BCGDan_0826 | 810 | 888066 | 888875 | BCGDan_sapMKO_0825 | BCGDan_sapMKO_0825 | 810 |
| 886719 | 887024 | Rv0793 | Rv0793 | 306 | 918501 | 918806 | BCG_0846 | BCG_0846 | 306 | 888832 | 889137 | BCGDan_0827 | BCGDan_0827 | 306 | 888948 | 889253 | BCGDan_sapMKO_0826 | BCGDan_sapMKO_0826 | 306 |
| 887137 | 888636 | Rv0794c | Rv0794c | 1500 | 918919 | 919530 | BCG_0847c | BCG_0847c | 612 | 889250 | 889861 | BCGDan_0828 | BCGDan_0828 | 612 | 889366 | 890067 | BCGDan_sapMKO_0827 | BCGDan_sapMKO_0827 | 702 |
| NA | NA | NA | NA | NA | 919518 | 920417 | BCG_0848c | BCG_0848c | 900 | 889849 | 890748 | BCGDan_0829 | BCGDan_0829 | 900 | 889965 | 890864 | BCGDan_sapMKO_0828 | BCGDan_sapMKO_0828 | 900 |
| 890388 | 891482 | Rv0797 | Rv0797 | 1095 | 920810 | 921904 | BCG_0849 | BCG_0849 | 1095 | 891141 | 892235 | BCGDan_0830 | BCGDan_0830 | 1095 | 891239 | 892351 | BCGDan_sapMKO_0829 | BCGDan_sapMKO_0829 | 1113 |
| 891472 | 892269 | cfp29 | Rv0798c | 798 | 921894 | 922691 | cfp29 | BCG_0850c | 798 | 892225 | 893022 | cfp29 | BCGDan_0831 | 798 | 892341 | 893138 | cfp29 | BCGDan_sapMKO_0830 | 798 |
| 892266 | 893273 | Rv0799c | Rv0799c | 1008 | 922688 | 923695 | BCG_0851c | BCG_0851c | 1008 | 893019 | 894026 | BCGDan_0832 | BCGDan_0832 | 1008 | 893135 | 894142 | BCGDan_sapMKO_0831 | BCGDan_sapMKO_0831 | 1008 |
| 893318 | 894619 | pepC | Rv0800 | 1302 | 923740 | 925041 | pepC | BCG_0852 | 1302 | 894071 | 895372 | pepC | BCGDan_0833 | 1302 | 894187 | 895488 | pepC | BCGDan_sapMKO_0832 | 1302 |
| 894631 | 894978 | Rv0801 | Rv0801 | 348 | 925053 | 925400 | BCG_0853 | BCG_0853 | 348 | 895384 | 895731 | BCGDan_0834 | BCGDan_0834 | 348 | 895500 | 895847 | BCGDan_sapMKO_0833 | BCGDan_sapMKO_0833 | 348 |
| 894972 | 895628 | Rv0802c | Rv0802c | 657 | 925394 | 926050 | BCG_0854c | BCG_0854c | 657 | 895725 | 896381 | BCGDan_0835 | BCGDan_0835 | 657 | 895841 | 896497 | BCGDan_sapMKO_0834 | BCGDan_sapMKO_0834 | 657 |
| 895820 | 898084 | purL | Rv0803 | 2265 | 926242 | 928506 | purL | BCG_0855 | 2265 | 896573 | 898837 | purL | BCGDan_0836 | 2265 | 896653 | 898953 | purL | BCGDan_sapMKO_0835 | 2301 |
| 898081 | 898710 | Rv0804 | Rv0804 | 630 | 928503 | 929132 | BCG_0856 | BCG_0856 | 630 | 898834 | 899463 | BCGDan_0837 | BCGDan_0837 | 630 | 898950 | 899579 | BCGDan_sapMKO_0836 | BCGDan_sapMKO_0836 | 630 |
| 898831 | 899787 | Rv0805 | Rv0805 | 957 | 929253 | 930209 | BCG_0857 | BCG_0857 | 957 | 899584 | 900540 | BCGDan_0838 | BCGDan_0838 | 957 | 899802 | 900656 | BCGDan_sapMKO_0837 | BCGDan_sapMKO_0837 | 855 |
| 899732 | 901330 | cpsY | Rv0806c | 1599 | 930154 | 931752 | cpsY | BCG_0858c | 1599 | 900485 | 902083 | cpsY | BCGDan_0839 | 1599 | 900601 | 902199 | cpsY | BCGDan_sapMKO_0838 | 1599 |
| 901635 | 902024 | Rv0807 | Rv0807 | 390 | 932057 | 932446 | BCG_0859 | BCG_0859 | 390 | 902388 | 902777 | BCGDan_0840 | BCGDan_0840 | 390 | 902504 | 902893 | BCGDan_sapMKO_0839 | BCGDan_sapMKO_0839 | 390 |
| 902111 | 903694 | purF | Rv0808 | 1584 | 932533 | 934116 | purF | BCG_0860 | 1584 | 902864 | 904447 | purF | BCGDan_0841 | 1584 | 902980 | 904563 | purF | BCGDan_sapMKO_0840 | 1584 |
| 903725 | 904819 | purM | Rv0809 | 1095 | 934147 | 935241 | purM | BCG_0861 | 1095 | 904478 | 905572 | purM | BCGDan_0842 | 1095 | 904594 | 905688 | purM | BCGDan_sapMKO_0841 | 1095 |
| 904905 | 905087 | Rv0810c | Rv0810c | 183 | 935327 | 935509 | BCG_0862c | BCG_0862c | 183 | 905658 | 905840 | BCGDan_0843 | BCGDan_0843 | 183 | 905774 | 905956 | BCGDan_sapMKO_0842 | BCGDan_sapMKO_0842 | 183 |
| 905234 | 906340 | Rv0811c | Rv0811c | 1107 | 935656 | 936762 | BCG_0863c | BCG_0863c | 1107 | 905987 | 907093 | BCGDan_0844 | BCGDan_0844 | 1107 | 906103 | 907209 | BCGDan_sapMKO_0843 | BCGDan_sapMKO_0843 | 1107 |
| 906423 | 907292 | Rv0812 | Rv0812 | 870 | 936845 | 937714 | BCG_0864 | BCG_0864 | 870 | 907176 | 908045 | BCGDan_0845 | BCGDan_0845 | 870 | 907274 | 908161 | BCGDan_sapMKO_0844 | BCGDan_sapMKO_0844 | 888 |
| 907338 | 908018 | Rv0813c | Rv0813c | 681 | 937760 | 938440 | BCG_0865c | BCG_0865c | 681 | 908091 | 908771 | BCGDan_0846 | BCGDan_0846 | 681 | 908207 | 908887 | BCGDan_sapMKO_0845 | BCGDan_sapMKO_0845 | 681 |
| 908181 | 908483 | sseC2 | Rv0814c | 303 | 938603 | 938905 | sseC2 | BCG_0866c | 303 | 908934 | 909236 | sseC2 | BCGDan_0847 | 303 | 909050 | 909352 | sseC2 | BCGDan_sapMKO_0846 | 303 |
| 908485 | 909318 | cysA2 | Rv0815c | 834 | 938907 | 939740 | cysA2 | BCG_0867c | 834 | 909238 | 910071 | cysA2 | BCGDan_0848 | 834 | 909354 | 910229 | cysA2 | BCGDan_sapMKO_0847 | 876 |
| 909611 | 910033 | thiX | Rv0816c | 423 | 940033 | 940455 | thiX | BCG_0868c | 423 | 910364 | 910786 | thiX | BCGDan_0849 | 423 | 910480 | 910902 | thiX | BCGDan_sapMKO_0848 | 423 |
| 910030 | 910842 | Rv0817c | Rv0817c | 813 | 940452 | 941264 | BCG_0869c | BCG_0869c | 813 | 910783 | 911595 | BCGDan_0850 | BCGDan_0850 | 813 | 910899 | 911711 | BCGDan_sapMKO_0849 | BCGDan_sapMKO_0849 | 813 |
| 910972 | 911739 | Rv0818 | Rv0818 | 768 | 941394 | 942161 | BCG_0870 | BCG_0870 | 768 | 911725 | 912492 | BCGDan_0851 | BCGDan_0851 | 768 | 911802 | 912608 | BCGDan_sapMKO_0850 | BCGDan_sapMKO_0850 | 807 |
| 911736 | 912683 | mshD | Rv0819 | 948 | 942158 | 943105 | BCG_0871 | BCG_0871 | 948 | 912489 | 913436 | BCGDan_0852 | BCGDan_0852 | 948 | 912605 | 913552 | BCGDan_sapMKO_0851 | BCGDan_sapMKO_0851 | 948 |
| 912726 | 913502 | phoT | Rv0820 | 777 | 943148 | 943660 | phoTa | BCG_0872 | 513 | 913479 | 913991 | phoTa | BCGDan_0853 | 513 | 913595 | 914107 | phoTa | BCGDan_sapMKO_0852 | 513 |
| NA | NA | NA | NA | NA | 943600 | 943923 | phoTb | BCG_0873 | 324 | 913931 | 914254 | phoTb | BCGDan_0854 | 324 | 914047 | 914367 | phoTb | BCGDan_sapMKO_0853 | 321 |
| 913558 | 914199 | phoY2 | Rv0821c | 642 | 943979 | 944620 | phoY2 | BCG_0874c | 642 | 914310 | 914951 | phoY2 | BCGDan_0855 | 642 | 914426 | 914989 | phoY2 | BCGDan_sapMKO_0854 | 564 |
| 914257 | 916311 | Rv0822c | Rv0822c | 2055 | 944678 | 946732 | BCG_0875c | BCG_0875c | 2055 | 915009 | 917063 | BCGDan_0856 | BCGDan_0856 | 2055 | 915125 | 917179 | BCGDan_sapMKO_0855 | BCGDan_sapMKO_0855 | 2055 |
| 916477 | 917646 | Rv0823c | Rv0823c | 1170 | 946898 | 948067 | BCG_0876c | BCG_0876c | 1170 | 917229 | 918398 | BCGDan_0857 | BCGDan_0857 | 1170 | 917345 | 918514 | BCGDan_sapMKO_0856 | BCGDan_sapMKO_0856 | 1170 |
| 917734 | 918750 | desA1 | Rv0824c | 1017 | 948155 | 949171 | desA1 | BCG_0877c | 1017 | 918486 | 919502 | desA1 | BCGDan_0858 | 1017 | 918602 | 919618 | desA1 | BCGDan_sapMKO_0857 | 1017 |
| 918912 | 919553 | Rv0825c | Rv0825c | 642 | 949333 | 949974 | BCG_0878c | BCG_0878c | 642 | 919664 | 920305 | BCGDan_0859 | BCGDan_0859 | 642 | 919780 | 920421 | BCGDan_sapMKO_0858 | BCGDan_sapMKO_0858 | 642 |
| 919634 | 920689 | Rv0826 | Rv0826 | 1056 | 950055 | 951110 | BCG_0879 | BCG_0879 | 1056 | 920386 | 921441 | BCGDan_0860 | BCGDan_0860 | 1056 | 920619 | 921557 | BCGDan_sapMKO_0859 | BCGDan_sapMKO_0859 | 939 |
| 920741 | 921133 | kmtR | Rv0827c | 393 | 951162 | 951554 | BCG_0880c | BCG_0880c | 393 | 921493 | 921885 | BCGDan_0861 | BCGDan_0861 | 393 | 921609 | 922001 | BCGDan_sapMKO_0860 | BCGDan_sapMKO_0860 | 393 |
| 921191 | 921613 | Rv0828c | Rv0828c | 423 | 951612 | 952034 | BCG_0881c | BCG_0881c | 423 | 921943 | 922365 | BCGDan_0862 | BCGDan_0862 | 423 | 922059 | 922481 | BCGDan_sapMKO_0861 | BCGDan_sapMKO_0861 | 423 |
| 921575 | 921865 | Rv0829 | Rv0829 | 291 | 951996 | 952286 | BCG_0882 | BCG_0882 | 291 | 922327 | 922617 | BCGDan_0863 | BCGDan_0863 | 291 | 922482 | 922733 | BCGDan_sapMKO_0862 | BCGDan_sapMKO_0862 | 252 |
| 921970 | 922875 | Rv0830 | Rv0830 | 906 | 952391 | 953296 | BCG_0883 | BCG_0883 | 906 | 922722 | 923627 | BCGDan_0864 | BCGDan_0864 | 906 | 922838 | 923743 | BCGDan_sapMKO_0863 | BCGDan_sapMKO_0863 | 906 |

| | | | | | | | | | | | | | | | | | | | |
|---|---|---|---|---|---|---|---|---|---|---|---|---|---|---|---|---|---|---|---|
| 922894 | 923709 | Rv0831c | Rv0831c | 816 | 953315 | 954130 | BCG_0884c | BCG_0884c | 816 | 923646 | 924461 | BCGDan_0865 | BCGDan_0865 | 816 | 923762 | 924577 | BCGDan_sapMKO_0864 | BCGDan_sapMKO_0864 | 816 |
| 924951 | 925364 | PE_PGRS12 | Rv0832 | 414 | 955372 | 958104 | PE_PGRS13 | BCG_0885 | 2733 | 925703 | 928435 | PE_PGRS13 | BCGDan_0870 | 2733 | 925870 | 928551 | PE_PGRS13 | BCGDan_sapMKO_0869 | 2682 |
| 927837 | 930485 | PE_PGRS14 | Rv0834c | 2649 | 958331 | 960979 | PE_PGRS14 | BCG_0886c | 2649 | 928662 | 931310 | PE_PGRS14 | BCGDan_0871 | 2649 | 928778 | 931426 | PE_PGRS14 | BCGDan_sapMKO_0870 | 2649 |
| 930953 | 931597 | lpqQ | Rv0835 | 645 | 961447 | 962091 | lpqQ | BCG_0887 | 645 | 931778 | 932422 | lpqQ | BCGDan_0872 | 645 | 931894 | 932538 | lpqQ | BCGDan_sapMKO_0871 | 645 |
| 932279 | 932932 | Rv0836c | Rv0836c | 654 | 962704 | 963426 | BCG_0888c | BCG_0888c | 723 | 933035 | 933757 | BCGDan_0873 | BCGDan_0873 | 723 | 933151 | 933966 | BCGDan_sapMKO_0872 | BCGDan_sapMKO_0872 | 816 |
| 933003 | 934031 | Rv0837c | Rv0837c | 1029 | 963497 | 964525 | BCG_0889c | BCG_0889c | 1029 | 933828 | 934856 | BCGDan_0874 | BCGDan_0874 | 1029 | 933944 | 934972 | BCGDan_sapMKO_0873 | BCGDan_sapMKO_0873 | 1029 |
| 934720 | 935490 | lpqR | Rv0838 | 771 | 965214 | 965996 | lpqR | BCG_0890 | 783 | 935545 | 936327 | lpqR | BCGDan_0875 | 783 | 935661 | 936443 | lpqR | BCGDan_sapMKO_0874 | 783 |
| 935577 | 936389 | Rv0839 | Rv0839 | 813 | 966083 | 966895 | BCG_0891 | BCG_0891 | 813 | 936414 | 937226 | BCGDan_0876 | BCGDan_0876 | 813 | 936530 | 937342 | BCGDan_sapMKO_0875 | BCGDan_sapMKO_0875 | 813 |
| 936457 | 937317 | pip | Rv0840c | 861 | 966963 | 967823 | pip | BCG_0892c | 861 | 937294 | 938154 | pip | BCGDan_0877 | 861 | 937410 | 938189 | pip | BCGDan_sapMKO_0876 | 780 |
| 937593 | 937835 | Rv0841 | Rv0841 | 243 | 968099 | 968341 | BCG_0893 | BCG_0893 | 243 | 938430 | 938672 | BCGDan_0878 | BCGDan_0878 | 243 | 938546 | 938788 | BCGDan_sapMKO_0877 | BCGDan_sapMKO_0877 | 243 |
| 938112 | 939404 | Rv0842 | Rv0842 | 1293 | 968618 | 969910 | BCG_0894 | BCG_0894 | 1293 | 938949 | 940241 | BCGDan_0879 | BCGDan_0879 | 1293 | 939029 | 940357 | BCGDan_sapMKO_0878 | BCGDan_sapMKO_0878 | 1329 |
| 939388 | 940392 | Rv0843 | Rv0843 | 1005 | 969894 | 970898 | BCG_0895 | BCG_0895 | 1005 | 940225 | 941229 | BCGDan_0880 | BCGDan_0880 | 1005 | 940362 | 941345 | BCGDan_sapMKO_0879 | BCGDan_sapMKO_0879 | 984 |
| 940456 | 941106 | narL | Rv0844c | 651 | 970962 | 971612 | narL | BCG_0896c | 651 | 941293 | 941943 | narL | BCGDan_0881 | 651 | 941409 | 942059 | narL | BCGDan_sapMKO_0880 | 651 |
| 941190 | 942467 | Rv0845 | Rv0845 | 1278 | 971696 | 972973 | BCG_0897 | BCG_0897 | 1278 | 942027 | 943304 | BCGDan_0882 | BCGDan_0882 | 1278 | 942143 | 943420 | BCGDan_sapMKO_0881 | BCGDan_sapMKO_0881 | 1278 |
| 942680 | 944194 | Rv0846c | Rv0846c | 1515 | 973186 | 974700 | BCG_0898c | BCG_0898c | 1515 | 943517 | 945031 | BCGDan_0883 | BCGDan_0883 | 1515 | 943633 | 945147 | BCGDan_sapMKO_0882 | BCGDan_sapMKO_0882 | 1515 |
| 944343 | 944735 | lpqS | Rv0847 | 393 | 974849 | 975241 | lpqS | BCG_0899 | 393 | 945180 | 945572 | lpqS | BCGDan_0884 | 393 | 945272 | 945688 | lpqS | BCGDan_sapMKO_0883 | 417 |
| 944938 | 946056 | cysK2 | Rv0848 | 1119 | 975444 | 976562 | cysK2 | BCG_0900 | 1119 | 945643 | 946893 | cysK2 | BCGDan_0885 | 1251 | 945759 | 947009 | cysK2 | BCGDan_sapMKO_0884 | 1251 |
| 946056 | 947315 | Rv0849 | Rv0849 | 1260 | 976562 | 977821 | BCG_0901 | BCG_0901 | 1260 | 946893 | 948152 | BCGDan_0886 | BCGDan_0886 | 1260 | 947009 | 948268 | BCGDan_sapMKO_0885 | BCGDan_sapMKO_0885 | 1260 |
| 947312 | 947644 | Rv0850 | Rv0850 | 333 | 977818 | 978150 | BCG_0902 | BCG_0902 | 333 | 948149 | 948481 | BCGDan_0887 | BCGDan_0887 | 333 | 948250 | 948597 | BCGDan_sapMKO_0886 | BCGDan_sapMKO_0886 | 348 |
| 947641 | 948468 | Rv0851c | Rv0851c | 828 | 978147 | 978974 | BCG_0903c | BCG_0903c | 828 | 948478 | 949305 | BCGDan_0888 | BCGDan_0888 | 828 | 948594 | 949421 | BCGDan_sapMKO_0887 | BCGDan_sapMKO_0887 | 828 |
| 948559 | 949395 | fadD16 | Rv0852 | 837 | 979065 | 979901 | fadD16 | BCG_0904 | 837 | 949396 | 950232 | fadD16 | BCGDan_0889 | 837 | 949614 | 950348 | fadD16 | BCGDan_sapMKO_0888 | 735 |
| 949436 | 951118 | pdc | Rv0853c | 1683 | 979942 | 981624 | pdc | BCG_0905c | 1683 | 950273 | 951955 | pdc | BCGDan_0890 | 1683 | 950389 | 952071 | pdc | BCGDan_sapMKO_0889 | 1683 |
| 951183 | 951626 | Rv0854 | Rv0854 | 444 | 981689 | 982132 | BCG_0906 | BCG_0906 | 444 | 952020 | 952463 | BCGDan_0891 | BCGDan_0891 | 444 | 952136 | 952579 | BCGDan_sapMKO_0890 | BCGDan_sapMKO_0890 | 444 |
| 951632 | 952711 | far | Rv0855 | 1080 | 982138 | 983217 | far | BCG_0907 | 1080 | 952469 | 953548 | far | BCGDan_0892 | 1080 | 952612 | 953664 | far | BCGDan_sapMKO_0891 | 1053 |
| 952825 | 953229 | Rv0856 | Rv0856 | 405 | 983330 | 983734 | BCG_0908 | BCG_0908 | 405 | 953661 | 954065 | BCGDan_0893 | BCGDan_0893 | 405 | 953777 | 954181 | BCGDan_sapMKO_0892 | BCGDan_sapMKO_0892 | 405 |
| 953257 | 953730 | Rv0857 | Rv0857 | 474 | 983762 | 984235 | BCG_0909 | BCG_0909 | 474 | 954093 | 954566 | BCGDan_0894 | BCGDan_0894 | 474 | 954230 | 954682 | BCGDan_sapMKO_0893 | BCGDan_sapMKO_0893 | 453 |
| 953727 | 954920 | dapC | Rv0858c | 1194 | 984232 | 985425 | BCG_0910c | BCG_0910c | 1194 | 954563 | 955756 | BCGDan_0895 | BCGDan_0895 | 1194 | 954679 | 955872 | BCGDan_sapMKO_0894 | BCGDan_sapMKO_0894 | 1194 |
| 955077 | 956288 | fadA | Rv0859 | 1212 | 985582 | 986793 | fadA | BCG_0911 | 1212 | 955913 | 957124 | fadA | BCGDan_0896 | 1212 | 956029 | 957240 | fadA | BCGDan_sapMKO_0895 | 1212 |
| 956293 | 958455 | fadB | Rv0860 | 2163 | 986798 | 988960 | fadB | BCG_0912 | 2163 | 957129 | 959291 | fadB | BCGDan_0897 | 2163 | 957245 | 959407 | fadB | BCGDan_sapMKO_0896 | 2163 |
| 958523 | 960151 | ercc3 | Rv0861c | 1629 | 989028 | 990656 | BCG_0913c | BCG_0913c | 1629 | 959359 | 960987 | BCGDan_0898 | BCGDan_0898 | 1629 | 959475 | 961103 | BCGDan_sapMKO_0897 | BCGDan_sapMKO_0897 | 1629 |
| 960342 | 962612 | Rv0862c | Rv0862c | 2271 | 990794 | 993064 | BCG_0914c | BCG_0914c | 2271 | 961125 | 963395 | BCGDan_0899 | BCGDan_0899 | 2271 | 961241 | 963511 | BCGDan_sapMKO_0898 | BCGDan_sapMKO_0898 | 2271 |
| 962599 | 962880 | Rv0863 | Rv0863 | 282 | 993051 | 993332 | BCG_0915 | BCG_0915 | 282 | 963382 | 963663 | BCGDan_0900 | BCGDan_0900 | 282 | 963498 | 963779 | BCGDan_sapMKO_0899 | BCGDan_sapMKO_0899 | 282 |
| 962890 | 963393 | moaC2 | Rv0864 | 504 | 993416 | 993844 | moaC2 | BCG_0916 | 429 | 963747 | 964175 | moaC2 | BCGDan_0901 | 429 | 963863 | 964291 | moaC2 | BCGDan_sapMKO_0900 | 429 |
| 963390 | 963872 | mog | Rv0865 | 483 | 993841 | 994323 | mog | BCG_0917 | 483 | 964172 | 964654 | mog | BCGDan_0902 | 483 | 964288 | 964770 | mog | BCGDan_sapMKO_0901 | 483 |
| 963869 | 964294 | moaE2 | Rv0866 | 426 | 994320 | 994745 | moaE2 | BCG_0918 | 426 | 964651 | 965076 | moaE2 | BCGDan_0903 | 426 | 964767 | 965192 | moaE2 | BCGDan_sapMKO_0902 | 426 |
| 964312 | 965535 | rpfA | Rv0867c | 1224 | 994763 | 995986 | rpfA | BCG_0919c | 1224 | 965094 | 966317 | rpfA | BCGDan_0904 | 1224 | 965210 | 966433 | rpfA | BCGDan_sapMKO_0903 | 1224 |
| 965983 | 966261 | moaD2 | Rv0868c | 279 | 996434 | 996712 | moaD2 | BCG_0920c | 279 | 966765 | 967043 | moaD2 | BCGDan_0905 | 279 | 966881 | 967159 | moaD2 | BCGDan_sapMKO_0904 | 279 |
| 966265 | 967347 | moaA2 | Rv0869c | 1083 | 996716 | 997798 | moaA2 | BCG_0921c | 1083 | 967047 | 968129 | moaA2 | BCGDan_0906 | 1083 | 967163 | 968245 | moaA2 | BCGDan_sapMKO_0905 | 1083 |
| 967344 | 967733 | Rv0870c | Rv0870c | 390 | 997795 | 998184 | BCG_0922c | BCG_0922c | 390 | 968126 | 968515 | BCGDan_0907 | BCGDan_0907 | 390 | 968242 | 968631 | BCGDan_sapMKO_0906 | BCGDan_sapMKO_0906 | 390 |
| 967898 | 968305 | cspB | Rv0871 | 408 | 998349 | 998756 | cspB | BCG_0923 | 408 | 968680 | 969087 | cspB | BCGDan_0908 | 408 | 968796 | 969203 | cspB | BCGDan_sapMKO_0907 | 408 |
| 968424 | 970244 | PE_PGRS15 | Rv0872c | 1821 | 998875 | 1000704 | PE_PGRS15 | BCG_0924c | 1830 | 969206 | 971035 | PE_PGRS15 | BCGDan_0909 | 1830 | 969322 | 971151 | PE_PGRS15 | BCGDan_sapMKO_0908 | 1830 |
| 970505 | 972457 | fadE10 | Rv0873 | 1953 | 1000965 | 1002917 | fadE10 | BCG_0925 | 1953 | 971296 | 973248 | fadE10 | BCGDan_0910 | 1953 | 971412 | 973364 | fadE10 | BCGDan_sapMKO_0909 | 1953 |
| 972546 | 973706 | Rv0874c | Rv0874c | 1161 | 1003006 | 1004166 | BCG_0926c | BCG_0926c | 1161 | 973337 | 974497 | BCGDan_0911 | BCGDan_0911 | 1161 | 973453 | 974613 | BCGDan_sapMKO_0910 | BCGDan_sapMKO_0910 | 1161 |
| 973806 | 974294 | Rv0875c | Rv0875c | 489 | 1004266 | 1004754 | BCG_0927c | BCG_0927c | 489 | 974597 | 975085 | BCGDan_0912 | BCGDan_0912 | 489 | 974713 | 975201 | BCGDan_sapMKO_0911 | BCGDan_sapMKO_0911 | 489 |
| 974291 | 975937 | Rv0876c | Rv0876c | 1647 | 1004751 | 1006397 | BCG_0928c | BCG_0928c | 1647 | 975082 | 976671 | BCGDan_0913 | BCGDan_0913 | 1590 | 975198 | 976787 | BCGDan_sapMKO_0912 | BCGDan_sapMKO_0912 | 1590 |
| 976075 | 976863 | Rv0877 | Rv0877 | 789 | 1006535 | 1007323 | BCG_0929 | BCG_0929 | 789 | 976866 | 977654 | BCGDan_0914 | BCGDan_0914 | 789 | 976982 | 977770 | BCGDan_sapMKO_0913 | BCGDan_sapMKO_0913 | 789 |
| 976872 | 978203 | PPE13 | Rv0878c | 1332 | 1007332 | 1008660 | PPE13 | BCG_0930c | 1329 | 977663 | 978991 | PPE13 | BCGDan_0915 | 1329 | 977779 | 979107 | PPE13 | BCGDan_sapMKO_0914 | 1329 |
| 978481 | 978756 | Rv0879c | Rv0879c | 276 | 1008938 | 1009213 | BCG_0931c | BCG_0931c | 276 | 979269 | 979544 | BCGDan_0916 | BCGDan_0916 | 276 | 979385 | 979660 | BCGDan_sapMKO_0915 | BCGDan_sapMKO_0915 | 276 |
| 978934 | 979365 | Rv0880 | Rv0880 | 432 | 1009391 | 1009822 | BCG_0932 | BCG_0932 | 432 | 979722 | 980153 | BCGDan_0917 | BCGDan_0917 | 432 | 979838 | 980269 | BCGDan_sapMKO_0916 | BCGDan_sapMKO_0916 | 432 |
| 979362 | 980228 | Rv0881 | Rv0881 | 867 | 1009819 | 1010685 | BCG_0933 | BCG_0933 | 867 | 980150 | 981016 | BCGDan_0918 | BCGDan_0918 | 867 | 980266 | 981132 | BCGDan_sapMKO_0917 | BCGDan_sapMKO_0917 | 867 |
| 980225 | 980509 | Rv0882 | Rv0882 | 285 | 1010682 | 1010966 | BCG_0934 | BCG_0934 | 285 | 981013 | 981297 | BCGDan_0919 | BCGDan_0919 | 285 | 981129 | 981413 | BCGDan_sapMKO_0918 | BCGDan_sapMKO_0918 | 285 |
| 980506 | 981267 | Rv0883c | Rv0883c | 762 | 1010963 | 1011724 | BCG_0935c | BCG_0935c | 762 | 981294 | 982055 | BCGDan_0920 | BCGDan_0920 | 762 | 981410 | 982171 | BCGDan_sapMKO_0919 | BCGDan_sapMKO_0919 | 762 |
| 981424 | 982554 | serC | Rv0884c | 1131 | 1011881 | 1013011 | serC | BCG_0936c | 1131 | 982212 | 983342 | serC | BCGDan_0921 | 1131 | 982328 | 983458 | serC | BCGDan_sapMKO_0920 | 1131 |
| 982762 | 983784 | Rv0885 | Rv0885 | 1023 | 1013219 | 1014241 | BCG_0937 | BCG_0937 | 1023 | 983550 | 984572 | BCGDan_0922 | BCGDan_0922 | 1023 | 983705 | 984688 | BCGDan_sapMKO_0921 | BCGDan_sapMKO_0921 | 984 |
| 983803 | 985530 | fprB | Rv0886 | 1728 | 1014260 | 1015987 | fprB | BCG_0938 | 1728 | 984591 | 986318 | fprB | BCGDan_0923 | 1728 | 984767 | 986434 | fprB | BCGDan_sapMKO_0922 | 1668 |
| 985513 | 985971 | Rv0887c | Rv0887c | 459 | 1015970 | 1016428 | BCG_0939c | BCG_0939c | 459 | 986301 | 986759 | BCGDan_0924 | BCGDan_0924 | 459 | 986417 | 986875 | BCGDan_sapMKO_0923 | BCGDan_sapMKO_0923 | 459 |
| 987233 | 988705 | Rv0888 | Rv0888 | 1473 | 1017690 | 1019162 | BCG_0940 | BCG_0940 | 1473 | 988021 | 989493 | BCGDan_0925 | BCGDan_0925 | 1473 | 988107 | 989609 | BCGDan_sapMKO_0924 | BCGDan_sapMKO_0924 | 1503 |
| 988740 | 989861 | citA | Rv0889c | 1122 | 1019197 | 1020318 | citA | BCG_0941c | 1122 | 989528 | 990649 | citA | BCGDan_0926 | 1122 | 989644 | 990765 | citA | BCGDan_sapMKO_0925 | 1122 |
| 989948 | 992596 | Rv0890c | Rv0890c | 2649 | 1020405 | 1023053 | BCG_0942c | BCG_0942c | 2649 | 990736 | 993384 | BCGDan_0927 | BCGDan_0927 | 2649 | 990852 | 993500 | BCGDan_sapMKO_0926 | BCGDan_sapMKO_0926 | 2649 |
| 992598 | 993455 | Rv0891c | Rv0891c | 858 | 1023055 | 1023912 | BCG_0943c | BCG_0943c | 858 | 993386 | 994243 | BCGDan_0928 | BCGDan_0928 | 858 | 993502 | 994359 | BCGDan_sapMKO_0927 | BCGDan_sapMKO_0927 | 858 |
| 993853 | 995340 | Rv0892 | Rv0892 | 1488 | 1024310 | 1025797 | BCG_0944 | BCG_0944 | 1488 | 994641 | 996128 | BCGDan_0929 | BCGDan_0929 | 1488 | 994715 | 996244 | BCGDan_sapMKO_0928 | BCGDan_sapMKO_0928 | 1530 |
| 995318 | 996295 | Rv0893c | Rv0893c | 978 | 1025775 | 1026752 | BCG_0945c | BCG_0945c | 978 | 996106 | 997083 | BCGDan_0930 | BCGDan_0930 | 978 | 996222 | 997199 | BCGDan_sapMKO_0929 | BCGDan_sapMKO_0929 | 978 |
| 996524 | 997705 | Rv0894 | Rv0894 | 1182 | 1026981 | 1028162 | BCG_0946 | BCG_0946 | 1182 | 997312 | 998493 | BCGDan_0931 | BCGDan_0931 | 1182 | 997545 | 998609 | BCGDan_sapMKO_0930 | BCGDan_sapMKO_0930 | 1065 |
| 997782 | 999299 | Rv0895 | Rv0895 | 1518 | 1028239 | 1029756 | BCG_0947 | BCG_0947 | 1518 | 998570 | 1000087 | BCGDan_0932 | BCGDan_0932 | 1518 | 998767 | 1000203 | BCGDan_sapMKO_0931 | BCGDan_sapMKO_0931 | 1437 |
| 999472 | 1000767 | gltA2 | Rv0896 | 1296 | 1029946 | 1031241 | gltA2 | BCG_0948 | 1296 | 1000277 | 1001572 | gltA2 | BCGDan_0933 | 1296 | 1000393 | 1001688 | gltA2 | BCGDan_sapMKO_0932 | 1296 |
| 1000808 | 1002415 | Rv0897c | Rv0897c | 1608 | 1031282 | 1032889 | BCG_0949c | BCG_0949c | 1608 | 1001613 | 1003220 | BCGDan_0934 | BCGDan_0934 | 1608 | 1001729 | 1003336 | BCGDan_sapMKO_0933 | BCGDan_sapMKO_0933 | 1608 |
| 1002441 | 1002704 | Rv0898c | Rv0898c | 264 | 1032915 | 1033178 | BCG_0950c | BCG_0950c | 264 | 1003246 | 1003509 | BCGDan_0935 | BCGDan_0935 | 264 | 1003362 | 1003625 | BCGDan_sapMKO_0934 | BCGDan_sapMKO_0934 | 264 |
| 1002812 | 1003792 | ompA | Rv0899 | 981 | 1033286 | 1034266 | ompA | BCG_0951 | 981 | 1003617 | 1004597 | ompA | BCGDan_0936 | 981 | 1003733 | 1004713 | ompA | BCGDan_sapMKO_0935 | 981 |
| 1003805 | 1003957 | Rv0900 | Rv0900 | 153 | 1034279 | 1034431 | BCG_0952 | BCG_0952 | 153 | 1004610 | 1004762 | BCGDan_0937 | BCGDan_0937 | 153 | 1004726 | 1004878 | BCGDan_sapMKO_0936 | BCGDan_sapMKO_0936 | 153 |
| 1003957 | 1004484 | Rv0901 | Rv0901 | 528 | 1034431 | 1034958 | BCG_0953 | BCG_0953 | 528 | 1004762 | 1005289 | BCGDan_0938 | BCGDan_0938 | 528 | 1004878 | 1005405 | BCGDan_sapMKO_0937 | BCGDan_sapMKO_0937 | 528 |
| 1004501 | 1005841 | prrB | Rv0902c | 1341 | 1034975 | 1036315 | prrB | BCG_0954c | 1341 | 1005306 | 1006646 | prrB | BCGDan_0939 | 1341 | 1005422 | 1006762 | prrB | BCGDan_sapMKO_0938 | 1341 |
| 1005852 | 1006562 | prrA | Rv0903c | 711 | 1036326 | 1037036 | prrA | BCG_0955c | 711 | 1006657 | 1007367 | prrA | BCGDan_0940 | 711 | 1006773 | 1007483 | prrA | BCGDan_sapMKO_0939 | 711 |
| 1006693 | 1008180 | accD3 | Rv0904c | 1488 | 1037167 | 1038654 | accD3 | BCG_0956c | 1488 | 1007498 | 1008985 | accD3 | BCGDan_0941 | 1488 | 1007614 | 1009101 | accD3 | BCGDan_sapMKO_0940 | 1488 |
| 1008207 | 1008938 | echA6 | Rv0905 | 732 | 1038681 | 1039412 | echA6 | BCG_0957 | 732 | 1009012 | 1009743 | echA6 | BCGDan_0942 | 732 | 1009128 | 1009859 | echA6 | BCGDan_sapMKO_0941 | 732 |
| 1008944 | 1010062 | Rv0906 | Rv0906 | 1119 | 1039418 | 1040536 | BCG_0958 | BCG_0958 | 1119 | 1009749 | 1010867 | BCGDan_0943 | BCGDan_0943 | 1119 | 1009865 | 1010983 | BCGDan_sapMKO_0942 | BCGDan_sapMKO_0942 | 1119 |
| 1010136 | 1011734 | Rv0907 | Rv0907 | 1599 | 1040629 | 1042209 | BCG_0959 | BCG_0959 | 1581 | 1010960 | 1012540 | BCGDan_0944 | BCGDan_0944 | 1581 | 1011076 | 1012656 | BCGDan_sapMKO_0943 | BCGDan_sapMKO_0943 | 1581 |
| 1011731 | 1014124 | ctpE | Rv0908 | 2394 | 1042206 | 1044599 | ctpE | BCG_0960 | 2394 | 1012537 | 1014930 | ctpE | BCGDan_0945 | 2394 | 1012653 | 1015046 | ctpE | BCGDan_sapMKO_0944 | 2394 |
| 1014681 | 1014860 | Rv0909 | Rv0909 | 180 | 1045156 | 1045335 | BCG_0961 | BCG_0961 | 180 | 1015487 | 1015666 | BCGDan_0946 | BCGDan_0946 | 180 | 1015603 | 1015782 | BCGDan_sapMKO_0945 | BCGDan_sapMKO_0945 | 180 |
| 1014866 | 1015300 | Rv0910 | Rv0910 | 435 | 1045341 | 1045775 | BCG_0962 | BCG_0962 | 435 | 1015672 | 1016106 | BCGDan_0947 | BCGDan_0947 | 435 | 1015788 | 1016222 | BCGDan_sapMKO_0946 | BCGDan_sapMKO_0946 | 435 |
| 1015398 | 1016171 | Rv0911 | Rv0911 | 774 | 1045873 | 1046646 | BCG_0963 | BCG_0963 | 774 | 1016204 | 1016977 | BCGDan_0948 | BCGDan_0948 | 774 | 1016368 | 1017093 | BCGDan_sapMKO_0947 | BCGDan_sapMKO_0947 | 726 |
| 1016236 | 1016685 | Rv0912 | Rv0912 | 450 | 1046711 | 1047160 | BCG_0964 | BCG_0964 | 450 | 1017042 | 1017491 | BCGDan_0949 | BCGDan_0949 | 450 | 1017188 | 1017607 | BCGDan_sapMKO_0948 | BCGDan_sapMKO_0948 | 420 |
| 1017217 | 1018725 | Rv0913c | Rv0913c | 1509 | 1047692 | 1049200 | BCG_0965c | BCG_0965c | 1509 | 1018023 | 1019531 | BCGDan_0950 | BCGDan_0950 | 1509 | 1018139 | 1019647 | BCGDan_sapMKO_0949 | BCGDan_sapMKO_0949 | 1509 |
| 1018727 | 1019965 | Rv0914c | Rv0914c | 1239 | 1049202 | 1050440 | BCG_0966c | BCG_0966c | 1239 | 1019533 | 1020771 | BCGDan_0951 | BCGDan_0951 | 1239 | 1019649 | 1020887 | BCGDan_sapMKO_0950 | BCGDan_sapMKO_0950 | 1239 |
| 1020058 | 1021329 | PPE14 | Rv0915c | 1272 | 1050533 | 1051804 | PPE14 | BCG_0967c | 1272 | 1020864 | 1022135 | PPE14 | BCGDan_0952 | 1272 | 1020980 | 1022209 | PPE14 | BCGDan_sapMKO_0951 | 1230 |
| 1021344 | 1021643 | PE7 | Rv0916c | 300 | 1051819 | 1052118 | PE7 | BCG_0968c | 300 | 1022150 | 1022449 | PE7 | BCGDan_0953 | 300 | 1022266 | 1022565 | PE7 | BCGDan_sapMKO_0952 | 300 |
| 1022087 | 1023868 | betP | Rv0917 | 1782 | 1052562 | 1054343 | betP | BCG_0969 | 1782 | 1022893 | 1024674 | betP | BCGDan_0954 | 1782 | 1023009 | 1024790 | betP | BCGDan_sapMKO_0953 | 1782 |

| | | | | | | | | | | | | | | | | | | | |
|---|---|---|---|---|---|---|---|---|---|---|---|---|---|---|---|---|---|---|---|
| 1024211 | 1024687 | Rv0918 | Rv0918 | 477 | 1054686 | 1055162 | BCG_0970 | BCG_0970 | 477 | 1025017 | 1025493 | BCGDan_0955 | BCGDan_0955 | 477 | 1025133 | 1025609 | BCGDan_sapMKO_0954 | BCGDan_sapMKO_0954 | 477 |
| 1024684 | 1025184 | Rv0919 | Rv0919 | 501 | 1055159 | 1055659 | BCG_0971 | BCG_0971 | 501 | 1025490 | 1025990 | BCGDan_0956 | BCGDan_0956 | 501 | 1025606 | 1026106 | BCGDan_sapMKO_0955 | BCGDan_sapMKO_0955 | 501 |
| 1025497 | 1026816 | Rv0920c | Rv0920c | 1320 | 1055972 | 1057291 | BCG_0972c | BCG_0972c | 1320 | 1026303 | 1027622 | BCGDan_0958 | BCGDan_0958 | 1320 | 1026419 | 1027738 | BCGDan_sapMKO_0957 | BCGDan_sapMKO_0957 | 1320 |
| 1027104 | 1027685 | Rv0921 | Rv0921 | 582 | 1057579 | 1058160 | BCG_0973 | BCG_0973 | 582 | 1027910 | 1028491 | BCGDan_0959 | BCGDan_0959 | 582 | 1028026 | 1028607 | BCGDan_sapMKO_0958 | BCGDan_sapMKO_0958 | 582 |
| 1027685 | 1029337 | Rv0922 | Rv0922 | 1653 | 1058160 | 1059812 | BCG_0974 | BCG_0974 | 1653 | 1028491 | 1030143 | BCGDan_0960 | BCGDan_0960 | 1653 | 1028619 | 1030259 | BCGDan_sapMKO_0959 | BCGDan_sapMKO_0959 | 1641 |
| 1029513 | 1030577 | Rv0923c | Rv0923c | 1065 | 1059988 | 1061052 | BCG_0975c | BCG_0975c | 1065 | 1030319 | 1031383 | BCGDan_0961 | BCGDan_0961 | 1065 | 1030435 | 1031499 | BCGDan_sapMKO_0960 | BCGDan_sapMKO_0960 | 1065 |
| 1030578 | 1031864 | mntH | Rv0924c | 1287 | 1061053 | 1062339 | mntH | BCG_0976c | 1287 | 1031384 | 1032670 | mntH | BCGDan_0962 | 1287 | 1031500 | 1032786 | mntH | BCGDan_sapMKO_0961 | 1287 |
| 1031896 | 1032633 | Rv0925c | Rv0925c | 738 | 1062371 | 1063108 | BCG_0977c | BCG_0977c | 738 | 1032702 | 1033439 | BCGDan_0963 | BCGDan_0963 | 738 | 1032818 | 1033555 | BCGDan_sapMKO_0962 | BCGDan_sapMKO_0962 | 738 |
| 1032710 | 1033786 | Rv0926c | Rv0926c | 1077 | 1063185 | 1064261 | BCG_0978c | BCG_0978c | 1077 | 1033516 | 1034592 | BCGDan_0964 | BCGDan_0964 | 1077 | 1033632 | 1034708 | BCGDan_sapMKO_0963 | BCGDan_sapMKO_0963 | 1077 |
| 1033840 | 1034631 | Rv0927c | Rv0927c | 792 | 1064315 | 1065106 | BCG_0979c | BCG_0979c | 792 | 1034646 | 1035437 | BCGDan_0965 | BCGDan_0965 | 792 | 1034762 | 1035553 | BCGDan_sapMKO_0964 | BCGDan_sapMKO_0964 | 792 |
| 1034903 | 1036015 | pstS3 | Rv0928 | 1113 | 1065378 | 1066490 | pstS3 | BCG_0980 | 1113 | 1035709 | 1036821 | pstS3 | BCGDan_0966 | 1113 | 1035825 | 1036937 | pstS3 | BCGDan_sapMKO_0965 | 1113 |
| 1036028 | 1037002 | pstC2 | Rv0929 | 975 | 1066503 | 1067477 | pstC2 | BCG_0981 | 975 | 1036834 | 1037808 | pstC2 | BCGDan_0967 | 975 | 1036950 | 1037924 | pstC2 | BCGDan_sapMKO_0966 | 975 |
| 1036999 | 1037925 | pstA1 | Rv0930 | 927 | 1067474 | 1068388 | pstA1 | BCG_0982 | 915 | 1037805 | 1038719 | pstA1 | BCGDan_0968 | 915 | 1037921 | 1038835 | pstA1 | BCGDan_sapMKO_0967 | 915 |
| 1037920 | 1039914 | pknD | Rv0931c | 1995 | 1068395 | 1069504 | pknDb | BCG_0983c | 1110 | 1038726 | 1039835 | pknDb | BCGDan_0969 | 1110 | 1038842 | 1040020 | pknDb | BCGDan_sapMKO_0968 | 1179 |
| NA | NA | NA | NA | NA | 1069515 | 1070390 | pknDa | BCG_0984c | 876 | 1039846 | 1040721 | pknDa | BCGDan_0970 | 876 | 1039962 | 1040837 | pknDa | BCGDan_sapMKO_0969 | 876 |
| 1039936 | 1041048 | pstS2 | Rv0932c | 1113 | 1070412 | 1071524 | pstS2 | BCG_0985c | 1113 | 1040743 | 1041855 | pstS2 | BCGDan_0971 | 1113 | 1040859 | 1041971 | pstS2 | BCGDan_sapMKO_0970 | 1113 |
| 1041264 | 1042094 | pstB | Rv0933 | 831 | 1071740 | 1071955 | pstBa | BCG_0986 | 216 | 1042071 | 1042286 | pstBa | BCGDan_0972 | 216 | 1042187 | 1042402 | pstBa | BCGDan_sapMKO_0971 | 216 |
| NA | NA | NA | NA | NA | 1071930 | 1072571 | pstBb | BCG_0987 | 642 | 1042261 | 1042902 | pstBb | BCGDan_0973 | 642 | 1042395 | 1043018 | pstBb | BCGDan_sapMKO_0972 | 624 |
| 1042115 | 1043239 | pstS1 | Rv0934 | 1125 | 1072592 | 1073716 | pstS1 | BCG_0988 | 1125 | 1042923 | 1044047 | pstS1 | BCGDan_0974 | 1125 | 1043039 | 1044163 | pstS1 | BCGDan_sapMKO_0973 | 1125 |
| 1043299 | 1044315 | pstC1 | Rv0935 | 1017 | 1073776 | 1074792 | pstC1 | BCG_0989 | 1017 | 1044107 | 1045123 | pstC1 | BCGDan_0975 | 1017 | 1044223 | 1045239 | pstC1 | BCGDan_sapMKO_0974 | 1017 |
| 1044317 | 1045222 | pstA2 | Rv0936 | 906 | 1074794 | 1075699 | pstA2 | BCG_0990 | 906 | 1045125 | 1046030 | pstA2 | BCGDan_0976 | 906 | 1045241 | 1046146 | pstA2 | BCGDan_sapMKO_0975 | 906 |
| 1045199 | 1046020 | mku | Rv0937c | 822 | 1075676 | 1076497 | BCG_0991c | BCG_0991c | 822 | 1046007 | 1046828 | BCGDan_0977 | BCGDan_0977 | 822 | 1046123 | 1046998 | BCGDan_sapMKO_0976 | BCGDan_sapMKO_0976 | 876 |
| 1046136 | 1048415 | ligD | Rv0938 | 2280 | 1076613 | 1078892 | BCG_0992 | BCG_0992 | 2280 | 1046944 | 1049223 | BCGDan_0978 | BCGDan_0978 | 2280 | 1047060 | 1049339 | BCGDan_sapMKO_0977 | BCGDan_sapMKO_0977 | 2280 |
| 1048412 | 1050346 | Rv0939 | Rv0939 | 1935 | 1078889 | 1080823 | BCG_0993 | BCG_0993 | 1935 | 1049220 | 1051154 | BCGDan_0979 | BCGDan_0979 | 1935 | 1049336 | 1051270 | BCGDan_sapMKO_0978 | BCGDan_sapMKO_0978 | 1935 |
| 1050593 | 1051459 | Rv0940c | Rv0940c | 867 | 1081070 | 1081936 | BCG_0994c | BCG_0994c | 867 | 1051401 | 1052267 | BCGDan_0980 | BCGDan_0980 | 867 | 1051517 | 1052383 | BCGDan_sapMKO_0979 | BCGDan_sapMKO_0979 | 867 |
| 1051544 | 1052317 | Rv0941c | Rv0941c | 774 | 1082021 | 1082794 | BCG_0995c | BCG_0995c | 774 | 1052352 | 1053125 | BCGDan_0981 | BCGDan_0981 | 774 | 1052468 | 1053241 | BCGDan_sapMKO_0980 | BCGDan_sapMKO_0980 | 774 |
| 1052360 | 1052638 | Rv0942 | Rv0942 | 279 | 1082837 | 1083115 | BCG_0996 | BCG_0996 | 279 | 1053168 | 1053446 | BCGDan_0982 | BCGDan_0982 | 279 | 1053284 | 1053562 | BCGDan_sapMKO_0981 | BCGDan_sapMKO_0981 | 279 |
| 1052696 | 1053736 | Rv0943c | Rv0943c | 1041 | 1083173 | 1084213 | BCG_0997c | BCG_0997c | 1041 | 1053504 | 1054544 | BCGDan_0983 | BCGDan_0983 | 1041 | 1053620 | 1054660 | BCGDan_sapMKO_0982 | BCGDan_sapMKO_0982 | 1041 |
| 1053765 | 1054241 | Rv0944 | Rv0944 | 477 | 1084242 | 1084718 | BCG_0998 | BCG_0998 | 477 | 1054573 | 1055049 | BCGDan_0984 | BCGDan_0984 | 477 | 1054683 | 1055165 | BCGDan_sapMKO_0983 | BCGDan_sapMKO_0983 | 483 |
| 1054247 | 1055008 | Rv0945 | Rv0945 | 762 | 1084724 | 1085485 | BCG_0999 | BCG_0999 | 762 | 1055055 | 1055816 | BCGDan_0985 | BCGDan_0985 | 762 | 1055171 | 1055932 | BCGDan_sapMKO_0984 | BCGDan_sapMKO_0984 | 762 |
| 1055024 | 1056685 | pgi | Rv0946c | 1662 | 1085501 | 1087162 | pgi | BCG_1000c | 1662 | 1055832 | 1057493 | pgi | BCGDan_0986 | 1662 | 1055948 | 1057609 | pgi | BCGDan_sapMKO_0985 | 1662 |
| 1057300 | 1057530 | Rv0947c | Rv0947c | 231 | 1087777 | 1088007 | BCG_1001c | BCG_1001c | 231 | 1058108 | 1058338 | BCGDan_0987 | BCGDan_0987 | 231 | 1058224 | 1058454 | BCGDan_sapMKO_0986 | BCGDan_sapMKO_0986 | 231 |
| 1057646 | 1057963 | Rv0948c | Rv0948c | 318 | 1088123 | 1088440 | BCG_1002c | BCG_1002c | 318 | 1058454 | 1058771 | BCGDan_0988 | BCGDan_0988 | 318 | 1058570 | 1058842 | BCGDan_sapMKO_0987 | BCGDan_sapMKO_0987 | 273 |
| 1058260 | 1060575 | uvrD1 | Rv0949 | 2316 | 1088737 | 1091052 | uvrD1 | BCG_1003 | 2316 | 1059068 | 1061383 | uvrD1 | BCGDan_0989 | 2316 | 1059184 | 1061499 | uvrD1 | BCGDan_sapMKO_0988 | 2316 |
| 1060656 | 1061654 | Rv0950c | Rv0950c | 999 | 1091133 | 1092131 | BCG_1004c | BCG_1004c | 999 | 1061464 | 1062462 | BCGDan_0990 | BCGDan_0990 | 999 | 1061580 | 1062578 | BCGDan_sapMKO_0989 | BCGDan_sapMKO_0989 | 999 |
| 1061964 | 1063127 | sucC | Rv0951 | 1164 | 1092441 | 1093604 | sucC | BCG_1005 | 1164 | 1062772 | 1063935 | sucC | BCGDan_0991 | 1164 | 1062888 | 1064051 | sucC | BCGDan_sapMKO_0990 | 1164 |
| 1063140 | 1064051 | sucD | Rv0952 | 912 | 1093617 | 1094528 | sucD | BCG_1006 | 912 | 1063948 | 1064859 | sucD | BCGDan_0992 | 912 | 1064073 | 1064975 | sucD | BCGDan_sapMKO_0991 | 903 |
| 1064114 | 1064962 | Rv0953c | Rv0953c | 849 | 1094591 | 1095439 | BCG_1007c | BCG_1007c | 849 | 1064922 | 1065770 | BCGDan_0993 | BCGDan_0993 | 849 | 1065038 | 1065886 | BCGDan_sapMKO_0992 | BCGDan_sapMKO_0992 | 849 |
| 1065127 | 1066038 | Rv0954 | Rv0954 | 912 | 1095604 | 1096515 | BCG_1008 | BCG_1008 | 912 | 1065935 | 1066846 | BCGDan_0994 | BCGDan_0994 | 912 | 1066051 | 1066962 | BCGDan_sapMKO_0993 | BCGDan_sapMKO_0993 | 912 |
| 1066078 | 1067445 | Rv0955 | Rv0955 | 1368 | 1096555 | 1097922 | BCG_1009 | BCG_1009 | 1368 | 1066886 | 1068253 | BCGDan_0995 | BCGDan_0995 | 1368 | 1067089 | 1068369 | BCGDan_sapMKO_0994 | BCGDan_sapMKO_0994 | 1281 |
| 1067561 | 1068208 | purN | Rv0956 | 648 | 1098038 | 1098685 | purN | BCG_1010 | 648 | 1068369 | 1069016 | purN | BCGDan_0996 | 648 | 1068485 | 1069132 | purN | BCGDan_sapMKO_0995 | 648 |
| 1068205 | 1069776 | purH | Rv0957 | 1572 | 1098682 | 1100253 | purH | BCG_1011 | 1572 | 1069013 | 1070584 | purH | BCGDan_0997 | 1572 | 1069129 | 1070700 | purH | BCGDan_sapMKO_0996 | 1572 |
| 1069883 | 1071262 | Rv0958 | Rv0958 | 1380 | 1100360 | 1101739 | BCG_1012 | BCG_1012 | 1380 | 1070691 | 1072070 | BCGDan_0998 | BCGDan_0998 | 1380 | 1070807 | 1072186 | BCGDan_sapMKO_0997 | BCGDan_sapMKO_0997 | 1380 |
| 1071255 | 1073273 | Rv0959 | Rv0959 | 2019 | 1101732 | 1103750 | BCG_1013 | BCG_1013 | 2019 | 1072063 | 1074081 | BCGDan_0999 | BCGDan_0999 | 2019 | 1072179 | 1074197 | BCGDan_sapMKO_0998 | BCGDan_sapMKO_0998 | 2019 |
| 1073327 | 1073548 | vapB9 | Rv0959A | 222 | 1103804 | 1104025 | BCG_1013A | BCG_1013A | 222 | 1074135 | 1074356 | BCGDan_1000 | BCGDan_1000 | 222 | 1074251 | 1074472 | BCGDan_sapMKO_0999 | BCGDan_sapMKO_0999 | 222 |
| 1073545 | 1073928 | vapC9 | Rv0960 | 384 | 1104022 | 1104405 | BCG_1014 | BCG_1014 | 384 | 1074353 | 1074736 | BCGDan_1001 | BCGDan_1001 | 384 | 1074469 | 1074852 | BCGDan_sapMKO_1000 | BCGDan_sapMKO_1000 | 384 |
| 1074074 | 1074421 | Rv0961 | Rv0961 | 348 | 1104551 | 1104898 | BCG_1015 | BCG_1015 | 348 | 1074882 | 1075229 | BCGDan_1002 | BCGDan_1002 | 348 | 1074998 | 1075345 | BCGDan_sapMKO_1001 | BCGDan_sapMKO_1001 | 348 |
| 1074440 | 1075114 | lprP | Rv0962c | 675 | 1104917 | 1105591 | lprP | BCG_1016c | 675 | 1075248 | 1075922 | lprP | BCGDan_1003 | 675 | 1075364 | 1076038 | lprP | BCGDan_sapMKO_1002 | 675 |
| 1075297 | 1076097 | Rv0963c | Rv0963c | 801 | 1105774 | 1106574 | BCG_1017c | BCG_1017c | 801 | 1076105 | 1076905 | BCGDan_1004 | BCGDan_1004 | 801 | 1076221 | 1077021 | BCGDan_sapMKO_1003 | BCGDan_sapMKO_1003 | 801 |
| 1076196 | 1076678 | Rv0964c | Rv0964c | 483 | 1106673 | 1107155 | BCG_1018c | BCG_1018c | 483 | 1077004 | 1077486 | BCGDan_1005 | BCGDan_1005 | 483 | 1077120 | 1077602 | BCGDan_sapMKO_1004 | BCGDan_sapMKO_1004 | 483 |
| 1076778 | 1077197 | Rv0965c | Rv0965c | 420 | 1107255 | 1107674 | BCG_1019c | BCG_1019c | 420 | 1077586 | 1078005 | BCGDan_1006 | BCGDan_1006 | 420 | 1077702 | 1078061 | BCGDan_sapMKO_1005 | BCGDan_sapMKO_1005 | 360 |
| 1077233 | 1077835 | Rv0966c | Rv0966c | 603 | 1107710 | 1108312 | BCG_1020c | BCG_1020c | 603 | 1078041 | 1078643 | BCGDan_1007 | BCGDan_1007 | 603 | 1078157 | 1078759 | BCGDan_sapMKO_1006 | BCGDan_sapMKO_1006 | 603 |
| 1077975 | 1078334 | csoR | Rv0967 | 360 | 1108451 | 1108810 | BCG_1021 | BCG_1021 | 360 | 1078782 | 1079141 | BCGDan_1008 | BCGDan_1008 | 360 | 1078898 | 1079257 | BCGDan_sapMKO_1007 | BCGDan_sapMKO_1007 | 360 |
| 1078391 | 1078687 | Rv0968 | Rv0968 | 297 | 1108867 | 1109163 | BCG_1022 | BCG_1022 | 297 | 1079198 | 1079494 | BCGDan_1009 | BCGDan_1009 | 297 | 1079314 | 1079610 | BCGDan_sapMKO_1008 | BCGDan_sapMKO_1008 | 297 |
| 1078743 | 1081055 | ctpV | Rv0969 | 2313 | 1109219 | 1111531 | ctpV | BCG_1023 | 2313 | 1079550 | 1081862 | ctpV | BCGDan_1010 | 2313 | 1079600 | 1081978 | ctpV | BCGDan_sapMKO_1009 | 2379 |
| 1081052 | 1081684 | Rv0970 | Rv0970 | 633 | 1111528 | 1112160 | BCG_1024 | BCG_1024 | 633 | 1081859 | 1082491 | BCGDan_1011 | BCGDan_1011 | 633 | 1081975 | 1082607 | BCGDan_sapMKO_1010 | BCGDan_sapMKO_1010 | 633 |
| 1081775 | 1082584 | echA7 | Rv0971c | 810 | 1112251 | 1113060 | echA7 | BCG_1025c | 810 | 1082582 | 1083391 | echA7 | BCGDan_1012 | 810 | 1082698 | 1083507 | echA7 | BCGDan_sapMKO_1011 | 810 |
| 1082584 | 1083750 | fadE12 | Rv0972c | 1167 | 1113060 | 1114226 | fadE12 | BCG_1026c | 1167 | 1083391 | 1084557 | fadE12 | BCGDan_1013 | 1167 | 1083507 | 1084673 | fadE12 | BCGDan_sapMKO_1012 | 1167 |
| 1083747 | 1085750 | accA2 | Rv0973c | 2004 | 1114223 | 1116226 | accA2 | BCG_1027c | 2004 | 1084554 | 1086557 | accA2 | BCGDan_1014 | 2004 | 1084670 | 1086679 | accA2 | BCGDan_sapMKO_1013 | 2010 |
| 1085756 | 1087345 | accD2 | Rv0974c | 1590 | 1116232 | 1117821 | accD2 | BCG_1028c | 1590 | 1086563 | 1088152 | accD2 | BCGDan_1015 | 1590 | 1086679 | 1088274 | accD2 | BCGDan_sapMKO_1014 | 1596 |
| 1087348 | 1088496 | fadE13 | Rv0975c | 1149 | 1117824 | 1118972 | fadE13 | BCG_1029c | 1149 | 1088155 | 1089303 | fadE13 | BCGDan_1016 | 1149 | 1088271 | 1089419 | fadE13 | BCGDan_sapMKO_1015 | 1149 |
| 1088493 | 1090175 | Rv0976c | Rv0976c | 1683 | 1118969 | 1120651 | BCG_1030c | BCG_1030c | 1683 | 1089300 | 1090982 | BCGDan_1017 | BCGDan_1017 | 1683 | 1089416 | 1091119 | BCGDan_sapMKO_1016 | BCGDan_sapMKO_1016 | 1704 |
| 1090373 | 1093144 | PE_PGRS16 | Rv0977 | 2772 | 1120850 | 1123621 | PE_PGRS16 | BCG_1031 | 2772 | 1091181 | 1093952 | PE_PGRS16 | BCGDan_1018 | 2772 | 1091297 | 1094068 | PE_PGRS16 | BCGDan_sapMKO_1017 | 2772 |
| 1093361 | 1094356 | PE_PGRS17 | Rv0978c | 996 | 1123838 | 1124845 | PE_PGRS17 | BCG_1032c | 1008 | 1094169 | 1095176 | PE_PGRS17 | BCGDan_1019 | 1008 | 1094285 | 1095337 | PE_PGRS17 | BCGDan_sapMKO_1018 | 1053 |
| 1094670 | 1094864 | Rv0979c | Rv0979c | 195 | 1125159 | 1125353 | BCG_1033c | BCG_1033c | 195 | 1095490 | 1095684 | BCGDan_1020 | BCGDan_1020 | 195 | 1095606 | 1095800 | BCGDan_sapMKO_1019 | BCGDan_sapMKO_1019 | 195 |
| 1094886 | 1095059 | rpmF | Rv0979A | 174 | 1125375 | 1125548 | rpmF | BCG_1034 | 174 | 1095706 | 1095879 | rpmF | BCGDan_1021 | 174 | 1095828 | 1095995 | rpmF | BCGDan_sapMKO_1020 | 168 |
| 1095078 | 1096451 | PE_PGRS18 | Rv0980c | 1374 | 1125567 | 1126940 | PE_PGRS18 | BCG_1035c | 1374 | 1095898 | 1097271 | PE_PGRS18 | BCGDan_1022 | 1374 | 1096014 | 1097387 | PE_PGRS18 | BCGDan_sapMKO_1021 | 1374 |
| 1096822 | 1097508 | mprA | Rv0981 | 687 | 1127305 | 1127997 | mprA | BCG_1036 | 693 | 1097636 | 1098328 | mprA | BCGDan_1023 | 693 | 1097752 | 1098444 | mprA | BCGDan_sapMKO_1022 | 693 |
| 1097508 | 1099022 | mprB | Rv0982 | 1515 | 1127997 | 1129511 | mprB | BCG_1037 | 1515 | 1098328 | 1099842 | mprB | BCGDan_1024 | 1515 | 1098516 | 1099958 | mprB | BCGDan_sapMKO_1023 | 1443 |
| 1099066 | 1100460 | pepD | Rv0983 | 1395 | 1129555 | 1130949 | BCG_1038 | BCG_1038 | 1395 | 1099886 | 1101280 | BCGDan_1025 | BCGDan_1025 | 1395 | 1100002 | 1101396 | BCGDan_sapMKO_1024 | BCGDan_sapMKO_1024 | 1395 |
| 1100460 | 1101005 | moaB2 | Rv0984 | 546 | 1130949 | 1131494 | moaB2 | BCG_1039 | 546 | 1101280 | 1101825 | moaB2 | BCGDan_1026 | 546 | 1101396 | 1101941 | moaB2 | BCGDan_sapMKO_1025 | 546 |
| 1101025 | 1101480 | mscL | Rv0985c | 456 | 1131514 | 1131969 | mscL | BCG_1040c | 456 | 1101845 | 1102300 | mscL | BCGDan_1027 | 456 | 1101961 | 1102464 | mscL | BCGDan_sapMKO_1026 | 504 |
| 1101803 | 1102549 | Rv0986 | Rv0986 | 747 | 1132292 | 1133038 | BCG_1041 | BCG_1041 | 747 | 1102623 | 1103369 | BCGDan_1028 | BCGDan_1028 | 747 | 1102739 | 1103485 | BCGDan_sapMKO_1027 | BCGDan_sapMKO_1027 | 747 |
| 1102542 | 1105109 | Rv0987 | Rv0987 | 2568 | 1133031 | 1135598 | BCG_1042 | BCG_1042 | 2568 | 1103362 | 1105929 | BCGDan_1029 | BCGDan_1029 | 2568 | 1103478 | 1106045 | BCGDan_sapMKO_1028 | BCGDan_sapMKO_1028 | 2568 |
| 1105116 | 1106276 | Rv0988 | Rv0988 | 1161 | 1135605 | 1136765 | BCG_1043 | BCG_1043 | 1161 | 1105936 | 1107096 | BCGDan_1030 | BCGDan_1030 | 1161 | 1106067 | 1107212 | BCGDan_sapMKO_1029 | BCGDan_sapMKO_1029 | 1146 |
| 1106405 | 1107382 | grcC2 | Rv0989c | 978 | 1136894 | 1137871 | grcC2 | BCG_1044c | 978 | 1107225 | 1108202 | grcC2 | BCGDan_1031 | 978 | 1107341 | 1108318 | grcC2 | BCGDan_sapMKO_1030 | 978 |
| 1107443 | 1108099 | Rv0990c | Rv0990c | 657 | 1137932 | 1138588 | BCG_1045c | BCG_1045c | 657 | 1108263 | 1108919 | BCGDan_1032 | BCGDan_1032 | 657 | 1108379 | 1109035 | BCGDan_sapMKO_1031 | BCGDan_sapMKO_1031 | 657 |
| 1108172 | 1108504 | Rv0991c | Rv0991c | 333 | 1138661 | 1138993 | BCG_1046c | BCG_1046c | 333 | 1108992 | 1109324 | BCGDan_1033 | BCGDan_1033 | 333 | 1109108 | 1109440 | BCGDan_sapMKO_1032 | BCGDan_sapMKO_1032 | 333 |
| 1108578 | 1109171 | Rv0992c | Rv0992c | 594 | 1139067 | 1139660 | BCG_1047c | BCG_1047c | 594 | 1109398 | 1109991 | BCGDan_1034 | BCGDan_1034 | 594 | 1109514 | 1110185 | BCGDan_sapMKO_1033 | BCGDan_sapMKO_1033 | 672 |
| 1109272 | 1110192 | galU | Rv0993 | 921 | 1139761 | 1140681 | galU | BCG_1048 | 921 | 1110092 | 1111012 | galU | BCGDan_1035 | 921 | 1110208 | 1111128 | galU | BCGDan_sapMKO_1034 | 921 |
| 1110269 | 1111549 | moeA1 | Rv0994 | 1281 | 1140758 | 1142038 | moeA1 | BCG_1049 | 1281 | 1111089 | 1112369 | moeA1 | BCGDan_1036 | 1281 | 1111214 | 1112485 | moeA1 | BCGDan_sapMKO_1035 | 1272 |
| 1111612 | 1112223 | rimJ | Rv0995 | 612 | 1142101 | 1142712 | rimJ | BCG_1050 | 612 | 1112432 | 1113043 | rimJ | BCGDan_1037 | 612 | 1112548 | 1113159 | rimJ | BCGDan_sapMKO_1036 | 612 |
| 1112384 | 1113460 | Rv0996 | Rv0996 | 1077 | 1142873 | 1143949 | BCG_1051 | BCG_1051 | 1077 | 1113204 | 1114280 | BCGDan_1038 | BCGDan_1038 | 1077 | 1113320 | 1114396 | BCGDan_sapMKO_1037 | BCGDan_sapMKO_1037 | 1077 |
| NA | NA | NA | NA | NA | 1144123 | 1144353 | BCG_1052 | BCG_1052 | 231 | 1114454 | 1114684 | BCGDan_1040 | BCGDan_1040 | 231 | 1114570 | 1114800 | BCGDan_sapMKO_1039 | BCGDan_sapMKO_1039 | 231 |
| 1332980 | 1333858 | Rv1190 | Rv1190 | 879 | 1144350 | 1144601 | BCG_1053 | BCG_1053 | 252 | 1114681 | 1114932 | BCGDan_1041 | BCGDan_1041 | 252 | 1114797 | 1115048 | BCGDan_sapMKO_1040 | BCGDan_sapMKO_1040 | 252 |
| 1114293 | 1114724 | Rv0997 | Rv0997 | 432 | 1144782 | 1145213 | BCG_1054 | BCG_1054 | 432 | 1115113 | 1115544 | BCGDan_1042 | BCGDan_1042 | 432 | 1115229 | 1115660 | BCGDan_sapMKO_1041 | BCGDan_sapMKO_1041 | 432 |

| | | | | | | | | | | | | | | | | | | | |
|---|---|---|---|---|---|---|---|---|---|---|---|---|---|---|---|---|---|---|---|
| 1114748 | 1115749 | Rv0998 | Rv0998 | 1002 | 1145237 | 1146238 | BCG_1055 | BCG_1055 | 1002 | 1115568 | 1116569 | BCGDan_1043 | BCGDan_1043 | 1002 | 1115735 | 1116685 | BCGDan_sapMKO_1042 | BCGDan_sapMKO_1042 | 951 |
| 1115767 | 1116525 | Rv0999 | Rv0999 | 759 | 1146256 | 1147014 | BCG_1056 | BCG_1056 | 759 | 1116587 | 1117345 | BCGDan_1044 | BCGDan_1044 | 759 | 1116703 | 1117461 | BCGDan_sapMKO_1043 | BCGDan_sapMKO_1043 | 759 |
| 1116531 | 1117148 | Rv1000c | Rv1000c | 618 | 1147020 | 1147637 | BCG_1057c | BCG_1057c | 618 | 1117351 | 1117968 | BCGDan_1045 | BCGDan_1045 | 618 | 1117467 | 1118084 | BCGDan_sapMKO_1044 | BCGDan_sapMKO_1044 | 618 |
| 1117185 | 1118393 | arcA | Rv1001 | 1209 | 1147674 | 1148882 | arcA | BCG_1058 | 1209 | 1118005 | 1119213 | arcA | BCGDan_1046 | 1209 | 1118121 | 1119329 | arcA | BCGDan_sapMKO_1045 | 1209 |
| 1118428 | 1119939 | Rv1002c | Rv1002c | 1512 | 1148917 | 1150428 | BCG_1059c | BCG_1059c | 1512 | 1119248 | 1120759 | BCGDan_1047 | BCGDan_1047 | 1512 | 1119364 | 1120932 | BCGDan_sapMKO_1046 | BCGDan_sapMKO_1046 | 1569 |
| 1120022 | 1120879 | Rv1003 | Rv1003 | 858 | 1150511 | 1151368 | BCG_1060 | BCG_1060 | 858 | 1120842 | 1121699 | BCGDan_1048 | BCGDan_1048 | 858 | 1120958 | 1121815 | BCGDan_sapMKO_1047 | BCGDan_sapMKO_1047 | 858 |
| 1120889 | 1122148 | Rv1004c | Rv1004c | 1260 | 1151378 | 1152637 | BCG_1061c | BCG_1061c | 1260 | 1121709 | 1122968 | BCGDan_1049 | BCGDan_1049 | 1260 | 1121825 | 1123048 | BCGDan_sapMKO_1048 | BCGDan_sapMKO_1048 | 1224 |
| 1122222 | 1123598 | pabB | Rv1005c | 1377 | 1152711 | 1154147 | pabB | BCG_1062c | 1437 | 1123042 | 1124478 | pabB | BCGDan_1050 | 1437 | 1123158 | 1124402 | pabB | BCGDan_sapMKO_1049 | 1245 |
| 1123714 | 1125417 | Rv1006 | Rv1006 | 1704 | 1154203 | 1155906 | BCG_1063 | BCG_1063 | 1704 | 1124534 | 1126237 | BCGDan_1051 | BCGDan_1051 | 1704 | 1124689 | 1126353 | BCGDan_sapMKO_1050 | BCGDan_sapMKO_1050 | 1665 |
| 1125444 | 1127003 | metS | Rv1007c | 1560 | 1155933 | 1157492 | metS | BCG_1064c | 1560 | 1126264 | 1127823 | metS | BCGDan_1052 | 1560 | 1126380 | 1127939 | metS | BCGDan_sapMKO_1051 | 1560 |
| 1127089 | 1127883 | tatD | Rv1008 | 795 | 1157578 | 1158372 | tatD | BCG_1065 | 795 | 1127909 | 1128703 | tatD | BCGDan_1053 | 795 | 1127965 | 1128819 | tatD | BCGDan_sapMKO_1052 | 855 |
| 1128091 | 1129179 | rpfB | Rv1009 | 1089 | 1158580 | 1159668 | rpfB | BCG_1066 | 1089 | 1128911 | 1129999 | rpfB | BCGDan_1054 | 1089 | 1129027 | 1130115 | rpfB | BCGDan_sapMKO_1053 | 1089 |
| 1129152 | 1130105 | ksgA | Rv1010 | 954 | 1159641 | 1160594 | ksgA | BCG_1067 | 954 | 1129972 | 1130925 | ksgA | BCGDan_1055 | 954 | 1130088 | 1131041 | ksgA | BCGDan_sapMKO_1054 | 954 |
| 1130191 | 1131111 | ispE | Rv1011 | 921 | 1160680 | 1161600 | ispE | BCG_1068 | 921 | 1131011 | 1131931 | ispE | BCGDan_1056 | 921 | 1131091 | 1132047 | ispE | BCGDan_sapMKO_1055 | 957 |
| 1131128 | 1131421 | Rv1012 | Rv1012 | 294 | 1161597 | 1161911 | BCG_1069 | BCG_1069 | 315 | 1131928 | 1132242 | BCGDan_1057 | BCGDan_1057 | 315 | 1132044 | 1132358 | BCGDan_sapMKO_1056 | BCGDan_sapMKO_1056 | 315 |
| 1131625 | 1133259 | pks16 | Rv1013 | 1635 | 1162115 | 1163749 | pks16 | BCG_1070 | 1635 | 1132446 | 1134080 | pks16 | BCGDan_1058 | 1635 | 1132562 | 1134196 | pks16 | BCGDan_sapMKO_1057 | 1635 |
| 1133333 | 1133908 | pth | Rv1014c | 576 | 1163823 | 1164398 | pth | BCG_1071c | 576 | 1134154 | 1134729 | pth | BCGDan_1059 | 576 | 1134270 | 1134845 | pth | BCGDan_sapMKO_1058 | 576 |
| 1133921 | 1134568 | rplY | Rv1015c | 648 | 1164411 | 1165058 | rplY | BCG_1072c | 648 | 1134742 | 1135389 | rplY | BCGDan_1060 | 648 | 1134858 | 1135505 | rplY | BCGDan_sapMKO_1059 | 648 |
| 1134785 | 1135465 | lpqT | Rv1016c | 681 | 1165192 | 1165950 | lpqT | BCG_1073c | 759 | 1135523 | 1136281 | lpqT | BCGDan_1061 | 759 | 1135639 | 1136397 | lpqT | BCGDan_sapMKO_1060 | 759 |
| 1135501 | 1136481 | prsA | Rv1017c | 981 | 1165986 | 1166966 | prsA | BCG_1074c | 981 | 1136317 | 1137297 | prsA | BCGDan_1062 | 981 | 1136433 | 1137413 | prsA | BCGDan_sapMKO_1061 | 981 |
| 1136573 | 1138060 | glmU | Rv1018c | 1488 | 1167058 | 1168545 | glmU | BCG_1075c | 1488 | 1137389 | 1138876 | glmU | BCGDan_1063 | 1488 | 1137505 | 1138992 | glmU | BCGDan_sapMKO_1062 | 1488 |
| 1138315 | 1138908 | Rv1019 | Rv1019 | 594 | 1168800 | 1169393 | BCG_1076 | BCG_1076 | 594 | 1139131 | 1139724 | BCGDan_1065 | BCGDan_1065 | 594 | 1139247 | 1139840 | BCGDan_sapMKO_1064 | BCGDan_sapMKO_1064 | 594 |
| 1138967 | 1142671 | mfd | Rv1020 | 3705 | 1169452 | 1173156 | mfd | BCG_1077 | 3705 | 1139783 | 1143487 | mfd | BCGDan_1066 | 3705 | 1139899 | 1143603 | mfd | BCGDan_sapMKO_1065 | 3705 |
| 1142671 | 1143648 | Rv1021 | Rv1021 | 978 | 1173156 | 1174133 | BCG_1078 | BCG_1078 | 978 | 1143487 | 1144464 | BCGDan_1067 | BCGDan_1067 | 978 | 1143603 | 1144580 | BCGDan_sapMKO_1066 | BCGDan_sapMKO_1066 | 978 |
| 1143736 | 1144467 | lpqU | Rv1022 | 732 | 1174221 | 1174952 | lpqU | BCG_1079 | 732 | 1144552 | 1145283 | lpqU | BCGDan_1068 | 732 | 1144719 | 1145399 | lpqU | BCGDan_sapMKO_1067 | 681 |
| 1144564 | 1145853 | eno | Rv1023 | 1290 | 1175049 | 1176338 | eno | BCG_1080 | 1290 | 1145380 | 1146669 | eno | BCGDan_1069 | 1290 | 1145496 | 1146785 | eno | BCGDan_sapMKO_1068 | 1290 |
| 1145858 | 1146544 | Rv1024 | Rv1024 | 687 | 1176343 | 1177029 | BCG_1081 | BCG_1081 | 687 | 1146674 | 1147360 | BCGDan_1070 | BCGDan_1070 | 687 | 1146790 | 1147476 | BCGDan_sapMKO_1069 | BCGDan_sapMKO_1069 | 687 |
| 1146561 | 1147028 | Rv1025 | Rv1025 | 468 | 1177046 | 1177513 | BCG_1082 | BCG_1082 | 468 | 1147377 | 1147844 | BCGDan_1071 | BCGDan_1071 | 468 | 1147469 | 1147960 | BCGDan_sapMKO_1070 | BCGDan_sapMKO_1070 | 492 |
| 1147019 | 1147978 | Rv1026 | Rv1026 | 960 | 1177504 | 1178463 | BCG_1083 | BCG_1083 | 960 | 1147835 | 1148794 | BCGDan_1072 | BCGDan_1072 | 960 | 1147951 | 1148910 | BCGDan_sapMKO_1071 | BCGDan_sapMKO_1071 | 960 |
| 1148427 | 1149107 | kdpE | Rv1027c | 681 | 1178912 | 1179592 | kdpE | BCG_1084c | 681 | 1149243 | 1149923 | kdpE | BCGDan_1073 | 681 | 1149359 | 1150039 | kdpE | BCGDan_sapMKO_1072 | 681 |
| 1149104 | 1151686 | kdpD | Rv1028c | 2583 | 1179589 | 1182171 | kdpD | BCG_1085c | 2583 | 1149920 | 1152502 | kdpD | BCGDan_1074 | 2583 | 1150036 | 1152618 | kdpD | BCGDan_sapMKO_1073 | 2583 |
| 1151920 | 1152012 | kdpF | Rv1028A | 93 | 1182405 | 1182497 | kdpF | BCG_1086 | 93 | 1152730 | 1152828 | kdpF | BCGDan_1075 | 99 | 1152846 | 1152944 | kdpF | BCGDan_sapMKO_1074 | 99 |
| 1152012 | 1153727 | kdpA | Rv1029 | 1716 | 1182497 | 1184212 | kdpA | BCG_1087 | 1716 | 1152828 | 1154543 | kdpA | BCGDan_1076 | 1716 | 1152944 | 1154659 | kdpA | BCGDan_sapMKO_1075 | 1716 |
| 1153724 | 1155853 | kdpB | Rv1030 | 2130 | 1184209 | 1186338 | kdpB | BCG_1088 | 2130 | 1154540 | 1156669 | kdpB | BCGDan_1077 | 2130 | 1154656 | 1156785 | kdpB | BCGDan_sapMKO_1076 | 2130 |
| 1155853 | 1156422 | kdpC | Rv1031 | 570 | 1186338 | 1186907 | kdpC | BCG_1089 | 570 | 1156669 | 1157238 | kdpC | BCGDan_1078 | 570 | 1156785 | 1157354 | kdpC | BCGDan_sapMKO_1077 | 570 |
| 1156426 | 1157955 | trcS | Rv1032c | 1530 | 1186911 | 1188440 | trcS | BCG_1090c | 1530 | 1157242 | 1158771 | trcS | BCGDan_1079 | 1530 | 1157358 | 1158818 | trcS | BCGDan_sapMKO_1078 | 1461 |
| 1157963 | 1158736 | trcR | Rv1033c | 774 | 1188448 | 1189221 | trcR | BCG_1091c | 774 | 1158779 | 1159552 | trcR | BCGDan_1080 | 774 | 1158895 | 1159659 | trcR | BCGDan_sapMKO_1079 | 765 |
| 1158918 | 1159307 | Rv1034c | Rv1034c | 390 | 1189403 | 1189792 | BCG_1092c | BCG_1092c | 390 | 1159734 | 1160123 | BCGDan_1081 | BCGDan_1081 | 390 | 1159850 | 1160239 | BCGDan_sapMKO_1080 | BCGDan_sapMKO_1080 | 390 |
| 1159375 | 1160061 | Rv1035c | Rv1035c | 687 | 1189860 | 1190546 | BCG_1093c | BCG_1093c | 687 | 1160191 | 1160877 | BCGDan_1082 | BCGDan_1082 | 687 | 1160307 | 1160972 | BCGDan_sapMKO_1081 | BCGDan_sapMKO_1081 | 666 |
| 1160095 | 1160433 | Rv1036c | Rv1036c | 339 | 1190580 | 1190918 | BCG_1094c | BCG_1094c | 339 | 1160911 | 1161249 | BCGDan_1083 | BCGDan_1083 | 339 | 1161027 | 1161350 | BCGDan_sapMKO_1082 | BCGDan_sapMKO_1082 | 324 |
| 1160544 | 1160828 | esxI | Rv1037c | 285 | 1191029 | 1191313 | esxI | BCG_1095c | 285 | 1161360 | 1161644 | esxI | BCGDan_1084 | 285 | 1161476 | 1161760 | esxI | BCGDan_sapMKO_1083 | 285 |
| 1160855 | 1161151 | esxJ | Rv1038c | 297 | 1191340 | 1191636 | esxJ | BCG_1096c | 297 | 1161671 | 1161967 | esxJ | BCGDan_1085 | 297 | 1161787 | 1162083 | esxJ | BCGDan_sapMKO_1084 | 297 |
| 1161297 | 1162472 | PPE15 | Rv1039c | 1176 | 1191782 | 1192957 | PPE15 | BCG_1097c | 1176 | 1162113 | 1163288 | PPE15 | BCGDan_1086 | 1176 | 1162229 | 1163362 | PPE15 | BCGDan_sapMKO_1085 | 1134 |
| 1162549 | 1163376 | PE8 | Rv1040c | 828 | 1193034 | 1193861 | PE8 | BCG_1098c | 828 | 1163365 | 1164192 | PE8 | BCGDan_1087 | 828 | 1163481 | 1164308 | PE8 | BCGDan_sapMKO_1086 | 828 |
| 1164572 | 1165435 | Rv1041c | Rv1041c | 864 | 1195057 | 1195608 | BCG_1099c | BCG_1099c | 552 | 1165388 | 1165939 | BCGDan_1088 | BCGDan_1088 | 552 | 1165507 | 1166055 | BCGDan_sapMKO_1087 | BCGDan_sapMKO_1087 | 549 |
| 1165092 | 1165499 | Rv1042c | Rv1042c | 408 | 1195577 | 1195984 | BCG_1100c | BCG_1100c | 408 | 1165908 | 1166315 | BCGDan_1089 | BCGDan_1089 | 408 | 1166024 | 1166422 | BCGDan_sapMKO_1088 | BCGDan_sapMKO_1088 | 399 |
| 1165781 | 1166806 | Rv1043c | Rv1043c | 1026 | 1196267 | 1197292 | BCG_1101c | BCG_1101c | 1026 | 1166598 | 1167623 | BCGDan_1090 | BCGDan_1090 | 1026 | 1166714 | 1167751 | BCGDan_sapMKO_1089 | BCGDan_sapMKO_1089 | 1038 |
| 1167053 | 1167676 | Rv1044 | Rv1044 | 624 | 1197539 | 1198162 | BCG_1102 | BCG_1102 | 624 | 1167870 | 1168493 | BCGDan_1091 | BCGDan_1091 | 624 | 1168025 | 1168609 | BCGDan_sapMKO_1090 | BCGDan_sapMKO_1090 | 585 |
| 1167673 | 1168554 | Rv1045 | Rv1045 | 882 | 1198159 | 1199040 | BCG_1103 | BCG_1103 | 882 | 1168490 | 1169371 | BCGDan_1092 | BCGDan_1092 | 882 | 1168606 | 1169487 | BCGDan_sapMKO_1091 | BCGDan_sapMKO_1091 | 882 |
| 1168704 | 1169228 | Rv1046c | Rv1046c | 525 | 1199122 | 1199715 | BCG_1104c | BCG_1104c | 594 | 1169453 | 1170046 | BCGDan_1093 | BCGDan_1093 | 594 | 1169569 | 1170162 | BCGDan_sapMKO_1092 | BCGDan_sapMKO_1092 | 594 |
| 1169423 | 1170670 | Rv1047 | Rv1047 | 1248 | 1199910 | 1201157 | BCG_1105 | BCG_1105 | 1248 | 1170241 | 1171488 | BCGDan_1094 | BCGDan_1094 | 1248 | 1170294 | 1171604 | BCGDan_sapMKO_1093 | BCGDan_sapMKO_1093 | 1311 |
| 1171038 | 1172153 | Rv1048c | Rv1048c | 1116 | 1201525 | 1202640 | BCG_1106c | BCG_1106c | 1116 | 1171856 | 1172971 | BCGDan_1095 | BCGDan_1095 | 1116 | 1171972 | 1172988 | BCGDan_sapMKO_1094 | BCGDan_sapMKO_1094 | 1017 |
| 1172386 | 1172832 | Rv1049 | Rv1049 | 447 | 1202873 | 1203319 | BCG_1107 | BCG_1107 | 447 | 1173204 | 1173650 | BCGDan_1096 | BCGDan_1096 | 447 | 1173320 | 1173766 | BCGDan_sapMKO_1095 | BCGDan_sapMKO_1095 | 447 |
| 1172881 | 1173786 | Rv1050 | Rv1050 | 906 | 1203367 | 1204272 | BCG_1108 | BCG_1108 | 906 | 1173698 | 1174603 | BCGDan_1097 | BCGDan_1097 | 906 | 1173841 | 1174719 | BCGDan_sapMKO_1096 | BCGDan_sapMKO_1096 | 879 |
| 1173945 | 1174700 | Rv1051c | Rv1051c | 756 | 1204431 | 1205186 | BCG_1109c | BCG_1109c | 756 | 1174762 | 1175517 | BCGDan_1098 | BCGDan_1098 | 756 | 1174878 | 1175633 | BCGDan_sapMKO_1097 | BCGDan_sapMKO_1097 | 756 |
| 1175723 | 1176112 | Rv1052 | Rv1052 | 390 | 1206209 | 1206598 | BCG_1110 | BCG_1110 | 390 | 1176540 | 1176929 | BCGDan_1099 | BCGDan_1099 | 390 | 1176656 | 1177045 | BCGDan_sapMKO_1098 | BCGDan_sapMKO_1098 | 390 |
| 1176011 | 1176286 | Rv1053c | Rv1053c | 276 | 1206497 | 1206772 | BCG_1111c | BCG_1111c | 276 | 1176828 | 1177103 | BCGDan_1100 | BCGDan_1100 | 276 | 1176944 | 1177219 | BCGDan_sapMKO_1099 | BCGDan_sapMKO_1099 | 276 |
| 1176928 | 1177242 | Rv1054 | Rv1054 | 315 | 1207414 | 1207728 | BCG_1112 | BCG_1112 | 315 | 1177745 | 1178059 | BCGDan_1101 | BCGDan_1101 | 315 | 1177861 | 1178175 | BCGDan_sapMKO_1100 | BCGDan_sapMKO_1100 | 315 |
| 1177239 | 1177373 | Rv1055 | Rv1055 | 135 | 1207725 | 1207859 | BCG_1113 | BCG_1113 | 135 | 1178056 | 1178190 | BCGDan_1102 | BCGDan_1102 | 135 | 1178172 | 1178306 | BCGDan_sapMKO_1101 | BCGDan_sapMKO_1101 | 135 |
| 1177628 | 1178392 | Rv1056 | Rv1056 | 765 | 1208114 | 1208878 | BCG_1114 | BCG_1114 | 765 | 1178445 | 1179209 | BCGDan_1104 | BCGDan_1104 | 765 | 1178645 | 1179325 | BCGDan_sapMKO_1103 | BCGDan_sapMKO_1103 | 681 |
| 1179396 | 1180577 | Rv1057 | Rv1057 | 1182 | 1209882 | 1211063 | BCG_1115 | BCG_1115 | 1182 | 1180213 | 1181394 | BCGDan_1105 | BCGDan_1105 | 1182 | 1180338 | 1181510 | BCGDan_sapMKO_1104 | BCGDan_sapMKO_1104 | 1173 |
| 1180684 | 1182315 | fadD14 | Rv1058 | 1632 | 1211170 | 1212801 | fadD14 | BCG_1116 | 1632 | 1181501 | 1183132 | fadD14 | BCGDan_1106 | 1632 | 1181629 | 1183248 | fadD14 | BCGDan_sapMKO_1105 | 1620 |
| 1182391 | 1183455 | Rv1059 | Rv1059 | 1065 | 1212877 | 1213941 | BCG_1117 | BCG_1117 | 1065 | 1183208 | 1184272 | BCGDan_1107 | BCGDan_1107 | 1065 | 1183330 | 1184388 | BCGDan_sapMKO_1106 | BCGDan_sapMKO_1106 | 1059 |
| 1183508 | 1183981 | Rv1060 | Rv1060 | 474 | 1213994 | 1214467 | BCG_1118 | BCG_1118 | 474 | 1184325 | 1184798 | BCGDan_1108 | BCGDan_1108 | 474 | 1184441 | 1184914 | BCGDan_sapMKO_1107 | BCGDan_sapMKO_1107 | 474 |
| 1184015 | 1184878 | Rv1061 | Rv1061 | 864 | 1214501 | 1215364 | BCG_1119 | BCG_1119 | 864 | 1184832 | 1185695 | BCGDan_1109 | BCGDan_1109 | 864 | 1184948 | 1185811 | BCGDan_sapMKO_1108 | BCGDan_sapMKO_1108 | 864 |
| 1184883 | 1185740 | Rv1062 | Rv1062 | 858 | 1215369 | 1216226 | BCG_1120 | BCG_1120 | 858 | 1185700 | 1186557 | BCGDan_1110 | BCGDan_1110 | 858 | 1185840 | 1186673 | BCGDan_sapMKO_1109 | BCGDan_sapMKO_1109 | 834 |
| 1185741 | 1186823 | Rv1063c | Rv1063c | 1083 | 1216227 | 1217309 | BCG_1121c | BCG_1121c | 1083 | 1186558 | 1187640 | BCGDan_1111 | BCGDan_1111 | 1083 | 1186674 | 1187732 | BCGDan_sapMKO_1110 | BCGDan_sapMKO_1110 | 1059 |
| 1186904 | 1187323 | lpqV | Rv1064c | 420 | 1217390 | 1217809 | lpqV | BCG_1122c | 420 | 1187721 | 1188140 | lpqV | BCGDan_1112 | 420 | 1187837 | 1188220 | lpqV | BCGDan_sapMKO_1111 | 384 |
| 1187435 | 1188001 | Rv1065 | Rv1065 | 567 | 1217921 | 1218487 | BCG_1123 | BCG_1123 | 567 | 1188252 | 1188818 | BCGDan_1113 | BCGDan_1113 | 567 | 1188374 | 1188934 | BCGDan_sapMKO_1112 | BCGDan_sapMKO_1112 | 561 |
| 1187998 | 1188393 | Rv1066 | Rv1066 | 396 | 1218484 | 1218879 | BCG_1124 | BCG_1124 | 396 | 1188815 | 1189210 | BCGDan_1114 | BCGDan_1114 | 396 | 1188931 | 1189326 | BCGDan_sapMKO_1113 | BCGDan_sapMKO_1113 | 396 |
| 1188421 | 1190424 | PE_PGRS19 | Rv1067c | 2004 | 1218907 | 1220922 | PE_PGRS19 | BCG_1125c | 2016 | 1189238 | 1191253 | PE_PGRS19 | BCGDan_1115 | 2016 | 1189354 | 1191369 | PE_PGRS19 | BCGDan_sapMKO_1114 | 2016 |
| 1190757 | 1192148 | PE_PGRS20 | Rv1068c | 1392 | 1221255 | 1223447 | PE_PGRS20 | BCG_1126c | 2193 | 1191586 | 1193778 | PE_PGRS20 | BCGDan_1116 | 2193 | 1191702 | 1193912 | PE_PGRS20 | BCGDan_sapMKO_1115 | 2211 |
| 1192510 | 1194273 | Rv1069c | Rv1069c | 1764 | 1223809 | 1225572 | BCG_1127c | BCG_1127c | 1764 | 1194140 | 1195903 | BCGDan_1117 | BCGDan_1117 | 1764 | 1194256 | 1196019 | BCGDan_sapMKO_1116 | BCGDan_sapMKO_1116 | 1764 |
| 1194270 | 1195043 | echA8 | Rv1070c | 774 | 1225569 | 1226342 | echA8 | BCG_1128c | 774 | 1195900 | 1196673 | echA8 | BCGDan_1118 | 774 | 1196016 | 1196789 | echA8 | BCGDan_sapMKO_1117 | 774 |
| 1195055 | 1196092 | echA9 | Rv1071c | 1038 | 1226354 | 1227391 | echA9 | BCG_1129c | 1038 | 1196685 | 1197722 | echA9 | BCGDan_1119 | 1038 | 1196801 | 1197838 | echA9 | BCGDan_sapMKO_1118 | 1038 |
| 1196279 | 1197115 | Rv1072 | Rv1072 | 837 | 1227578 | 1228414 | BCG_1130 | BCG_1130 | 837 | 1197909 | 1198745 | BCGDan_1120 | BCGDan_1120 | 837 | 1198025 | 1198861 | BCGDan_sapMKO_1119 | BCGDan_sapMKO_1119 | 837 |
| 1197231 | 1198082 | Rv1073 | Rv1073 | 852 | 1228530 | 1229381 | BCG_1131 | BCG_1131 | 852 | 1198861 | 1199712 | BCGDan_1121 | BCGDan_1121 | 852 | 1198968 | 1199828 | BCGDan_sapMKO_1120 | BCGDan_sapMKO_1120 | 861 |
| 1198156 | 1199373 | fadA3 | Rv1074c | 1218 | 1229455 | 1230672 | fadA3 | BCG_1132c | 1218 | 1199786 | 1201018 | fadA3 | BCGDan_1122 | 1233 | 1199902 | 1201134 | fadA3 | BCGDan_sapMKO_1121 | 1233 |
| 1199426 | 1200370 | Rv1075c | Rv1075c | 945 | 1230725 | 1231669 | BCG_1133c | BCG_1133c | 945 | 1201056 | 1202000 | BCGDan_1123 | BCGDan_1123 | 945 | 1201172 | 1202122 | BCGDan_sapMKO_1122 | BCGDan_sapMKO_1122 | 951 |
| 1200767 | 1201660 | lipU | Rv1076 | 894 | 1232066 | 1232959 | lipU | BCG_1134 | 894 | 1202397 | 1203290 | lipU | BCGDan_1124 | 894 | 1202396 | 1203406 | lipU | BCGDan_sapMKO_1123 | 1011 |
| 1201717 | 1203111 | cbs | Rv1077 | 1395 | 1233016 | 1234410 | cbs | BCG_1135 | 1395 | 1203347 | 1204741 | cbs | BCGDan_1125 | 1395 | 1203463 | 1204857 | cbs | BCGDan_sapMKO_1124 | 1395 |
| 1203313 | 1204035 | pra | Rv1078 | 723 | 1234612 | 1235334 | pra | BCG_1136 | 723 | 1204943 | 1205665 | pra | BCGDan_1126 | 723 | 1205059 | 1205781 | pra | BCGDan_sapMKO_1125 | 723 |
| 1204067 | 1205233 | metB | Rv1079 | 1167 | 1235366 | 1236532 | metB | BCG_1137 | 1167 | 1205697 | 1206863 | metB | BCGDan_1127 | 1167 | 1205813 | 1206979 | metB | BCGDan_sapMKO_1126 | 1167 |
| 1205304 | 1205798 | greA | Rv1080c | 495 | 1236603 | 1237097 | greA | BCG_1138c | 495 | 1206934 | 1207428 | greA | BCGDan_1128 | 495 | 1207050 | 1207544 | greA | BCGDan_sapMKO_1127 | 495 |
| 1205984 | 1206418 | Rv1081c | Rv1081c | 435 | 1237283 | 1237717 | BCG_1139c | BCG_1139c | 435 | 1207614 | 1208048 | BCGDan_1129 | BCGDan_1129 | 435 | 1207730 | 1208164 | BCGDan_sapMKO_1128 | BCGDan_sapMKO_1128 | 435 |
| 1206520 | 1207386 | mca | Rv1082 | 867 | 1237819 | 1238685 | mca | BCG_1140 | 867 | 1208150 | 1209016 | mca | BCGDan_1130 | 867 | 1208266 | 1209132 | mca | BCGDan_sapMKO_1129 | 867 |

| | | | | | | | | | | | | | | | | | | | |
|---|---|---|---|---|---|---|---|---|---|---|---|---|---|---|---|---|---|---|---|
| 1207383 | 1207649 | Rv1083 | Rv1083 | 267 | 1238682 | 1238948 | BCG_1141 | BCG_1141 | 267 | 1209013 | 1209279 | BCGDan_1131 | BCGDan_1131 | 267 | 1209129 | 1209395 | BCGDan_sapMKO_1130 | BCGDan_sapMKO_1130 | 267 |
| 1207636 | 1209657 | Rv1084 | Rv1084 | 2022 | 1238935 | 1240956 | BCG_1142 | BCG_1142 | 2022 | 1209266 | 1211287 | BCGDan_1132 | BCGDan_1132 | 2022 | 1209382 | 1211403 | BCGDan_sapMKO_1131 | BCGDan_sapMKO_1131 | 2022 |
| 1209756 | 1210484 | Rv1085c | Rv1085c | 729 | 1241055 | 1241783 | BCG_1143c | BCG_1143c | 729 | 1211386 | 1212114 | BCGDan_1133 | BCGDan_1133 | 729 | 1211502 | 1212278 | BCGDan_sapMKO_1132 | BCGDan_sapMKO_1132 | 777 |
| 1210595 | 1211383 | Rv1086 | Rv1086 | 789 | 1241894 | 1242682 | BCG_1144 | BCG_1144 | 789 | 1212225 | 1213013 | BCGDan_1134 | BCGDan_1134 | 789 | 1212341 | 1213129 | BCGDan_sapMKO_1133 | BCGDan_sapMKO_1133 | 789 |
| 1211560 | 1213863 | PE_PGRS21 | Rv1087 | 2304 | 1242859 | 1245288 | PE_PGRS21 | BCG_1145 | 2430 | 1213190 | 1215619 | PE_PGRS21 | BCGDan_1135 | 2430 | 1213306 | 1215735 | PE_PGRS21 | BCGDan_sapMKO_1134 | 2430 |
| 1214040 | 1214360 | Rv1087A | Rv1087A | 321 | 1245465 | 1245785 | BCG_1146 | BCG_1146 | 321 | 1215796 | 1216116 | BCGDan_1136 | BCGDan_1136 | 321 | 1215912 | 1216232 | BCGDan_sapMKO_1135 | BCGDan_sapMKO_1135 | 321 |
| 1214513 | 1214947 | PE9 | Rv1088 | 435 | 1245938 | 1246372 | PE9 | BCG_1147 | 435 | 1216269 | 1216703 | PE9 | BCGDan_1137 | 435 | 1216385 | 1216819 | PE9 | BCGDan_sapMKO_1136 | 435 |
| 1214769 | 1215131 | PE10 | Rv1089 | 363 | 1246251 | 1246556 | PE10 | BCG_1148 | 306 | 1216582 | 1216887 | PE10 | BCGDan_1138 | 306 | 1216698 | 1217003 | PE10 | BCGDan_sapMKO_1137 | 306 |
| 1215517 | 1215621 | celA2a | Rv1089A | 105 | 1246942 | 1247046 | celA2a | BCG_1149 | 105 | 1217273 | 1217377 | celA2a | BCGDan_1139 | 105 | 1217389 | 1217493 | celA2a | BCGDan_sapMKO_1138 | 105 |
| 1215599 | 1216054 | celA2b | Rv1090 | 456 | 1247024 | 1247479 | celA2b | BCG_1150 | 456 | 1217355 | 1217810 | celA2b | BCGDan_1140 | 456 | 1217471 | 1217926 | celA2b | BCGDan_sapMKO_1139 | 456 |
| 1216469 | 1219030 | PE_PGRS22 | Rv1091 | 2562 | 1247894 | 1250446 | PE_PGRS22 | BCG_1151 | 2553 | 1218225 | 1220777 | PE_PGRS22 | BCGDan_1141 | 2553 | 1218341 | 1220893 | PE_PGRS22 | BCGDan_sapMKO_1140 | 2553 |
| 1219248 | 1220186 | coaA | Rv1092c | 939 | 1250664 | 1251602 | coaA | BCG_1152c | 939 | 1220995 | 1221867 | coaA | BCGDan_1142 | 873 | 1221111 | 1221983 | coaA | BCGDan_sapMKO_1141 | 873 |
| 1220574 | 1221854 | glyA1 | Rv1093 | 1281 | 1251990 | 1253270 | glyA1 | BCG_1153 | 1281 | 1222321 | 1223601 | glyA1 | BCGDan_1143 | 1281 | 1222437 | 1223717 | glyA1 | BCGDan_sapMKO_1142 | 1281 |
| 1221959 | 1222786 | desA2 | Rv1094 | 828 | 1253375 | 1254202 | desA2 | BCG_1154 | 828 | 1223706 | 1224533 | desA2 | BCGDan_1144 | 828 | 1223822 | 1224649 | desA2 | BCGDan_sapMKO_1143 | 828 |
| 1222997 | 1224298 | phoH2 | Rv1095 | 1302 | 1254413 | 1255714 | phoH2 | BCG_1155 | 1302 | 1224744 | 1226045 | phoH2 | BCGDan_1145 | 1302 | 1224860 | 1226161 | phoH2 | BCGDan_sapMKO_1144 | 1302 |
| 1224385 | 1225260 | Rv1096 | Rv1096 | 876 | 1255801 | 1256676 | BCG_1156 | BCG_1156 | 876 | 1226132 | 1227007 | BCGDan_1146 | BCGDan_1146 | 876 | 1226248 | 1227123 | BCGDan_sapMKO_1145 | BCGDan_sapMKO_1145 | 876 |
| 1225263 | 1226144 | Rv1097c | Rv1097c | 882 | 1256679 | 1257560 | BCG_1157c | BCG_1157c | 882 | 1227010 | 1227891 | BCGDan_1147 | BCGDan_1147 | 882 | 1227126 | 1228007 | BCGDan_sapMKO_1146 | BCGDan_sapMKO_1146 | 882 |
| 1226141 | 1227565 | fum | Rv1098c | 1425 | 1257557 | 1258981 | fum | BCG_1158c | 1425 | 1227888 | 1229312 | fum | BCGDan_1148 | 1425 | 1228004 | 1229428 | fum | BCGDan_sapMKO_1147 | 1425 |
| 1227596 | 1228684 | glpX | Rv1099c | 1089 | 1259012 | 1259998 | BCG_1159c | BCG_1159c | 987 | 1229343 | 1230431 | glpX | BCGDan_1149 | 1089 | 1229459 | 1230547 | BCGDan_sapMKO_1148 | BCGDan_sapMKO_1148 | 1089 |
| 1228683 | 1229384 | Rv1100 | Rv1100 | 702 | 1260099 | 1260800 | BCG_1160 | BCG_1160 | 702 | 1230430 | 1231131 | BCGDan_1150 | BCGDan_1150 | 702 | 1230648 | 1231247 | BCGDan_sapMKO_1149 | BCGDan_sapMKO_1149 | 600 |
| 1229391 | 1230548 | Rv1101c | Rv1101c | 1158 | 1260807 | 1261835 | BCG_1161c | BCG_1161c | 1029 | 1231138 | 1232166 | BCGDan_1151 | BCGDan_1151 | 1029 | 1231254 | 1232282 | BCGDan_sapMKO_1150 | BCGDan_sapMKO_1150 | 1029 |
| 1230660 | 1230971 | mazF3 | Rv1102c | 312 | 1262075 | 1262386 | BCG_1162c | BCG_1162c | 312 | 1232406 | 1232717 | BCGDan_1152 | BCGDan_1152 | 312 | 1232522 | 1232833 | BCGDan_sapMKO_1151 | BCGDan_sapMKO_1151 | 312 |
| 1230971 | 1231291 | mazE3 | Rv1103c | 321 | 1262386 | 1262706 | BCG_1163c | BCG_1163c | 321 | 1232717 | 1233037 | BCGDan_1153 | BCGDan_1153 | 321 | 1232833 | 1233153 | BCGDan_sapMKO_1152 | BCGDan_sapMKO_1152 | 321 |
| 1231301 | 1231990 | Rv1104 | Rv1104 | 690 | 1262716 | 1263405 | BCG_1164 | BCG_1164 | 690 | 1233047 | 1233736 | BCGDan_1154 | BCGDan_1154 | 690 | 1233163 | 1233852 | BCGDan_sapMKO_1153 | BCGDan_sapMKO_1153 | 690 |
| 1232311 | 1232826 | Rv1105 | Rv1105 | 516 | 1263726 | 1264241 | BCG_1165 | BCG_1165 | 516 | 1234057 | 1234572 | BCGDan_1155 | BCGDan_1155 | 516 | 1234173 | 1234688 | BCGDan_sapMKO_1154 | BCGDan_sapMKO_1154 | 516 |
| 1232844 | 1233956 | Rv1106c | Rv1106c | 1113 | 1264259 | 1265371 | BCG_1166c | BCG_1166c | 1113 | 1234590 | 1235702 | BCGDan_1156 | BCGDan_1156 | 1113 | 1234706 | 1235806 | BCGDan_sapMKO_1155 | BCGDan_sapMKO_1155 | 1101 |
| 1233966 | 1234223 | xseB | Rv1107c | 258 | 1265381 | 1265638 | xseB | BCG_1167c | 258 | 1235712 | 1235969 | xseB | BCGDan_1157 | 258 | 1235828 | 1236085 | xseB | BCGDan_sapMKO_1156 | 258 |
| 1234213 | 1235460 | xseA | Rv1108c | 1248 | 1265628 | 1266875 | xseA | BCG_1168c | 1248 | 1235959 | 1237206 | xseA | BCGDan_1158 | 1248 | 1236075 | 1237322 | xseA | BCGDan_sapMKO_1157 | 1248 |
| 1235457 | 1236095 | Rv1109c | Rv1109c | 639 | 1266872 | 1267510 | BCG_1169c | BCG_1169c | 639 | 1237203 | 1237841 | BCGDan_1159 | BCGDan_1159 | 639 | 1237319 | 1237891 | BCGDan_sapMKO_1158 | BCGDan_sapMKO_1158 | 573 |
| 1236185 | 1237192 | lytB2 | Rv1110 | 1008 | 1267600 | 1268607 | lytB2 | BCG_1170 | 1008 | 1237931 | 1238938 | lytB2 | BCGDan_1160 | 1008 | 1238047 | 1239054 | lytB2 | BCGDan_sapMKO_1159 | 1008 |
| 1237209 | 1238192 | Rv1111c | Rv1111c | 984 | 1268624 | 1269607 | BCG_1171c | BCG_1171c | 984 | 1238955 | 1239938 | BCGDan_1161 | BCGDan_1161 | 984 | 1239071 | 1239985 | BCGDan_sapMKO_1160 | BCGDan_sapMKO_1160 | 915 |
| 1238255 | 1239328 | Rv1112 | Rv1112 | 1074 | 1269670 | 1270743 | BCG_1172 | BCG_1172 | 1074 | 1240001 | 1241074 | BCGDan_1162 | BCGDan_1162 | 1074 | 1240117 | 1241190 | BCGDan_sapMKO_1161 | BCGDan_sapMKO_1161 | 1074 |
| 1239416 | 1239613 | vapB32 | Rv1113 | 198 | 1270831 | 1271028 | BCG_1173 | BCG_1173 | 198 | 1241162 | 1241359 | BCGDan_1163 | BCGDan_1163 | 198 | 1241278 | 1241475 | BCGDan_sapMKO_1162 | BCGDan_sapMKO_1162 | 198 |
| 1239610 | 1239984 | vapC32 | Rv1114 | 375 | 1271025 | 1271399 | BCG_1174 | BCG_1174 | 375 | 1241356 | 1241730 | BCGDan_1164 | BCGDan_1164 | 375 | 1241472 | 1241846 | BCGDan_sapMKO_1163 | BCGDan_sapMKO_1163 | 375 |
| 1240187 | 1240885 | Rv1115 | Rv1115 | 699 | 1271602 | 1272300 | BCG_1175 | BCG_1175 | 699 | 1241933 | 1242631 | BCGDan_1165 | BCGDan_1165 | 699 | 1242106 | 1242747 | BCGDan_sapMKO_1164 | BCGDan_sapMKO_1164 | 642 |
| 1241003 | 1241188 | Rv1116 | Rv1116 | 186 | 1272418 | 1272603 | BCG_1176 | BCG_1176 | 186 | 1242749 | 1242934 | BCGDan_1166 | BCGDan_1166 | 186 | 1242865 | 1243050 | BCGDan_sapMKO_1165 | BCGDan_sapMKO_1165 | 186 |
| 1241115 | 1241390 | Rv1116A | Rv1116A | 276 | 1272530 | 1272805 | BCG_1177c | BCG_1177c | 276 | 1242861 | 1243136 | BCGDan_1167 | BCGDan_1167 | 276 | 1242977 | 1243252 | BCGDan_sapMKO_1166 | BCGDan_sapMKO_1166 | 276 |
| 1241633 | 1241956 | Rv1117 | Rv1117 | 324 | 1273048 | 1273371 | BCG_1178 | BCG_1178 | 324 | 1243379 | 1243702 | BCGDan_1168 | BCGDan_1168 | 324 | 1243495 | 1243818 | BCGDan_sapMKO_1167 | BCGDan_sapMKO_1167 | 324 |
| 1241971 | 1242831 | Rv1118c | Rv1118c | 861 | 1273386 | 1274246 | BCG_1179c | BCG_1179c | 861 | 1243717 | 1244577 | BCGDan_1169 | BCGDan_1169 | 861 | 1243833 | 1244693 | BCGDan_sapMKO_1168 | BCGDan_sapMKO_1168 | 861 |
| 1242864 | 1243013 | Rv1119c | Rv1119c | 150 | 1274279 | 1274428 | BCG_1180c | BCG_1180c | 150 | 1244610 | 1244759 | BCGDan_1170 | BCGDan_1170 | 150 | 1244726 | 1244875 | BCGDan_sapMKO_1169 | BCGDan_sapMKO_1169 | 150 |
| 1243010 | 1243504 | Rv1120c | Rv1120c | 495 | 1274425 | 1274919 | BCG_1181c | BCG_1181c | 495 | 1244756 | 1245250 | BCGDan_1171 | BCGDan_1171 | 495 | 1244872 | 1245366 | BCGDan_sapMKO_1170 | BCGDan_sapMKO_1170 | 495 |
| 1243707 | 1245107 | zwf1 | Rv1121 | 1401 | 1275122 | 1276522 | zwf1 | BCG_1182 | 1401 | 1245453 | 1246853 | zwf1 | BCGDan_1172 | 1401 | 1245569 | 1246969 | zwf1 | BCGDan_sapMKO_1171 | 1401 |
| 1245129 | 1246151 | gnd2 | Rv1122 | 1023 | 1276544 | 1277566 | gnd2 | BCG_1183 | 1023 | 1246875 | 1247897 | gnd2 | BCGDan_1173 | 1023 | 1247003 | 1248013 | gnd2 | BCGDan_sapMKO_1172 | 1011 |
| 1246144 | 1247052 | bpoB | Rv1123c | 909 | 1277559 | 1278467 | bpoB | BCG_1184c | 909 | 1247890 | 1248798 | bpoB | BCGDan_1174 | 909 | 1248006 | 1248872 | bpoB | BCGDan_sapMKO_1173 | 867 |
| 1247127 | 1248077 | ephC | Rv1124 | 951 | 1278542 | 1279492 | ephC | BCG_1185 | 951 | 1248873 | 1249823 | ephC | BCGDan_1175 | 951 | 1249058 | 1249939 | ephC | BCGDan_sapMKO_1174 | 882 |
| 1248082 | 1249326 | Rv1125 | Rv1125 | 1245 | 1279497 | 1280741 | BCG_1186 | BCG_1186 | 1245 | 1249828 | 1251072 | BCGDan_1176 | BCGDan_1176 | 1245 | 1249944 | 1251188 | BCGDan_sapMKO_1175 | BCGDan_sapMKO_1175 | 1245 |
| 1249330 | 1249935 | Rv1126c | Rv1126c | 606 | 1280745 | 1281350 | BCG_1187c | BCG_1187c | 606 | 1251076 | 1251681 | BCGDan_1177 | BCGDan_1177 | 606 | 1251192 | 1251797 | BCGDan_sapMKO_1176 | BCGDan_sapMKO_1176 | 606 |
| 1249932 | 1251404 | ppdK | Rv1127c | 1473 | 1281347 | 1282819 | ppdK | BCG_1188c | 1473 | 1251678 | 1253150 | ppdK | BCGDan_1178 | 1473 | 1251794 | 1253266 | ppdK | BCGDan_sapMKO_1177 | 1473 |
| 1251617 | 1252972 | Rv1128c | Rv1128c | 1356 | 1283032 | 1284387 | BCG_1189c | BCG_1189c | 1356 | 1253363 | 1254718 | BCGDan_1179 | BCGDan_1179 | 1356 | 1253479 | 1254834 | BCGDan_sapMKO_1178 | BCGDan_sapMKO_1178 | 1356 |
| 1253074 | 1254534 | Rv1129c | Rv1129c | 1461 | 1284489 | 1285949 | BCG_1190c | BCG_1190c | 1461 | 1254820 | 1256280 | BCGDan_1180 | BCGDan_1180 | 1461 | 1254936 | 1256297 | BCGDan_sapMKO_1179 | BCGDan_sapMKO_1179 | 1362 |
| 1254555 | 1256135 | prpD | Rv1130 | 1581 | 1285970 | 1287550 | BCG_1191 | BCG_1191 | 1581 | 1256301 | 1257881 | BCGDan_1181 | BCGDan_1181 | 1581 | 1256480 | 1257997 | BCGDan_sapMKO_1180 | BCGDan_sapMKO_1180 | 1518 |
| 1256132 | 1257313 | prpC | Rv1131 | 1182 | 1287547 | 1288728 | gltA1 | BCG_1192 | 1182 | 1257878 | 1259059 | gltA1 | BCGDan_1182 | 1182 | 1257994 | 1259175 | gltA1 | BCGDan_sapMKO_1181 | 1182 |
| 1257325 | 1259055 | Rv1132 | Rv1132 | 1731 | 1288740 | 1290470 | BCG_1193 | BCG_1193 | 1731 | 1259071 | 1260801 | BCGDan_1183 | BCGDan_1183 | 1731 | 1259187 | 1260917 | BCGDan_sapMKO_1182 | BCGDan_sapMKO_1182 | 1731 |
| 1259067 | 1261346 | metE | Rv1133c | 2280 | 1290482 | 1292761 | metE | BCG_1194c | 2280 | 1260813 | 1263092 | metE | BCGDan_1184 | 2280 | 1260929 | 1263313 | metE | BCGDan_sapMKO_1183 | 2385 |
| 1261922 | 1262158 | Rv1134 | Rv1134 | 237 | 1293337 | 1293573 | BCG_1195 | BCG_1195 | 237 | 1263668 | 1263904 | BCGDan_1185 | BCGDan_1185 | 237 | 1263784 | 1264020 | BCGDan_sapMKO_1184 | BCGDan_sapMKO_1184 | 237 |
| 1262272 | 1264128 | PPE16 | Rv1135c | 1857 | 1293687 | 1295543 | PPE16 | BCG_1196c | 1857 | 1264018 | 1265874 | PPE16 | BCGDan_1186 | 1857 | 1264134 | 1265990 | PPE16 | BCGDan_sapMKO_1185 | 1857 |
| 1264314 | 1264556 | Rv1135A | Rv1135A | 243 | 1295729 | 1295971 | BCG_1197 | BCG_1197 | 243 | 1266060 | 1266302 | BCGDan_1187 | BCGDan_1187 | 243 | 1266176 | 1266418 | BCGDan_sapMKO_1186 | BCGDan_sapMKO_1186 | 243 |
| 1264606 | 1264947 | Rv1136 | Rv1136 | 342 | 1296021 | 1296362 | BCG_1198 | BCG_1198 | 342 | 1266352 | 1266693 | BCGDan_1188 | BCGDan_1188 | 342 | 1266468 | 1266809 | BCGDan_sapMKO_1187 | BCGDan_sapMKO_1187 | 342 |
| 1265087 | 1265455 | Rv1137c | Rv1137c | 369 | 1296502 | 1296870 | BCG_1199c | BCG_1199c | 369 | 1266833 | 1267201 | BCGDan_1189 | BCGDan_1189 | 369 | 1266949 | 1267317 | BCGDan_sapMKO_1188 | BCGDan_sapMKO_1188 | 369 |
| 1265472 | 1266488 | Rv1138c | Rv1138c | 1017 | 1296887 | 1297903 | BCG_1200c | BCG_1200c | 1017 | 1267218 | 1268234 | BCGDan_1190 | BCGDan_1190 | 1017 | 1267334 | 1268350 | BCGDan_sapMKO_1189 | BCGDan_sapMKO_1189 | 1017 |
| 1266485 | 1266985 | Rv1139c | Rv1139c | 501 | 1297900 | 1298400 | BCG_1201c | BCG_1201c | 501 | 1268231 | 1268731 | BCGDan_1191 | BCGDan_1191 | 501 | 1268347 | 1268847 | BCGDan_sapMKO_1190 | BCGDan_sapMKO_1190 | 501 |
| 1267347 | 1268195 | Rv1140 | Rv1140 | 849 | 1298762 | 1299610 | BCG_1202 | BCG_1202 | 849 | 1269093 | 1269941 | BCGDan_1192 | BCGDan_1192 | 849 | 1269335 | 1270057 | BCGDan_sapMKO_1191 | BCGDan_sapMKO_1191 | 723 |
| 1268203 | 1269009 | echA11 | Rv1141c | 807 | 1299618 | 1300424 | echA11 | BCG_1203c | 807 | 1269949 | 1270755 | echA11 | BCGDan_1193 | 807 | 1270065 | 1270880 | echA11 | BCGDan_sapMKO_1192 | 816 |
| 1269152 | 1269958 | echA10 | Rv1142c | 807 | 1300567 | 1301373 | echA10 | BCG_1204c | 807 | 1270898 | 1271704 | echA10 | BCGDan_1194 | 807 | 1271014 | 1271820 | echA10 | BCGDan_sapMKO_1193 | 807 |
| 1270062 | 1271144 | mcr | Rv1143 | 1083 | 1301477 | 1302559 | mcr | BCG_1205 | 1083 | 1271808 | 1272890 | mcr | BCGDan_1195 | 1083 | 1271819 | 1273006 | mcr | BCGDan_sapMKO_1194 | 1188 |
| 1271156 | 1271908 | Rv1144 | Rv1144 | 753 | 1302571 | 1303323 | BCG_1206 | BCG_1206 | 753 | 1272902 | 1273654 | BCGDan_1196 | BCGDan_1196 | 753 | 1273018 | 1273770 | BCGDan_sapMKO_1195 | BCGDan_sapMKO_1195 | 753 |
| 1272423 | 1273334 | mmpL13a | Rv1145 | 912 | 1303838 | 1306183 | mmpL13 | BCG_1207 | 2346 | 1274169 | 1276514 | mmpL13 | BCGDan_1197 | 2346 | 1274285 | 1276630 | mmpL13 | BCGDan_sapMKO_1196 | 2346 |
| 1274900 | 1275550 | Rv1147 | Rv1147 | 651 | 1306316 | 1306966 | BCG_1208 | BCG_1208 | 651 | 1276647 | 1277297 | BCGDan_1198 | BCGDan_1198 | 651 | 1276763 | 1277413 | BCGDan_sapMKO_1197 | BCGDan_sapMKO_1197 | 651 |
| 1276300 | 1277748 | Rv1148c | Rv1148c | 1449 | 1307716 | 1309164 | BCG_1209c | BCG_1209c | 1449 | 1278047 | 1279495 | BCGDan_1199 | BCGDan_1199 | 1449 | 1278163 | 1279527 | BCGDan_sapMKO_1198 | BCGDan_sapMKO_1198 | 1365 |
| 1277893 | 1278300 | Rv1149 | Rv1149 | 408 | 1309309 | 1309716 | BCG_1210 | BCG_1210 | 408 | 1279640 | 1280047 | BCGDan_1200 | BCGDan_1200 | 408 | 1279765 | 1280163 | BCGDan_sapMKO_1199 | BCGDan_sapMKO_1199 | 399 |
| 1278269 | 1278820 | Rv1150 | Rv1150 | 552 | 1309685 | 1310236 | BCG_1211 | BCG_1211 | 552 | 1280016 | 1280567 | BCGDan_1201 | BCGDan_1201 | 552 | 1280132 | 1280680 | BCGDan_sapMKO_1200 | BCGDan_sapMKO_1200 | 549 |
| 1278904 | 1279617 | Rv1151c | Rv1151c | 714 | 1310320 | 1311033 | BCG_1212c | BCG_1212c | 714 | 1280651 | 1281364 | BCGDan_1202 | BCGDan_1202 | 714 | 1280767 | 1281480 | BCGDan_sapMKO_1201 | BCGDan_sapMKO_1201 | 714 |
| 1279655 | 1280020 | Rv1152 | Rv1152 | 366 | 1311071 | 1311436 | BCG_1213 | BCG_1213 | 366 | 1281432 | 1281767 | BCGDan_1203 | BCGDan_1203 | 336 | 1281548 | 1281883 | BCGDan_sapMKO_1202 | BCGDan_sapMKO_1202 | 336 |
| 1279998 | 1280846 | omt | Rv1153c | 849 | 1311414 | 1312262 | omt | BCG_1214c | 849 | 1281745 | 1282593 | omt | BCGDan_1204 | 849 | 1281861 | 1282709 | omt | BCGDan_sapMKO_1203 | 849 |
| 1280843 | 1281484 | Rv1154c | Rv1154c | 642 | 1312259 | 1312900 | BCG_1215c | BCG_1215c | 642 | 1282590 | 1283231 | BCGDan_1205 | BCGDan_1205 | 642 | 1282706 | 1283293 | BCGDan_sapMKO_1204 | BCGDan_sapMKO_1204 | 588 |
| 1281429 | 1281872 | Rv1155 | Rv1155 | 444 | 1312845 | 1313288 | BCG_1216 | BCG_1216 | 444 | 1283176 | 1283619 | BCGDan_1206 | BCGDan_1206 | 444 | 1283292 | 1283735 | BCGDan_sapMKO_1205 | BCGDan_sapMKO_1205 | 444 |
| NA | NA | NA | NA | NA | NA | NA | NA | NA | NA | 1283777 | 1283965 | BCGDan_1207 | BCGDan_1207 | 189 | 1283893 | 1284081 | BCGDan_sapMKO_1206 | BCGDan_sapMKO_1206 | 189 |
| 1282306 | 1282893 | Rv1156 | Rv1156 | 588 | 1313722 | 1314309 | BCG_1217 | BCG_1217 | 588 | 1284053 | 1284640 | BCGDan_1208 | BCGDan_1208 | 588 | 1284169 | 1284756 | BCGDan_sapMKO_1207 | BCGDan_sapMKO_1207 | 588 |
| 1283056 | 1284171 | Rv1157c | Rv1157c | 1116 | 1314472 | 1315515 | BCG_1218c | BCG_1218c | 1044 | 1284803 | 1285846 | BCGDan_1209 | BCGDan_1209 | 1044 | 1284919 | 1285962 | BCGDan_sapMKO_1208 | BCGDan_sapMKO_1208 | 1044 |
| 1284179 | 1284862 | Rv1158c | Rv1158c | 684 | 1315523 | 1316206 | BCG_1219c | BCG_1219c | 684 | 1285854 | 1286537 | BCGDan_1210 | BCGDan_1210 | 684 | 1285970 | 1286653 | BCGDan_sapMKO_1209 | BCGDan_sapMKO_1209 | 684 |
| 1284992 | 1286287 | pimE | Rv1159 | 1296 | 1316336 | 1317631 | BCG_1220 | BCG_1220 | 1296 | 1286667 | 1287962 | BCGDan_1211 | BCGDan_1211 | 1296 | 1286783 | 1288078 | BCGDan_sapMKO_1210 | BCGDan_sapMKO_1210 | 1296 |
| 1286284 | 1286568 | Rv1159A | Rv1159A | 285 | 1317628 | 1317912 | BCG_1221c | BCG_1221c | 285 | 1287959 | 1288243 | BCGDan_1212 | BCGDan_1212 | 285 | 1288075 | 1288359 | BCGDan_sapMKO_1211 | BCGDan_sapMKO_1211 | 285 |
| 1286595 | 1287020 | mutT2 | Rv1160 | 426 | 1317939 | 1318364 | mutT2 | BCG_1222 | 426 | 1288270 | 1288695 | mutT2 | BCGDan_1213 | 426 | 1288386 | 1288811 | mutT2 | BCGDan_sapMKO_1212 | 426 |
| 1287328 | 1291026 | narG | Rv1161 | 3699 | 1318672 | 1322370 | narG | BCG_1223 | 3699 | 1289003 | 1292701 | narG | BCGDan_1214 | 3699 | 1289119 | 1292817 | narG | BCGDan_sapMKO_1213 | 3699 |
| 1291065 | 1292741 | narH | Rv1162 | 1677 | 1322409 | 1324085 | narH | BCG_1224 | 1677 | 1292740 | 1294416 | narH | BCGDan_1215 | 1677 | 1292856 | 1294532 | narH | BCGDan_sapMKO_1214 | 1677 |
| 1292798 | 1293403 | narJ | Rv1163 | 606 | 1324142 | 1324747 | narJ | BCG_1225 | 606 | 1294473 | 1295078 | narJ | BCGDan_1216 | 606 | 1294589 | 1295194 | narJ | BCGDan_sapMKO_1215 | 606 |

| | | | | | | | | | | | | | | | | | | | |
|---|---|---|---|---|---|---|---|---|---|---|---|---|---|---|---|---|---|---|---|
| 1293406 | 1294146 | narI | Rv1164 | 741 | 1324750 | 1325490 | narI | BCG_1226 | 741 | 1295081 | 1295821 | narI | BCGDan_1217 | 741 | 1295197 | 1295937 | narI | BCGDan_sapMKO_1216 | 741 |
| 1294168 | 1296054 | typA | Rv1165 | 1887 | 1325512 | 1327398 | typA | BCG_1227 | 1887 | 1295843 | 1297729 | typA | BCGDan_1218 | 1887 | 1295959 | 1297845 | typA | BCGDan_sapMKO_1217 | 1887 |
| 1296152 | 1298059 | lpqW | Rv1166 | 1908 | 1327496 | 1329403 | lpqW | BCG_1228 | 1908 | 1297827 | 1299734 | lpqW | BCGDan_1219 | 1908 | 1297976 | 1299850 | lpqW | BCGDan_sapMKO_1218 | 1875 |
| 1298087 | 1298692 | Rv1167c | Rv1167c | 606 | 1329431 | 1330036 | BCG_1229c | BCG_1229c | 606 | 1299762 | 1300367 | BCGDan_1220 | BCGDan_1220 | 606 | 1299878 | 1300483 | BCGDan_sapMKO_1219 | BCGDan_sapMKO_1219 | 606 |
| 1298764 | 1299804 | PPE17 | Rv1168c | 1041 | 1330108 | 1330653 | PPE17b | BCG_1230c | 546 | 1300439 | 1300984 | PPE17b | BCGDan_1221 | 546 | 1300555 | 1301040 | BCGDan_sapMKO_1220 | BCGDan_sapMKO_1220 | 486 |
| NA | NA | NA | NA | NA | 1330607 | 1331149 | PPE17a | BCG_1231c | 543 | 1300938 | 1301480 | PPE17a | BCGDan_1222 | 543 | 1301054 | 1301596 | PPE17a | BCGDan_sapMKO_1221 | 543 |
| 1299822 | 1300124 | lipX | Rv1169c | 303 | 1331167 | 1331469 | PE11 | BCG_1232c | 303 | 1301498 | 1301800 | PE11 | BCGDan_1223 | 303 | 1301614 | 1301934 | PE11 | BCGDan_sapMKO_1222 | 321 |
| 1300304 | 1301215 | mshB | Rv1170 | 912 | 1331649 | 1332560 | mshB | BCG_1233 | 912 | 1301980 | 1302891 | mshB | BCGDan_1224 | 912 | 1302096 | 1303007 | mshB | BCGDan_sapMKO_1223 | 912 |
| 1301307 | 1301747 | Rv1171 | Rv1171 | 441 | 1332652 | 1333092 | BCG_1234 | BCG_1234 | 441 | 1302983 | 1303423 | BCGDan_1225 | BCGDan_1225 | 441 | 1303099 | 1303539 | BCGDan_sapMKO_1224 | BCGDan_sapMKO_1224 | 441 |
| 1301755 | 1302681 | PE12 | Rv1172c | 927 | 1333100 | 1334026 | PE12 | BCG_1235c | 927 | 1303431 | 1304357 | PE12 | BCGDan_1226 | 927 | 1303547 | 1304473 | PE12 | BCGDan_sapMKO_1225 | 927 |
| 1302931 | 1305501 | fbiC | Rv1173 | 2571 | 1334276 | 1336846 | fbiC | BCG_1236 | 2571 | 1304607 | 1307177 | fbiC | BCGDan_1227 | 2571 | 1304843 | 1307293 | fbiC | BCGDan_sapMKO_1226 | 2451 |
| 1305669 | 1306001 | TB8.4 | Rv1174c | 333 | 1336952 | 1337284 | BCG_1237c | BCG_1237c | 333 | 1307283 | 1307615 | BCGDan_1228 | BCGDan_1228 | 333 | 1307399 | 1307731 | BCGDan_sapMKO_1227 | BCGDan_sapMKO_1227 | 333 |
| 1306202 | 1308226 | fadH | Rv1175c | 2025 | 1337485 | 1339509 | fadH | BCG_1238c | 2025 | 1307816 | 1309840 | fadH | BCGDan_1229 | 2025 | 1307932 | 1309956 | fadH | BCGDan_sapMKO_1228 | 2025 |
| 1308223 | 1308792 | Rv1176c | Rv1176c | 570 | 1339506 | 1340075 | BCG_1239c | BCG_1239c | 570 | 1309837 | 1310406 | BCGDan_1230 | BCGDan_1230 | 570 | 1309953 | 1310522 | BCGDan_sapMKO_1229 | BCGDan_sapMKO_1229 | 570 |
| 1309005 | 1309331 | fdxC | Rv1177 | 327 | 1340288 | 1340614 | fdxC | BCG_1240 | 327 | 1310619 | 1310945 | fdxC | BCGDan_1231 | 327 | 1310735 | 1311061 | fdxC | BCGDan_sapMKO_1230 | 327 |
| 1309364 | 1310452 | Rv1178 | Rv1178 | 1089 | 1340647 | 1341735 | BCG_1241 | BCG_1241 | 1089 | 1310978 | 1312066 | BCGDan_1232 | BCGDan_1232 | 1089 | 1311094 | 1312182 | BCGDan_sapMKO_1231 | BCGDan_sapMKO_1231 | 1089 |
| 1310480 | 1313299 | Rv1179c | Rv1179c | 2820 | 1341763 | 1344582 | BCG_1242c | BCG_1242c | 2820 | 1312094 | 1314913 | BCGDan_1233 | BCGDan_1233 | 2820 | 1312210 | 1315029 | BCGDan_sapMKO_1232 | BCGDan_sapMKO_1232 | 2820 |
| 1313725 | 1315191 | pks3 | Rv1180 | 1467 | 1345009 | 1351266 | pks3 | BCG_1243 | 6258 | 1315340 | 1321597 | pks3 | BCGDan_1234 | 6258 | 1315456 | 1321713 | pks3 | BCGDan_sapMKO_1233 | 6258 |
| 1320035 | 1321453 | papA3 | Rv1182 | 1419 | 1351319 | 1352737 | papA3 | BCG_1244 | 1419 | 1321650 | 1323068 | papA3 | BCGDan_1235 | 1419 | 1321766 | 1323184 | papA3 | BCGDan_sapMKO_1234 | 1419 |
| 1321520 | 1324528 | mmpL10 | Rv1183 | 3009 | 1352804 | 1355812 | mmpL10 | BCG_1245 | 3009 | 1323135 | 1326143 | mmpL10 | BCGDan_1236 | 3009 | 1323251 | 1326259 | mmpL10 | BCGDan_sapMKO_1235 | 3009 |
| 1324532 | 1325611 | Rv1184c | Rv1184c | 1080 | 1355816 | 1356895 | BCG_1246c | BCG_1246c | 1080 | 1326147 | 1327226 | BCGDan_1237 | BCGDan_1237 | 1080 | 1326263 | 1327342 | BCGDan_sapMKO_1236 | BCGDan_sapMKO_1236 | 1080 |
| 1325776 | 1327512 | fadD21 | Rv1185c | 1737 | 1357060 | 1358796 | fadD21 | BCG_1247c | 1737 | 1327391 | 1329127 | fadD21 | BCGDan_1238 | 1737 | 1327507 | 1329243 | fadD21 | BCGDan_sapMKO_1237 | 1737 |
| 1327689 | 1329305 | Rv1186c | Rv1186c | 1617 | 1358973 | 1360589 | BCG_1248c | BCG_1248c | 1617 | 1329304 | 1330920 | BCGDan_1239 | BCGDan_1239 | 1617 | 1329420 | 1331036 | BCGDan_sapMKO_1238 | BCGDan_sapMKO_1238 | 1617 |
| 1329390 | 1331021 | rocA | Rv1187 | 1632 | 1360674 | 1362305 | rocA | BCG_1249 | 1632 | 1331005 | 1332636 | rocA | BCGDan_1240 | 1632 | 1331109 | 1332752 | rocA | BCGDan_sapMKO_1239 | 1644 |
| 1331021 | 1332010 | Rv1188 | Rv1188 | 990 | 1362305 | 1363294 | BCG_1250 | BCG_1250 | 990 | 1332636 | 1333625 | BCGDan_1241 | BCGDan_1241 | 990 | 1332752 | 1333741 | BCGDan_sapMKO_1240 | BCGDan_sapMKO_1240 | 990 |
| 1332092 | 1332964 | sigI | Rv1189 | 873 | 1363376 | 1364581 | sigI | BCG_1251 | 1206 | 1333707 | 1334579 | sigI | BCGDan_1242 | 873 | 1333823 | 1334695 | sigI | BCGDan_sapMKO_1241 | 873 |
| NA | NA | NA | NA | NA | NA | NA | NA | NA | NA | 1334595 | 1335473 | BCGDan_1243 | BCGDan_1243 | 879 | 1334711 | 1335589 | BCGDan_sapMKO_1242 | BCGDan_sapMKO_1242 | 879 |
| NA | NA | NA | NA | NA | NA | NA | NA | NA | NA | 1335546 | 1335974 | BCGDan_1244 | BCGDan_1244 | 429 | 1335690 | 1336577 | BCGDan_sapMKO_1243 | BCGDan_sapMKO_1243 | 888 |
| 1334927 | 1335754 | Rv1192 | Rv1192 | 828 | 1364663 | 1365490 | BCG_1252 | BCG_1252 | 828 | 1336543 | 1337370 | BCGDan_1245 | BCGDan_1245 | 828 | 1336659 | 1337486 | BCGDan_sapMKO_1244 | BCGDan_sapMKO_1244 | 828 |
| 1335794 | 1337215 | fadD36 | Rv1193 | 1422 | 1365530 | 1366951 | fadD36 | BCG_1253 | 1422 | 1337410 | 1338831 | fadD36 | BCGDan_1246 | 1422 | 1337526 | 1338947 | fadD36 | BCGDan_sapMKO_1245 | 1422 |
| 1337248 | 1338513 | Rv1194c | Rv1194c | 1266 | 1366984 | 1368249 | BCG_1254c | BCG_1254c | 1266 | 1338864 | 1340129 | BCGDan_1247 | BCGDan_1247 | 1266 | 1338980 | 1340266 | BCGDan_sapMKO_1246 | BCGDan_sapMKO_1246 | 1287 |
| 1339003 | 1339302 | PE13 | Rv1195 | 300 | 1368725 | 1369024 | PE13 | BCG_1255 | 300 | 1340605 | 1340904 | PE13 | BCGDan_1248 | 300 | 1340691 | 1341020 | PE13 | BCGDan_sapMKO_1247 | 330 |
| 1339349 | 1340524 | PPE18 | Rv1196 | 1176 | 1369071 | 1370243 | PPE18 | BCG_1256 | 1173 | 1340951 | 1342123 | PPE18 | BCGDan_1249 | 1173 | 1341067 | 1342239 | PPE18 | BCGDan_sapMKO_1248 | 1173 |
| 1340659 | 1340955 | esxK | Rv1197 | 297 | 1370378 | 1370674 | esxK | BCG_1257 | 297 | 1342258 | 1342554 | esxK | BCGDan_1250 | 297 | 1342374 | 1342670 | esxK | BCGDan_sapMKO_1249 | 297 |
| 1341006 | 1341290 | esxL | Rv1198 | 285 | 1370725 | 1371009 | esxL | BCG_1258 | 285 | 1342605 | 1342889 | esxL | BCGDan_1251 | 285 | 1342721 | 1343005 | esxL | BCGDan_sapMKO_1250 | 285 |
| 1341358 | 1342605 | Rv1199c | Rv1199c | 1248 | 1371077 | 1372324 | BCG_1259c | BCG_1259c | 1248 | 1342957 | 1344204 | BCGDan_1252 | BCGDan_1252 | 1248 | 1343073 | 1344383 | BCGDan_sapMKO_1251 | BCGDan_sapMKO_1251 | 1311 |
| 1342942 | 1344219 | Rv1200 | Rv1200 | 1278 | 1372661 | 1373938 | BCG_1260 | BCG_1260 | 1278 | 1344541 | 1345818 | BCGDan_1253 | BCGDan_1253 | 1278 | 1344657 | 1345934 | BCGDan_sapMKO_1252 | BCGDan_sapMKO_1252 | 1278 |
| 1344216 | 1345169 | dapD | Rv1201c | 954 | 1373935 | 1374888 | BCG_1261c | BCG_1261c | 954 | 1345815 | 1346768 | BCGDan_1254 | BCGDan_1254 | 954 | 1345931 | 1346884 | BCGDan_sapMKO_1253 | BCGDan_sapMKO_1253 | 954 |
| 1345260 | 1346324 | dapE | Rv1202 | 1065 | 1374979 | 1376043 | dapE | BCG_1262 | 1065 | 1346859 | 1347923 | dapE | BCGDan_1255 | 1065 | 1346975 | 1348039 | dapE | BCGDan_sapMKO_1254 | 1065 |
| 1346321 | 1346905 | Rv1203c | Rv1203c | 585 | 1376040 | 1376624 | BCG_1263c | BCG_1263c | 585 | 1347920 | 1348504 | BCGDan_1256 | BCGDan_1256 | 585 | 1348036 | 1348620 | BCGDan_sapMKO_1255 | BCGDan_sapMKO_1255 | 585 |
| 1346936 | 1348624 | Rv1204c | Rv1204c | 1689 | 1376655 | 1378343 | BCG_1264c | BCG_1264c | 1689 | 1348535 | 1350223 | BCGDan_1257 | BCGDan_1257 | 1689 | 1348651 | 1350339 | BCGDan_sapMKO_1256 | BCGDan_sapMKO_1256 | 1689 |
| 1348719 | 1349282 | Rv1205 | Rv1205 | 564 | 1378438 | 1379001 | BCG_1265 | BCG_1265 | 564 | 1350318 | 1350881 | BCGDan_1258 | BCGDan_1258 | 564 | 1350434 | 1350997 | BCGDan_sapMKO_1257 | BCGDan_sapMKO_1257 | 564 |
| 1349332 | 1351125 | fadD6 | Rv1206 | 1794 | 1379051 | 1380844 | fadD6 | BCG_1266 | 1794 | 1350931 | 1352724 | fadD6 | BCGDan_1259 | 1794 | 1351047 | 1352840 | fadD6 | BCGDan_sapMKO_1258 | 1794 |
| 1351191 | 1352147 | folP2 | Rv1207 | 957 | 1380910 | 1381866 | folP2 | BCG_1267 | 957 | 1352790 | 1353746 | folP2 | BCGDan_1260 | 957 | 1353038 | 1353862 | folP2 | BCGDan_sapMKO_1259 | 825 |
| 1352144 | 1353118 | gpgS | Rv1208 | 975 | 1381863 | 1382837 | BCG_1268 | BCG_1268 | 975 | 1353743 | 1354717 | BCGDan_1261 | BCGDan_1261 | 975 | 1353859 | 1354833 | BCGDan_sapMKO_1260 | BCGDan_sapMKO_1260 | 975 |
| 1353157 | 1353525 | Rv1209 | Rv1209 | 369 | 1382876 | 1383244 | BCG_1269 | BCG_1269 | 369 | 1354756 | 1355124 | BCGDan_1262 | BCGDan_1262 | 369 | 1354872 | 1355240 | BCGDan_sapMKO_1261 | BCGDan_sapMKO_1261 | 369 |
| 1353522 | 1354136 | tagA | Rv1210 | 615 | 1383241 | 1383855 | tagA | BCG_1270 | 615 | 1355121 | 1355735 | tagA | BCGDan_1263 | 615 | 1355237 | 1355851 | tagA | BCGDan_sapMKO_1262 | 615 |
| 1354243 | 1354470 | Rv1211 | Rv1211 | 228 | 1383963 | 1384190 | BCG_1271 | BCG_1271 | 228 | 1355842 | 1356069 | BCGDan_1264 | BCGDan_1264 | 228 | 1355958 | 1356185 | BCGDan_sapMKO_1263 | BCGDan_sapMKO_1263 | 228 |
| 1354498 | 1355661 | glgA | Rv1212c | 1164 | 1384218 | 1385381 | BCG_1272c | BCG_1272c | 1164 | 1356097 | 1357260 | BCGDan_1265 | BCGDan_1265 | 1164 | 1356213 | 1357376 | BCGDan_sapMKO_1264 | BCGDan_sapMKO_1264 | 1164 |
| 1355836 | 1357050 | glgC | Rv1213 | 1215 | 1385556 | 1386770 | glgC | BCG_1273 | 1215 | 1357435 | 1358649 | glgC | BCGDan_1266 | 1215 | 1357560 | 1358765 | glgC | BCGDan_sapMKO_1265 | 1206 |
| 1357293 | 1357625 | PE14 | Rv1214c | 333 | 1387013 | 1387345 | PE14 | BCG_1274c | 333 | 1358892 | 1359224 | PE14 | BCGDan_1267 | 333 | 1359008 | 1359340 | PE14 | BCGDan_sapMKO_1266 | 333 |
| 1357759 | 1359444 | Rv1215c | Rv1215c | 1686 | 1387479 | 1389164 | BCG_1275c | BCG_1275c | 1686 | 1359358 | 1361043 | BCGDan_1268 | BCGDan_1268 | 1686 | 1359474 | 1361135 | BCGDan_sapMKO_1267 | BCGDan_sapMKO_1267 | 1662 |
| 1359472 | 1360146 | Rv1216c | Rv1216c | 675 | 1389192 | 1389866 | BCG_1276c | BCG_1276c | 675 | 1361071 | 1361745 | BCGDan_1269 | BCGDan_1269 | 675 | 1361187 | 1361861 | BCGDan_sapMKO_1268 | BCGDan_sapMKO_1268 | 675 |
| 1360155 | 1361801 | Rv1217c | Rv1217c | 1647 | 1389875 | 1391521 | BCG_1277c | BCG_1277c | 1647 | 1361754 | 1363400 | BCGDan_1270 | BCGDan_1270 | 1647 | 1361870 | 1363516 | BCGDan_sapMKO_1269 | BCGDan_sapMKO_1269 | 1647 |
| 1361798 | 1362733 | Rv1218c | Rv1218c | 936 | 1391518 | 1392453 | BCG_1278c | BCG_1278c | 936 | 1363397 | 1364332 | BCGDan_1271 | BCGDan_1271 | 936 | 1363513 | 1364448 | BCGDan_sapMKO_1270 | BCGDan_sapMKO_1270 | 936 |
| 1362723 | 1363361 | Rv1219c | Rv1219c | 639 | 1392443 | 1393081 | BCG_1279c | BCG_1279c | 639 | 1364322 | 1364960 | BCGDan_1272 | BCGDan_1272 | 639 | 1364438 | 1365076 | BCGDan_sapMKO_1271 | BCGDan_sapMKO_1271 | 639 |
| 1363503 | 1364150 | Rv1220c | Rv1220c | 648 | 1393223 | 1393870 | BCG_1280c | BCG_1280c | 648 | 1365102 | 1365749 | BCGDan_1273 | BCGDan_1273 | 648 | 1365218 | 1365961 | BCGDan_sapMKO_1272 | BCGDan_sapMKO_1272 | 744 |
| 1364413 | 1365186 | sigE | Rv1221 | 774 | 1394133 | 1394906 | sigE | BCG_1281 | 774 | 1366012 | 1366785 | sigE | BCGDan_1274 | 774 | 1366128 | 1366901 | sigE | BCGDan_sapMKO_1273 | 774 |
| 1365344 | 1365808 | rseA | Rv1222 | 465 | 1395064 | 1395528 | BCG_1282 | BCG_1282 | 465 | 1366943 | 1367407 | BCGDan_1275 | BCGDan_1275 | 465 | 1367059 | 1367523 | BCGDan_sapMKO_1274 | BCGDan_sapMKO_1274 | 465 |
| 1365875 | 1367461 | htrA | Rv1223 | 1587 | 1395597 | 1397183 | htrA | BCG_1283 | 1587 | 1367488 | 1369062 | htrA | BCGDan_1276 | 1575 | 1367604 | 1369178 | htrA | BCGDan_sapMKO_1275 | 1575 |
| 1367463 | 1367858 | tatB | Rv1224 | 396 | 1397185 | 1397580 | tatB | BCG_1284 | 396 | 1369064 | 1369459 | tatB | BCGDan_1277 | 396 | 1369180 | 1369575 | tatB | BCGDan_sapMKO_1276 | 396 |
| 1367891 | 1368721 | Rv1225c | Rv1225c | 831 | 1397613 | 1398443 | BCG_1285c | BCG_1285c | 831 | 1369492 | 1370322 | BCGDan_1278 | BCGDan_1278 | 831 | 1369608 | 1370438 | BCGDan_sapMKO_1277 | BCGDan_sapMKO_1277 | 831 |
| 1368832 | 1370295 | Rv1226c | Rv1226c | 1464 | 1398554 | 1400017 | BCG_1286c | BCG_1286c | 1464 | 1370433 | 1371896 | BCGDan_1279 | BCGDan_1279 | 1464 | 1370549 | 1372012 | BCGDan_sapMKO_1278 | BCGDan_sapMKO_1278 | 1464 |
| 1370292 | 1370825 | Rv1227c | Rv1227c | 534 | 1400014 | 1400547 | BCG_1287c | BCG_1287c | 534 | 1371893 | 1372426 | BCGDan_1280 | BCGDan_1280 | 534 | 1372009 | 1372542 | BCGDan_sapMKO_1279 | BCGDan_sapMKO_1279 | 534 |
| 1370920 | 1371477 | lpqX | Rv1228 | 558 | 1400642 | 1401199 | lpqX | BCG_1288 | 558 | 1372521 | 1373078 | lpqX | BCGDan_1281 | 558 | 1372637 | 1373194 | lpqX | BCGDan_sapMKO_1280 | 558 |
| 1371777 | 1372949 | mrp | Rv1229c | 1173 | 1401499 | 1402671 | mrp | BCG_1289c | 1173 | 1373378 | 1374550 | mrp | BCGDan_1282 | 1173 | 1373494 | 1374639 | mrp | BCGDan_sapMKO_1281 | 1146 |
| 1372962 | 1374197 | Rv1230c | Rv1230c | 1236 | 1402684 | 1403919 | BCG_1290c | BCG_1290c | 1236 | 1374563 | 1375798 | BCGDan_1283 | BCGDan_1283 | 1236 | 1374679 | 1375824 | BCGDan_sapMKO_1282 | BCGDan_sapMKO_1282 | 1146 |
| 1374322 | 1374864 | Rv1231c | Rv1231c | 543 | 1404044 | 1404586 | BCG_1291c | BCG_1291c | 543 | 1375923 | 1376465 | BCGDan_1284 | BCGDan_1284 | 543 | 1376039 | 1376581 | BCGDan_sapMKO_1283 | BCGDan_sapMKO_1283 | 543 |
| 1374861 | 1376168 | Rv1232c | Rv1232c | 1308 | 1404583 | 1405890 | BCG_1292c | BCG_1292c | 1308 | 1376462 | 1377769 | BCGDan_1285 | BCGDan_1285 | 1308 | 1376578 | 1377885 | BCGDan_sapMKO_1284 | BCGDan_sapMKO_1284 | 1308 |
| 1376230 | 1376826 | Rv1233c | Rv1233c | 597 | 1405952 | 1406548 | BCG_1293c | BCG_1293c | 597 | 1377831 | 1378427 | BCGDan_1286 | BCGDan_1286 | 597 | 1377947 | 1378543 | BCGDan_sapMKO_1285 | BCGDan_sapMKO_1285 | 597 |
| 1376976 | 1377503 | Rv1234 | Rv1234 | 528 | 1406698 | 1407225 | BCG_1294 | BCG_1294 | 528 | 1378577 | 1379104 | BCGDan_1287 | BCGDan_1287 | 528 | 1378765 | 1379220 | BCGDan_sapMKO_1286 | BCGDan_sapMKO_1286 | 456 |
| 1377524 | 1378930 | lpqY | Rv1235 | 1407 | 1407246 | 1408652 | lpqY | BCG_1295 | 1407 | 1379125 | 1380531 | lpqY | BCGDan_1288 | 1407 | 1379247 | 1380647 | lpqY | BCGDan_sapMKO_1287 | 1401 |
| 1378927 | 1379850 | sugA | Rv1236 | 924 | 1408649 | 1409572 | sugA | BCG_1296 | 924 | 1380528 | 1381451 | sugA | BCGDan_1289 | 924 | 1380644 | 1381567 | sugA | BCGDan_sapMKO_1288 | 924 |
| 1379855 | 1380679 | sugB | Rv1237 | 825 | 1409577 | 1410401 | sugB | BCG_1297 | 825 | 1381456 | 1382280 | sugB | BCGDan_1290 | 825 | 1381557 | 1382396 | sugB | BCGDan_sapMKO_1289 | 840 |
| 1380684 | 1381865 | sugC | Rv1238 | 1182 | 1410406 | 1411587 | sugC | BCG_1298 | 1182 | 1382285 | 1383466 | sugC | BCGDan_1291 | 1182 | 1382401 | 1383582 | sugC | BCGDan_sapMKO_1290 | 1182 |
| 1381942 | 1383042 | corA | Rv1239c | 1101 | 1411664 | 1412764 | corA | BCG_1299c | 1101 | 1383543 | 1384643 | corA | BCGDan_1292 | 1101 | 1383659 | 1384759 | corA | BCGDan_sapMKO_1291 | 1101 |
| 1383213 | 1384202 | mdh | Rv1240 | 990 | 1412935 | 1413924 | mdh | BCG_1300 | 990 | 1384814 | 1385803 | mdh | BCGDan_1293 | 990 | 1384930 | 1385919 | mdh | BCGDan_sapMKO_1292 | 990 |
| 1384278 | 1384538 | vapB33 | Rv1241 | 261 | 1414000 | 1414260 | BCG_1301 | BCG_1301 | 261 | 1385879 | 1386139 | BCGDan_1294 | BCGDan_1294 | 261 | 1385995 | 1386255 | BCGDan_sapMKO_1293 | BCGDan_sapMKO_1293 | 261 |
| 1384535 | 1384966 | vapC33 | Rv1242 | 432 | 1414257 | 1414688 | BCG_1302 | BCG_1302 | 432 | 1386136 | 1386567 | BCGDan_1295 | BCGDan_1295 | 432 | 1386252 | 1386683 | BCGDan_sapMKO_1294 | BCGDan_sapMKO_1294 | 432 |
| 1384989 | 1386677 | PE_PGRS23 | Rv1243c | 1689 | 1414711 | 1416399 | PE_PGRS23 | BCG_1303c | 1689 | 1386590 | 1388278 | PE_PGRS23 | BCGDan_1296 | 1689 | 1386706 | 1388394 | PE_PGRS23 | BCGDan_sapMKO_1295 | 1689 |
| 1386857 | 1387717 | lpqZ | Rv1244 | 861 | 1416579 | 1417439 | lpqZ | BCG_1304 | 861 | 1388458 | 1389318 | lpqZ | BCGDan_1297 | 861 | 1388451 | 1389434 | lpqZ | BCGDan_sapMKO_1296 | 984 |
| 1387798 | 1388628 | Rv1245c | Rv1245c | 831 | 1417520 | 1418350 | BCG_1305c | BCG_1305c | 831 | 1389399 | 1390229 | BCGDan_1298 | BCGDan_1298 | 831 | 1389515 | 1390345 | BCGDan_sapMKO_1297 | BCGDan_sapMKO_1297 | 831 |
| 1388685 | 1388978 | relE | Rv1246c | 294 | 1418407 | 1418700 | BCG_1306c | BCG_1306c | 294 | 1390286 | 1390579 | BCGDan_1299 | BCGDan_1299 | 294 | 1390402 | 1390695 | BCGDan_sapMKO_1298 | BCGDan_sapMKO_1298 | 294 |
| 1388975 | 1389244 | relB | Rv1247c | 270 | 1418697 | 1418966 | BCG_1307c | BCG_1307c | 270 | 1390576 | 1390845 | BCGDan_1300 | BCGDan_1300 | 270 | 1390692 | 1390961 | BCGDan_sapMKO_1299 | BCGDan_sapMKO_1299 | 270 |
| 1389357 | 1393052 | Rv1248c | Rv1248c | 3696 | 1419079 | 1422723 | kgd | BCG_1308c | 3645 | 1390958 | 1394653 | kgd | BCGDan_1301 | 3696 | 1391074 | 1394769 | kgd | BCGDan_sapMKO_1300 | 3696 |
| 1393194 | 1393982 | Rv1249c | Rv1249c | 789 | 1422916 | 1423704 | BCG_1309c | BCG_1309c | 789 | 1394795 | 1395583 | BCGDan_1302 | BCGDan_1302 | 789 | 1394911 | 1395699 | BCGDan_sapMKO_1301 | BCGDan_sapMKO_1301 | 789 |

| | | | | | | | | | | | | | | | | | | | |
|---|---|---|---|---|---|---|---|---|---|---|---|---|---|---|---|---|---|---|---|
| 1394179 | 1395918 | Rv1250 | Rv1250 | 1740 | 1423901 | 1425640 | BCG_1310 | BCG_1310 | 1740 | 1395780 | 1397519 | BCGDan_1303 | BCGDan_1303 | 1740 | 1395896 | 1397635 | BCGDan_sapMKO_1302 | BCGDan_sapMKO_1302 | 1740 |
| 1395821 | 1399240 | Rv1251c | Rv1251c | 3420 | 1425543 | 1428962 | BCG_1311c | BCG_1311c | 3420 | 1397422 | 1400841 | BCGDan_1304 | BCGDan_1304 | 3420 | 1397538 | 1400957 | BCGDan_sapMKO_1303 | BCGDan_sapMKO_1303 | 3420 |
| 1399296 | 1399904 | lprE | Rv1252c | 609 | 1429018 | 1429626 | lprE | BCG_1312c | 609 | 1400897 | 1401505 | lprE | BCGDan_1305 | 609 | 1401013 | 1401621 | lprE | BCGDan_sapMKO_1304 | 609 |
| 1399970 | 1401661 | deaD | Rv1253 | 1692 | 1429692 | 1431383 | deaD | BCG_1313 | 1692 | 1401571 | 1403262 | deaD | BCGDan_1306 | 1692 | 1401687 | 1403378 | deaD | BCGDan_sapMKO_1305 | 1692 |
| 1401658 | 1402809 | Rv1254 | Rv1254 | 1152 | 1431380 | 1432531 | BCG_1314 | BCG_1314 | 1152 | 1403259 | 1404410 | BCGDan_1307 | BCGDan_1307 | 1152 | 1403375 | 1404526 | BCGDan_sapMKO_1306 | BCGDan_sapMKO_1306 | 1152 |
| 1402778 | 1403386 | Rv1255c | Rv1255c | 609 | 1432500 | 1432799 | BCG_1315c | BCG_1315c | 300 | 1404379 | 1404678 | BCGDan_1308 | BCGDan_1308 | 300 | 1404495 | 1404794 | BCGDan_sapMKO_1307 | BCGDan_sapMKO_1307 | 300 |
| 1406081 | 1407340 | Rv1258c | Rv1258c | 1260 | 1432796 | 1434055 | BCG_1316c | BCG_1316c | 1260 | 1404675 | 1405934 | BCGDan_1309 | BCGDan_1309 | 1260 | 1404791 | 1406050 | BCGDan_sapMKO_1308 | BCGDan_sapMKO_1308 | 1260 |
| 1407339 | 1408238 | udgB | Rv1259 | 900 | 1434054 | 1434953 | BCG_1317 | BCG_1317 | 900 | 1405933 | 1406832 | BCGDan_1310 | BCGDan_1310 | 900 | 1406121 | 1406948 | BCGDan_sapMKO_1309 | BCGDan_sapMKO_1309 | 828 |
| 1408240 | 1409358 | Rv1260 | Rv1260 | 1119 | 1434955 | 1435779 | BCG_1318 | BCG_1318 | 825 | 1406834 | 1407658 | BCGDan_1311 | BCGDan_1311 | 825 | 1406950 | 1407774 | BCGDan_sapMKO_1310 | BCGDan_sapMKO_1310 | 825 |
| NA | NA | NA | NA | NA | 1435764 | 1436072 | BCG_1319 | BCG_1319 | 309 | 1407643 | 1407951 | BCGDan_1312 | BCGDan_1312 | 309 | 1407759 | 1408067 | BCGDan_sapMKO_1311 | BCGDan_sapMKO_1311 | 309 |
| 1409484 | 1409933 | Rv1261c | Rv1261c | 450 | 1436198 | 1436647 | BCG_1320c | BCG_1320c | 450 | 1408077 | 1408526 | BCGDan_1313 | BCGDan_1313 | 450 | 1408193 | 1408642 | BCGDan_sapMKO_1312 | BCGDan_sapMKO_1312 | 450 |
| 1409938 | 1410372 | Rv1262c | Rv1262c | 435 | 1436652 | 1437086 | BCG_1321c | BCG_1321c | 435 | 1408531 | 1408965 | BCGDan_1314 | BCGDan_1314 | 435 | 1408647 | 1409081 | BCGDan_sapMKO_1313 | BCGDan_sapMKO_1313 | 435 |
| 1410431 | 1411819 | amiB2 | Rv1263 | 1389 | 1437145 | 1438533 | amiB2 | BCG_1322 | 1389 | 1409024 | 1410412 | amiB2 | BCGDan_1315 | 1389 | 1409140 | 1410528 | amiB2 | BCGDan_sapMKO_1314 | 1389 |
| 1411894 | 1413087 | Rv1264 | Rv1264 | 1194 | 1438608 | 1439801 | BCG_1323 | BCG_1323 | 1194 | 1410487 | 1411680 | BCGDan_1316 | BCGDan_1316 | 1194 | 1410615 | 1411796 | BCGDan_sapMKO_1315 | BCGDan_sapMKO_1315 | 1182 |
| 1413260 | 1413940 | Rv1265 | Rv1265 | 681 | 1439974 | 1440654 | BCG_1324 | BCG_1324 | 681 | 1411853 | 1412533 | BCGDan_1317 | BCGDan_1317 | 681 | 1411969 | 1412649 | BCGDan_sapMKO_1316 | BCGDan_sapMKO_1316 | 681 |
| 1413960 | 1415840 | pknH | Rv1266c | 1881 | 1440674 | 1442464 | pknH | BCG_1325c | 1791 | 1412553 | 1414343 | pknH | BCGDan_1318 | 1791 | 1412669 | 1414459 | pknH | BCGDan_sapMKO_1317 | 1791 |
| 1416181 | 1417347 | embR | Rv1267c | 1167 | 1442805 | 1443971 | embR | BCG_1326c | 1167 | 1414684 | 1415850 | embR | BCGDan_1319 | 1167 | 1414800 | 1415966 | embR | BCGDan_sapMKO_1318 | 1167 |
| 1417658 | 1418356 | Rv1268c | Rv1268c | 699 | 1444282 | 1444980 | BCG_1327c | BCG_1327c | 699 | 1416161 | 1416859 | BCGDan_1320 | BCGDan_1320 | 699 | 1416277 | 1416975 | BCGDan_sapMKO_1319 | BCGDan_sapMKO_1319 | 699 |
| 1418579 | 1418953 | Rv1269c | Rv1269c | 375 | 1445203 | 1445577 | BCG_1328c | BCG_1328c | 375 | 1417082 | 1417456 | BCGDan_1321 | BCGDan_1321 | 375 | 1417198 | 1417620 | BCGDan_sapMKO_1320 | BCGDan_sapMKO_1320 | 423 |
| 1419014 | 1419748 | lprA | Rv1270c | 735 | 1445638 | 1446372 | lprA | BCG_1329c | 735 | 1417517 | 1418251 | lprA | BCGDan_1322 | 735 | 1417633 | 1418406 | lprA | BCGDan_sapMKO_1321 | 774 |
| 1419961 | 1420302 | Rv1271c | Rv1271c | 342 | 1446585 | 1446926 | BCG_1330c | BCG_1330c | 342 | 1418464 | 1418805 | BCGDan_1323 | BCGDan_1323 | 342 | 1418580 | 1418921 | BCGDan_sapMKO_1322 | BCGDan_sapMKO_1322 | 342 |
| 1420410 | 1422305 | Rv1272c | Rv1272c | 1896 | 1447034 | 1448929 | BCG_1331c | BCG_1331c | 1896 | 1418913 | 1420808 | BCGDan_1324 | BCGDan_1324 | 1896 | 1419029 | 1420834 | BCGDan_sapMKO_1323 | BCGDan_sapMKO_1323 | 1806 |
| 1422302 | 1424050 | Rv1273c | Rv1273c | 1749 | 1448926 | 1450674 | BCG_1332c | BCG_1332c | 1749 | 1420805 | 1422553 | BCGDan_1325 | BCGDan_1325 | 1749 | 1420921 | 1422669 | BCGDan_sapMKO_1324 | BCGDan_sapMKO_1324 | 1749 |
| 1424197 | 1424754 | lprB | Rv1274 | 558 | 1450821 | 1451378 | lprB | BCG_1333 | 558 | 1422700 | 1423257 | lprB | BCGDan_1326 | 558 | 1422801 | 1423373 | lprB | BCGDan_sapMKO_1325 | 573 |
| 1424751 | 1425293 | lprC | Rv1275 | 543 | 1451375 | 1451917 | lprC | BCG_1334 | 543 | 1423254 | 1423796 | lprC | BCGDan_1327 | 543 | 1423370 | 1423912 | lprC | BCGDan_sapMKO_1326 | 543 |
| 1425438 | 1425914 | Rv1276c | Rv1276c | 477 | 1452062 | 1452538 | BCG_1335c | BCG_1335c | 477 | 1423941 | 1424417 | BCGDan_1328 | BCGDan_1328 | 477 | 1424057 | 1424563 | BCGDan_sapMKO_1327 | BCGDan_sapMKO_1327 | 507 |
| 1426164 | 1427417 | Rv1277 | Rv1277 | 1254 | 1452788 | 1454041 | BCG_1336 | BCG_1336 | 1254 | 1424667 | 1425920 | BCGDan_1329 | BCGDan_1329 | 1254 | 1424885 | 1426036 | BCGDan_sapMKO_1328 | BCGDan_sapMKO_1328 | 1152 |
| 1427414 | 1430041 | Rv1278 | Rv1278 | 2628 | 1454038 | 1456665 | BCG_1337 | BCG_1337 | 2628 | 1425917 | 1428544 | BCGDan_1330 | BCGDan_1330 | 2628 | 1426033 | 1428660 | BCGDan_sapMKO_1329 | BCGDan_sapMKO_1329 | 2628 |
| 1430062 | 1431648 | Rv1279 | Rv1279 | 1587 | 1456686 | 1458272 | BCG_1338 | BCG_1338 | 1587 | 1428565 | 1430151 | BCGDan_1331 | BCGDan_1331 | 1587 | 1428657 | 1430267 | BCGDan_sapMKO_1330 | BCGDan_sapMKO_1330 | 1611 |
| 1431665 | 1433440 | oppA | Rv1280c | 1776 | 1458289 | 1460064 | oppA | BCG_1339c | 1776 | 1430168 | 1431943 | oppA | BCGDan_1332 | 1776 | 1430284 | 1431921 | oppA | BCGDan_sapMKO_1331 | 1638 |
| 1433433 | 1435271 | oppD | Rv1281c | 1839 | 1460057 | 1461895 | oppD | BCG_1340c | 1839 | 1431936 | 1433774 | oppD | BCGDan_1333 | 1839 | 1432052 | 1433890 | oppD | BCGDan_sapMKO_1332 | 1839 |
| 1435268 | 1436143 | oppC | Rv1282c | 876 | 1461892 | 1462767 | oppC | BCG_1341c | 876 | 1433771 | 1434646 | oppC | BCGDan_1334 | 876 | 1433887 | 1434762 | oppC | BCGDan_sapMKO_1333 | 876 |
| 1436140 | 1437117 | oppB | Rv1283c | 978 | 1462764 | 1463741 | oppB | BCG_1342c | 978 | 1434643 | 1435620 | oppB | BCGDan_1335 | 978 | 1434759 | 1435736 | oppB | BCGDan_sapMKO_1334 | 978 |
| 1437324 | 1437815 | canA | Rv1284 | 492 | 1463948 | 1464439 | canA | BCG_1343 | 492 | 1435827 | 1436318 | canA | BCGDan_1336 | 492 | 1435943 | 1436434 | canA | BCGDan_sapMKO_1335 | 492 |
| 1437909 | 1438907 | cysD | Rv1285 | 999 | 1464533 | 1465531 | cysD | BCG_1344 | 999 | 1436412 | 1437410 | cysD | BCGDan_1337 | 999 | 1436528 | 1437526 | cysD | BCGDan_sapMKO_1336 | 999 |
| 1438907 | 1440751 | cysN | Rv1286 | 1845 | 1465531 | 1467375 | cysN | BCG_1345 | 1845 | 1437410 | 1439254 | cysN | BCGDan_1338 | 1845 | 1437526 | 1439370 | cysN | BCGDan_sapMKO_1337 | 1845 |
| 1440805 | 1441290 | Rv1287 | Rv1287 | 486 | 1467429 | 1467914 | BCG_1346 | BCG_1346 | 486 | 1439308 | 1439793 | BCGDan_1339 | BCGDan_1339 | 486 | 1439424 | 1439909 | BCGDan_sapMKO_1338 | BCGDan_sapMKO_1338 | 486 |
| 1441348 | 1442718 | Rv1288 | Rv1288 | 1371 | 1467972 | 1469189 | BCG_1347 | BCG_1347 | 1218 | 1439851 | 1441068 | BCGDan_1340 | BCGDan_1340 | 1218 | 1439967 | 1441184 | BCGDan_sapMKO_1339 | BCGDan_sapMKO_1339 | 1218 |
| 1442767 | 1443399 | Rv1289 | Rv1289 | 633 | 1469238 | 1469870 | BCG_1348 | BCG_1348 | 633 | 1441129 | 1441749 | BCGDan_1341 | BCGDan_1341 | 621 | 1441245 | 1441865 | BCGDan_sapMKO_1340 | BCGDan_sapMKO_1340 | 621 |
| 1443482 | 1445047 | Rv1290c | Rv1290c | 1566 | 1470009 | 1471574 | BCG_1349c | BCG_1349c | 1566 | 1441888 | 1443453 | BCGDan_1342 | BCGDan_1342 | 1566 | 1442004 | 1443437 | BCGDan_sapMKO_1341 | BCGDan_sapMKO_1341 | 1434 |
| 1445058 | 1445372 | Rv1290A | Rv1290A | 315 | 1471585 | 1471899 | BCG_1350 | BCG_1350 | 315 | 1443464 | 1443778 | BCGDan_1343 | BCGDan_1343 | 315 | 1443580 | 1443894 | BCGDan_sapMKO_1342 | BCGDan_sapMKO_1342 | 315 |
| 1445499 | 1445834 | Rv1291c | Rv1291c | 336 | 1472026 | 1472361 | BCG_1351c | BCG_1351c | 336 | 1443905 | 1444240 | BCGDan_1344 | BCGDan_1344 | 336 | 1444021 | 1444356 | BCGDan_sapMKO_1343 | BCGDan_sapMKO_1343 | 336 |
| 1446379 | 1448031 | argS | Rv1292 | 1653 | 1472906 | 1474558 | argS | BCG_1352 | 1653 | 1444785 | 1446437 | argS | BCGDan_1346 | 1653 | 1444901 | 1446553 | argS | BCGDan_sapMKO_1345 | 1653 |
| 1448028 | 1449371 | lysA | Rv1293 | 1344 | 1474555 | 1475898 | lysA | BCG_1353 | 1344 | 1446434 | 1447777 | lysA | BCGDan_1347 | 1344 | 1446550 | 1447893 | lysA | BCGDan_sapMKO_1346 | 1344 |
| 1449375 | 1450700 | thrA | Rv1294 | 1326 | 1475902 | 1477227 | thrA | BCG_1354 | 1326 | 1447781 | 1449106 | thrA | BCGDan_1348 | 1326 | 1447897 | 1449222 | thrA | BCGDan_sapMKO_1347 | 1326 |
| 1450697 | 1451779 | thrC | Rv1295 | 1083 | 1477224 | 1478306 | thrC | BCG_1355 | 1083 | 1449103 | 1450185 | thrC | BCGDan_1349 | 1083 | 1449261 | 1450301 | thrC | BCGDan_sapMKO_1348 | 1041 |
| 1451997 | 1452947 | thrB | Rv1296 | 951 | 1478524 | 1479474 | thrB | BCG_1356 | 951 | 1450346 | 1451296 | thrB | BCGDan_1350 | 951 | 1450462 | 1451412 | thrB | BCGDan_sapMKO_1349 | 951 |
| 1453204 | 1455012 | rho | Rv1297 | 1809 | 1479731 | 1481539 | rho | BCG_1357 | 1809 | 1451553 | 1453361 | rho | BCGDan_1351 | 1809 | 1451669 | 1453477 | rho | BCGDan_sapMKO_1350 | 1809 |
| 1455163 | 1455405 | rpmE | Rv1298 | 243 | 1481690 | 1481932 | rpmE | BCG_1358 | 243 | 1453512 | 1453754 | rpmE | BCGDan_1352 | 243 | 1453628 | 1453870 | rpmE | BCGDan_sapMKO_1351 | 243 |
| 1455495 | 1456568 | prfA | Rv1299 | 1074 | 1482022 | 1483095 | prfA | BCG_1359 | 1074 | 1453844 | 1454917 | prfA | BCGDan_1353 | 1074 | 1453960 | 1455033 | prfA | BCGDan_sapMKO_1352 | 1074 |
| 1456565 | 1457542 | hemK | Rv1300 | 978 | 1483092 | 1484069 | hemK | BCG_1360 | 978 | 1454914 | 1455891 | hemK | BCGDan_1354 | 978 | 1455093 | 1456007 | hemK | BCGDan_sapMKO_1353 | 915 |
| 1457558 | 1458211 | Rv1301 | Rv1301 | 654 | 1484085 | 1484738 | BCG_1361 | BCG_1361 | 654 | 1455907 | 1456560 | BCGDan_1355 | BCGDan_1355 | 654 | 1456023 | 1456676 | BCGDan_sapMKO_1354 | BCGDan_sapMKO_1354 | 654 |
| 1458295 | 1459509 | rfe | Rv1302 | 1215 | 1484822 | 1486036 | rfe | BCG_1362 | 1215 | 1456644 | 1457858 | rfe | BCGDan_1356 | 1215 | 1456760 | 1457974 | rfe | BCGDan_sapMKO_1355 | 1215 |
| 1459766 | 1460251 | Rv1303 | Rv1303 | 486 | 1486293 | 1486778 | BCG_1363 | BCG_1363 | 486 | 1458115 | 1458600 | BCGDan_1357 | BCGDan_1357 | 486 | 1458231 | 1458716 | BCGDan_sapMKO_1356 | BCGDan_sapMKO_1356 | 486 |
| 1460244 | 1460996 | atpB | Rv1304 | 753 | 1486771 | 1487523 | atpB | BCG_1364 | 753 | 1458593 | 1459345 | atpB | BCGDan_1358 | 753 | 1458709 | 1459461 | atpB | BCGDan_sapMKO_1357 | 753 |
| 1461045 | 1461290 | atpE | Rv1305 | 246 | 1487572 | 1487817 | atpE | BCG_1365 | 246 | 1459394 | 1459639 | atpE | BCGDan_1359 | 246 | 1459468 | 1459755 | atpE | BCGDan_sapMKO_1358 | 288 |
| 1461321 | 1461836 | atpF | Rv1306 | 516 | 1487848 | 1488363 | atpF | BCG_1366 | 516 | 1459670 | 1460185 | atpF | BCGDan_1360 | 516 | 1459786 | 1460301 | atpF | BCGDan_sapMKO_1359 | 516 |
| 1461843 | 1463183 | atpH | Rv1307 | 1341 | 1488370 | 1489710 | atpH | BCG_1367 | 1341 | 1460192 | 1461532 | atpH | BCGDan_1361 | 1341 | 1460308 | 1461648 | atpH | BCGDan_sapMKO_1360 | 1341 |
| 1463228 | 1464877 | atpA | Rv1308 | 1650 | 1489755 | 1491404 | atpA | BCG_1368 | 1650 | 1461577 | 1463226 | atpA | BCGDan_1362 | 1650 | 1461693 | 1463342 | atpA | BCGDan_sapMKO_1361 | 1650 |
| 1464884 | 1465801 | atpG | Rv1309 | 918 | 1491411 | 1492328 | atpG | BCG_1369 | 918 | 1463233 | 1464150 | atpG | BCGDan_1363 | 918 | 1463349 | 1464266 | atpG | BCGDan_sapMKO_1362 | 918 |
| 1465841 | 1467301 | atpD | Rv1310 | 1461 | 1492368 | 1493828 | atpD | BCG_1370 | 1461 | 1464190 | 1465650 | atpD | BCGDan_1364 | 1461 | 1464306 | 1465766 | atpD | BCGDan_sapMKO_1363 | 1461 |
| 1467315 | 1467680 | atpC | Rv1311 | 366 | 1493842 | 1494207 | atpC | BCG_1371 | 366 | 1465664 | 1466029 | atpC | BCGDan_1365 | 366 | 1465780 | 1466145 | atpC | BCGDan_sapMKO_1364 | 366 |
| 1467688 | 1468131 | Rv1312 | Rv1312 | 444 | 1494215 | 1494649 | BCG_1372 | BCG_1372 | 435 | 1466037 | 1466471 | BCGDan_1366 | BCGDan_1366 | 435 | 1466165 | 1466587 | BCGDan_sapMKO_1365 | BCGDan_sapMKO_1365 | 423 |
| 1468171 | 1469505 | Rv1313c | Rv1313c | 1335 | 1494696 | 1495427 | BCG_1373c | BCG_1373c | 732 | 1466518 | 1467249 | BCGDan_1367 | BCGDan_1367 | 732 | 1466634 | 1467365 | BCGDan_sapMKO_1366 | BCGDan_sapMKO_1366 | 732 |
| NA | NA | NA | NA | NA | 1495402 | 1496019 | BCG_1374c | BCG_1374c | 618 | 1467224 | 1467841 | BCGDan_1368 | BCGDan_1368 | 618 | 1467340 | 1467957 | BCGDan_sapMKO_1367 | BCGDan_sapMKO_1367 | 618 |
| 1469671 | 1470252 | Rv1314c | Rv1314c | 582 | 1496185 | 1496766 | BCG_1375c | BCG_1375c | 582 | 1468007 | 1468588 | BCGDan_1369 | BCGDan_1369 | 582 | 1468123 | 1468704 | BCGDan_sapMKO_1368 | BCGDan_sapMKO_1368 | 582 |
| 1470321 | 1471577 | murA | Rv1315 | 1257 | 1496835 | 1498091 | murA | BCG_1376 | 1257 | 1468657 | 1469913 | murA | BCGDan_1370 | 1257 | 1468773 | 1470029 | murA | BCGDan_sapMKO_1369 | 1257 |
| 1477134 | 1477631 | ogt | Rv1316c | 498 | 1503648 | 1504145 | ogt | BCG_1377c | 498 | 1475470 | 1475967 | ogt | BCGDan_1374 | 498 | 1475586 | 1476083 | ogt | BCGDan_sapMKO_1373 | 498 |
| 1477628 | 1479118 | alkA | Rv1317c | 1491 | 1504142 | 1505632 | alkA | BCG_1378c | 1491 | 1475964 | 1477454 | alkA | BCGDan_1375 | 1491 | 1476080 | 1477570 | alkA | BCGDan_sapMKO_1374 | 1491 |
| 1479199 | 1480824 | Rv1318c | Rv1318c | 1626 | 1505713 | 1507338 | BCG_1379c | BCG_1379c | 1626 | 1477535 | 1479160 | BCGDan_1376 | BCGDan_1376 | 1626 | 1477651 | 1479276 | BCGDan_sapMKO_1375 | BCGDan_sapMKO_1375 | 1626 |
| 1480894 | 1482501 | Rv1319c | Rv1319c | 1608 | 1507408 | 1509015 | BCG_1380c | BCG_1380c | 1608 | 1479230 | 1480837 | BCGDan_1377 | BCGDan_1377 | 1608 | 1479346 | 1480953 | BCGDan_sapMKO_1376 | BCGDan_sapMKO_1376 | 1608 |
| 1482514 | 1484217 | Rv1320c | Rv1320c | 1704 | 1509028 | 1510731 | BCG_1381c | BCG_1381c | 1704 | 1480850 | 1482553 | BCGDan_1378 | BCGDan_1378 | 1704 | 1480966 | 1482513 | BCGDan_sapMKO_1377 | BCGDan_sapMKO_1377 | 1548 |
| 1484279 | 1484959 | Rv1321 | Rv1321 | 681 | 1510793 | 1511473 | BCG_1382 | BCG_1382 | 681 | 1482615 | 1483295 | BCGDan_1379 | BCGDan_1379 | 681 | 1482740 | 1483411 | BCGDan_sapMKO_1378 | BCGDan_sapMKO_1378 | 672 |
| 1484982 | 1485278 | Rv1322 | Rv1322 | 297 | 1511496 | 1511792 | BCG_1383 | BCG_1383 | 297 | 1483318 | 1483614 | BCGDan_1380 | BCGDan_1380 | 297 | 1483434 | 1483730 | BCGDan_sapMKO_1379 | BCGDan_sapMKO_1379 | 297 |
| 1485313 | 1485771 | Rv1322A | Rv1322A | 459 | 1511827 | 1512285 | BCG_1384c | BCG_1384c | 459 | 1483649 | 1484107 | BCGDan_1381 | BCGDan_1381 | 459 | 1483765 | 1484220 | BCGDan_sapMKO_1380 | BCGDan_sapMKO_1380 | 456 |
| 1485862 | 1487031 | fadA4 | Rv1323 | 1170 | 1512376 | 1513545 | fadA4 | BCG_1385 | 1170 | 1484198 | 1485367 | fadA4 | BCGDan_1382 | 1170 | 1484314 | 1485483 | fadA4 | BCGDan_sapMKO_1381 | 1170 |
| 1487161 | 1488075 | Rv1324 | Rv1324 | 915 | 1513675 | 1514589 | BCG_1386 | BCG_1386 | 915 | 1485497 | 1486411 | BCGDan_1383 | BCGDan_1383 | 915 | 1485646 | 1486527 | BCGDan_sapMKO_1382 | BCGDan_sapMKO_1382 | 882 |
| 1488154 | 1489965 | PE_PGRS24 | Rv1325c | 1812 | 1514668 | 1516479 | PE_PGRS24 | BCG_1387c | 1812 | 1486490 | 1488301 | PE_PGRS24 | BCGDan_1384 | 1812 | 1486606 | 1486767 | PE_PGRS24 | BCGDan_sapMKO_1383 | 162 |
| 1490117 | 1492312 | glgB | Rv1326c | 2196 | 1516631 | 1518826 | glgB | BCG_1388c | 2196 | 1488453 | 1490648 | glgB | BCGDan_1385 | 2196 | 1488569 | 1490764 | glgB | BCGDan_sapMKO_1384 | 2196 |
| 1492320 | 1494425 | glgE | Rv1327c | 2106 | 1518834 | 1520939 | glgE | BCG_1389c | 2106 | 1490656 | 1492761 | glgE | BCGDan_1386 | 2106 | 1490772 | 1492877 | glgE | BCGDan_sapMKO_1385 | 2106 |
| 1494564 | 1497155 | glgP | Rv1328 | 2592 | 1521078 | 1523669 | glgP | BCG_1390 | 2592 | 1492900 | 1495491 | glgP | BCGDan_1387 | 2592 | 1493016 | 1495607 | glgP | BCGDan_sapMKO_1386 | 2592 |
| 1497195 | 1499189 | dinG | Rv1329c | 1995 | 1523709 | 1525703 | dinG | BCG_1391c | 1995 | 1495531 | 1497525 | dinG | BCGDan_1388 | 1995 | 1495647 | 1497623 | dinG | BCGDan_sapMKO_1387 | 1977 |
| 1499213 | 1500559 | pncB1 | Rv1330c | 1347 | 1525727 | 1527073 | BCG_1392c | BCG_1392c | 1347 | 1497549 | 1498895 | BCGDan_1389 | BCGDan_1389 | 1347 | 1497665 | 1499011 | BCGDan_sapMKO_1388 | BCGDan_sapMKO_1388 | 1347 |
| 1500661 | 1500966 | Rv1331 | Rv1331 | 306 | 1527175 | 1527480 | BCG_1393 | BCG_1393 | 306 | 1498997 | 1499302 | BCGDan_1390 | BCGDan_1390 | 306 | 1499113 | 1499418 | BCGDan_sapMKO_1389 | BCGDan_sapMKO_1389 | 306 |
| 1500926 | 1501582 | Rv1332 | Rv1332 | 657 | 1527440 | 1528096 | BCG_1394 | BCG_1394 | 657 | 1499262 | 1499918 | BCGDan_1391 | BCGDan_1391 | 657 | 1499378 | 1500034 | BCGDan_sapMKO_1390 | BCGDan_sapMKO_1390 | 657 |
| 1501599 | 1502633 | Rv1333 | Rv1333 | 1035 | 1528113 | 1529147 | BCG_1395 | BCG_1395 | 1035 | 1499935 | 1500969 | BCGDan_1392 | BCGDan_1392 | 1035 | 1500051 | 1501085 | BCGDan_sapMKO_1391 | BCGDan_sapMKO_1391 | 1035 |

| | | | | | | | | | | | | | | | | | | | |
|---|---|---|---|---|---|---|---|---|---|---|---|---|---|---|---|---|---|---|---|
| 1502641 | 1503081 | mec | Rv1334 | 441 | 1529155 | 1529595 | BCG_1396 | BCG_1396 | 441 | 1500977 | 1501417 | BCGDan_1393 | BCGDan_1393 | 441 | 1501120 | 1501533 | BCGDan_sapMKO_1392 | BCGDan_sapMKO_1392 | 414 |
| 1503103 | 1503384 | cysO | Rv1335 | 282 | 1529617 | 1529898 | BCG_1397 | BCG_1397 | 282 | 1501439 | 1501720 | BCGDan_1394 | BCGDan_1394 | 282 | 1501555 | 1501836 | BCGDan_sapMKO_1393 | BCGDan_sapMKO_1393 | 282 |
| 1503394 | 1504365 | cysM | Rv1336 | 972 | 1529908 | 1530879 | cysM | BCG_1398 | 972 | 1501730 | 1502701 | cysM | BCGDan_1395 | 972 | 1501846 | 1502817 | cysM | BCGDan_sapMKO_1394 | 972 |
| 1504356 | 1505078 | Rv1337 | Rv1337 | 723 | 1530870 | 1531592 | BCG_1399 | BCG_1399 | 723 | 1502692 | 1503414 | BCGDan_1396 | BCGDan_1396 | 723 | 1502808 | 1503530 | BCGDan_sapMKO_1395 | BCGDan_sapMKO_1395 | 723 |
| 1505075 | 1505890 | murI | Rv1338 | 816 | 1531589 | 1532404 | murI | BCG_1400 | 816 | 1503411 | 1504226 | murI | BCGDan_1397 | 816 | 1503527 | 1504342 | murI | BCGDan_sapMKO_1396 | 816 |
| 1505917 | 1506738 | Rv1339 | Rv1339 | 822 | 1532431 | 1533252 | BCG_1401 | BCG_1401 | 822 | 1504253 | 1505074 | BCGDan_1398 | BCGDan_1398 | 822 | 1504414 | 1505190 | BCGDan_sapMKO_1397 | BCGDan_sapMKO_1397 | 777 |
| 1506755 | 1507534 | rphA | Rv1340 | 780 | 1533269 | 1534048 | rphA | BCG_1402 | 780 | 1505091 | 1505870 | rphA | BCGDan_1399 | 780 | 1505207 | 1505986 | rphA | BCGDan_sapMKO_1398 | 780 |
| 1507573 | 1508187 | Rv1341 | Rv1341 | 615 | 1534087 | 1534701 | BCG_1403 | BCG_1403 | 615 | 1505909 | 1506523 | BCGDan_1400 | BCGDan_1400 | 615 | 1506025 | 1506639 | BCGDan_sapMKO_1399 | BCGDan_sapMKO_1399 | 615 |
| 1508184 | 1508546 | Rv1342c | Rv1342c | 363 | 1534698 | 1535060 | BCG_1404c | BCG_1404c | 363 | 1506520 | 1506882 | BCGDan_1401 | BCGDan_1401 | 363 | 1506636 | 1506998 | BCGDan_sapMKO_1400 | BCGDan_sapMKO_1400 | 363 |
| 1508543 | 1508923 | lprD | Rv1343c | 381 | 1535057 | 1535437 | lprD | BCG_1405c | 381 | 1506879 | 1507259 | lprD | BCGDan_1402 | 381 | 1506995 | 1507375 | lprD | BCGDan_sapMKO_1401 | 381 |
| 1508968 | 1509288 | mbtL | Rv1344 | 321 | 1535482 | 1535802 | BCG_1406 | BCG_1406 | 321 | 1507304 | 1507624 | BCGDan_1403 | BCGDan_1403 | 321 | 1507420 | 1507740 | BCGDan_sapMKO_1402 | BCGDan_sapMKO_1402 | 321 |
| 1509281 | 1510846 | mbtM | Rv1345 | 1566 | 1535795 | 1537360 | fadD33 | BCG_1407 | 1566 | 1507617 | 1509182 | fadD33 | BCGDan_1404 | 1566 | 1507733 | 1509298 | fadD33 | BCGDan_sapMKO_1403 | 1566 |
| 1510846 | 1512006 | mbtN | Rv1346 | 1161 | 1537360 | 1538520 | fadE14 | BCG_1408 | 1161 | 1509182 | 1510342 | fadE14 | BCGDan_1405 | 1161 | 1509298 | 1510458 | fadE14 | BCGDan_sapMKO_1404 | 1161 |
| 1511973 | 1512605 | mbtK | Rv1347c | 633 | 1538487 | 1539119 | BCG_1409c | BCG_1409c | 633 | 1510309 | 1510941 | BCGDan_1406 | BCGDan_1406 | 633 | 1510425 | 1511057 | BCGDan_sapMKO_1405 | BCGDan_sapMKO_1405 | 633 |
| 1513047 | 1515626 | irtA | Rv1348 | 2580 | 1539561 | 1542140 | BCG_1410 | BCG_1410 | 2580 | 1511383 | 1513962 | BCGDan_1408 | BCGDan_1408 | 2580 | 1511499 | 1514078 | BCGDan_sapMKO_1407 | BCGDan_sapMKO_1407 | 2580 |
| 1515623 | 1517362 | irtB | Rv1349 | 1740 | 1542137 | 1543876 | BCG_1411 | BCG_1411 | 1740 | 1513959 | 1515698 | BCGDan_1409 | BCGDan_1409 | 1740 | 1514075 | 1515814 | BCGDan_sapMKO_1408 | BCGDan_sapMKO_1408 | 1740 |
| 1517491 | 1518234 | fabG2 | Rv1350 | 744 | 1544005 | 1544748 | fabG2 | BCG_1412 | 744 | 1515827 | 1516570 | fabG2 | BCGDan_1410 | 744 | 1515943 | 1516686 | fabG2 | BCGDan_sapMKO_1409 | 744 |
| 1518231 | 1518560 | Rv1351 | Rv1351 | 330 | 1544745 | 1545074 | BCG_1413 | BCG_1413 | 330 | 1516567 | 1516896 | BCGDan_1411 | BCGDan_1411 | 330 | 1516683 | 1517012 | BCGDan_sapMKO_1410 | BCGDan_sapMKO_1410 | 330 |
| 1518763 | 1519134 | Rv1352 | Rv1352 | 372 | 1545277 | 1545648 | BCG_1414 | BCG_1414 | 372 | 1517099 | 1517470 | BCGDan_1412 | BCGDan_1412 | 372 | 1517269 | 1517586 | BCGDan_sapMKO_1411 | BCGDan_sapMKO_1411 | 318 |
| 1519200 | 1519985 | Rv1353c | Rv1353c | 786 | 1545714 | 1546499 | BCG_1415c | BCG_1415c | 786 | 1517536 | 1518321 | BCGDan_1413 | BCGDan_1413 | 786 | 1517652 | 1518437 | BCGDan_sapMKO_1412 | BCGDan_sapMKO_1412 | 786 |
| 1520005 | 1521876 | Rv1354c | Rv1354c | 1872 | 1546519 | 1548390 | BCG_1416c | BCG_1416c | 1872 | 1518341 | 1520212 | BCGDan_1414 | BCGDan_1414 | 1872 | 1518457 | 1520328 | BCGDan_sapMKO_1413 | BCGDan_sapMKO_1413 | 1872 |
| 1521885 | 1524032 | moeY | Rv1355c | 2148 | 1548399 | 1550546 | moeY | BCG_1417c | 2148 | 1520221 | 1522368 | moeY | BCGDan_1415 | 2148 | 1520337 | 1522484 | moeY | BCGDan_sapMKO_1414 | 2148 |
| 1524029 | 1524820 | Rv1356c | Rv1356c | 792 | 1550543 | 1551334 | BCG_1418c | BCG_1418c | 792 | 1522365 | 1523156 | BCGDan_1416 | BCGDan_1416 | 792 | 1522481 | 1523272 | BCGDan_sapMKO_1415 | BCGDan_sapMKO_1415 | 792 |
| 1525293 | 1526216 | Rv1357c | Rv1357c | 924 | 1551807 | 1552730 | BCG_1419c | BCG_1419c | 924 | 1523629 | 1524552 | BCGDan_1417 | BCGDan_1417 | 924 | 1523745 | 1524578 | BCGDan_sapMKO_1416 | BCGDan_sapMKO_1416 | 834 |
| 1526612 | 1530091 | Rv1358 | Rv1358 | 3480 | 1553126 | 1556605 | BCG_1420 | BCG_1420 | 3480 | 1524948 | 1528427 | BCGDan_1418 | BCGDan_1418 | 3480 | 1525169 | 1528543 | BCGDan_sapMKO_1417 | BCGDan_sapMKO_1417 | 3375 |
| 1530173 | 1530925 | Rv1359 | Rv1359 | 753 | 1556687 | 1557439 | BCG_1421 | BCG_1421 | 753 | 1528509 | 1529261 | BCGDan_1419 | BCGDan_1419 | 753 | 1528625 | 1529377 | BCGDan_sapMKO_1418 | BCGDan_sapMKO_1418 | 753 |
| 1531348 | 1532370 | Rv1360 | Rv1360 | 1023 | 1557862 | 1558884 | BCG_1422 | BCG_1422 | 1023 | 1529684 | 1530706 | BCGDan_1420 | BCGDan_1420 | 1023 | 1529800 | 1530822 | BCGDan_sapMKO_1419 | BCGDan_sapMKO_1419 | 1023 |
| 1532443 | 1533633 | PPE19 | Rv1361c | 1191 | 1558957 | 1560147 | PPE19 | BCG_1423c | 1191 | 1530779 | 1531969 | PPE19 | BCGDan_1421 | 1191 | 1530895 | 1532085 | PPE19 | BCGDan_sapMKO_1420 | 1191 |
| 1533948 | 1534610 | Rv1362c | Rv1362c | 663 | 1560462 | 1561124 | BCG_1424c | BCG_1424c | 663 | 1532284 | 1532946 | BCGDan_1422 | BCGDan_1422 | 663 | 1532400 | 1533062 | BCGDan_sapMKO_1421 | BCGDan_sapMKO_1421 | 663 |
| 1534607 | 1535392 | Rv1363c | Rv1363c | 786 | 1561121 | 1561906 | BCG_1425c | BCG_1425c | 786 | 1532943 | 1533728 | BCGDan_1423 | BCGDan_1423 | 786 | 1533059 | 1533844 | BCGDan_sapMKO_1422 | BCGDan_sapMKO_1422 | 786 |
| 1535683 | 1537644 | Rv1364c | Rv1364c | 1962 | 1562197 | 1564158 | BCG_1426c | BCG_1426c | 1962 | 1534019 | 1535980 | BCGDan_1424 | BCGDan_1424 | 1962 | 1534135 | 1536096 | BCGDan_sapMKO_1423 | BCGDan_sapMKO_1423 | 1962 |
| 1537783 | 1538169 | rsfA | Rv1365c | 387 | 1564297 | 1564683 | BCG_1427c | BCG_1427c | 387 | 1536119 | 1536505 | BCGDan_1425 | BCGDan_1425 | 387 | 1536235 | 1536585 | BCGDan_sapMKO_1424 | BCGDan_sapMKO_1424 | 351 |
| 1538390 | 1539211 | Rv1366 | Rv1366 | 822 | 1564904 | 1565725 | BCG_1428 | BCG_1428 | 822 | 1536726 | 1537547 | BCGDan_1426 | BCGDan_1426 | 822 | 1536926 | 1537663 | BCGDan_sapMKO_1425 | BCGDan_sapMKO_1425 | 738 |
| 1539512 | 1540645 | Rv1367c | Rv1367c | 1134 | 1566026 | 1567159 | BCG_1429c | BCG_1429c | 1134 | 1537848 | 1538981 | BCGDan_1427 | BCGDan_1427 | 1134 | 1537964 | 1539169 | BCGDan_sapMKO_1426 | BCGDan_sapMKO_1426 | 1206 |
| 1541020 | 1541805 | lprF | Rv1368 | 786 | 1567534 | 1568319 | lprF | BCG_1430 | 786 | 1539356 | 1540141 | lprF | BCGDan_1428 | 786 | 1539472 | 1540257 | lprF | BCGDan_sapMKO_1427 | 786 |
| 1543359 | 1544828 | Rv1371 | Rv1371 | 1470 | 1568515 | 1569468 | BCG_1431 | BCG_1431 | 954 | 1540337 | 1541290 | BCGDan_1429 | BCGDan_1429 | 954 | 1540453 | 1541406 | BCGDan_sapMKO_1428 | BCGDan_sapMKO_1428 | 954 |
| NA | NA | NA | NA | NA | 1568990 | 1569985 | BCG_1432 | BCG_1432 | 996 | 1540812 | 1541807 | BCGDan_1430 | BCGDan_1430 | 996 | 1540928 | 1541923 | BCGDan_sapMKO_1429 | BCGDan_sapMKO_1429 | 996 |
| 1544825 | 1546006 | Rv1372 | Rv1372 | 1182 | 1569982 | 1571163 | BCG_1433 | BCG_1433 | 1182 | 1541804 | 1542985 | BCGDan_1431 | BCGDan_1431 | 1182 | 1541920 | 1543101 | BCGDan_sapMKO_1430 | BCGDan_sapMKO_1430 | 1182 |
| 1546012 | 1546992 | Rv1373 | Rv1373 | 981 | 1571169 | 1572152 | BCG_1434 | BCG_1434 | 984 | 1542991 | 1543974 | BCGDan_1432 | BCGDan_1432 | 984 | 1543107 | 1544090 | BCGDan_sapMKO_1431 | BCGDan_sapMKO_1431 | 984 |
| 1547072 | 1547530 | Rv1374c | Rv1374c | 459 | 1572232 | 1572690 | BCG_1435c | BCG_1435c | 459 | 1544054 | 1544512 | BCGDan_1433 | BCGDan_1433 | 459 | 1544170 | 1544628 | BCGDan_sapMKO_1432 | BCGDan_sapMKO_1432 | 459 |
| 1547832 | 1549151 | Rv1375 | Rv1375 | 1320 | 1572992 | 1574311 | BCG_1436 | BCG_1436 | 1320 | 1544814 | 1546133 | BCGDan_1434 | BCGDan_1434 | 1320 | 1544930 | 1546249 | BCGDan_sapMKO_1433 | BCGDan_sapMKO_1433 | 1320 |
| 1549148 | 1550641 | Rv1376 | Rv1376 | 1494 | 1574308 | 1575801 | BCG_1437 | BCG_1437 | 1494 | 1546130 | 1547623 | BCGDan_1435 | BCGDan_1435 | 1494 | 1546246 | 1547739 | BCGDan_sapMKO_1434 | BCGDan_sapMKO_1434 | 1494 |
| 1550579 | 1551217 | Rv1377c | Rv1377c | 639 | 1575739 | 1576377 | BCG_1438c | BCG_1438c | 639 | 1547561 | 1548199 | BCGDan_1436 | BCGDan_1436 | 639 | 1547677 | 1548315 | BCGDan_sapMKO_1435 | BCGDan_sapMKO_1435 | 639 |
| 1551228 | 1552655 | Rv1378c | Rv1378c | 1428 | 1576388 | 1577815 | BCG_1439c | BCG_1439c | 1428 | 1548210 | 1549637 | BCGDan_1437 | BCGDan_1437 | 1428 | 1548326 | 1549588 | BCGDan_sapMKO_1436 | BCGDan_sapMKO_1436 | 1263 |
| 1552654 | 1553235 | pyrR | Rv1379 | 582 | 1577814 | 1578395 | pyrR | BCG_1440 | 582 | 1549636 | 1550217 | pyrR | BCGDan_1438 | 582 | 1549752 | 1550333 | pyrR | BCGDan_sapMKO_1437 | 582 |
| 1553232 | 1554191 | pyrB | Rv1380 | 960 | 1578392 | 1579351 | pyrB | BCG_1441 | 960 | 1550214 | 1551173 | pyrB | BCGDan_1439 | 960 | 1550330 | 1551289 | pyrB | BCGDan_sapMKO_1438 | 960 |
| 1554188 | 1555480 | pyrC | Rv1381 | 1293 | 1579348 | 1580640 | pyrC | BCG_1442 | 1293 | 1551170 | 1552462 | pyrC | BCGDan_1440 | 1293 | 1551286 | 1552578 | pyrC | BCGDan_sapMKO_1439 | 1293 |
| 1555477 | 1555974 | Rv1382 | Rv1382 | 498 | 1580637 | 1581134 | BCG_1443 | BCG_1443 | 498 | 1552459 | 1552956 | BCGDan_1441 | BCGDan_1441 | 498 | 1552575 | 1553072 | BCGDan_sapMKO_1440 | BCGDan_sapMKO_1440 | 498 |
| 1555971 | 1557101 | carA | Rv1383 | 1131 | 1581131 | 1582261 | carA | BCG_1444 | 1131 | 1552953 | 1554083 | carA | BCGDan_1442 | 1131 | 1553069 | 1554199 | carA | BCGDan_sapMKO_1441 | 1131 |
| 1557101 | 1560448 | carB | Rv1384 | 3348 | 1582261 | 1585608 | carB | BCG_1445 | 3348 | 1554083 | 1557430 | carB | BCGDan_1443 | 3348 | 1554199 | 1557546 | carB | BCGDan_sapMKO_1442 | 3348 |
| 1560445 | 1561269 | pyrF | Rv1385 | 825 | 1585605 | 1586429 | pyrF | BCG_1446 | 825 | 1557427 | 1558251 | pyrF | BCGDan_1444 | 825 | 1557543 | 1558367 | pyrF | BCGDan_sapMKO_1443 | 825 |
| 1561464 | 1561772 | PE15 | Rv1386 | 309 | 1586624 | 1586932 | PE15 | BCG_1447 | 309 | 1558446 | 1558754 | PE15 | BCGDan_1445 | 309 | 1558562 | 1558870 | PE15 | BCGDan_sapMKO_1444 | 309 |
| 1561769 | 1563388 | PPE20 | Rv1387 | 1620 | 1586929 | 1588548 | PPE20 | BCG_1448 | 1620 | 1558751 | 1560370 | PPE20 | BCGDan_1446 | 1620 | 1558867 | 1560486 | PPE20 | BCGDan_sapMKO_1445 | 1620 |
| 1563694 | 1564266 | mihF | Rv1388 | 573 | 1588854 | 1589426 | mihF | BCG_1449 | 573 | 1560676 | 1561248 | mihF | BCGDan_1447 | 573 | 1560792 | 1561364 | mihF | BCGDan_sapMKO_1446 | 573 |
| 1564401 | 1565027 | gmk | Rv1389 | 627 | 1589561 | 1590187 | gmk | BCG_1450 | 627 | 1561383 | 1562009 | gmk | BCGDan_1448 | 627 | 1561499 | 1562125 | gmk | BCGDan_sapMKO_1447 | 627 |
| 1565093 | 1565425 | rpoZ | Rv1390 | 333 | 1590253 | 1590585 | rpoZ | BCG_1451 | 333 | 1562075 | 1562407 | rpoZ | BCGDan_1449 | 333 | 1562191 | 1562523 | rpoZ | BCGDan_sapMKO_1448 | 333 |
| 1565441 | 1566697 | dfp | Rv1391 | 1257 | 1590601 | 1591857 | dfp | BCG_1452 | 1257 | 1562423 | 1563679 | dfp | BCGDan_1450 | 1257 | 1562539 | 1563795 | dfp | BCGDan_sapMKO_1449 | 1257 |
| 1566825 | 1568036 | metK | Rv1392 | 1212 | 1591985 | 1593196 | metK | BCG_1453 | 1212 | 1563807 | 1565018 | metK | BCGDan_1451 | 1212 | 1563914 | 1565134 | metK | BCGDan_sapMKO_1450 | 1221 |
| 1568109 | 1569587 | Rv1393c | Rv1393c | 1479 | 1593269 | 1594747 | BCG_1454c | BCG_1454c | 1479 | 1565091 | 1566569 | BCGDan_1452 | BCGDan_1452 | 1479 | 1565207 | 1566682 | BCGDan_sapMKO_1451 | BCGDan_sapMKO_1451 | 1476 |
| 1569584 | 1570969 | cyp132 | Rv1394c | 1386 | 1594744 | 1596129 | cyp132 | BCG_1455c | 1386 | 1566566 | 1567951 | cyp132 | BCGDan_1453 | 1386 | 1566682 | 1568067 | cyp132 | BCGDan_sapMKO_1452 | 1386 |
| 1571047 | 1572081 | Rv1395 | Rv1395 | 1035 | 1596207 | 1597241 | BCG_1456 | BCG_1456 | 1035 | 1568029 | 1569063 | BCGDan_1454 | BCGDan_1454 | 1035 | 1568184 | 1569179 | BCGDan_sapMKO_1453 | BCGDan_sapMKO_1453 | 996 |
| 1572127 | 1573857 | PE_PGRS25 | Rv1396c | 1731 | 1597287 | 1598645 | PE_PGRS25 | BCG_1457c | 1359 | 1569109 | 1570467 | PE_PGRS25 | BCGDan_1455 | 1359 | 1569225 | 1570583 | PE_PGRS25 | BCGDan_sapMKO_1454 | 1359 |
| 1574112 | 1574513 | vapC10 | Rv1397c | 402 | 1598900 | 1599301 | BCG_1458c | BCG_1458c | 402 | 1570722 | 1571123 | BCGDan_1456 | BCGDan_1456 | 402 | 1570838 | 1571254 | BCGDan_sapMKO_1455 | BCGDan_sapMKO_1455 | 417 |
| 1574510 | 1574767 | vapB10 | Rv1398c | 258 | 1599298 | 1599555 | BCG_1459c | BCG_1459c | 258 | 1571120 | 1571377 | BCGDan_1457 | BCGDan_1457 | 258 | 1571236 | 1571493 | BCGDan_sapMKO_1456 | BCGDan_sapMKO_1456 | 258 |
| 1574850 | 1575809 | nlhH | Rv1399c | 960 | 1599638 | 1600597 | lipH | BCG_1460c | 960 | 1571460 | 1572419 | lipH | BCGDan_1458 | 960 | 1571576 | 1572496 | lipH | BCGDan_sapMKO_1457 | 921 |
| 1575834 | 1576796 | lipI | Rv1400c | 963 | 1600622 | 1601584 | lipI | BCG_1461c | 963 | 1572444 | 1573406 | lipI | BCGDan_1459 | 963 | 1572560 | 1573522 | lipI | BCGDan_sapMKO_1458 | 963 |
| 1576930 | 1577532 | Rv1401 | Rv1401 | 603 | 1601718 | 1602320 | BCG_1462 | BCG_1462 | 603 | 1573540 | 1574142 | BCGDan_1460 | BCGDan_1460 | 603 | 1573656 | 1574258 | BCGDan_sapMKO_1459 | BCGDan_sapMKO_1459 | 603 |
| 1577613 | 1579580 | priA | Rv1402 | 1968 | 1602401 | 1604368 | priA | BCG_1463 | 1968 | 1574223 | 1576190 | priA | BCGDan_1461 | 1968 | 1574330 | 1576306 | priA | BCGDan_sapMKO_1460 | 1977 |
| 1579598 | 1580422 | Rv1403c | Rv1403c | 825 | 1604386 | 1605210 | BCG_1464c | BCG_1464c | 825 | 1576208 | 1577032 | BCGDan_1462 | BCGDan_1462 | 825 | 1576324 | 1577148 | BCGDan_sapMKO_1461 | BCGDan_sapMKO_1461 | 825 |
| 1580591 | 1581073 | Rv1404 | Rv1404 | 483 | 1605379 | 1605861 | BCG_1465 | BCG_1465 | 483 | 1577201 | 1577683 | BCGDan_1463 | BCGDan_1463 | 483 | 1577316 | 1577798 | BCGDan_sapMKO_1462 | BCGDan_sapMKO_1462 | 483 |
| 1581145 | 1581969 | Rv1405c | Rv1405c | 825 | 1605933 | 1606757 | BCG_1466c | BCG_1466c | 825 | 1577755 | 1578579 | BCGDan_1464 | BCGDan_1464 | 825 | 1577870 | 1578694 | BCGDan_sapMKO_1463 | BCGDan_sapMKO_1463 | 825 |
| 1582166 | 1583104 | fmt | Rv1406 | 939 | 1606954 | 1607892 | fmt | BCG_1467 | 939 | 1578776 | 1579714 | fmt | BCGDan_1465 | 939 | 1578891 | 1579829 | fmt | BCGDan_sapMKO_1464 | 939 |
| 1583101 | 1584474 | fmu | Rv1407 | 1374 | 1607889 | 1609262 | fmu | BCG_1468 | 1374 | 1579711 | 1581084 | fmu | BCGDan_1466 | 1374 | 1579826 | 1581199 | fmu | BCGDan_sapMKO_1465 | 1374 |
| 1584499 | 1585197 | rpe | Rv1408 | 699 | 1609287 | 1609985 | rpe | BCG_1469 | 699 | 1581109 | 1581807 | rpe | BCGDan_1467 | 699 | 1581233 | 1581922 | rpe | BCGDan_sapMKO_1466 | 690 |
| 1585194 | 1586213 | ribG | Rv1409 | 1020 | 1609982 | 1611001 | ribG | BCG_1470 | 1020 | 1581804 | 1582823 | ribG | BCGDan_1468 | 1020 | 1581919 | 1582938 | ribG | BCGDan_sapMKO_1467 | 1020 |
| 1586210 | 1587766 | Rv1410c | Rv1410c | 1557 | 1610998 | 1612554 | BCG_1471c | BCG_1471c | 1557 | 1582820 | 1584376 | BCGDan_1469 | BCGDan_1469 | 1557 | 1582935 | 1584539 | BCGDan_sapMKO_1468 | BCGDan_sapMKO_1468 | 1605 |
| 1587772 | 1588482 | lprG | Rv1411c | 711 | 1612560 | 1613270 | lprG | BCG_1472c | 711 | 1584382 | 1585092 | lprG | BCGDan_1470 | 711 | 1584497 | 1585216 | lprG | BCGDan_sapMKO_1469 | 720 |
| 1588567 | 1589172 | ribC | Rv1412 | 606 | 1613355 | 1613960 | ribC | BCG_1473 | 606 | 1585177 | 1585782 | ribC | BCGDan_1471 | 606 | 1585292 | 1585897 | ribC | BCGDan_sapMKO_1470 | 606 |
| 1589386 | 1589901 | Rv1413 | Rv1413 | 516 | 1614174 | 1614689 | BCG_1474 | BCG_1474 | 516 | 1585996 | 1586511 | BCGDan_1472 | BCGDan_1472 | 516 | 1586111 | 1586626 | BCGDan_sapMKO_1471 | BCGDan_sapMKO_1471 | 516 |
| 1589891 | 1590292 | Rv1414 | Rv1414 | 402 | 1614679 | 1615080 | BCG_1475 | BCG_1475 | 402 | 1586501 | 1586902 | BCGDan_1473 | BCGDan_1473 | 402 | 1586616 | 1587017 | BCGDan_sapMKO_1472 | BCGDan_sapMKO_1472 | 402 |
| 1590397 | 1591674 | ribA2 | Rv1415 | 1278 | 1615185 | 1616462 | ribA2 | BCG_1476 | 1278 | 1587007 | 1588284 | ribA2 | BCGDan_1474 | 1278 | 1587122 | 1588399 | ribA2 | BCGDan_sapMKO_1473 | 1278 |
| 1591671 | 1592153 | ribH | Rv1416 | 483 | 1616477 | 1616941 | ribH | BCG_1477 | 465 | 1588281 | 1588763 | ribH | BCGDan_1475 | 483 | 1588396 | 1588878 | ribH | BCGDan_sapMKO_1474 | 483 |
| 1592150 | 1592614 | Rv1417 | Rv1417 | 465 | 1616938 | 1617402 | BCG_1478 | BCG_1478 | 465 | 1588760 | 1589224 | BCGDan_1476 | BCGDan_1476 | 465 | 1588875 | 1589339 | BCGDan_sapMKO_1475 | BCGDan_sapMKO_1475 | 465 |
| 1592639 | 1593325 | lprH | Rv1418 | 687 | 1617427 | 1618113 | lprH | BCG_1479 | 687 | 1589249 | 1589935 | lprH | BCGDan_1477 | 687 | 1589421 | 1590050 | lprH | BCGDan_sapMKO_1476 | 630 |
| 1593505 | 1593978 | Rv1419 | Rv1419 | 474 | 1618293 | 1618766 | BCG_1480 | BCG_1480 | 474 | 1590115 | 1590588 | BCGDan_1478 | BCGDan_1478 | 474 | 1590230 | 1590703 | BCGDan_sapMKO_1477 | BCGDan_sapMKO_1477 | 474 |
| 1594042 | 1595982 | uvrC | Rv1420 | 1941 | 1618830 | 1620770 | uvrC | BCG_1481 | 1941 | 1590652 | 1592592 | uvrC | BCGDan_1479 | 1941 | 1590824 | 1592707 | uvrC | BCGDan_sapMKO_1478 | 1884 |

| | | | | | | | | | | | | | | | | | | | |
|---|---|---|---|---|---|---|---|---|---|---|---|---|---|---|---|---|---|---|---|
| 1595979 | 1596884 | Rv1421 | Rv1421 | 906 | 1620767 | 1621672 | BCG_1482 | BCG_1482 | 906 | 1592589 | 1593494 | BCGDan_1480 | BCGDan_1480 | 906 | 1592707 | 1593609 | BCGDan_sapMKO_1479 | BCGDan_sapMKO_1479 | 903 |
| 1596881 | 1597909 | Rv1422 | Rv1422 | 1029 | 1621669 | 1622697 | BCG_1483 | BCG_1483 | 1029 | 1593491 | 1594519 | BCGDan_1481 | BCGDan_1481 | 1029 | 1593606 | 1594634 | BCGDan_sapMKO_1480 | BCGDan_sapMKO_1480 | 1029 |
| 1597906 | 1598883 | whiA | Rv1423 | 978 | 1622694 | 1623671 | whiA | BCG_1484 | 978 | 1594516 | 1595493 | whiA | BCGDan_1482 | 978 | 1594625 | 1595608 | whiA | BCGDan_sapMKO_1481 | 984 |
| 1598893 | 1599654 | Rv1424c | Rv1424c | 762 | 1623681 | 1624442 | BCG_1485c | BCG_1485c | 762 | 1595503 | 1596264 | BCGDan_1483 | BCGDan_1483 | 762 | 1595618 | 1596379 | BCGDan_sapMKO_1482 | BCGDan_sapMKO_1482 | 762 |
| 1599658 | 1601037 | Rv1425 | Rv1425 | 1380 | 1624446 | 1625825 | BCG_1486 | BCG_1486 | 1380 | 1596268 | 1597647 | BCGDan_1484 | BCGDan_1484 | 1380 | 1596443 | 1597762 | BCGDan_sapMKO_1483 | BCGDan_sapMKO_1483 | 1320 |
| 1601059 | 1602321 | lipO | Rv1426c | 1263 | 1625847 | 1627109 | lipO | BCG_1487c | 1263 | 1597669 | 1598931 | lipO | BCGDan_1485 | 1263 | 1597784 | 1598998 | lipO | BCGDan_sapMKO_1484 | 1215 |
| 1602321 | 1603928 | fadD12 | Rv1427c | 1608 | 1627109 | 1628716 | fadD12 | BCG_1488c | 1608 | 1598931 | 1600538 | fadD12 | BCGDan_1486 | 1608 | 1599046 | 1600653 | fadD12 | BCGDan_sapMKO_1485 | 1608 |
| 1603932 | 1604759 | Rv1428c | Rv1428c | 828 | 1628720 | 1629547 | BCG_1489c | BCG_1489c | 828 | 1600542 | 1601369 | BCGDan_1487 | BCGDan_1487 | 828 | 1600657 | 1601391 | BCGDan_sapMKO_1486 | BCGDan_sapMKO_1486 | 735 |
| 1604878 | 1606146 | Rv1429 | Rv1429 | 1269 | 1629666 | 1630934 | BCG_1490 | BCG_1490 | 1269 | 1601488 | 1602756 | BCGDan_1488 | BCGDan_1488 | 1269 | 1601648 | 1602871 | BCGDan_sapMKO_1487 | BCGDan_sapMKO_1487 | 1224 |
| 1606386 | 1607972 | PE16 | Rv1430 | 1587 | 1631174 | 1632760 | PE16 | BCG_1491 | 1587 | 1602996 | 1604582 | PE16 | BCGDan_1489 | 1587 | 1603111 | 1604697 | PE16 | BCGDan_sapMKO_1488 | 1587 |
| 1608083 | 1609852 | Rv1431 | Rv1431 | 1770 | 1632871 | 1634640 | BCG_1492 | BCG_1492 | 1770 | 1604693 | 1606462 | BCGDan_1490 | BCGDan_1490 | 1770 | 1604808 | 1606577 | BCGDan_sapMKO_1489 | BCGDan_sapMKO_1489 | 1770 |
| 1609849 | 1611270 | Rv1432 | Rv1432 | 1422 | 1634637 | 1636058 | BCG_1493 | BCG_1493 | 1422 | 1606459 | 1607880 | BCGDan_1491 | BCGDan_1491 | 1422 | 1606574 | 1607995 | BCGDan_sapMKO_1490 | BCGDan_sapMKO_1490 | 1422 |
| 1611434 | 1612249 | Rv1433 | Rv1433 | 816 | 1636222 | 1637043 | BCG_1494 | BCG_1494 | 822 | 1608044 | 1608865 | BCGDan_1492 | BCGDan_1492 | 822 | 1608159 | 1608980 | BCGDan_sapMKO_1491 | BCGDan_sapMKO_1491 | 822 |
| 1612256 | 1612393 | Rv1434 | Rv1434 | 138 | 1637043 | 1637180 | BCG_1495 | BCG_1495 | 138 | 1608865 | 1609002 | BCGDan_1493 | BCGDan_1493 | 138 | 1608980 | 1609117 | BCGDan_sapMKO_1492 | BCGDan_sapMKO_1492 | 138 |
| 1612342 | 1612950 | Rv1435c | Rv1435c | 609 | 1637129 | 1637758 | BCG_1496c | BCG_1496c | 630 | 1608951 | 1609580 | BCGDan_1494 | BCGDan_1494 | 630 | 1609066 | 1609686 | BCGDan_sapMKO_1493 | BCGDan_sapMKO_1493 | 621 |
| 1613307 | 1614326 | gap | Rv1436 | 1020 | 1638115 | 1639134 | gap | BCG_1497 | 1020 | 1609937 | 1610956 | gap | BCGDan_1495 | 1020 | 1610052 | 1611071 | gap | BCGDan_sapMKO_1494 | 1020 |
| 1614329 | 1615567 | pgk | Rv1437 | 1239 | 1639137 | 1640375 | pgk | BCG_1498 | 1239 | 1610959 | 1612197 | pgk | BCGDan_1496 | 1239 | 1611074 | 1612312 | pgk | BCGDan_sapMKO_1495 | 1239 |
| 1615564 | 1616349 | tpi | Rv1438 | 786 | 1640372 | 1641157 | tpi | BCG_1499 | 786 | 1612194 | 1612979 | tpi | BCGDan_1497 | 786 | 1612309 | 1613094 | tpi | BCGDan_sapMKO_1496 | 786 |
| 1616961 | 1617386 | Rv1439c | Rv1439c | 426 | 1641769 | 1642194 | BCG_1500c | BCG_1500c | 426 | 1613591 | 1614016 | BCGDan_1498 | BCGDan_1498 | 426 | 1613706 | 1614131 | BCGDan_sapMKO_1497 | BCGDan_sapMKO_1497 | 426 |
| 1617837 | 1618070 | secG | Rv1440 | 234 | 1642645 | 1642878 | secG | BCG_1501 | 234 | 1614467 | 1614700 | secG | BCGDan_1499 | 234 | 1614582 | 1614815 | secG | BCGDan_sapMKO_1498 | 234 |
| 1618209 | 1619684 | PE_PGRS26 | Rv1441c | 1476 | 1643017 | 1644492 | PE_PGRS26 | BCG_1502c | 1476 | 1614839 | 1616314 | PE_PGRS26 | BCGDan_1500 | 1476 | 1614954 | 1616429 | PE_PGRS26 | BCGDan_sapMKO_1499 | 1476 |
| 1619791 | 1622091 | bisC | Rv1442 | 2301 | 1644599 | 1646899 | bisC | BCG_1503 | 2301 | 1616421 | 1618721 | bisC | BCGDan_1501 | 2301 | 1616536 | 1618836 | bisC | BCGDan_sapMKO_1500 | 2301 |
| 1622207 | 1622692 | Rv1443c | Rv1443c | 486 | 1647015 | 1647500 | BCG_1504c | BCG_1504c | 486 | 1618837 | 1619322 | BCGDan_1502 | BCGDan_1502 | 486 | 1618952 | 1619437 | BCGDan_sapMKO_1501 | BCGDan_sapMKO_1501 | 486 |
| 1623287 | 1623697 | Rv1444c | Rv1444c | 411 | 1648099 | 1648509 | BCG_1505c | BCG_1505c | 411 | 1619929 | 1620339 | BCGDan_1503 | BCGDan_1503 | 411 | 1620041 | 1620400 | BCGDan_sapMKO_1502 | BCGDan_sapMKO_1502 | 360 |
| 1623714 | 1624457 | devB | Rv1445c | 744 | 1648526 | 1649269 | devB | BCG_1506c | 744 | 1620356 | 1621099 | devB | BCGDan_1504 | 744 | 1620468 | 1621211 | devB | BCGDan_sapMKO_1503 | 744 |
| 1624454 | 1625365 | opcA | Rv1446c | 912 | 1649266 | 1650177 | opcA | BCG_1507c | 912 | 1621096 | 1622007 | opcA | BCGDan_1505 | 912 | 1621208 | 1622119 | opcA | BCGDan_sapMKO_1504 | 912 |
| 1625418 | 1626962 | zwf2 | Rv1447c | 1545 | 1650230 | 1651774 | zwf2 | BCG_1508c | 1545 | 1622060 | 1623604 | zwf2 | BCGDan_1506 | 1545 | 1622172 | 1623716 | zwf2 | BCGDan_sapMKO_1505 | 1545 |
| 1626959 | 1628080 | tal | Rv1448c | 1122 | 1651771 | 1652892 | tal | BCG_1509c | 1122 | 1623601 | 1624722 | tal | BCGDan_1507 | 1122 | 1623713 | 1624834 | tal | BCGDan_sapMKO_1506 | 1122 |
| 1628097 | 1630199 | tkt | Rv1449c | 2103 | 1652909 | 1655011 | tkt | BCG_1510c | 2103 | 1624739 | 1626841 | tkt | BCGDan_1508 | 2103 | 1624851 | 1626953 | tkt | BCGDan_sapMKO_1507 | 2103 |
| 1630638 | 1634627 | PE_PGRS27 | Rv1450c | 3990 | 1655450 | 1659649 | PE_PGRS27 | BCG_1511c | 4200 | 1627280 | 1631479 | PE_PGRS27 | BCGDan_1509 | 4200 | 1627392 | 1631594 | PE_PGRS27 | BCGDan_sapMKO_1508 | 4203 |
| 1635029 | 1635955 | ctaB | Rv1451 | 927 | 1660051 | 1660977 | ctaB | BCG_1512 | 927 | 1631881 | 1632807 | ctaB | BCGDan_1510 | 927 | 1632041 | 1632919 | ctaB | BCGDan_sapMKO_1509 | 879 |
| 1636004 | 1638229 | PE_PGRS28 | Rv1452c | 2226 | 1661026 | 1663248 | PE_PGRS28 | BCG_1513c | 2223 | 1632856 | 1635078 | PE_PGRS28 | BCGDan_1511 | 2223 | 1632968 | 1635190 | PE_PGRS28 | BCGDan_sapMKO_1510 | 2223 |
| 1638381 | 1639646 | Rv1453 | Rv1453 | 1266 | 1663400 | 1664665 | BCG_1514 | BCG_1514 | 1266 | 1635230 | 1636495 | BCGDan_1512 | BCGDan_1512 | 1266 | 1635342 | 1636607 | BCGDan_sapMKO_1511 | BCGDan_sapMKO_1511 | 1266 |
| 1639674 | 1640660 | qor | Rv1454c | 987 | 1664693 | 1665679 | qor | BCG_1515c | 987 | 1636523 | 1637509 | qor | BCGDan_1513 | 987 | 1636635 | 1637621 | qor | BCGDan_sapMKO_1512 | 987 |
| 1640680 | 1641543 | Rv1455 | Rv1455 | 864 | 1665699 | 1666562 | BCG_1516 | BCG_1516 | 864 | 1637529 | 1638392 | BCGDan_1514 | BCGDan_1514 | 864 | 1637641 | 1638504 | BCGDan_sapMKO_1513 | BCGDan_sapMKO_1513 | 864 |
| 1641493 | 1642425 | Rv1456c | Rv1456c | 933 | 1666512 | 1667444 | BCG_1517c | BCG_1517c | 933 | 1638342 | 1639274 | BCGDan_1515 | BCGDan_1515 | 933 | 1638454 | 1639386 | BCGDan_sapMKO_1514 | BCGDan_sapMKO_1514 | 933 |
| 1642537 | 1643322 | Rv1457c | Rv1457c | 786 | 1667556 | 1668341 | BCG_1518c | BCG_1518c | 786 | 1639386 | 1640171 | BCGDan_1516 | BCGDan_1516 | 786 | 1639498 | 1640283 | BCGDan_sapMKO_1515 | BCGDan_sapMKO_1515 | 786 |
| 1643319 | 1644260 | Rv1458c | Rv1458c | 942 | 1668338 | 1669279 | BCG_1519c | BCG_1519c | 942 | 1640168 | 1641109 | BCGDan_1517 | BCGDan_1517 | 942 | 1640280 | 1641122 | BCGDan_sapMKO_1516 | BCGDan_sapMKO_1516 | 843 |
| 1644363 | 1646138 | Rv1459c | Rv1459c | 1776 | 1669435 | 1671210 | BCG_1520c | BCG_1520c | 1776 | 1641265 | 1643040 | BCGDan_1518 | BCGDan_1518 | 1776 | 1641376 | 1643151 | BCGDan_sapMKO_1517 | BCGDan_sapMKO_1517 | 1776 |
| 1646186 | 1646992 | Rv1460 | Rv1460 | 807 | 1671258 | 1672064 | BCG_1521 | BCG_1521 | 807 | 1643088 | 1643894 | BCGDan_1519 | BCGDan_1519 | 807 | 1643271 | 1644005 | BCGDan_sapMKO_1518 | BCGDan_sapMKO_1518 | 735 |
| 1646989 | 1649529 | Rv1461 | Rv1461 | 2541 | 1672061 | 1674601 | BCG_1522 | BCG_1522 | 2541 | 1643891 | 1646431 | BCGDan_1520 | BCGDan_1520 | 2541 | 1644002 | 1646542 | BCGDan_sapMKO_1519 | BCGDan_sapMKO_1519 | 2541 |
| 1649526 | 1650719 | Rv1462 | Rv1462 | 1194 | 1674598 | 1675791 | BCG_1523 | BCG_1523 | 1194 | 1646428 | 1647621 | BCGDan_1521 | BCGDan_1521 | 1194 | 1646539 | 1647732 | BCGDan_sapMKO_1520 | BCGDan_sapMKO_1520 | 1194 |
| 1650716 | 1651516 | Rv1463 | Rv1463 | 801 | 1675788 | 1676588 | BCG_1524 | BCG_1524 | 801 | 1647618 | 1648418 | BCGDan_1522 | BCGDan_1522 | 801 | 1647729 | 1648529 | BCGDan_sapMKO_1521 | BCGDan_sapMKO_1521 | 801 |
| 1651518 | 1652771 | csd | Rv1464 | 1254 | 1676590 | 1677843 | csd | BCG_1525 | 1254 | 1648420 | 1649673 | csd | BCGDan_1523 | 1254 | 1648531 | 1649784 | csd | BCGDan_sapMKO_1522 | 1254 |
| 1652768 | 1653256 | Rv1465 | Rv1465 | 489 | 1677840 | 1678328 | BCG_1526 | BCG_1526 | 489 | 1649670 | 1650158 | BCGDan_1524 | BCGDan_1524 | 489 | 1649781 | 1650269 | BCGDan_sapMKO_1523 | BCGDan_sapMKO_1523 | 489 |
| 1653231 | 1653578 | Rv1466 | Rv1466 | 348 | 1678303 | 1678650 | BCG_1527 | BCG_1527 | 348 | 1650133 | 1650480 | BCGDan_1525 | BCGDan_1525 | 348 | 1650244 | 1650591 | BCGDan_sapMKO_1524 | BCGDan_sapMKO_1524 | 348 |
| 1653673 | 1655502 | fadE15 | Rv1467c | 1830 | 1678745 | 1680574 | fadE15 | BCG_1528c | 1830 | 1650575 | 1652404 | fadE15 | BCGDan_1526 | 1830 | 1650686 | 1652515 | fadE15 | BCGDan_sapMKO_1525 | 1830 |
| 1655609 | 1656721 | PE_PGRS29 | Rv1468c | 1113 | 1680681 | 1681793 | PE_PGRS29 | BCG_1529c | 1113 | 1652511 | 1653623 | PE_PGRS29 | BCGDan_1527 | 1113 | 1652622 | 1653734 | PE_PGRS29 | BCGDan_sapMKO_1526 | 1113 |
| 1656963 | 1658936 | ctpD | Rv1469 | 1974 | 1682035 | 1684008 | ctpD | BCG_1530 | 1974 | 1653865 | 1655838 | ctpD | BCGDan_1528 | 1974 | 1654051 | 1655949 | ctpD | BCGDan_sapMKO_1527 | 1899 |
| 1658980 | 1659354 | trxA | Rv1470 | 375 | 1684052 | 1684426 | trxA | BCG_1531 | 375 | 1655882 | 1656256 | trxA | BCGDan_1529 | 375 | 1655993 | 1656367 | trxA | BCGDan_sapMKO_1528 | 375 |
| 1659370 | 1659741 | trxB1 | Rv1471 | 372 | 1684442 | 1684813 | trxB1 | BCG_1532 | 372 | 1656272 | 1656643 | trxB1 | BCGDan_1530 | 372 | 1656383 | 1656754 | trxB1 | BCGDan_sapMKO_1529 | 372 |
| 1659763 | 1660620 | echA12 | Rv1472 | 858 | 1684835 | 1685692 | echA12 | BCG_1533 | 858 | 1656665 | 1657522 | echA12 | BCGDan_1531 | 858 | 1656824 | 1657633 | echA12 | BCGDan_sapMKO_1530 | 810 |
| 1660656 | 1662284 | Rv1473 | Rv1473 | 1629 | 1685728 | 1687356 | BCG_1534 | BCG_1534 | 1629 | 1657558 | 1659186 | BCGDan_1532 | BCGDan_1532 | 1629 | 1657669 | 1659297 | BCGDan_sapMKO_1531 | BCGDan_sapMKO_1531 | 1629 |
| 1662381 | 1662572 | Rv1473A | Rv1473A | 192 | 1687453 | 1687644 | BCG_1535 | BCG_1535 | 192 | 1659283 | 1659474 | BCGDan_1533 | BCGDan_1533 | 192 | 1659394 | 1659585 | BCGDan_sapMKO_1532 | BCGDan_sapMKO_1532 | 192 |
| 1662641 | 1663204 | Rv1474c | Rv1474c | 564 | 1687713 | 1688276 | BCG_1536c | BCG_1536c | 564 | 1659543 | 1660106 | BCGDan_1534 | BCGDan_1534 | 564 | 1659654 | 1660217 | BCGDan_sapMKO_1533 | BCGDan_sapMKO_1533 | 564 |
| 1663215 | 1666046 | acn | Rv1475c | 2832 | 1688287 | 1691118 | acn | BCG_1537c | 2832 | 1660117 | 1662948 | acn | BCGDan_1535 | 2832 | 1660228 | 1663059 | acn | BCGDan_sapMKO_1534 | 2832 |
| 1666204 | 1666764 | Rv1476 | Rv1476 | 561 | 1691276 | 1691836 | BCG_1538 | BCG_1538 | 561 | 1663106 | 1663666 | BCGDan_1536 | BCGDan_1536 | 561 | 1663217 | 1663777 | BCGDan_sapMKO_1535 | BCGDan_sapMKO_1535 | 561 |
| 1666990 | 1668408 | ripA | Rv1477 | 1419 | 1692062 | 1693480 | BCG_1539 | BCG_1539 | 1419 | 1663892 | 1665310 | BCGDan_1537 | BCGDan_1537 | 1419 | 1664081 | 1665421 | BCGDan_sapMKO_1536 | BCGDan_sapMKO_1536 | 1341 |
| 1668419 | 1669144 | Rv1478 | Rv1478 | 726 | 1693491 | 1694216 | BCG_1540 | BCG_1540 | 726 | 1665321 | 1666046 | BCGDan_1538 | BCGDan_1538 | 726 | 1665432 | 1666157 | BCGDan_sapMKO_1537 | BCGDan_sapMKO_1537 | 726 |
| 1669283 | 1670416 | moxR1 | Rv1479 | 1134 | 1694355 | 1695488 | moxR1 | BCG_1541 | 1134 | 1666185 | 1667318 | moxR1 | BCGDan_1539 | 1134 | 1666296 | 1667429 | moxR1 | BCGDan_sapMKO_1538 | 1134 |
| 1670413 | 1671366 | Rv1480 | Rv1480 | 954 | 1695485 | 1696438 | BCG_1542 | BCG_1542 | 954 | 1667315 | 1668268 | BCGDan_1540 | BCGDan_1540 | 954 | 1667426 | 1668379 | BCGDan_sapMKO_1539 | BCGDan_sapMKO_1539 | 954 |
| 1671377 | 1672384 | Rv1481 | Rv1481 | 1008 | 1696449 | 1697456 | BCG_1543 | BCG_1543 | 1008 | 1668279 | 1669286 | BCGDan_1541 | BCGDan_1541 | 1008 | 1668390 | 1669397 | BCGDan_sapMKO_1540 | BCGDan_sapMKO_1540 | 1008 |
| 1672457 | 1673299 | Rv1482c | Rv1482c | 843 | 1697529 | 1698371 | BCG_1544c | BCG_1544c | 843 | 1669359 | 1670201 | BCGDan_1542 | BCGDan_1542 | 843 | 1669470 | 1670312 | BCGDan_sapMKO_1541 | BCGDan_sapMKO_1541 | 843 |
| 1673440 | 1674183 | fabG1 | Rv1483 | 744 | 1698512 | 1699255 | fabG1 | BCG_1545 | 744 | 1670342 | 1671085 | fabG1 | BCGDan_1543 | 744 | 1670453 | 1671196 | fabG1 | BCGDan_sapMKO_1542 | 744 |
| 1674202 | 1675011 | inhA | Rv1484 | 810 | 1699274 | 1700083 | inhA | BCG_1546 | 810 | 1671104 | 1671913 | inhA | BCGDan_1544 | 810 | 1671215 | 1672024 | inhA | BCGDan_sapMKO_1543 | 810 |
| 1675017 | 1676051 | hemZ | Rv1485 | 1035 | 1700089 | 1701123 | hemZ | BCG_1547 | 1035 | 1671919 | 1672953 | hemZ | BCGDan_1545 | 1035 | 1672030 | 1673064 | hemZ | BCGDan_sapMKO_1544 | 1035 |
| 1676017 | 1676883 | Rv1486c | Rv1486c | 867 | 1700985 | 1701956 | BCG_1548c | BCG_1548c | 972 | 1672815 | 1673786 | BCGDan_1546 | BCGDan_1546 | 972 | 1672926 | 1673897 | BCGDan_sapMKO_1545 | BCGDan_sapMKO_1545 | 972 |
| 1676941 | 1677375 | Rv1487 | Rv1487 | 435 | 1702014 | 1702448 | BCG_1549 | BCG_1549 | 435 | 1673844 | 1674278 | BCGDan_1547 | BCGDan_1547 | 435 | 1673955 | 1674389 | BCGDan_sapMKO_1546 | BCGDan_sapMKO_1546 | 435 |
| 1677397 | 1678542 | Rv1488 | Rv1488 | 1146 | 1702470 | 1703615 | BCG_1550 | BCG_1550 | 1146 | 1674300 | 1675445 | BCGDan_1548 | BCGDan_1548 | 1146 | 1674435 | 1675547 | BCGDan_sapMKO_1547 | BCGDan_sapMKO_1547 | 1113 |
| 1678552 | 1678908 | Rv1489 | Rv1489 | 357 | 1703625 | 1703981 | BCG_1551 | BCG_1551 | 357 | 1675455 | 1675811 | BCGDan_1549 | BCGDan_1549 | 357 | 1675565 | 1675921 | BCGDan_sapMKO_1548 | BCGDan_sapMKO_1548 | 357 |
| 1678942 | 1679172 | Rv1489A | Rv1489A | 231 | 1704015 | 1704245 | BCG_1552 | BCG_1552 | 231 | 1675845 | 1676075 | BCGDan_1550 | BCGDan_1550 | 231 | 1675955 | 1676185 | BCGDan_sapMKO_1549 | BCGDan_sapMKO_1549 | 231 |
| 1679322 | 1680629 | Rv1490 | Rv1490 | 1308 | 1704395 | 1705702 | BCG_1553 | BCG_1553 | 1308 | 1676225 | 1677532 | BCGDan_1551 | BCGDan_1551 | 1308 | 1676335 | 1677642 | BCGDan_sapMKO_1550 | BCGDan_sapMKO_1550 | 1308 |
| 1681208 | 1681966 | Rv1491c | Rv1491c | 759 | 1706281 | 1707039 | BCG_1554c | BCG_1554c | 759 | 1678111 | 1678869 | BCGDan_1552 | BCGDan_1552 | 759 | 1678221 | 1678979 | BCGDan_sapMKO_1551 | BCGDan_sapMKO_1551 | 759 |
| 1682157 | 1684004 | mutA | Rv1492 | 1848 | 1707230 | 1709077 | mutA | BCG_1555 | 1848 | 1679060 | 1680907 | mutA | BCGDan_1553 | 1848 | 1679146 | 1681017 | mutA | BCGDan_sapMKO_1552 | 1872 |
| 1684005 | 1686257 | mutB | Rv1493 | 2253 | 1709078 | 1711330 | mutB | BCG_1556 | 2253 | 1680908 | 1683160 | mutB | BCGDan_1554 | 2253 | 1681018 | 1683270 | mutB | BCGDan_sapMKO_1553 | 2253 |
| 1686271 | 1686573 | mazE4 | Rv1494 | 303 | 1711344 | 1711646 | BCG_1557 | BCG_1557 | 303 | 1683174 | 1683476 | BCGDan_1555 | BCGDan_1555 | 303 | 1683281 | 1683586 | BCGDan_sapMKO_1554 | BCGDan_sapMKO_1554 | 306 |
| 1686570 | 1686887 | mazF4 | Rv1495 | 318 | 1711643 | 1711960 | BCG_1558 | BCG_1558 | 318 | 1683473 | 1683790 | BCGDan_1556 | BCGDan_1556 | 318 | 1683583 | 1683900 | BCGDan_sapMKO_1555 | BCGDan_sapMKO_1555 | 318 |
| 1686884 | 1687888 | Rv1496 | Rv1496 | 1005 | 1711957 | 1712961 | BCG_1559 | BCG_1559 | 1005 | 1683787 | 1684791 | BCGDan_1557 | BCGDan_1557 | 1005 | 1683891 | 1684901 | BCGDan_sapMKO_1556 | BCGDan_sapMKO_1556 | 1011 |
| 1687941 | 1689230 | lipL | Rv1497 | 1290 | 1713014 | 1714303 | lipL | BCG_1560 | 1290 | 1684844 | 1686133 | lipL | BCGDan_1558 | 1290 | 1684954 | 1686243 | lipL | BCGDan_sapMKO_1557 | 1290 |
| 1689303 | 1689920 | Rv1498c | Rv1498c | 618 | 1714376 | 1714993 | BCG_1561c | BCG_1561c | 618 | 1686206 | 1686823 | BCGDan_1559 | BCGDan_1559 | 618 | 1686316 | 1686933 | BCGDan_sapMKO_1558 | BCGDan_sapMKO_1558 | 618 |
| 1690134 | 1690346 | Rv1498A | Rv1498A | 213 | 1715207 | 1715398 | BCG_1562c | BCG_1562c | 192 | 1687037 | 1687228 | BCGDan_1560 | BCGDan_1560 | 192 | 1687147 | 1687338 | BCGDan_sapMKO_1559 | BCGDan_sapMKO_1559 | 192 |
| 1690407 | 1690805 | Rv1499 | Rv1499 | 399 | 1715480 | 1715878 | BCG_1563 | BCG_1563 | 399 | 1687310 | 1687708 | BCGDan_1561 | BCGDan_1561 | 399 | 1687462 | 1687818 | BCGDan_sapMKO_1560 | BCGDan_sapMKO_1560 | 357 |
| 1690850 | 1691878 | Rv1500 | Rv1500 | 1029 | 1715923 | 1716951 | BCG_1564 | BCG_1564 | 1029 | 1687753 | 1688781 | BCGDan_1562 | BCGDan_1562 | 1029 | 1687863 | 1688891 | BCGDan_sapMKO_1561 | BCGDan_sapMKO_1561 | 1029 |
| 1691890 | 1692711 | Rv1501 | Rv1501 | 822 | 1716963 | 1717784 | BCG_1565 | BCG_1565 | 822 | 1688793 | 1689614 | BCGDan_1563 | BCGDan_1563 | 822 | 1688903 | 1689724 | BCGDan_sapMKO_1562 | BCGDan_sapMKO_1562 | 822 |
| 1692924 | 1693823 | Rv1502 | Rv1502 | 900 | 1717997 | 1718896 | BCG_1566 | BCG_1566 | 900 | 1689827 | 1690726 | BCGDan_1564 | BCGDan_1564 | 900 | 1689937 | 1690836 | BCGDan_sapMKO_1563 | BCGDan_sapMKO_1563 | 900 |
| 1693996 | 1694544 | Rv1503c | Rv1503c | 549 | 1719069 | 1720217 | BCG_1567c | BCG_1567c | 1149 | 1690899 | 1692047 | BCGDan_1565 | BCGDan_1565 | 1149 | 1691009 | 1692157 | BCGDan_sapMKO_1564 | BCGDan_sapMKO_1564 | 1149 |

| | | | | | | | | | | | | | | | | | | | |
|---|---|---|---|---|---|---|---|---|---|---|---|---|---|---|---|---|---|---|---|
| 1695281 | 1695946 | Rv1505c | Rv1505c | 666 | 1720354 | 1721019 | BCG_1568c | BCG_1568c | 666 | 1692184 | 1692849 | BCGDan_1566 | BCGDan_1566 | 666 | 1692294 | 1692959 | BCGDan_sapMKO_1565 | BCGDan_sapMKO_1565 | 666 |
| 1708871 | 1709635 | Rv1517 | Rv1517 | 765 | 1721212 | 1721976 | BCG_1569 | BCG_1569 | 765 | 1693042 | 1693806 | BCGDan_1567 | BCGDan_1567 | 765 | 1693185 | 1693916 | BCGDan_sapMKO_1566 | BCGDan_sapMKO_1566 | 732 |
| 1709644 | 1710603 | Rv1518 | Rv1518 | 960 | 1721985 | 1722944 | BCG_1570 | BCG_1570 | 960 | 1693839 | 1694774 | BCGDan_1568 | BCGDan_1568 | 936 | 1693925 | 1694884 | BCGDan_sapMKO_1567 | BCGDan_sapMKO_1567 | 960 |
| 1710733 | 1711002 | Rv1519 | Rv1519 | 270 | 1723074 | 1723343 | BCG_1571 | BCG_1571 | 270 | 1694904 | 1695173 | BCGDan_1569 | BCGDan_1569 | 270 | 1695014 | 1695283 | BCGDan_sapMKO_1568 | BCGDan_sapMKO_1568 | 270 |
| 1711028 | 1712068 | Rv1520 | Rv1520 | 1041 | 1723369 | 1724409 | BCG_1572 | BCG_1572 | 1041 | 1695199 | 1696239 | BCGDan_1570 | BCGDan_1570 | 1041 | 1695276 | 1696349 | BCGDan_sapMKO_1569 | BCGDan_sapMKO_1569 | 1074 |
| 1712302 | 1714053 | fadD25 | Rv1521 | 1752 | 1724643 | 1726394 | fadD25 | BCG_1573 | 1752 | 1696473 | 1698224 | fadD25 | BCGDan_1571 | 1752 | 1696580 | 1698334 | fadD25 | BCGDan_sapMKO_1570 | 1755 |
| 1714172 | 1717612 | mmpL12 | Rv1522c | 3441 | 1726630 | 1729953 | mmpL12 | BCG_1574c | 3324 | 1698460 | 1701783 | mmpL12 | BCGDan_1572 | 3324 | 1698570 | 1701893 | mmpL12 | BCGDan_sapMKO_1571 | 3324 |
| 1717653 | 1718696 | Rv1523 | Rv1523 | 1044 | 1729994 | 1731037 | BCG_1575 | BCG_1575 | 1044 | 1701824 | 1702867 | BCGDan_1573 | BCGDan_1573 | 1044 | 1701934 | 1702977 | BCGDan_sapMKO_1572 | BCGDan_sapMKO_1572 | 1044 |
| 1718726 | 1719970 | Rv1524 | Rv1524 | 1245 | 1731067 | 1732311 | BCG_1576 | BCG_1576 | 1245 | 1702897 | 1704141 | BCGDan_1574 | BCGDan_1574 | 1245 | 1703007 | 1704251 | BCGDan_sapMKO_1573 | BCGDan_sapMKO_1573 | 1245 |
| 1720017 | 1720802 | wbbL2 | Rv1525 | 786 | 1732358 | 1733143 | wbbL2 | BCG_1577 | 786 | 1704188 | 1704973 | wbbL2 | BCGDan_1575 | 786 | 1704322 | 1705083 | wbbL2 | BCGDan_sapMKO_1574 | 762 |
| 1720780 | 1722060 | Rv1526c | Rv1526c | 1281 | 1733121 | 1734401 | BCG_1578c | BCG_1578c | 1281 | 1704951 | 1706231 | BCGDan_1576 | BCGDan_1576 | 1281 | 1705061 | 1706341 | BCGDan_sapMKO_1575 | BCGDan_sapMKO_1575 | 1281 |
| 1722083 | 1728409 | pks5 | Rv1527c | 6327 | 1734424 | 1740750 | pks5 | BCG_1579c | 6327 | 1706254 | 1712580 | pks5 | BCGDan_1577 | 6327 | 1706364 | 1712690 | pks5 | BCGDan_sapMKO_1576 | 6327 |
| 1728953 | 1729450 | papA4 | Rv1528c | 498 | 1741294 | 1741791 | papA4 | BCG_1580c | 498 | 1713124 | 1713621 | papA4 | BCGDan_1578 | 498 | 1713234 | 1713731 | papA4 | BCGDan_sapMKO_1577 | 498 |
| 1729502 | 1731256 | fadD24 | Rv1529 | 1755 | 1741843 | 1743597 | fadD24 | BCG_1581 | 1755 | 1713673 | 1715427 | fadD24 | BCGDan_1579 | 1755 | 1713783 | 1715537 | fadD24 | BCGDan_sapMKO_1578 | 1755 |
| 1731373 | 1732476 | adh | Rv1530 | 1104 | 1743714 | 1744817 | adh | BCG_1582 | 1104 | 1715544 | 1716647 | adh | BCGDan_1580 | 1104 | 1715654 | 1716757 | adh | BCGDan_sapMKO_1579 | 1104 |
| 1732473 | 1733039 | Rv1531 | Rv1531 | 567 | 1744814 | 1745380 | BCG_1583 | BCG_1583 | 567 | 1716644 | 1717210 | BCGDan_1581 | BCGDan_1581 | 567 | 1716769 | 1717320 | BCGDan_sapMKO_1580 | BCGDan_sapMKO_1580 | 552 |
| 1733116 | 1733550 | Rv1532c | Rv1532c | 435 | 1745457 | 1745891 | BCG_1584c | BCG_1584c | 435 | 1717287 | 1717721 | BCGDan_1582 | BCGDan_1582 | 435 | 1717397 | 1717831 | BCGDan_sapMKO_1581 | BCGDan_sapMKO_1581 | 435 |
| 1733610 | 1734737 | Rv1533 | Rv1533 | 1128 | 1745951 | 1747078 | BCG_1585 | BCG_1585 | 1128 | 1717781 | 1718908 | BCGDan_1583 | BCGDan_1583 | 1128 | 1717891 | 1719018 | BCGDan_sapMKO_1582 | BCGDan_sapMKO_1582 | 1128 |
| 1734734 | 1735411 | Rv1534 | Rv1534 | 678 | 1747075 | 1747752 | BCG_1586 | BCG_1586 | 678 | 1718905 | 1719582 | BCGDan_1584 | BCGDan_1584 | 678 | 1719015 | 1719692 | BCGDan_sapMKO_1583 | BCGDan_sapMKO_1583 | 678 |
| 1735976 | 1736212 | Rv1535 | Rv1535 | 237 | 1748317 | 1748553 | BCG_1587 | BCG_1587 | 237 | 1720147 | 1720383 | BCGDan_1585 | BCGDan_1585 | 237 | 1720257 | 1720493 | BCGDan_sapMKO_1584 | BCGDan_sapMKO_1584 | 237 |
| 1736519 | 1739644 | ileS | Rv1536 | 3126 | 1748860 | 1751985 | ileS | BCG_1588 | 3126 | 1720690 | 1723815 | ileS | BCGDan_1586 | 3126 | 1720800 | 1723925 | ileS | BCGDan_sapMKO_1585 | 3126 |
| 1739856 | 1741247 | dinX | Rv1537 | 1392 | 1752197 | 1753588 | dinX | BCG_1589 | 1392 | 1724027 | 1725418 | dinX | BCGDan_1587 | 1392 | 1724122 | 1725528 | dinX | BCGDan_sapMKO_1586 | 1407 |
| 1741212 | 1742192 | ansA | Rv1538c | 981 | 1753553 | 1754533 | ansA | BCG_1590c | 981 | 1725383 | 1726363 | ansA | BCGDan_1588 | 981 | 1725493 | 1726473 | ansA | BCGDan_sapMKO_1587 | 981 |
| 1742244 | 1742852 | lspA | Rv1539 | 609 | 1754585 | 1755193 | lspA | BCG_1591 | 609 | 1726415 | 1727023 | lspA | BCGDan_1589 | 609 | 1726525 | 1727133 | lspA | BCGDan_sapMKO_1588 | 609 |
| 1742845 | 1743771 | Rv1540 | Rv1540 | 927 | 1755186 | 1756112 | BCG_1592 | BCG_1592 | 927 | 1727016 | 1727942 | BCGDan_1590 | BCGDan_1590 | 927 | 1727168 | 1728052 | BCGDan_sapMKO_1589 | BCGDan_sapMKO_1589 | 885 |
| 1743778 | 1744371 | lprI | Rv1541c | 594 | 1756119 | 1756712 | lprI | BCG_1593c | 594 | 1727949 | 1728542 | lprI | BCGDan_1591 | 594 | 1728059 | 1728652 | lprI | BCGDan_sapMKO_1590 | 594 |
| 1744426 | 1744836 | glbN | Rv1542c | 411 | 1756767 | 1757177 | glbN | BCG_1594c | 411 | 1728597 | 1729007 | glbN | BCGDan_1592 | 411 | 1728707 | 1729117 | glbN | BCGDan_sapMKO_1591 | 411 |
| 1745064 | 1746089 | Rv1543 | Rv1543 | 1026 | 1757405 | 1758430 | BCG_1595 | BCG_1595 | 1026 | 1729235 | 1730260 | BCGDan_1593 | BCGDan_1593 | 1026 | 1729345 | 1730370 | BCGDan_sapMKO_1592 | BCGDan_sapMKO_1592 | 1026 |
| 1746094 | 1746897 | Rv1544 | Rv1544 | 804 | 1758435 | 1758533 | BCG_1596 | BCG_1596 | 99 | 1730265 | 1731068 | BCGDan_1594 | BCGDan_1594 | 804 | 1730375 | 1731178 | BCGDan_sapMKO_1593 | BCGDan_sapMKO_1593 | 804 |
| 1746919 | 1747146 | Rv1545 | Rv1545 | 228 | 1759250 | 1759477 | BCG_1598 | BCG_1598 | 228 | 1731090 | 1731317 | BCGDan_1595 | BCGDan_1595 | 228 | 1731200 | 1731427 | BCGDan_sapMKO_1594 | BCGDan_sapMKO_1594 | 228 |
| 1747195 | 1747626 | Rv1546 | Rv1546 | 432 | 1759526 | 1759957 | BCG_1599 | BCG_1599 | 432 | 1731366 | 1731797 | BCGDan_1596 | BCGDan_1596 | 432 | 1731476 | 1731907 | BCGDan_sapMKO_1595 | BCGDan_sapMKO_1595 | 432 |
| 1747694 | 1751248 | dnaE1 | Rv1547 | 3555 | 1760025 | 1763579 | dnaE1 | BCG_1600 | 3555 | 1731865 | 1735419 | dnaE1 | BCGDan_1597 | 3555 | 1732005 | 1735529 | dnaE1 | BCGDan_sapMKO_1596 | 3525 |
| 1751297 | 1753333 | PPE21 | Rv1548c | 2037 | 1763628 | 1765664 | PPE21 | BCG_1601c | 2037 | 1735468 | 1737504 | PPE21 | BCGDan_1598 | 2037 | 1735578 | 1737623 | PPE21 | BCGDan_sapMKO_1597 | 2046 |
| 1753510 | 1754037 | fadD11.1 | Rv1549 | 528 | 1765821 | 1767764 | fadD11 | BCG_1602 | 1944 | 1737661 | 1739604 | fadD11 | BCGDan_1599 | 1944 | 1737915 | 1739714 | fadD11 | BCGDan_sapMKO_1598 | 1800 |
| 1755445 | 1757310 | plsB1 | Rv1551 | 1866 | 1767778 | 1769643 | plsB1 | BCG_1603 | 1866 | 1739618 | 1741483 | plsB1 | BCGDan_1600 | 1866 | 1739728 | 1741593 | plsB1 | BCGDan_sapMKO_1599 | 1866 |
| 1757681 | 1759432 | frdA | Rv1552 | 1752 | 1770013 | 1771764 | frdA | BCG_1604 | 1752 | 1741853 | 1743604 | frdA | BCGDan_1601 | 1752 | 1741962 | 1743713 | frdA | BCGDan_sapMKO_1600 | 1752 |
| 1759435 | 1760178 | frdB | Rv1553 | 744 | 1771767 | 1772897 | frdBC | BCG_1605 | 1131 | 1743607 | 1744737 | frdBC | BCGDan_1602 | 1131 | 1743716 | 1744846 | frdBC | BCGDan_sapMKO_1601 | 1131 |
| 1760552 | 1760929 | frdD | Rv1555 | 378 | 1772894 | 1773271 | frdD | BCG_1606 | 378 | 1744734 | 1745111 | frdD | BCGDan_1603 | 378 | 1744843 | 1745220 | frdD | BCGDan_sapMKO_1602 | 378 |
| 1760997 | 1761605 | Rv1556 | Rv1556 | 609 | 1773339 | 1773947 | BCG_1607 | BCG_1607 | 609 | 1745179 | 1745787 | BCGDan_1604 | BCGDan_1604 | 609 | 1745369 | 1745896 | BCGDan_sapMKO_1603 | BCGDan_sapMKO_1603 | 528 |
| NA | NA | NA | NA | NA | 1774086 | 1774532 | mmpS6 | BCG_1608 | 447 | 1745941 | 1746372 | mmpS6 | BCGDan_1605 | 432 | 1746050 | 1746481 | mmpS6 | BCGDan_sapMKO_1604 | 432 |
| 1761744 | 1762937 | mmpL6 | Rv1557 | 1194 | 1774529 | 1777432 | mmpL6 | BCG_1609 | 2904 | 1746369 | 1749272 | mmpL6 | BCGDan_1606 | 2904 | 1746478 | 1749381 | mmpL6 | BCGDan_sapMKO_1605 | 2904 |
| 1762947 | 1763393 | Rv1558 | Rv1558 | 447 | 1777442 | 1777888 | BCG_1610 | BCG_1610 | 447 | 1749282 | 1749728 | BCGDan_1607 | BCGDan_1607 | 447 | 1749391 | 1749837 | BCGDan_sapMKO_1606 | BCGDan_sapMKO_1606 | 447 |
| 1763428 | 1764717 | ilvA | Rv1559 | 1290 | 1777923 | 1779212 | ilvA | BCG_1611 | 1290 | 1749763 | 1751052 | ilvA | BCGDan_1608 | 1290 | 1749872 | 1751161 | ilvA | BCGDan_sapMKO_1607 | 1290 |
| 1764755 | 1764973 | vapB11 | Rv1560 | 219 | 1779250 | 1779468 | BCG_1612 | BCG_1612 | 219 | 1751090 | 1751308 | BCGDan_1609 | BCGDan_1609 | 219 | 1751214 | 1751417 | BCGDan_sapMKO_1608 | BCGDan_sapMKO_1608 | 204 |
| 1764979 | 1765383 | vapC11 | Rv1561 | 405 | 1779474 | 1779878 | BCG_1613 | BCG_1613 | 405 | 1751314 | 1751718 | BCGDan_1610 | BCGDan_1610 | 405 | 1751423 | 1751827 | BCGDan_sapMKO_1609 | BCGDan_sapMKO_1609 | 405 |
| 1765400 | 1767142 | treZ | Rv1562c | 1743 | 1779895 | 1780455 | treZb | BCG_1614c | 561 | 1751735 | 1752295 | treZb | BCGDan_1611 | 561 | 1751844 | 1752344 | treZb | BCGDan_sapMKO_1610 | 501 |
| NA | NA | NA | NA | NA | 1780452 | 1781636 | treZa | BCG_1615c | 1185 | 1752292 | 1753476 | treZa | BCGDan_1612 | 1185 | 1752401 | 1753585 | treZa | BCGDan_sapMKO_1611 | 1185 |
| 1767135 | 1769432 | treY | Rv1563c | 2298 | 1781629 | 1783926 | treY | BCG_1616c | 2298 | 1753469 | 1755766 | treY | BCGDan_1613 | 2298 | 1753578 | 1755875 | treY | BCGDan_sapMKO_1612 | 2298 |
| 1769436 | 1771601 | treX | Rv1564c | 2166 | 1783930 | 1786095 | treX | BCG_1617c | 2166 | 1755770 | 1757935 | treX | BCGDan_1614 | 2166 | 1755879 | 1758044 | treX | BCGDan_sapMKO_1613 | 2166 |
| 1771640 | 1773829 | Rv1565c | Rv1565c | 2190 | 1786134 | 1788323 | BCG_1618c | BCG_1618c | 2190 | 1757974 | 1760163 | BCGDan_1615 | BCGDan_1615 | 2190 | 1758083 | 1760206 | BCGDan_sapMKO_1614 | BCGDan_sapMKO_1614 | 2124 |
| 1773928 | 1774620 | Rv1566c | Rv1566c | 693 | 1788422 | 1789114 | BCG_1619c | BCG_1619c | 693 | 1760262 | 1760954 | BCGDan_1616 | BCGDan_1616 | 693 | 1760371 | 1761063 | BCGDan_sapMKO_1615 | BCGDan_sapMKO_1615 | 693 |
| 1774860 | 1775144 | Rv1567c | Rv1567c | 285 | 1789354 | 1789638 | BCG_1620c | BCG_1620c | 285 | 1761194 | 1761478 | BCGDan_1617 | BCGDan_1617 | 285 | 1761303 | 1761587 | BCGDan_sapMKO_1616 | BCGDan_sapMKO_1616 | 285 |
| 1775392 | 1776705 | bioA | Rv1568 | 1314 | 1789886 | 1791199 | bioA | BCG_1621 | 1314 | 1761726 | 1763039 | bioA | BCGDan_1618 | 1314 | 1761799 | 1763148 | bioA | BCGDan_sapMKO_1617 | 1350 |
| 1776702 | 1777862 | bioF1 | Rv1569 | 1161 | 1791196 | 1792356 | bioF1 | BCG_1622 | 1161 | 1763036 | 1764196 | bioF1 | BCGDan_1619 | 1161 | 1763145 | 1764305 | bioF1 | BCGDan_sapMKO_1618 | 1161 |
| 1777859 | 1778539 | bioD | Rv1570 | 681 | 1792353 | 1793033 | bioD | BCG_1623 | 681 | 1764193 | 1764873 | bioD | BCGDan_1620 | 681 | 1764302 | 1764982 | bioD | BCGDan_sapMKO_1619 | 681 |
| 1778539 | 1779048 | Rv1571 | Rv1571 | 510 | 1793033 | 1793542 | BCG_1624 | BCG_1624 | 510 | 1764873 | 1765382 | BCGDan_1621 | BCGDan_1621 | 510 | 1764982 | 1765491 | BCGDan_sapMKO_1620 | BCGDan_sapMKO_1620 | 510 |
| 1779194 | 1779298 | Rv1572c | Rv1572c | 105 | 1793688 | 1794644 | BCG_1625c | BCG_1625c | 957 | 1765528 | 1766484 | BCGDan_1622 | BCGDan_1622 | 957 | 1765637 | 1766599 | BCGDan_sapMKO_1621 | BCGDan_sapMKO_1621 | 963 |
| 1789168 | 1789836 | Rv1588c | Rv1588c | 669 | 1794415 | 1795083 | BCG_1626c | BCG_1626c | 669 | 1766255 | 1766923 | BCGDan_1623 | BCGDan_1623 | 669 | 1766364 | 1767032 | BCGDan_sapMKO_1622 | BCGDan_sapMKO_1622 | 669 |
| 1790284 | 1791333 | bioB | Rv1589 | 1050 | 1795531 | 1796580 | bioB | BCG_1627 | 1050 | 1767371 | 1768420 | bioB | BCGDan_1624 | 1050 | 1767432 | 1768529 | bioB | BCGDan_sapMKO_1623 | 1098 |
| 1791334 | 1791573 | Rv1590 | Rv1590 | 240 | 1796581 | 1796820 | BCG_1628 | BCG_1628 | 240 | 1768421 | 1768660 | BCGDan_1625 | BCGDan_1625 | 240 | 1768530 | 1768769 | BCGDan_sapMKO_1624 | BCGDan_sapMKO_1624 | 240 |
| 1791570 | 1792235 | Rv1591 | Rv1591 | 666 | 1796817 | 1797482 | BCG_1629 | BCG_1629 | 666 | 1768657 | 1769322 | BCGDan_1626 | BCGDan_1626 | 666 | 1768709 | 1769431 | BCGDan_sapMKO_1625 | BCGDan_sapMKO_1625 | 723 |
| 1792400 | 1793740 | Rv1592c | Rv1592c | 1341 | 1797647 | 1798987 | BCG_1630c | BCG_1630c | 1341 | 1769487 | 1770827 | BCGDan_1627 | BCGDan_1627 | 1341 | 1769596 | 1770936 | BCGDan_sapMKO_1626 | BCGDan_sapMKO_1626 | 1341 |
| 1793997 | 1794707 | Rv1593c | Rv1593c | 711 | 1799244 | 1799954 | BCG_1631c | BCG_1631c | 711 | 1771084 | 1771794 | BCGDan_1628 | BCGDan_1628 | 711 | 1771193 | 1771903 | BCGDan_sapMKO_1627 | BCGDan_sapMKO_1627 | 711 |
| 1794756 | 1795805 | nadA | Rv1594 | 1050 | 1800003 | 1801052 | nadA | BCG_1632 | 1050 | 1771843 | 1772892 | nadA | BCGDan_1629 | 1050 | 1771952 | 1773001 | nadA | BCGDan_sapMKO_1628 | 1050 |
| 1795805 | 1797388 | nadB | Rv1595 | 1584 | 1801052 | 1802635 | nadB | BCG_1633 | 1584 | 1772892 | 1774475 | nadB | BCGDan_1630 | 1584 | 1772998 | 1774584 | nadB | BCGDan_sapMKO_1629 | 1587 |
| 1797388 | 1798245 | nadC | Rv1596 | 858 | 1802635 | 1803492 | nadC | BCG_1634 | 858 | 1774475 | 1775332 | nadC | BCGDan_1631 | 858 | 1774578 | 1775441 | nadC | BCGDan_sapMKO_1630 | 864 |
| 1798294 | 1799052 | Rv1597 | Rv1597 | 759 | 1803541 | 1804299 | BCG_1635 | BCG_1635 | 759 | 1775381 | 1776139 | BCGDan_1632 | BCGDan_1632 | 759 | 1775490 | 1776248 | BCGDan_sapMKO_1631 | BCGDan_sapMKO_1631 | 759 |
| 1799073 | 1799483 | Rv1598c | Rv1598c | 411 | 1804320 | 1804730 | BCG_1636c | BCG_1636c | 411 | 1776160 | 1776570 | BCGDan_1633 | BCGDan_1633 | 411 | 1776269 | 1776637 | BCGDan_sapMKO_1632 | BCGDan_sapMKO_1632 | 369 |
| 1799583 | 1800899 | hisD | Rv1599 | 1317 | 1804830 | 1806146 | hisD | BCG_1637 | 1317 | 1776670 | 1777986 | hisD | BCGDan_1634 | 1317 | 1776779 | 1778095 | hisD | BCGDan_sapMKO_1633 | 1317 |
| 1800896 | 1802038 | hisC1 | Rv1600 | 1143 | 1806143 | 1807285 | hisC1 | BCG_1638 | 1143 | 1777983 | 1779125 | hisC1 | BCGDan_1635 | 1143 | 1778092 | 1779234 | hisC1 | BCGDan_sapMKO_1634 | 1143 |
| 1802035 | 1802667 | hisB | Rv1601 | 633 | 1807282 | 1807914 | hisB | BCG_1639 | 633 | 1779122 | 1779754 | hisB | BCGDan_1636 | 633 | 1779231 | 1779863 | hisB | BCGDan_sapMKO_1635 | 633 |
| 1802664 | 1803284 | hisH | Rv1602 | 621 | 1807911 | 1808531 | hisH | BCG_1640 | 621 | 1779751 | 1780371 | hisH | BCGDan_1637 | 621 | 1779860 | 1780480 | hisH | BCGDan_sapMKO_1636 | 621 |
| 1803294 | 1804031 | hisA | Rv1603 | 738 | 1808541 | 1809278 | hisA | BCG_1641 | 738 | 1780381 | 1781118 | hisA | BCGDan_1638 | 738 | 1780493 | 1781227 | hisA | BCGDan_sapMKO_1637 | 735 |
| 1804039 | 1804851 | impA | Rv1604 | 813 | 1809286 | 1810098 | impA | BCG_1642 | 813 | 1781126 | 1781938 | impA | BCGDan_1639 | 813 | 1781235 | 1782047 | impA | BCGDan_sapMKO_1638 | 813 |
| 1804853 | 1805656 | hisF | Rv1605 | 804 | 1810100 | 1810903 | hisF | BCG_1643 | 804 | 1781940 | 1782743 | hisF | BCGDan_1640 | 804 | 1782049 | 1782852 | hisF | BCGDan_sapMKO_1639 | 804 |
| 1805653 | 1806000 | hisI | Rv1606 | 348 | 1810900 | 1811247 | hisI | BCG_1644 | 348 | 1782740 | 1783087 | hisI | BCGDan_1641 | 348 | 1782849 | 1783196 | hisI | BCGDan_sapMKO_1640 | 348 |
| 1806181 | 1807263 | chaA | Rv1607 | 1083 | 1811428 | 1812510 | chaA | BCG_1645 | 1083 | 1783268 | 1784350 | chaA | BCGDan_1642 | 1083 | 1783377 | 1784459 | chaA | BCGDan_sapMKO_1641 | 1083 |
| 1807298 | 1807762 | bcpB | Rv1608c | 465 | 1812545 | 1813009 | bcpB | BCG_1646c | 465 | 1784385 | 1784849 | bcpB | BCGDan_1643 | 465 | 1784494 | 1784958 | bcpB | BCGDan_sapMKO_1642 | 465 |
| 1807903 | 1809453 | trpE | Rv1609 | 1551 | 1813150 | 1814700 | trpE | BCG_1647 | 1551 | 1784990 | 1786540 | trpE | BCGDan_1644 | 1551 | 1785015 | 1786649 | trpE | BCGDan_sapMKO_1643 | 1635 |
| 1809443 | 1810150 | Rv1610 | Rv1610 | 708 | 1814690 | 1815397 | BCG_1648 | BCG_1648 | 708 | 1786530 | 1787237 | BCGDan_1645 | BCGDan_1645 | 708 | 1786687 | 1787346 | BCGDan_sapMKO_1644 | BCGDan_sapMKO_1644 | 660 |
| 1810240 | 1811058 | trpC | Rv1611 | 819 | 1815487 | 1816305 | trpC | BCG_1649 | 819 | 1787327 | 1788145 | trpC | BCGDan_1646 | 819 | 1787436 | 1788254 | trpC | BCGDan_sapMKO_1645 | 819 |
| 1811127 | 1812359 | trpB | Rv1612 | 1233 | 1816374 | 1817606 | trpB | BCG_1650 | 1233 | 1788214 | 1789446 | trpB | BCGDan_1647 | 1233 | 1788290 | 1789555 | trpB | BCGDan_sapMKO_1646 | 1266 |
| 1812359 | 1813171 | trpA | Rv1613 | 813 | 1817606 | 1818418 | trpA | BCG_1651 | 813 | 1789446 | 1790258 | trpA | BCGDan_1648 | 813 | 1789555 | 1790367 | trpA | BCGDan_sapMKO_1647 | 813 |
| 1813171 | 1814577 | lgt | Rv1614 | 1407 | 1818418 | 1819824 | lgt | BCG_1652 | 1407 | 1790258 | 1791664 | lgt | BCGDan_1649 | 1407 | 1790367 | 1791773 | lgt | BCGDan_sapMKO_1648 | 1407 |
| 1815253 | 1815693 | Rv1615 | Rv1615 | 441 | 1820500 | 1820940 | BCG_1653 | BCG_1653 | 441 | 1792340 | 1792780 | BCGDan_1650 | BCGDan_1650 | 441 | 1792494 | 1792889 | BCGDan_sapMKO_1649 | BCGDan_sapMKO_1649 | 396 |
| 1815683 | 1816081 | Rv1616 | Rv1616 | 399 | 1820930 | 1821328 | BCG_1654 | BCG_1654 | 399 | 1792770 | 1793168 | BCGDan_1651 | BCGDan_1651 | 399 | 1792879 | 1793277 | BCGDan_sapMKO_1650 | BCGDan_sapMKO_1650 | 399 |

| | | | | | | | | | | | | | | | | | | | |
|---|---|---|---|---|---|---|---|---|---|---|---|---|---|---|---|---|---|---|---|
| 1816189 | 1817607 | pykA | Rv1617 | 1419 | 1821436 | 1822854 | pykA | BCG_1655 | 1419 | 1793276 | 1794694 | pykA | BCGDan_1652 | 1419 | 1793385 | 1794803 | pykA | BCGDan_sapMKO_1651 | 1419 |
| 1817615 | 1818517 | tesB1 | Rv1618 | 903 | 1822862 | 1823764 | tesB1 | BCG_1656 | 903 | 1794702 | 1795604 | tesB1 | BCGDan_1653 | 903 | 1794793 | 1795713 | tesB1 | BCGDan_sapMKO_1652 | 921 |
| 1818575 | 1820029 | Rv1619 | Rv1619 | 1455 | 1823822 | 1825276 | BCG_1657 | BCG_1657 | 1455 | 1795662 | 1797116 | BCGDan_1654 | BCGDan_1654 | 1455 | 1795771 | 1797225 | BCGDan_sapMKO_1653 | BCGDan_sapMKO_1653 | 1455 |
| 1819963 | 1821693 | cydC | Rv1620c | 1731 | 1825210 | 1826940 | cydC | BCG_1658c | 1731 | 1797050 | 1798780 | cydC | BCGDan_1655 | 1731 | 1797159 | 1798889 | cydC | BCGDan_sapMKO_1654 | 1731 |
| 1821690 | 1823273 | cydD | Rv1621c | 1584 | 1826937 | 1828520 | cydD | BCG_1659c | 1584 | 1798777 | 1800360 | cydD | BCGDan_1656 | 1584 | 1798886 | 1800469 | cydD | BCGDan_sapMKO_1655 | 1584 |
| 1823360 | 1824400 | cydB | Rv1622c | 1041 | 1828607 | 1829647 | cydB | BCG_1660c | 1041 | 1800447 | 1801487 | cydB | BCGDan_1657 | 1041 | 1800556 | 1801596 | cydB | BCGDan_sapMKO_1656 | 1041 |
| 1824430 | 1825887 | cydA | Rv1623c | 1458 | 1829677 | 1831134 | cydA | BCG_1661c | 1458 | 1801517 | 1802974 | cydA | BCGDan_1658 | 1458 | 1801626 | 1803083 | cydA | BCGDan_sapMKO_1657 | 1458 |
| 1825998 | 1826585 | Rv1624c | Rv1624c | 588 | 1831245 | 1831832 | BCG_1662c | BCG_1662c | 588 | 1803085 | 1803672 | BCGDan_1659 | BCGDan_1659 | 588 | 1803194 | 1803727 | BCGDan_sapMKO_1658 | BCGDan_sapMKO_1658 | 534 |
| 1826614 | 1827945 | cya | Rv1625c | 1332 | 1831861 | 1833117 | cya | BCG_1663c | 1257 | 1803701 | 1804957 | cya | BCGDan_1660 | 1257 | 1803810 | 1805066 | cya | BCGDan_sapMKO_1659 | 1257 |
| 1828180 | 1828797 | Rv1626 | Rv1626 | 618 | 1833427 | 1834044 | BCG_1664 | BCG_1664 | 618 | 1805267 | 1805884 | BCGDan_1662 | BCGDan_1662 | 618 | 1805376 | 1805993 | BCGDan_sapMKO_1661 | BCGDan_sapMKO_1661 | 618 |
| 1828865 | 1830073 | Rv1627c | Rv1627c | 1209 | 1834112 | 1835320 | BCG_1665c | BCG_1665c | 1209 | 1805952 | 1807160 | BCGDan_1663 | BCGDan_1663 | 1209 | 1806061 | 1807263 | BCGDan_sapMKO_1662 | BCGDan_sapMKO_1662 | 1203 |
| 1830070 | 1830561 | Rv1628c | Rv1628c | 492 | 1835317 | 1835808 | BCG_1666c | BCG_1666c | 492 | 1807157 | 1807648 | BCGDan_1664 | BCGDan_1664 | 492 | 1807266 | 1807757 | BCGDan_sapMKO_1663 | BCGDan_sapMKO_1663 | 492 |
| 1830665 | 1833379 | polA | Rv1629 | 2715 | 1835912 | 1838626 | polA | BCG_1667 | 2715 | 1807752 | 1810466 | polA | BCGDan_1665 | 2715 | 1807861 | 1810575 | polA | BCGDan_sapMKO_1664 | 2715 |
| 1833542 | 1834987 | rpsA | Rv1630 | 1446 | 1838789 | 1840234 | rpsA | BCG_1668 | 1446 | 1810629 | 1812074 | rpsA | BCGDan_1666 | 1446 | 1810738 | 1812183 | rpsA | BCGDan_sapMKO_1665 | 1446 |
| 1835013 | 1836236 | coaE | Rv1631 | 1224 | 1840260 | 1841483 | coaE | BCG_1669 | 1224 | 1812100 | 1813323 | coaE | BCGDan_1667 | 1224 | 1812209 | 1813432 | coaE | BCGDan_sapMKO_1666 | 1224 |
| 1836387 | 1836830 | Rv1632c | Rv1632c | 444 | 1841634 | 1842077 | BCG_1670c | BCG_1670c | 444 | 1813474 | 1813917 | BCGDan_1668 | BCGDan_1668 | 444 | 1813583 | 1814026 | BCGDan_sapMKO_1667 | BCGDan_sapMKO_1667 | 444 |
| 1837075 | 1839171 | uvrB | Rv1633 | 2097 | 1842322 | 1844418 | uvrB | BCG_1671 | 2097 | 1814162 | 1816258 | uvrB | BCGDan_1669 | 2097 | 1814091 | 1816367 | uvrB | BCGDan_sapMKO_1668 | 2277 |
| 1839168 | 1840583 | Rv1634 | Rv1634 | 1416 | 1844415 | 1845830 | BCG_1672 | BCG_1672 | 1416 | 1816255 | 1817670 | BCGDan_1670 | BCGDan_1670 | 1416 | 1816364 | 1817779 | BCGDan_sapMKO_1669 | BCGDan_sapMKO_1669 | 1416 |
| 1840572 | 1842242 | Rv1635c | Rv1635c | 1671 | 1845819 | 1847489 | BCG_1673c | BCG_1673c | 1671 | 1817659 | 1819329 | BCGDan_1671 | BCGDan_1671 | 1671 | 1817768 | 1819465 | BCGDan_sapMKO_1670 | BCGDan_sapMKO_1670 | 1698 |
| 1842451 | 1842891 | TB15.3 | Rv1636 | 441 | 1847698 | 1848138 | TB15.3 | BCG_1674 | 441 | 1819538 | 1819978 | TB15.3 | BCGDan_1672 | 441 | 1819647 | 1820087 | TB15.3 | BCGDan_sapMKO_1671 | 441 |
| 1842898 | 1843692 | Rv1637c | Rv1637c | 795 | 1848145 | 1848939 | BCG_1675c | BCG_1675c | 795 | 1819985 | 1820779 | BCGDan_1673 | BCGDan_1673 | 795 | 1820094 | 1820828 | BCGDan_sapMKO_1672 | BCGDan_sapMKO_1672 | 735 |
| 1843741 | 1846659 | uvrA | Rv1638 | 2919 | 1848988 | 1851906 | uvrA | BCG_1676 | 2919 | 1820828 | 1823746 | uvrA | BCGDan_1674 | 2919 | 1820901 | 1823855 | uvrA | BCGDan_sapMKO_1673 | 2955 |
| 1846716 | 1846973 | Rv1638A | Rv1638A | 258 | 1851963 | 1852220 | BCG_1677c | BCG_1677c | 258 | 1823803 | 1824060 | BCGDan_1675 | BCGDan_1675 | 258 | 1823912 | 1824169 | BCGDan_sapMKO_1674 | BCGDan_sapMKO_1674 | 258 |
| 1846989 | 1848458 | Rv1639c | Rv1639c | 1470 | 1852236 | 1853705 | BCG_1678c | BCG_1678c | 1470 | 1824076 | 1825545 | BCGDan_1676 | BCGDan_1676 | 1470 | 1824185 | 1825654 | BCGDan_sapMKO_1675 | BCGDan_sapMKO_1675 | 1470 |
| 1848517 | 1852035 | lysX | Rv1640c | 3519 | 1853764 | 1857282 | lysX | BCG_1679c | 3519 | 1825604 | 1829122 | lysX | BCGDan_1677 | 3519 | 1825713 | 1828964 | lysX | BCGDan_sapMKO_1676 | 3252 |
| 1852273 | 1852878 | infC | Rv1641 | 606 | 1857519 | 1858124 | infC | BCG_1680 | 606 | 1829359 | 1829964 | infC | BCGDan_1678 | 606 | 1829552 | 1830073 | infC | BCGDan_sapMKO_1677 | 522 |
| 1852928 | 1853122 | rpmI | Rv1642 | 195 | 1858174 | 1858368 | rpmI | BCG_1681 | 195 | 1830014 | 1830208 | rpmI | BCGDan_1679 | 195 | 1830123 | 1830317 | rpmI | BCGDan_sapMKO_1678 | 195 |
| 1853184 | 1853573 | rplT | Rv1643 | 390 | 1858430 | 1858819 | rplT | BCG_1682 | 390 | 1830270 | 1830659 | rplT | BCGDan_1680 | 390 | 1830379 | 1830768 | rplT | BCGDan_sapMKO_1679 | 390 |
| 1853606 | 1854388 | tsnR | Rv1644 | 783 | 1858852 | 1859634 | tsnR | BCG_1683 | 783 | 1830692 | 1831474 | tsnR | BCGDan_1681 | 783 | 1830801 | 1831583 | tsnR | BCGDan_sapMKO_1680 | 783 |
| 1854399 | 1855454 | Rv1645c | Rv1645c | 1056 | 1859645 | 1860700 | BCG_1684c | BCG_1684c | 1056 | 1831485 | 1832540 | BCGDan_1682 | BCGDan_1682 | 1056 | 1831594 | 1832649 | BCGDan_sapMKO_1681 | BCGDan_sapMKO_1681 | 1056 |
| 1855764 | 1856696 | PE17 | Rv1646 | 933 | 1861010 | 1861942 | PE17 | BCG_1685 | 933 | 1832850 | 1833782 | PE17 | BCGDan_1683 | 933 | 1832959 | 1833891 | PE17 | BCGDan_sapMKO_1682 | 933 |
| 1856774 | 1857724 | Rv1647 | Rv1647 | 951 | 1862020 | 1862970 | BCG_1686 | BCG_1686 | 951 | 1833860 | 1834810 | BCGDan_1684 | BCGDan_1684 | 951 | 1834005 | 1834919 | BCGDan_sapMKO_1683 | BCGDan_sapMKO_1683 | 915 |
| 1857731 | 1858537 | Rv1648 | Rv1648 | 807 | 1862977 | 1863783 | BCG_1687 | BCG_1687 | 807 | 1834817 | 1835623 | BCGDan_1685 | BCGDan_1685 | 807 | 1834926 | 1835732 | BCGDan_sapMKO_1684 | BCGDan_sapMKO_1684 | 807 |
| 1858733 | 1859758 | pheS | Rv1649 | 1026 | 1863979 | 1865004 | pheS | BCG_1688 | 1026 | 1835819 | 1836844 | pheS | BCGDan_1686 | 1026 | 1835895 | 1836953 | pheS | BCGDan_sapMKO_1685 | 1059 |
| 1859758 | 1862253 | pheT | Rv1650 | 2496 | 1865004 | 1867499 | pheT | BCG_1689 | 2496 | 1836844 | 1839339 | pheT | BCGDan_1687 | 2496 | 1836953 | 1839328 | pheT | BCGDan_sapMKO_1686 | 2376 |
| 1862347 | 1865382 | PE_PGRS30 | Rv1651c | 3036 | 1867593 | 1870649 | PE_PGRS30 | BCG_1690c | 3057 | 1839433 | 1842489 | PE_PGRS30 | BCGDan_1688 | 3057 | 1839541 | 1842597 | PE_PGRS30 | BCGDan_sapMKO_1687 | 3057 |
| 1865576 | 1866634 | argC | Rv1652 | 1059 | 1870843 | 1871901 | argC | BCG_1691 | 1059 | 1842683 | 1843741 | argC | BCGDan_1689 | 1059 | 1842791 | 1843849 | argC | BCGDan_sapMKO_1688 | 1059 |
| 1866631 | 1867845 | argJ | Rv1653 | 1215 | 1871898 | 1873112 | argJ | BCG_1692 | 1215 | 1843738 | 1844952 | argJ | BCGDan_1690 | 1215 | 1843846 | 1845060 | argJ | BCGDan_sapMKO_1689 | 1215 |
| 1867842 | 1868726 | argB | Rv1654 | 885 | 1873109 | 1873993 | argB | BCG_1693 | 885 | 1844949 | 1845833 | argB | BCGDan_1691 | 885 | 1845057 | 1845941 | argB | BCGDan_sapMKO_1690 | 885 |
| 1868723 | 1869925 | argD | Rv1655 | 1203 | 1873990 | 1875192 | argD | BCG_1694 | 1203 | 1845830 | 1847032 | argD | BCGDan_1692 | 1203 | 1845938 | 1847140 | argD | BCGDan_sapMKO_1691 | 1203 |
| 1869922 | 1870845 | argF | Rv1656 | 924 | 1875189 | 1876112 | argF | BCG_1695 | 924 | 1847029 | 1847952 | argF | BCGDan_1693 | 924 | 1847137 | 1848060 | argF | BCGDan_sapMKO_1692 | 924 |
| 1870842 | 1871354 | argR | Rv1657 | 513 | 1876109 | 1876621 | argR | BCG_1696 | 513 | 1847949 | 1848461 | argR | BCGDan_1694 | 513 | 1848057 | 1848569 | argR | BCGDan_sapMKO_1693 | 513 |
| 1871363 | 1872559 | argG | Rv1658 | 1197 | 1876630 | 1877826 | argG | BCG_1697 | 1197 | 1848470 | 1849666 | argG | BCGDan_1695 | 1197 | 1848578 | 1849774 | argG | BCGDan_sapMKO_1694 | 1197 |
| 1872639 | 1874051 | argH | Rv1659 | 1413 | 1877906 | 1879318 | argH | BCG_1698 | 1413 | 1849746 | 1851158 | argH | BCGDan_1696 | 1413 | 1849854 | 1851266 | argH | BCGDan_sapMKO_1695 | 1413 |
| 1874160 | 1875221 | pks10 | Rv1660 | 1062 | 1879427 | 1880488 | pks10 | BCG_1699 | 1062 | 1851267 | 1852328 | pks10 | BCGDan_1697 | 1062 | 1851375 | 1852436 | pks10 | BCGDan_sapMKO_1696 | 1062 |
| 1875304 | 1881684 | pks7 | Rv1661 | 6381 | 1880571 | 1886951 | pks7 | BCG_1700 | 6381 | 1852411 | 1858791 | pks7 | BCGDan_1698 | 6381 | 1852519 | 1858899 | pks7 | BCGDan_sapMKO_1697 | 6381 |
| 1881704 | 1886512 | pks8 | Rv1662 | 4809 | 1886971 | 1891776 | pks8 | BCG_1701 | 4806 | 1858811 | 1863616 | pks8 | BCGDan_1699 | 4806 | 1858919 | 1863724 | pks8 | BCGDan_sapMKO_1698 | 4806 |
| 1886512 | 1888020 | pks17 | Rv1663 | 1509 | 1891776 | 1893284 | pks17 | BCG_1702 | 1509 | 1863616 | 1865124 | pks17 | BCGDan_1700 | 1509 | 1863700 | 1865232 | pks17 | BCGDan_sapMKO_1699 | 1533 |
| 1888026 | 1891079 | pks9 | Rv1664 | 3054 | 1893290 | 1896343 | pks9 | BCG_1703 | 3054 | 1865130 | 1868183 | pks9 | BCGDan_1701 | 3054 | 1865238 | 1868291 | pks9 | BCGDan_sapMKO_1700 | 3054 |
| 1891226 | 1892287 | pks11 | Rv1665 | 1062 | 1896490 | 1897551 | pks11 | BCG_1704 | 1062 | 1868330 | 1869391 | pks11 | BCGDan_1702 | 1062 | 1868438 | 1869499 | pks11 | BCGDan_sapMKO_1701 | 1062 |
| 1892270 | 1893562 | cyp139 | Rv1666c | 1293 | 1897534 | 1898826 | cyp139 | BCG_1705c | 1293 | 1869374 | 1870666 | cyp139 | BCGDan_1703 | 1293 | 1869482 | 1870792 | cyp139 | BCGDan_sapMKO_1702 | 1311 |
| 1893577 | 1894230 | Rv1667c | Rv1667c | 654 | 1898841 | 1900616 | BCG_1706c | BCG_1706c | 1776 | 1870681 | 1872456 | BCGDan_1704 | BCGDan_1704 | 1776 | 1870789 | 1872564 | BCGDan_sapMKO_1703 | BCGDan_sapMKO_1703 | 1776 |
| 1895725 | 1896087 | Rv1669 | Rv1669 | 363 | 1900999 | 1901361 | BCG_1707 | BCG_1707 | 363 | 1872782 | 1873144 | BCGDan_1705 | BCGDan_1705 | 363 | 1872890 | 1873252 | BCGDan_sapMKO_1704 | BCGDan_sapMKO_1704 | 363 |
| 1896120 | 1896467 | Rv1670 | Rv1670 | 348 | 1901394 | 1901741 | BCG_1708 | BCG_1708 | 348 | 1873177 | 1873524 | BCGDan_1706 | BCGDan_1706 | 348 | 1873249 | 1873632 | BCGDan_sapMKO_1705 | BCGDan_sapMKO_1705 | 384 |
| 1896475 | 1896867 | Rv1671 | Rv1671 | 393 | 1901749 | 1902141 | BCG_1709 | BCG_1709 | 393 | 1873532 | 1873924 | BCGDan_1707 | BCGDan_1707 | 393 | 1873640 | 1874032 | BCGDan_sapMKO_1706 | BCGDan_sapMKO_1706 | 393 |
| 1896876 | 1898207 | Rv1672c | Rv1672c | 1332 | 1902150 | 1903481 | BCG_1710c | BCG_1710c | 1332 | 1873933 | 1875264 | BCGDan_1708 | BCGDan_1708 | 1332 | 1874041 | 1875372 | BCGDan_sapMKO_1707 | BCGDan_sapMKO_1707 | 1332 |
| 1898300 | 1899232 | Rv1673c | Rv1673c | 933 | 1903574 | 1904506 | BCG_1711c | BCG_1711c | 933 | 1875357 | 1876289 | BCGDan_1709 | BCGDan_1709 | 933 | 1875465 | 1876397 | BCGDan_sapMKO_1708 | BCGDan_sapMKO_1708 | 933 |
| 1899260 | 1899916 | Rv1674c | Rv1674c | 657 | 1904534 | 1905190 | BCG_1712c | BCG_1712c | 657 | 1876317 | 1876973 | BCGDan_1710 | BCGDan_1710 | 657 | 1876425 | 1877081 | BCGDan_sapMKO_1709 | BCGDan_sapMKO_1709 | 657 |
| 1900241 | 1900975 | cmr | Rv1675c | 735 | 1905515 | 1906249 | BCG_1713c | BCG_1713c | 735 | 1877298 | 1878032 | BCGDan_1711 | BCGDan_1711 | 735 | 1877406 | 1878140 | BCGDan_sapMKO_1710 | BCGDan_sapMKO_1710 | 735 |
| 1901047 | 1901751 | Rv1676 | Rv1676 | 705 | 1906321 | 1907025 | BCG_1714 | BCG_1714 | 705 | 1878104 | 1878808 | BCGDan_1712 | BCGDan_1712 | 705 | 1878212 | 1878916 | BCGDan_sapMKO_1711 | BCGDan_sapMKO_1711 | 705 |
| 1901748 | 1902296 | dsbF | Rv1677 | 549 | 1907022 | 1907570 | dsbF | BCG_1715 | 549 | 1878805 | 1879353 | dsbF | BCGDan_1713 | 549 | 1878913 | 1879461 | dsbF | BCGDan_sapMKO_1712 | 549 |
| 1902397 | 1903299 | Rv1678 | Rv1678 | 903 | 1907671 | 1908573 | BCG_1716 | BCG_1716 | 903 | 1879454 | 1880356 | BCGDan_1714 | BCGDan_1714 | 903 | 1879556 | 1880464 | BCGDan_sapMKO_1713 | BCGDan_sapMKO_1713 | 909 |
| 1903299 | 1904420 | fadE16 | Rv1679 | 1122 | 1908573 | 1909694 | fadE16 | BCG_1717 | 1122 | 1880356 | 1881477 | fadE16 | BCGDan_1715 | 1122 | 1880461 | 1881585 | fadE16 | BCGDan_sapMKO_1714 | 1125 |
| 1904429 | 1905253 | Rv1680 | Rv1680 | 825 | 1909703 | 1910527 | BCG_1718 | BCG_1718 | 825 | 1881486 | 1882310 | BCGDan_1716 | BCGDan_1716 | 825 | 1881594 | 1882418 | BCGDan_sapMKO_1715 | BCGDan_sapMKO_1715 | 825 |
| 1905250 | 1906242 | moeX | Rv1681 | 993 | 1910524 | 1911516 | moeX | BCG_1719 | 993 | 1882307 | 1883299 | moeX | BCGDan_1717 | 993 | 1882412 | 1883407 | moeX | BCGDan_sapMKO_1716 | 996 |
| 1906403 | 1907320 | Rv1682 | Rv1682 | 918 | 1911677 | 1912594 | BCG_1720 | BCG_1720 | 918 | 1883460 | 1884377 | BCGDan_1718 | BCGDan_1718 | 918 | 1883637 | 1884485 | BCGDan_sapMKO_1717 | BCGDan_sapMKO_1717 | 849 |
| 1907594 | 1910593 | Rv1683 | Rv1683 | 3000 | 1912812 | 1915811 | BCG_1721 | BCG_1721 | 3000 | 1884595 | 1887594 | BCGDan_1719 | BCGDan_1719 | 3000 | 1884703 | 1887702 | BCGDan_sapMKO_1718 | BCGDan_sapMKO_1718 | 3000 |
| 1910586 | 1910810 | Rv1684 | Rv1684 | 225 | 1915804 | 1916028 | BCG_1722 | BCG_1722 | 225 | 1887587 | 1887811 | BCGDan_1720 | BCGDan_1720 | 225 | 1887695 | 1887919 | BCGDan_sapMKO_1719 | BCGDan_sapMKO_1719 | 225 |
| 1910776 | 1911399 | Rv1685c | Rv1685c | 624 | 1915994 | 1916617 | BCG_1723c | BCG_1723c | 624 | 1887777 | 1888400 | BCGDan_1721 | BCGDan_1721 | 624 | 1887885 | 1888508 | BCGDan_sapMKO_1720 | BCGDan_sapMKO_1720 | 624 |
| 1911401 | 1912081 | Rv1686c | Rv1686c | 681 | 1916619 | 1917299 | BCG_1724c | BCG_1724c | 681 | 1888402 | 1889082 | BCGDan_1722 | BCGDan_1722 | 681 | 1888510 | 1889190 | BCGDan_sapMKO_1721 | BCGDan_sapMKO_1721 | 681 |
| 1912153 | 1912920 | Rv1687c | Rv1687c | 768 | 1917371 | 1918138 | BCG_1725c | BCG_1725c | 768 | 1889154 | 1889921 | BCGDan_1723 | BCGDan_1723 | 768 | 1889262 | 1890029 | BCGDan_sapMKO_1722 | BCGDan_sapMKO_1722 | 768 |
| 1912979 | 1913590 | mpg | Rv1688 | 612 | 1918197 | 1918808 | BCG_1726 | BCG_1726 | 612 | 1889980 | 1890591 | BCGDan_1724 | BCGDan_1724 | 612 | 1890088 | 1890699 | BCGDan_sapMKO_1723 | BCGDan_sapMKO_1723 | 612 |
| 1913602 | 1914876 | tyrS | Rv1689 | 1275 | 1918820 | 1920094 | tyrS | BCG_1727 | 1275 | 1890603 | 1891877 | tyrS | BCGDan_1725 | 1275 | 1890720 | 1891985 | tyrS | BCGDan_sapMKO_1724 | 1266 |
| 1915527 | 1915910 | lprJ | Rv1690 | 384 | 1920745 | 1921128 | lprJ | BCG_1728 | 384 | 1892528 | 1892911 | lprJ | BCGDan_1726 | 384 | 1892636 | 1893019 | lprJ | BCGDan_sapMKO_1725 | 384 |
| 1915949 | 1916701 | Rv1691 | Rv1691 | 753 | 1921167 | 1921919 | BCG_1729 | BCG_1729 | 753 | 1892950 | 1893702 | BCGDan_1727 | BCGDan_1727 | 753 | 1893058 | 1893810 | BCGDan_sapMKO_1726 | BCGDan_sapMKO_1726 | 753 |
| 1916698 | 1917759 | Rv1692 | Rv1692 | 1062 | 1921916 | 1922977 | BCG_1730 | BCG_1730 | 1062 | 1893699 | 1894760 | BCGDan_1728 | BCGDan_1728 | 1062 | 1893807 | 1894868 | BCGDan_sapMKO_1727 | BCGDan_sapMKO_1727 | 1062 |
| 1917756 | 1917932 | Rv1693 | Rv1693 | 177 | 1922974 | 1923150 | BCG_1731 | BCG_1731 | 177 | 1894757 | 1894933 | BCGDan_1729 | BCGDan_1729 | 177 | 1894865 | 1895041 | BCGDan_sapMKO_1728 | BCGDan_sapMKO_1728 | 177 |
| 1917940 | 1918746 | tlyA | Rv1694 | 807 | 1923158 | 1923964 | tlyA | BCG_1732 | 807 | 1894941 | 1895747 | tlyA | BCGDan_1730 | 807 | 1895049 | 1895855 | tlyA | BCGDan_sapMKO_1729 | 807 |
| 1918746 | 1919669 | ppnK | Rv1695 | 924 | 1923964 | 1924887 | ppnK | BCG_1733 | 924 | 1895747 | 1896670 | ppnK | BCGDan_1731 | 924 | 1895855 | 1896778 | ppnK | BCGDan_sapMKO_1730 | 924 |
| 1919683 | 1921446 | recN | Rv1696 | 1764 | 1924901 | 1926664 | recN | BCG_1734 | 1764 | 1896684 | 1898447 | recN | BCGDan_1732 | 1764 | 1896792 | 1898555 | recN | BCGDan_sapMKO_1731 | 1764 |
| 1921542 | 1922723 | Rv1697 | Rv1697 | 1182 | 1926760 | 1927941 | BCG_1735 | BCG_1735 | 1182 | 1898543 | 1899724 | BCGDan_1733 | BCGDan_1733 | 1182 | 1898657 | 1899832 | BCGDan_sapMKO_1732 | BCGDan_sapMKO_1732 | 1176 |
| 1922745 | 1923689 | mctB | Rv1698 | 945 | 1927963 | 1928907 | BCG_1736 | BCG_1736 | 945 | 1899746 | 1900690 | BCGDan_1734 | BCGDan_1734 | 945 | 1899854 | 1900798 | BCGDan_sapMKO_1733 | BCGDan_sapMKO_1733 | 945 |
| 1923829 | 1925589 | pyrG | Rv1699 | 1761 | 1929047 | 1930807 | pyrG | BCG_1737 | 1761 | 1900830 | 1902590 | pyrG | BCGDan_1735 | 1761 | 1901067 | 1902698 | pyrG | BCGDan_sapMKO_1734 | 1632 |
| 1925582 | 1926205 | Rv1700 | Rv1700 | 624 | 1930800 | 1931423 | BCG_1738 | BCG_1738 | 624 | 1902583 | 1903206 | BCGDan_1736 | BCGDan_1736 | 624 | 1902691 | 1903314 | BCGDan_sapMKO_1735 | BCGDan_sapMKO_1735 | 624 |
| 1926202 | 1927137 | Rv1701 | Rv1701 | 936 | 1931420 | 1932355 | BCG_1739 | BCG_1739 | 936 | 1903203 | 1904138 | BCGDan_1737 | BCGDan_1737 | 936 | 1903311 | 1904246 | BCGDan_sapMKO_1736 | BCGDan_sapMKO_1736 | 936 |
| 1927211 | 1928575 | Rv1702c | Rv1702c | 1365 | 1932429 | 1933793 | BCG_1740c | BCG_1740c | 1365 | 1904212 | 1905576 | BCGDan_1738 | BCGDan_1738 | 1365 | 1904320 | 1905684 | BCGDan_sapMKO_1737 | BCGDan_sapMKO_1737 | 1365 |

| | | | | | | | | | | | | | | | | | | | |
|---|---|---|---|---|---|---|---|---|---|---|---|---|---|---|---|---|---|---|---|
| 1929131 | 1929721 | Rv1703c | Rv1703c | 591 | 1934348 | 1934938 | BCG_1741c | BCG_1741c | 591 | 1906131 | 1906721 | BCGDan_1739 | BCGDan_1739 | 591 | 1906239 | 1906829 | BCGDan_sapMKO_1738 | BCGDan_sapMKO_1738 | 591 |
| 1929786 | 1931456 | cycA | Rv1704c | 1671 | 1935003 | 1936673 | cycA | BCG_1742c | 1671 | 1906786 | 1908456 | cycA | BCGDan_1740 | 1671 | 1906894 | 1908564 | cycA | BCGDan_sapMKO_1739 | 1671 |
| 1931497 | 1932654 | PPE22 | Rv1705c | 1158 | 1936714 | 1937871 | PPE22 | BCG_1743c | 1158 | 1908497 | 1909654 | PPE22 | BCGDan_1741 | 1158 | 1908605 | 1909720 | PPE22 | BCGDan_sapMKO_1740 | 1116 |
| 1932694 | 1933878 | PPE23 | Rv1706c | 1185 | 1937911 | 1939095 | PPE23 | BCG_1744c | 1185 | 1909694 | 1910878 | PPE23 | BCGDan_1742 | 1185 | 1909802 | 1910986 | PPE23 | BCGDan_sapMKO_1741 | 1185 |
| 1934482 | 1934649 | Rv1706A | Rv1706A | 168 | 1939699 | 1939866 | BCG_1745c | BCG_1745c | 168 | 1911482 | 1911649 | BCGDan_1743 | BCGDan_1743 | 168 | 1911590 | 1911757 | BCGDan_sapMKO_1742 | BCGDan_sapMKO_1742 | 168 |
| 1934882 | 1936342 | Rv1707 | Rv1707 | 1461 | 1940099 | 1941559 | BCG_1746 | BCG_1746 | 1461 | 1911882 | 1913342 | BCGDan_1744 | BCGDan_1744 | 1461 | 1912026 | 1913450 | BCGDan_sapMKO_1743 | BCGDan_sapMKO_1743 | 1425 |
| 1936360 | 1937316 | Rv1708 | Rv1708 | 957 | 1941577 | 1942533 | BCG_1747 | BCG_1747 | 957 | 1913360 | 1914316 | BCGDan_1745 | BCGDan_1745 | 957 | 1913561 | 1914424 | BCGDan_sapMKO_1744 | BCGDan_sapMKO_1744 | 864 |
| 1937313 | 1938149 | scpA | Rv1709 | 837 | 1942530 | 1943366 | BCG_1748 | BCG_1748 | 837 | 1914313 | 1915149 | BCGDan_1746 | BCGDan_1746 | 837 | 1914421 | 1915257 | BCGDan_sapMKO_1745 | BCGDan_sapMKO_1745 | 837 |
| 1938146 | 1938841 | scpB | Rv1710 | 696 | 1943363 | 1944058 | BCG_1749 | BCG_1749 | 696 | 1915146 | 1915841 | BCGDan_1747 | BCGDan_1747 | 696 | 1915254 | 1915949 | BCGDan_sapMKO_1746 | BCGDan_sapMKO_1746 | 696 |
| 1938838 | 1939602 | Rv1711 | Rv1711 | 765 | 1944055 | 1944819 | BCG_1750 | BCG_1750 | 765 | 1915841 | 1916602 | BCGDan_1748 | BCGDan_1748 | 762 | 1915949 | 1916710 | BCGDan_sapMKO_1747 | BCGDan_sapMKO_1747 | 762 |
| 1939599 | 1940291 | cmk | Rv1712 | 693 | 1944816 | 1945508 | cmk | BCG_1751 | 693 | 1916599 | 1917291 | cmk | BCGDan_1749 | 693 | 1916707 | 1917399 | cmk | BCGDan_sapMKO_1748 | 693 |
| 1940288 | 1941679 | engA | Rv1713 | 1392 | 1945505 | 1946896 | engA | BCG_1752 | 1392 | 1917288 | 1918679 | engA | BCGDan_1750 | 1392 | 1917396 | 1918787 | engA | BCGDan_sapMKO_1749 | 1392 |
| 1941853 | 1942665 | Rv1714 | Rv1714 | 813 | 1947070 | 1947882 | BCG_1753 | BCG_1753 | 813 | 1918853 | 1919665 | BCGDan_1751 | BCGDan_1751 | 813 | 1918970 | 1919773 | BCGDan_sapMKO_1750 | BCGDan_sapMKO_1750 | 804 |
| 1942659 | 1943573 | fadB3 | Rv1715 | 915 | 1947876 | 1948790 | fadB3 | BCG_1754 | 915 | 1919659 | 1920573 | fadB3 | BCGDan_1752 | 915 | 1919767 | 1920681 | fadB3 | BCGDan_sapMKO_1751 | 915 |
| 1943576 | 1944406 | Rv1716 | Rv1716 | 831 | 1948793 | 1949623 | BCG_1755 | BCG_1755 | 831 | 1920576 | 1921406 | BCGDan_1753 | BCGDan_1753 | 831 | 1920684 | 1921514 | BCGDan_sapMKO_1752 | BCGDan_sapMKO_1752 | 831 |
| 1944406 | 1944756 | Rv1717 | Rv1717 | 351 | 1949623 | 1949973 | BCG_1756 | BCG_1756 | 351 | 1921406 | 1921756 | BCGDan_1754 | BCGDan_1754 | 351 | 1921550 | 1921864 | BCGDan_sapMKO_1753 | BCGDan_sapMKO_1753 | 315 |
| 1944809 | 1945627 | Rv1718 | Rv1718 | 819 | 1950026 | 1950649 | BCG_1757 | BCG_1757 | 624 | 1921809 | 1922432 | BCGDan_1755 | BCGDan_1755 | 624 | 1921917 | 1922540 | BCGDan_sapMKO_1754 | BCGDan_sapMKO_1754 | 624 |
| 1945641 | 1946420 | Rv1719 | Rv1719 | 780 | 1950857 | 1951636 | BCG_1758 | BCG_1758 | 780 | 1922640 | 1923419 | BCGDan_1756 | BCGDan_1756 | 780 | 1922748 | 1923527 | BCGDan_sapMKO_1755 | BCGDan_sapMKO_1755 | 780 |
| 1947030 | 1947419 | vapC12 | Rv1720c | 390 | 1952246 | 1952635 | BCG_1759c | BCG_1759c | 390 | 1924029 | 1924418 | BCGDan_1758 | BCGDan_1758 | 390 | 1924137 | 1924526 | BCGDan_sapMKO_1757 | BCGDan_sapMKO_1757 | 390 |
| 1947416 | 1947643 | vapB12 | Rv1721c | 228 | 1952632 | 1952859 | BCG_1760c | BCG_1760c | 228 | 1924415 | 1924642 | BCGDan_1759 | BCGDan_1759 | 228 | 1924523 | 1924750 | BCGDan_sapMKO_1758 | BCGDan_sapMKO_1758 | 228 |
| 1947861 | 1949345 | Rv1722 | Rv1722 | 1485 | 1953077 | 1954561 | BCG_1761 | BCG_1761 | 1485 | 1924860 | 1926344 | BCGDan_1760 | BCGDan_1760 | 1485 | 1924968 | 1926452 | BCGDan_sapMKO_1759 | BCGDan_sapMKO_1759 | 1485 |
| 1949342 | 1950589 | Rv1723 | Rv1723 | 1248 | 1954558 | 1955805 | BCG_1762 | BCG_1762 | 1248 | 1926341 | 1927588 | BCGDan_1761 | BCGDan_1761 | 1248 | 1926449 | 1927696 | BCGDan_sapMKO_1760 | BCGDan_sapMKO_1760 | 1248 |
| 1950632 | 1951051 | Rv1724c | Rv1724c | 420 | 1955848 | 1956267 | BCG_1763c | BCG_1763c | 420 | 1927631 | 1928050 | BCGDan_1762 | BCGDan_1762 | 420 | 1927739 | 1928158 | BCGDan_sapMKO_1761 | BCGDan_sapMKO_1761 | 420 |
| 1951041 | 1951751 | Rv1725c | Rv1725c | 711 | 1956257 | 1956967 | BCG_1764c | BCG_1764c | 711 | 1928040 | 1928750 | BCGDan_1763 | BCGDan_1763 | 711 | 1928148 | 1928858 | BCGDan_sapMKO_1762 | BCGDan_sapMKO_1762 | 711 |
| 1951852 | 1953237 | Rv1726 | Rv1726 | 1386 | 1957068 | 1958453 | BCG_1765 | BCG_1765 | 1386 | 1928851 | 1930236 | BCGDan_1764 | BCGDan_1764 | 1386 | 1928959 | 1930344 | BCGDan_sapMKO_1763 | BCGDan_sapMKO_1763 | 1386 |
| 1953270 | 1953839 | Rv1727 | Rv1727 | 570 | 1958486 | 1959055 | BCG_1766 | BCG_1766 | 570 | 1930269 | 1930838 | BCGDan_1765 | BCGDan_1765 | 570 | 1930377 | 1930946 | BCGDan_sapMKO_1764 | BCGDan_sapMKO_1764 | 570 |
| 1953864 | 1954634 | Rv1728c | Rv1728c | 771 | 1959080 | 1959850 | BCG_1767c | BCG_1767c | 771 | 1930863 | 1931633 | BCGDan_1766 | BCGDan_1766 | 771 | 1930971 | 1931741 | BCGDan_sapMKO_1765 | BCGDan_sapMKO_1765 | 771 |
| 1954631 | 1955569 | Rv1729c | Rv1729c | 939 | 1959847 | 1960785 | BCG_1768c | BCG_1768c | 939 | 1931630 | 1932568 | BCGDan_1767 | BCGDan_1767 | 939 | 1931738 | 1932676 | BCGDan_sapMKO_1766 | BCGDan_sapMKO_1766 | 939 |
| 1955692 | 1957245 | Rv1730c | Rv1730c | 1554 | 1960851 | 1962404 | BCG_1769c | BCG_1769c | 1554 | 1932634 | 1934187 | BCGDan_1768 | BCGDan_1768 | 1554 | 1932742 | 1934166 | BCGDan_sapMKO_1767 | BCGDan_sapMKO_1767 | 1425 |
| 1957677 | 1959233 | gabD2 | Rv1731 | 1557 | 1962836 | 1964392 | gabD2 | BCG_1770 | 1557 | 1934619 | 1936175 | gabD2 | BCGDan_1769 | 1557 | 1934727 | 1936283 | gabD2 | BCGDan_sapMKO_1768 | 1557 |
| 1959243 | 1959791 | Rv1732c | Rv1732c | 549 | 1964402 | 1964950 | BCG_1771c | BCG_1771c | 549 | 1936185 | 1936733 | BCGDan_1770 | BCGDan_1770 | 549 | 1936293 | 1936841 | BCGDan_sapMKO_1769 | BCGDan_sapMKO_1769 | 549 |
| 1959855 | 1960487 | Rv1733c | Rv1733c | 633 | 1965014 | 1965646 | BCG_1772c | BCG_1772c | 633 | 1936797 | 1937429 | BCGDan_1771 | BCGDan_1771 | 633 | 1936905 | 1937537 | BCGDan_sapMKO_1770 | BCGDan_sapMKO_1770 | 633 |
| 1960774 | 1961016 | Rv1734c | Rv1734c | 243 | 1965933 | 1966175 | BCG_1773c | BCG_1773c | 243 | 1937716 | 1937958 | BCGDan_1772 | BCGDan_1772 | 243 | 1937824 | 1938066 | BCGDan_sapMKO_1771 | BCGDan_sapMKO_1771 | 243 |
| 1961291 | 1961788 | Rv1735c | Rv1735c | 498 | 1966450 | 1966947 | BCG_1774c | BCG_1774c | 498 | 1938233 | 1938730 | BCGDan_1773 | BCGDan_1773 | 498 | 1938341 | 1938838 | BCGDan_sapMKO_1772 | BCGDan_sapMKO_1772 | 498 |
| 1962228 | 1964186 | narX | Rv1736c | 1959 | 1967387 | 1969345 | narX | BCG_1775c | 1959 | 1939170 | 1941128 | narX | BCGDan_1774 | 1959 | 1939278 | 1941236 | narX | BCGDan_sapMKO_1773 | 1959 |
| 1964183 | 1965370 | narK2 | Rv1737c | 1188 | 1969342 | 1970529 | narK2 | BCG_1776c | 1188 | 1941125 | 1942312 | narK2 | BCGDan_1775 | 1188 | 1941233 | 1942396 | narK2 | BCGDan_sapMKO_1774 | 1164 |
| 1965657 | 1965941 | Rv1738 | Rv1738 | 285 | 1970816 | 1971100 | BCG_1777 | BCG_1777 | 285 | 1942599 | 1942883 | BCGDan_1776 | BCGDan_1776 | 285 | 1942659 | 1942991 | BCGDan_sapMKO_1775 | BCGDan_sapMKO_1775 | 333 |
| 1965955 | 1967637 | Rv1739c | Rv1739c | 1683 | 1971114 | 1972796 | BCG_1778c | BCG_1778c | 1683 | 1942897 | 1944579 | BCGDan_1777 | BCGDan_1777 | 1683 | 1943005 | 1944687 | BCGDan_sapMKO_1776 | BCGDan_sapMKO_1776 | 1683 |
| 1967705 | 1967917 | vapB34 | Rv1740 | 213 | 1972864 | 1973076 | BCG_1779 | BCG_1779 | 213 | 1944647 | 1944859 | BCGDan_1778 | BCGDan_1778 | 213 | 1944773 | 1944967 | BCGDan_sapMKO_1777 | BCGDan_sapMKO_1777 | 195 |
| 1967917 | 1968165 | vapC34 | Rv1741 | 249 | 1973076 | 1973324 | BCG_1780 | BCG_1780 | 249 | 1944859 | 1945107 | BCGDan_1779 | BCGDan_1779 | 249 | 1944967 | 1945215 | BCGDan_sapMKO_1778 | BCGDan_sapMKO_1778 | 249 |
| 1968173 | 1968910 | Rv1742 | Rv1742 | 738 | 1973405 | 1974070 | BCG_1781 | BCG_1781 | 666 | 1945188 | 1945853 | BCGDan_1780 | BCGDan_1780 | 666 | 1945296 | 1945961 | BCGDan_sapMKO_1779 | BCGDan_sapMKO_1779 | 666 |
| 1969004 | 1970704 | pknE | Rv1743 | 1701 | 1974164 | 1975864 | pknE | BCG_1782 | 1701 | 1945947 | 1947647 | pknE | BCGDan_1781 | 1701 | 1946055 | 1947755 | pknE | BCGDan_sapMKO_1780 | 1701 |
| 1970989 | 1971390 | Rv1744c | Rv1744c | 402 | 1976149 | 1976550 | BCG_1783c | BCG_1783c | 402 | 1947932 | 1948333 | BCGDan_1782 | BCGDan_1782 | 402 | 1948040 | 1948441 | BCGDan_sapMKO_1781 | BCGDan_sapMKO_1781 | 402 |
| 1971380 | 1971991 | idi | Rv1745c | 612 | 1976621 | 1977151 | idi | BCG_1784c | 531 | 1948404 | 1948934 | idi | BCGDan_1783 | 531 | 1948512 | 1949042 | idi | BCGDan_sapMKO_1782 | 531 |
| 1972138 | 1973568 | pknF | Rv1746 | 1431 | 1977298 | 1978728 | pknF | BCG_1785 | 1431 | 1949081 | 1950511 | pknF | BCGDan_1784 | 1431 | 1949189 | 1950619 | pknF | BCGDan_sapMKO_1783 | 1431 |
| 1973630 | 1976227 | Rv1747 | Rv1747 | 2598 | 1978790 | 1981387 | BCG_1786 | BCG_1786 | 2598 | 1950573 | 1953170 | BCGDan_1785 | BCGDan_1785 | 2598 | 1950687 | 1953278 | BCGDan_sapMKO_1784 | BCGDan_sapMKO_1784 | 2592 |
| 1976600 | 1977331 | Rv1748 | Rv1748 | 732 | 1981760 | 1982491 | BCG_1787 | BCG_1787 | 732 | 1953543 | 1954274 | BCGDan_1786 | BCGDan_1786 | 732 | 1953711 | 1954382 | BCGDan_sapMKO_1785 | BCGDan_sapMKO_1785 | 672 |
| 1977328 | 1977885 | Rv1749c | Rv1749c | 558 | 1982488 | 1983045 | BCG_1788c | BCG_1788c | 558 | 1954271 | 1954828 | BCGDan_1787 | BCGDan_1787 | 558 | 1954379 | 1954870 | BCGDan_sapMKO_1786 | BCGDan_sapMKO_1786 | 492 |
| 1977969 | 1979567 | fadD1 | Rv1750c | 1599 | 1983129 | 1984727 | fadD1 | BCG_1789c | 1599 | 1954912 | 1956510 | fadD1 | BCGDan_1788 | 1599 | 1955020 | 1956618 | fadD1 | BCGDan_sapMKO_1787 | 1599 |
| 1979621 | 1981003 | Rv1751 | Rv1751 | 1383 | 1984781 | 1986163 | BCG_1790 | BCG_1790 | 1383 | 1956564 | 1957946 | BCGDan_1789 | BCGDan_1789 | 1383 | 1956816 | 1958054 | BCGDan_sapMKO_1788 | BCGDan_sapMKO_1788 | 1239 |
| 1981130 | 1981579 | Rv1752 | Rv1752 | 450 | 1986290 | 1986739 | BCG_1791 | BCG_1791 | 450 | 1958073 | 1958522 | BCGDan_1790 | BCGDan_1790 | 450 | 1958181 | 1958630 | BCGDan_sapMKO_1789 | BCGDan_sapMKO_1789 | 450 |
| 1981614 | 1984775 | PPE24 | Rv1753c | 3162 | 1986774 | 1989854 | PPE24 | BCG_1792c | 3081 | 1958557 | 1961484 | PPE24 | BCGDan_1791 | 2928 | 1958665 | 1961745 | PPE24 | BCGDan_sapMKO_1790 | 3081 |
| 1984979 | 1986670 | Rv1754c | Rv1754c | 1692 | 1990058 | 1991749 | BCG_1793c | BCG_1793c | 1692 | 1961688 | 1963379 | BCGDan_1792 | BCGDan_1792 | 1692 | 1961949 | 1963640 | BCGDan_sapMKO_1791 | BCGDan_sapMKO_1791 | 1692 |
| 1986854 | 1987696 | plcD | Rv1755c | 843 | 1991933 | 1993477 | plcD | BCG_1794c | 1545 | 1963563 | 1965107 | plcD | BCGDan_1793 | 1545 | 1963824 | 1965401 | plcD | BCGDan_sapMKO_1792 | 1578 |
| NA | NA | NA | NA | NA | 1993606 | 1994790 | BCG_1795c | BCG_1795c | 1185 | 1965236 | 1966420 | BCGDan_1794 | BCGDan_1794 | 1185 | 1965497 | 1966681 | BCGDan_sapMKO_1793 | BCGDan_sapMKO_1793 | 1185 |
| NA | NA | NA | NA | NA | 1994940 | 1996079 | BCG_1796 | BCG_1796 | 1140 | 1966609 | 1967709 | BCGDan_1795 | BCGDan_1795 | 1101 | 1966867 | 1967970 | BCGDan_sapMKO_1794 | BCGDan_sapMKO_1794 | 1104 |
| NA | NA | NA | NA | NA | 1996280 | 1999117 | mmpL14 | BCG_1797 | 2838 | 1967910 | 1970747 | mmpL14 | BCGDan_1796 | 2838 | 1968171 | 1971008 | mmpL14 | BCGDan_sapMKO_1795 | 2838 |
| 1989042 | 1989566 | cut1 | Rv1758 | 525 | 1999438 | 2000094 | cut1 | BCG_1798 | 657 | 1971068 | 1971724 | cut1 | BCGDan_1797 | 657 | 1971329 | 1971985 | cut1 | BCGDan_sapMKO_1796 | 657 |
| 1989833 | 1992577 | wag22 | Rv1759c | 2745 | 2000361 | 2002769 | wag22b | BCG_1799c | 2409 | 1971991 | 1974402 | wag22b | BCGDan_1798 | 2412 | 1972252 | 1974663 | wag22b | BCGDan_sapMKO_1797 | 2412 |
| NA | NA | NA | NA | NA | 2002766 | 2003050 | wag22a | BCG_1800c | 285 | 1974399 | 1974683 | wag22a | BCGDan_1799 | 285 | 1974660 | 1974944 | wag22a | BCGDan_sapMKO_1798 | 285 |
| 1993153 | 1994661 | Rv1760 | Rv1760 | 1509 | 2003626 | 2005155 | BCG_1801 | BCG_1801 | 1530 | 1975259 | 1976788 | BCGDan_1800 | BCGDan_1800 | 1530 | 1975622 | 1977049 | BCGDan_sapMKO_1799 | BCGDan_sapMKO_1799 | 1428 |
| 1994671 | 1995054 | Rv1761c | Rv1761c | 384 | 2005143 | 2005526 | BCG_1802c | BCG_1802c | 384 | 1976776 | 1977159 | BCGDan_1801 | BCGDan_1801 | 384 | 1977037 | 1977420 | BCGDan_sapMKO_1800 | BCGDan_sapMKO_1800 | 384 |
| 1995054 | 1995842 | Rv1762c | Rv1762c | 789 | 2005526 | 2006314 | BCG_1803c | BCG_1803c | 789 | 1977159 | 1977947 | BCGDan_1802 | BCGDan_1802 | 789 | 1977420 | 1978208 | BCGDan_sapMKO_1801 | BCGDan_sapMKO_1801 | 789 |
| NA | NA | NA | NA | NA | 2006454 | 2006921 | BCG_1804c | BCG_1804c | 468 | 1978087 | 1978554 | BCGDan_1803 | BCGDan_1803 | 468 | 1978348 | 1978815 | BCGDan_sapMKO_1802 | BCGDan_sapMKO_1802 | 468 |
| NA | NA | NA | NA | NA | 2007019 | 2007171 | BCG_1805c | BCG_1805c | 153 | 1978652 | 1978804 | BCGDan_1804 | BCGDan_1804 | 153 | 1978913 | 1979065 | BCGDan_sapMKO_1803 | BCGDan_sapMKO_1803 | 153 |
| 1997418 | 1998515 | Rv1765c | Rv1765c | 1098 | NA | NA | NA | NA | NA | 1978965 | 1980221 | Rv1313c | BCGDan_1805 | 1257 | 1979226 | 1980482 | Rv1313c | BCGDan_sapMKO_1804 | 1257 |
| 1999142 | 1999357 | Rv1765A | Rv1765A | 216 | NA | NA | NA | NA | NA | 1980848 | 1981063 | BCGDan_1806 | BCGDan_1806 | 216 | 1981109 | 1981324 | BCGDan_sapMKO_1805 | BCGDan_sapMKO_1805 | 216 |
| 1999737 | 2000006 | Rv1766 | Rv1766 | 270 | NA | NA | NA | NA | NA | 1981443 | 1981712 | BCGDan_1807 | BCGDan_1807 | 270 | 1981704 | 1981973 | BCGDan_sapMKO_1806 | BCGDan_sapMKO_1806 | 270 |
| 2000074 | 2000433 | Rv1767 | Rv1767 | 360 | NA | NA | NA | NA | NA | 1981780 | 1982139 | BCGDan_1808 | BCGDan_1808 | 360 | 1982041 | 1982400 | BCGDan_sapMKO_1807 | BCGDan_sapMKO_1807 | 360 |
| 2000614 | 2002470 | PE_PGRS31 | Rv1768 | 1857 | NA | NA | NA | NA | NA | 1982320 | 1984194 | PE_PGRS31 | BCGDan_1809 | 1875 | 1982581 | 1984455 | PE_PGRS31 | BCGDan_sapMKO_1808 | 1875 |
| 2002626 | 2003870 | Rv1769 | Rv1769 | 1245 | NA | NA | NA | NA | NA | 1984350 | 1985594 | BCGDan_1810 | BCGDan_1810 | 1245 | 1984611 | 1985855 | BCGDan_sapMKO_1809 | BCGDan_sapMKO_1809 | 1245 |
| 2003878 | 2005164 | Rv1770 | Rv1770 | 1287 | NA | NA | NA | NA | NA | 1985602 | 1986888 | BCGDan_1811 | BCGDan_1811 | 1287 | 1985863 | 1987149 | BCGDan_sapMKO_1810 | BCGDan_sapMKO_1810 | 1287 |
| 2005161 | 2006447 | Rv1771 | Rv1771 | 1287 | NA | NA | NA | NA | NA | 1986885 | 1988171 | BCGDan_1812 | BCGDan_1812 | 1287 | 1987146 | 1988432 | BCGDan_sapMKO_1811 | BCGDan_sapMKO_1811 | 1287 |
| 2006636 | 2006947 | Rv1772 | Rv1772 | 312 | NA | NA | NA | NA | NA | 1988360 | 1988671 | BCGDan_1813 | BCGDan_1813 | 312 | 1988621 | 1988932 | BCGDan_sapMKO_1812 | BCGDan_sapMKO_1812 | 312 |
| 2007020 | 2007766 | Rv1773c | Rv1773c | 747 | 2007332 | 2008771 | BCG_1806c | BCG_1806c | 1440 | 1988744 | 1989490 | BCGDan_1814 | BCGDan_1814 | 747 | 1989005 | 1989751 | BCGDan_sapMKO_1813 | BCGDan_sapMKO_1813 | 747 |
| 2007832 | 2009172 | Rv1774 | Rv1774 | 1341 | 2008837 | 2010177 | BCG_1807 | BCG_1807 | 1341 | 1989556 | 1990896 | BCGDan_1815 | BCGDan_1815 | 1341 | 1989817 | 1991157 | BCGDan_sapMKO_1814 | BCGDan_sapMKO_1814 | 1341 |
| 2009172 | 2009990 | Rv1775 | Rv1775 | 819 | 2010177 | 2010995 | BCG_1808 | BCG_1808 | 819 | 1990896 | 1991714 | BCGDan_1816 | BCGDan_1816 | 819 | 1991157 | 1991975 | BCGDan_sapMKO_1815 | BCGDan_sapMKO_1815 | 819 |
| 2009995 | 2010555 | Rv1776c | Rv1776c | 561 | 2011000 | 2011560 | BCG_1809c | BCG_1809c | 561 | 1991719 | 1992279 | BCGDan_1817 | BCGDan_1817 | 561 | 1991980 | 1992540 | BCGDan_sapMKO_1816 | BCGDan_sapMKO_1816 | 561 |
| 2010656 | 2011960 | cyp144 | Rv1777 | 1305 | 2011661 | 2012965 | cyp144 | BCG_1810 | 1305 | 1992380 | 1993684 | cyp144 | BCGDan_1818 | 1305 | 1992731 | 1993945 | cyp144 | BCGDan_sapMKO_1817 | 1215 |
| 2012081 | 2012530 | Rv1778c | Rv1778c | 450 | 2013086 | 2013535 | BCG_1811c | BCG_1811c | 450 | 1993805 | 1994254 | BCGDan_1819 | BCGDan_1819 | 450 | 1994066 | 1994572 | BCGDan_sapMKO_1818 | BCGDan_sapMKO_1818 | 507 |
| 2012686 | 2014479 | Rv1779c | Rv1779c | 1794 | 2013691 | 2015484 | BCG_1812c | BCG_1812c | 1794 | 1994410 | 1996203 | BCGDan_1820 | BCGDan_1820 | 1794 | 1994671 | 1996464 | BCGDan_sapMKO_1819 | BCGDan_sapMKO_1819 | 1794 |
| 2014699 | 2015262 | Rv1780 | Rv1780 | 564 | 2015704 | 2016267 | BCG_1813 | BCG_1813 | 564 | 1996423 | 1996986 | BCGDan_1821 | BCGDan_1821 | 564 | 1996684 | 1997247 | BCGDan_sapMKO_1820 | BCGDan_sapMKO_1820 | 564 |
| 2015302 | 2017476 | malQ | Rv1781c | 2175 | 2016307 | 2018481 | malQ | BCG_1814c | 2175 | 1997026 | 1999200 | malQ | BCGDan_1822 | 2175 | 1997287 | 1999470 | malQ | BCGDan_sapMKO_1821 | 2184 |
| 2017740 | 2019260 | eccB5 | Rv1782 | 1521 | 2018745 | 2020265 | BCG_1815 | BCG_1815 | 1521 | 1999464 | 2000984 | BCGDan_1823 | BCGDan_1823 | 1521 | 1999854 | 2001245 | BCGDan_sapMKO_1822 | BCGDan_sapMKO_1822 | 1392 |
| 2019257 | 2023432 | eccC5 | Rv1783 | 4176 | 2020262 | 2024437 | BCG_1816 | BCG_1816 | 4176 | 2000981 | 2005156 | BCGDan_1824 | BCGDan_1824 | 4176 | 2001242 | 2005417 | BCGDan_sapMKO_1823 | BCGDan_sapMKO_1823 | 4176 |
| 2023447 | 2024628 | cyp143 | Rv1785c | 1182 | 2024452 | 2025633 | cyp143 | BCG_1817c | 1182 | 2005171 | 2006352 | cyp143 | BCGDan_1825 | 1182 | 2005432 | 2006613 | cyp143 | BCGDan_sapMKO_1824 | 1182 |

| | | | | | | | | | | | | | | | | | | | |
|---|---|---|---|---|---|---|---|---|---|---|---|---|---|---|---|---|---|---|---|
| 2024828 | 2025031 | Rv1786 | Rv1786 | 204 | 2025833 | 2026036 | BCG_1818 | BCG_1818 | 204 | 2006552 | 2006755 | BCGDan_1826 | BCGDan_1826 | 204 | 2006843 | 2007016 | BCGDan_sapMKO_1825 | BCGDan_sapMKO_1825 | 174 |
| 2025301 | 2026398 | PPE25 | Rv1787 | 1098 | 2026306 | 2027400 | PPE25 | BCG_1819 | 1095 | 2007025 | 2008119 | PPE25 | BCGDan_1827 | 1095 | 2007268 | 2008380 | PPE25 | BCGDan_sapMKO_1826 | 1113 |
| 2026477 | 2026776 | PE18 | Rv1788 | 300 | 2027479 | 2027778 | PE18 | BCG_1820 | 300 | 2008198 | 2008497 | PE18 | BCGDan_1828 | 300 | 2008459 | 2008758 | PE18 | BCGDan_sapMKO_1827 | 300 |
| 2026790 | 2027971 | PPE26 | Rv1789 | 1182 | 2027792 | 2028973 | PPE26 | BCG_1821 | 1182 | 2008511 | 2009692 | PPE26 | BCGDan_1829 | 1182 | 2008772 | 2009953 | PPE26 | BCGDan_sapMKO_1828 | 1182 |
| 2028425 | 2029477 | PPE27 | Rv1790 | 1053 | 2029427 | 2030479 | PPE27 | BCG_1822 | 1053 | 2010146 | 2011198 | PPE27 | BCGDan_1830 | 1053 | 2010407 | 2011459 | PPE27 | BCGDan_sapMKO_1829 | 1053 |
| 2029904 | 2030203 | PE19 | Rv1791 | 300 | 2030906 | 2031205 | PE19 | BCG_1823 | 300 | 2011625 | 2011924 | PE19 | BCGDan_1831 | 300 | 2011886 | 2012185 | PE19 | BCGDan_sapMKO_1830 | 300 |
| 2030347 | 2030643 | esxM | Rv1792 | 297 | 2031349 | 2031645 | esxM | BCG_1824 | 297 | 2012068 | 2012364 | esxM | BCGDan_1832 | 297 | 2012344 | 2012625 | esxM | BCGDan_sapMKO_1831 | 282 |
| 2030694 | 2030978 | esxN | Rv1793 | 285 | 2031696 | 2031980 | esxN | BCG_1825 | 285 | 2012415 | 2012699 | esxN | BCGDan_1833 | 285 | 2012676 | 2012960 | esxN | BCGDan_sapMKO_1832 | 285 |
| 2031066 | 2031968 | Rv1794 | Rv1794 | 903 | 2032068 | 2032970 | BCG_1826 | BCG_1826 | 903 | 2012787 | 2013689 | BCGDan_1834 | BCGDan_1834 | 903 | 2013048 | 2013950 | BCGDan_sapMKO_1833 | BCGDan_sapMKO_1833 | 903 |
| 2032240 | 2033751 | eccD5 | Rv1795 | 1512 | 2033242 | 2034753 | BCG_1827 | BCG_1827 | 1512 | 2013961 | 2015472 | BCGDan_1835 | BCGDan_1835 | 1512 | 2014288 | 2015733 | BCGDan_sapMKO_1834 | BCGDan_sapMKO_1834 | 1446 |
| 2033729 | 2035486 | mycP5 | Rv1796 | 1758 | 2034731 | 2036488 | BCG_1828 | BCG_1828 | 1758 | 2015450 | 2017207 | BCGDan_1836 | BCGDan_1836 | 1758 | 2015711 | 2017468 | BCGDan_sapMKO_1835 | BCGDan_sapMKO_1835 | 1758 |
| 2035483 | 2036703 | eccE5 | Rv1797 | 1221 | 2036485 | 2037705 | BCG_1829 | BCG_1829 | 1221 | 2017204 | 2018424 | BCGDan_1837 | BCGDan_1837 | 1221 | 2017465 | 2018685 | BCGDan_sapMKO_1836 | BCGDan_sapMKO_1836 | 1221 |
| 2036700 | 2038532 | eccA5 | Rv1798 | 1833 | 2037702 | 2039534 | BCG_1830 | BCG_1830 | 1833 | 2018421 | 2020253 | BCGDan_1838 | BCGDan_1838 | 1833 | 2018682 | 2020514 | BCGDan_sapMKO_1837 | BCGDan_sapMKO_1837 | 1833 |
| 2039159 | 2039350 | lppT | Rv1799 | 192 | 2040161 | 2040352 | lppT | BCG_1831 | 192 | 2020880 | 2021071 | lppT | BCGDan_1839 | 192 | 2021141 | 2021332 | lppT | BCGDan_sapMKO_1838 | 192 |
| 2039453 | 2041420 | PPE28 | Rv1800 | 1968 | 2040455 | 2042422 | PPE28 | BCG_1832 | 1968 | 2021174 | 2023141 | PPE28 | BCGDan_1840 | 1968 | 2021435 | 2023402 | PPE28 | BCGDan_sapMKO_1839 | 1968 |
| 2042001 | 2043272 | PPE29 | Rv1801 | 1272 | 2043003 | 2044274 | PPE29 | BCG_1833 | 1272 | 2023722 | 2024993 | PPE29 | BCGDan_1841 | 1272 | 2023983 | 2025254 | PPE29 | BCGDan_sapMKO_1840 | 1272 |
| 2043384 | 2044775 | PPE30 | Rv1802 | 1392 | 2044386 | 2045777 | PPE30 | BCG_1834 | 1392 | 2025105 | 2026496 | PPE30 | BCGDan_1842 | 1392 | 2025366 | 2026757 | PPE30 | BCGDan_sapMKO_1841 | 1392 |
| 2044923 | 2046842 | PE_PGRS32 | Rv1803c | 1920 | 2045925 | 2047547 | PE_PGRS32b | BCG_1835c | 1623 | 2026644 | 2028266 | PE_PGRS32b | BCGDan_1843 | 1623 | 2026905 | 2028518 | PE_PGRS32b | BCGDan_sapMKO_1842 | 1614 |
| NA | NA | NA | NA | NA | 2047581 | 2047844 | PE_PGRS32a | BCG_1836c | 264 | 2028300 | 2028611 | PE_PGRS32a | BCGDan_1844 | 312 | 2028561 | 2028857 | PE_PGRS32a | BCGDan_sapMKO_1843 | 297 |
| 2047023 | 2047349 | Rv1804c | Rv1804c | 327 | 2048025 | 2048351 | BCG_1837c | BCG_1837c | 327 | 2028744 | 2029070 | BCGDan_1845 | BCGDan_1845 | 327 | 2029005 | 2029322 | BCGDan_sapMKO_1844 | BCGDan_sapMKO_1844 | 318 |
| 2047687 | 2048034 | Rv1805c | Rv1805c | 348 | 2048689 | 2049036 | BCG_1838c | BCG_1838c | 348 | 2029408 | 2029755 | BCGDan_1846 | BCGDan_1846 | 348 | 2029669 | 2030016 | BCGDan_sapMKO_1845 | BCGDan_sapMKO_1845 | 348 |
| 2048072 | 2048371 | PE20 | Rv1806 | 300 | 2049074 | 2049373 | PE20 | BCG_1839 | 300 | 2029793 | 2030092 | PE20 | BCGDan_1847 | 300 | 2030054 | 2030353 | PE20 | BCGDan_sapMKO_1846 | 300 |
| 2048398 | 2049597 | PPE31 | Rv1807 | 1200 | 2049400 | 2050599 | PPE31 | BCG_1840 | 1200 | 2030119 | 2031318 | PPE31 | BCGDan_1848 | 1200 | 2030380 | 2031579 | PPE31 | BCGDan_sapMKO_1847 | 1200 |
| 2049921 | 2051150 | PPE32 | Rv1808 | 1230 | 2050923 | 2052152 | PPE32 | BCG_1841 | 1230 | 2031642 | 2032871 | PPE32 | BCGDan_1849 | 1230 | 2031900 | 2033132 | PPE32 | BCGDan_sapMKO_1848 | 1233 |
| 2051282 | 2052688 | PPE33 | Rv1809 | 1407 | 2052284 | 2052847 | PPE33a | BCG_1842 | 564 | 2033003 | 2033566 | PPE33a | BCGDan_1850 | 564 | 2033264 | 2033827 | PPE33a | BCGDan_sapMKO_1849 | 564 |
| 2053443 | 2054147 | mgtC | Rv1811 | 705 | 2054445 | 2055149 | BCG_1845 | BCG_1845 | 705 | 2034436 | 2035140 | BCGDan_1851 | BCGDan_1851 | 705 | 2034697 | 2035401 | BCGDan_sapMKO_1851 | BCGDan_sapMKO_1851 | 705 |
| 2054157 | 2055359 | Rv1812c | Rv1812c | 1203 | 2055159 | 2056361 | BCG_1846c | BCG_1846c | 1203 | 2035150 | 2036352 | BCGDan_1852 | BCGDan_1852 | 1203 | 2035411 | 2036604 | BCGDan_sapMKO_1852 | BCGDan_sapMKO_1852 | 1194 |
| 2055681 | 2056112 | Rv1813c | Rv1813c | 432 | 2056683 | 2057114 | BCG_1847c | BCG_1847c | 432 | 2036674 | 2037105 | BCGDan_1853 | BCGDan_1853 | 432 | 2036935 | 2037366 | BCGDan_sapMKO_1853 | BCGDan_sapMKO_1853 | 432 |
| 2056521 | 2057423 | erg3 | Rv1814 | 903 | 2057523 | 2058425 | erg3 | BCG_1848 | 903 | 2037514 | 2038416 | erg3 | BCGDan_1854 | 903 | 2037733 | 2038677 | erg3 | BCGDan_sapMKO_1854 | 945 |
| 2057528 | 2058193 | Rv1815 | Rv1815 | 666 | 2058530 | 2059195 | BCG_1849 | BCG_1849 | 666 | 2038521 | 2039186 | BCGDan_1855 | BCGDan_1855 | 666 | 2038782 | 2039447 | BCGDan_sapMKO_1855 | BCGDan_sapMKO_1855 | 666 |
| 2058256 | 2058960 | Rv1816 | Rv1816 | 705 | 2059258 | 2059962 | BCG_1850 | BCG_1850 | 705 | 2039249 | 2039953 | BCGDan_1856 | BCGDan_1856 | 705 | 2039528 | 2040214 | BCGDan_sapMKO_1856 | BCGDan_sapMKO_1856 | 687 |
| 2059595 | 2061058 | Rv1817 | Rv1817 | 1464 | 2060597 | 2062060 | BCG_1852 | BCG_1852 | 1464 | 2040588 | 2042051 | BCGDan_1857 | BCGDan_1857 | 1464 | 2040849 | 2042312 | BCGDan_sapMKO_1859 | BCGDan_sapMKO_1859 | 1464 |
| 2061178 | 2062674 | PE_PGRS33 | Rv1818c | 1497 | 2062180 | 2063685 | PE_PGRS33 | BCG_1853c | 1506 | 2042171 | 2043676 | PE_PGRS33 | BCGDan_1858 | 1506 | 2042432 | 2043937 | PE_PGRS33 | BCGDan_sapMKO_1860 | 1506 |
| 2062809 | 2064728 | bacA | Rv1819c | 1920 | 2063820 | 2065739 | BCG_1854c | BCG_1854c | 1920 | 2043811 | 2045730 | BCGDan_1859 | BCGDan_1859 | 1920 | 2044072 | 2045934 | BCGDan_sapMKO_1861 | BCGDan_sapMKO_1861 | 1863 |
| 2064799 | 2066442 | ilvG | Rv1820 | 1644 | 2065810 | 2067453 | ilvG | BCG_1855 | 1644 | 2045801 | 2047444 | ilvG | BCGDan_1860 | 1644 | 2046062 | 2047705 | ilvG | BCGDan_sapMKO_1862 | 1644 |
| 2066457 | 2068883 | secA2 | Rv1821 | 2427 | 2067468 | 2069894 | secA2 | BCG_1856 | 2427 | 2047459 | 2049885 | secA2 | BCGDan_1861 | 2427 | 2047810 | 2050146 | secA2 | BCGDan_sapMKO_1863 | 2337 |
| 2069080 | 2069709 | pgsA2 | Rv1822 | 630 | 2070091 | 2070720 | pgsA2 | BCG_1857 | 630 | 2050082 | 2050711 | pgsA2 | BCGDan_1862 | 630 | 2050343 | 2050972 | pgsA2 | BCGDan_sapMKO_1864 | 630 |
| 2069702 | 2070625 | Rv1823 | Rv1823 | 924 | 2070713 | 2071636 | BCG_1858 | BCG_1858 | 924 | 2050704 | 2051627 | BCGDan_1863 | BCGDan_1863 | 924 | 2050959 | 2051888 | BCGDan_sapMKO_1865 | BCGDan_sapMKO_1865 | 930 |
| 2070654 | 2071019 | Rv1824 | Rv1824 | 366 | 2071665 | 2072030 | BCG_1859 | BCG_1859 | 366 | 2051656 | 2052021 | BCGDan_1864 | BCGDan_1864 | 366 | 2051950 | 2052282 | BCGDan_sapMKO_1866 | BCGDan_sapMKO_1866 | 333 |
| 2071036 | 2071914 | Rv1825 | Rv1825 | 879 | 2072047 | 2072925 | BCG_1860 | BCG_1860 | 879 | 2052038 | 2052916 | BCGDan_1865 | BCGDan_1865 | 879 | 2052299 | 2053177 | BCGDan_sapMKO_1867 | BCGDan_sapMKO_1867 | 879 |
| 2071952 | 2072356 | gcvH | Rv1826 | 405 | 2072963 | 2073367 | gcvH | BCG_1861 | 405 | 2052954 | 2053358 | gcvH | BCGDan_1866 | 405 | 2053215 | 2053619 | gcvH | BCGDan_sapMKO_1868 | 405 |
| 2072596 | 2073084 | garA | Rv1827 | 489 | 2073607 | 2074095 | cfp17 | BCG_1862 | 489 | 2053598 | 2054086 | cfp17 | BCGDan_1867 | 489 | 2053868 | 2054347 | cfp17 | BCGDan_sapMKO_1869 | 480 |
| 2073081 | 2073824 | Rv1828 | Rv1828 | 744 | 2074092 | 2074835 | BCG_1863 | BCG_1863 | 744 | 2054083 | 2054826 | BCGDan_1868 | BCGDan_1868 | 744 | 2054344 | 2055087 | BCGDan_sapMKO_1870 | BCGDan_sapMKO_1870 | 744 |
| 2073943 | 2074437 | Rv1829 | Rv1829 | 495 | 2074954 | 2075448 | BCG_1864 | BCG_1864 | 495 | 2054945 | 2055439 | BCGDan_1869 | BCGDan_1869 | 495 | 2055206 | 2055700 | BCGDan_sapMKO_1871 | BCGDan_sapMKO_1871 | 495 |
| 2074841 | 2075518 | Rv1830 | Rv1830 | 678 | 2075796 | 2076473 | BCG_1865 | BCG_1865 | 678 | 2055787 | 2056464 | BCGDan_1870 | BCGDan_1870 | 678 | 2056093 | 2056725 | BCGDan_sapMKO_1872 | BCGDan_sapMKO_1872 | 633 |
| 2075571 | 2075828 | Rv1831 | Rv1831 | 258 | 2076526 | 2076783 | BCG_1866 | BCG_1866 | 258 | 2056517 | 2056774 | BCGDan_1871 | BCGDan_1871 | 258 | 2056778 | 2057035 | BCGDan_sapMKO_1873 | BCGDan_sapMKO_1873 | 258 |
| 2075877 | 2078702 | gcvB | Rv1832 | 2826 | 2076832 | 2079657 | gcvB | BCG_1867 | 2826 | 2056823 | 2059648 | gcvB | BCGDan_1872 | 2826 | 2057084 | 2059909 | gcvB | BCGDan_sapMKO_1874 | 2826 |
| 2078929 | 2079789 | Rv1833c | Rv1833c | 861 | 2079884 | 2080744 | BCG_1868c | BCG_1868c | 861 | 2059875 | 2060735 | BCGDan_1873 | BCGDan_1873 | 861 | 2060136 | 2060996 | BCGDan_sapMKO_1875 | BCGDan_sapMKO_1875 | 861 |
| 2079830 | 2080696 | lipZ | Rv1834 | 867 | 2080785 | 2081651 | BCG_1869 | BCG_1869 | 867 | 2060776 | 2061642 | BCGDan_1874 | BCGDan_1874 | 867 | 2061139 | 2061903 | BCGDan_sapMKO_1876 | BCGDan_sapMKO_1876 | 765 |
| 2080701 | 2082587 | Rv1835c | Rv1835c | 1887 | 2081656 | 2083542 | BCG_1870c | BCG_1870c | 1887 | 2061647 | 2063533 | BCGDan_1875 | BCGDan_1875 | 1887 | 2061908 | 2063716 | BCGDan_sapMKO_1877 | BCGDan_sapMKO_1877 | 1809 |
| 2082603 | 2084636 | Rv1836c | Rv1836c | 2034 | 2083558 | 2085591 | BCG_1871c | BCG_1871c | 2034 | 2063549 | 2065582 | BCGDan_1876 | BCGDan_1876 | 2034 | 2063810 | 2065843 | BCGDan_sapMKO_1878 | BCGDan_sapMKO_1878 | 2034 |
| 2084756 | 2086981 | glcB | Rv1837c | 2226 | 2085711 | 2087936 | glcB | BCG_1872c | 2226 | 2065702 | 2067927 | glcB | BCGDan_1877 | 2226 | 2065963 | 2068188 | glcB | BCGDan_sapMKO_1879 | 2226 |
| 2087257 | 2087652 | vapC13 | Rv1838c | 396 | 2088212 | 2088607 | BCG_1873c | BCG_1873c | 396 | 2068203 | 2068598 | BCGDan_1878 | BCGDan_1878 | 396 | 2068464 | 2068832 | BCGDan_sapMKO_1880 | BCGDan_sapMKO_1880 | 369 |
| 2087649 | 2087912 | vapB13 | Rv1839c | 264 | 2088604 | 2088867 | BCG_1874c | BCG_1874c | 264 | 2068595 | 2068858 | BCGDan_1879 | BCGDan_1879 | 264 | 2068856 | 2069119 | BCGDan_sapMKO_1881 | BCGDan_sapMKO_1881 | 264 |
| NA | NA | NA | NA | NA | 2088926 | 2089972 | PE_PGRS34b | BCG_1875c | 1047 | 2068917 | 2069963 | PE_PGRS34b | BCGDan_1880 | 1047 | 2069178 | 2069747 | BCGDan_sapMKO_1882 | BCGDan_sapMKO_1882 | 570 |
| 2087971 | 2089518 | PE_PGRS34 | Rv1840c | 1548 | 2089654 | 2090472 | PE_PGRS34a | BCG_1876c | 819 | 2069645 | 2070463 | PE_PGRS34a | BCGDan_1881 | 819 | 2069906 | 2070724 | PE_PGRS34a | BCGDan_sapMKO_1883 | 819 |
| 2089681 | 2090718 | Rv1841c | Rv1841c | 1038 | 2090635 | 2091684 | BCG_1877c | BCG_1877c | 1050 | 2070626 | 2071675 | BCGDan_1882 | BCGDan_1882 | 1050 | 2070887 | 2071936 | BCGDan_sapMKO_1884 | BCGDan_sapMKO_1884 | 1050 |
| 2090718 | 2092085 | Rv1842c | Rv1842c | 1368 | 2091684 | 2093051 | BCG_1878c | BCG_1878c | 1368 | 2071675 | 2073042 | BCGDan_1883 | BCGDan_1883 | 1368 | 2071936 | 2073303 | BCGDan_sapMKO_1885 | BCGDan_sapMKO_1885 | 1368 |
| 2092259 | 2093698 | guaB1 | Rv1843c | 1440 | 2093225 | 2094664 | guaB1 | BCG_1879c | 1440 | 2073216 | 2074655 | guaB1 | BCGDan_1884 | 1440 | 2073477 | 2074862 | guaB1 | BCGDan_sapMKO_1886 | 1386 |
| 2093731 | 2095188 | gnd1 | Rv1844c | 1458 | 2094030 | 2096087 | gnd1 | BCG_1880c | 2058 | 2074021 | 2076078 | gnd1 | BCGDan_1885 | 2058 | 2074282 | 2076339 | gnd1 | BCGDan_sapMKO_1887 | 2058 |
| 2095218 | 2096168 | blaR | Rv1845c | 951 | 2096177 | 2097127 | BCG_1881c | BCG_1881c | 951 | 2076168 | 2077118 | BCGDan_1886 | BCGDan_1886 | 951 | 2076429 | 2077379 | BCGDan_sapMKO_1888 | BCGDan_sapMKO_1888 | 951 |
| 2096183 | 2096599 | blaI | Rv1846c | 417 | 2097142 | 2097558 | BCG_1882c | BCG_1882c | 417 | 2077133 | 2077549 | BCGDan_1887 | BCGDan_1887 | 417 | 2077394 | 2077810 | BCGDan_sapMKO_1889 | BCGDan_sapMKO_1889 | 417 |
| 2096877 | 2097299 | Rv1847 | Rv1847 | 423 | 2097836 | 2098258 | BCG_1883 | BCG_1883 | 423 | 2077827 | 2078249 | BCGDan_1888 | BCGDan_1888 | 423 | 2078088 | 2078510 | BCGDan_sapMKO_1890 | BCGDan_sapMKO_1890 | 423 |
| 2097348 | 2097650 | ureA | Rv1848 | 303 | 2098307 | 2098609 | ureA | BCG_1884 | 303 | 2078298 | 2078600 | ureA | BCGDan_1889 | 303 | 2078559 | 2078861 | ureA | BCGDan_sapMKO_1891 | 303 |
| 2097647 | 2097961 | ureB | Rv1849 | 315 | 2098606 | 2098920 | ureB | BCG_1885 | 315 | 2078597 | 2078911 | ureB | BCGDan_1890 | 315 | 2078858 | 2079172 | ureB | BCGDan_sapMKO_1892 | 315 |
| 2097961 | 2099694 | ureC | Rv1850 | 1734 | 2098920 | 2100653 | ureC | BCG_1886 | 1734 | 2078911 | 2080644 | ureC | BCGDan_1891 | 1734 | 2079172 | 2080905 | ureC | BCGDan_sapMKO_1893 | 1734 |
| 2099694 | 2100329 | ureF | Rv1851 | 636 | 2100653 | 2101288 | ureF | BCG_1887 | 636 | 2080644 | 2081279 | ureF | BCGDan_1892 | 636 | 2080905 | 2081540 | ureF | BCGDan_sapMKO_1894 | 636 |
| 2100340 | 2101014 | ureG | Rv1852 | 675 | 2101299 | 2101973 | ureG | BCG_1888 | 675 | 2081290 | 2081964 | ureG | BCGDan_1893 | 675 | 2081551 | 2082225 | ureG | BCGDan_sapMKO_1895 | 675 |
| 2101022 | 2101648 | ureD | Rv1853 | 627 | 2101981 | 2102607 | ureD | BCG_1889 | 627 | 2081972 | 2082598 | ureD | BCGDan_1894 | 627 | 2082233 | 2082859 | ureD | BCGDan_sapMKO_1896 | 627 |
| 2101651 | 2103042 | ndh | Rv1854c | 1392 | 2102610 | 2104001 | ndh | BCG_1890c | 1392 | 2082601 | 2083992 | ndh | BCGDan_1895 | 1392 | 2082862 | 2084202 | ndh | BCGDan_sapMKO_1897 | 1341 |
| 2103184 | 2104107 | Rv1855c | Rv1855c | 924 | 2104142 | 2105065 | BCG_1891c | BCG_1891c | 924 | 2084133 | 2085056 | BCGDan_1896 | BCGDan_1896 | 924 | 2084394 | 2085317 | BCGDan_sapMKO_1898 | BCGDan_sapMKO_1898 | 924 |
| 2104146 | 2104823 | Rv1856c | Rv1856c | 678 | 2105104 | 2105781 | BCG_1892c | BCG_1892c | 678 | 2085095 | 2085772 | BCGDan_1897 | BCGDan_1897 | 678 | 2085356 | 2086033 | BCGDan_sapMKO_1899 | BCGDan_sapMKO_1899 | 678 |
| 2104985 | 2105770 | modA | Rv1857 | 786 | 2105943 | 2106728 | modA | BCG_1893 | 786 | 2085934 | 2086719 | modA | BCGDan_1898 | 786 | 2086234 | 2086980 | modA | BCGDan_sapMKO_1900 | 747 |
| 2105773 | 2106567 | modB | Rv1858 | 795 | 2106731 | 2107525 | modB | BCG_1894 | 795 | 2086722 | 2087516 | modB | BCGDan_1899 | 795 | 2086983 | 2087777 | modB | BCGDan_sapMKO_1901 | 795 |
| 2106574 | 2107683 | modC | Rv1859 | 1110 | 2107532 | 2108641 | modC | BCG_1895 | 1110 | 2087523 | 2088632 | modC | BCGDan_1900 | 1110 | 2087784 | 2088893 | modC | BCGDan_sapMKO_1902 | 1110 |
| 2107736 | 2108713 | apa | Rv1860 | 978 | 2108694 | 2109671 | apa | BCG_1896 | 978 | 2088685 | 2089662 | apa | BCGDan_1901 | 978 | 2088946 | 2089923 | apa | BCGDan_sapMKO_1903 | 978 |
| 2109165 | 2109470 | Rv1861 | Rv1861 | 306 | 2110123 | 2110428 | BCG_1897 | BCG_1897 | 306 | 2090114 | 2090419 | BCGDan_1902 | BCGDan_1902 | 306 | 2090375 | 2090680 | BCGDan_sapMKO_1904 | BCGDan_sapMKO_1904 | 306 |
| 2109544 | 2110584 | adhA | Rv1862 | 1041 | 2110503 | 2111543 | adhA | BCG_1898 | 1041 | 2090494 | 2091534 | adhA | BCGDan_1903 | 1041 | 2090755 | 2091795 | adhA | BCGDan_sapMKO_1905 | 1041 |
| 2110591 | 2111361 | Rv1863c | Rv1863c | 771 | 2111550 | 2112320 | BCG_1899c | BCG_1899c | 771 | 2091541 | 2092311 | BCGDan_1904 | BCGDan_1904 | 771 | 2091802 | 2092524 | BCGDan_sapMKO_1906 | BCGDan_sapMKO_1906 | 723 |
| 2111354 | 2112109 | Rv1864c | Rv1864c | 756 | 2112313 | 2113068 | BCG_1900c | BCG_1900c | 756 | 2092304 | 2093059 | BCGDan_1905 | BCGDan_1905 | 756 | 2092565 | 2093266 | BCGDan_sapMKO_1907 | BCGDan_sapMKO_1907 | 702 |
| 2112106 | 2112966 | Rv1865c | Rv1865c | 861 | 2113065 | 2113925 | BCG_1901c | BCG_1901c | 861 | 2093056 | 2093916 | BCGDan_1906 | BCGDan_1906 | 861 | 2093317 | 2094177 | BCGDan_sapMKO_1908 | BCGDan_sapMKO_1908 | 861 |
| 2113140 | 2115476 | Rv1866 | Rv1866 | 2337 | 2114099 | 2116435 | BCG_1902 | BCG_1902 | 2337 | 2094090 | 2096426 | BCGDan_1907 | BCGDan_1907 | 2337 | 2094273 | 2096687 | BCGDan_sapMKO_1909 | BCGDan_sapMKO_1909 | 2415 |
| 2115764 | 2117248 | Rv1867 | Rv1867 | 1485 | 2116723 | 2118207 | BCG_1903 | BCG_1903 | 1485 | 2096714 | 2098198 | BCGDan_1908 | BCGDan_1908 | 1485 | 2096975 | 2098459 | BCGDan_sapMKO_1910 | BCGDan_sapMKO_1910 | 1485 |
| 2117347 | 2119446 | Rv1868 | Rv1868 | 2100 | 2118306 | 2120405 | BCG_1904 | BCG_1904 | 2100 | 2098297 | 2100396 | BCGDan_1909 | BCGDan_1909 | 2100 | 2098558 | 2100657 | BCGDan_sapMKO_1911 | BCGDan_sapMKO_1911 | 2100 |
| 2119460 | 2120695 | Rv1869c | Rv1869c | 1236 | 2120419 | 2121654 | BCG_1905c | BCG_1905c | 1236 | 2100410 | 2101645 | BCGDan_1910 | BCGDan_1910 | 1236 | 2100671 | 2101906 | BCGDan_sapMKO_1912 | BCGDan_sapMKO_1912 | 1236 |
| 2120795 | 2121430 | Rv1870c | Rv1870c | 636 | 2121721 | 2122389 | BCG_1906c | BCG_1906c | 669 | 2101712 | 2102380 | BCGDan_1911 | BCGDan_1911 | 669 | 2101973 | 2102620 | BCGDan_sapMKO_1913 | BCGDan_sapMKO_1913 | 648 |

| | | | | | | | | | | | | | | | | | | | |
|---|---|---|---|---|---|---|---|---|---|---|---|---|---|---|---|---|---|---|---|
| 2121495 | 2121884 | Rv1871c | Rv1871c | 390 | 2122454 | 2122843 | BCG_1907c | BCG_1907c | 390 | 2102445 | 2102834 | BCGDan_1912 | BCGDan_1912 | 390 | 2102706 | 2103083 | BCGDan_sapMKO_1914 | BCGDan_sapMKO_1914 | 378 |
| 2121907 | 2123151 | lldD2 | Rv1872c | 1245 | 2122866 | 2124110 | lldD2 | BCG_1908c | 1245 | 2102857 | 2104101 | lldD2 | BCGDan_1913 | 1245 | 2103118 | 2104362 | lldD2 | BCGDan_sapMKO_1915 | 1245 |
| 2123174 | 2123611 | Rv1873 | Rv1873 | 438 | 2124133 | 2124570 | BCG_1909 | BCG_1909 | 438 | 2104124 | 2104561 | BCGDan_1914 | BCGDan_1914 | 438 | 2104385 | 2104822 | BCGDan_sapMKO_1916 | BCGDan_sapMKO_1916 | 438 |
| 2123684 | 2124370 | Rv1874 | Rv1874 | 687 | 2124643 | 2125329 | BCG_1910 | BCG_1910 | 687 | 2104634 | 2105320 | BCGDan_1915 | BCGDan_1915 | 687 | 2104877 | 2105581 | BCGDan_sapMKO_1917 | BCGDan_sapMKO_1917 | 705 |
| 2124381 | 2124824 | Rv1875 | Rv1875 | 444 | 2125340 | 2125783 | BCG_1911 | BCG_1911 | 444 | 2105331 | 2105774 | BCGDan_1916 | BCGDan_1916 | 444 | 2105592 | 2106035 | BCGDan_sapMKO_1918 | BCGDan_sapMKO_1918 | 444 |
| 2125340 | 2125819 | bfrA | Rv1876 | 480 | 2126299 | 2126778 | bfrA | BCG_1912 | 480 | 2106290 | 2106769 | bfrA | BCGDan_1917 | 480 | 2106545 | 2107030 | bfrA | BCGDan_sapMKO_1919 | 486 |
| 2125904 | 2127967 | Rv1877 | Rv1877 | 2064 | 2126863 | 2128398 | BCG_1913 | BCG_1913 | 1536 | 2106854 | 2108389 | BCGDan_1918 | BCGDan_1918 | 1536 | 2107115 | 2108650 | BCGDan_sapMKO_1920 | BCGDan_sapMKO_1920 | 1536 |
| NA | NA | NA | NA | NA | 2127713 | 2128927 | BCG_1914 | BCG_1914 | 1215 | 2107704 | 2108918 | BCGDan_1919 | BCGDan_1919 | 1215 | 2107965 | 2109179 | BCGDan_sapMKO_1921 | BCGDan_sapMKO_1921 | 1215 |
| 2128022 | 2129374 | glnA3 | Rv1878 | 1353 | 2128982 | 2130334 | glnA3 | BCG_1915 | 1353 | 2108973 | 2110325 | glnA3 | BCGDan_1920 | 1353 | 2109234 | 2110586 | glnA3 | BCGDan_sapMKO_1922 | 1353 |
| 2129377 | 2130513 | Rv1879 | Rv1879 | 1137 | 2130337 | 2131473 | BCG_1916 | BCG_1916 | 1137 | 2110328 | 2111464 | BCGDan_1921 | BCGDan_1921 | 1137 | 2110589 | 2111725 | BCGDan_sapMKO_1923 | BCGDan_sapMKO_1923 | 1137 |
| 2130541 | 2131857 | cyp140 | Rv1880c | 1317 | 2131501 | 2132817 | cyp140 | BCG_1917c | 1317 | 2111492 | 2112808 | cyp140 | BCGDan_1922 | 1317 | 2111753 | 2113117 | cyp140 | BCGDan_sapMKO_1924 | 1365 |
| 2131907 | 2132329 | lppE | Rv1881c | 423 | 2132867 | 2133289 | lppE | BCG_1918c | 423 | 2112858 | 2113280 | lppE | BCGDan_1923 | 423 | 2113119 | 2113505 | lppE | BCGDan_sapMKO_1925 | 387 |
| 2132370 | 2133203 | Rv1882c | Rv1882c | 834 | 2133330 | 2134163 | BCG_1919c | BCG_1919c | 834 | 2113321 | 2114154 | BCGDan_1924 | BCGDan_1924 | 834 | 2113582 | 2114415 | BCGDan_sapMKO_1926 | BCGDan_sapMKO_1926 | 834 |
| 2133231 | 2133692 | Rv1883c | Rv1883c | 462 | 2134191 | 2134667 | BCG_1920c | BCG_1920c | 477 | 2114182 | 2114658 | BCGDan_1925 | BCGDan_1925 | 477 | 2114443 | 2114919 | BCGDan_sapMKO_1927 | BCGDan_sapMKO_1927 | 477 |
| 2133731 | 2134261 | rpfC | Rv1884c | 531 | 2134706 | 2135236 | rpfC | BCG_1921c | 531 | 2114697 | 2115227 | rpfC | BCGDan_1926 | 531 | 2114958 | 2115488 | rpfC | BCGDan_sapMKO_1928 | 531 |
| 2134273 | 2134872 | Rv1885c | Rv1885c | 600 | 2135248 | 2135847 | BCG_1922c | BCG_1922c | 600 | 2115239 | 2115838 | BCGDan_1927 | BCGDan_1927 | 600 | 2115500 | 2116099 | BCGDan_sapMKO_1929 | BCGDan_sapMKO_1929 | 600 |
| 2134890 | 2135867 | fbpB | Rv1886c | 978 | 2135865 | 2136842 | fbpB | BCG_1923c | 978 | 2115856 | 2116833 | fbpB | BCGDan_1928 | 978 | 2116117 | 2117049 | fbpB | BCGDan_sapMKO_1930 | 933 |
| 2136258 | 2137400 | Rv1887 | Rv1887 | 1143 | 2137233 | 2138375 | BCG_1924 | BCG_1924 | 1143 | 2117224 | 2118366 | BCGDan_1929 | BCGDan_1929 | 1143 | 2117569 | 2118627 | BCGDan_sapMKO_1931 | BCGDan_sapMKO_1931 | 1059 |
| 2137519 | 2138079 | Rv1888c | Rv1888c | 561 | 2138345 | 2139064 | BCG_1925c | BCG_1925c | 720 | 2118336 | 2119055 | BCGDan_1930 | BCGDan_1930 | 720 | 2118597 | 2119316 | BCGDan_sapMKO_1932 | BCGDan_sapMKO_1932 | 720 |
| 2138444 | 2138617 | Rv1888A | Rv1888A | 174 | 2139429 | 2139602 | BCG_1926c | BCG_1926c | 174 | 2119420 | 2119593 | BCGDan_1931 | BCGDan_1931 | 174 | 2119681 | 2119854 | BCGDan_sapMKO_1933 | BCGDan_sapMKO_1933 | 174 |
| 2138661 | 2139017 | Rv1889c | Rv1889c | 357 | 2139646 | 2140002 | BCG_1927c | BCG_1927c | 357 | 2119637 | 2119993 | BCGDan_1932 | BCGDan_1932 | 357 | 2119898 | 2120254 | BCGDan_sapMKO_1934 | BCGDan_sapMKO_1934 | 357 |
| 2139076 | 2139687 | Rv1890c | Rv1890c | 612 | 2140061 | 2140672 | BCG_1928c | BCG_1928c | 612 | 2120052 | 2120663 | BCGDan_1933 | BCGDan_1933 | 612 | 2120313 | 2120864 | BCGDan_sapMKO_1935 | BCGDan_sapMKO_1935 | 552 |
| 2139741 | 2140148 | Rv1891 | Rv1891 | 408 | 2140726 | 2141133 | BCG_1929 | BCG_1929 | 408 | 2120717 | 2121124 | BCGDan_1934 | BCGDan_1934 | 408 | 2120957 | 2121385 | BCGDan_sapMKO_1936 | BCGDan_sapMKO_1936 | 429 |
| 2140165 | 2140476 | Rv1892 | Rv1892 | 312 | 2141150 | 2141461 | BCG_1930 | BCG_1930 | 312 | 2121141 | 2121452 | BCGDan_1935 | BCGDan_1935 | 312 | 2121402 | 2121713 | BCGDan_sapMKO_1937 | BCGDan_sapMKO_1937 | 312 |
| 2140486 | 2140704 | Rv1893 | Rv1893 | 219 | 2141471 | 2141689 | BCG_1931 | BCG_1931 | 219 | 2121462 | 2121680 | BCGDan_1936 | BCGDan_1936 | 219 | 2121723 | 2121941 | BCGDan_sapMKO_1938 | BCGDan_sapMKO_1938 | 219 |
| 2140739 | 2141869 | Rv1894c | Rv1894c | 1131 | 2141724 | 2142854 | BCG_1932c | BCG_1932c | 1131 | 2121715 | 2122845 | BCGDan_1937 | BCGDan_1937 | 1131 | 2121976 | 2123106 | BCGDan_sapMKO_1939 | BCGDan_sapMKO_1939 | 1131 |
| 2142521 | 2143675 | Rv1895 | Rv1895 | 1155 | 2143506 | 2143829 | BCG_1933 | BCG_1933 | 324 | 2123497 | 2123820 | BCGDan_1938 | BCGDan_1938 | 324 | 2123758 | 2124081 | BCGDan_sapMKO_1940 | BCGDan_sapMKO_1940 | 324 |
| NA | NA | NA | NA | NA | 2143826 | 2144530 | BCG_1934 | BCG_1934 | 705 | 2123817 | 2124521 | BCGDan_1939 | BCGDan_1939 | 705 | 2124078 | 2124782 | BCGDan_sapMKO_1941 | BCGDan_sapMKO_1941 | 705 |
| 2143535 | 2144446 | Rv1896c | Rv1896c | 912 | 2144520 | 2145431 | BCG_1935c | BCG_1935c | 912 | 2124511 | 2125422 | BCGDan_1940 | BCGDan_1940 | 912 | 2124772 | 2125683 | BCGDan_sapMKO_1942 | BCGDan_sapMKO_1942 | 912 |
| 2144451 | 2144882 | Rv1897c | Rv1897c | 432 | 2145436 | 2145867 | BCG_1936c | BCG_1936c | 432 | 2125427 | 2125858 | BCGDan_1941 | BCGDan_1941 | 432 | 2125688 | 2126119 | BCGDan_sapMKO_1943 | BCGDan_sapMKO_1943 | 432 |
| 2144940 | 2145248 | Rv1898 | Rv1898 | 309 | 2145925 | 2146233 | BCG_1937 | BCG_1937 | 309 | 2125922 | 2126224 | BCGDan_1942 | BCGDan_1942 | 303 | 2126183 | 2126485 | BCGDan_sapMKO_1944 | BCGDan_sapMKO_1944 | 303 |
| 2145214 | 2146245 | lppD | Rv1899c | 1032 | 2146199 | 2147275 | lppD | BCG_1938c | 1077 | 2126190 | 2127266 | lppD | BCGDan_1943 | 1077 | 2126451 | 2127527 | lppD | BCGDan_sapMKO_1945 | 1077 |
| 2146245 | 2147633 | lipJ | Rv1900c | 1389 | 2147275 | 2148663 | lipJ | BCG_1939c | 1389 | 2127266 | 2128654 | lipJ | BCGDan_1944 | 1389 | 2127527 | 2128864 | lipJ | BCGDan_sapMKO_1946 | 1338 |
| 2147662 | 2148954 | cinA | Rv1901 | 1293 | 2148692 | 2149984 | cinA | BCG_1940 | 1293 | 2128683 | 2129975 | cinA | BCGDan_1945 | 1293 | 2128944 | 2130236 | cinA | BCGDan_sapMKO_1947 | 1293 |
| 2149006 | 2150274 | nanT | Rv1902c | 1269 | 2150036 | 2151304 | nanT | BCG_1941c | 1269 | 2130027 | 2131295 | nanT | BCGDan_1946 | 1269 | 2130288 | 2131625 | nanT | BCGDan_sapMKO_1948 | 1338 |
| 2150364 | 2150768 | Rv1903 | Rv1903 | 405 | 2151394 | 2151798 | BCG_1942 | BCG_1942 | 405 | 2131385 | 2131789 | BCGDan_1947 | BCGDan_1947 | 405 | 2131646 | 2132050 | BCGDan_sapMKO_1949 | BCGDan_sapMKO_1949 | 405 |
| 2150954 | 2151385 | Rv1904 | Rv1904 | 432 | 2151984 | 2152415 | BCG_1943 | BCG_1943 | 432 | 2131975 | 2132406 | BCGDan_1948 | BCGDan_1948 | 432 | 2132236 | 2132667 | BCGDan_sapMKO_1950 | BCGDan_sapMKO_1950 | 432 |
| 2151433 | 2152395 | aao | Rv1905c | 963 | 2152463 | 2153425 | aao | BCG_1944c | 963 | 2132454 | 2133416 | aao | BCGDan_1949 | 963 | 2132715 | 2133677 | aao | BCGDan_sapMKO_1951 | 963 |
| 2152425 | 2152895 | Rv1906c | Rv1906c | 471 | 2153455 | 2153925 | BCG_1945c | BCG_1945c | 471 | 2133446 | 2133916 | BCGDan_1950 | BCGDan_1950 | 471 | 2133707 | 2134177 | BCGDan_sapMKO_1952 | BCGDan_sapMKO_1952 | 471 |
| 2153235 | 2153882 | Rv1907c | Rv1907c | 648 | 2154265 | 2154912 | BCG_1946c | BCG_1946c | 648 | 2134256 | 2134903 | BCGDan_1951 | BCGDan_1951 | 648 | 2134517 | 2135164 | BCGDan_sapMKO_1953 | BCGDan_sapMKO_1953 | 648 |
| 2153889 | 2156111 | katG | Rv1908c | 2223 | 2154919 | 2157141 | katG | BCG_1947c | 2223 | 2134910 | 2137132 | katG | BCGDan_1952 | 2223 | 2135171 | 2137318 | katG | BCGDan_sapMKO_1954 | 2148 |
| 2156149 | 2156592 | furA | Rv1909c | 444 | 2157179 | 2157631 | furA | BCG_1948c | 453 | 2137170 | 2137622 | furA | BCGDan_1953 | 453 | 2137431 | 2137874 | furA | BCGDan_sapMKO_1955 | 444 |
| 2156706 | 2157299 | Rv1910c | Rv1910c | 594 | 2157736 | 2158329 | BCG_1949c | BCG_1949c | 594 | 2137727 | 2138320 | BCGDan_1954 | BCGDan_1954 | 594 | 2137988 | 2138602 | BCGDan_sapMKO_1956 | BCGDan_sapMKO_1956 | 615 |
| 2157382 | 2157987 | lppC | Rv1911c | 606 | 2158412 | 2159017 | lppC | BCG_1950c | 606 | 2138403 | 2139008 | lppC | BCGDan_1955 | 606 | 2138664 | 2139269 | lppC | BCGDan_sapMKO_1957 | 606 |
| 2158087 | 2159091 | fadB5 | Rv1912c | 1005 | 2159117 | 2160121 | fadB5 | BCG_1951c | 1005 | 2139108 | 2140112 | fadB5 | BCGDan_1956 | 1005 | 2139369 | 2140373 | fadB5 | BCGDan_sapMKO_1958 | 1005 |
| 2159191 | 2159943 | Rv1913 | Rv1913 | 753 | 2160221 | 2160973 | BCG_1952 | BCG_1952 | 753 | 2140212 | 2140964 | BCGDan_1957 | BCGDan_1957 | 753 | 2140473 | 2141225 | BCGDan_sapMKO_1959 | BCGDan_sapMKO_1959 | 753 |
| 2159921 | 2160328 | Rv1914c | Rv1914c | 408 | 2160951 | 2161358 | BCG_1953c | BCG_1953c | 408 | 2140942 | 2141349 | BCGDan_1958 | BCGDan_1958 | 408 | 2141203 | 2141610 | BCGDan_sapMKO_1960 | BCGDan_sapMKO_1960 | 408 |
| 2160463 | 2161566 | aceAa | Rv1915 | 1104 | 2161493 | 2163793 | aceA | BCG_1954 | 2301 | 2141484 | 2143784 | aceA | BCGDan_1959 | 2301 | 2141745 | 2144045 | aceA | BCGDan_sapMKO_1961 | 2301 |
| 2162932 | 2167311 | PPE34 | Rv1917c | 4380 | 2163963 | 2169470 | PPE34 | BCG_1955c | 5508 | 2143954 | 2149461 | PPE34 | BCGDan_1960 | 5508 | 2144215 | 2149722 | PPE34 | BCGDan_sapMKO_1962 | 5508 |
| NA | NA | NA | NA | NA | 2169808 | 2170812 | PPE35b | BCG_1956c | 1005 | 2149799 | 2150803 | PPE35b | BCGDan_1961 | 1005 | 2150063 | 2151064 | PPE35b | BCGDan_sapMKO_1963 | 1002 |
| 2167649 | 2170612 | PPE35 | Rv1918c | 2964 | 2170895 | 2172772 | PPE35a | BCG_1957c | 1878 | 2150886 | 2152763 | PPE35a | BCGDan_1962 | 1878 | 2151147 | 2153024 | PPE35a | BCGDan_sapMKO_1964 | 1878 |
| 2171061 | 2171525 | Rv1919c | Rv1919c | 465 | 2173221 | 2173685 | BCG_1958c | BCG_1958c | 465 | 2153212 | 2153676 | BCGDan_1963 | BCGDan_1963 | 465 | 2153473 | 2153937 | BCGDan_sapMKO_1965 | BCGDan_sapMKO_1965 | 465 |
| 2171623 | 2172486 | Rv1920 | Rv1920 | 864 | 2173783 | 2174646 | BCG_1959 | BCG_1959 | 864 | 2153774 | 2154637 | BCGDan_1964 | BCGDan_1964 | 864 | 2154098 | 2154898 | BCGDan_sapMKO_1966 | BCGDan_sapMKO_1966 | 801 |
| 2172524 | 2173795 | lppF | Rv1921c | 1272 | 2174684 | 2175955 | lppF | BCG_1960c | 1272 | 2154675 | 2155946 | lppF | BCGDan_1965 | 1272 | 2154936 | 2156207 | lppF | BCGDan_sapMKO_1967 | 1272 |
| 2174067 | 2175182 | Rv1922 | Rv1922 | 1116 | 2176227 | 2177342 | BCG_1961 | BCG_1961 | 1116 | 2156218 | 2157333 | BCGDan_1966 | BCGDan_1966 | 1116 | 2156512 | 2157594 | BCGDan_sapMKO_1968 | BCGDan_sapMKO_1968 | 1083 |
| 2175173 | 2176513 | lipD | Rv1923 | 1341 | 2177333 | 2178673 | lipD | BCG_1962 | 1341 | 2157324 | 2158664 | lipD | BCGDan_1967 | 1341 | 2157645 | 2158925 | lipD | BCGDan_sapMKO_1969 | 1281 |
| 2176550 | 2176930 | Rv1924c | Rv1924c | 381 | 2178710 | 2179090 | BCG_1963c | BCG_1963c | 381 | 2158701 | 2159081 | BCGDan_1968 | BCGDan_1968 | 381 | 2158962 | 2159342 | BCGDan_sapMKO_1970 | BCGDan_sapMKO_1970 | 381 |
| 2177087 | 2178949 | fadD31 | Rv1925 | 1863 | 2179247 | 2181109 | fadD31 | BCG_1964 | 1863 | 2159238 | 2161100 | fadD31 | BCGDan_1969 | 1863 | 2159499 | 2161361 | fadD31 | BCGDan_sapMKO_1971 | 1863 |
| 2178957 | 2179436 | mpt63 | Rv1926c | 480 | 2181117 | 2181596 | mpt63 | BCG_1965c | 480 | 2161108 | 2161587 | mpt63 | BCGDan_1970 | 480 | 2161369 | 2161848 | mpt63 | BCGDan_sapMKO_1972 | 480 |
| 2179673 | 2180446 | Rv1927 | Rv1927 | 774 | 2181833 | 2182606 | BCG_1966 | BCG_1966 | 774 | 2161824 | 2162597 | BCGDan_1971 | BCGDan_1971 | 774 | 2162082 | 2162858 | BCGDan_sapMKO_1973 | BCGDan_sapMKO_1973 | 777 |
| 2180450 | 2181217 | Rv1928c | Rv1928c | 768 | 2182610 | 2183377 | BCG_1967c | BCG_1967c | 768 | 2162601 | 2163368 | BCGDan_1972 | BCGDan_1972 | 768 | 2162862 | 2163629 | BCGDan_sapMKO_1974 | BCGDan_sapMKO_1974 | 768 |
| 2181262 | 2181906 | Rv1929c | Rv1929c | 645 | 2183422 | 2184066 | BCG_1968c | BCG_1968c | 645 | 2163413 | 2164057 | BCGDan_1973 | BCGDan_1973 | 645 | 2163674 | 2164318 | BCGDan_sapMKO_1975 | BCGDan_sapMKO_1975 | 645 |
| 2181918 | 2182442 | Rv1930c | Rv1930c | 525 | 2184078 | 2184602 | BCG_1969c | BCG_1969c | 525 | 2164069 | 2164593 | BCGDan_1974 | BCGDan_1974 | 525 | 2164330 | 2164854 | BCGDan_sapMKO_1976 | BCGDan_sapMKO_1976 | 525 |
| 2182460 | 2183239 | Rv1931c | Rv1931c | 780 | 2184620 | 2185399 | BCG_1970c | BCG_1970c | 780 | 2164611 | 2165390 | BCGDan_1975 | BCGDan_1975 | 780 | 2164872 | 2165651 | BCGDan_sapMKO_1977 | BCGDan_sapMKO_1977 | 780 |
| 2183372 | 2183869 | tpx | Rv1932 | 498 | 2185532 | 2186029 | tpx | BCG_1971 | 498 | 2165523 | 2166020 | tpx | BCGDan_1976 | 498 | 2165784 | 2166281 | tpx | BCGDan_sapMKO_1978 | 498 |
| 2183866 | 2184957 | fadE18 | Rv1933c | 1092 | 2186026 | 2187117 | fadE18 | BCG_1972c | 1092 | 2166017 | 2167108 | fadE18 | BCGDan_1977 | 1092 | 2166278 | 2167369 | fadE18 | BCGDan_sapMKO_1979 | 1092 |
| 2184959 | 2186188 | fadE17 | Rv1934c | 1230 | 2187119 | 2188348 | fadE17 | BCG_1973c | 1230 | 2167110 | 2168339 | fadE17 | BCGDan_1978 | 1230 | 2167371 | 2168600 | fadE17 | BCGDan_sapMKO_1980 | 1230 |
| 2186203 | 2187159 | echA13 | Rv1935c | 957 | 2188363 | 2189319 | echA13 | BCG_1974c | 957 | 2168354 | 2169310 | echA13 | BCGDan_1979 | 957 | 2168615 | 2169571 | echA13 | BCGDan_sapMKO_1981 | 957 |
| 2187384 | 2188493 | Rv1936 | Rv1936 | 1110 | 2189544 | 2190653 | BCG_1975 | BCG_1975 | 1110 | 2169535 | 2170644 | BCGDan_1980 | BCGDan_1980 | 1110 | 2169796 | 2170905 | BCGDan_sapMKO_1982 | BCGDan_sapMKO_1982 | 1110 |
| 2188496 | 2191015 | Rv1937 | Rv1937 | 2520 | 2190656 | 2193175 | BCG_1976 | BCG_1976 | 2520 | 2170647 | 2173166 | BCGDan_1981 | BCGDan_1981 | 2520 | 2170908 | 2173427 | BCGDan_sapMKO_1983 | BCGDan_sapMKO_1983 | 2520 |
| 2191027 | 2192097 | ephB | Rv1938 | 1071 | 2193187 | 2194257 | ephB | BCG_1977 | 1071 | 2173178 | 2174248 | ephB | BCGDan_1982 | 1071 | 2173439 | 2174509 | ephB | BCGDan_sapMKO_1984 | 1071 |
| 2192094 | 2192609 | Rv1939 | Rv1939 | 516 | 2194254 | 2194769 | BCG_1978 | BCG_1978 | 516 | 2174245 | 2174760 | BCGDan_1983 | BCGDan_1983 | 516 | 2174506 | 2175021 | BCGDan_sapMKO_1985 | BCGDan_sapMKO_1985 | 516 |
| 2192606 | 2193667 | ribA1 | Rv1940 | 1062 | 2194766 | 2195827 | ribA1 | BCG_1979 | 1062 | 2174757 | 2175818 | ribA1 | BCGDan_1984 | 1062 | 2175018 | 2176079 | ribA1 | BCGDan_sapMKO_1986 | 1062 |
| 2193664 | 2194434 | Rv1941 | Rv1941 | 771 | 2195824 | 2196594 | BCG_1980 | BCG_1980 | 771 | 2175815 | 2176585 | BCGDan_1985 | BCGDan_1985 | 771 | 2176076 | 2176846 | BCGDan_sapMKO_1987 | BCGDan_sapMKO_1987 | 771 |
| 2194644 | 2194973 | mazF5 | Rv1942c | 330 | 2196804 | 2197133 | BCG_1981c | BCG_1981c | 330 | 2176795 | 2177148 | BCGDan_1986 | BCGDan_1986 | 354 | 2177056 | 2177409 | BCGDan_sapMKO_1988 | BCGDan_sapMKO_1988 | 354 |
| 2194970 | 2195347 | mazE5 | Rv1943c | 378 | 2197130 | 2197507 | BCG_1982c | BCG_1982c | 378 | 2177121 | 2177498 | BCGDan_1987 | BCGDan_1987 | 378 | 2177382 | 2177759 | BCGDan_sapMKO_1989 | BCGDan_sapMKO_1989 | 378 |
| 2195344 | 2195934 | Rv1944c | Rv1944c | 591 | 2197504 | 2198094 | BCG_1983c | BCG_1983c | 591 | 2177495 | 2178085 | BCGDan_1988 | BCGDan_1988 | 591 | 2177756 | 2178346 | BCGDan_sapMKO_1990 | BCGDan_sapMKO_1990 | 591 |
| 2195989 | 2197353 | Rv1945 | Rv1945 | 1365 | 2198149 | 2199513 | BCG_1984 | BCG_1984 | 1365 | 2178140 | 2179504 | BCGDan_1989 | BCGDan_1989 | 1365 | 2178401 | 2179765 | BCGDan_sapMKO_1991 | BCGDan_sapMKO_1991 | 1365 |
| 2197508 | 2197960 | lppG | Rv1946c | 453 | 2199668 | 2200120 | lppG | BCG_1985c | 453 | 2179659 | 2180111 | lppG | BCGDan_1990 | 453 | 2179920 | 2180372 | lppG | BCGDan_sapMKO_1992 | 453 |
| 2198024 | 2198425 | Rv1947 | Rv1947 | 402 | 2200184 | 2200585 | BCG_1986 | BCG_1986 | 402 | 2180175 | 2180576 | BCGDan_1991 | BCGDan_1991 | 402 | 2180436 | 2180837 | BCGDan_sapMKO_1993 | BCGDan_sapMKO_1993 | 402 |
| 2198714 | 2199064 | Rv1948c | Rv1948c | 351 | 2200874 | 2201224 | BCG_1987c | BCG_1987c | 351 | 2180865 | 2181215 | BCGDan_1992 | BCGDan_1992 | 351 | 2181126 | 2181476 | BCGDan_sapMKO_1994 | BCGDan_sapMKO_1994 | 351 |
| 2199075 | 2200034 | Rv1949c | Rv1949c | 960 | 2201235 | 2202137 | BCG_1988c | BCG_1988c | 903 | 2181226 | 2182128 | BCGDan_1993 | BCGDan_1993 | 903 | 2181487 | 2182389 | BCGDan_sapMKO_1995 | BCGDan_sapMKO_1995 | 903 |
| 2199998 | 2200189 | Rv1950c | Rv1950c | 192 | 2202158 | 2202349 | BCG_1989c | BCG_1989c | 192 | 2182149 | 2182340 | BCGDan_1994 | BCGDan_1994 | 192 | 2182410 | 2182601 | BCGDan_sapMKO_1996 | BCGDan_sapMKO_1996 | 192 |
| 2200190 | 2200486 | Rv1951c | Rv1951c | 297 | 2202350 | 2202646 | BCG_1990c | BCG_1990c | 297 | 2182341 | 2182637 | BCGDan_1995 | BCGDan_1995 | 297 | 2182602 | 2182898 | BCGDan_sapMKO_1997 | BCGDan_sapMKO_1997 | 297 |
| 2200726 | 2200941 | vapB14 | Rv1952 | 216 | 2202886 | 2203101 | BCG_1991 | BCG_1991 | 216 | 2182877 | 2183092 | BCGDan_1996 | BCGDan_1996 | 216 | 2183126 | 2183353 | BCGDan_sapMKO_1998 | BCGDan_sapMKO_1998 | 228 |
| 2200938 | 2201249 | vapC14 | Rv1953 | 312 | 2203098 | 2203409 | BCG_1992 | BCG_1992 | 312 | 2183089 | 2183400 | BCGDan_1997 | BCGDan_1997 | 312 | 2183350 | 2183661 | BCGDan_sapMKO_1999 | BCGDan_sapMKO_1999 | 312 |

| | | | | | | | | | | | | | | | | | | | |
|---|---|---|---|---|---|---|---|---|---|---|---|---|---|---|---|---|---|---|---|
| 2201223 | 2201744 | Rv1954c | Rv1954c | 522 | 2203383 | 2203904 | BCG_1993c | BCG_1993c | 522 | 2183374 | 2183895 | BCGDan_1998 | BCGDan_1998 | 522 | 2183635 | 2184156 | BCGDan_sapMKO_2000 | BCGDan_sapMKO_2000 | 522 |
| 2201719 | 2202096 | higB | Rv1955 | 378 | 2203744 | 2204256 | BCG_1994 | BCG_1994 | 513 | 2183735 | 2184247 | BCGDan_1999 | BCGDan_1999 | 513 | 2183996 | 2184508 | BCGDan_sapMKO_2001 | BCGDan_sapMKO_2001 | 513 |
| 2202138 | 2202587 | higA | Rv1956 | 450 | 2204298 | 2204747 | BCG_1995 | BCG_1995 | 450 | 2184289 | 2184738 | BCGDan_2000 | BCGDan_2000 | 450 | 2184550 | 2184999 | BCGDan_sapMKO_2002 | BCGDan_sapMKO_2002 | 450 |
| 2202584 | 2203129 | Rv1957 | Rv1957 | 546 | 2204744 | 2205289 | BCG_1996 | BCG_1996 | 546 | 2184735 | 2185280 | BCGDan_2001 | BCGDan_2001 | 546 | 2184996 | 2185541 | BCGDan_sapMKO_2003 | BCGDan_sapMKO_2003 | 546 |
| 2203018 | 2203632 | Rv1958c | Rv1958c | 615 | 2205178 | 2205792 | BCG_1997c | BCG_1997c | 615 | 2185169 | 2185783 | BCGDan_2002 | BCGDan_2002 | 615 | 2185430 | 2186044 | BCGDan_sapMKO_2004 | BCGDan_sapMKO_2004 | 615 |
| 2203681 | 2203977 | parE1 | Rv1959c | 297 | 2205841 | 2206137 | BCG_1998c | BCG_1998c | 297 | 2185832 | 2186128 | BCGDan_2003 | BCGDan_2003 | 297 | 2186093 | 2186389 | BCGDan_sapMKO_2005 | BCGDan_sapMKO_2005 | 297 |
| 2203974 | 2204225 | parD1 | Rv1960c | 252 | 2206134 | 2206385 | BCG_1999c | BCG_1999c | 252 | 2186125 | 2186376 | BCGDan_2004 | BCGDan_2004 | 252 | 2186386 | 2186637 | BCGDan_sapMKO_2006 | BCGDan_sapMKO_2006 | 252 |
| 2204212 | 2204706 | Rv1961 | Rv1961 | 495 | 2206372 | 2206866 | BCG_2000 | BCG_2000 | 495 | 2186363 | 2186857 | BCGDan_2005 | BCGDan_2005 | 495 | 2186624 | 2187118 | BCGDan_sapMKO_2007 | BCGDan_sapMKO_2007 | 495 |
| 2204866 | 2205273 | vapC35 | Rv1962c | 408 | 2207026 | 2207433 | BCG_2001c | BCG_2001c | 408 | 2187017 | 2187424 | BCGDan_2006 | BCGDan_2006 | 408 | 2187278 | 2187685 | BCGDan_sapMKO_2008 | BCGDan_sapMKO_2008 | 408 |
| 2205277 | 2205549 | vapB35 | Rv1962A | 273 | 2207437 | 2207709 | BCG_2001A | BCG_2001A | 273 | 2187428 | 2187700 | BCGDan_2007 | BCGDan_2007 | 273 | 2187689 | 2187961 | BCGDan_sapMKO_2009 | BCGDan_sapMKO_2009 | 273 |
| 2205582 | 2206802 | mce3R | Rv1963c | 1221 | 2207742 | 2208962 | BCG_2002c | BCG_2002c | 1221 | 2187733 | 2188953 | BCGDan_2008 | BCGDan_2008 | 1221 | 2187994 | 2189157 | BCGDan_sapMKO_2010 | BCGDan_sapMKO_2010 | 1164 |
| 2207700 | 2208497 | yrbE3A | Rv1964 | 798 | 2209861 | 2210265 | yrbE3A | BCG_2003 | 405 | 2189852 | 2190256 | yrbE3A | BCGDan_2009 | 405 | 2190113 | 2190517 | yrbE3A | BCGDan_sapMKO_2011 | 405 |
| 2231680 | 2232219 | erm(37) | Rv1988 | 540 | 2210350 | 2210874 | BCG_2004 | BCG_2004 | 525 | 2190341 | 2190865 | BCGDan_2010 | BCGDan_2010 | 525 | 2190602 | 2191126 | BCGDan_sapMKO_2012 | BCGDan_sapMKO_2012 | 525 |
| 2232739 | 2233299 | Rv1989c | Rv1989c | 561 | 2211394 | 2211954 | BCG_2005c | BCG_2005c | 561 | 2191385 | 2191945 | BCGDan_2011 | BCGDan_2011 | 561 | 2191646 | 2192206 | BCGDan_sapMKO_2013 | BCGDan_sapMKO_2013 | 561 |
| 2233296 | 2233637 | Rv1990c | Rv1990c | 342 | 2211951 | 2212292 | BCG_2006c | BCG_2006c | 342 | 2191942 | 2192283 | BCGDan_2012 | BCGDan_2012 | 342 | 2192203 | 2192544 | BCGDan_sapMKO_2014 | BCGDan_sapMKO_2014 | 342 |
| 2233881 | 2234216 | Rv1990A | Rv1990A | 336 | 2212536 | 2212871 | BCG_2007c | BCG_2007c | 336 | 2192527 | 2192862 | BCGDan_2013 | BCGDan_2013 | 336 | 2192788 | 2193123 | BCGDan_sapMKO_2015 | BCGDan_sapMKO_2015 | 336 |
| 2234305 | 2234649 | mazF6 | Rv1991c | 345 | 2212960 | 2213304 | BCG_2008c | BCG_2008c | 345 | 2192951 | 2193295 | mazF6 | BCGDan_2014 | 345 | 2193212 | 2193556 | BCGDan_sapMKO_2016 | BCGDan_sapMKO_2016 | 345 |
| 2234643 | 2234891 | mazE6 | Rv1991A | 249 | NA | NA | NA | NA | NA | 2193289 | 2193537 | mazE6 | BCGDan_2015 | 249 | 2193550 | 2193798 | mazE6 | BCGDan_sapMKO_2017 | 249 |
| 2234991 | 2237306 | ctpG | Rv1992c | 2316 | 2213646 | 2215961 | ctpG | BCG_2009c | 2316 | 2193637 | 2195952 | ctpG | BCGDan_2016 | 2316 | 2193898 | 2196213 | ctpG | BCGDan_sapMKO_2018 | 2316 |
| 2237303 | 2237575 | Rv1993c | Rv1993c | 273 | 2215958 | 2216230 | BCG_2010c | BCG_2010c | 273 | 2195949 | 2196221 | BCGDan_2017 | BCGDan_2017 | 273 | 2196210 | 2196482 | BCGDan_sapMKO_2019 | BCGDan_sapMKO_2019 | 273 |
| 2237628 | 2237984 | cmtR | Rv1994c | 357 | 2216283 | 2216639 | BCG_2011c | BCG_2011c | 357 | 2196274 | 2196630 | BCGDan_2018 | BCGDan_2018 | 357 | 2196535 | 2196891 | BCGDan_sapMKO_2020 | BCGDan_sapMKO_2020 | 357 |
| 2238141 | 2238908 | Rv1995 | Rv1995 | 768 | 2216796 | 2217563 | BCG_2012 | BCG_2012 | 768 | 2196787 | 2197554 | BCGDan_2019 | BCGDan_2019 | 768 | 2197048 | 2197815 | BCGDan_sapMKO_2021 | BCGDan_sapMKO_2021 | 768 |
| 2239004 | 2239957 | Rv1996 | Rv1996 | 954 | 2217659 | 2218612 | BCG_2013 | BCG_2013 | 954 | 2197650 | 2198603 | BCGDan_2020 | BCGDan_2020 | 954 | 2197911 | 2198864 | BCGDan_sapMKO_2022 | BCGDan_sapMKO_2022 | 954 |
| 2240159 | 2242876 | ctpF | Rv1997 | 2718 | 2218814 | 2221531 | ctpF | BCG_2014 | 2718 | 2198805 | 2201522 | ctpF | BCGDan_2021 | 2718 | 2199105 | 2201783 | ctpF | BCGDan_sapMKO_2023 | 2679 |
| 2242945 | 2243721 | Rv1998c | Rv1998c | 777 | 2221600 | 2222376 | BCG_2015c | BCG_2015c | 777 | 2201591 | 2202367 | BCGDan_2022 | BCGDan_2022 | 777 | 2201852 | 2202628 | BCGDan_sapMKO_2024 | BCGDan_sapMKO_2024 | 777 |
| 2243816 | 2245138 | Rv1999c | Rv1999c | 1323 | 2222471 | 2223793 | BCG_2016c | BCG_2016c | 1323 | 2202462 | 2203784 | BCGDan_2023 | BCGDan_2023 | 1323 | 2202723 | 2204045 | BCGDan_sapMKO_2025 | BCGDan_sapMKO_2025 | 1323 |
| 2245209 | 2246822 | Rv2000 | Rv2000 | 1614 | 2223864 | 2225477 | BCG_2017 | BCG_2017 | 1614 | 2203855 | 2205468 | BCGDan_2024 | BCGDan_2024 | 1614 | 2204287 | 2205729 | BCGDan_sapMKO_2026 | BCGDan_sapMKO_2026 | 1443 |
| 2246832 | 2247584 | Rv2001 | Rv2001 | 753 | 2225487 | 2226239 | BCG_2018 | BCG_2018 | 753 | 2205478 | 2206230 | BCGDan_2025 | BCGDan_2025 | 753 | 2205739 | 2206491 | BCGDan_sapMKO_2027 | BCGDan_sapMKO_2027 | 753 |
| 2247660 | 2248442 | fabG3 | Rv2002 | 783 | 2226315 | 2227097 | fabG3 | BCG_2019 | 783 | 2206306 | 2207088 | fabG3 | BCGDan_2026 | 783 | 2206513 | 2207349 | fabG3 | BCGDan_sapMKO_2028 | 837 |
| 2248563 | 2249420 | Rv2003c | Rv2003c | 858 | 2227218 | 2228075 | BCG_2020c | BCG_2020c | 858 | 2207209 | 2208066 | BCGDan_2027 | BCGDan_2027 | 858 | 2207470 | 2208327 | BCGDan_sapMKO_2029 | BCGDan_sapMKO_2029 | 858 |
| 2249478 | 2250974 | Rv2004c | Rv2004c | 1497 | 2228133 | 2229629 | BCG_2021c | BCG_2021c | 1497 | 2208124 | 2209620 | BCGDan_2028 | BCGDan_2028 | 1497 | 2208385 | 2209881 | BCGDan_sapMKO_2030 | BCGDan_sapMKO_2030 | 1497 |
| 2250996 | 2251883 | Rv2005c | Rv2005c | 888 | 2229651 | 2230538 | BCG_2022c | BCG_2022c | 888 | 2209642 | 2210529 | BCGDan_2029 | BCGDan_2029 | 888 | 2209903 | 2210790 | BCGDan_sapMKO_2031 | BCGDan_sapMKO_2031 | 888 |
| 2252002 | 2255985 | otsB1 | Rv2006 | 3984 | 2230657 | 2234640 | otsB1 | BCG_2023 | 3984 | 2210648 | 2214631 | otsB1 | BCGDan_2030 | 3984 | 2210909 | 2214892 | otsB1 | BCGDan_sapMKO_2032 | 3984 |
| 2256084 | 2256428 | fdxA | Rv2007c | 345 | 2234739 | 2235083 | fdxA | BCG_2024c | 345 | 2214730 | 2215074 | fdxA | BCGDan_2031 | 345 | 2214991 | 2215335 | fdxA | BCGDan_sapMKO_2033 | 345 |
| 2256617 | 2257942 | Rv2008c | Rv2008c | 1326 | 2235272 | 2236597 | BCG_2025c | BCG_2025c | 1326 | 2215263 | 2216588 | BCGDan_2032 | BCGDan_2032 | 1326 | 2215524 | 2216693 | BCGDan_sapMKO_2034 | BCGDan_sapMKO_2034 | 1170 |
| 2258030 | 2258272 | vapB15 | Rv2009 | 243 | 2236685 | 2236927 | BCG_2026 | BCG_2026 | 243 | 2216676 | 2216918 | BCGDan_2033 | BCGDan_2033 | 243 | 2216952 | 2217179 | BCGDan_sapMKO_2035 | BCGDan_sapMKO_2035 | 228 |
| 2258273 | 2258671 | vapC15 | Rv2010 | 399 | 2236928 | 2237326 | BCG_2027 | BCG_2027 | 399 | 2216919 | 2217317 | BCGDan_2034 | BCGDan_2034 | 399 | 2217180 | 2217578 | BCGDan_sapMKO_2036 | BCGDan_sapMKO_2036 | 399 |
| 2258854 | 2259285 | Rv2011c | Rv2011c | 432 | 2237509 | 2237940 | BCG_2028c | BCG_2028c | 432 | 2217500 | 2217931 | BCGDan_2035 | BCGDan_2035 | 432 | 2217761 | 2218192 | BCGDan_sapMKO_2037 | BCGDan_sapMKO_2037 | 432 |
| 2259326 | 2259820 | Rv2012 | Rv2012 | 495 | 2237981 | 2238475 | BCG_2029 | BCG_2029 | 495 | 2217972 | 2218466 | BCGDan_2036 | BCGDan_2036 | 495 | 2218293 | 2218727 | BCGDan_sapMKO_2038 | BCGDan_sapMKO_2038 | 435 |
| 2260665 | 2261144 | Rv2013 | Rv2013 | 480 | 2239179 | 2239796 | BCG_2030 | BCG_2030 | 618 | 2219170 | 2219787 | BCGDan_2037 | BCGDan_2037 | 618 | 2219431 | 2220048 | BCGDan_sapMKO_2039 | BCGDan_sapMKO_2039 | 618 |
| 2261098 | 2261688 | Rv2014 | Rv2014 | 591 | 2239750 | 2240340 | BCG_2031 | BCG_2031 | 591 | 2219741 | 2220331 | BCGDan_2038 | BCGDan_2038 | 591 | 2220002 | 2220592 | BCGDan_sapMKO_2040 | BCGDan_sapMKO_2040 | 591 |
| 2261816 | 2263072 | Rv2015c | Rv2015c | 1257 | 2240468 | 2241724 | BCG_2032c | BCG_2032c | 1257 | 2220459 | 2221715 | BCGDan_2039 | BCGDan_2039 | 1257 | 2220720 | 2221976 | BCGDan_sapMKO_2041 | BCGDan_sapMKO_2041 | 1257 |
| 2263426 | 2264001 | Rv2016 | Rv2016 | 576 | 2242078 | 2242653 | BCG_2033 | BCG_2033 | 576 | 2222069 | 2222644 | BCGDan_2040 | BCGDan_2040 | 576 | 2222294 | 2222905 | BCGDan_sapMKO_2042 | BCGDan_sapMKO_2042 | 612 |
| 2263998 | 2265038 | Rv2017 | Rv2017 | 1041 | 2242650 | 2243690 | BCG_2034 | BCG_2034 | 1041 | 2222641 | 2223681 | BCGDan_2041 | BCGDan_2041 | 1041 | 2222902 | 2223942 | BCGDan_sapMKO_2043 | BCGDan_sapMKO_2043 | 1041 |
| 2265280 | 2265999 | Rv2018 | Rv2018 | 720 | 2243932 | 2244651 | BCG_2035 | BCG_2035 | 720 | 2223923 | 2224642 | BCGDan_2042 | BCGDan_2042 | 720 | 2224175 | 2224903 | BCGDan_sapMKO_2044 | BCGDan_sapMKO_2044 | 729 |
| 2265989 | 2266405 | Rv2019 | Rv2019 | 417 | 2244641 | 2245057 | BCG_2036 | BCG_2036 | 417 | 2224632 | 2225048 | BCGDan_2043 | BCGDan_2043 | 417 | 2224893 | 2225309 | BCGDan_sapMKO_2045 | BCGDan_sapMKO_2045 | 417 |
| 2266421 | 2266720 | Rv2020c | Rv2020c | 300 | 2245073 | 2245372 | BCG_2037c | BCG_2037c | 300 | 2225064 | 2225363 | BCGDan_2044 | BCGDan_2044 | 300 | 2225325 | 2225612 | BCGDan_sapMKO_2046 | BCGDan_sapMKO_2046 | 288 |
| 2266805 | 2267110 | Rv2021c | Rv2021c | 306 | 2245457 | 2245762 | BCG_2038c | BCG_2038c | 306 | 2225448 | 2225753 | BCGDan_2045 | BCGDan_2045 | 306 | 2225709 | 2225993 | BCGDan_sapMKO_2047 | BCGDan_sapMKO_2047 | 285 |
| 2267119 | 2267724 | Rv2022c | Rv2022c | 606 | 2245771 | 2246376 | BCG_2039c | BCG_2039c | 606 | 2225762 | 2226367 | BCGDan_2046 | BCGDan_2046 | 606 | 2226023 | 2226628 | BCGDan_sapMKO_2048 | BCGDan_sapMKO_2048 | 606 |
| 2267749 | 2268108 | Rv2023c | Rv2023c | 360 | 2246401 | 2246760 | BCG_2040c | BCG_2040c | 360 | 2226392 | 2226751 | BCGDan_2047 | BCGDan_2047 | 360 | 2226653 | 2227012 | BCGDan_sapMKO_2049 | BCGDan_sapMKO_2049 | 360 |
| 2268268 | 2268726 | Rv2023A | Rv2023A | 459 | 2246920 | 2247597 | BCG_2041c | BCG_2041c | 678 | 2226911 | 2227588 | BCGDan_2048 | BCGDan_2048 | 678 | 2227172 | 2227849 | BCGDan_sapMKO_2050 | BCGDan_sapMKO_2050 | 678 |
| NA | NA | NA | NA | NA | 2247992 | 2248888 | BCG_2042c | BCG_2042c | 897 | 2227983 | 2228879 | BCGDan_2049 | BCGDan_2049 | 897 | 2228244 | 2229140 | BCGDan_sapMKO_2051 | BCGDan_sapMKO_2051 | 897 |
| 2268693 | 2270240 | Rv2024c | Rv2024c | 1548 | 2249072 | 2253892 | BCG_2043c | BCG_2043c | 4821 | 2229063 | 2233883 | BCGDan_2050 | BCGDan_2050 | 4821 | 2229324 | 2234144 | BCGDan_sapMKO_2052 | BCGDan_sapMKO_2052 | 4821 |
| 2270750 | 2271748 | Rv2025c | Rv2025c | 999 | 2254402 | 2255400 | BCG_2044c | BCG_2044c | 999 | 2234393 | 2235391 | BCGDan_2051 | BCGDan_2051 | 999 | 2234654 | 2235652 | BCGDan_sapMKO_2053 | BCGDan_sapMKO_2053 | 999 |
| 2271863 | 2272747 | Rv2026c | Rv2026c | 885 | 2255515 | 2256399 | BCG_2045c | BCG_2045c | 885 | 2235506 | 2236390 | BCGDan_2052 | BCGDan_2052 | 885 | 2235767 | 2236651 | BCGDan_sapMKO_2054 | BCGDan_sapMKO_2054 | 885 |
| 2272787 | 2274508 | dosT | Rv2027c | 1722 | 2256439 | 2258160 | BCG_2046c | BCG_2046c | 1722 | 2236430 | 2238151 | DosT | BCGDan_2053 | 1722 | 2236691 | 2238367 | BCGDan_sapMKO_2055 | BCGDan_sapMKO_2055 | 1677 |
| 2274569 | 2275408 | Rv2028c | Rv2028c | 840 | 2258221 | 2259060 | BCG_2047c | BCG_2047c | 840 | 2238212 | 2239051 | BCGDan_2054 | BCGDan_2054 | 840 | 2238473 | 2239312 | BCGDan_sapMKO_2056 | BCGDan_sapMKO_2056 | 840 |
| 2275405 | 2276424 | pfkB | Rv2029c | 1020 | 2259057 | 2260076 | pfkB | BCG_2048c | 1020 | 2239048 | 2240067 | pfkB | BCGDan_2055 | 1020 | 2239309 | 2240328 | pfkB | BCGDan_sapMKO_2057 | 1020 |
| 2276441 | 2278486 | Rv2030c | Rv2030c | 2046 | 2260093 | 2262138 | BCG_2049c | BCG_2049c | 2046 | 2240084 | 2242129 | BCGDan_2056 | BCGDan_2056 | 2046 | 2240345 | 2242390 | BCGDan_sapMKO_2058 | BCGDan_sapMKO_2058 | 2046 |
| 2278498 | 2278932 | hspX | Rv2031c | 435 | 2262150 | 2262584 | hspX | BCG_2050c | 435 | 2242141 | 2242575 | hspX | BCGDan_2057 | 435 | 2242402 | 2242836 | hspX | BCGDan_sapMKO_2059 | 435 |
| 2279129 | 2280124 | acg | Rv2032 | 996 | 2262781 | 2263776 | BCG_2051 | BCG_2051 | 996 | 2242772 | 2243767 | BCGDan_2058 | BCGDan_2058 | 996 | 2243033 | 2244028 | BCGDan_sapMKO_2060 | BCGDan_sapMKO_2060 | 996 |
| 2280240 | 2281082 | Rv2033c | Rv2033c | 843 | 2263892 | 2264734 | BCG_2052c | BCG_2052c | 843 | 2243883 | 2244725 | BCGDan_2059 | BCGDan_2059 | 843 | 2244144 | 2244986 | BCGDan_sapMKO_2061 | BCGDan_sapMKO_2061 | 843 |
| 2281294 | 2281617 | Rv2034 | Rv2034 | 324 | 2264946 | 2265269 | BCG_2053 | BCG_2053 | 324 | 2244937 | 2245260 | BCGDan_2060 | BCGDan_2060 | 324 | 2245198 | 2245521 | BCGDan_sapMKO_2062 | BCGDan_sapMKO_2062 | 324 |
| 2281614 | 2282102 | Rv2035 | Rv2035 | 489 | 2265266 | 2265754 | BCG_2054 | BCG_2054 | 489 | 2245257 | 2245745 | BCGDan_2061 | BCGDan_2061 | 489 | 2245518 | 2246006 | BCGDan_sapMKO_2063 | BCGDan_sapMKO_2063 | 489 |
| 2282099 | 2282740 | Rv2036 | Rv2036 | 642 | 2265751 | 2266392 | BCG_2055 | BCG_2055 | 642 | 2245742 | 2246383 | BCGDan_2062 | BCGDan_2062 | 642 | 2246003 | 2246644 | BCGDan_sapMKO_2064 | BCGDan_sapMKO_2064 | 642 |
| 2282747 | 2283721 | Rv2037c | Rv2037c | 975 | 2266399 | 2267373 | BCG_2056c | BCG_2056c | 975 | 2246390 | 2247364 | BCGDan_2063 | BCGDan_2063 | 975 | 2246651 | 2247625 | BCGDan_sapMKO_2065 | BCGDan_sapMKO_2065 | 975 |
| 2283723 | 2284796 | Rv2038c | Rv2038c | 1074 | 2267375 | 2268448 | BCG_2057c | BCG_2057c | 1074 | 2247366 | 2248439 | BCGDan_2064 | BCGDan_2064 | 1074 | 2247627 | 2248700 | BCGDan_sapMKO_2066 | BCGDan_sapMKO_2066 | 1074 |
| 2284799 | 2285641 | Rv2039c | Rv2039c | 843 | 2268451 | 2269293 | BCG_2058c | BCG_2058c | 843 | 2248442 | 2249284 | BCGDan_2065 | BCGDan_2065 | 843 | 2248703 | 2249545 | BCGDan_sapMKO_2067 | BCGDan_sapMKO_2067 | 843 |
| 2285628 | 2286530 | Rv2040c | Rv2040c | 903 | 2269280 | 2270182 | BCG_2059c | BCG_2059c | 903 | 2249271 | 2250173 | BCGDan_2066 | BCGDan_2066 | 903 | 2249532 | 2250395 | BCGDan_sapMKO_2068 | BCGDan_sapMKO_2068 | 864 |
| 2286527 | 2287846 | Rv2041c | Rv2041c | 1320 | 2270179 | 2271498 | BCG_2060c | BCG_2060c | 1320 | 2250170 | 2251489 | BCGDan_2067 | BCGDan_2067 | 1320 | 2250431 | 2251750 | BCGDan_sapMKO_2069 | BCGDan_sapMKO_2069 | 1320 |
| 2287884 | 2288681 | Rv2042c | Rv2042c | 798 | 2271536 | 2272333 | BCG_2061c | BCG_2061c | 798 | 2251527 | 2252324 | BCGDan_2068 | BCGDan_2068 | 798 | 2251788 | 2252585 | BCGDan_sapMKO_2070 | BCGDan_sapMKO_2070 | 798 |
| 2288681 | 2289241 | pncA | Rv2043c | 561 | 2272333 | 2272893 | pncA | BCG_2062c | 561 | 2252324 | 2252884 | pncA | BCGDan_2069 | 561 | 2252585 | 2253145 | pncA | BCGDan_sapMKO_2071 | 561 |
| 2289282 | 2289599 | Rv2044c | Rv2044c | 318 | 2272934 | 2273251 | BCG_2063c | BCG_2063c | 318 | 2252925 | 2253242 | BCGDan_2070 | BCGDan_2070 | 318 | 2253186 | 2253536 | BCGDan_sapMKO_2072 | BCGDan_sapMKO_2072 | 351 |
| 2289685 | 2291220 | lipT | Rv2045c | 1536 | 2273337 | 2274872 | lipT | BCG_2064c | 1536 | 2253328 | 2254863 | lipT | BCGDan_2071 | 1536 | 2253589 | 2255094 | lipT | BCGDan_sapMKO_2073 | 1506 |
| 2291269 | 2291925 | lppI | Rv2046 | 657 | 2274921 | 2275577 | lppI | BCG_2065 | 657 | 2254912 | 2255568 | lppI | BCGDan_2072 | 657 | 2255173 | 2255829 | lppI | BCGDan_sapMKO_2074 | 657 |
| 2291962 | 2294526 | Rv2047c | Rv2047c | 2565 | 2275614 | 2278178 | BCG_2066c | BCG_2066c | 2565 | 2255605 | 2258169 | BCGDan_2073 | BCGDan_2073 | 2565 | 2255866 | 2258430 | BCGDan_sapMKO_2075 | BCGDan_sapMKO_2075 | 2565 |
| 2294531 | 2306986 | pks12 | Rv2048c | 12456 | 2278183 | 2290638 | pks12 | BCG_2067c | 12456 | 2258174 | 2270629 | pks12 | BCGDan_2074 | 12456 | 2258435 | 2270878 | pks12 | BCGDan_sapMKO_2076 | 12444 |
| 2307293 | 2307517 | Rv2049c | Rv2049c | 225 | 2290945 | 2291169 | BCG_2068c | BCG_2068c | 225 | 2270936 | 2271160 | BCGDan_2075 | BCGDan_2075 | 225 | 2271185 | 2271409 | BCGDan_sapMKO_2077 | BCGDan_sapMKO_2077 | 225 |
| 2307821 | 2308156 | Rv2050 | Rv2050 | 336 | 2291473 | 2291808 | BCG_2069 | BCG_2069 | 336 | 2271464 | 2271799 | BCGDan_2076 | BCGDan_2076 | 336 | 2271752 | 2272048 | BCGDan_sapMKO_2078 | BCGDan_sapMKO_2078 | 297 |
| 2308131 | 2310755 | ppm1 | Rv2051c | 2625 | 2291783 | 2294407 | ppm1 | BCG_2070c | 2625 | 2271774 | 2274398 | ppm1 | BCGDan_2077 | 2625 | 2272023 | 2274812 | ppm1 | BCGDan_sapMKO_2079 | 2790 |
| 2310913 | 2312517 | Rv2052c | Rv2052c | 1605 | 2294565 | 2296169 | BCG_2071c | BCG_2071c | 1605 | 2274556 | 2276160 | BCGDan_2078 | BCGDan_2078 | 1605 | 2274805 | 2276409 | BCGDan_sapMKO_2080 | BCGDan_sapMKO_2080 | 1605 |
| 2312522 | 2313049 | fxsA | Rv2053c | 528 | 2296174 | 2296701 | BCG_2072c | BCG_2072c | 528 | 2276165 | 2276692 | fxsA | BCGDan_2079 | 528 | 2276414 | 2276944 | BCGDan_sapMKO_2081 | BCGDan_sapMKO_2081 | 531 |
| 2313125 | 2313838 | Rv2054 | Rv2054 | 714 | 2296777 | 2297490 | BCG_2073 | BCG_2073 | 714 | 2276768 | 2277481 | BCGDan_2080 | BCGDan_2080 | 714 | 2277017 | 2277730 | BCGDan_sapMKO_2082 | BCGDan_sapMKO_2082 | 714 |
| 2314087 | 2314353 | rpsR2 | Rv2055c | 267 | 2297739 | 2298005 | rpsR2 | BCG_2074c | 267 | 2277730 | 2277996 | rpsR2 | BCGDan_2081 | 267 | 2277979 | 2278245 | rpsR2 | BCGDan_sapMKO_2083 | 267 |
| 2314354 | 2314659 | rpsN2 | Rv2056c | 306 | 2298006 | 2298311 | rpsN2 | BCG_2075c | 306 | 2277997 | 2278302 | rpsN2 | BCGDan_2082 | 306 | 2278246 | 2278551 | rpsN2 | BCGDan_sapMKO_2084 | 306 |
| 2314661 | 2314825 | rpmG1 | Rv2057c | 165 | 2298313 | 2298477 | rpmG1 | BCG_2076c | 165 | 2278304 | 2278468 | rpmG1 | BCGDan_2083 | 165 | 2278553 | 2278717 | rpmG1 | BCGDan_sapMKO_2085 | 165 |

| | | | | | | | | | | | | | | | | | | | |
|---|---|---|---|---|---|---|---|---|---|---|---|---|---|---|---|---|---|---|---|
| 2314825 | 2315061 | rpmB2 | Rv2058c | 237 | 2298477 | 2298713 | rpmB2 | BCG_2077c | 237 | 2278468 | 2278704 | rpmB2 | BCGDan_2084 | 237 | 2278717 | 2278953 | rpmB2 | BCGDan_sapMKO_2086 | 237 |
| 2315174 | 2316709 | Rv2059 | Rv2059 | 1536 | 2298826 | 2300361 | BCG_2078 | BCG_2078 | 1536 | 2278817 | 2280352 | BCGDan_2085 | BCGDan_2085 | 1536 | 2279066 | 2280601 | BCGDan_sapMKO_2087 | BCGDan_sapMKO_2087 | 1536 |
| 2316279 | 2316680 | Rv2060 | Rv2060 | 402 | 2299931 | 2300332 | BCG_2079 | BCG_2079 | 402 | 2279922 | 2280323 | BCGDan_2086 | BCGDan_2086 | 402 | 2280171 | 2280572 | BCGDan_sapMKO_2088 | BCGDan_sapMKO_2088 | 402 |
| 2316681 | 2317085 | Rv2061c | Rv2061c | 405 | 2300333 | 2300737 | BCG_2080c | BCG_2080c | 405 | 2280324 | 2280728 | BCGDan_2087 | BCGDan_2087 | 405 | 2280573 | 2280977 | BCGDan_sapMKO_2089 | BCGDan_sapMKO_2089 | 405 |
| 2317169 | 2320753 | cobN | Rv2062c | 3585 | 2300821 | 2304405 | cobN | BCG_2081c | 3585 | 2280812 | 2284396 | cobN | BCGDan_2088 | 3585 | 2281061 | 2284645 | cobN | BCGDan_sapMKO_2090 | 3585 |
| 2320831 | 2321064 | mazE7 | Rv2063 | 234 | 2304483 | 2304716 | BCG_2082 | BCG_2082 | 234 | 2284474 | 2284707 | BCGDan_2089 | BCGDan_2089 | 234 | 2284723 | 2284956 | BCGDan_sapMKO_2091 | BCGDan_sapMKO_2091 | 234 |
| 2321451 | 2322542 | cobG | Rv2064 | 1092 | 2305103 | 2306194 | cobG | BCG_2083 | 1092 | 2285094 | 2286185 | cobG | BCGDan_2090 | 1092 | 2285343 | 2286434 | cobG | BCGDan_sapMKO_2092 | 1092 |
| 2322552 | 2323178 | cobH | Rv2065 | 627 | 2306204 | 2306830 | cobH | BCG_2084 | 627 | 2286195 | 2286821 | cobH | BCGDan_2091 | 627 | 2286444 | 2287070 | cobH | BCGDan_sapMKO_2093 | 627 |
| 2323175 | 2324701 | cobI | Rv2066 | 1527 | 2306827 | 2308353 | cobI | BCG_2085 | 1527 | 2286818 | 2288344 | cobI | BCGDan_2092 | 1527 | 2287067 | 2288593 | cobI | BCGDan_sapMKO_2094 | 1527 |
| 2324647 | 2325870 | Rv2067c | Rv2067c | 1224 | 2308299 | 2309522 | BCG_2086c | BCG_2086c | 1224 | 2288290 | 2289513 | BCGDan_2093 | BCGDan_2093 | 1224 | 2288539 | 2289762 | BCGDan_sapMKO_2095 | BCGDan_sapMKO_2095 | 1224 |
| 2325886 | 2326809 | blaC | Rv2068c | 924 | 2309538 | 2310461 | blaC | BCG_2087c | 924 | 2289529 | 2290452 | blaC | BCGDan_2094 | 924 | 2289778 | 2290701 | blaC | BCGDan_sapMKO_2096 | 924 |
| 2326944 | 2327501 | sigC | Rv2069 | 558 | 2310596 | 2311153 | sigC | BCG_2088 | 558 | 2290587 | 2291144 | sigC | BCGDan_2095 | 558 | 2290836 | 2291393 | sigC | BCGDan_sapMKO_2097 | 558 |
| 2327491 | 2328225 | cobK | Rv2070c | 735 | 2311143 | 2311838 | cobK | BCG_2089c | 696 | 2291134 | 2291829 | cobK | BCGDan_2096 | 696 | 2291383 | 2292078 | cobK | BCGDan_sapMKO_2098 | 696 |
| 2328222 | 2328977 | cobM | Rv2071c | 756 | 2311875 | 2312630 | cobM | BCG_2090c | 756 | 2291866 | 2292621 | cobM | BCGDan_2097 | 756 | 2292115 | 2292888 | cobM | BCGDan_sapMKO_2099 | 774 |
| NA | NA | NA | NA | NA | 2312627 | 2313511 | cobLb | BCG_2091c | 885 | 2292618 | 2293451 | cobLb | BCGDan_2098 | 834 | 2292867 | 2293700 | cobLb | BCGDan_sapMKO_2100 | 834 |
| 2328974 | 2330146 | cobL | Rv2072c | 1173 | 2313453 | 2313641 | cobLa | BCG_2092c | 189 | 2293444 | 2293632 | cobLa | BCGDan_2099 | 189 | 2293693 | 2293881 | cobLa | BCGDan_sapMKO_2101 | 189 |
| 2331416 | 2332879 | Rv2075c | Rv2075c | 1464 | 2313716 | 2314504 | BCG_2093c | BCG_2093c | 789 | 2293707 | 2294495 | BCGDan_2100 | BCGDan_2100 | 789 | 2293956 | 2294744 | BCGDan_sapMKO_2102 | BCGDan_sapMKO_2102 | 789 |
| 2333037 | 2333288 | Rv2076c | Rv2076c | 252 | 2314662 | 2314913 | BCG_2094c | BCG_2094c | 252 | 2294653 | 2294904 | BCGDan_2101 | BCGDan_2101 | 252 | 2294902 | 2295153 | BCGDan_sapMKO_2103 | BCGDan_sapMKO_2103 | 252 |
| 2333323 | 2334294 | Rv2077c | Rv2077c | 972 | 2314948 | 2315919 | BCG_2095c | BCG_2095c | 972 | 2294939 | 2295910 | BCGDan_2102 | BCGDan_2102 | 972 | 2295188 | 2296159 | BCGDan_sapMKO_2104 | BCGDan_sapMKO_2104 | 972 |
| 2334295 | 2334594 | Rv2077A | Rv2077A | 300 | 2315920 | 2316219 | BCG_2096c | BCG_2096c | 300 | 2295911 | 2296210 | BCGDan_2103 | BCGDan_2103 | 300 | 2296160 | 2296459 | BCGDan_sapMKO_2105 | BCGDan_sapMKO_2105 | 300 |
| 2335059 | 2335373 | Rv2078 | Rv2078 | 315 | 2316684 | 2316998 | BCG_2097 | BCG_2097 | 315 | 2296675 | 2296989 | BCGDan_2104 | BCGDan_2104 | 315 | 2296924 | 2297238 | BCGDan_sapMKO_2106 | BCGDan_sapMKO_2106 | 315 |
| 2335355 | 2337325 | Rv2079 | Rv2079 | 1971 | 2316980 | 2318950 | BCG_2098 | BCG_2098 | 1971 | 2296971 | 2298941 | BCGDan_2105 | BCGDan_2105 | 1971 | 2297220 | 2299190 | BCGDan_sapMKO_2107 | BCGDan_sapMKO_2107 | 1971 |
| 2337306 | 2337869 | lppJ | Rv2080 | 564 | 2318931 | 2319494 | lppJ | BCG_2099 | 564 | 2298922 | 2299485 | lppJ | BCGDan_2106 | 564 | 2299171 | 2299734 | lppJ | BCGDan_sapMKO_2108 | 564 |
| 2338065 | 2338505 | Rv2081c | Rv2081c | 441 | 2319530 | 2320141 | BCG_2100c | BCG_2100c | 612 | 2299521 | 2300126 | BCGDan_2107 | BCGDan_2107 | 606 | 2299770 | 2300375 | BCGDan_sapMKO_2109 | BCGDan_sapMKO_2109 | 606 |
| 2338709 | 2340874 | Rv2082 | Rv2082 | 2166 | 2320345 | 2322432 | BCG_2101 | BCG_2101 | 2088 | 2300330 | 2302417 | BCGDan_2108 | BCGDan_2108 | 2088 | 2300579 | 2302666 | BCGDan_sapMKO_2110 | BCGDan_sapMKO_2110 | 2088 |
| 2340871 | 2341815 | Rv2083 | Rv2083 | 945 | 2322429 | 2323373 | BCG_2102 | BCG_2102 | 945 | 2302414 | 2303358 | BCGDan_2109 | BCGDan_2109 | 945 | 2302663 | 2303607 | BCGDan_sapMKO_2111 | BCGDan_sapMKO_2111 | 945 |
| 2341808 | 2342944 | Rv2084 | Rv2084 | 1137 | 2323366 | 2324361 | BCG_2103 | BCG_2103 | 996 | 2303351 | 2304346 | BCGDan_2110 | BCGDan_2110 | 996 | 2303600 | 2304595 | BCGDan_sapMKO_2112 | BCGDan_sapMKO_2112 | 996 |
| NA | NA | NA | NA | NA | 2324181 | 2324513 | BCG_2104 | BCG_2104 | 333 | 2304166 | 2304498 | BCGDan_2111 | BCGDan_2111 | 333 | 2304415 | 2304744 | BCGDan_sapMKO_2113 | BCGDan_sapMKO_2113 | 330 |
| 2343027 | 2343332 | Rv2085 | Rv2085 | 306 | 2324596 | 2324901 | BCG_2105 | BCG_2105 | 306 | 2304581 | 2304886 | BCGDan_2112 | BCGDan_2112 | 306 | 2304830 | 2305135 | BCGDan_sapMKO_2114 | BCGDan_sapMKO_2114 | 306 |
| 2343311 | 2343916 | Rv2086 | Rv2086 | 606 | 2324880 | 2325485 | BCG_2106 | BCG_2106 | 606 | 2304865 | 2305470 | BCGDan_2113 | BCGDan_2113 | 606 | 2305114 | 2305719 | BCGDan_sapMKO_2115 | BCGDan_sapMKO_2115 | 606 |
| 2343994 | 2344224 | Rv2087 | Rv2087 | 231 | 2325563 | 2325793 | BCG_2107 | BCG_2107 | 231 | 2305548 | 2305778 | BCGDan_2114 | BCGDan_2114 | 231 | 2305797 | 2306027 | BCGDan_sapMKO_2116 | BCGDan_sapMKO_2116 | 231 |
| 2344411 | 2346180 | pknJ | Rv2088 | 1770 | 2325980 | 2327749 | pknJ | BCG_2108 | 1770 | 2305965 | 2307734 | pknJ | BCGDan_2115 | 1770 | 2306214 | 2307983 | pknJ | BCGDan_sapMKO_2117 | 1770 |
| 2346197 | 2347324 | pepE | Rv2089c | 1128 | 2327766 | 2328893 | pepE | BCG_2109c | 1128 | 2307751 | 2308878 | pepE | BCGDan_2116 | 1128 | 2308000 | 2309214 | pepE | BCGDan_sapMKO_2118 | 1215 |
| 2347373 | 2348554 | Rv2090 | Rv2090 | 1182 | 2328942 | 2330009 | BCG_2110 | BCG_2110 | 1068 | 2308927 | 2309994 | BCGDan_2117 | BCGDan_2117 | 1068 | 2309176 | 2310243 | BCGDan_sapMKO_2119 | BCGDan_sapMKO_2119 | 1068 |
| 2348558 | 2349292 | Rv2091c | Rv2091c | 735 | 2330013 | 2330747 | BCG_2111c | BCG_2111c | 735 | 2309998 | 2310732 | BCGDan_2118 | BCGDan_2118 | 735 | 2310247 | 2310981 | BCGDan_sapMKO_2120 | BCGDan_sapMKO_2120 | 735 |
| 2349334 | 2352054 | helY | Rv2092c | 2721 | 2330789 | 2333509 | helY | BCG_2112c | 2721 | 2310774 | 2313494 | helY | BCGDan_2119 | 2721 | 2311023 | 2313743 | helY | BCGDan_sapMKO_2121 | 2721 |
| 2352103 | 2353029 | tatC | Rv2093c | 927 | 2333558 | 2334484 | tatC | BCG_2113c | 927 | 2313543 | 2314469 | tatC | BCGDan_2120 | 927 | 2313792 | 2314646 | tatC | BCGDan_sapMKO_2122 | 855 |
| 2353046 | 2353297 | tatA | Rv2094c | 252 | 2334501 | 2334752 | tatA | BCG_2114c | 252 | 2314486 | 2314737 | tatA | BCGDan_2121 | 252 | 2314735 | 2314986 | tatA | BCGDan_sapMKO_2123 | 252 |
| 2353365 | 2354315 | pafC | Rv2095c | 951 | 2334820 | 2335770 | BCG_2115c | BCG_2115c | 951 | 2314805 | 2315755 | BCGDan_2122 | BCGDan_2122 | 951 | 2315054 | 2316004 | BCGDan_sapMKO_2124 | BCGDan_sapMKO_2124 | 951 |
| 2354312 | 2355310 | pafB | Rv2096c | 999 | 2335767 | 2336765 | BCG_2116c | BCG_2116c | 999 | 2315752 | 2316750 | BCGDan_2123 | BCGDan_2123 | 999 | 2316001 | 2316999 | BCGDan_sapMKO_2125 | BCGDan_sapMKO_2125 | 999 |
| 2355319 | 2356677 | pafA | Rv2097c | 1359 | 2336774 | 2338132 | BCG_2117c | BCG_2117c | 1359 | 2316759 | 2318117 | BCGDan_2124 | BCGDan_2124 | 1359 | 2317008 | 2318351 | BCGDan_sapMKO_2126 | BCGDan_sapMKO_2126 | 1344 |
| 2356729 | 2358033 | PE_PGRS36 | Rv2098c | 1305 | 2338184 | 2339659 | PE_PGRS36 | BCG_2118c | 1476 | 2318169 | 2319644 | PE_PGRS36 | BCGDan_2125 | 1476 | 2318418 | 2319893 | PE_PGRS36 | BCGDan_sapMKO_2127 | 1476 |
| 2358389 | 2360041 | Rv2100 | Rv2100 | 1653 | 2339842 | 2341494 | BCG_2119 | BCG_2119 | 1653 | 2319827 | 2321479 | BCGDan_2126 | BCGDan_2126 | 1653 | 2320076 | 2321728 | BCGDan_sapMKO_2128 | BCGDan_sapMKO_2128 | 1653 |
| 2360240 | 2363281 | helZ | Rv2101 | 3042 | 2341693 | 2344734 | helZ | BCG_2120 | 3042 | 2321678 | 2324719 | helZ | BCGDan_2127 | 3042 | 2321927 | 2324968 | helZ | BCGDan_sapMKO_2129 | 3042 |
| 2363391 | 2364107 | Rv2102 | Rv2102 | 717 | 2344844 | 2345560 | BCG_2121 | BCG_2121 | 717 | 2324829 | 2325545 | BCGDan_2128 | BCGDan_2128 | 717 | 2324961 | 2325794 | BCGDan_sapMKO_2130 | BCGDan_sapMKO_2130 | 834 |
| 2364086 | 2364520 | vapC37 | Rv2103c | 435 | 2345539 | 2345973 | BCG_2122c | BCG_2122c | 435 | 2325524 | 2325958 | BCGDan_2129 | BCGDan_2129 | 435 | 2325773 | 2326207 | BCGDan_sapMKO_2131 | BCGDan_sapMKO_2131 | 435 |
| 2364527 | 2364781 | vapB37 | Rv2104c | 255 | 2345980 | 2346234 | BCG_2123c | BCG_2123c | 255 | 2325965 | 2326219 | BCGDan_2130 | BCGDan_2130 | 255 | 2326214 | 2326468 | BCGDan_sapMKO_2132 | BCGDan_sapMKO_2132 | 255 |
| 2367359 | 2367655 | PE22 | Rv2107 | 297 | 2347454 | 2347750 | PE22 | BCG_2124 | 297 | 2327439 | 2327735 | PE22 | BCGDan_2131 | 297 | 2327688 | 2327984 | PE22 | BCGDan_sapMKO_2133 | 297 |
| 2367711 | 2368442 | PPE36 | Rv2108 | 732 | 2347806 | 2348537 | PPE36 | BCG_2125 | 732 | 2327791 | 2328522 | PPE36 | BCGDan_2132 | 732 | 2328040 | 2328771 | PPE36 | BCGDan_sapMKO_2134 | 732 |
| 2368983 | 2369729 | prcA | Rv2109c | 747 | 2349077 | 2349823 | prcA | BCG_2126c | 747 | 2329062 | 2329808 | prcA | BCGDan_2133 | 747 | 2329311 | 2330057 | prcA | BCGDan_sapMKO_2135 | 747 |
| 2369726 | 2370601 | prcB | Rv2110c | 876 | 2349820 | 2350695 | prcB | BCG_2127c | 876 | 2329805 | 2330680 | prcB | BCGDan_2134 | 876 | 2330054 | 2330929 | prcB | BCGDan_sapMKO_2136 | 876 |
| 2370598 | 2370792 | pup | Rv2111c | 195 | 2350692 | 2350886 | BCG_2128c | BCG_2128c | 195 | 2330677 | 2330871 | BCGDan_2135 | BCGDan_2135 | 195 | 2330926 | 2331141 | BCGDan_sapMKO_2137 | BCGDan_sapMKO_2137 | 216 |
| 2370905 | 2372569 | dop | Rv2112c | 1665 | 2350999 | 2352606 | BCG_2129c | BCG_2129c | 1608 | 2330984 | 2332591 | BCGDan_2136 | BCGDan_2136 | 1608 | 2331233 | 2332750 | BCGDan_sapMKO_2138 | BCGDan_sapMKO_2138 | 1518 |
| 2372630 | 2373823 | Rv2113 | Rv2113 | 1194 | 2352667 | 2353860 | BCG_2130 | BCG_2130 | 1194 | 2332652 | 2333845 | BCGDan_2137 | BCGDan_2137 | 1194 | 2332949 | 2334094 | BCGDan_sapMKO_2139 | BCGDan_sapMKO_2139 | 1146 |
| 2373834 | 2374457 | Rv2114 | Rv2114 | 624 | 2353871 | 2354494 | BCG_2131 | BCG_2131 | 624 | 2333856 | 2334479 | BCGDan_2138 | BCGDan_2138 | 624 | 2334105 | 2334728 | BCGDan_sapMKO_2140 | BCGDan_sapMKO_2140 | 624 |
| 2374461 | 2376290 | mpa | Rv2115c | 1830 | 2354498 | 2356327 | BCG_2132c | BCG_2132c | 1830 | 2334483 | 2336312 | BCGDan_2139 | BCGDan_2139 | 1830 | 2334732 | 2336342 | BCGDan_sapMKO_2141 | BCGDan_sapMKO_2141 | 1611 |
| 2376571 | 2377140 | lppK | Rv2116 | 570 | 2356608 | 2357177 | lppK | BCG_2133 | 570 | 2336593 | 2337162 | lppK | BCGDan_2140 | 570 | 2336842 | 2337411 | lppK | BCGDan_sapMKO_2142 | 570 |
| 2377148 | 2377441 | Rv2117 | Rv2117 | 294 | 2357185 | 2357478 | BCG_2134 | BCG_2134 | 294 | 2337170 | 2337463 | BCGDan_2141 | BCGDan_2141 | 294 | 2337419 | 2337712 | BCGDan_sapMKO_2143 | BCGDan_sapMKO_2143 | 294 |
| 2377470 | 2378312 | Rv2118c | Rv2118c | 843 | 2357507 | 2358349 | BCG_2135c | BCG_2135c | 843 | 2337492 | 2338334 | BCGDan_2142 | BCGDan_2142 | 843 | 2337741 | 2338583 | BCGDan_sapMKO_2144 | BCGDan_sapMKO_2144 | 843 |
| 2378386 | 2379222 | Rv2119 | Rv2119 | 837 | 2358423 | 2359259 | BCG_2136 | BCG_2136 | 837 | 2338408 | 2339244 | BCGDan_2143 | BCGDan_2143 | 837 | 2338657 | 2339493 | BCGDan_sapMKO_2145 | BCGDan_sapMKO_2145 | 837 |
| 2379245 | 2379727 | Rv2120c | Rv2120c | 483 | 2359282 | 2359764 | BCG_2137c | BCG_2137c | 483 | 2339267 | 2339749 | BCGDan_2144 | BCGDan_2144 | 483 | 2339516 | 2339998 | BCGDan_sapMKO_2146 | BCGDan_sapMKO_2146 | 483 |
| 2379806 | 2380660 | hisG | Rv2121c | 855 | 2359843 | 2360697 | hisG | BCG_2138c | 855 | 2339828 | 2340682 | hisG | BCGDan_2145 | 855 | 2340077 | 2340931 | hisG | BCGDan_sapMKO_2147 | 855 |
| 2380663 | 2380944 | hisE | Rv2122c | 282 | 2360700 | 2360981 | hisE | BCG_2139c | 282 | 2340685 | 2340966 | hisE | BCGDan_2146 | 282 | 2340934 | 2341215 | hisE | BCGDan_sapMKO_2148 | 282 |
| 2381071 | 2382492 | PPE37 | Rv2123 | 1422 | 2361108 | 2362502 | PPE37 | BCG_2140 | 1395 | 2341093 | 2342487 | PPE37 | BCGDan_2147 | 1395 | 2341342 | 2342736 | PPE37 | BCGDan_sapMKO_2149 | 1395 |
| 2382489 | 2386067 | metH | Rv2124c | 3579 | 2362499 | 2366077 | metH | BCG_2141c | 3579 | 2342484 | 2346062 | metH | BCGDan_2148 | 3579 | 2342733 | 2346311 | metH | BCGDan_sapMKO_2150 | 3579 |
| 2386293 | 2387171 | Rv2125 | Rv2125 | 879 | 2366303 | 2367181 | BCG_2142 | BCG_2142 | 879 | 2346288 | 2347166 | BCGDan_2149 | BCGDan_2149 | 879 | 2346537 | 2347415 | BCGDan_sapMKO_2151 | BCGDan_sapMKO_2151 | 879 |
| 2387202 | 2387972 | PE_PGRS37 | Rv2126c | 771 | 2367212 | 2367982 | PE_PGRS37 | BCG_2143c | 771 | 2347197 | 2347967 | PE_PGRS37 | BCGDan_2150 | 771 | 2347446 | 2348216 | PE_PGRS37 | BCGDan_sapMKO_2152 | 771 |
| 2388616 | 2390085 | ansP1 | Rv2127 | 1470 | 2368626 | 2370095 | ansP1 | BCG_2144 | 1470 | 2348611 | 2350080 | ansP1 | BCGDan_2151 | 1470 | 2348860 | 2350329 | ansP1 | BCGDan_sapMKO_2153 | 1470 |
| 2390085 | 2390288 | Rv2128 | Rv2128 | 204 | 2370095 | 2370298 | BCG_2145 | BCG_2145 | 204 | 2350080 | 2350283 | BCGDan_2152 | BCGDan_2152 | 204 | 2350329 | 2350532 | BCGDan_sapMKO_2154 | BCGDan_sapMKO_2154 | 204 |
| 2390308 | 2391189 | Rv2129c | Rv2129c | 882 | 2370318 | 2371199 | BCG_2146c | BCG_2146c | 882 | 2350303 | 2351184 | BCGDan_2153 | BCGDan_2153 | 882 | 2350552 | 2351433 | BCGDan_sapMKO_2155 | BCGDan_sapMKO_2155 | 882 |
| 2391215 | 2392459 | mshC | Rv2130c | 1245 | 2371225 | 2372469 | cysS2 | BCG_2147c | 1245 | 2351210 | 2352454 | cysS2 | BCGDan_2154 | 1245 | 2351459 | 2352673 | cysS2 | BCGDan_sapMKO_2156 | 1215 |
| 2392517 | 2393320 | cysQ | Rv2131c | 804 | 2372527 | 2373330 | cysQ | BCG_2148c | 804 | 2352512 | 2353315 | cysQ | BCGDan_2155 | 804 | 2352761 | 2353564 | cysQ | BCGDan_sapMKO_2157 | 804 |
| 2393411 | 2393641 | Rv2132 | Rv2132 | 231 | 2373421 | 2373651 | BCG_2149 | BCG_2149 | 231 | 2353406 | 2353636 | BCGDan_2156 | BCGDan_2156 | 231 | 2353655 | 2353885 | BCGDan_sapMKO_2158 | BCGDan_sapMKO_2158 | 231 |
| 2393851 | 2394639 | Rv2133c | Rv2133c | 789 | 2373861 | 2374649 | BCG_2150c | BCG_2150c | 789 | 2353846 | 2354634 | BCGDan_2157 | BCGDan_2157 | 789 | 2354095 | 2354910 | BCGDan_sapMKO_2159 | BCGDan_sapMKO_2159 | 816 |
| 2394650 | 2395237 | Rv2134c | Rv2134c | 588 | 2374660 | 2375247 | BCG_2151c | BCG_2151c | 588 | 2354645 | 2355232 | BCGDan_2158 | BCGDan_2158 | 588 | 2354894 | 2355481 | BCGDan_sapMKO_2160 | BCGDan_sapMKO_2160 | 588 |
| 2395301 | 2396011 | Rv2135c | Rv2135c | 711 | 2375311 | 2376021 | BCG_2152c | BCG_2152c | 711 | 2355296 | 2356006 | BCGDan_2159 | BCGDan_2159 | 711 | 2355545 | 2356255 | BCGDan_sapMKO_2161 | BCGDan_sapMKO_2161 | 711 |
| 2396008 | 2396838 | Rv2136c | Rv2136c | 831 | 2376018 | 2376848 | BCG_2153c | BCG_2153c | 831 | 2356003 | 2356833 | BCGDan_2160 | BCGDan_2160 | 831 | 2356252 | 2357100 | BCGDan_sapMKO_2162 | BCGDan_sapMKO_2162 | 849 |
| 2396902 | 2397315 | Rv2137c | Rv2137c | 414 | 2376912 | 2377325 | BCG_2154c | BCG_2154c | 414 | 2356897 | 2357310 | BCGDan_2161 | BCGDan_2161 | 414 | 2357146 | 2357559 | BCGDan_sapMKO_2163 | BCGDan_sapMKO_2163 | 414 |
| 2397330 | 2398406 | lppL | Rv2138 | 1077 | 2377340 | 2378416 | lppL | BCG_2155 | 1077 | 2357325 | 2358401 | lppL | BCGDan_2162 | 1077 | 2357625 | 2358650 | lppL | BCGDan_sapMKO_2164 | 1026 |
| 2398720 | 2399793 | pyrD | Rv2139 | 1074 | 2378730 | 2379803 | pyrD | BCG_2156 | 1074 | 2358715 | 2359788 | pyrD | BCGDan_2163 | 1074 | 2358919 | 2360037 | pyrD | BCGDan_sapMKO_2165 | 1119 |
| 2399798 | 2400328 | TB18.6 | Rv2140c | 531 | 2379808 | 2380338 | TB18.6 | BCG_2157c | 531 | 2359793 | 2360323 | TB18.6 | BCGDan_2164 | 531 | 2360042 | 2360572 | TB18.6 | BCGDan_sapMKO_2166 | 531 |
| 2400376 | 2401722 | Rv2141c | Rv2141c | 1347 | 2380386 | 2381732 | BCG_2158c | BCG_2158c | 1347 | 2360371 | 2361717 | BCGDan_2165 | BCGDan_2165 | 1347 | 2360620 | 2361966 | BCGDan_sapMKO_2167 | BCGDan_sapMKO_2167 | 1347 |
| 2402193 | 2402510 | parE2 | Rv2142c | 318 | 2382203 | 2382520 | BCG_2159c | BCG_2159c | 318 | 2362188 | 2362505 | parE2 | BCGDan_2167 | 318 | 2362437 | 2362754 | parE2 | BCGDan_sapMKO_2169 | 318 |
| 2402507 | 2402722 | parD2 | Rv2142A | 216 | NA | NA | NA | NA | NA | 2362502 | 2362717 | parD2 | BCGDan_2168 | 216 | 2362751 | 2362966 | parD2 | BCGDan_sapMKO_2170 | 216 |
| 2402977 | 2404035 | Rv2143 | Rv2143 | 1059 | 2382987 | 2384045 | BCG_2160 | BCG_2160 | 1059 | 2362972 | 2364030 | BCGDan_2169 | BCGDan_2169 | 1059 | 2363221 | 2364279 | BCGDan_sapMKO_2171 | BCGDan_sapMKO_2171 | 1059 |
| 2404165 | 2404521 | Rv2144c | Rv2144c | 357 | 2384175 | 2384531 | BCG_2161c | BCG_2161c | 357 | 2364160 | 2364516 | BCGDan_2170 | BCGDan_2170 | 357 | 2364409 | 2364765 | BCGDan_sapMKO_2172 | BCGDan_sapMKO_2172 | 357 |

| | | | | | | | | | | | | | | | | | | | |
|---|---|---|---|---|---|---|---|---|---|---|---|---|---|---|---|---|---|---|---|
| 2404616 | 2405398 | wag31 | Rv2145c | 783 | 2384626 | 2385408 | wag31 | BCG_2162c | 783 | 2364611 | 2365393 | wag31 | BCGDan_2171 | 783 | 2364860 | 2365642 | wag31 | BCGDan_sapMKO_2173 | 783 |
| 2405666 | 2405956 | Rv2146c | Rv2146c | 291 | 2385676 | 2385966 | BCG_2163c | BCG_2163c | 291 | 2365661 | 2365951 | BCGDan_2172 | BCGDan_2172 | 291 | 2365910 | 2366197 | BCGDan_sapMKO_2174 | BCGDan_sapMKO_2174 | 288 |
| 2406118 | 2406843 | Rv2147c | Rv2147c | 726 | 2386128 | 2386853 | BCG_2164c | BCG_2164c | 726 | 2366113 | 2366838 | BCGDan_2173 | BCGDan_2173 | 726 | 2366362 | 2367018 | BCGDan_sapMKO_2175 | BCGDan_sapMKO_2175 | 657 |
| 2406840 | 2407616 | Rv2148c | Rv2148c | 777 | 2386850 | 2387626 | BCG_2165c | BCG_2165c | 777 | 2366835 | 2367611 | BCGDan_2174 | BCGDan_2174 | 777 | 2367084 | 2367869 | BCGDan_sapMKO_2176 | BCGDan_sapMKO_2176 | 786 |
| 2407622 | 2408374 | yfiH | Rv2149c | 753 | 2387632 | 2388384 | yfiH | BCG_2166c | 753 | 2367617 | 2368369 | yfiH | BCGDan_2175 | 753 | 2367866 | 2368573 | yfiH | BCGDan_sapMKO_2177 | 708 |
| 2408385 | 2409524 | ftsZ | Rv2150c | 1140 | 2388395 | 2389534 | ftsZ | BCG_2167c | 1140 | 2368380 | 2369519 | ftsZ | BCGDan_2176 | 1140 | 2368629 | 2369768 | ftsZ | BCGDan_sapMKO_2178 | 1140 |
| 2409697 | 2410641 | ftsQ | Rv2151c | 945 | 2389707 | 2390651 | ftsQ | BCG_2168c | 945 | 2369692 | 2370636 | ftsQ | BCGDan_2177 | 945 | 2369941 | 2370885 | ftsQ | BCGDan_sapMKO_2179 | 945 |
| 2410638 | 2412122 | murC | Rv2152c | 1485 | 2390648 | 2392132 | murC | BCG_2169c | 1485 | 2370633 | 2372117 | murC | BCGDan_2178 | 1485 | 2370882 | 2372366 | murC | BCGDan_sapMKO_2180 | 1485 |
| 2412119 | 2413351 | murG | Rv2153c | 1233 | 2392129 | 2393361 | murG | BCG_2170c | 1233 | 2372114 | 2373346 | murG | BCGDan_2179 | 1233 | 2372363 | 2373490 | murG | BCGDan_sapMKO_2181 | 1128 |
| 2413348 | 2414922 | ftsW | Rv2154c | 1575 | 2393358 | 2394932 | ftsW | BCG_2171c | 1575 | 2373343 | 2374917 | ftsW | BCGDan_2180 | 1575 | 2373592 | 2375007 | ftsW | BCGDan_sapMKO_2182 | 1416 |
| 2414934 | 2416394 | murD | Rv2155c | 1461 | 2394944 | 2396404 | BCG_2172c | BCG_2172c | 1461 | 2374929 | 2376389 | BCGDan_2181 | BCGDan_2181 | 1461 | 2375178 | 2376668 | BCGDan_sapMKO_2183 | BCGDan_sapMKO_2183 | 1491 |
| 2416396 | 2417475 | murX | Rv2156c | 1080 | 2396406 | 2397485 | murX | BCG_2173c | 1080 | 2376391 | 2377470 | murX | BCGDan_2182 | 1080 | 2376640 | 2377719 | murX | BCGDan_sapMKO_2184 | 1080 |
| 2417472 | 2419004 | murF | Rv2157c | 1533 | 2397482 | 2399014 | murF | BCG_2174c | 1533 | 2377467 | 2378999 | murF | BCGDan_2183 | 1533 | 2377716 | 2379248 | murF | BCGDan_sapMKO_2185 | 1533 |
| 2419001 | 2420608 | murE | Rv2158c | 1608 | 2399011 | 2400618 | murE | BCG_2175c | 1608 | 2378996 | 2380606 | murE | BCGDan_2184 | 1611 | 2379245 | 2380810 | murE | BCGDan_sapMKO_2186 | 1566 |
| 2420631 | 2421665 | Rv2159c | Rv2159c | 1035 | 2400641 | 2401675 | BCG_2176c | BCG_2176c | 1035 | 2380626 | 2381660 | BCGDan_2185 | BCGDan_2185 | 1035 | 2380875 | 2381909 | BCGDan_sapMKO_2187 | BCGDan_sapMKO_2187 | 1035 |
| 2421643 | 2422278 | Rv2160A | Rv2160A | 636 | 2401672 | 2402292 | BCG_2177c | BCG_2177c | 621 | 2381657 | 2382277 | BCGDan_2186 | BCGDan_2186 | 621 | 2381906 | 2382526 | BCGDan_sapMKO_2188 | BCGDan_sapMKO_2188 | 621 |
| 2422271 | 2423137 | Rv2161c | Rv2161c | 867 | 2402285 | 2403151 | BCG_2178c | BCG_2178c | 867 | 2382270 | 2383136 | BCGDan_2187 | BCGDan_2187 | 867 | 2382519 | 2383385 | BCGDan_sapMKO_2189 | BCGDan_sapMKO_2189 | 867 |
| 2423240 | 2424838 | PE_PGRS38 | Rv2162c | 1599 | 2403254 | 2404726 | PE_PGRS38 | BCG_2179c | 1473 | 2383239 | 2384711 | PE_PGRS38 | BCGDan_2188 | 1473 | 2383488 | 2384960 | PE_PGRS38 | BCGDan_sapMKO_2190 | 1473 |
| 2425048 | 2427087 | pbpB | Rv2163c | 2040 | 2404936 | 2406975 | pbpB | BCG_2180c | 2040 | 2384921 | 2386960 | pbpB | BCGDan_2189 | 2040 | 2385170 | 2386990 | pbpB | BCGDan_sapMKO_2191 | 1821 |
| 2427084 | 2428238 | Rv2164c | Rv2164c | 1155 | 2406972 | 2408126 | BCG_2181c | BCG_2181c | 1155 | 2386957 | 2388111 | BCGDan_2190 | BCGDan_2190 | 1155 | 2387206 | 2388360 | BCGDan_sapMKO_2192 | BCGDan_sapMKO_2192 | 1155 |
| 2428235 | 2429425 | Rv2165c | Rv2165c | 1191 | 2408123 | 2409313 | BCG_2182c | BCG_2182c | 1191 | 2388108 | 2389298 | mraW | BCGDan_2191 | 1191 | 2388357 | 2389547 | BCGDan_sapMKO_2193 | BCGDan_sapMKO_2193 | 1191 |
| 2429427 | 2429858 | Rv2166c | Rv2166c | 432 | 2409315 | 2409746 | BCG_2183c | BCG_2183c | 432 | 2389300 | 2389731 | BCGDan_2192 | BCGDan_2192 | 432 | 2389549 | 2389980 | BCGDan_sapMKO_2194 | BCGDan_sapMKO_2194 | 432 |
| 2431565 | 2431969 | Rv2169c | Rv2169c | 405 | 2410095 | 2410499 | BCG_2184c | BCG_2184c | 405 | 2390080 | 2390484 | BCGDan_2193 | BCGDan_2193 | 405 | 2390329 | 2390733 | BCGDan_sapMKO_2195 | BCGDan_sapMKO_2195 | 405 |
| 2432235 | 2432855 | Rv2170 | Rv2170 | 621 | 2410765 | 2411385 | BCG_2185 | BCG_2185 | 621 | 2390750 | 2391370 | BCGDan_2194 | BCGDan_2194 | 621 | 2391035 | 2391619 | BCGDan_sapMKO_2196 | BCGDan_sapMKO_2196 | 585 |
| 2432951 | 2433634 | lppM | Rv2171 | 684 | 2411481 | 2412164 | lppM | BCG_2186 | 684 | 2391466 | 2392149 | lppM | BCGDan_2195 | 684 | 2391739 | 2392398 | lppM | BCGDan_sapMKO_2197 | 660 |
| 2433631 | 2434536 | Rv2172c | Rv2172c | 906 | 2412161 | 2413066 | BCG_2187c | BCG_2187c | 906 | 2392146 | 2393051 | BCGDan_2196 | BCGDan_2196 | 906 | 2392395 | 2393300 | BCGDan_sapMKO_2198 | BCGDan_sapMKO_2198 | 906 |
| 2434847 | 2435905 | idsA2 | Rv2173 | 1059 | 2413377 | 2414435 | idsA2 | BCG_2188 | 1059 | 2393362 | 2394420 | idsA2 | BCGDan_2197 | 1059 | 2393611 | 2394669 | idsA2 | BCGDan_sapMKO_2199 | 1059 |
| 2435909 | 2437459 | mptA | Rv2174 | 1551 | 2414439 | 2416142 | BCG_2189 | BCG_2189 | 1704 | 2394424 | 2396127 | BCGDan_2198 | BCGDan_2198 | 1704 | 2394706 | 2396376 | BCGDan_sapMKO_2200 | BCGDan_sapMKO_2200 | 1671 |
| 2437446 | 2437886 | Rv2175c | Rv2175c | 441 | 2415977 | 2416417 | BCG_2190c | BCG_2190c | 441 | 2395962 | 2396402 | BCGDan_2199 | BCGDan_2199 | 441 | 2396211 | 2396651 | BCGDan_sapMKO_2201 | BCGDan_sapMKO_2201 | 441 |
| 2437941 | 2439140 | pknL | Rv2176 | 1200 | 2416472 | 2417671 | pknL | BCG_2191 | 1200 | 2396457 | 2397656 | pknL | BCGDan_2200 | 1200 | 2396706 | 2397905 | pknL | BCGDan_sapMKO_2202 | 1200 |
| 2439282 | 2439947 | Rv2177c | Rv2177c | 666 | 2417813 | 2418478 | BCG_2192c | BCG_2192c | 666 | 2397798 | 2398463 | BCGDan_2201 | BCGDan_2201 | 666 | 2398047 | 2398712 | BCGDan_sapMKO_2203 | BCGDan_sapMKO_2203 | 666 |
| 2440332 | 2441720 | aroG | Rv2178c | 1389 | 2418863 | 2420251 | aroG | BCG_2193c | 1389 | 2398848 | 2400236 | aroG | BCGDan_2202 | 1389 | 2399097 | 2400485 | aroG | BCGDan_sapMKO_2204 | 1389 |
| 2441811 | 2442317 | Rv2179c | Rv2179c | 507 | 2420342 | 2420848 | BCG_2194c | BCG_2194c | 507 | 2400327 | 2400833 | BCGDan_2203 | BCGDan_2203 | 507 | 2400576 | 2401082 | BCGDan_sapMKO_2205 | BCGDan_sapMKO_2205 | 507 |
| 2442327 | 2443214 | Rv2180c | Rv2180c | 888 | 2420858 | 2421745 | BCG_2195c | BCG_2195c | 888 | 2400843 | 2401730 | BCGDan_2204 | BCGDan_2204 | 888 | 2401092 | 2402030 | BCGDan_sapMKO_2206 | BCGDan_sapMKO_2206 | 939 |
| 2443302 | 2444585 | Rv2181 | Rv2181 | 1284 | 2421833 | 2423116 | BCG_2196 | BCG_2196 | 1284 | 2401818 | 2403101 | BCGDan_2205 | BCGDan_2205 | 1284 | 2402115 | 2403350 | BCGDan_sapMKO_2207 | BCGDan_sapMKO_2207 | 1236 |
| 2444586 | 2445329 | Rv2182c | Rv2182c | 744 | 2423117 | 2423860 | BCG_2197c | BCG_2197c | 744 | 2403102 | 2403845 | BCGDan_2206 | BCGDan_2206 | 744 | 2403351 | 2404094 | BCGDan_sapMKO_2208 | BCGDan_sapMKO_2208 | 744 |
| 2445415 | 2445810 | Rv2183c | Rv2183c | 396 | 2423946 | 2424341 | BCG_2198c | BCG_2198c | 396 | 2403931 | 2404326 | BCGDan_2207 | BCGDan_2207 | 396 | 2404180 | 2404584 | BCGDan_sapMKO_2209 | BCGDan_sapMKO_2209 | 405 |
| 2445807 | 2446946 | Rv2184c | Rv2184c | 1140 | 2424338 | 2425477 | BCG_2199c | BCG_2199c | 1140 | 2404323 | 2405462 | BCGDan_2208 | BCGDan_2208 | 1140 | 2404572 | 2405714 | BCGDan_sapMKO_2210 | BCGDan_sapMKO_2210 | 1143 |
| 2447066 | 2447500 | TB16.3 | Rv2185c | 435 | 2425597 | 2426031 | TB16.3 | BCG_2200c | 435 | 2405582 | 2406016 | TB16.3 | BCGDan_2209 | 435 | 2405831 | 2406265 | TB16.3 | BCGDan_sapMKO_2211 | 435 |
| 2447605 | 2447994 | Rv2186c | Rv2186c | 390 | 2426136 | 2426525 | BCG_2201c | BCG_2201c | 390 | 2406121 | 2406510 | BCGDan_2210 | BCGDan_2210 | 390 | 2406370 | 2406759 | BCGDan_sapMKO_2212 | BCGDan_sapMKO_2212 | 390 |
| 2448160 | 2449962 | fadD15 | Rv2187 | 1803 | 2426691 | 2427203 | BCG_2202 | BCG_2202 | 513 | 2406676 | 2407188 | BCGDan_2211 | BCGDan_2211 | 513 | 2406925 | 2407437 | BCGDan_sapMKO_2213 | BCGDan_sapMKO_2213 | 513 |
| NA | NA | NA | NA | NA | 2426966 | 2428492 | fadD15 | BCG_2203 | 1527 | 2406951 | 2408477 | fadD15 | BCGDan_2212 | 1527 | 2407170 | 2408726 | fadD15 | BCGDan_sapMKO_2214 | 1557 |
| 2449993 | 2451150 | pimB | Rv2188c | 1158 | 2428523 | 2429680 | BCG_2204c | BCG_2204c | 1158 | 2408508 | 2409557 | BCGDan_2213 | BCGDan_2213 | 1050 | 2408757 | 2409806 | BCGDan_sapMKO_2215 | BCGDan_sapMKO_2215 | 1050 |
| 2451247 | 2452020 | Rv2189c | Rv2189c | 774 | 2429777 | 2430550 | BCG_2205c | BCG_2205c | 774 | 2409762 | 2410535 | BCGDan_2214 | BCGDan_2214 | 774 | 2410011 | 2410844 | BCGDan_sapMKO_2216 | BCGDan_sapMKO_2216 | 834 |
| 2452115 | 2453272 | Rv2190c | Rv2190c | 1158 | 2430645 | 2431802 | BCG_2206c | BCG_2206c | 1158 | 2410630 | 2411787 | BCGDan_2215 | BCGDan_2215 | 1158 | 2410879 | 2411997 | BCGDan_sapMKO_2217 | BCGDan_sapMKO_2217 | 1119 |
| 2453819 | 2455756 | Rv2191 | Rv2191 | 1938 | 2432349 | 2434286 | BCG_2207 | BCG_2207 | 1938 | 2412334 | 2414271 | BCGDan_2216 | BCGDan_2216 | 1938 | 2412607 | 2414520 | BCGDan_sapMKO_2218 | BCGDan_sapMKO_2218 | 1914 |
| 2455631 | 2456743 | trpD | Rv2192c | 1113 | 2434161 | 2435273 | trpD | BCG_2208c | 1113 | 2414146 | 2415258 | trpD | BCGDan_2217 | 1113 | 2414395 | 2415438 | trpD | BCGDan_sapMKO_2219 | 1044 |
| 2456901 | 2457512 | ctaE | Rv2193 | 612 | 2435431 | 2436042 | ctaE | BCG_2209 | 612 | 2415416 | 2416027 | ctaE | BCGDan_2218 | 612 | 2415710 | 2416276 | ctaE | BCGDan_sapMKO_2220 | 567 |
| 2457553 | 2458395 | qcrC | Rv2194 | 843 | 2436083 | 2436925 | qcrC | BCG_2210 | 843 | 2416068 | 2416910 | qcrC | BCGDan_2219 | 843 | 2416317 | 2417159 | qcrC | BCGDan_sapMKO_2221 | 843 |
| 2458392 | 2459681 | qcrA | Rv2195 | 1290 | 2436922 | 2438211 | qcrA | BCG_2211 | 1290 | 2416907 | 2418196 | qcrA | BCGDan_2220 | 1290 | 2417234 | 2418445 | qcrA | BCGDan_sapMKO_2222 | 1212 |
| 2459678 | 2461327 | qcrB | Rv2196 | 1650 | 2438208 | 2439857 | qcrB | BCG_2212 | 1650 | 2418193 | 2419842 | qcrB | BCGDan_2221 | 1650 | 2418442 | 2420343 | qcrB | BCGDan_sapMKO_2223 | 1902 |
| 2461504 | 2462148 | Rv2197c | Rv2197c | 645 | 2440148 | 2440792 | BCG_2213c | BCG_2213c | 645 | 2420247 | 2420891 | BCGDan_2222 | BCGDan_2222 | 645 | 2420267 | 2420911 | BCGDan_sapMKO_2224 | BCGDan_sapMKO_2224 | 645 |
| 2462148 | 2463047 | mmpS3 | Rv2198c | 900 | 2440792 | 2441691 | mmpS3 | BCG_2214c | 900 | 2420891 | 2421790 | mmpS3 | BCGDan_2223 | 900 | 2420911 | 2421810 | mmpS3 | BCGDan_sapMKO_2225 | 900 |
| 2463233 | 2463652 | Rv2199c | Rv2199c | 420 | 2441877 | 2442296 | BCG_2215c | BCG_2215c | 420 | 2421976 | 2422395 | BCGDan_2224 | BCGDan_2224 | 420 | 2421996 | 2422415 | BCGDan_sapMKO_2226 | BCGDan_sapMKO_2226 | 420 |
| 2463660 | 2464751 | ctaC | Rv2200c | 1092 | 2442304 | 2443395 | ctaC | BCG_2216c | 1092 | 2422403 | 2423494 | ctaC | BCGDan_2225 | 1092 | 2422423 | 2423409 | ctaC | BCGDan_sapMKO_2227 | 987 |
| 2464997 | 2466955 | asnB | Rv2201 | 1959 | 2443641 | 2445599 | asnB | BCG_2217 | 1959 | 2423740 | 2425698 | asnB | BCGDan_2226 | 1959 | 2423760 | 2425718 | asnB | BCGDan_sapMKO_2228 | 1959 |
| 2467053 | 2468027 | adoK | Rv2202c | 975 | 2445697 | 2446671 | adoK | BCG_2218c | 975 | 2425796 | 2426770 | adoK | BCGDan_2227 | 975 | 2425816 | 2426790 | adoK | BCGDan_sapMKO_2229 | 975 |
| 2468231 | 2468923 | Rv2203 | Rv2203 | 693 | 2446875 | 2447567 | BCG_2219 | BCG_2219 | 693 | 2426974 | 2427666 | BCGDan_2228 | BCGDan_2228 | 693 | 2426994 | 2427686 | BCGDan_sapMKO_2230 | BCGDan_sapMKO_2230 | 693 |
| 2468931 | 2469287 | Rv2204c | Rv2204c | 357 | 2447575 | 2447931 | BCG_2220c | BCG_2220c | 357 | 2427674 | 2428030 | BCGDan_2229 | BCGDan_2229 | 357 | 2427694 | 2428050 | BCGDan_sapMKO_2231 | BCGDan_sapMKO_2231 | 357 |
| 2469387 | 2470463 | Rv2205c | Rv2205c | 1077 | 2448031 | 2449107 | BCG_2221c | BCG_2221c | 1077 | 2428130 | 2429206 | BCGDan_2230 | BCGDan_2230 | 1077 | 2428150 | 2429295 | BCGDan_sapMKO_2232 | BCGDan_sapMKO_2232 | 1146 |
| 2470622 | 2471332 | Rv2206 | Rv2206 | 711 | 2449266 | 2449976 | BCG_2222 | BCG_2222 | 711 | 2429365 | 2430075 | BCGDan_2231 | BCGDan_2231 | 711 | 2429385 | 2430095 | BCGDan_sapMKO_2233 | BCGDan_sapMKO_2233 | 711 |
| 2471411 | 2472496 | cobT | Rv2207 | 1086 | 2450055 | 2451140 | cobT | BCG_2223 | 1086 | 2430154 | 2431239 | cobT | BCGDan_2232 | 1086 | 2430174 | 2431259 | cobT | BCGDan_sapMKO_2234 | 1086 |
| 2472493 | 2473242 | cobS | Rv2208 | 750 | 2451137 | 2451886 | cobS | BCG_2224 | 750 | 2431236 | 2431985 | cobS | BCGDan_2233 | 750 | 2431259 | 2432005 | cobS | BCGDan_sapMKO_2235 | 747 |
| 2473400 | 2474938 | Rv2209 | Rv2209 | 1539 | 2452044 | 2453582 | BCG_2225 | BCG_2225 | 1539 | 2432143 | 2433681 | BCGDan_2234 | BCGDan_2234 | 1539 | 2432190 | 2433701 | BCGDan_sapMKO_2236 | BCGDan_sapMKO_2236 | 1512 |
| 2474864 | 2475970 | ilvE | Rv2210c | 1107 | 2453508 | 2454614 | ilvE | BCG_2226c | 1107 | 2433607 | 2434713 | ilvE | BCGDan_2235 | 1107 | 2433627 | 2434733 | ilvE | BCGDan_sapMKO_2237 | 1107 |
| 2476042 | 2477181 | gcvT | Rv2211c | 1140 | 2454686 | 2455825 | gcvT | BCG_2227c | 1140 | 2434785 | 2435924 | gcvT | BCGDan_2236 | 1140 | 2434805 | 2435908 | gcvT | BCGDan_sapMKO_2238 | 1104 |
| 2477190 | 2478326 | Rv2212 | Rv2212 | 1137 | 2455834 | 2456970 | BCG_2228 | BCG_2228 | 1137 | 2435933 | 2437069 | BCGDan_2237 | BCGDan_2237 | 1137 | 2435953 | 2437089 | BCGDan_sapMKO_2239 | BCGDan_sapMKO_2239 | 1137 |
| 2478338 | 2479885 | pepB | Rv2213 | 1548 | 2456982 | 2458529 | pepB | BCG_2229 | 1548 | 2437081 | 2438628 | pepB | BCGDan_2238 | 1548 | 2437101 | 2438648 | pepB | BCGDan_sapMKO_2240 | 1548 |
| 2479923 | 2481701 | ephD | Rv2214c | 1779 | 2458567 | 2460345 | ephD | BCG_2230c | 1779 | 2438666 | 2440444 | ephD | BCGDan_2239 | 1779 | 2438686 | 2440509 | ephD | BCGDan_sapMKO_2241 | 1824 |
| 2481965 | 2483626 | dlaT | Rv2215 | 1662 | 2460609 | 2462270 | dlaT | BCG_2231 | 1662 | 2440708 | 2442369 | dlaT | BCGDan_2240 | 1662 | 2440728 | 2442389 | dlaT | BCGDan_sapMKO_2242 | 1662 |
| 2483626 | 2484531 | Rv2216 | Rv2216 | 906 | 2462270 | 2463175 | BCG_2232 | BCG_2232 | 906 | 2442369 | 2443274 | BCGDan_2241 | BCGDan_2241 | 906 | 2442389 | 2443294 | BCGDan_sapMKO_2243 | BCGDan_sapMKO_2243 | 906 |
| 2484584 | 2485276 | lipB | Rv2217 | 693 | 2463228 | 2463920 | lipB | BCG_2233 | 693 | 2443327 | 2444019 | lipB | BCGDan_2242 | 693 | 2443347 | 2444039 | lipB | BCGDan_sapMKO_2244 | 693 |
| 2485273 | 2486208 | lipA | Rv2218 | 936 | 2463917 | 2464852 | lipA | BCG_2234 | 936 | 2444016 | 2444951 | lipA | BCGDan_2243 | 936 | 2444036 | 2444971 | lipA | BCGDan_sapMKO_2245 | 936 |
| 2486235 | 2486987 | Rv2219 | Rv2219 | 753 | 2464879 | 2465631 | BCG_2235 | BCG_2235 | 753 | 2444978 | 2445730 | BCGDan_2244 | BCGDan_2244 | 753 | 2444998 | 2445750 | BCGDan_sapMKO_2246 | BCGDan_sapMKO_2246 | 753 |
| 2486994 | 2487416 | Rv2219A | Rv2219A | 423 | 2465638 | 2466060 | BCG_2236c | BCG_2236c | 423 | 2445737 | 2446159 | BCGDan_2245 | BCGDan_2245 | 423 | 2445757 | 2446179 | BCGDan_sapMKO_2247 | BCGDan_sapMKO_2247 | 423 |
| 2487615 | 2489051 | glnA1 | Rv2220 | 1437 | 2466259 | 2467695 | glnA1 | BCG_2237 | 1437 | 2446358 | 2447794 | glnA1 | BCGDan_2246 | 1437 | 2446378 | 2447814 | glnA1 | BCGDan_sapMKO_2248 | 1437 |
| 2489369 | 2492353 | glnE | Rv2221c | 2985 | 2468013 | 2470997 | glnE | BCG_2238c | 2985 | 2448112 | 2451096 | glnE | BCGDan_2247 | 2985 | 2448132 | 2451116 | glnE | BCGDan_sapMKO_2249 | 2985 |
| 2492402 | 2493742 | glnA2 | Rv2222c | 1341 | 2471046 | 2472386 | glnA2 | BCG_2239c | 1341 | 2451145 | 2452485 | glnA2 | BCGDan_2248 | 1341 | 2451165 | 2452505 | glnA2 | BCGDan_sapMKO_2250 | 1341 |
| 2493837 | 2495399 | Rv2223c | Rv2223c | 1563 | 2472481 | 2474043 | BCG_2240c | BCG_2240c | 1563 | 2452580 | 2454142 | BCGDan_2249 | BCGDan_2249 | 1563 | 2452600 | 2454162 | BCGDan_sapMKO_2251 | BCGDan_sapMKO_2251 | 1563 |
| 2495461 | 2497023 | caeA | Rv2224c | 1563 | 2474105 | 2475667 | BCG_2241c | BCG_2241c | 1563 | 2454204 | 2455766 | BCGDan_2250 | BCGDan_2250 | 1563 | 2454224 | 2455780 | BCGDan_sapMKO_2252 | BCGDan_sapMKO_2252 | 1557 |
| 2497742 | 2498587 | panB | Rv2225 | 846 | 2476386 | 2477231 | panB | BCG_2242 | 846 | 2456485 | 2457330 | panB | BCGDan_2251 | 846 | 2456505 | 2457350 | panB | BCGDan_sapMKO_2253 | 846 |
| 2498832 | 2500373 | Rv2226 | Rv2226 | 1542 | 2477476 | 2479017 | BCG_2243 | BCG_2243 | 1542 | 2457575 | 2459116 | BCGDan_2252 | BCGDan_2252 | 1542 | 2457652 | 2459136 | BCGDan_sapMKO_2254 | BCGDan_sapMKO_2254 | 1485 |
| 2500931 | 2501632 | Rv2227 | Rv2227 | 702 | 2479575 | 2479895 | BCG_2244 | BCG_2244 | 321 | 2459674 | 2459994 | BCGDan_2254 | BCGDan_2254 | 321 | 2459694 | 2460014 | BCGDan_sapMKO_2256 | BCGDan_sapMKO_2256 | 321 |
| NA | NA | NA | NA | NA | 2479903 | 2480277 | BCG_2245 | BCG_2245 | 375 | 2460002 | 2460376 | BCGDan_2255 | BCGDan_2255 | 375 | 2460022 | 2460396 | BCGDan_sapMKO_2257 | BCGDan_sapMKO_2257 | 375 |
| 2501644 | 2502738 | Rv2228c | Rv2228c | 1095 | 2480289 | 2481383 | BCG_2246c | BCG_2246c | 1095 | 2460388 | 2461482 | BCGDan_2256 | BCGDan_2256 | 1095 | 2460408 | 2461502 | BCGDan_sapMKO_2258 | BCGDan_sapMKO_2258 | 1095 |
| 2502735 | 2503472 | Rv2229c | Rv2229c | 738 | 2481380 | 2482117 | BCG_2247c | BCG_2247c | 738 | 2461479 | 2462216 | BCGDan_2257 | BCGDan_2257 | 738 | 2461499 | 2462236 | BCGDan_sapMKO_2259 | BCGDan_sapMKO_2259 | 738 |

| | | | | | | | | | | | | | | | | | | | |
|---|---|---|---|---|---|---|---|---|---|---|---|---|---|---|---|---|---|---|---|
| 2503469 | 2504608 | Rv2230c | Rv2230c | 1140 | 2482114 | 2483253 | BCG_2248c | BCG_2248c | 1140 | 2462213 | 2463352 | BCGDan_2258 | BCGDan_2258 | 1140 | 2462233 | 2463366 | BCGDan_sapMKO_2260 | BCGDan_sapMKO_2260 | 1134 |
| 2504605 | 2505699 | cobC | Rv2231c | 1095 | 2483250 | 2484344 | cobC | BCG_2249c | 1095 | 2463349 | 2464443 | cobC | BCGDan_2259 | 1095 | 2463369 | 2464463 | cobC | BCGDan_sapMKO_2261 | 1095 |
| 2505736 | 2506161 | vapC16 | Rv2231A | 426 | NA | NA | NA | NA | NA | 2464480 | 2464905 | vapC16 | BCGDan_2260 | 426 | 2464503 | 2464925 | vapC16 | BCGDan_sapMKO_2262 | 423 |
| 2506207 | 2506383 | vapB16 | Rv2231B | 177 | NA | NA | NA | NA | NA | 2464951 | 2465127 | vapB16 | BCGDan_2261 | 177 | 2464971 | 2465147 | vapB16 | BCGDan_sapMKO_2263 | 177 |
| 2506278 | 2507153 | ptkA | Rv2232 | 876 | 2484923 | 2485798 | BCG_2250 | BCG_2250 | 876 | 2465022 | 2465897 | BCGDan_2262 | BCGDan_2262 | 876 | 2465159 | 2465917 | BCGDan_sapMKO_2264 | BCGDan_sapMKO_2264 | 759 |
| 2507146 | 2507637 | ptpA | Rv2234 | 492 | 2485791 | 2486282 | ptpA | BCG_2251 | 492 | 2465890 | 2466381 | ptpA | BCGDan_2263 | 492 | 2465967 | 2466401 | ptpA | BCGDan_sapMKO_2265 | 435 |
| 2507637 | 2508452 | Rv2235 | Rv2235 | 816 | 2486282 | 2487097 | BCG_2252 | BCG_2252 | 816 | 2466381 | 2467196 | BCGDan_2264 | BCGDan_2264 | 816 | 2466401 | 2467216 | BCGDan_sapMKO_2266 | BCGDan_sapMKO_2266 | 816 |
| 2508434 | 2509375 | cobD | Rv2236c | 942 | 2487079 | 2488020 | cobD | BCG_2253c | 942 | 2467178 | 2468119 | cobD | BCGDan_2265 | 942 | 2467198 | 2468157 | cobD | BCGDan_sapMKO_2267 | 960 |
| 2509489 | 2510256 | Rv2237 | Rv2237 | 768 | 2488134 | 2488901 | BCG_2254 | BCG_2254 | 768 | 2468233 | 2469000 | BCGDan_2266 | BCGDan_2266 | 768 | 2468205 | 2469020 | BCGDan_sapMKO_2268 | BCGDan_sapMKO_2268 | 816 |
| 2510715 | 2511176 | ahpE | Rv2238c | 462 | 2489360 | 2489821 | ahpE | BCG_2255c | 462 | 2469459 | 2469920 | ahpE | BCGDan_2268 | 462 | 2469479 | 2469940 | ahpE | BCGDan_sapMKO_2270 | 462 |
| 2511176 | 2511652 | Rv2239c | Rv2239c | 477 | 2489821 | 2490297 | BCG_2256c | BCG_2256c | 477 | 2469920 | 2470396 | BCGDan_2269 | BCGDan_2269 | 477 | 2469940 | 2470416 | BCGDan_sapMKO_2271 | BCGDan_sapMKO_2271 | 477 |
| 2511690 | 2512280 | Rv2240c | Rv2240c | 591 | 2490335 | 2491132 | BCG_2257c | BCG_2257c | 798 | 2470434 | 2471024 | BCGDan_2270 | BCGDan_2270 | 591 | 2470454 | 2471251 | BCGDan_sapMKO_2272 | BCGDan_sapMKO_2272 | 798 |
| 2512539 | 2515244 | aceE | Rv2241 | 2706 | 2491184 | 2493889 | aceE | BCG_2258 | 2706 | 2471283 | 2473988 | aceE | BCGDan_2271 | 2706 | 2471216 | 2474008 | aceE | BCGDan_sapMKO_2273 | 2793 |
| 2515304 | 2516548 | Rv2242 | Rv2242 | 1245 | 2493949 | 2495193 | BCG_2259 | BCG_2259 | 1245 | 2474048 | 2475292 | BCGDan_2272 | BCGDan_2272 | 1245 | 2474068 | 2475312 | BCGDan_sapMKO_2274 | BCGDan_sapMKO_2274 | 1245 |
| 2516787 | 2517695 | fabD | Rv2243 | 909 | 2495432 | 2496340 | fabD | BCG_2260 | 909 | 2475531 | 2476439 | fabD | BCGDan_2273 | 909 | 2475551 | 2476459 | fabD | BCGDan_sapMKO_2275 | 909 |
| 2517771 | 2518118 | acpM | Rv2244 | 348 | 2496416 | 2496763 | acpM | BCG_2261 | 348 | 2476515 | 2476862 | acpM | BCGDan_2274 | 348 | 2476535 | 2476882 | acpM | BCGDan_sapMKO_2276 | 348 |
| 2518115 | 2519365 | kasA | Rv2245 | 1251 | 2496760 | 2498010 | kasA | BCG_2262 | 1251 | 2476859 | 2478109 | kasA | BCGDan_2275 | 1251 | 2476921 | 2478129 | kasA | BCGDan_sapMKO_2277 | 1209 |
| 2519396 | 2520712 | kasB | Rv2246 | 1317 | 2498041 | 2499357 | kasB | BCG_2263 | 1317 | 2478140 | 2479456 | kasB | BCGDan_2276 | 1317 | 2478223 | 2479476 | kasB | BCGDan_sapMKO_2278 | 1254 |
| 2520743 | 2522164 | accD6 | Rv2247 | 1422 | 2499388 | 2500809 | accD6 | BCG_2264 | 1422 | 2479487 | 2480908 | accD6 | BCGDan_2277 | 1422 | 2479507 | 2480928 | accD6 | BCGDan_sapMKO_2279 | 1422 |
| NA | NA | NA | NA | NA | 2500917 | 2501321 | BCG_2265 | BCG_2265 | 405 | 2481016 | 2481420 | BCGDan_2278 | BCGDan_2278 | 405 | 2481036 | 2481440 | BCGDan_sapMKO_2280 | BCGDan_sapMKO_2280 | 405 |
| 2522360 | 2523175 | Rv2248 | Rv2248 | 816 | 2501029 | 2501733 | BCG_2266 | BCG_2266 | 705 | 2481128 | 2481832 | BCGDan_2279 | BCGDan_2279 | 705 | 2481148 | 2481852 | BCGDan_sapMKO_2281 | BCGDan_sapMKO_2281 | 705 |
| 2523241 | 2524791 | glpD1 | Rv2249c | 1551 | 2501802 | 2503352 | glpD1 | BCG_2267c | 1551 | 2481901 | 2483451 | glpD1 | BCGDan_2280 | 1551 | 2481921 | 2483471 | glpD1 | BCGDan_sapMKO_2282 | 1551 |
| 2524785 | 2525354 | Rv2250c | Rv2250c | 570 | 2503346 | 2503915 | BCG_2268c | BCG_2268c | 570 | 2483445 | 2484014 | BCGDan_2281 | BCGDan_2281 | 570 | 2483465 | 2484025 | BCGDan_sapMKO_2283 | BCGDan_sapMKO_2283 | 561 |
| 2525402 | 2525821 | Rv2250A | Rv2250A | 420 | 2503963 | 2505552 | BCG_2269 | BCG_2269 | 1590 | 2484062 | 2485651 | BCGDan_2282 | BCGDan_2282 | 1590 | 2484052 | 2485671 | BCGDan_sapMKO_2284 | BCGDan_sapMKO_2284 | 1620 |
| 2526989 | 2527918 | Rv2252 | Rv2252 | 930 | 2505549 | 2506478 | BCG_2270 | BCG_2270 | 930 | 2485648 | 2486577 | BCGDan_2283 | BCGDan_2283 | 930 | 2485668 | 2486597 | BCGDan_sapMKO_2285 | BCGDan_sapMKO_2285 | 930 |
| 2527984 | 2528487 | Rv2253 | Rv2253 | 504 | 2506544 | 2507047 | BCG_2271 | BCG_2271 | 504 | 2486643 | 2487146 | BCGDan_2284 | BCGDan_2284 | 504 | 2486687 | 2487166 | BCGDan_sapMKO_2286 | BCGDan_sapMKO_2286 | 480 |
| 2528520 | 2528975 | Rv2254c | Rv2254c | 456 | 2507080 | 2507535 | BCG_2272c | BCG_2272c | 456 | 2487179 | 2487634 | BCGDan_2285 | BCGDan_2285 | 456 | 2487199 | 2487654 | BCGDan_sapMKO_2287 | BCGDan_sapMKO_2287 | 456 |
| 2528980 | 2529174 | Rv2255c | Rv2255c | 195 | 2507540 | 2507734 | BCG_2273c | BCG_2273c | 195 | 2487639 | 2487833 | BCGDan_2286 | BCGDan_2286 | 195 | 2487659 | 2487853 | BCGDan_sapMKO_2288 | BCGDan_sapMKO_2288 | 195 |
| 2529341 | 2529874 | Rv2256c | Rv2256c | 534 | 2507901 | 2508434 | BCG_2274c | BCG_2274c | 534 | 2488000 | 2488533 | BCGDan_2287 | BCGDan_2287 | 534 | 2488020 | 2488553 | BCGDan_sapMKO_2289 | BCGDan_sapMKO_2289 | 534 |
| 2530004 | 2530822 | Rv2257c | Rv2257c | 819 | 2508564 | 2509382 | BCG_2275c | BCG_2275c | 819 | 2488663 | 2489481 | BCGDan_2288 | BCGDan_2288 | 819 | 2488683 | 2489501 | BCGDan_sapMKO_2290 | BCGDan_sapMKO_2290 | 819 |
| 2530836 | 2531897 | Rv2258c | Rv2258c | 1062 | 2509396 | 2510457 | BCG_2276c | BCG_2276c | 1062 | 2489495 | 2490556 | BCGDan_2289 | BCGDan_2289 | 1062 | 2489515 | 2490576 | BCGDan_sapMKO_2291 | BCGDan_sapMKO_2291 | 1062 |
| 2532245 | 2533330 | mscR | Rv2259 | 1086 | 2510699 | 2511784 | adhE2 | BCG_2277 | 1086 | 2490798 | 2491883 | adhE2 | BCGDan_2290 | 1086 | 2490818 | 2491903 | adhE2 | BCGDan_sapMKO_2292 | 1086 |
| 2533330 | 2533965 | Rv2260 | Rv2260 | 636 | 2511784 | 2512419 | BCG_2278 | BCG_2278 | 636 | 2491883 | 2492518 | BCGDan_2291 | BCGDan_2291 | 636 | 2491903 | 2492538 | BCGDan_sapMKO_2293 | BCGDan_sapMKO_2293 | 636 |
| 2534042 | 2534464 | Rv2261c | Rv2261c | 423 | 2512496 | 2514004 | BCG_2279c | BCG_2279c | 1509 | 2492595 | 2494103 | BCGDan_2292 | BCGDan_2292 | 1509 | 2492615 | 2494144 | BCGDan_sapMKO_2294 | BCGDan_sapMKO_2294 | 1530 |
| 2535641 | 2536594 | Rv2263 | Rv2263 | 954 | 2514093 | 2515046 | BCG_2280 | BCG_2280 | 954 | 2494192 | 2495145 | BCGDan_2293 | BCGDan_2293 | 954 | 2494212 | 2495165 | BCGDan_sapMKO_2295 | BCGDan_sapMKO_2295 | 954 |
| 2536572 | 2538350 | Rv2264c | Rv2264c | 1779 | 2515024 | 2516802 | BCG_2281c | BCG_2281c | 1779 | 2495123 | 2496901 | BCGDan_2294 | BCGDan_2294 | 1779 | 2495143 | 2496921 | BCGDan_sapMKO_2296 | BCGDan_sapMKO_2296 | 1779 |
| 2538700 | 2539929 | Rv2265 | Rv2265 | 1230 | 2517152 | 2518381 | BCG_2282 | BCG_2282 | 1230 | 2497251 | 2498480 | BCGDan_2295 | BCGDan_2295 | 1230 | 2497271 | 2498500 | BCGDan_sapMKO_2297 | BCGDan_sapMKO_2297 | 1230 |
| 2540104 | 2541390 | cyp124 | Rv2266 | 1287 | 2518556 | 2519842 | cyp124 | BCG_2283 | 1287 | 2498655 | 2499941 | cyp124 | BCGDan_2296 | 1287 | 2498675 | 2499961 | cyp124 | BCGDan_sapMKO_2298 | 1287 |
| 2541644 | 2542810 | Rv2267c | Rv2267c | 1167 | 2520096 | 2521262 | BCG_2284c | BCG_2284c | 1167 | 2500195 | 2501361 | BCGDan_2297 | BCGDan_2297 | 1167 | 2500215 | 2501294 | BCGDan_sapMKO_2299 | BCGDan_sapMKO_2299 | 1080 |
| 2542807 | 2544276 | cyp128 | Rv2268c | 1470 | 2521259 | 2522728 | cyp128 | BCG_2285c | 1470 | 2501358 | 2502827 | cyp128 | BCGDan_2298 | 1470 | 2501378 | 2502847 | cyp128 | BCGDan_sapMKO_2300 | 1470 |
| 2544289 | 2544621 | Rv2269c | Rv2269c | 333 | 2522741 | 2523073 | BCG_2286c | BCG_2286c | 333 | 2502840 | 2503172 | BCGDan_2299 | BCGDan_2299 | 333 | 2502860 | 2503192 | BCGDan_sapMKO_2301 | BCGDan_sapMKO_2301 | 333 |
| 2544698 | 2545225 | lppN | Rv2270 | 528 | 2523150 | 2523677 | lppN | BCG_2287 | 528 | 2503249 | 2503776 | lppN | BCGDan_2300 | 528 | 2503290 | 2503796 | lppN | BCGDan_sapMKO_2302 | 507 |
| 2545332 | 2545631 | Rv2271 | Rv2271 | 300 | 2523784 | 2524083 | BCG_2288 | BCG_2288 | 300 | 2503883 | 2504182 | BCGDan_2301 | BCGDan_2301 | 300 | 2503903 | 2504202 | BCGDan_sapMKO_2303 | BCGDan_sapMKO_2303 | 300 |
| 2545737 | 2546105 | Rv2272 | Rv2272 | 369 | 2524189 | 2524557 | BCG_2289 | BCG_2289 | 369 | 2504288 | 2504656 | BCGDan_2302 | BCGDan_2302 | 369 | 2504308 | 2504676 | BCGDan_sapMKO_2304 | BCGDan_sapMKO_2304 | 369 |
| 2546102 | 2546431 | Rv2273 | Rv2273 | 330 | 2524554 | 2524883 | BCG_2290 | BCG_2290 | 330 | 2504653 | 2504982 | BCGDan_2303 | BCGDan_2303 | 330 | 2504673 | 2505002 | BCGDan_sapMKO_2305 | BCGDan_sapMKO_2305 | 330 |
| 2546488 | 2546805 | mazF8 | Rv2274c | 318 | 2524940 | 2525257 | BCG_2291c | BCG_2291c | 318 | 2505039 | 2505356 | BCGDan_2304 | BCGDan_2304 | 318 | 2505059 | 2505376 | BCGDan_sapMKO_2306 | BCGDan_sapMKO_2306 | 318 |
| 2546883 | 2547752 | Rv2275 | Rv2275 | 870 | 2525335 | 2526204 | BCG_2292 | BCG_2292 | 870 | 2505434 | 2506303 | BCGDan_2305 | BCGDan_2305 | 870 | 2505454 | 2506323 | BCGDan_sapMKO_2307 | BCGDan_sapMKO_2307 | 870 |
| 2547749 | 2548939 | cyp121 | Rv2276 | 1191 | 2526201 | 2527391 | cyp121 | BCG_2293 | 1191 | 2506300 | 2507490 | cyp121 | BCGDan_2306 | 1191 | 2506320 | 2507510 | cyp121 | BCGDan_sapMKO_2308 | 1191 |
| 2549124 | 2550029 | Rv2277c | Rv2277c | 906 | 2527576 | 2528475 | BCG_2294c | BCG_2294c | 900 | 2507675 | 2508574 | BCGDan_2307 | BCGDan_2307 | 900 | 2507695 | 2508594 | BCGDan_sapMKO_2309 | BCGDan_sapMKO_2309 | 900 |
| 2551560 | 2552939 | Rv2280 | Rv2280 | 1380 | 2528654 | 2530033 | BCG_2295 | BCG_2295 | 1380 | 2508753 | 2510132 | BCGDan_2308 | BCGDan_2308 | 1380 | 2508782 | 2510152 | BCGDan_sapMKO_2310 | BCGDan_sapMKO_2310 | 1371 |
| 2553173 | 2554831 | pitB | Rv2281 | 1659 | 2530267 | 2531925 | pitB | BCG_2296 | 1659 | 2510366 | 2512024 | pitB | BCGDan_2309 | 1659 | 2510386 | 2512044 | pitB | BCGDan_sapMKO_2311 | 1659 |
| 2554938 | 2555876 | Rv2282c | Rv2282c | 939 | 2532032 | 2532970 | BCG_2297c | BCG_2297c | 939 | 2512131 | 2513051 | BCGDan_2310 | BCGDan_2310 | 921 | 2512151 | 2513071 | BCGDan_sapMKO_2312 | BCGDan_sapMKO_2312 | 921 |
| 2555941 | 2556135 | Rv2283 | Rv2283 | 195 | 2533035 | 2533229 | BCG_2298 | BCG_2298 | 195 | 2513134 | 2513328 | BCGDan_2311 | BCGDan_2311 | 195 | 2513154 | 2513348 | BCGDan_sapMKO_2313 | BCGDan_sapMKO_2313 | 195 |
| 2556145 | 2557440 | lipM | Rv2284 | 1296 | 2533239 | 2534534 | lipM | BCG_2299 | 1296 | 2513338 | 2514633 | lipM | BCGDan_2312 | 1296 | 2513358 | 2514653 | lipM | BCGDan_sapMKO_2314 | 1296 |
| 2557473 | 2558810 | Rv2285 | Rv2285 | 1338 | 2534567 | 2535904 | BCG_2300 | BCG_2300 | 1338 | 2514666 | 2516003 | BCGDan_2313 | BCGDan_2313 | 1338 | 2514686 | 2516023 | BCGDan_sapMKO_2315 | BCGDan_sapMKO_2315 | 1338 |
| 2558877 | 2559569 | Rv2286c | Rv2286c | 693 | 2535971 | 2536234 | BCG_2301c | BCG_2301c | 264 | 2516070 | 2516333 | BCGDan_2314 | BCGDan_2314 | 264 | 2516090 | 2516353 | BCGDan_sapMKO_2316 | BCGDan_sapMKO_2316 | 264 |
| NA | NA | NA | NA | NA | 2536262 | 2536663 | BCG_2302c | BCG_2302c | 402 | 2516361 | 2516762 | BCGDan_2315 | BCGDan_2315 | 402 | 2516381 | 2516782 | BCGDan_sapMKO_2317 | BCGDan_sapMKO_2317 | 402 |
| 2559703 | 2561331 | yjcE | Rv2287 | 1629 | 2536797 | 2538425 | yjcE | BCG_2303 | 1629 | 2516896 | 2518524 | yjcE | BCGDan_2316 | 1629 | 2516916 | 2518544 | yjcE | BCGDan_sapMKO_2318 | 1629 |
| 2561328 | 2561705 | Rv2288 | Rv2288 | 378 | 2538422 | 2538799 | BCG_2304 | BCG_2304 | 378 | 2518521 | 2518898 | BCGDan_2317 | BCGDan_2317 | 378 | 2518541 | 2518918 | BCGDan_sapMKO_2319 | BCGDan_sapMKO_2319 | 378 |
| 2561675 | 2562457 | cdh | Rv2289 | 783 | 2538769 | 2539551 | cdh | BCG_2305 | 783 | 2518868 | 2519650 | cdh | BCGDan_2318 | 783 | 2518888 | 2519670 | cdh | BCGDan_sapMKO_2320 | 783 |
| 2562599 | 2563114 | lppO | Rv2290 | 516 | 2539693 | 2539848 | lppOa | BCG_2306 | 156 | 2519792 | 2519947 | lppOa | BCGDan_2319 | 156 | 2519812 | 2519967 | lppOa | BCGDan_sapMKO_2321 | 156 |
| NA | NA | NA | NA | NA | 2539842 | 2540207 | lppOb | BCG_2307 | 366 | 2519941 | 2520306 | lppOb | BCGDan_2320 | 366 | 2519961 | 2520326 | lppOb | BCGDan_sapMKO_2322 | 366 |
| 2563174 | 2564028 | sseB | Rv2291 | 855 | 2540267 | 2541121 | sseB | BCG_2308 | 855 | 2520366 | 2521220 | sseB | BCGDan_2321 | 855 | 2520386 | 2521240 | sseB | BCGDan_sapMKO_2323 | 855 |
| 2564029 | 2564253 | Rv2292c | Rv2292c | 225 | 2541122 | 2542126 | BCG_2309c | BCG_2309c | 1005 | 2521221 | 2522225 | BCGDan_2322 | BCGDan_2322 | 1005 | 2521241 | 2522215 | BCGDan_sapMKO_2324 | BCGDan_sapMKO_2324 | 975 |
| 2565327 | 2566550 | Rv2294 | Rv2294 | 1224 | 2542421 | 2543644 | BCG_2310 | BCG_2310 | 1224 | 2522520 | 2523743 | BCGDan_2323 | BCGDan_2323 | 1224 | 2522600 | 2523763 | BCGDan_sapMKO_2325 | BCGDan_sapMKO_2325 | 1164 |
| 2566772 | 2567410 | Rv2295 | Rv2295 | 639 | 2543867 | 2544505 | BCG_2311 | BCG_2311 | 639 | 2523966 | 2524604 | BCGDan_2324 | BCGDan_2324 | 639 | 2523956 | 2524624 | BCGDan_sapMKO_2326 | BCGDan_sapMKO_2326 | 669 |
| 2567504 | 2568406 | Rv2296 | Rv2296 | 903 | 2544599 | 2545501 | BCG_2312 | BCG_2312 | 903 | 2524698 | 2525600 | BCGDan_2325 | BCGDan_2325 | 903 | 2524670 | 2525620 | BCGDan_sapMKO_2327 | BCGDan_sapMKO_2327 | 951 |
| 2568438 | 2568890 | Rv2297 | Rv2297 | 453 | 2545533 | 2545985 | BCG_2313 | BCG_2313 | 453 | 2525632 | 2526084 | BCGDan_2326 | BCGDan_2326 | 453 | 2525652 | 2526104 | BCGDan_sapMKO_2328 | BCGDan_sapMKO_2328 | 453 |
| 2569082 | 2570053 | Rv2298 | Rv2298 | 972 | 2546177 | 2547148 | BCG_2314 | BCG_2314 | 972 | 2526276 | 2527247 | BCGDan_2327 | BCGDan_2327 | 972 | 2526200 | 2527267 | BCGDan_sapMKO_2329 | BCGDan_sapMKO_2329 | 1068 |
| 2570059 | 2572002 | htpG | Rv2299c | 1944 | 2547154 | 2549097 | htpG | BCG_2315c | 1944 | 2527253 | 2529196 | htpG | BCGDan_2328 | 1944 | 2527273 | 2529216 | htpG | BCGDan_sapMKO_2330 | 1944 |
| 2572076 | 2573008 | Rv2300c | Rv2300c | 933 | 2549171 | 2550103 | BCG_2316c | BCG_2316c | 933 | 2529270 | 2530202 | BCGDan_2329 | BCGDan_2329 | 933 | 2529290 | 2530108 | BCGDan_sapMKO_2331 | BCGDan_sapMKO_2331 | 819 |
| 2573015 | 2573707 | cut2 | Rv2301 | 693 | 2550110 | 2550802 | cut2 | BCG_2317 | 693 | 2530209 | 2530901 | cut2 | BCGDan_2330 | 693 | 2530262 | 2530921 | cut2 | BCGDan_sapMKO_2332 | 660 |
| 2573813 | 2574055 | Rv2302 | Rv2302 | 243 | 2550908 | 2551150 | BCG_2318 | BCG_2318 | 243 | 2531007 | 2531249 | BCGDan_2331 | BCGDan_2331 | 243 | 2531027 | 2531269 | BCGDan_sapMKO_2333 | BCGDan_sapMKO_2333 | 243 |
| 2574096 | 2575019 | Rv2303c | Rv2303c | 924 | 2551191 | 2552114 | BCG_2319c | BCG_2319c | 924 | 2531290 | 2532213 | BCGDan_2332 | BCGDan_2332 | 924 | 2531310 | 2532233 | BCGDan_sapMKO_2334 | BCGDan_sapMKO_2334 | 924 |
| 2575016 | 2575225 | Rv2304c | Rv2304c | 210 | 2552111 | 2552320 | BCG_2320c | BCG_2320c | 210 | 2532210 | 2532419 | BCGDan_2333 | BCGDan_2333 | 210 | 2532230 | 2532439 | BCGDan_sapMKO_2335 | BCGDan_sapMKO_2335 | 210 |
| 2575809 | 2577098 | Rv2305 | Rv2305 | 1290 | 2552904 | 2554193 | BCG_2321 | BCG_2321 | 1290 | 2533003 | 2534292 | BCGDan_2334 | BCGDan_2334 | 1290 | 2533023 | 2534312 | BCGDan_sapMKO_2336 | BCGDan_sapMKO_2336 | 1290 |
| 2577108 | 2577701 | Rv2306A | Rv2306A | 594 | 2554203 | 2554796 | BCG_2322 | BCG_2322 | 594 | 2534302 | 2534895 | BCGDan_2335 | BCGDan_2335 | 594 | 2534322 | 2534915 | BCGDan_sapMKO_2337 | BCGDan_sapMKO_2337 | 594 |
| 2577488 | 2577922 | Rv2306B | Rv2306B | 435 | 2554583 | 2555017 | BCG_2323 | BCG_2323 | 435 | 2534682 | 2535116 | BCGDan_2336 | BCGDan_2336 | 435 | 2534702 | 2535136 | BCGDan_sapMKO_2338 | BCGDan_sapMKO_2338 | 435 |
| 2577851 | 2578696 | Rv2307c | Rv2307c | 846 | 2554946 | 2555791 | BCG_2324c | BCG_2324c | 846 | 2535045 | 2535890 | BCGDan_2337 | BCGDan_2337 | 846 | 2535065 | 2535910 | BCGDan_sapMKO_2339 | BCGDan_sapMKO_2339 | 846 |
| 2579228 | 2579419 | Rv2307A | Rv2307A | 192 | 2556323 | 2556514 | BCG_2325c | BCG_2325c | 192 | 2536422 | 2536613 | BCGDan_2338 | BCGDan_2338 | 192 | 2536442 | 2536633 | BCGDan_sapMKO_2340 | BCGDan_sapMKO_2340 | 192 |
| 2579504 | 2579935 | Rv2307B | Rv2307B | 432 | 2556599 | 2557030 | BCG_2326c | BCG_2326c | 432 | 2536698 | 2537129 | BCGDan_2339 | BCGDan_2339 | 432 | 2536718 | 2537149 | BCGDan_sapMKO_2341 | BCGDan_sapMKO_2341 | 432 |
| 2580028 | 2580210 | Rv2307D | Rv2307D | 183 | 2557123 | 2557305 | BCG_2327c | BCG_2327c | 183 | 2537222 | 2537404 | BCGDan_2340 | BCGDan_2340 | 183 | 2537242 | 2537424 | BCGDan_sapMKO_2342 | BCGDan_sapMKO_2342 | 183 |
| 2580419 | 2581135 | Rv2308 | Rv2308 | 717 | 2557514 | 2558230 | BCG_2328 | BCG_2328 | 717 | 2537613 | 2538329 | BCGDan_2341 | BCGDan_2341 | 717 | 2537633 | 2538349 | BCGDan_sapMKO_2343 | BCGDan_sapMKO_2343 | 717 |
| 2581843 | 2582298 | Rv2309c | Rv2309c | 456 | 2558938 | 2559393 | BCG_2329c | BCG_2329c | 456 | 2539037 | 2539492 | BCGDan_2343 | BCGDan_2343 | 456 | 2539057 | 2539512 | BCGDan_sapMKO_2345 | BCGDan_sapMKO_2345 | 456 |
| 2583045 | 2583332 | Rv2309A | Rv2309A | 288 | 2560140 | 2560427 | BCG_2330 | BCG_2330 | 288 | 2540239 | 2540526 | BCGDan_2344 | BCGDan_2344 | 288 | 2540259 | 2540546 | BCGDan_sapMKO_2346 | BCGDan_sapMKO_2346 | 288 |
| 2583435 | 2583779 | Rv2310 | Rv2310 | 345 | 2560530 | 2560874 | BCG_2331 | BCG_2331 | 345 | 2540629 | 2540973 | BCGDan_2345 | BCGDan_2345 | 345 | 2540649 | 2540993 | BCGDan_sapMKO_2347 | BCGDan_sapMKO_2347 | 345 |

| | | | | | | | | | | | | | | | | | | | |
|---|---|---|---|---|---|---|---|---|---|---|---|---|---|---|---|---|---|---|---|
| 2583884 | 2584408 | Rv2311 | Rv2311 | 525 | 2560979 | 2561503 | BCG_2332 | BCG_2332 | 525 | 2541078 | 2541602 | BCGDan_2346 | BCGDan_2346 | 525 | 2541026 | 2541622 | BCGDan_sapMKO_2348 | BCGDan_sapMKO_2348 | 597 |
| 2584486 | 2584755 | Rv2312 | Rv2312 | 270 | 2561581 | 2561850 | BCG_2333 | BCG_2333 | 270 | 2541680 | 2541949 | BCGDan_2347 | BCGDan_2347 | 270 | 2541700 | 2541969 | BCGDan_sapMKO_2349 | BCGDan_sapMKO_2349 | 270 |
| 2585052 | 2585906 | Rv2313c | Rv2313c | 855 | 2562147 | 2563001 | BCG_2334c | BCG_2334c | 855 | 2542246 | 2543100 | BCGDan_2348 | BCGDan_2348 | 855 | 2542266 | 2543120 | BCGDan_sapMKO_2350 | BCGDan_sapMKO_2350 | 855 |
| 2585917 | 2587290 | Rv2314c | Rv2314c | 1374 | 2563012 | 2564385 | BCG_2335c | BCG_2335c | 1374 | 2543111 | 2544484 | BCGDan_2349 | BCGDan_2349 | 1374 | 2543131 | 2544504 | BCGDan_sapMKO_2351 | BCGDan_sapMKO_2351 | 1374 |
| 2587287 | 2588804 | Rv2315c | Rv2315c | 1518 | 2564382 | 2565899 | BCG_2336c | BCG_2336c | 1518 | 2544481 | 2545998 | BCGDan_2350 | BCGDan_2350 | 1518 | 2544501 | 2546018 | BCGDan_sapMKO_2352 | BCGDan_sapMKO_2352 | 1518 |
| 2588838 | 2589710 | uspA | Rv2316 | 873 | 2565933 | 2566805 | uspA | BCG_2337 | 873 | 2546032 | 2546904 | uspA | BCGDan_2351 | 873 | 2546052 | 2546924 | uspA | BCGDan_sapMKO_2353 | 873 |
| 2589697 | 2590521 | uspB | Rv2317 | 825 | 2566792 | 2567616 | uspB | BCG_2338 | 825 | 2546891 | 2547715 | uspB | BCGDan_2352 | 825 | 2546911 | 2547735 | uspB | BCGDan_sapMKO_2354 | 825 |
| 2590518 | 2591840 | uspC | Rv2318 | 1323 | 2567613 | 2568935 | uspC | BCG_2339 | 1323 | 2547712 | 2549034 | uspC | BCGDan_2353 | 1323 | 2547732 | 2549054 | uspC | BCGDan_sapMKO_2355 | 1323 |
| 2591848 | 2592726 | Rv2319c | Rv2319c | 879 | 2568943 | 2569821 | BCG_2340c | BCG_2340c | 879 | 2549042 | 2549920 | BCGDan_2354 | BCGDan_2354 | 879 | 2549062 | 2549940 | BCGDan_sapMKO_2356 | BCGDan_sapMKO_2356 | 879 |
| 2592723 | 2594153 | rocE | Rv2320c | 1431 | 2569818 | 2571248 | rocE | BCG_2341c | 1431 | 2549917 | 2551347 | rocE | BCGDan_2355 | 1431 | 2549937 | 2551334 | rocE | BCGDan_sapMKO_2357 | 1398 |
| 2594154 | 2594699 | rocD2 | Rv2321c | 546 | 2571249 | 2571794 | rocD2 | BCG_2342c | 546 | 2551348 | 2551893 | rocD2 | BCGDan_2356 | 546 | 2551368 | 2551919 | rocD2 | BCGDan_sapMKO_2358 | 552 |
| 2594699 | 2595364 | rocD1 | Rv2322c | 666 | 2571794 | 2572459 | rocD1 | BCG_2343c | 666 | 2551893 | 2552558 | rocD1 | BCGDan_2357 | 666 | 2551913 | 2552578 | rocD1 | BCGDan_sapMKO_2359 | 666 |
| 2595361 | 2596269 | Rv2323c | Rv2323c | 909 | 2572456 | 2573364 | BCG_2344c | BCG_2344c | 909 | 2552555 | 2553463 | BCGDan_2358 | BCGDan_2358 | 909 | 2552575 | 2553348 | BCGDan_sapMKO_2360 | BCGDan_sapMKO_2360 | 774 |
| 2596334 | 2596780 | Rv2324 | Rv2324 | 447 | 2573429 | 2573875 | BCG_2345 | BCG_2345 | 447 | 2553528 | 2553974 | BCGDan_2359 | BCGDan_2359 | 447 | 2553548 | 2553994 | BCGDan_sapMKO_2361 | BCGDan_sapMKO_2361 | 447 |
| 2597009 | 2597857 | Rv2325c | Rv2325c | 849 | 2574104 | 2574952 | BCG_2346c | BCG_2346c | 849 | 2554203 | 2555051 | BCGDan_2360 | BCGDan_2360 | 849 | 2554223 | 2555014 | BCGDan_sapMKO_2362 | BCGDan_sapMKO_2362 | 792 |
| 2597854 | 2599947 | Rv2326c | Rv2326c | 2094 | 2574949 | 2577042 | BCG_2347c | BCG_2347c | 2094 | 2555048 | 2557141 | BCGDan_2361 | BCGDan_2361 | 2094 | 2555068 | 2557038 | BCGDan_sapMKO_2363 | BCGDan_sapMKO_2363 | 1971 |
| 2599988 | 2600479 | Rv2327 | Rv2327 | 492 | 2577083 | 2577574 | BCG_2348 | BCG_2348 | 492 | 2557182 | 2557673 | BCGDan_2362 | BCGDan_2362 | 492 | 2557202 | 2557693 | BCGDan_sapMKO_2364 | BCGDan_sapMKO_2364 | 492 |
| 2600731 | 2601879 | PE23 | Rv2328 | 1149 | 2577826 | 2578974 | PE23 | BCG_2349 | 1149 | 2557925 | 2559073 | PE23 | BCGDan_2363 | 1149 | 2557945 | 2559093 | PE23 | BCGDan_sapMKO_2365 | 1149 |
| 2601914 | 2603461 | narK1 | Rv2329c | 1548 | 2579009 | 2580556 | narK1 | BCG_2350c | 1548 | 2559108 | 2560655 | narK1 | BCGDan_2364 | 1548 | 2559128 | 2560675 | narK1 | BCGDan_sapMKO_2366 | 1548 |
| 2603695 | 2604222 | lppP | Rv2330c | 528 | 2580790 | 2581317 | lppP | BCG_2351c | 528 | 2560889 | 2561416 | lppP | BCGDan_2365 | 528 | 2560909 | 2561397 | lppP | BCGDan_sapMKO_2367 | 489 |
| 2604297 | 2604683 | Rv2331 | Rv2331 | 387 | 2581392 | 2581778 | BCG_2352 | BCG_2352 | 387 | 2561491 | 2561877 | BCGDan_2366 | BCGDan_2366 | 387 | 2561511 | 2561897 | BCGDan_sapMKO_2368 | BCGDan_sapMKO_2368 | 387 |
| 2604740 | 2605078 | Rv2331A | Rv2331A | 339 | 2581835 | 2582173 | BCG_2353 | BCG_2353 | 339 | 2561934 | 2562272 | BCGDan_2367 | BCGDan_2367 | 339 | 2561954 | 2562292 | BCGDan_sapMKO_2369 | BCGDan_sapMKO_2369 | 339 |
| 2605108 | 2606754 | mez | Rv2332 | 1647 | 2582203 | 2583849 | mez | BCG_2354 | 1647 | 2562302 | 2563948 | mez | BCGDan_2368 | 1647 | 2562322 | 2563968 | mez | BCGDan_sapMKO_2370 | 1647 |
| 2606708 | 2608321 | stp | Rv2333c | 1614 | 2583846 | 2585417 | BCG_2355c | BCG_2355c | 1572 | 2563945 | 2565516 | BCGDan_2369 | BCGDan_2369 | 1572 | 2563965 | 2565536 | BCGDan_sapMKO_2371 | BCGDan_sapMKO_2371 | 1572 |
| 2608796 | 2609728 | cysK1 | Rv2334 | 933 | 2585892 | 2586824 | cysK1 | BCG_2356 | 933 | 2565991 | 2566923 | cysK1 | BCGDan_2370 | 933 | 2566011 | 2566943 | cysK1 | BCGDan_sapMKO_2372 | 933 |
| 2609732 | 2610421 | cysE | Rv2335 | 690 | 2586828 | 2587517 | BCG_2357 | BCG_2357 | 690 | 2566927 | 2567616 | BCGDan_2371 | BCGDan_2371 | 690 | 2566947 | 2567636 | BCGDan_sapMKO_2373 | BCGDan_sapMKO_2373 | 690 |
| 2610837 | 2611805 | Rv2336 | Rv2336 | 969 | 2587933 | 2588901 | BCG_2358 | BCG_2358 | 969 | 2568032 | 2569000 | BCGDan_2372 | BCGDan_2372 | 969 | 2568052 | 2569020 | BCGDan_sapMKO_2374 | BCGDan_sapMKO_2374 | 969 |
| 2611869 | 2612987 | Rv2337c | Rv2337c | 1119 | 2588965 | 2590083 | BCG_2359c | BCG_2359c | 1119 | 2569064 | 2570182 | BCGDan_2373 | BCGDan_2373 | 1119 | 2569084 | 2570202 | BCGDan_sapMKO_2375 | BCGDan_sapMKO_2375 | 1119 |
| 2613107 | 2614063 | moeW | Rv2338c | 957 | 2590203 | 2591159 | moeW | BCG_2360c | 957 | 2570302 | 2571258 | moeW | BCGDan_2374 | 957 | 2570322 | 2571278 | moeW | BCGDan_sapMKO_2376 | 957 |
| 2614693 | 2617581 | mmpL9 | Rv2339 | 2889 | 2591790 | 2594270 | mmpL9a | BCG_2361 | 2481 | 2571889 | 2574369 | mmpL9a | BCGDan_2375 | 2481 | 2571909 | 2574389 | mmpL9a | BCGDan_sapMKO_2377 | 2481 |
| NA | NA | NA | NA | NA | 2594283 | 2594678 | mmpL9b | BCG_2362 | 396 | 2574382 | 2574777 | mmpL9b | BCGDan_2376 | 396 | 2574402 | 2574797 | mmpL9b | BCGDan_sapMKO_2378 | 396 |
| 2617667 | 2618908 | PE_PGRS39 | Rv2340c | 1242 | 2594764 | 2596005 | PE_PGRS39 | BCG_2363c | 1242 | 2574863 | 2576104 | PE_PGRS39 | BCGDan_2377 | 1242 | 2574883 | 2576124 | PE_PGRS39 | BCGDan_sapMKO_2379 | 1242 |
| 2619597 | 2620016 | lppQ | Rv2341 | 420 | 2596694 | 2597113 | lppQ | BCG_2364 | 420 | 2576793 | 2577212 | lppQ | BCGDan_2379 | 420 | 2576798 | 2577232 | lppQ | BCGDan_sapMKO_2381 | 435 |
| 2620272 | 2620529 | Rv2342 | Rv2342 | 258 | 2597369 | 2597626 | BCG_2365 | BCG_2365 | 258 | 2577468 | 2577725 | BCGDan_2380 | BCGDan_2380 | 258 | 2577506 | 2577745 | BCGDan_sapMKO_2382 | BCGDan_sapMKO_2382 | 240 |
| 2620533 | 2622452 | dnaG | Rv2343c | 1920 | 2597630 | 2599549 | dnaG | BCG_2366c | 1920 | 2577729 | 2579648 | dnaG | BCGDan_2381 | 1920 | 2577749 | 2579668 | dnaG | BCGDan_sapMKO_2383 | 1920 |
| 2622457 | 2623752 | dgt | Rv2344c | 1296 | 2599554 | 2600849 | dgt | BCG_2367c | 1296 | 2579653 | 2580948 | dgt | BCGDan_2382 | 1296 | 2579673 | 2580968 | dgt | BCGDan_sapMKO_2384 | 1296 |
| 2623821 | 2625803 | Rv2345 | Rv2345 | 1983 | 2600918 | 2602900 | BCG_2368 | BCG_2368 | 1983 | 2581017 | 2582999 | BCGDan_2383 | BCGDan_2383 | 1983 | 2581037 | 2583019 | BCGDan_sapMKO_2385 | BCGDan_sapMKO_2385 | 1983 |
| 2625888 | 2626172 | esxO | Rv2346c | 285 | 2602985 | 2603167 | esxO | BCG_2369c | 183 | 2583084 | 2583266 | esxO | BCGDan_2384 | 183 | 2583104 | 2583286 | esxO | BCGDan_sapMKO_2386 | 183 |
| NA | NA | NA | NA | NA | 2603151 | 2604557 | PPE71 | BCG_2370c | 1407 | 2583250 | 2584656 | PPE71 | BCGDan_2385 | 1407 | 2583270 | 2584676 | PPE71 | BCGDan_sapMKO_2387 | 1407 |
| 2639673 | 2641064 | glyS | Rv2357c | 1392 | 2604695 | 2606086 | glyS | BCG_2371c | 1392 | 2584794 | 2586185 | glyS | BCGDan_2386 | 1392 | 2584814 | 2586268 | glyS | BCGDan_sapMKO_2388 | 1455 |
| 2641246 | 2641653 | smtB | Rv2358 | 408 | 2606268 | 2606675 | BCG_2372 | BCG_2372 | 408 | 2586367 | 2586774 | BCGDan_2387 | BCGDan_2387 | 408 | 2586429 | 2586794 | BCGDan_sapMKO_2389 | BCGDan_sapMKO_2389 | 366 |
| 2641650 | 2642042 | zur | Rv2359 | 393 | 2606672 | 2607064 | furB | BCG_2373 | 393 | 2586771 | 2587163 | furB | BCGDan_2388 | 393 | 2586791 | 2587183 | furB | BCGDan_sapMKO_2390 | 393 |
| 2642150 | 2642578 | Rv2360c | Rv2360c | 429 | 2607172 | 2607600 | BCG_2374c | BCG_2374c | 429 | 2587271 | 2587699 | BCGDan_2389 | BCGDan_2389 | 429 | 2587291 | 2587719 | BCGDan_sapMKO_2391 | BCGDan_sapMKO_2391 | 429 |
| 2642578 | 2643468 | Rv2361c | Rv2361c | 891 | 2607600 | 2608490 | BCG_2375c | BCG_2375c | 891 | 2587699 | 2588589 | BCGDan_2390 | BCGDan_2390 | 891 | 2587719 | 2588609 | BCGDan_sapMKO_2392 | BCGDan_sapMKO_2392 | 891 |
| 2643461 | 2644258 | recO | Rv2362c | 798 | 2608483 | 2609280 | BCG_2376c | BCG_2376c | 798 | 2588582 | 2589379 | BCGDan_2391 | BCGDan_2391 | 798 | 2588602 | 2589399 | BCGDan_sapMKO_2393 | BCGDan_sapMKO_2393 | 798 |
| 2644320 | 2645774 | amiA2 | Rv2363 | 1455 | 2609342 | 2610796 | amiA2 | BCG_2377 | 1455 | 2589441 | 2590895 | amiA2 | BCGDan_2392 | 1455 | 2589461 | 2590915 | amiA2 | BCGDan_sapMKO_2394 | 1455 |
| 2645771 | 2646673 | era | Rv2364c | 903 | 2610793 | 2611695 | era | BCG_2378c | 903 | 2590892 | 2591794 | era | BCGDan_2393 | 903 | 2590912 | 2591841 | era | BCGDan_sapMKO_2395 | 930 |
| 2646747 | 2647088 | Rv2365c | Rv2365c | 342 | 2611769 | 2612110 | BCG_2379c | BCG_2379c | 342 | 2591868 | 2592209 | BCGDan_2394 | BCGDan_2394 | 342 | 2591888 | 2592226 | BCGDan_sapMKO_2396 | BCGDan_sapMKO_2396 | 339 |
| 2647060 | 2648367 | Rv2366c | Rv2366c | 1308 | 2612082 | 2613389 | BCG_2380c | BCG_2380c | 1308 | 2592181 | 2593488 | BCGDan_2395 | BCGDan_2395 | 1308 | 2592201 | 2593508 | BCGDan_sapMKO_2397 | BCGDan_sapMKO_2397 | 1308 |
| 2648364 | 2648912 | Rv2367c | Rv2367c | 549 | 2613386 | 2613934 | BCG_2381c | BCG_2381c | 549 | 2593485 | 2594033 | BCGDan_2396 | BCGDan_2396 | 549 | 2593505 | 2594038 | BCGDan_sapMKO_2398 | BCGDan_sapMKO_2398 | 534 |
| 2648916 | 2649974 | phoH1 | Rv2368c | 1059 | 2613938 | 2614996 | phoH1 | BCG_2382c | 1059 | 2594037 | 2595095 | phoH1 | BCGDan_2397 | 1059 | 2594057 | 2595115 | phoH1 | BCGDan_sapMKO_2399 | 1059 |
| 2649946 | 2650248 | Rv2369c | Rv2369c | 303 | 2614968 | 2615270 | BCG_2383c | BCG_2383c | 303 | 2595067 | 2595369 | BCGDan_2398 | BCGDan_2398 | 303 | 2595087 | 2595389 | BCGDan_sapMKO_2400 | BCGDan_sapMKO_2400 | 303 |
| 2650245 | 2651558 | Rv2370c | Rv2370c | 1314 | 2615267 | 2616580 | BCG_2384c | BCG_2384c | 1314 | 2595366 | 2596679 | BCGDan_2399 | BCGDan_2399 | 1314 | 2595386 | 2596699 | BCGDan_sapMKO_2401 | BCGDan_sapMKO_2401 | 1314 |
| 2651753 | 2651938 | PE_PGRS40 | Rv2371 | 186 | 2616775 | 2616960 | PE_PGRS40 | BCG_2385 | 186 | 2596874 | 2597059 | PE_PGRS40 | BCGDan_2400 | 186 | 2596894 | 2597079 | PE_PGRS40 | BCGDan_sapMKO_2402 | 186 |
| 2652037 | 2652825 | Rv2372c | Rv2372c | 789 | 2617059 | 2617847 | BCG_2386c | BCG_2386c | 789 | 2597158 | 2597946 | BCGDan_2401 | BCGDan_2401 | 789 | 2597178 | 2597966 | BCGDan_sapMKO_2403 | BCGDan_sapMKO_2403 | 789 |
| 2652839 | 2653987 | dnaJ2 | Rv2373c | 1149 | 2617861 | 2619009 | dnaJ2 | BCG_2387c | 1149 | 2597960 | 2599108 | dnaJ2 | BCGDan_2402 | 1149 | 2597980 | 2599128 | dnaJ2 | BCGDan_sapMKO_2404 | 1149 |
| 2654062 | 2655093 | hrcA | Rv2374c | 1032 | 2619084 | 2620115 | hrcA | BCG_2388c | 1032 | 2599183 | 2600214 | hrcA | BCGDan_2403 | 1032 | 2599203 | 2600234 | hrcA | BCGDan_sapMKO_2405 | 1032 |
| 2655265 | 2655582 | Rv2375 | Rv2375 | 318 | 2620287 | 2620604 | BCG_2389 | BCG_2389 | 318 | 2600386 | 2600703 | BCGDan_2404 | BCGDan_2404 | 318 | 2600406 | 2600723 | BCGDan_sapMKO_2406 | BCGDan_sapMKO_2406 | 318 |
| 2655609 | 2656115 | cfp2 | Rv2376c | 507 | 2620631 | 2621137 | cfp2 | BCG_2390c | 507 | 2600730 | 2601236 | cfp2 | BCGDan_2405 | 507 | 2600750 | 2601256 | cfp2 | BCGDan_sapMKO_2407 | 507 |
| 2656215 | 2656430 | mbtH | Rv2377c | 216 | 2621237 | 2621452 | mbtH | BCG_2391c | 216 | 2601336 | 2601551 | mbtH | BCGDan_2406 | 216 | 2601356 | 2601571 | mbtH | BCGDan_sapMKO_2408 | 216 |
| 2656408 | 2657703 | mbtG | Rv2378c | 1296 | 2621430 | 2622725 | mbtG | BCG_2392c | 1296 | 2601529 | 2602824 | mbtG | BCGDan_2407 | 1296 | 2601549 | 2602844 | mbtG | BCGDan_sapMKO_2409 | 1296 |
| 2657700 | 2662085 | mbtF | Rv2379c | 4386 | 2622722 | 2627107 | mbtF | BCG_2393c | 4386 | 2602821 | 2607206 | mbtF | BCGDan_2408 | 4386 | 2602841 | 2607217 | mbtF | BCGDan_sapMKO_2410 | 4377 |
| 2662067 | 2667115 | mbtE | Rv2380c | 5049 | 2627089 | 2632137 | mbtE | BCG_2394c | 5049 | 2607188 | 2612236 | mbtE | BCGDan_2409 | 5049 | 2607208 | 2612403 | mbtE | BCGDan_sapMKO_2411 | 5196 |
| 2667255 | 2670269 | mbtD | Rv2381c | 3015 | 2632277 | 2635291 | mbtD | BCG_2395c | 3015 | 2612376 | 2615390 | mbtD | BCGDan_2410 | 3015 | 2612396 | 2615425 | mbtD | BCGDan_sapMKO_2412 | 3030 |
| 2670269 | 2671603 | mbtC | Rv2382c | 1335 | 2635291 | 2636625 | mbtC | BCG_2396c | 1335 | 2615390 | 2616724 | mbtC | BCGDan_2411 | 1335 | 2615410 | 2616750 | mbtC | BCGDan_sapMKO_2413 | 1341 |
| 2671593 | 2675837 | mbtB | Rv2383c | 4245 | 2636615 | 2640859 | mbtB | BCG_2397c | 4245 | 2616714 | 2620958 | mbtB | BCGDan_2412 | 4245 | 2616734 | 2620978 | mbtB | BCGDan_sapMKO_2414 | 4245 |
| 2675936 | 2677633 | mbtA | Rv2384 | 1698 | 2640958 | 2642655 | mbtA | BCG_2398 | 1698 | 2621057 | 2622754 | mbtA | BCGDan_2413 | 1698 | 2621076 | 2622773 | mbtA | BCGDan_sapMKO_2415 | 1698 |
| 2677729 | 2678649 | mbtJ | Rv2385 | 921 | 2642751 | 2643671 | mbtJ | BCG_2399 | 921 | 2622850 | 2623770 | mbtJ | BCGDan_2414 | 921 | 2622806 | 2623789 | mbtJ | BCGDan_sapMKO_2416 | 984 |
| 2678653 | 2680005 | mbtI | Rv2386c | 1353 | 2643675 | 2645027 | mbtI | BCG_2400c | 1353 | 2623774 | 2625126 | mbtI | BCGDan_2415 | 1353 | 2623793 | 2625145 | mbtI | BCGDan_sapMKO_2417 | 1353 |
| NA | NA | NA | NA | NA | NA | NA | NA | NA | NA | 2625579 | 2625788 | BCGDan_2416 | BCGDan_2416 | 210 | 2625597 | 2625806 | BCGDan_sapMKO_2418 | BCGDan_sapMKO_2418 | 210 |
| 2680765 | 2682018 | Rv2387 | Rv2387 | 1254 | 2645787 | 2647040 | BCG_2401 | BCG_2401 | 1254 | 2625886 | 2627139 | BCGDan_2417 | BCGDan_2417 | 1254 | 2625904 | 2627157 | BCGDan_sapMKO_2419 | BCGDan_sapMKO_2419 | 1254 |
| 2682015 | 2683142 | hemN | Rv2388c | 1128 | 2647037 | 2648164 | hemN | BCG_2402c | 1128 | 2627136 | 2628263 | hemN | BCGDan_2418 | 1128 | 2627154 | 2628281 | hemN | BCGDan_sapMKO_2420 | 1128 |
| 2683248 | 2683712 | rpfD | Rv2389c | 465 | 2648270 | 2648734 | rpfD | BCG_2403c | 465 | 2628369 | 2628833 | rpfD | BCGDan_2419 | 465 | 2628387 | 2628851 | rpfD | BCGDan_sapMKO_2421 | 465 |
| 2683709 | 2684266 | Rv2390c | Rv2390c | 558 | 2648731 | 2649288 | BCG_2404c | BCG_2404c | 558 | 2628830 | 2629387 | BCGDan_2420 | BCGDan_2420 | 558 | 2628848 | 2629405 | BCGDan_sapMKO_2422 | BCGDan_sapMKO_2422 | 558 |
| 2684679 | 2686370 | sirA | Rv2391 | 1692 | 2649701 | 2651392 | nirA | BCG_2405 | 1692 | 2629800 | 2631491 | nirA | BCGDan_2421 | 1692 | 2629842 | 2631509 | nirA | BCGDan_sapMKO_2423 | 1668 |
| 2686367 | 2687131 | cysH | Rv2392 | 765 | 2651389 | 2652153 | cysH | BCG_2406 | 765 | 2631488 | 2632252 | cysH | BCGDan_2422 | 765 | 2631506 | 2632270 | cysH | BCGDan_sapMKO_2424 | 765 |
| 2687128 | 2687973 | che1 | Rv2393 | 846 | 2652150 | 2653049 | BCG_2407 | BCG_2407 | 900 | 2632249 | 2633148 | BCGDan_2423 | BCGDan_2423 | 900 | 2632267 | 2633166 | BCGDan_sapMKO_2425 | BCGDan_sapMKO_2425 | 900 |
| 2688010 | 2689941 | ggtB | Rv2394 | 1932 | 2653086 | 2655017 | ggtB | BCG_2408 | 1932 | 2633185 | 2635116 | ggtB | BCGDan_2424 | 1932 | 2633245 | 2635134 | ggtB | BCGDan_sapMKO_2426 | 1890 |
| 2690072 | 2692075 | Rv2395 | Rv2395 | 2004 | 2655148 | 2655588 | BCG_2409 | BCG_2409 | 441 | 2635247 | 2635687 | BCGDan_2425 | BCGDan_2425 | 441 | 2635265 | 2635705 | BCGDan_sapMKO_2427 | BCGDan_sapMKO_2427 | 441 |
| NA | NA | NA | NA | NA | 2655602 | 2657152 | BCG_2410 | BCG_2410 | 1551 | 2635701 | 2637251 | BCGDan_2426 | BCGDan_2426 | 1551 | 2635719 | 2637269 | BCGDan_sapMKO_2428 | BCGDan_sapMKO_2428 | 1551 |
| 2692799 | 2693884 | PE_PGRS41 | Rv2396 | 1086 | 2657876 | 2658958 | PE_PGRS41 | BCG_2411 | 1083 | 2637975 | 2639057 | PE_PGRS41 | BCGDan_2427 | 1083 | 2637993 | 2639075 | PE_PGRS41 | BCGDan_sapMKO_2429 | 1083 |
| 2693909 | 2694964 | cysA1 | Rv2397c | 1056 | 2658983 | 2660038 | cysA1 | BCG_2412c | 1056 | 2639082 | 2640137 | cysA1 | BCGDan_2428 | 1056 | 2639100 | 2640155 | cysA1 | BCGDan_sapMKO_2430 | 1056 |
| 2694981 | 2695799 | cysW | Rv2398c | 819 | 2660055 | 2660873 | cysW | BCG_2413c | 819 | 2640154 | 2640972 | cysW | BCGDan_2429 | 819 | 2640172 | 2640990 | cysW | BCGDan_sapMKO_2431 | 819 |
| 2695796 | 2696647 | cysT | Rv2399c | 852 | 2660870 | 2661721 | cysT | BCG_2414c | 852 | 2640969 | 2641820 | cysT | BCGDan_2430 | 852 | 2640987 | 2641838 | cysT | BCGDan_sapMKO_2432 | 852 |
| 2696644 | 2697714 | subI | Rv2400c | 1071 | 2661718 | 2662788 | subI | BCG_2415c | 1071 | 2641817 | 2642887 | subI | BCGDan_2431 | 1071 | 2641835 | 2642830 | subI | BCGDan_sapMKO_2433 | 996 |
| 2697728 | 2698057 | Rv2401 | Rv2401 | 330 | 2662802 | 2663131 | BCG_2416 | BCG_2416 | 330 | 2642901 | 2643230 | BCGDan_2432 | BCGDan_2432 | 330 | 2642919 | 2643248 | BCGDan_sapMKO_2434 | BCGDan_sapMKO_2434 | 330 |

| | | | | | | | | | | | | | | | | | | | |
|---|---|---|---|---|---|---|---|---|---|---|---|---|---|---|---|---|---|---|---|
| 2698042 | 2698245 | Rv2401A | Rv2401A | 204 | 2663116 | 2663319 | BCG_2417c | BCG_2417c | 204 | 2643215 | 2643418 | BCGDan_2433 | BCGDan_2433 | 204 | 2643233 | 2643427 | BCGDan_sapMKO_2435 | BCGDan_sapMKO_2435 | 195 |
| 2698529 | 2700457 | Rv2402 | Rv2402 | 1929 | 2663603 | 2665531 | BCG_2418 | BCG_2418 | 1929 | 2643702 | 2645630 | BCGDan_2434 | BCGDan_2434 | 1929 | 2643606 | 2645648 | BCGDan_sapMKO_2436 | BCGDan_sapMKO_2436 | 2043 |
| 2700535 | 2701290 | lppR | Rv2403c | 756 | 2665609 | 2666364 | lppR | BCG_2419c | 756 | 2645708 | 2646463 | lppR | BCGDan_2435 | 756 | 2645726 | 2646481 | lppR | BCGDan_sapMKO_2437 | 756 |
| 2701287 | 2703248 | lepA | Rv2404c | 1962 | 2666361 | 2668322 | lepA | BCG_2420c | 1962 | 2646460 | 2648421 | lepA | BCGDan_2436 | 1962 | 2646478 | 2648226 | lepA | BCGDan_sapMKO_2438 | 1749 |
| 2703269 | 2703838 | Rv2405 | Rv2405 | 570 | 2668343 | 2668912 | BCG_2421 | BCG_2421 | 570 | 2648442 | 2649011 | BCGDan_2437 | BCGDan_2437 | 570 | 2648427 | 2649029 | BCGDan_sapMKO_2439 | BCGDan_sapMKO_2439 | 603 |
| 2704009 | 2704437 | Rv2406c | Rv2406c | 429 | 2669083 | 2669511 | BCG_2422c | BCG_2422c | 429 | 2649182 | 2649610 | BCGDan_2438 | BCGDan_2438 | 429 | 2649200 | 2649628 | BCGDan_sapMKO_2440 | BCGDan_sapMKO_2440 | 429 |
| 2704697 | 2705518 | Rv2407 | Rv2407 | 822 | 2669771 | 2670613 | BCG_2423 | BCG_2423 | 843 | 2649870 | 2650712 | BCGDan_2439 | BCGDan_2439 | 843 | 2649888 | 2650730 | BCGDan_sapMKO_2441 | BCGDan_sapMKO_2441 | 843 |
| 2706017 | 2706736 | PE24 | Rv2408 | 720 | 2671206 | 2671925 | PE24 | BCG_2424 | 720 | 2651305 | 2652024 | PE24 | BCGDan_2440 | 720 | 2651323 | 2652042 | PE24 | BCGDan_sapMKO_2442 | 720 |
| 2706494 | 2707333 | Rv2409c | Rv2409c | 840 | 2671683 | 2672522 | BCG_2425c | BCG_2425c | 840 | 2651782 | 2652621 | BCGDan_2441 | BCGDan_2441 | 840 | 2651800 | 2652639 | BCGDan_sapMKO_2443 | BCGDan_sapMKO_2443 | 840 |
| 2707333 | 2708310 | Rv2410c | Rv2410c | 978 | 2672522 | 2673499 | BCG_2426c | BCG_2426c | 978 | 2652621 | 2653598 | BCGDan_2442 | BCGDan_2442 | 978 | 2652639 | 2653616 | BCGDan_sapMKO_2444 | BCGDan_sapMKO_2444 | 978 |
| 2708310 | 2709965 | Rv2411c | Rv2411c | 1656 | 2673499 | 2675154 | BCG_2427c | BCG_2427c | 1656 | 2653598 | 2655253 | BCGDan_2443 | BCGDan_2443 | 1656 | 2653616 | 2655262 | BCGDan_sapMKO_2445 | BCGDan_sapMKO_2445 | 1647 |
| 2710075 | 2710335 | rpsT | Rv2412 | 261 | 2675264 | 2675524 | rpsT | BCG_2428 | 261 | 2655363 | 2655623 | rpsT | BCGDan_2444 | 261 | 2655381 | 2655641 | rpsT | BCGDan_sapMKO_2446 | 261 |
| 2710351 | 2711301 | Rv2413c | Rv2413c | 951 | 2675540 | 2676490 | BCG_2429c | BCG_2429c | 951 | 2655639 | 2656589 | BCGDan_2445 | BCGDan_2445 | 951 | 2655657 | 2656625 | BCGDan_sapMKO_2447 | BCGDan_sapMKO_2447 | 969 |
| 2711332 | 2712876 | Rv2414c | Rv2414c | 1545 | 2676521 | 2678065 | BCG_2430c | BCG_2430c | 1545 | 2656620 | 2658164 | BCGDan_2446 | BCGDan_2446 | 1545 | 2656638 | 2658182 | BCGDan_sapMKO_2448 | BCGDan_sapMKO_2448 | 1545 |
| 2712891 | 2713784 | Rv2415c | Rv2415c | 894 | 2678080 | 2678973 | BCG_2431c | BCG_2431c | 894 | 2658179 | 2659072 | BCGDan_2447 | BCGDan_2447 | 894 | 2658197 | 2659132 | BCGDan_sapMKO_2449 | BCGDan_sapMKO_2449 | 936 |
| 2714124 | 2715332 | eis | Rv2416c | 1209 | 2679313 | 2680539 | BCG_2432c | BCG_2432c | 1227 | 2659412 | 2660638 | BCGDan_2448 | BCGDan_2448 | 1227 | 2659430 | 2660656 | BCGDan_sapMKO_2450 | BCGDan_sapMKO_2450 | 1227 |
| 2715472 | 2716314 | Rv2417c | Rv2417c | 843 | 2680661 | 2681503 | BCG_2433c | BCG_2433c | 843 | 2660760 | 2661602 | BCGDan_2449 | BCGDan_2449 | 843 | 2660778 | 2661620 | BCGDan_sapMKO_2451 | BCGDan_sapMKO_2451 | 843 |
| 2716395 | 2717138 | Rv2418c | Rv2418c | 744 | 2681584 | 2682327 | BCG_2434c | BCG_2434c | 744 | 2661683 | 2662426 | BCGDan_2450 | BCGDan_2450 | 744 | 2661701 | 2662414 | BCGDan_sapMKO_2452 | BCGDan_sapMKO_2452 | 714 |
| 2717128 | 2717799 | gpgP | Rv2419c | 672 | 2682317 | 2682988 | BCG_2435c | BCG_2435c | 672 | 2662416 | 2663087 | BCGDan_2451 | BCGDan_2451 | 672 | 2662434 | 2663105 | BCGDan_sapMKO_2453 | BCGDan_sapMKO_2453 | 672 |
| 2717796 | 2718176 | Rv2420c | Rv2420c | 381 | 2682985 | 2683368 | BCG_2436c | BCG_2436c | 384 | 2663084 | 2663467 | BCGDan_2452 | BCGDan_2452 | 384 | 2663102 | 2663485 | BCGDan_sapMKO_2454 | BCGDan_sapMKO_2454 | 384 |
| 2718173 | 2718808 | nadD | Rv2421c | 636 | 2683365 | 2684000 | nadD | BCG_2437c | 636 | 2663464 | 2664099 | nadD | BCGDan_2453 | 636 | 2663482 | 2664117 | nadD | BCGDan_sapMKO_2455 | 636 |
| 2719083 | 2719355 | Rv2422 | Rv2422 | 273 | 2684275 | 2684547 | BCG_2438 | BCG_2438 | 273 | 2664374 | 2664646 | BCGDan_2454 | BCGDan_2454 | 273 | 2664392 | 2664664 | BCGDan_sapMKO_2456 | BCGDan_sapMKO_2456 | 273 |
| 2719597 | 2720643 | Rv2423 | Rv2423 | 1047 | 2684789 | 2685835 | BCG_2439 | BCG_2439 | 1047 | 2664888 | 2665934 | BCGDan_2455 | BCGDan_2455 | 1047 | 2664906 | 2665952 | BCGDan_sapMKO_2457 | BCGDan_sapMKO_2457 | 1047 |
| 2720776 | 2721777 | Rv2424c | Rv2424c | 1002 | 2685968 | 2686660 | BCG_2440c | BCG_2440c | 693 | 2666067 | 2666759 | BCGDan_2456 | BCGDan_2456 | 693 | 2666085 | 2666825 | BCGDan_sapMKO_2458 | BCGDan_sapMKO_2458 | 741 |
| NA | NA | NA | NA | NA | 2686674 | 2686967 | BCG_2441c | BCG_2441c | 294 | 2666773 | 2667066 | BCGDan_2457 | BCGDan_2457 | 294 | 2666791 | 2667084 | BCGDan_sapMKO_2459 | BCGDan_sapMKO_2459 | 294 |
| 2721866 | 2723308 | Rv2425c | Rv2425c | 1443 | 2687056 | 2688498 | BCG_2442c | BCG_2442c | 1443 | 2667155 | 2668597 | BCGDan_2458 | BCGDan_2458 | 1443 | 2667173 | 2668552 | BCGDan_sapMKO_2460 | BCGDan_sapMKO_2460 | 1380 |
| 2723308 | 2724183 | Rv2426c | Rv2426c | 876 | 2688498 | 2689373 | BCG_2443c | BCG_2443c | 876 | 2668597 | 2669472 | BCGDan_2459 | BCGDan_2459 | 876 | 2668615 | 2669490 | BCGDan_sapMKO_2461 | BCGDan_sapMKO_2461 | 876 |
| 2724230 | 2725477 | proA | Rv2427c | 1248 | 2689400 | 2690647 | proA | BCG_2444c | 1248 | 2669499 | 2670746 | proA | BCGDan_2460 | 1248 | 2669517 | 2670764 | proA | BCGDan_sapMKO_2462 | 1248 |
| 2725571 | 2726087 | oxyR' | Rv2427A | 517 | 2690730 | 2691035 | BCG_2445c | BCG_2445c | 306 | 2670829 | 2671134 | BCGDan_2461 | BCGDan_2461 | 306 | 2670850 | 2671152 | BCGDan_sapMKO_2463 | BCGDan_sapMKO_2463 | 303 |
| NA | NA | NA | NA | NA | 2691036 | 2691158 | BCG_2446c | BCG_2446c | 123 | 2671135 | 2671257 | BCGDan_2462 | BCGDan_2462 | 123 | 2671156 | 2671275 | BCGDan_sapMKO_2464 | BCGDan_sapMKO_2464 | 120 |
| 2726193 | 2726780 | ahpC | Rv2428 | 588 | 2691363 | 2691950 | ahpC | BCG_2447 | 588 | 2671462 | 2672049 | ahpC | BCGDan_2463 | 588 | 2671480 | 2672067 | ahpC | BCGDan_sapMKO_2465 | 588 |
| 2726806 | 2727339 | ahpD | Rv2429 | 534 | 2691976 | 2692509 | ahpD | BCG_2448 | 534 | 2672075 | 2672608 | ahpD | BCGDan_2464 | 534 | 2672093 | 2672626 | ahpD | BCGDan_sapMKO_2466 | 534 |
| 2727336 | 2727920 | PPE41 | Rv2430c | 585 | 2692506 | 2693090 | PPE41 | BCG_2449c | 585 | 2672605 | 2673189 | PPE41 | BCGDan_2465 | 585 | 2672623 | 2673207 | PPE41 | BCGDan_sapMKO_2467 | 585 |
| 2727967 | 2728266 | PE25 | Rv2431c | 300 | 2693137 | 2693436 | PE25 | BCG_2450c | 300 | 2673236 | 2673535 | PE25 | BCGDan_2466 | 300 | 2673254 | 2673553 | PE25 | BCGDan_sapMKO_2468 | 300 |
| 2728437 | 2728847 | Rv2432c | Rv2432c | 411 | 2693607 | 2694017 | BCG_2451c | BCG_2451c | 411 | 2673706 | 2674116 | BCGDan_2467 | BCGDan_2467 | 411 | 2673724 | 2674134 | BCGDan_sapMKO_2469 | BCGDan_sapMKO_2469 | 411 |
| 2728844 | 2729134 | Rv2433c | Rv2433c | 291 | 2694014 | 2694304 | BCG_2452c | BCG_2452c | 291 | 2674113 | 2674403 | BCGDan_2468 | BCGDan_2468 | 291 | 2674131 | 2674421 | BCGDan_sapMKO_2470 | BCGDan_sapMKO_2470 | 291 |
| 2729115 | 2730560 | Rv2434c | Rv2434c | 1446 | 2694285 | 2695730 | BCG_2453c | BCG_2453c | 1446 | 2674384 | 2675829 | BCGDan_2469 | BCGDan_2469 | 1446 | 2674402 | 2675847 | BCGDan_sapMKO_2471 | BCGDan_sapMKO_2471 | 1446 |
| 2730557 | 2732749 | Rv2435c | Rv2435c | 2193 | 2695727 | 2697919 | BCG_2454c | BCG_2454c | 2193 | 2675826 | 2678018 | BCGDan_2470 | BCGDan_2470 | 2193 | 2675844 | 2677916 | BCGDan_sapMKO_2472 | BCGDan_sapMKO_2472 | 2073 |
| 2733230 | 2734144 | rbsK | Rv2436 | 915 | 2698400 | 2699314 | rbsK | BCG_2455 | 915 | 2678499 | 2679413 | rbsK | BCGDan_2471 | 915 | 2678550 | 2679431 | rbsK | BCGDan_sapMKO_2473 | 882 |
| 2734376 | 2734795 | Rv2437 | Rv2437 | 420 | 2699546 | 2699965 | BCG_2456 | BCG_2456 | 420 | 2679645 | 2680064 | BCGDan_2472 | BCGDan_2472 | 420 | 2679645 | 2680082 | BCGDan_sapMKO_2474 | BCGDan_sapMKO_2474 | 438 |
| 2734792 | 2736831 | nadE | Rv2438c | 2040 | 2699962 | 2702001 | nadE | BCG_2457c | 2040 | 2680061 | 2682100 | nadE | BCGDan_2473 | 2040 | 2680079 | 2682118 | nadE | BCGDan_sapMKO_2475 | 2040 |
| 2736709 | 2736987 | Rv2438A | Rv2438A | 279 | 2701879 | 2702157 | BCG_2458 | BCG_2458 | 279 | 2681978 | 2682256 | BCGDan_2474 | BCGDan_2474 | 279 | 2681996 | 2682274 | BCGDan_sapMKO_2476 | BCGDan_sapMKO_2476 | 279 |
| 2737117 | 2738247 | proB | Rv2439c | 1131 | 2702287 | 2703417 | proB | BCG_2459c | 1131 | 2682386 | 2683516 | proB | BCGDan_2475 | 1131 | 2682404 | 2683534 | proB | BCGDan_sapMKO_2477 | 1131 |
| 2738247 | 2739686 | obg | Rv2440c | 1440 | 2703417 | 2704856 | obg | BCG_2460c | 1440 | 2683516 | 2684955 | obg | BCGDan_2476 | 1440 | 2683534 | 2684988 | obg | BCGDan_sapMKO_2478 | 1455 |
| 2739772 | 2740032 | rpmA | Rv2441c | 261 | 2704942 | 2705202 | rpmA | BCG_2461c | 261 | 2685041 | 2685301 | rpmA | BCGDan_2477 | 261 | 2685059 | 2685319 | rpmA | BCGDan_sapMKO_2479 | 261 |
| 2740047 | 2740361 | rplU | Rv2442c | 315 | 2705217 | 2705531 | rplU | BCG_2462c | 315 | 2685316 | 2685630 | rplU | BCGDan_2478 | 315 | 2685334 | 2685678 | rplU | BCGDan_sapMKO_2480 | 345 |
| 2740709 | 2742184 | dctA | Rv2443 | 1476 | 2705879 | 2707354 | dctA | BCG_2463 | 1476 | 2685978 | 2687453 | dctA | BCGDan_2479 | 1476 | 2685996 | 2687471 | dctA | BCGDan_sapMKO_2481 | 1476 |
| 2742123 | 2744984 | rne | Rv2444c | 2862 | 2707293 | 2710154 | rne | BCG_2464c | 2862 | 2687392 | 2690253 | rne | BCGDan_2480 | 2862 | 2687410 | 2690157 | rne | BCGDan_sapMKO_2482 | 2748 |
| 2745314 | 2745724 | ndkA | Rv2445c | 411 | 2710484 | 2710894 | ndkA | BCG_2465c | 411 | 2690583 | 2690993 | ndkA | BCGDan_2481 | 411 | 2690601 | 2691023 | ndkA | BCGDan_sapMKO_2483 | 423 |
| 2745767 | 2746138 | Rv2446c | Rv2446c | 372 | 2710937 | 2711308 | BCG_2466c | BCG_2466c | 372 | 2691036 | 2691407 | BCGDan_2482 | BCGDan_2482 | 372 | 2691054 | 2691425 | BCGDan_sapMKO_2484 | BCGDan_sapMKO_2484 | 372 |
| 2746135 | 2747598 | folC | Rv2447c | 1464 | 2711305 | 2712768 | folC | BCG_2467c | 1464 | 2691404 | 2692867 | folC | BCGDan_2483 | 1464 | 2691422 | 2692885 | folC | BCGDan_sapMKO_2485 | 1464 |
| 2747595 | 2750225 | valS | Rv2448c | 2631 | 2712765 | 2715395 | valS | BCG_2468c | 2631 | 2692864 | 2695494 | valS | BCGDan_2484 | 2631 | 2692882 | 2695512 | valS | BCGDan_sapMKO_2486 | 2631 |
| 2750313 | 2751572 | Rv2449c | Rv2449c | 1260 | 2715483 | 2716742 | BCG_2469c | BCG_2469c | 1260 | 2695582 | 2696841 | BCGDan_2485 | BCGDan_2485 | 1260 | 2695600 | 2696859 | BCGDan_sapMKO_2487 | BCGDan_sapMKO_2487 | 1260 |
| 2751662 | 2752180 | rpfE | Rv2450c | 519 | 2716832 | 2717350 | rpfE | BCG_2470c | 519 | 2696931 | 2697449 | rpfE | BCGDan_2486 | 519 | 2696949 | 2697467 | rpfE | BCGDan_sapMKO_2488 | 519 |
| 2752262 | 2752660 | Rv2451 | Rv2451 | 399 | 2717432 | 2717830 | BCG_2471 | BCG_2471 | 399 | 2697531 | 2697929 | BCGDan_2487 | BCGDan_2487 | 399 | 2697549 | 2697947 | BCGDan_sapMKO_2489 | BCGDan_sapMKO_2489 | 399 |
| 2752848 | 2752994 | Rv2452c | Rv2452c | 147 | 2718018 | 2718164 | BCG_2472c | BCG_2472c | 147 | 2698117 | 2698263 | BCGDan_2488 | BCGDan_2488 | 147 | 2698135 | 2698281 | BCGDan_sapMKO_2490 | BCGDan_sapMKO_2490 | 147 |
| 2753018 | 2753623 | mobA | Rv2453c | 606 | 2718188 | 2718793 | mobA | BCG_2473c | 606 | 2698287 | 2698892 | mobA | BCGDan_2489 | 606 | 2698305 | 2698910 | mobA | BCGDan_sapMKO_2491 | 606 |
| 2753625 | 2754746 | Rv2454c | Rv2454c | 1122 | 2718795 | 2719916 | BCG_2474c | BCG_2474c | 1122 | 2698894 | 2700015 | BCGDan_2490 | BCGDan_2490 | 1122 | 2698912 | 2700033 | BCGDan_sapMKO_2492 | BCGDan_sapMKO_2492 | 1122 |
| 2754743 | 2756704 | Rv2455c | Rv2455c | 1962 | 2719913 | 2721874 | BCG_2475c | BCG_2475c | 1962 | 2700012 | 2701973 | BCGDan_2491 | BCGDan_2491 | 1962 | 2700030 | 2702006 | BCGDan_sapMKO_2493 | BCGDan_sapMKO_2493 | 1977 |
| 2756936 | 2758192 | Rv2456c | Rv2456c | 1257 | 2722106 | 2723362 | BCG_2476c | BCG_2476c | 1257 | 2702205 | 2703461 | BCGDan_2492 | BCGDan_2492 | 1257 | 2702087 | 2703478 | BCGDan_sapMKO_2494 | BCGDan_sapMKO_2494 | 1392 |
| 2758208 | 2759488 | clpX | Rv2457c | 1281 | 2723378 | 2724658 | clpX | BCG_2477c | 1281 | 2703477 | 2704757 | clpX | BCGDan_2493 | 1281 | 2703494 | 2704774 | clpX_1 | BCGDan_sapMKO_2495 | 1281 |
| 2759779 | 2760687 | mmuM | Rv2458 | 909 | 2724949 | 2725857 | mmuM | BCG_2478 | 909 | 2705048 | 2705956 | mmuM | BCGDan_2494 | 909 | 2705065 | 2705973 | mmuM | BCGDan_sapMKO_2496 | 909 |
| 2760854 | 2762380 | Rv2459 | Rv2459 | 1527 | 2726024 | 2727550 | BCG_2479 | BCG_2479 | 1527 | 2706123 | 2707649 | BCGDan_2495 | BCGDan_2495 | 1527 | 2706050 | 2707666 | BCGDan_sapMKO_2497 | BCGDan_sapMKO_2497 | 1617 |
| 2762531 | 2763175 | clpP2 | Rv2460c | 645 | 2727701 | 2728345 | clpP2 | BCG_2480c | 645 | 2707800 | 2708444 | clpP2 | BCGDan_2496 | 645 | 2707817 | 2708461 | clpP2 | BCGDan_sapMKO_2498 | 645 |
| 2763172 | 2763774 | clpP1 | Rv2461c | 603 | 2728342 | 2728944 | clpP1 | BCG_2481c | 603 | 2708441 | 2709043 | clpP1 | BCGDan_2497 | 603 | 2708458 | 2709042 | clpP1 | BCGDan_sapMKO_2499 | 585 |
| 2763891 | 2765291 | tig | Rv2462c | 1401 | 2729061 | 2730461 | tig | BCG_2482c | 1401 | 2709160 | 2710560 | tig | BCGDan_2498 | 1401 | 2709177 | 2710577 | tig | BCGDan_sapMKO_2500 | 1401 |
| 2765655 | 2766839 | lipP | Rv2463 | 1185 | 2730825 | 2732009 | lipP | BCG_2483 | 1185 | 2710924 | 2712108 | lipP | BCGDan_2501 | 1185 | 2710941 | 2712125 | lipP | BCGDan_sapMKO_2503 | 1185 |
| 2766859 | 2767665 | Rv2464c | Rv2464c | 807 | 2732029 | 2732835 | BCG_2484c | BCG_2484c | 807 | 2712128 | 2712934 | BCGDan_2502 | BCGDan_2502 | 807 | 2712145 | 2712951 | BCGDan_sapMKO_2504 | BCGDan_sapMKO_2504 | 807 |
| 2767671 | 2768159 | rpiB | Rv2465c | 489 | 2732841 | 2733329 | rpiB | BCG_2485c | 489 | 2712940 | 2713428 | rpiB | BCGDan_2503 | 489 | 2712957 | 2713457 | rpiB | BCGDan_sapMKO_2505 | 501 |
| 2768261 | 2768884 | Rv2466c | Rv2466c | 624 | 2733431 | 2734054 | BCG_2486c | BCG_2486c | 624 | 2713530 | 2714153 | BCGDan_2504 | BCGDan_2504 | 624 | 2713547 | 2714170 | BCGDan_sapMKO_2506 | BCGDan_sapMKO_2506 | 624 |
| 2768986 | 2771571 | pepN | Rv2467 | 2586 | 2734156 | 2736741 | pepN | BCG_2487 | 2586 | 2714255 | 2716840 | pepN | BCGDan_2505 | 2586 | 2714272 | 2716857 | pepN | BCGDan_sapMKO_2507 | 2586 |
| 2771644 | 2772147 | Rv2468c | Rv2468c | 504 | 2736814 | 2737317 | BCG_2488c | BCG_2488c | 504 | 2716913 | 2717416 | BCGDan_2506 | BCGDan_2506 | 504 | 2716930 | 2717397 | BCGDan_sapMKO_2508 | BCGDan_sapMKO_2508 | 468 |
| 2772098 | 2772331 | Rv2468A | Rv2468A | 234 | NA | NA | NA | NA | NA | 2717367 | 2717600 | BCGDan_2507 | BCGDan_2507 | 234 | 2717387 | 2717617 | BCGDan_sapMKO_2509 | BCGDan_sapMKO_2509 | 231 |
| 2772367 | 2773035 | Rv2469c | Rv2469c | 669 | 2737537 | 2738205 | BCG_2489c | BCG_2489c | 669 | 2717636 | 2718367 | BCGDan_2508 | BCGDan_2508 | 732 | 2717653 | 2718384 | BCGDan_sapMKO_2510 | BCGDan_sapMKO_2510 | 732 |
| 2773178 | 2773564 | glbO | Rv2470 | 387 | 2738348 | 2738734 | glbO | BCG_2490 | 387 | 2718447 | 2718833 | glbO | BCGDan_2509 | 387 | 2718455 | 2718850 | glbO | BCGDan_sapMKO_2511 | 396 |
| 2773564 | 2775204 | aglA | Rv2471 | 1641 | 2738734 | 2740374 | aglA | BCG_2491 | 1641 | 2718833 | 2720473 | aglA | BCGDan_2510 | 1641 | 2718928 | 2720490 | aglA | BCGDan_sapMKO_2512 | 1563 |
| 2775272 | 2775565 | Rv2472 | Rv2472 | 294 | 2740442 | 2740735 | BCG_2492 | BCG_2492 | 294 | 2720541 | 2720834 | BCGDan_2511 | BCGDan_2511 | 294 | 2720558 | 2720851 | BCGDan_sapMKO_2513 | BCGDan_sapMKO_2513 | 294 |
| 2775568 | 2776284 | Rv2473 | Rv2473 | 717 | 2740738 | 2741454 | BCG_2493 | BCG_2493 | 717 | 2720837 | 2721553 | BCGDan_2512 | BCGDan_2512 | 717 | 2720854 | 2721570 | BCGDan_sapMKO_2514 | BCGDan_sapMKO_2514 | 717 |
| 2776316 | 2776969 | Rv2474c | Rv2474c | 654 | 2741486 | 2742139 | BCG_2494c | BCG_2494c | 654 | 2721585 | 2722238 | BCGDan_2513 | BCGDan_2513 | 654 | 2721602 | 2722276 | BCGDan_sapMKO_2515 | BCGDan_sapMKO_2515 | 675 |
| 2776975 | 2777391 | Rv2475c | Rv2475c | 417 | 2742145 | 2742561 | BCG_2495c | BCG_2495c | 417 | 2722244 | 2722660 | BCGDan_2514 | BCGDan_2514 | 417 | 2722261 | 2722677 | BCGDan_sapMKO_2516 | BCGDan_sapMKO_2516 | 417 |
| 2777388 | 2782262 | gdh | Rv2476c | 4875 | 2742558 | 2747429 | gdh | BCG_2496c | 4872 | 2722657 | 2727528 | gdh | BCGDan_2515 | 4872 | 2722674 | 2727545 | gdh | BCGDan_sapMKO_2517 | 4872 |
| 2782366 | 2784042 | Rv2477c | Rv2477c | 1677 | 2747533 | 2749209 | BCG_2497c | BCG_2497c | 1677 | 2727632 | 2729308 | BCGDan_2516 | BCGDan_2516 | 1677 | 2727649 | 2729325 | BCGDan_sapMKO_2518 | BCGDan_sapMKO_2518 | 1677 |
| 2784123 | 2784608 | Rv2478c | Rv2478c | 486 | 2749290 | 2749775 | BCG_2498 | BCG_2498 | 486 | 2729389 | 2729874 | BCGDan_2517 | BCGDan_2517 | 486 | 2729406 | 2729912 | BCGDan_sapMKO_2519 | BCGDan_sapMKO_2519 | 507 |
| 2786575 | 2786898 | Rv2481c | Rv2481c | 324 | 2750384 | 2750707 | BCG_2499c | BCG_2499c | 324 | 2730483 | 2730806 | BCGDan_2518 | BCGDan_2518 | 324 | 2730500 | 2730823 | BCGDan_sapMKO_2520 | BCGDan_sapMKO_2520 | 324 |
| 2786914 | 2789283 | plsB2 | Rv2482c | 2370 | 2750723 | 2753092 | plsB2 | BCG_2500c | 2370 | 2730822 | 2733191 | plsB2 | BCGDan_2519 | 2370 | 2730839 | 2733208 | plsB2 | BCGDan_sapMKO_2521 | 2370 |
| 2789280 | 2791022 | plsC | Rv2483c | 1743 | 2753089 | 2754831 | plsC | BCG_2501c | 1743 | 2733188 | 2734930 | plsC | BCGDan_2520 | 1743 | 2733205 | 2734947 | plsC | BCGDan_sapMKO_2522 | 1743 |

| | | | | | | | | | | | | | | | | | | | |
|---|---|---|---|---|---|---|---|---|---|---|---|---|---|---|---|---|---|---|---|
| 2791019 | 2792494 | Rv2484c | Rv2484c | 1476 | 2754828 | 2756303 | BCG_2502c | BCG_2502c | 1476 | 2734927 | 2736402 | BCGDan_2521 | BCGDan_2521 | 1476 | 2734944 | 2736458 | BCGDan_sapMKO_2523 | BCGDan_sapMKO_2523 | 1515 |
| 2792723 | 2793988 | lipQ | Rv2485c | 1266 | 2756532 | 2757797 | lipQ | BCG_2503c | 1266 | 2736631 | 2737896 | lipQ | BCGDan_2522 | 1266 | 2736648 | 2737913 | lipQ | BCGDan_sapMKO_2524 | 1266 |
| 2794350 | 2795120 | echA14 | Rv2486 | 771 | 2758159 | 2758929 | echA14 | BCG_2504 | 771 | 2738258 | 2739028 | echA14 | BCGDan_2524 | 771 | 2738275 | 2739045 | echA14 | BCGDan_sapMKO_2526 | 771 |
| NA | NA | NA | NA | NA | 2759109 | 2759933 | PE_PGRS42d | BCG_2505c | 825 | 2739208 | 2740032 | PE_PGRS42d | BCGDan_2525 | 825 | NA | NA | NA | NA | NA |
| 2795301 | 2797385 | PE_PGRS42 | Rv2487c | 2085 | 2759609 | 2761192 | PE_PGRS42ab | BCG_2506c | 1584 | 2739708 | 2741291 | PE_PGRS42ab | BCGDan_2526 | 1584 | 2739724 | 2741307 | PE_PGRS42ab | BCGDan_sapMKO_2527 | 1584 |
| 2797467 | 2800880 | Rv2488c | Rv2488c | 3414 | 2761274 | 2764687 | BCG_2507c | BCG_2507c | 3414 | 2741373 | 2744786 | BCGDan_2527 | BCGDan_2527 | 3414 | 2741389 | 2744802 | BCGDan_sapMKO_2528 | BCGDan_sapMKO_2528 | 3414 |
| 2800846 | 2801145 | Rv2489c | Rv2489c | 300 | 2764653 | 2764952 | BCG_2508c | BCG_2508c | 300 | 2744752 | 2745051 | BCGDan_2528 | BCGDan_2528 | 300 | 2744768 | 2745067 | BCGDan_sapMKO_2529 | BCGDan_sapMKO_2529 | 300 |
| NA | NA | NA | NA | NA | 2765061 | 2768513 | PE_PGRS43b | BCG_2509c | 3453 | 2745160 | 2748615 | PE_PGRS43b | BCGDan_2529 | 3456 | 2745176 | 2748898 | PE_PGRS43b | BCGDan_sapMKO_2530 | 3723 |
| 2801254 | 2806236 | PE_PGRS43 | Rv2490c | 4983 | 2768605 | 2770032 | PE_PGRS43a | BCG_2510c | 1428 | 2748707 | 2750134 | PE_PGRS43a | BCGDan_2530 | 1428 | 2748723 | 2750150 | PE_PGRS43a | BCGDan_sapMKO_2531 | 1428 |
| 2806665 | 2807288 | Rv2491 | Rv2491 | 624 | 2770461 | 2771084 | BCG_2511 | BCG_2511 | 624 | 2750563 | 2751186 | BCGDan_2531 | BCGDan_2531 | 624 | 2750579 | 2751202 | BCGDan_sapMKO_2532 | BCGDan_sapMKO_2532 | 624 |
| 2807278 | 2808030 | Rv2492 | Rv2492 | 753 | 2771074 | 2771826 | BCG_2512 | BCG_2512 | 753 | 2751176 | 2751928 | BCGDan_2532 | BCGDan_2532 | 753 | 2751192 | 2751944 | BCGDan_sapMKO_2533 | BCGDan_sapMKO_2533 | 753 |
| 2808083 | 2808304 | vapB38 | Rv2493 | 222 | 2771879 | 2772100 | BCG_2513 | BCG_2513 | 222 | 2751981 | 2752202 | BCGDan_2533 | BCGDan_2533 | 222 | 2751997 | 2752218 | BCGDan_sapMKO_2534 | BCGDan_sapMKO_2534 | 222 |
| 2808310 | 2808735 | vapC38 | Rv2494 | 426 | 2772106 | 2772531 | BCG_2514 | BCG_2514 | 426 | 2752208 | 2752633 | BCGDan_2534 | BCGDan_2534 | 426 | 2752224 | 2752649 | BCGDan_sapMKO_2535 | BCGDan_sapMKO_2535 | 426 |
| 2808758 | 2809939 | bkdC | Rv2495c | 1182 | 2772554 | 2773735 | pdhC | BCG_2515c | 1182 | 2752656 | 2753837 | pdhC | BCGDan_2535 | 1182 | 2752672 | 2753853 | pdhC | BCGDan_sapMKO_2536 | 1182 |
| 2809936 | 2810982 | bkdB | Rv2496c | 1047 | 2773732 | 2774778 | pdhB | BCG_2516c | 1047 | 2753834 | 2754880 | pdhB | BCGDan_2536 | 1047 | 2753850 | 2754896 | pdhB | BCGDan_sapMKO_2537 | 1047 |
| 2810993 | 2812096 | bkdA | Rv2497c | 1104 | 2774789 | 2775892 | pdhA | BCG_2517c | 1104 | 2754891 | 2755994 | pdhA | BCGDan_2537 | 1104 | 2754907 | 2755980 | pdhA | BCGDan_sapMKO_2538 | 1074 |
| 2812355 | 2813176 | citE | Rv2498c | 822 | 2776151 | 2776972 | citE | BCG_2518c | 822 | 2756253 | 2757074 | citE | BCGDan_2538 | 822 | 2756269 | 2757090 | citE | BCGDan_sapMKO_2539 | 822 |
| 2813173 | 2813730 | Rv2499c | Rv2499c | 558 | 2776969 | 2777526 | BCG_2519c | BCG_2519c | 558 | 2757071 | 2757628 | BCGDan_2539 | BCGDan_2539 | 558 | 2757087 | 2757644 | BCGDan_sapMKO_2540 | BCGDan_sapMKO_2540 | 558 |
| 2813727 | 2814911 | fadE19 | Rv2500c | 1185 | 2777523 | 2778707 | fadE19 | BCG_2520c | 1185 | 2757625 | 2758809 | fadE19 | BCGDan_2540 | 1185 | 2757641 | 2758756 | fadE19 | BCGDan_sapMKO_2541 | 1116 |
| 2814916 | 2816880 | accA1 | Rv2501c | 1965 | 2778712 | 2780676 | accA1 | BCG_2521c | 1965 | 2758814 | 2760778 | accA1 | BCGDan_2541 | 1965 | 2758830 | 2760794 | accA1 | BCGDan_sapMKO_2542 | 1965 |
| 2816885 | 2818474 | accD1 | Rv2502c | 1590 | 2780681 | 2782270 | accD1 | BCG_2522c | 1590 | 2760783 | 2762372 | accD1 | BCGDan_2542 | 1590 | 2760799 | 2762388 | accD1 | BCGDan_sapMKO_2543 | 1590 |
| 2818471 | 2819127 | scoB | Rv2503c | 657 | 2782267 | 2782923 | scoB | BCG_2523c | 657 | 2762369 | 2763025 | scoB | BCGDan_2543 | 657 | 2762385 | 2763011 | scoB | BCGDan_sapMKO_2544 | 627 |
| 2819124 | 2819870 | scoA | Rv2504c | 747 | 2782920 | 2783666 | scoA | BCG_2524c | 747 | 2763022 | 2763768 | scoA | BCGDan_2544 | 747 | 2763038 | 2763784 | scoA | BCGDan_sapMKO_2545 | 747 |
| 2819953 | 2821596 | fadD35 | Rv2505c | 1644 | 2783749 | 2785392 | fadD35 | BCG_2525c | 1644 | 2763851 | 2765494 | fadD35 | BCGDan_2545 | 1644 | 2763867 | 2765444 | fadD35 | BCGDan_sapMKO_2546 | 1578 |
| 2821712 | 2822359 | Rv2506 | Rv2506 | 648 | 2785508 | 2786155 | BCG_2526 | BCG_2526 | 648 | 2765610 | 2766257 | BCGDan_2546 | BCGDan_2546 | 648 | 2765626 | 2766273 | BCGDan_sapMKO_2547 | BCGDan_sapMKO_2547 | 648 |
| 2822438 | 2823259 | Rv2507 | Rv2507 | 822 | 2786234 | 2787055 | BCG_2527 | BCG_2527 | 822 | 2766336 | 2767157 | BCGDan_2547 | BCGDan_2547 | 822 | 2766352 | 2767173 | BCGDan_sapMKO_2548 | BCGDan_sapMKO_2548 | 822 |
| 2823256 | 2824593 | Rv2508c | Rv2508c | 1338 | 2787052 | 2788389 | BCG_2528c | BCG_2528c | 1338 | 2767154 | 2768491 | BCGDan_2548 | BCGDan_2548 | 1338 | 2767170 | 2768507 | BCGDan_sapMKO_2549 | BCGDan_sapMKO_2549 | 1338 |
| 2824678 | 2825484 | Rv2509 | Rv2509 | 807 | 2788474 | 2789280 | BCG_2529 | BCG_2529 | 807 | 2768576 | 2769382 | BCGDan_2549 | BCGDan_2549 | 807 | 2768592 | 2769398 | BCGDan_sapMKO_2550 | BCGDan_sapMKO_2550 | 807 |
| 2825488 | 2827089 | Rv2510c | Rv2510c | 1602 | 2789284 | 2790885 | BCG_2530c | BCG_2530c | 1602 | 2769386 | 2770987 | BCGDan_2550 | BCGDan_2550 | 1602 | 2769402 | 2771003 | BCGDan_sapMKO_2551 | BCGDan_sapMKO_2551 | 1602 |
| 2827157 | 2827804 | orn | Rv2511 | 648 | 2790953 | 2791600 | orn | BCG_2531 | 648 | 2771055 | 2771702 | orn | BCGDan_2551 | 648 | 2771071 | 2771718 | orn | BCGDan_sapMKO_2552 | 648 |
| 2828556 | 2829803 | Rv2512c | Rv2512c | 1248 | 2792352 | 2793599 | BCG_2532c | BCG_2532c | 1248 | 2772454 | 2773701 | BCGDan_2553 | BCGDan_2553 | 1248 | 2772470 | 2773780 | BCGDan_sapMKO_2554 | BCGDan_sapMKO_2554 | 1311 |
| NA | NA | NA | NA | NA | 2793599 | 2793736 | BCG_2533 | BCG_2533 | 138 | 2773701 | 2773838 | BCGDan_2554 | BCGDan_2554 | 138 | 2773717 | 2773854 | BCGDan_sapMKO_2555 | BCGDan_sapMKO_2555 | 138 |
| 2830161 | 2830583 | Rv2513 | Rv2513 | 423 | 2793957 | 2794379 | BCG_2534 | BCG_2534 | 423 | 2774059 | 2774481 | BCGDan_2555 | BCGDan_2555 | 423 | 2774075 | 2774497 | BCGDan_sapMKO_2556 | BCGDan_sapMKO_2556 | 423 |
| 2830877 | 2831338 | Rv2514c | Rv2514c | 462 | 2794673 | 2795134 | BCG_2535c | BCG_2535c | 462 | 2774775 | 2775236 | BCGDan_2556 | BCGDan_2556 | 462 | 2774791 | 2775252 | BCGDan_sapMKO_2557 | BCGDan_sapMKO_2557 | 462 |
| 2831344 | 2832591 | Rv2515c | Rv2515c | 1248 | 2795140 | 2796387 | BCG_2536c | BCG_2536c | 1248 | 2775242 | 2776489 | BCGDan_2557 | BCGDan_2557 | 1248 | 2775258 | 2776487 | BCGDan_sapMKO_2558 | BCGDan_sapMKO_2558 | 1230 |
| 2832710 | 2833513 | Rv2516c | Rv2516c | 804 | 2796506 | 2797309 | BCG_2537c | BCG_2537c | 804 | 2776608 | 2777411 | BCGDan_2558 | BCGDan_2558 | 804 | 2776624 | 2777427 | BCGDan_sapMKO_2559 | BCGDan_sapMKO_2559 | 804 |
| 2833510 | 2833761 | Rv2517c | Rv2517c | 252 | 2797306 | 2797557 | BCG_2538c | BCG_2538c | 252 | 2777408 | 2777659 | BCGDan_2559 | BCGDan_2559 | 252 | 2777424 | 2777675 | BCGDan_sapMKO_2560 | BCGDan_sapMKO_2560 | 252 |
| 2834109 | 2835335 | ldtB | Rv2518c | 1227 | 2797905 | 2799131 | lppS | BCG_2539c | 1227 | 2778007 | 2779233 | lppS | BCGDan_2560 | 1227 | 2778023 | 2779180 | lppS | BCGDan_sapMKO_2561 | 1158 |
| 2835785 | 2837263 | PE26 | Rv2519 | 1479 | 2799581 | 2801059 | PE26 | BCG_2540 | 1479 | 2779683 | 2781161 | PE26 | BCGDan_2562 | 1479 | 2779699 | 2781177 | PE26 | BCGDan_sapMKO_2563 | 1479 |
| 2837388 | 2837615 | Rv2520c | Rv2520c | 228 | 2801184 | 2801411 | BCG_2541c | BCG_2541c | 228 | 2781286 | 2781513 | BCGDan_2563 | BCGDan_2563 | 228 | 2781302 | 2781529 | BCGDan_sapMKO_2564 | BCGDan_sapMKO_2564 | 228 |
| 2837684 | 2838157 | bcp | Rv2521 | 474 | 2801480 | 2801953 | bcp | BCG_2542 | 474 | 2781582 | 2782055 | bcp | BCGDan_2564 | 474 | 2781598 | 2782071 | bcp | BCGDan_sapMKO_2565 | 474 |
| 2838129 | 2839541 | Rv2522c | Rv2522c | 1413 | 2801925 | 2803337 | BCG_2543c | BCG_2543c | 1413 | 2782027 | 2783439 | BCGDan_2565 | BCGDan_2565 | 1413 | 2782043 | 2783365 | BCGDan_sapMKO_2566 | BCGDan_sapMKO_2566 | 1323 |
| 2839538 | 2839930 | acpS | Rv2523c | 393 | 2803334 | 2803726 | acpS | BCG_2544c | 393 | 2783436 | 2783828 | acpS | BCGDan_2566 | 393 | 2783452 | 2783844 | acpS | BCGDan_sapMKO_2567 | 393 |
| 2840123 | 2849332 | fas | Rv2524c | 9210 | 2803919 | 2813128 | fas | BCG_2545c | 9210 | 2784021 | 2793230 | fas | BCGDan_2567 | 9210 | 2784037 | 2793246 | fas | BCGDan_sapMKO_2568 | 9210 |
| 2849852 | 2850574 | Rv2525c | Rv2525c | 723 | 2813648 | 2814370 | BCG_2546c | BCG_2546c | 723 | 2793750 | 2794472 | BCGDan_2568 | BCGDan_2568 | 723 | 2793766 | 2794488 | BCGDan_sapMKO_2569 | BCGDan_sapMKO_2569 | 723 |
| 2851091 | 2851318 | vapB17 | Rv2526 | 228 | 2814887 | 2815114 | BCG_2547 | BCG_2547 | 228 | 2794989 | 2795216 | BCGDan_2569 | BCGDan_2569 | 228 | 2795005 | 2795232 | BCGDan_sapMKO_2570 | BCGDan_sapMKO_2570 | 228 |
| 2851315 | 2851716 | vapC17 | Rv2527 | 402 | 2815111 | 2815512 | BCG_2548 | BCG_2548 | 402 | 2795213 | 2795614 | BCGDan_2570 | BCGDan_2570 | 402 | 2795229 | 2795630 | BCGDan_sapMKO_2571 | BCGDan_sapMKO_2571 | 402 |
| 2851751 | 2852671 | mrr | Rv2528c | 921 | 2815547 | 2816467 | mrr | BCG_2549c | 921 | 2795649 | 2796569 | mrr | BCGDan_2571 | 921 | 2795665 | 2796585 | mrr | BCGDan_sapMKO_2572 | 921 |
| 2852875 | 2854266 | Rv2529 | Rv2529 | 1392 | 2816671 | 2818062 | BCG_2550 | BCG_2550 | 1392 | 2796773 | 2798164 | BCGDan_2572 | BCGDan_2572 | 1392 | 2796789 | 2798180 | BCGDan_sapMKO_2573 | BCGDan_sapMKO_2573 | 1392 |
| 2854267 | 2854686 | vapC39 | Rv2530c | 420 | 2818063 | 2818482 | BCG_2551c | BCG_2551c | 420 | 2798165 | 2798584 | BCGDan_2573 | BCGDan_2573 | 420 | 2798181 | 2798600 | BCGDan_sapMKO_2574 | BCGDan_sapMKO_2574 | 420 |
| 2854683 | 2854907 | vapB39 | Rv2530A | 225 | 2818479 | 2818703 | BCG_2552c | BCG_2552c | 225 | 2798581 | 2798805 | BCGDan_2574 | BCGDan_2574 | 225 | 2798597 | 2798821 | BCGDan_sapMKO_2575 | BCGDan_sapMKO_2575 | 225 |
| 2854938 | 2857781 | Rv2531c | Rv2531c | 2844 | 2818734 | 2821577 | BCG_2553c | BCG_2553c | 2844 | 2798836 | 2801679 | BCGDan_2575 | BCGDan_2575 | 2844 | 2798852 | 2801695 | BCGDan_sapMKO_2576 | BCGDan_sapMKO_2576 | 2844 |
| 2857853 | 2858254 | Rv2532c | Rv2532c | 402 | 2821649 | 2822050 | BCG_2554c | BCG_2554c | 402 | 2801751 | 2802152 | BCGDan_2576 | BCGDan_2576 | 402 | 2801767 | 2802168 | BCGDan_sapMKO_2577 | BCGDan_sapMKO_2577 | 402 |
| 2858254 | 2858724 | nusB | Rv2533c | 471 | 2822050 | 2822520 | nusB | BCG_2555c | 471 | 2802152 | 2802622 | nusB | BCGDan_2577 | 471 | 2802168 | 2802638 | nusB | BCGDan_sapMKO_2578 | 471 |
| 2858727 | 2859290 | efp | Rv2534c | 564 | 2822523 | 2823086 | efp | BCG_2556c | 564 | 2802625 | 2803188 | efp | BCGDan_2578 | 564 | 2802641 | 2803204 | efp | BCGDan_sapMKO_2579 | 564 |
| 2859300 | 2860418 | pepQ | Rv2535c | 1119 | 2823096 | 2824214 | pepQ | BCG_2557c | 1119 | 2803198 | 2804316 | pepQ | BCGDan_2579 | 1119 | 2803214 | 2804269 | pepQ | BCGDan_sapMKO_2580 | 1056 |
| 2860452 | 2861144 | Rv2536 | Rv2536 | 693 | 2824248 | 2824940 | BCG_2558 | BCG_2558 | 693 | 2804350 | 2805042 | BCGDan_2580 | BCGDan_2580 | 693 | 2804366 | 2805058 | BCGDan_sapMKO_2581 | BCGDan_sapMKO_2581 | 693 |
| 2861148 | 2861591 | aroD | Rv2537c | 444 | 2824944 | 2825387 | aroD | BCG_2559c | 444 | 2805046 | 2805489 | aroD | BCGDan_2581 | 444 | 2805062 | 2805505 | aroD | BCGDan_sapMKO_2582 | 444 |
| 2861588 | 2862676 | aroB | Rv2538c | 1089 | 2825384 | 2826472 | aroB | BCG_2560c | 1089 | 2805486 | 2806574 | aroB | BCGDan_2582 | 1089 | 2805502 | 2806590 | aroB | BCGDan_sapMKO_2583 | 1089 |
| 2862673 | 2863203 | aroK | Rv2539c | 531 | 2826469 | 2826999 | aroK | BCG_2561c | 531 | 2806571 | 2807101 | aroK | BCGDan_2583 | 531 | 2806587 | 2807117 | aroK | BCGDan_sapMKO_2584 | 531 |
| 2863207 | 2864412 | aroF | Rv2540c | 1206 | 2827003 | 2828208 | aroF | BCG_2562c | 1206 | 2807105 | 2808310 | aroF | BCGDan_2584 | 1206 | 2807121 | 2808326 | aroF | BCGDan_sapMKO_2585 | 1206 |
| 2864427 | 2864834 | Rv2541 | Rv2541 | 408 | 2828223 | 2828630 | BCG_2563 | BCG_2563 | 408 | 2808325 | 2808732 | BCGDan_2585 | BCGDan_2585 | 408 | 2808341 | 2808748 | BCGDan_sapMKO_2586 | BCGDan_sapMKO_2586 | 408 |
| 2865130 | 2866341 | Rv2542 | Rv2542 | 1212 | 2828926 | 2830137 | BCG_2564 | BCG_2564 | 1212 | 2809028 | 2810239 | BCGDan_2586 | BCGDan_2586 | 1212 | 2809044 | 2810255 | BCGDan_sapMKO_2587 | BCGDan_sapMKO_2587 | 1212 |
| 2866468 | 2867127 | lppA | Rv2543 | 660 | 2830264 | 2830923 | lppA | BCG_2565 | 660 | 2810366 | 2811025 | lppA | BCGDan_2587 | 660 | 2810382 | 2811041 | lppA | BCGDan_sapMKO_2588 | 660 |
| NA | NA | NA | NA | NA | 2830920 | 2831579 | lprR | BCG_2566 | 660 | 2811022 | 2811681 | lprR | BCGDan_2588 | 660 | 2811038 | 2811697 | lprR | BCGDan_sapMKO_2589 | 660 |
| 2867124 | 2867786 | lppB | Rv2544 | 663 | 2831576 | 2832238 | lppB | BCG_2567 | 663 | 2811678 | 2812340 | lppB | BCGDan_2589 | 663 | 2811694 | 2812356 | lppB | BCGDan_sapMKO_2590 | 663 |
| 2867783 | 2868061 | vapB18 | Rv2545 | 279 | 2832235 | 2832513 | BCG_2568 | BCG_2568 | 279 | 2812337 | 2812615 | BCGDan_2590 | BCGDan_2590 | 279 | 2812353 | 2812631 | BCGDan_sapMKO_2591 | BCGDan_sapMKO_2591 | 279 |
| 2868154 | 2868567 | vapC18 | Rv2546 | 414 | 2832606 | 2833019 | BCG_2569 | BCG_2569 | 414 | 2812708 | 2813121 | BCGDan_2591 | BCGDan_2591 | 414 | 2812721 | 2813137 | BCGDan_sapMKO_2592 | BCGDan_sapMKO_2592 | 417 |
| 2868606 | 2868863 | vapB19 | Rv2547 | 258 | 2833058 | 2833315 | BCG_2570 | BCG_2570 | 258 | 2813160 | 2813417 | BCGDan_2592 | BCGDan_2592 | 258 | 2813161 | 2813433 | BCGDan_sapMKO_2593 | BCGDan_sapMKO_2593 | 273 |
| 2868860 | 2869237 | vapC19 | Rv2548 | 378 | 2833312 | 2833689 | BCG_2571 | BCG_2571 | 378 | 2813414 | 2813791 | BCGDan_2593 | BCGDan_2593 | 378 | 2813430 | 2813807 | BCGDan_sapMKO_2594 | BCGDan_sapMKO_2594 | 378 |
| 2869727 | 2870122 | vapC20 | Rv2549c | 396 | 2834179 | 2834574 | BCG_2572c | BCG_2572c | 396 | 2814281 | 2814676 | BCGDan_2594 | BCGDan_2594 | 396 | 2814297 | 2814692 | BCGDan_sapMKO_2595 | BCGDan_sapMKO_2595 | 396 |
| 2870119 | 2870364 | vapB20 | Rv2550c | 246 | 2834571 | 2834816 | BCG_2573c | BCG_2573c | 246 | 2814673 | 2814918 | BCGDan_2595 | BCGDan_2595 | 246 | 2814689 | 2814934 | BCGDan_sapMKO_2596 | BCGDan_sapMKO_2596 | 246 |
| 2870775 | 2871194 | Rv2551c | Rv2551c | 420 | 2835227 | 2835646 | BCG_2574c | BCG_2574c | 420 | 2815329 | 2815748 | BCGDan_2596 | BCGDan_2596 | 420 | 2815345 | 2815764 | BCGDan_sapMKO_2597 | BCGDan_sapMKO_2597 | 420 |
| 2871206 | 2872015 | aroE | Rv2552c | 810 | 2835658 | 2836440 | aroE | BCG_2575c | 783 | 2815760 | 2816542 | aroE | BCGDan_2597 | 783 | 2815776 | 2816558 | aroE | BCGDan_sapMKO_2598 | 783 |
| 2872012 | 2873265 | Rv2553c | Rv2553c | 1254 | 2836463 | 2837716 | BCG_2576c | BCG_2576c | 1254 | 2816565 | 2817818 | BCGDan_2598 | BCGDan_2598 | 1254 | 2816581 | 2817678 | BCGDan_sapMKO_2599 | BCGDan_sapMKO_2599 | 1098 |
| 2873258 | 2873770 | Rv2554c | Rv2554c | 513 | 2837709 | 2838221 | BCG_2577c | BCG_2577c | 513 | 2817811 | 2818323 | BCGDan_2599 | BCGDan_2599 | 513 | 2817827 | 2818339 | BCGDan_sapMKO_2600 | BCGDan_sapMKO_2600 | 513 |
| 2873771 | 2876485 | alaS | Rv2555c | 2715 | 2838222 | 2840936 | alaS | BCG_2578c | 2715 | 2818324 | 2821038 | alaS | BCGDan_2600 | 2715 | 2818340 | 2821054 | alaS | BCGDan_sapMKO_2601 | 2715 |
| 2876576 | 2876965 | Rv2556c | Rv2556c | 390 | 2841027 | 2841416 | BCG_2579c | BCG_2579c | 390 | 2821129 | 2821518 | BCGDan_2601 | BCGDan_2601 | 390 | 2821145 | 2821534 | BCGDan_sapMKO_2602 | BCGDan_sapMKO_2602 | 390 |
| 2877072 | 2877746 | Rv2557 | Rv2557 | 675 | 2841523 | 2842197 | BCG_2580 | BCG_2580 | 675 | 2821625 | 2822299 | BCGDan_2602 | BCGDan_2602 | 675 | 2821677 | 2822315 | BCGDan_sapMKO_2603 | BCGDan_sapMKO_2603 | 639 |
| 2877831 | 2878541 | Rv2558 | Rv2558 | 711 | 2842282 | 2842992 | BCG_2581 | BCG_2581 | 711 | 2822384 | 2823094 | BCGDan_2603 | BCGDan_2603 | 711 | 2822487 | 2823110 | BCGDan_sapMKO_2604 | BCGDan_sapMKO_2604 | 624 |
| 2878571 | 2879929 | Rv2559c | Rv2559c | 1359 | 2843022 | 2844380 | BCG_2582c | BCG_2582c | 1359 | 2823124 | 2824482 | BCGDan_2604 | BCGDan_2604 | 1359 | 2823140 | 2824498 | BCGDan_sapMKO_2605 | BCGDan_sapMKO_2605 | 1359 |
| 2880075 | 2881052 | Rv2560 | Rv2560 | 978 | 2844526 | 2845503 | BCG_2583 | BCG_2583 | 978 | 2824628 | 2825605 | BCGDan_2605 | BCGDan_2605 | 978 | 2824644 | 2825621 | BCGDan_sapMKO_2606 | BCGDan_sapMKO_2606 | 978 |
| 2881409 | 2881702 | Rv2561 | Rv2561 | 294 | 2845860 | 2846597 | BCG_2584 | BCG_2584 | 738 | 2825962 | 2826699 | BCGDan_2606 | BCGDan_2606 | 738 | 2825978 | 2826715 | BCGDan_sapMKO_2607 | BCGDan_sapMKO_2607 | 738 |
| 2882290 | 2883339 | Rv2563 | Rv2563 | 1050 | 2846740 | 2847789 | BCG_2585 | BCG_2585 | 1050 | 2826842 | 2827891 | BCGDan_2607 | BCGDan_2607 | 1050 | 2826741 | 2827907 | BCGDan_sapMKO_2608 | BCGDan_sapMKO_2608 | 1167 |
| 2883342 | 2884334 | glnQ | Rv2564 | 993 | 2847792 | 2848784 | glnQ | BCG_2586 | 993 | 2827894 | 2828886 | glnQ | BCGDan_2608 | 993 | 2827910 | 2828902 | glnQ | BCGDan_sapMKO_2609 | 993 |
| 2884611 | 2886362 | Rv2565 | Rv2565 | 1752 | 2849061 | 2850812 | BCG_2587 | BCG_2587 | 1752 | 2829163 | 2830914 | BCGDan_2609 | BCGDan_2609 | 1752 | 2829179 | 2830930 | BCGDan_sapMKO_2610 | BCGDan_sapMKO_2610 | 1752 |

| | | | | | | | | | | | | | | | | | | | |
|---|---|---|---|---|---|---|---|---|---|---|---|---|---|---|---|---|---|---|---|
| 2886373 | 2889795 | Rv2566 | Rv2566 | 3423 | 2850823 | 2852424 | BCG_2588 | BCG_2588 | 1602 | 2830925 | 2832526 | BCGDan_2610 | BCGDan_2610 | 1602 | 2831022 | 2832542 | BCGDan_sapMKO_2611 | BCGDan_sapMKO_2611 | 1521 |
| NA | NA | NA | NA | NA | 2852450 | 2854243 | BCG_2589 | BCG_2589 | 1794 | 2832552 | 2834345 | BCGDan_2611 | BCGDan_2611 | 1794 | 2832472 | 2834361 | BCGDan_sapMKO_2612 | BCGDan_sapMKO_2612 | 1890 |
| 2889795 | 2892449 | Rv2567 | Rv2567 | 2655 | 2854243 | 2856897 | BCG_2590 | BCG_2590 | 2655 | 2834345 | 2836999 | BCGDan_2612 | BCGDan_2612 | 2655 | 2834361 | 2837015 | BCGDan_sapMKO_2613 | BCGDan_sapMKO_2613 | 2655 |
| 2892446 | 2893471 | Rv2568c | Rv2568c | 1026 | 2856894 | 2857919 | BCG_2591c | BCG_2591c | 1026 | 2836996 | 2838021 | BCGDan_2613 | BCGDan_2613 | 1026 | 2837012 | 2838046 | BCGDan_sapMKO_2614 | BCGDan_sapMKO_2614 | 1035 |
| 2893464 | 2894408 | Rv2569c | Rv2569c | 945 | 2857912 | 2858856 | BCG_2592c | BCG_2592c | 945 | 2838014 | 2838958 | BCGDan_2614 | BCGDan_2614 | 945 | 2838030 | 2838956 | BCGDan_sapMKO_2615 | BCGDan_sapMKO_2615 | 927 |
| 2894512 | 2894901 | Rv2570 | Rv2570 | 390 | 2858960 | 2859349 | BCG_2593 | BCG_2593 | 390 | 2839062 | 2839451 | BCGDan_2615 | BCGDan_2615 | 390 | 2839078 | 2839467 | BCGDan_sapMKO_2616 | BCGDan_sapMKO_2616 | 390 |
| 2894893 | 2895960 | Rv2571c | Rv2571c | 1068 | 2859341 | 2860408 | BCG_2594c | BCG_2594c | 1068 | 2839443 | 2840510 | BCGDan_2616 | BCGDan_2616 | 1068 | 2839459 | 2840526 | BCGDan_sapMKO_2617 | BCGDan_sapMKO_2617 | 1068 |
| 2896013 | 2897803 | aspS | Rv2572c | 1791 | 2860415 | 2862250 | aspS | BCG_2595c | 1836 | 2840517 | 2842352 | aspS | BCGDan_2617 | 1836 | 2840533 | 2842368 | aspS | BCGDan_sapMKO_2618 | 1836 |
| 2898083 | 2898783 | Rv2573 | Rv2573 | 741 | 2862490 | 2863230 | BCG_2596 | BCG_2596 | 741 | 2842592 | 2843332 | BCGDan_2618 | BCGDan_2618 | 741 | 2842521 | 2843348 | BCGDan_sapMKO_2619 | BCGDan_sapMKO_2619 | 828 |
| 2898806 | 2899309 | Rv2574 | Rv2574 | 504 | 2863253 | 2863756 | BCG_2597 | BCG_2597 | 504 | 2843355 | 2843858 | BCGDan_2619 | BCGDan_2619 | 504 | 2843371 | 2843874 | BCGDan_sapMKO_2620 | BCGDan_sapMKO_2620 | 504 |
| 2899339 | 2900220 | Rv2575 | Rv2575 | 882 | 2863786 | 2864667 | BCG_2598 | BCG_2598 | 882 | 2843888 | 2844769 | BCGDan_2620 | BCGDan_2620 | 882 | 2843904 | 2844785 | BCGDan_sapMKO_2621 | BCGDan_sapMKO_2621 | 882 |
| 2900226 | 2900690 | Rv2576c | Rv2576c | 465 | 2864673 | 2865137 | BCG_2599c | BCG_2599c | 465 | 2844775 | 2845239 | BCGDan_2621 | BCGDan_2621 | 465 | 2844791 | 2845213 | BCGDan_sapMKO_2622 | BCGDan_sapMKO_2622 | 423 |
| 2900918 | 2902507 | Rv2577 | Rv2577 | 1590 | 2865365 | 2866954 | BCG_2600 | BCG_2600 | 1590 | 2845467 | 2847056 | BCGDan_2622 | BCGDan_2622 | 1590 | 2845674 | 2847071 | BCGDan_sapMKO_2623 | BCGDan_sapMKO_2623 | 1398 |
| 2902509 | 2903531 | Rv2578c | Rv2578c | 1023 | 2866956 | 2867978 | BCG_2601c | BCG_2601c | 1023 | 2847058 | 2848080 | BCGDan_2623 | BCGDan_2623 | 1023 | 2847073 | 2848095 | BCGDan_sapMKO_2624 | BCGDan_sapMKO_2624 | 1023 |
| 2903639 | 2904541 | dhaA | Rv2579 | 903 | 2868086 | 2868988 | dhaA | BCG_2602 | 903 | 2848188 | 2849090 | dhaA | BCGDan_2624 | 903 | 2848203 | 2849105 | dhaA | BCGDan_sapMKO_2625 | 903 |
| 2904821 | 2906092 | hisS | Rv2580c | 1272 | 2869268 | 2870539 | hisS | BCG_2603c | 1272 | 2849370 | 2850641 | hisS | BCGDan_2625 | 1272 | 2849385 | 2850656 | hisS | BCGDan_sapMKO_2626 | 1272 |
| 2906089 | 2906763 | Rv2581c | Rv2581c | 675 | 2870536 | 2871210 | BCG_2604c | BCG_2604c | 675 | 2850638 | 2851312 | BCGDan_2626 | BCGDan_2626 | 675 | 2850653 | 2851327 | BCGDan_sapMKO_2627 | BCGDan_sapMKO_2627 | 675 |
| 2906814 | 2907740 | ppiB | Rv2582 | 927 | 2871261 | 2872187 | ppiB | BCG_2605 | 927 | 2851363 | 2852289 | ppiB | BCGDan_2627 | 927 | 2851426 | 2852304 | ppiB | BCGDan_sapMKO_2628 | 879 |
| 2907826 | 2910198 | relA | Rv2583c | 2373 | 2872273 | 2874645 | relA | BCG_2606c | 2373 | 2852375 | 2854747 | relA | BCGDan_2628 | 2373 | 2852390 | 2854771 | relA | BCGDan_sapMKO_2629 | 2382 |
| 2910229 | 2910900 | apt | Rv2584c | 672 | 2874676 | 2875347 | apt | BCG_2607c | 672 | 2854778 | 2855449 | apt | BCGDan_2629 | 672 | 2854793 | 2855464 | apt | BCGDan_sapMKO_2630 | 672 |
| 2911004 | 2912677 | Rv2585c | Rv2585c | 1674 | 2875451 | 2877124 | BCG_2608c | BCG_2608c | 1674 | 2855553 | 2857226 | BCGDan_2630 | BCGDan_2630 | 1674 | 2855568 | 2857241 | BCGDan_sapMKO_2631 | BCGDan_sapMKO_2631 | 1674 |
| 2912683 | 2914011 | secF | Rv2586c | 1329 | 2877130 | 2878458 | secF | BCG_2609c | 1329 | 2857232 | 2858560 | secF | BCGDan_2631 | 1329 | 2857247 | 2858575 | secF | BCGDan_sapMKO_2632 | 1329 |
| 2914015 | 2915736 | secD | Rv2587c | 1722 | 2878462 | 2880183 | secD | BCG_2610c | 1722 | 2858564 | 2860285 | secD | BCGDan_2632 | 1722 | 2858579 | 2860255 | secD | BCGDan_sapMKO_2633 | 1677 |
| 2915846 | 2916193 | yajC | Rv2588c | 348 | 2880293 | 2880640 | BCG_2611c | BCG_2611c | 348 | 2860395 | 2860742 | BCGDan_2633 | BCGDan_2633 | 348 | 2860410 | 2860757 | BCGDan_sapMKO_2634 | BCGDan_sapMKO_2634 | 348 |
| 2916360 | 2917709 | gabT | Rv2589 | 1350 | 2880807 | 2882156 | gabT | BCG_2612 | 1350 | 2860909 | 2862258 | gabT | BCGDan_2634 | 1350 | 2860924 | 2862273 | gabT | BCGDan_sapMKO_2635 | 1350 |
| 2917871 | 2921377 | fadD9 | Rv2590 | 3507 | 2882318 | 2885824 | fadD9 | BCG_2613 | 3507 | 2862420 | 2865926 | fadD9 | BCGDan_2635 | 3507 | 2862435 | 2865941 | fadD9 | BCGDan_sapMKO_2636 | 3507 |
| 2921551 | 2923182 | PE_PGRS44 | Rv2591 | 1632 | 2885998 | 2887638 | PE_PGRS44 | BCG_2614 | 1641 | 2866100 | 2867740 | PE_PGRS44 | BCGDan_2636 | 1641 | 2866142 | 2867755 | PE_PGRS44 | BCGDan_sapMKO_2637 | 1614 |
| 2923199 | 2924233 | ruvB | Rv2592c | 1035 | 2887655 | 2888689 | ruvB | BCG_2615c | 1035 | 2867757 | 2868791 | ruvB | BCGDan_2637 | 1035 | 2867772 | 2868806 | ruvB | BCGDan_sapMKO_2638 | 1035 |
| 2924230 | 2924820 | ruvA | Rv2593c | 591 | 2888686 | 2889276 | ruvA | BCG_2616c | 591 | 2868788 | 2869378 | ruvA | BCGDan_2638 | 591 | 2868803 | 2869393 | ruvA | BCGDan_sapMKO_2639 | 591 |
| 2924817 | 2925383 | ruvC | Rv2594c | 567 | 2889273 | 2889839 | ruvC | BCG_2617c | 567 | 2869375 | 2869941 | ruvC | BCGDan_2639 | 567 | 2869390 | 2869947 | ruvC | BCGDan_sapMKO_2640 | 558 |
| 2925492 | 2925737 | vapB40 | Rv2595 | 246 | 2889948 | 2890193 | BCG_2618 | BCG_2618 | 246 | 2870050 | 2870295 | BCGDan_2640 | BCGDan_2640 | 246 | 2870098 | 2870310 | BCGDan_sapMKO_2641 | BCGDan_sapMKO_2641 | 213 |
| 2925734 | 2926138 | vapC40 | Rv2596 | 405 | 2890190 | 2890594 | BCG_2619 | BCG_2619 | 405 | 2870292 | 2870696 | BCGDan_2641 | BCGDan_2641 | 405 | 2870307 | 2870711 | BCGDan_sapMKO_2642 | BCGDan_sapMKO_2642 | 405 |
| 2926355 | 2926975 | Rv2597 | Rv2597 | 621 | 2890811 | 2891431 | BCG_2620 | BCG_2620 | 621 | 2870913 | 2871533 | BCGDan_2642 | BCGDan_2642 | 621 | 2870928 | 2871548 | BCGDan_sapMKO_2643 | BCGDan_sapMKO_2643 | 621 |
| 2926986 | 2927480 | Rv2598 | Rv2598 | 495 | 2891442 | 2891936 | BCG_2621 | BCG_2621 | 495 | 2871544 | 2872038 | BCGDan_2643 | BCGDan_2643 | 495 | 2871589 | 2872053 | BCGDan_sapMKO_2644 | BCGDan_sapMKO_2644 | 465 |
| 2927477 | 2927908 | Rv2599 | Rv2599 | 432 | 2891933 | 2892364 | BCG_2622 | BCG_2622 | 432 | 2872035 | 2872466 | BCGDan_2644 | BCGDan_2644 | 432 | 2872050 | 2872481 | BCGDan_sapMKO_2645 | BCGDan_sapMKO_2645 | 432 |
| 2927990 | 2928391 | Rv2600 | Rv2600 | 402 | 2892446 | 2892652 | BCG_2623 | BCG_2623 | 207 | 2872548 | 2872754 | BCGDan_2645 | BCGDan_2645 | 207 | 2872563 | 2872769 | BCGDan_sapMKO_2646 | BCGDan_sapMKO_2646 | 207 |
| NA | NA | NA | NA | NA | 2892658 | 2892837 | BCG_2624 | BCG_2624 | 180 | 2872760 | 2872939 | BCGDan_2646 | BCGDan_2646 | 180 | 2872775 | 2872954 | BCGDan_sapMKO_2647 | BCGDan_sapMKO_2647 | 180 |
| 2928388 | 2929959 | speE | Rv2601 | 1572 | 2892834 | 2894405 | speE | BCG_2625 | 1572 | 2872936 | 2874507 | speE | BCGDan_2647 | 1572 | 2872951 | 2874522 | speE | BCGDan_sapMKO_2648 | 1572 |
| 2930070 | 2930357 | vapB41 | Rv2601A | 288 | 2894516 | 2894803 | BCG_2626 | BCG_2626 | 288 | 2874618 | 2874905 | BCGDan_2648 | BCGDan_2648 | 288 | 2874633 | 2874920 | BCGDan_sapMKO_2649 | BCGDan_sapMKO_2649 | 288 |
| 2930344 | 2930784 | vapC41 | Rv2602 | 441 | 2894790 | 2895230 | BCG_2627 | BCG_2627 | 441 | 2874892 | 2875332 | BCGDan_2649 | BCGDan_2649 | 441 | 2874907 | 2875347 | BCGDan_sapMKO_2650 | BCGDan_sapMKO_2650 | 441 |
| 2930805 | 2931560 | Rv2603c | Rv2603c | 756 | 2895251 | 2896006 | BCG_2628c | BCG_2628c | 756 | 2875353 | 2876108 | BCGDan_2650 | BCGDan_2650 | 756 | 2875368 | 2876123 | BCGDan_sapMKO_2651 | BCGDan_sapMKO_2651 | 756 |
| 2931693 | 2932289 | snoP | Rv2604c | 597 | 2896139 | 2896735 | BCG_2629c | BCG_2629c | 597 | 2876241 | 2876837 | BCGDan_2651 | BCGDan_2651 | 597 | 2876256 | 2876852 | BCGDan_sapMKO_2652 | BCGDan_sapMKO_2652 | 597 |
| 2932297 | 2933142 | tesB2 | Rv2605c | 846 | 2896743 | 2897588 | tesB2 | BCG_2630c | 846 | 2876845 | 2877690 | tesB2 | BCGDan_2652 | 846 | 2876860 | 2877579 | tesB2 | BCGDan_sapMKO_2653 | 720 |
| 2933171 | 2934070 | snzP | Rv2606c | 900 | 2897617 | 2898516 | BCG_2631c | BCG_2631c | 900 | 2877719 | 2878618 | BCGDan_2653 | BCGDan_2653 | 900 | 2877733 | 2878632 | BCGDan_sapMKO_2654 | BCGDan_sapMKO_2654 | 900 |
| 2934198 | 2934872 | pdxH | Rv2607 | 675 | 2898644 | 2899318 | pdxH | BCG_2632 | 675 | 2878746 | 2879420 | pdxH | BCGDan_2654 | 675 | 2878760 | 2879434 | pdxH | BCGDan_sapMKO_2655 | 675 |
| 2935046 | 2936788 | PPE42 | Rv2608 | 1743 | 2899492 | 2901234 | PPE42 | BCG_2633 | 1743 | 2879594 | 2881336 | PPE42 | BCGDan_2655 | 1743 | 2879608 | 2881350 | PPE42 | BCGDan_sapMKO_2656 | 1743 |
| 2936810 | 2937865 | Rv2609c | Rv2609c | 1056 | 2901256 | 2902311 | BCG_2634c | BCG_2634c | 1056 | 2881358 | 2882413 | BCGDan_2656 | BCGDan_2656 | 1056 | 2881372 | 2882427 | BCGDan_sapMKO_2657 | BCGDan_sapMKO_2657 | 1056 |
| 2937865 | 2939001 | pimA | Rv2610c | 1137 | 2902311 | 2903447 | pimA | BCG_2635c | 1137 | 2882413 | 2883549 | pimA | BCGDan_2657 | 1137 | 2882427 | 2883551 | pimA | BCGDan_sapMKO_2658 | 1125 |
| 2939012 | 2939962 | Rv2611c | Rv2611c | 951 | 2903458 | 2904408 | BCG_2636c | BCG_2636c | 951 | 2883560 | 2884510 | BCGDan_2658 | BCGDan_2658 | 951 | 2883574 | 2884524 | BCGDan_sapMKO_2659 | BCGDan_sapMKO_2659 | 951 |
| 2939959 | 2940612 | pgsA1 | Rv2612c | 654 | 2904405 | 2905058 | pgsA1 | BCG_2637c | 654 | 2884507 | 2885160 | pgsA1 | BCGDan_2659 | 654 | 2884521 | 2885174 | pgsA1 | BCGDan_sapMKO_2660 | 654 |
| 2940609 | 2941196 | Rv2613c | Rv2613c | 588 | 2905055 | 2905642 | BCG_2638c | BCG_2638c | 588 | 2885157 | 2885744 | BCGDan_2660 | BCGDan_2660 | 588 | 2885171 | 2885758 | BCGDan_sapMKO_2661 | BCGDan_sapMKO_2661 | 588 |
| 2941189 | 2943267 | thrS | Rv2614c | 2079 | 2905635 | 2907713 | thrS | BCG_2639c | 2079 | 2885737 | 2887815 | thrS | BCGDan_2661 | 2079 | 2885751 | 2887829 | thrS | BCGDan_sapMKO_2662 | 2079 |
| 2943376 | 2943603 | Rv2614A | Rv2614A | 228 | 2907822 | 2908049 | BCG_2640 | BCG_2640 | 228 | 2887924 | 2888151 | BCGDan_2662 | BCGDan_2662 | 228 | 2887938 | 2888165 | BCGDan_sapMKO_2663 | BCGDan_sapMKO_2663 | 228 |
| NA | NA | NA | NA | NA | 2908046 | 2909215 | PE_PGRS45b | BCG_2641c | 1170 | 2888148 | 2889317 | PE_PGRS45b | BCGDan_2663 | 1170 | 2888162 | 2889331 | PE_PGRS45b | BCGDan_sapMKO_2664 | 1170 |
| 2943600 | 2944985 | PE_PGRS45 | Rv2615c | 1386 | 2909203 | 2909433 | PE_PGRS45a | BCG_2642c | 231 | 2889305 | 2889535 | PE_PGRS45a | BCGDan_2664 | 231 | 2889319 | 2889549 | PE_PGRS45a | BCGDan_sapMKO_2665 | 231 |
| 2945330 | 2945830 | Rv2616 | Rv2616 | 501 | 2909778 | 2910278 | BCG_2643 | BCG_2643 | 501 | 2889880 | 2890380 | BCGDan_2665 | BCGDan_2665 | 501 | 2889894 | 2890394 | BCGDan_sapMKO_2666 | BCGDan_sapMKO_2666 | 501 |
| 2945847 | 2946287 | Rv2617c | Rv2617c | 441 | 2910295 | 2910735 | BCG_2644c | BCG_2644c | 441 | 2890397 | 2890837 | BCGDan_2666 | BCGDan_2666 | 441 | 2890411 | 2890851 | BCGDan_sapMKO_2667 | BCGDan_sapMKO_2667 | 441 |
| 2946434 | 2947111 | Rv2618 | Rv2618 | 678 | 2910882 | 2911559 | BCG_2645 | BCG_2645 | 678 | 2890984 | 2891661 | BCGDan_2667 | BCGDan_2667 | 678 | 2890947 | 2891675 | BCGDan_sapMKO_2668 | BCGDan_sapMKO_2668 | 729 |
| 2947096 | 2947449 | Rv2619c | Rv2619c | 354 | 2911544 | 2911897 | BCG_2646c | BCG_2646c | 354 | 2891646 | 2891999 | BCGDan_2668 | BCGDan_2668 | 354 | 2891660 | 2892013 | BCGDan_sapMKO_2669 | BCGDan_sapMKO_2669 | 354 |
| 2947462 | 2947887 | Rv2620c | Rv2620c | 426 | 2911910 | 2912335 | BCG_2647c | BCG_2647c | 426 | 2892012 | 2892437 | BCGDan_2669 | BCGDan_2669 | 426 | 2892026 | 2892451 | BCGDan_sapMKO_2670 | BCGDan_sapMKO_2670 | 426 |
| 2947884 | 2948558 | Rv2621c | Rv2621c | 675 | 2912332 | 2913003 | BCG_2648c | BCG_2648c | 672 | 2892434 | 2893105 | BCGDan_2670 | BCGDan_2670 | 672 | 2892448 | 2893119 | BCGDan_sapMKO_2671 | BCGDan_sapMKO_2671 | 672 |
| 2948636 | 2949457 | Rv2622 | Rv2622 | 822 | 2913081 | 2913902 | BCG_2649 | BCG_2649 | 822 | 2893183 | 2894004 | BCGDan_2671 | BCGDan_2671 | 822 | 2893197 | 2894018 | BCGDan_sapMKO_2672 | BCGDan_sapMKO_2672 | 822 |
| 2949593 | 2950486 | TB31.7 | Rv2623 | 894 | 2914038 | 2914931 | TB31.7 | BCG_2650 | 894 | 2894140 | 2895033 | TB31.7 | BCGDan_2672 | 894 | 2894154 | 2895047 | TB31.7 | BCGDan_sapMKO_2673 | 894 |
| 2950489 | 2951307 | Rv2624c | Rv2624c | 819 | 2914934 | 2915752 | BCG_2651c | BCG_2651c | 819 | 2895036 | 2895830 | BCGDan_2673 | BCGDan_2673 | 795 | 2895050 | 2895844 | BCGDan_sapMKO_2674 | BCGDan_sapMKO_2674 | 795 |
| 2951322 | 2952503 | Rv2625c | Rv2625c | 1182 | 2915767 | 2916948 | BCG_2652c | BCG_2652c | 1182 | 2895869 | 2897050 | BCGDan_2674 | BCGDan_2674 | 1182 | 2895883 | 2897064 | BCGDan_sapMKO_2675 | BCGDan_sapMKO_2675 | 1182 |
| 2952562 | 2952993 | hrp1 | Rv2626c | 432 | 2917007 | 2917438 | BCG_2653c | BCG_2653c | 432 | 2897109 | 2897540 | BCGDan_2675 | BCGDan_2675 | 432 | 2897123 | 2897554 | BCGDan_sapMKO_2676 | BCGDan_sapMKO_2676 | 432 |
| 2953507 | 2954748 | Rv2627c | Rv2627c | 1242 | 2917952 | 2919193 | BCG_2654c | BCG_2654c | 1242 | 2898054 | 2899295 | BCGDan_2676 | BCGDan_2676 | 1242 | 2898068 | 2899309 | BCGDan_sapMKO_2677 | BCGDan_sapMKO_2677 | 1242 |
| 2955058 | 2955420 | Rv2628 | Rv2628 | 363 | 2919503 | 2919865 | BCG_2655 | BCG_2655 | 363 | 2899605 | 2899967 | BCGDan_2677 | BCGDan_2677 | 363 | 2899619 | 2899981 | BCGDan_sapMKO_2678 | BCGDan_sapMKO_2678 | 363 |
| 2955767 | 2956891 | Rv2629 | Rv2629 | 1125 | 2920212 | 2921336 | BCG_2656 | BCG_2656 | 1125 | 2900314 | 2901438 | BCGDan_2678 | BCGDan_2678 | 1125 | 2900328 | 2901452 | BCGDan_sapMKO_2679 | BCGDan_sapMKO_2679 | 1125 |
| 2956893 | 2957432 | Rv2630 | Rv2630 | 540 | 2921338 | 2921877 | BCG_2657 | BCG_2657 | 540 | 2901440 | 2901979 | BCGDan_2679 | BCGDan_2679 | 540 | 2901454 | 2901993 | BCGDan_sapMKO_2680 | BCGDan_sapMKO_2680 | 540 |
| 2957572 | 2958870 | Rv2631 | Rv2631 | 1299 | 2922016 | 2923314 | BCG_2658 | BCG_2658 | 1299 | 2902118 | 2903416 | BCGDan_2680 | BCGDan_2680 | 1299 | 2902009 | 2903430 | BCGDan_sapMKO_2681 | BCGDan_sapMKO_2681 | 1422 |
| 2958909 | 2959190 | Rv2632c | Rv2632c | 282 | 2923353 | 2923634 | BCG_2659c | BCG_2659c | 282 | 2903455 | 2903736 | BCGDan_2681 | BCGDan_2681 | 282 | 2903469 | 2903750 | BCGDan_sapMKO_2682 | BCGDan_sapMKO_2682 | 282 |
| 2959335 | 2959820 | Rv2633c | Rv2633c | 486 | 2923779 | 2924264 | BCG_2660c | BCG_2660c | 486 | 2903881 | 2904366 | BCGDan_2682 | BCGDan_2682 | 486 | 2903895 | 2904380 | BCGDan_sapMKO_2683 | BCGDan_sapMKO_2683 | 486 |
| 2960105 | 2962441 | PE_PGRS46 | Rv2634c | 2337 | 2924549 | 2926951 | PE_PGRS46 | BCG_2661c | 2403 | 2904651 | 2907053 | PE_PGRS46 | BCGDan_2683 | 2403 | 2904665 | 2907067 | PE_PGRS46 | BCGDan_sapMKO_2684 | 2403 |
| 2962470 | 2962712 | Rv2635 | Rv2635 | 243 | 2926980 | 2927222 | BCG_2662 | BCG_2662 | 243 | 2907082 | 2907324 | BCGDan_2684 | BCGDan_2684 | 243 | 2907096 | 2907338 | BCGDan_sapMKO_2685 | BCGDan_sapMKO_2685 | 243 |
| 2962713 | 2963390 | Rv2636 | Rv2636 | 678 | 2927223 | 2927900 | BCG_2663 | BCG_2663 | 678 | 2907325 | 2908002 | BCGDan_2685 | BCGDan_2685 | 678 | 2907387 | 2908016 | BCGDan_sapMKO_2686 | BCGDan_sapMKO_2686 | 630 |
| 2963586 | 2964242 | dedA | Rv2637 | 657 | 2928096 | 2928752 | dedA | BCG_2664 | 657 | 2908198 | 2908854 | dedA | BCGDan_2686 | 657 | 2908212 | 2908868 | dedA | BCGDan_sapMKO_2687 | 657 |
| 2964405 | 2964851 | Rv2638 | Rv2638 | 447 | 2928915 | 2929361 | BCG_2665 | BCG_2665 | 447 | 2909017 | 2909463 | BCGDan_2687 | BCGDan_2687 | 447 | 2909031 | 2909477 | BCGDan_sapMKO_2688 | BCGDan_sapMKO_2688 | 447 |
| 2965026 | 2965358 | Rv2639c | Rv2639c | 333 | 2929536 | 2929868 | BCG_2666c | BCG_2666c | 333 | 2909638 | 2909970 | BCGDan_2688 | BCGDan_2688 | 333 | 2909652 | 2909984 | BCGDan_sapMKO_2689 | BCGDan_sapMKO_2689 | 333 |
| 2965478 | 2965837 | Rv2640c | Rv2640c | 360 | 2929988 | 2930347 | BCG_2667c | BCG_2667c | 360 | 2910090 | 2910449 | BCGDan_2689 | BCGDan_2689 | 360 | 2910104 | 2910463 | BCGDan_sapMKO_2690 | BCGDan_sapMKO_2690 | 360 |
| 2965939 | 2966397 | cadI | Rv2641 | 459 | 2930449 | 2930907 | cadI | BCG_2668 | 459 | 2910551 | 2911009 | cadI | BCGDan_2690 | 459 | 2910565 | 2911023 | cadI | BCGDan_sapMKO_2691 | 459 |
| 2966533 | 2966913 | Rv2642 | Rv2642 | 381 | 2931043 | 2931423 | BCG_2669 | BCG_2669 | 381 | 2911145 | 2911525 | BCGDan_2691 | BCGDan_2691 | 381 | 2911159 | 2911539 | BCGDan_sapMKO_2692 | BCGDan_sapMKO_2692 | 381 |
| 2966910 | 2968406 | arsC | Rv2643 | 1497 | 2931420 | 2932916 | arsC | BCG_2670 | 1497 | 2911522 | 2913018 | arsC | BCGDan_2692 | 1497 | 2911536 | 2913032 | arsC | BCGDan_sapMKO_2693 | 1497 |
| 2968533 | 2968850 | Rv2644c | Rv2644c | 318 | 2933043 | 2933360 | BCG_2671c | BCG_2671c | 318 | 2913145 | 2913462 | BCGDan_2693 | BCGDan_2693 | 318 | 2913159 | 2913476 | BCGDan_sapMKO_2694 | BCGDan_sapMKO_2694 | 318 |
| 2980963 | 2981190 | Rv2660c | Rv2660c | 228 | 2934497 | 2934718 | BCG_2672c | BCG_2672c | 222 | 2914599 | 2914820 | BCGDan_2698 | BCGDan_2698 | 222 | 2914613 | 2914834 | BCGDan_sapMKO_2699 | BCGDan_sapMKO_2699 | 222 |
| 2981187 | 2981576 | Rv2661c | Rv2661c | 390 | 2934715 | 2935104 | BCG_2673c | BCG_2673c | 390 | 2914817 | 2915206 | BCGDan_2699 | BCGDan_2699 | 390 | 2914831 | 2915220 | BCGDan_sapMKO_2700 | BCGDan_sapMKO_2700 | 390 |

| | | | | | | | | | | | | | | | | | | | |
|---|---|---|---|---|---|---|---|---|---|---|---|---|---|---|---|---|---|---|---|
| 2981482 | 2981754 | Rv2662 | Rv2662 | 273 | 2935010 | 2935282 | BCG_2674 | BCG_2674 | 273 | 2915112 | 2915384 | BCGDan_2700 | BCGDan_2700 | 273 | 2915126 | 2915398 | BCGDan_sapMKO_2701 | BCGDan_sapMKO_2701 | 273 |
| 2981853 | 2982086 | Rv2663 | Rv2663 | 234 | 2935381 | 2935614 | BCG_2675 | BCG_2675 | 234 | 2915483 | 2915716 | BCGDan_2701 | BCGDan_2701 | 234 | 2915497 | 2915730 | BCGDan_sapMKO_2702 | BCGDan_sapMKO_2702 | 234 |
| 2982097 | 2982351 | Rv2664 | Rv2664 | 255 | 2935625 | 2935879 | BCG_2676 | BCG_2676 | 255 | 2915727 | 2915981 | BCGDan_2702 | BCGDan_2702 | 255 | 2915741 | 2915995 | BCGDan_sapMKO_2703 | BCGDan_sapMKO_2703 | 255 |
| NA | NA | NA | NA | NA | 2936108 | 2936599 | BCG_2677c | BCG_2677c | 492 | 2916210 | 2916701 | BCGDan_2703 | BCGDan_2703 | 492 | 2916224 | 2916715 | BCGDan_sapMKO_2704 | BCGDan_sapMKO_2704 | 492 |
| 2982699 | 2982980 | Rv2665 | Rv2665 | 282 | 2936227 | 2936508 | BCG_2678 | BCG_2678 | 282 | 2916329 | 2916610 | BCGDan_2704 | BCGDan_2704 | 282 | 2916343 | 2916624 | BCGDan_sapMKO_2705 | BCGDan_sapMKO_2705 | 282 |
| 2983071 | 2983874 | Rv2666 | Rv2666 | 804 | 2936599 | 2937402 | BCG_2679 | BCG_2679 | 804 | 2916701 | 2917504 | BCGDan_2705 | BCGDan_2705 | 804 | 2916652 | 2917518 | BCGDan_sapMKO_2706 | BCGDan_sapMKO_2706 | 867 |
| 2983896 | 2984654 | clpC2 | Rv2667 | 759 | 2937424 | 2938182 | clpX' | BCG_2680 | 759 | 2917526 | 2918284 | clpX | BCGDan_2706 | 759 | 2917540 | 2918298 | clpX_2 | BCGDan_sapMKO_2707 | 759 |
| 2984733 | 2985254 | Rv2668 | Rv2668 | 522 | 2938261 | 2938782 | BCG_2681 | BCG_2681 | 522 | 2918363 | 2918884 | BCGDan_2707 | BCGDan_2707 | 522 | 2918332 | 2918898 | BCGDan_sapMKO_2708 | BCGDan_sapMKO_2708 | 567 |
| 2985283 | 2985753 | Rv2669 | Rv2669 | 471 | 2938811 | 2939281 | BCG_2682 | BCG_2682 | 471 | 2918913 | 2919383 | BCGDan_2708 | BCGDan_2708 | 471 | 2918927 | 2919397 | BCGDan_sapMKO_2709 | BCGDan_sapMKO_2709 | 471 |
| 2985731 | 2986840 | Rv2670c | Rv2670c | 1110 | 2939259 | 2940368 | BCG_2683c | BCG_2683c | 1110 | 2919361 | 2920470 | BCGDan_2709 | BCGDan_2709 | 1110 | 2919375 | 2920484 | BCGDan_sapMKO_2710 | BCGDan_sapMKO_2710 | 1110 |
| 2986839 | 2987615 | ribD | Rv2671 | 777 | 2940367 | 2941143 | ribD | BCG_2684 | 777 | 2920469 | 2921245 | ribD | BCGDan_2710 | 777 | 2920483 | 2921259 | ribD | BCGDan_sapMKO_2711 | 777 |
| 2987682 | 2989268 | Rv2672 | Rv2672 | 1587 | 2941210 | 2942796 | BCG_2685 | BCG_2685 | 1587 | 2921312 | 2922898 | BCGDan_2711 | BCGDan_2711 | 1587 | 2921356 | 2922912 | BCGDan_sapMKO_2712 | BCGDan_sapMKO_2712 | 1557 |
| 2989291 | 2990592 | aftC | Rv2673 | 1302 | 2942819 | 2944120 | BCG_2686 | BCG_2686 | 1302 | 2922921 | 2924222 | BCGDan_2712 | BCGDan_2712 | 1302 | 2922935 | 2924236 | BCGDan_sapMKO_2713 | BCGDan_sapMKO_2713 | 1302 |
| 2990706 | 2991116 | msrB | Rv2674 | 411 | 2944289 | 2944699 | BCG_2687 | BCG_2687 | 411 | 2924391 | 2924801 | BCGDan_2713 | BCGDan_2713 | 411 | 2924405 | 2924815 | BCGDan_sapMKO_2714 | BCGDan_sapMKO_2714 | 411 |
| 2991184 | 2991936 | Rv2675c | Rv2675c | 753 | 2944767 | 2945519 | BCG_2688c | BCG_2688c | 753 | 2924869 | 2925621 | BCGDan_2714 | BCGDan_2714 | 753 | 2924883 | 2925635 | BCGDan_sapMKO_2715 | BCGDan_sapMKO_2715 | 753 |
| 2991933 | 2992628 | Rv2676c | Rv2676c | 696 | 2945516 | 2946211 | BCG_2689c | BCG_2689c | 696 | 2925618 | 2926313 | BCGDan_2715 | BCGDan_2715 | 696 | 2925632 | 2926327 | BCGDan_sapMKO_2716 | BCGDan_sapMKO_2716 | 696 |
| 2992634 | 2993992 | hemY | Rv2677c | 1359 | 2946217 | 2947575 | hemY | BCG_2690c | 1359 | 2926319 | 2927677 | hemY | BCGDan_2716 | 1359 | 2926333 | 2927646 | hemY | BCGDan_sapMKO_2717 | 1314 |
| 2993989 | 2995062 | hemE | Rv2678c | 1074 | 2947572 | 2948645 | hemE | BCG_2691c | 1074 | 2927674 | 2928747 | hemE | BCGDan_2717 | 1074 | 2927688 | 2928788 | hemE | BCGDan_sapMKO_2718 | 1101 |
| 2995115 | 2995945 | echA15 | Rv2679 | 831 | 2948698 | 2949528 | BCG_2692 | BCG_2692 | 831 | 2928800 | 2929630 | BCGDan_2718 | BCGDan_2718 | 831 | 2928814 | 2929644 | BCGDan_sapMKO_2719 | BCGDan_sapMKO_2719 | 831 |
| 2996105 | 2996737 | Rv2680 | Rv2680 | 633 | 2949790 | 2950422 | BCG_2693 | BCG_2693 | 633 | 2929790 | 2930422 | BCGDan_2719 | BCGDan_2719 | 633 | 2929855 | 2930487 | BCGDan_sapMKO_2720 | BCGDan_sapMKO_2720 | 633 |
| 2996739 | 2998055 | Rv2681 | Rv2681 | 1317 | 2950424 | 2951740 | BCG_2694 | BCG_2694 | 1317 | 2930424 | 2931740 | BCGDan_2720 | BCGDan_2720 | 1317 | 2930489 | 2931805 | BCGDan_sapMKO_2721 | BCGDan_sapMKO_2721 | 1317 |
| 2998052 | 2999968 | dxs1 | Rv2682c | 1917 | 2951737 | 2953653 | dxs1 | BCG_2695c | 1917 | 2931737 | 2933653 | dxs1 | BCGDan_2721 | 1917 | 2931802 | 2933718 | dxs1 | BCGDan_sapMKO_2722 | 1917 |
| 3000112 | 3000609 | Rv2683 | Rv2683 | 498 | 2953797 | 2954294 | BCG_2696 | BCG_2696 | 498 | 2933797 | 2934294 | BCGDan_2722 | BCGDan_2722 | 498 | 2933862 | 2934359 | BCGDan_sapMKO_2723 | BCGDan_sapMKO_2723 | 498 |
| 3000614 | 3001903 | arsA | Rv2684 | 1290 | 2954299 | 2955588 | arsA | BCG_2697 | 1290 | 2934299 | 2935588 | arsA | BCGDan_2723 | 1290 | 2934364 | 2935653 | arsA | BCGDan_sapMKO_2724 | 1290 |
| 3001983 | 3003269 | arsB1 | Rv2685 | 1287 | 2955668 | 2956954 | arsB1 | BCG_2698 | 1287 | 2935668 | 2936954 | arsB1 | BCGDan_2724 | 1287 | 2935733 | 2937019 | arsB1 | BCGDan_sapMKO_2725 | 1287 |
| 3003280 | 3004038 | Rv2686c | Rv2686c | 759 | 2956965 | 2957723 | BCG_2699c | BCG_2699c | 759 | 2936965 | 2937723 | BCGDan_2725 | BCGDan_2725 | 759 | 2937030 | 2937752 | BCGDan_sapMKO_2726 | BCGDan_sapMKO_2726 | 723 |
| 3004035 | 3004748 | Rv2687c | Rv2687c | 714 | 2957720 | 2958433 | BCG_2700c | BCG_2700c | 714 | 2937720 | 2938433 | BCGDan_2726 | BCGDan_2726 | 714 | 2937785 | 2938498 | BCGDan_sapMKO_2727 | BCGDan_sapMKO_2727 | 714 |
| 3004745 | 3005650 | Rv2688c | Rv2688c | 906 | 2958430 | 2959335 | BCG_2701c | BCG_2701c | 906 | 2938430 | 2939335 | BCGDan_2727 | BCGDan_2727 | 906 | 2938495 | 2939445 | BCGDan_sapMKO_2728 | BCGDan_sapMKO_2728 | 951 |
| 3005845 | 3007062 | Rv2689c | Rv2689c | 1218 | 2959530 | 2960747 | BCG_2702c | BCG_2702c | 1218 | 2939530 | 2940747 | BCGDan_2728 | BCGDan_2728 | 1218 | 2939595 | 2940812 | BCGDan_sapMKO_2729 | BCGDan_sapMKO_2729 | 1218 |
| 3007236 | 3009209 | Rv2690c | Rv2690c | 1974 | 2960903 | 2962894 | BCG_2703c | BCG_2703c | 1992 | 2940850 | 2942841 | BCGDan_2729 | BCGDan_2729 | 1992 | 2940915 | 2942906 | BCGDan_sapMKO_2730 | BCGDan_sapMKO_2730 | 1992 |
| 3009344 | 3010027 | ceoB | Rv2691 | 684 | 2963029 | 2963712 | ceoB | BCG_2704 | 684 | 2942976 | 2943659 | ceoB | BCGDan_2730 | 684 | 2943041 | 2943724 | ceoB | BCGDan_sapMKO_2731 | 684 |
| 3010024 | 3010686 | ceoC | Rv2692 | 663 | 2963709 | 2964371 | ceoC | BCG_2705 | 663 | 2943656 | 2944318 | ceoC | BCGDan_2731 | 663 | 2943721 | 2944383 | ceoC | BCGDan_sapMKO_2732 | 663 |
| 3010697 | 3011368 | Rv2693c | Rv2693c | 672 | 2964382 | 2965053 | BCG_2706c | BCG_2706c | 672 | 2944329 | 2945000 | BCGDan_2732 | BCGDan_2732 | 672 | 2944394 | 2945065 | BCGDan_sapMKO_2733 | BCGDan_sapMKO_2733 | 672 |
| 3011399 | 3011767 | Rv2694c | Rv2694c | 369 | 2965084 | 2965452 | BCG_2707c | BCG_2707c | 369 | 2945031 | 2945399 | BCGDan_2733 | BCGDan_2733 | 369 | 2945096 | 2945464 | BCGDan_sapMKO_2734 | BCGDan_sapMKO_2734 | 369 |
| 3011916 | 3012623 | Rv2695 | Rv2695 | 708 | 2965601 | 2966308 | BCG_2708 | BCG_2708 | 708 | 2945548 | 2946255 | BCGDan_2734 | BCGDan_2734 | 708 | 2945613 | 2946320 | BCGDan_sapMKO_2735 | BCGDan_sapMKO_2735 | 708 |
| 3012829 | 3013608 | Rv2696c | Rv2696c | 780 | 2966514 | 2967293 | BCG_2709c | BCG_2709c | 780 | 2946461 | 2947240 | BCGDan_2735 | BCGDan_2735 | 780 | 2946526 | 2947305 | BCGDan_sapMKO_2736 | BCGDan_sapMKO_2736 | 780 |
| 3013683 | 3014147 | dut | Rv2697c | 465 | 2967368 | 2967832 | dut | BCG_2710c | 465 | 2947315 | 2947779 | dut | BCGDan_2736 | 465 | 2947380 | 2947844 | dut | BCGDan_sapMKO_2737 | 465 |
| 3014173 | 3014658 | Rv2698 | Rv2698 | 486 | 2967858 | 2968343 | BCG_2711 | BCG_2711 | 486 | 2947805 | 2948290 | BCGDan_2737 | BCGDan_2737 | 486 | 2947900 | 2948355 | BCGDan_sapMKO_2738 | BCGDan_sapMKO_2738 | 456 |
| 3014663 | 3014965 | Rv2699c | Rv2699c | 303 | 2968348 | 2968650 | BCG_2712c | BCG_2712c | 303 | 2948295 | 2948597 | BCGDan_2738 | BCGDan_2738 | 303 | 2948360 | 2948662 | BCGDan_sapMKO_2739 | BCGDan_sapMKO_2739 | 303 |
| 3015203 | 3015853 | Rv2700 | Rv2700 | 651 | 2968888 | 2969538 | BCG_2713 | BCG_2713 | 651 | 2948835 | 2949485 | BCGDan_2739 | BCGDan_2739 | 651 | 2948900 | 2949550 | BCGDan_sapMKO_2740 | BCGDan_sapMKO_2740 | 651 |
| 3015863 | 3016735 | suhB | Rv2701c | 873 | 2969548 | 2970420 | suhB | BCG_2714c | 873 | 2949495 | 2950367 | suhB | BCGDan_2740 | 873 | 2949560 | 2950432 | suhB | BCGDan_sapMKO_2741 | 873 |
| 3016858 | 3017655 | ppgK | Rv2702 | 798 | 2970543 | 2971340 | ppgK | BCG_2715 | 798 | 2950490 | 2951287 | ppgK | BCGDan_2741 | 798 | 2950555 | 2951352 | ppgK | BCGDan_sapMKO_2742 | 798 |
| 3017835 | 3019421 | sigA | Rv2703 | 1587 | 2971520 | 2973106 | sigA | BCG_2716 | 1587 | 2951467 | 2953053 | sigA | BCGDan_2742 | 1587 | 2951526 | 2953118 | sigA | BCGDan_sapMKO_2743 | 1593 |
| 3019458 | 3019886 | Rv2704 | Rv2704 | 429 | 2973143 | 2973571 | BCG_2717 | BCG_2717 | 429 | 2953090 | 2953518 | BCGDan_2743 | BCGDan_2743 | 429 | 2953173 | 2953583 | BCGDan_sapMKO_2744 | BCGDan_sapMKO_2744 | 411 |
| 3019814 | 3020203 | Rv2705c | Rv2705c | 390 | 2973499 | 2973888 | BCG_2718c | BCG_2718c | 390 | 2953446 | 2953835 | BCGDan_2744 | BCGDan_2744 | 390 | 2953511 | 2953900 | BCGDan_sapMKO_2745 | BCGDan_sapMKO_2745 | 390 |
| 3020200 | 3020457 | Rv2706c | Rv2706c | 258 | 2973885 | 2974142 | BCG_2719c | BCG_2719c | 258 | 2953832 | 2954089 | BCGDan_2745 | BCGDan_2745 | 258 | 2953897 | 2954154 | BCGDan_sapMKO_2746 | BCGDan_sapMKO_2746 | 258 |
| 3020573 | 3021547 | Rv2707 | Rv2707 | 975 | 2974258 | 2975232 | BCG_2720 | BCG_2720 | 975 | 2954205 | 2955179 | BCGDan_2746 | BCGDan_2746 | 975 | 2954270 | 2955244 | BCGDan_sapMKO_2747 | BCGDan_sapMKO_2747 | 975 |
| 3021548 | 3021796 | Rv2708c | Rv2708c | 249 | 2975233 | 2975481 | BCG_2721c | BCG_2721c | 249 | 2955180 | 2955428 | BCGDan_2747 | BCGDan_2747 | 249 | 2955245 | 2955484 | BCGDan_sapMKO_2748 | BCGDan_sapMKO_2748 | 240 |
| 3021839 | 3022285 | Rv2709 | Rv2709 | 447 | 2975524 | 2975970 | BCG_2722 | BCG_2722 | 447 | 2955471 | 2955917 | BCGDan_2748 | BCGDan_2748 | 447 | 2955554 | 2955982 | BCGDan_sapMKO_2749 | BCGDan_sapMKO_2749 | 429 |
| 3022461 | 3023432 | sigB | Rv2710 | 972 | 2976146 | 2977117 | sigB | BCG_2723 | 972 | 2956093 | 2957064 | sigB | BCGDan_2749 | 972 | 2956128 | 2957129 | sigB | BCGDan_sapMKO_2750 | 1002 |
| 3023565 | 3024257 | ideR | Rv2711 | 693 | 2977250 | 2977942 | ideR | BCG_2724 | 693 | 2957197 | 2957889 | ideR | BCGDan_2750 | 693 | 2957262 | 2957954 | ideR | BCGDan_sapMKO_2751 | 693 |
| 3024270 | 3025328 | Rv2712c | Rv2712c | 1059 | 2977955 | 2979013 | BCG_2725c | BCG_2725c | 1059 | 2957902 | 2958960 | BCGDan_2751 | BCGDan_2751 | 1059 | 2957967 | 2959049 | BCGDan_sapMKO_2752 | BCGDan_sapMKO_2752 | 1083 |
| 3025441 | 3026847 | sthA | Rv2713 | 1407 | 2979126 | 2980532 | sthA | BCG_2726 | 1407 | 2959073 | 2960479 | sthA | BCGDan_2752 | 1407 | 2959129 | 2960544 | sthA | BCGDan_sapMKO_2753 | 1416 |
| 3027065 | 3028039 | Rv2714 | Rv2714 | 975 | 2980750 | 2981724 | BCG_2727 | BCG_2727 | 975 | 2960697 | 2961671 | BCGDan_2753 | BCGDan_2753 | 975 | 2960762 | 2961736 | BCGDan_sapMKO_2754 | BCGDan_sapMKO_2754 | 975 |
| 3028098 | 3029123 | Rv2715 | Rv2715 | 1026 | 2981783 | 2982808 | BCG_2728 | BCG_2728 | 1026 | 2961730 | 2962755 | BCGDan_2754 | BCGDan_2754 | 1026 | 2961825 | 2962820 | BCGDan_sapMKO_2755 | BCGDan_sapMKO_2755 | 996 |
| 3029172 | 3029858 | Rv2716 | Rv2716 | 687 | 2982857 | 2983543 | BCG_2729 | BCG_2729 | 687 | 2962804 | 2963490 | BCGDan_2755 | BCGDan_2755 | 687 | 2962869 | 2963555 | BCGDan_sapMKO_2756 | BCGDan_sapMKO_2756 | 687 |
| 3029867 | 3030361 | Rv2717c | Rv2717c | 495 | 2983552 | 2984046 | BCG_2730c | BCG_2730c | 495 | 2963499 | 2963993 | BCGDan_2756 | BCGDan_2756 | 495 | 2963564 | 2964082 | BCGDan_sapMKO_2757 | BCGDan_sapMKO_2757 | 519 |
| 3030413 | 3030877 | nrdR | Rv2718c | 465 | 2984098 | 2984562 | BCG_2731c | BCG_2731c | 465 | 2964045 | 2964509 | BCGDan_2757 | BCGDan_2757 | 465 | 2964110 | 2964538 | BCGDan_sapMKO_2758 | BCGDan_sapMKO_2758 | 429 |
| 3031040 | 3031537 | Rv2719c | Rv2719c | 498 | 2984725 | 2985222 | BCG_2732c | BCG_2732c | 498 | 2964672 | 2965169 | BCGDan_2758 | BCGDan_2758 | 498 | 2964737 | 2965234 | BCGDan_sapMKO_2759 | BCGDan_sapMKO_2759 | 498 |
| 3031788 | 3032498 | lexA | Rv2720 | 711 | 2985530 | 2986183 | lexA | BCG_2733 | 654 | 2965477 | 2966130 | lexA | BCGDan_2759 | 654 | 2965485 | 2966195 | lexA | BCGDan_sapMKO_2760 | 711 |
| 3032520 | 3034619 | Rv2721c | Rv2721c | 2100 | 2986205 | 2988304 | BCG_2734c | BCG_2734c | 2100 | 2966152 | 2968251 | BCGDan_2760 | BCGDan_2760 | 2100 | 2966217 | 2968295 | BCGDan_sapMKO_2761 | BCGDan_sapMKO_2761 | 2079 |
| 3034635 | 3034883 | Rv2722 | Rv2722 | 249 | 2988320 | 2988568 | BCG_2735 | BCG_2735 | 249 | 2968267 | 2968515 | BCGDan_2761 | BCGDan_2761 | 249 | 2968332 | 2968580 | BCGDan_sapMKO_2762 | BCGDan_sapMKO_2762 | 249 |
| 3034909 | 3036102 | Rv2723 | Rv2723 | 1194 | 2988594 | 2989787 | BCG_2736 | BCG_2736 | 1194 | 2968541 | 2969734 | BCGDan_2762 | BCGDan_2762 | 1194 | 2968606 | 2969799 | BCGDan_sapMKO_2763 | BCGDan_sapMKO_2763 | 1194 |
| 3036131 | 3037291 | fadE20 | Rv2724c | 1161 | 2989816 | 2990976 | fadE20 | BCG_2737c | 1161 | 2969763 | 2970923 | fadE20 | BCGDan_2763 | 1161 | 2969828 | 2970988 | fadE20 | BCGDan_sapMKO_2764 | 1161 |
| 3037427 | 3038914 | hflX | Rv2725c | 1488 | 2991112 | 2992599 | hflX | BCG_2738c | 1488 | 2971059 | 2972489 | hflX | BCGDan_2764 | 1431 | 2971124 | 2972554 | hflX | BCGDan_sapMKO_2765 | 1431 |
| 3038931 | 3039800 | dapF | Rv2726c | 870 | 2992616 | 2993485 | dapF | BCG_2739c | 870 | 2972563 | 2973432 | dapF | BCGDan_2765 | 870 | 2972628 | 2973506 | dapF | BCGDan_sapMKO_2766 | 879 |
| 3039825 | 3040769 | miaA | Rv2727c | 945 | 2993510 | 2994454 | miaA | BCG_2740c | 945 | 2973457 | 2974401 | miaA | BCGDan_2766 | 945 | 2973522 | 2974466 | miaA | BCGDan_sapMKO_2767 | 945 |
| 3040766 | 3041461 | Rv2728c | Rv2728c | 696 | 2994451 | 2995146 | BCG_2741c | BCG_2741c | 696 | 2974398 | 2975093 | BCGDan_2767 | BCGDan_2767 | 696 | 2974463 | 2975158 | BCGDan_sapMKO_2768 | BCGDan_sapMKO_2768 | 696 |
| 3041570 | 3042475 | Rv2729c | Rv2729c | 906 | 2995255 | 2996160 | BCG_2742c | BCG_2742c | 906 | 2975202 | 2976107 | BCGDan_2768 | BCGDan_2768 | 906 | 2975267 | 2976172 | BCGDan_sapMKO_2769 | BCGDan_sapMKO_2769 | 906 |
| 3042542 | 3043018 | Rv2730 | Rv2730 | 477 | 2996227 | 2996703 | BCG_2743 | BCG_2743 | 477 | 2976174 | 2976650 | BCGDan_2769 | BCGDan_2769 | 477 | 2976239 | 2976715 | BCGDan_sapMKO_2770 | BCGDan_sapMKO_2770 | 477 |
| 3043026 | 3044378 | Rv2731 | Rv2731 | 1353 | 2996711 | 2998063 | BCG_2744 | BCG_2744 | 1353 | 2976658 | 2978010 | BCGDan_2770 | BCGDan_2770 | 1353 | 2976678 | 2978075 | BCGDan_sapMKO_2771 | BCGDan_sapMKO_2771 | 1398 |
| 3044375 | 3044989 | Rv2732c | Rv2732c | 615 | 2998060 | 2998674 | BCG_2745c | BCG_2745c | 615 | 2978007 | 2978621 | BCGDan_2771 | BCGDan_2771 | 615 | 2978072 | 2978692 | BCGDan_sapMKO_2772 | BCGDan_sapMKO_2772 | 621 |
| 3044986 | 3046524 | Rv2733c | Rv2733c | 1539 | 2998671 | 3000209 | BCG_2746c | BCG_2746c | 1539 | 2978618 | 2980156 | BCGDan_2772 | BCGDan_2772 | 1539 | 2978683 | 2980218 | BCGDan_sapMKO_2773 | BCGDan_sapMKO_2773 | 1536 |
| 3046821 | 3047675 | Rv2734 | Rv2734 | 855 | 3000506 | 3001360 | BCG_2747 | BCG_2747 | 855 | 2980453 | 2981307 | BCGDan_2773 | BCGDan_2773 | 855 | 2980518 | 2981372 | BCGDan_sapMKO_2774 | BCGDan_sapMKO_2774 | 855 |
| 3047560 | 3048552 | Rv2735c | Rv2735c | 993 | 3001245 | 3002237 | BCG_2748c | BCG_2748c | 993 | 2981192 | 2982184 | BCGDan_2774 | BCGDan_2774 | 993 | 2981257 | 2982249 | BCGDan_sapMKO_2775 | BCGDan_sapMKO_2775 | 993 |
| 3048562 | 3049086 | recX | Rv2736c | 525 | 3002247 | 3002771 | recX | BCG_2749c | 525 | 2982194 | 2982718 | recX | BCGDan_2775 | 525 | 2982259 | 2982783 | recX | BCGDan_sapMKO_2776 | 525 |
| 3049052 | 3051424 | recA | Rv2737c | 2373 | 3002737 | 3005109 | recA | BCG_2750c | 2373 | 2982684 | 2985056 | recA | BCGDan_2776 | 2373 | 2982749 | 2985121 | recA | BCGDan_sapMKO_2777 | 2373 |
| 3051619 | 3051792 | Rv2737A | Rv2737A | 174 | 3005304 | 3005477 | BCG_2751 | BCG_2751 | 174 | 2985251 | 2985424 | BCGDan_2777 | BCGDan_2777 | 174 | 2985316 | 2985489 | BCGDan_sapMKO_2778 | BCGDan_sapMKO_2778 | 174 |
| 3051806 | 3052012 | Rv2738c | Rv2738c | 207 | 3005491 | 3005697 | BCG_2752c | BCG_2752c | 207 | 2985438 | 2985644 | BCGDan_2778 | BCGDan_2778 | 207 | 2985503 | 2985709 | BCGDan_sapMKO_2779 | BCGDan_sapMKO_2779 | 207 |
| 3052023 | 3053189 | Rv2739c | Rv2739c | 1167 | 3005708 | 3006874 | BCG_2753c | BCG_2753c | 1167 | 2985655 | 2986821 | BCGDan_2779 | BCGDan_2779 | 1167 | 2985720 | 2986886 | BCGDan_sapMKO_2780 | BCGDan_sapMKO_2780 | 1167 |
| 3053233 | 3053682 | ephG | Rv2740 | 450 | 3006918 | 3007367 | ephG | BCG_2754 | 450 | 2986865 | 2987314 | ephG | BCGDan_2780 | 450 | 2986930 | 2987379 | ephG | BCGDan_sapMKO_2781 | 450 |
| 3053914 | 3055491 | PE_PGRS47 | Rv2741 | 1578 | 3007643 | 3008917 | PE_PGRS47 | BCG_2756 | 1275 | 2987590 | 2988864 | PE_PGRS47 | BCGDan_2781 | 1275 | 2987655 | 2988929 | PE_PGRS47 | BCGDan_sapMKO_2782 | 1275 |
| 3055515 | 3056348 | Rv2742c | Rv2742c | 834 | 3008941 | 3009189 | BCG_2757c | BCG_2757c | 249 | 2988888 | 2989136 | BCGDan_2782 | BCGDan_2782 | 249 | 2988953 | 2989201 | BCGDan_sapMKO_2783 | BCGDan_sapMKO_2783 | 249 |
| NA | NA | NA | NA | NA | 3009210 | 3009797 | BCG_2758c | BCG_2758c | 588 | 2989157 | 2989744 | BCGDan_2783 | BCGDan_2783 | 588 | 2989222 | 2989809 | BCGDan_sapMKO_2784 | BCGDan_sapMKO_2784 | 588 |
| 3056420 | 3057232 | Rv2743c | Rv2743c | 813 | 3009869 | 3010681 | BCG_2759c | BCG_2759c | 813 | 2989816 | 2990628 | BCGDan_2784 | BCGDan_2784 | 813 | 2989881 | 2990693 | BCGDan_sapMKO_2785 | BCGDan_sapMKO_2785 | 813 |
| 3057251 | 3058063 | 35kd_ag | Rv2744c | 813 | 3010700 | 3011512 | 35kd_ag | BCG_2760c | 813 | 2990647 | 2991459 | 35kd_ag | BCGDan_2785 | 813 | 2990712 | 2991524 | 35kd_ag | BCGDan_sapMKO_2786 | 813 |

| | | | | | | | | | | | | | | | | | | | |
|---|---|---|---|---|---|---|---|---|---|---|---|---|---|---|---|---|---|---|---|
| 3058193 | 3058531 | clgR | Rv2745c | 339 | 3011642 | 3011980 | BCG_2761c | BCG_2761c | 339 | 2991589 | 2991927 | BCGDan_2786 | BCGDan_2786 | 339 | 2991654 | 2992010 | BCGDan_sapMKO_2787 | BCGDan_sapMKO_2787 | 357 |
| 3058602 | 3059231 | pgsA3 | Rv2746c | 630 | 3012051 | 3012680 | pgsA3 | BCG_2762c | 630 | 2991998 | 2992627 | pgsA3 | BCGDan_2787 | 630 | 2992063 | 2992662 | pgsA3 | BCGDan_sapMKO_2788 | 600 |
| 3059262 | 3059786 | argA | Rv2747 | 525 | 3012711 | 3013235 | BCG_2763 | BCG_2763 | 525 | 2992658 | 2993182 | BCGDan_2788 | BCGDan_2788 | 525 | 2992723 | 2993247 | BCGDan_sapMKO_2789 | BCGDan_sapMKO_2789 | 525 |
| 3059855 | 3062506 | ftsK | Rv2748c | 2652 | 3013304 | 3015955 | ftsK | BCG_2764c | 2652 | 2993251 | 2995902 | ftsK | BCGDan_2789 | 2652 | 2993316 | 2995619 | ftsK | BCGDan_sapMKO_2790 | 2304 |
| 3062505 | 3062819 | Rv2749 | Rv2749 | 315 | 3015954 | 3016268 | BCG_2765 | BCG_2765 | 315 | 2995901 | 2996215 | BCGDan_2790 | BCGDan_2790 | 315 | 2995966 | 2996280 | BCGDan_sapMKO_2791 | BCGDan_sapMKO_2791 | 315 |
| 3062816 | 3063634 | Rv2750 | Rv2750 | 819 | 3016265 | 3017083 | BCG_2766 | BCG_2766 | 819 | 2996212 | 2997030 | BCGDan_2791 | BCGDan_2791 | 819 | 2996277 | 2997095 | BCGDan_sapMKO_2792 | BCGDan_sapMKO_2792 | 819 |
| 3063638 | 3064528 | Rv2751 | Rv2751 | 891 | 3017087 | 3017977 | BCG_2767 | BCG_2767 | 891 | 2997034 | 2997924 | BCGDan_2792 | BCGDan_2792 | 891 | 2997099 | 2997989 | BCGDan_sapMKO_2793 | BCGDan_sapMKO_2793 | 891 |
| 3064515 | 3066191 | Rv2752c | Rv2752c | 1677 | 3017964 | 3019640 | BCG_2768c | BCG_2768c | 1677 | 2997911 | 2999587 | BCGDan_2793 | BCGDan_2793 | 1677 | 2997976 | 2999652 | BCGDan_sapMKO_2794 | BCGDan_sapMKO_2794 | 1677 |
| 3066222 | 3067124 | dapA | Rv2753c | 903 | 3019671 | 3020573 | dapA | BCG_2769c | 903 | 2999618 | 3000520 | dapA | BCGDan_2794 | 903 | 2999683 | 3000585 | dapA | BCGDan_sapMKO_2795 | 903 |
| 3067193 | 3067945 | thyX | Rv2754c | 753 | 3020642 | 3021394 | BCG_2770c | BCG_2770c | 753 | 3000589 | 3001341 | BCGDan_2795 | BCGDan_2795 | 753 | 3000654 | 3001415 | BCGDan_sapMKO_2796 | BCGDan_sapMKO_2796 | 762 |
| 3068189 | 3068464 | hsdS.1 | Rv2755c | 276 | 3021638 | 3021913 | hsdS' | BCG_2771c | 276 | 3001585 | 3001860 | hsdS | BCGDan_2796 | 276 | 3001650 | 3001925 | hsdS_1 | BCGDan_sapMKO_2797 | 276 |
| 3068461 | 3070083 | hsdM | Rv2756c | 1623 | 3021910 | 3023532 | hsdM | BCG_2772c | 1623 | 3001857 | 3003479 | hsdM | BCGDan_2797 | 1623 | 3001922 | 3003502 | hsdM | BCGDan_sapMKO_2798 | 1581 |
| 3070170 | 3070586 | vapC21 | Rv2757c | 417 | 3023619 | 3024035 | BCG_2773c | BCG_2773c | 417 | 3003566 | 3003982 | BCGDan_2798 | BCGDan_2798 | 417 | 3003631 | 3004047 | BCGDan_sapMKO_2799 | BCGDan_sapMKO_2799 | 417 |
| 3070583 | 3070849 | vapB21 | Rv2758c | 267 | 3024032 | 3024298 | BCG_2774c | BCG_2774c | 267 | 3003979 | 3004245 | BCGDan_2799 | BCGDan_2799 | 267 | 3004044 | 3004271 | BCGDan_sapMKO_2800 | BCGDan_sapMKO_2800 | 228 |
| 3070875 | 3071270 | vapC42 | Rv2759c | 396 | 3024324 | 3024719 | BCG_2775c | BCG_2775c | 396 | 3004271 | 3004666 | BCGDan_2800 | BCGDan_2800 | 396 | 3004336 | 3004731 | BCGDan_sapMKO_2801 | BCGDan_sapMKO_2801 | 396 |
| 3071267 | 3071536 | vapB42 | Rv2760c | 270 | 3024716 | 3024985 | BCG_2776c | BCG_2776c | 270 | 3004663 | 3004932 | BCGDan_2801 | BCGDan_2801 | 270 | 3004728 | 3004997 | BCGDan_sapMKO_2802 | BCGDan_sapMKO_2802 | 270 |
| 3071546 | 3072640 | hsdS | Rv2761c | 1095 | 3024995 | 3026089 | hsdS | BCG_2777c | 1095 | 3004942 | 3006036 | hsdS | BCGDan_2802 | 1095 | 3005007 | 3006101 | hsdS_2 | BCGDan_sapMKO_2803 | 1095 |
| 3072637 | 3073056 | Rv2762c | Rv2762c | 420 | 3026086 | 3026358 | BCG_2778c | BCG_2778c | 273 | 3006033 | 3006305 | BCGDan_2803 | BCGDan_2803 | 273 | 3006098 | 3006370 | BCGDan_sapMKO_2804 | BCGDan_sapMKO_2804 | 273 |
| NA | NA | NA | NA | NA | 3026374 | 3026505 | BCG_2779c | BCG_2779c | 132 | 3006321 | 3006452 | BCGDan_2804 | BCGDan_2804 | 132 | 3006386 | 3006517 | BCGDan_sapMKO_2805 | BCGDan_sapMKO_2805 | 132 |
| 3073130 | 3073609 | dfrA | Rv2763c | 480 | 3026579 | 3027058 | dfrA | BCG_2780c | 480 | 3006526 | 3007005 | dfrA | BCGDan_2805 | 480 | 3006591 | 3007085 | dfrA | BCGDan_sapMKO_2806 | 495 |
| 3073680 | 3074471 | thyA | Rv2764c | 792 | 3027129 | 3027920 | thyA | BCG_2781c | 792 | 3007076 | 3007867 | thyA | BCGDan_2806 | 792 | 3007141 | 3007941 | thyA | BCGDan_sapMKO_2807 | 801 |
| 3074636 | 3075373 | Rv2765 | Rv2765 | 738 | 3028085 | 3028822 | BCG_2782 | BCG_2782 | 738 | 3008032 | 3008769 | BCGDan_2807 | BCGDan_2807 | 738 | 3008097 | 3008834 | BCGDan_sapMKO_2808 | BCGDan_sapMKO_2808 | 738 |
| 3075588 | 3076370 | Rv2766c | Rv2766c | 783 | 3029037 | 3029819 | BCG_2783c | BCG_2783c | 783 | 3008984 | 3009766 | BCGDan_2808 | BCGDan_2808 | 783 | 3009049 | 3009831 | BCGDan_sapMKO_2809 | BCGDan_sapMKO_2809 | 783 |
| 3076367 | 3076720 | Rv2767c | Rv2767c | 354 | 3029816 | 3030169 | BCG_2784c | BCG_2784c | 354 | 3009763 | 3010116 | BCGDan_2809 | BCGDan_2809 | 354 | 3009828 | 3010181 | BCGDan_sapMKO_2810 | BCGDan_sapMKO_2810 | 354 |
| 3076894 | 3078078 | PPE43 | Rv2768c | 1185 | 3030343 | 3031527 | PPE43 | BCG_2785c | 1185 | 3010290 | 3011474 | PPE43 | BCGDan_2810 | 1185 | 3010355 | 3011497 | PPE43 | BCGDan_sapMKO_2811 | 1143 |
| 3078158 | 3078985 | PE27 | Rv2769c | 828 | 3031607 | 3032434 | PE27 | BCG_2786c | 828 | 3011554 | 3012381 | PE27 | BCGDan_2811 | 828 | 3011619 | 3012446 | PE27 | BCGDan_sapMKO_2812 | 828 |
| 3079309 | 3080457 | PPE44 | Rv2770c | 1149 | 3032758 | 3033906 | PPE44 | BCG_2787c | 1149 | 3012705 | 3013853 | PPE44 | BCGDan_2812 | 1149 | 3012770 | 3013876 | PPE44 | BCGDan_sapMKO_2813 | 1107 |
| 3080581 | 3081033 | Rv2771c | Rv2771c | 453 | 3034030 | 3034482 | BCG_2788c | BCG_2788c | 453 | 3013977 | 3014429 | BCGDan_2813 | BCGDan_2813 | 453 | 3014042 | 3014494 | BCGDan_sapMKO_2814 | BCGDan_sapMKO_2814 | 453 |
| 3081119 | 3081592 | Rv2772c | Rv2772c | 474 | 3034568 | 3035041 | BCG_2789c | BCG_2789c | 474 | 3014515 | 3014988 | BCGDan_2814 | BCGDan_2814 | 474 | 3014580 | 3015053 | BCGDan_sapMKO_2815 | BCGDan_sapMKO_2815 | 474 |
| 3081604 | 3082341 | dapB | Rv2773c | 738 | 3035053 | 3035790 | dapB | BCG_2790c | 738 | 3015000 | 3015737 | dapB | BCGDan_2815 | 738 | 3015065 | 3015811 | dapB | BCGDan_sapMKO_2816 | 747 |
| 3082352 | 3082756 | Rv2774c | Rv2774c | 405 | 3035801 | 3036205 | BCG_2791c | BCG_2791c | 405 | 3015748 | 3016152 | BCGDan_2816 | BCGDan_2816 | 405 | 3015813 | 3016217 | BCGDan_sapMKO_2817 | BCGDan_sapMKO_2817 | 405 |
| 3082909 | 3083370 | Rv2775 | Rv2775 | 462 | 3036358 | 3036819 | BCG_2792 | BCG_2792 | 462 | 3016305 | 3016766 | BCGDan_2817 | BCGDan_2817 | 462 | 3016307 | 3016831 | BCGDan_sapMKO_2818 | BCGDan_sapMKO_2818 | 525 |
| 3083374 | 3084303 | Rv2776c | Rv2776c | 930 | 3036823 | 3037752 | BCG_2793c | BCG_2793c | 930 | 3016770 | 3017699 | BCGDan_2818 | BCGDan_2818 | 930 | 3016835 | 3017926 | BCGDan_sapMKO_2819 | BCGDan_sapMKO_2819 | 1092 |
| 3084485 | 3085555 | Rv2777c | Rv2777c | 1071 | 3037934 | 3039004 | BCG_2794c | BCG_2794c | 1071 | 3017881 | 3018951 | BCGDan_2819 | BCGDan_2819 | 1071 | 3017946 | 3018863 | BCGDan_sapMKO_2820 | BCGDan_sapMKO_2820 | 918 |
| 3085713 | 3086183 | Rv2778c | Rv2778c | 471 | 3039162 | 3039632 | BCG_2795c | BCG_2795c | 471 | 3019109 | 3019579 | BCGDan_2820 | BCGDan_2820 | 471 | 3019174 | 3019620 | BCGDan_sapMKO_2821 | BCGDan_sapMKO_2821 | 447 |
| 3086215 | 3086754 | Rv2779c | Rv2779c | 540 | 3039664 | 3040203 | BCG_2796c | BCG_2796c | 540 | 3019611 | 3020150 | BCGDan_2821 | BCGDan_2821 | 540 | 3019676 | 3020215 | BCGDan_sapMKO_2822 | BCGDan_sapMKO_2822 | 540 |
| 3086820 | 3087935 | ald | Rv2780 | 1116 | 3040269 | 3040667 | ald_a | BCG_2797 | 399 | 3020216 | 3020614 | ald_a | BCGDan_2822 | 399 | 3020281 | 3020679 | ald_a | BCGDan_sapMKO_2823 | 399 |
| NA | NA | NA | NA | NA | 3040664 | 3041383 | aldb | BCG_2798 | 720 | 3020611 | 3021330 | aldb | BCGDan_2823 | 720 | 3020550 | 3021395 | aldb | BCGDan_sapMKO_2824 | 846 |
| 3087950 | 3088984 | Rv2781c | Rv2781c | 1035 | 3041398 | 3042432 | BCG_2799c | BCG_2799c | 1035 | 3021345 | 3022379 | BCGDan_2824 | BCGDan_2824 | 1035 | 3021410 | 3022444 | BCGDan_sapMKO_2825 | BCGDan_sapMKO_2825 | 1035 |
| 3089045 | 3090361 | pepR | Rv2782c | 1317 | 3042493 | 3043809 | pepR | BCG_2800c | 1317 | 3022440 | 3023756 | pepR | BCGDan_2825 | 1317 | 3022505 | 3023821 | pepR | BCGDan_sapMKO_2826 | 1317 |
| 3090339 | 3092597 | gpsI | Rv2783c | 2259 | 3043787 | 3046045 | gpsI | BCG_2801c | 2259 | 3023734 | 3025992 | gpsI | BCGDan_2826 | 2259 | 3023799 | 3026057 | gpsI | BCGDan_sapMKO_2827 | 2259 |
| 3092951 | 3093466 | lppU | Rv2784c | 516 | 3046399 | 3046914 | lppU | BCG_2802c | 516 | 3026346 | 3026861 | lppU | BCGDan_2827 | 516 | 3026411 | 3026926 | lppU | BCGDan_sapMKO_2828 | 516 |
| 3093479 | 3093748 | rpsO | Rv2785c | 270 | 3046927 | 3047196 | rpsO | BCG_2803c | 270 | 3026874 | 3027143 | rpsO | BCGDan_2828 | 270 | 3026939 | 3027208 | rpsO | BCGDan_sapMKO_2829 | 270 |
| 3093905 | 3094900 | ribF | Rv2786c | 996 | 3047353 | 3048348 | ribF | BCG_2804c | 996 | 3027300 | 3028295 | ribF | BCGDan_2829 | 996 | 3027365 | 3028360 | ribF | BCGDan_sapMKO_2830 | 996 |
| 3095111 | 3096874 | Rv2787 | Rv2787 | 1764 | 3048559 | 3050322 | BCG_2805 | BCG_2805 | 1764 | 3028506 | 3030269 | BCGDan_2830 | BCGDan_2830 | 1764 | 3028598 | 3030334 | BCGDan_sapMKO_2831 | BCGDan_sapMKO_2831 | 1737 |
| 3096959 | 3097645 | sirR | Rv2788 | 687 | 3050407 | 3051093 | sirR | BCG_2806 | 687 | 3030354 | 3031040 | sirR | BCGDan_2831 | 687 | 3030419 | 3031105 | sirR | BCGDan_sapMKO_2832 | 687 |
| 3097706 | 3098938 | fadE21 | Rv2789c | 1233 | 3051154 | 3052386 | fadE21 | BCG_2807c | 1233 | 3031101 | 3032333 | fadE21 | BCGDan_2832 | 1233 | 3031166 | 3032398 | fadE21 | BCGDan_sapMKO_2833 | 1233 |
| 3098964 | 3100169 | ltp1 | Rv2790c | 1206 | 3052412 | 3053617 | ltp1 | BCG_2808c | 1206 | 3032359 | 3033564 | ltp1 | BCGDan_2833 | 1206 | 3032424 | 3033581 | ltp1 | BCGDan_sapMKO_2834 | 1158 |
| 3100202 | 3101581 | Rv2791c | Rv2791c | 1380 | 3053650 | 3055029 | BCG_2809c | BCG_2809c | 1380 | 3033597 | 3034976 | BCGDan_2834 | BCGDan_2834 | 1380 | 3033662 | 3035041 | BCGDan_sapMKO_2835 | BCGDan_sapMKO_2835 | 1380 |
| 3101581 | 3102162 | Rv2792c | Rv2792c | 582 | 3055029 | 3055610 | BCG_2810c | BCG_2810c | 582 | 3034976 | 3035557 | BCGDan_2835 | BCGDan_2835 | 582 | 3035041 | 3035622 | BCGDan_sapMKO_2836 | BCGDan_sapMKO_2836 | 582 |
| 3102364 | 3103260 | truB | Rv2793c | 897 | 3055812 | 3056708 | truB | BCG_2811c | 897 | 3035759 | 3036655 | truB | BCGDan_2836 | 897 | 3035824 | 3036720 | truB | BCGDan_sapMKO_2837 | 897 |
| 3103257 | 3103940 | pptT | Rv2794c | 684 | 3056705 | 3057388 | BCG_2812c | BCG_2812c | 684 | 3036652 | 3037335 | BCGDan_2837 | BCGDan_2837 | 684 | 3036717 | 3037400 | BCGDan_sapMKO_2838 | BCGDan_sapMKO_2838 | 684 |
| 3103937 | 3104911 | Rv2795c | Rv2795c | 975 | 3057385 | 3058359 | BCG_2813c | BCG_2813c | 975 | 3037332 | 3038306 | BCGDan_2838 | BCGDan_2838 | 975 | 3037397 | 3038371 | BCGDan_sapMKO_2839 | BCGDan_sapMKO_2839 | 975 |
| 3105056 | 3105619 | lppV | Rv2796c | 564 | 3058504 | 3059067 | lppV | BCG_2814c | 564 | 3038451 | 3039014 | lppV | BCGDan_2839 | 564 | 3038516 | 3039079 | lppV | BCGDan_sapMKO_2840 | 564 |
| 3105619 | 3107307 | Rv2797c | Rv2797c | 1689 | 3059067 | 3060755 | BCG_2815c | BCG_2815c | 1689 | 3039014 | 3040702 | BCGDan_2840 | BCGDan_2840 | 1689 | 3039079 | 3040767 | BCGDan_sapMKO_2841 | BCGDan_sapMKO_2841 | 1689 |
| 3107311 | 3107637 | Rv2798c | Rv2798c | 327 | 3060759 | 3061085 | BCG_2816c | BCG_2816c | 327 | 3040706 | 3041032 | BCGDan_2841 | BCGDan_2841 | 327 | 3040771 | 3041118 | BCGDan_sapMKO_2842 | BCGDan_sapMKO_2842 | 348 |
| 3107768 | 3108397 | Rv2799 | Rv2799 | 630 | 3061216 | 3061845 | BCG_2817 | BCG_2817 | 630 | 3041163 | 3041792 | BCGDan_2842 | BCGDan_2842 | 630 | 3041270 | 3041857 | BCGDan_sapMKO_2843 | BCGDan_sapMKO_2843 | 588 |
| 3108416 | 3110065 | Rv2800 | Rv2800 | 1650 | 3061864 | 3063513 | BCG_2818 | BCG_2818 | 1650 | 3041811 | 3043460 | BCGDan_2843 | BCGDan_2843 | 1650 | 3041876 | 3043525 | BCGDan_sapMKO_2844 | BCGDan_sapMKO_2844 | 1650 |
| 3110167 | 3110523 | mazF9 | Rv2801c | 357 | 3063615 | 3063971 | BCG_2819c | BCG_2819c | 357 | 3043562 | 3043918 | mazF9 | BCGDan_2844 | 357 | 3043627 | 3043980 | BCGDan_sapMKO_2845 | BCGDan_sapMKO_2845 | 354 |
| 3110507 | 3110737 | mazE9 | Rv2801A | 231 | NA | NA | NA | NA | NA | 3043902 | 3044132 | mazE9 | BCGDan_2845 | 231 | 3043967 | 3044197 | mazE9 | BCGDan_sapMKO_2846 | 231 |
| 3110780 | 3111823 | Rv2802c | Rv2802c | 1044 | 3064228 | 3065271 | BCG_2820c | BCG_2820c | 1044 | 3044175 | 3045218 | BCGDan_2846 | BCGDan_2846 | 1044 | 3044240 | 3045283 | BCGDan_sapMKO_2847 | BCGDan_sapMKO_2847 | 1044 |
| 3111822 | 3112289 | Rv2803 | Rv2803 | 468 | 3065270 | 3065737 | BCG_2821 | BCG_2821 | 468 | 3045217 | 3045684 | BCGDan_2847 | BCGDan_2847 | 468 | 3045282 | 3045749 | BCGDan_sapMKO_2848 | BCGDan_sapMKO_2848 | 468 |
| 3112465 | 3113094 | Rv2804c | Rv2804c | 630 | 3065913 | 3066542 | BCG_2822c | BCG_2822c | 630 | 3045860 | 3046489 | BCGDan_2848 | BCGDan_2848 | 630 | 3045925 | 3046554 | BCGDan_sapMKO_2849 | BCGDan_sapMKO_2849 | 630 |
| 3112867 | 3113271 | Rv2805 | Rv2805 | 405 | 3066315 | 3066719 | BCG_2823 | BCG_2823 | 405 | 3046262 | 3046666 | BCGDan_2849 | BCGDan_2849 | 405 | 3046300 | 3046731 | BCGDan_sapMKO_2850 | BCGDan_sapMKO_2850 | 432 |
| 3113268 | 3113459 | Rv2806 | Rv2806 | 192 | 3066716 | 3066907 | BCG_2824 | BCG_2824 | 192 | 3046663 | 3046854 | BCGDan_2850 | BCGDan_2850 | 192 | 3046728 | 3046919 | BCGDan_sapMKO_2851 | BCGDan_sapMKO_2851 | 192 |
| 3113658 | 3114812 | Rv2807 | Rv2807 | 1155 | 3067106 | 3068260 | BCG_2825 | BCG_2825 | 1155 | 3047053 | 3048207 | BCGDan_2851 | BCGDan_2851 | 1155 | 3047118 | 3048272 | BCGDan_sapMKO_2852 | BCGDan_sapMKO_2852 | 1155 |
| 3115046 | 3115303 | Rv2808 | Rv2808 | 258 | 3068494 | 3068751 | BCG_2826 | BCG_2826 | 258 | 3048441 | 3048698 | BCGDan_2852 | BCGDan_2852 | 258 | 3048506 | 3048763 | BCGDan_sapMKO_2853 | BCGDan_sapMKO_2853 | 258 |
| 3115408 | 3115719 | Rv2809 | Rv2809 | 312 | 3068856 | 3069167 | BCG_2827 | BCG_2827 | 312 | 3048803 | 3049114 | BCGDan_2853 | BCGDan_2853 | 312 | 3048868 | 3049179 | BCGDan_sapMKO_2854 | BCGDan_sapMKO_2854 | 312 |
| 3115741 | 3116142 | Rv2810c | Rv2810c | 402 | 3069189 | 3069590 | BCG_2828c | BCG_2828c | 402 | 3049136 | 3049537 | BCGDan_2854 | BCGDan_2854 | 402 | 3049204 | 3049602 | BCGDan_sapMKO_2855 | BCGDan_sapMKO_2855 | 399 |
| 3116139 | 3116747 | Rv2811 | Rv2811 | 609 | 3069587 | 3070195 | BCG_2829 | BCG_2829 | 609 | 3049534 | 3050142 | BCGDan_2855 | BCGDan_2855 | 609 | 3049599 | 3050207 | BCGDan_sapMKO_2856 | BCGDan_sapMKO_2856 | 609 |
| 3116818 | 3118227 | Rv2812 | Rv2812 | 1410 | 3070266 | 3071675 | BCG_2830 | BCG_2830 | 1410 | 3050213 | 3051622 | BCGDan_2856 | BCGDan_2856 | 1410 | 3050278 | 3051687 | BCGDan_sapMKO_2857 | BCGDan_sapMKO_2857 | 1410 |
| 3118224 | 3119036 | Rv2813 | Rv2813 | 813 | 3071672 | 3072484 | BCG_2831 | BCG_2831 | 813 | 3051619 | 3052431 | BCGDan_2857 | BCGDan_2857 | 813 | 3051687 | 3052496 | BCGDan_sapMKO_2858 | BCGDan_sapMKO_2858 | 810 |
| 3120566 | 3121552 | Rv2814c | Rv2814c | 987 | 3074087 | 3074923 | BCG_2832c | BCG_2832c | 837 | 3054034 | 3054870 | BCGDan_2858 | BCGDan_2858 | 837 | 3054099 | 3054983 | BCGDan_sapMKO_2859 | BCGDan_sapMKO_2859 | 885 |
| 3121501 | 3121827 | Rv2815c | Rv2815c | 327 | 3075022 | 3075348 | BCG_2833c | BCG_2833c | 327 | 3054969 | 3055295 | BCGDan_2859 | BCGDan_2859 | 327 | 3055034 | 3055360 | BCGDan_sapMKO_2860 | BCGDan_sapMKO_2860 | 327 |
| 3123625 | 3123966 | Rv2816c | Rv2816c | 342 | 3077602 | 3077943 | BCG_2834c | BCG_2834c | 342 | 3057549 | 3057890 | BCGDan_2860 | BCGDan_2860 | 342 | 3057614 | 3057955 | BCGDan_sapMKO_2861 | BCGDan_sapMKO_2861 | 342 |
| 3123967 | 3124983 | Rv2817c | Rv2817c | 1017 | 3077944 | 3078960 | BCG_2835c | BCG_2835c | 1017 | 3057891 | 3058907 | BCGDan_2861 | BCGDan_2861 | 1017 | 3057956 | 3058981 | BCGDan_sapMKO_2862 | BCGDan_sapMKO_2862 | 1026 |
| 3124996 | 3126144 | Rv2818c | Rv2818c | 1149 | 3078973 | 3079185 | BCG_2836c | BCG_2836c | 213 | 3058920 | 3059132 | BCGDan_2862 | BCGDan_2862 | 213 | 3058988 | 3059197 | BCGDan_sapMKO_2863 | BCGDan_sapMKO_2863 | 210 |
| NA | NA | NA | NA | NA | 3079210 | 3080220 | BCG_2837c | BCG_2837c | 1011 | 3059157 | 3060167 | BCGDan_2863 | BCGDan_2863 | 1011 | 3059222 | 3060232 | BCGDan_sapMKO_2864 | BCGDan_sapMKO_2864 | 1011 |
| 3126240 | 3127367 | Rv2819c | Rv2819c | 1128 | 3080217 | 3081344 | BCG_2838c | BCG_2838c | 1128 | 3060164 | 3061291 | BCGDan_2864 | BCGDan_2864 | 1128 | 3060229 | 3061356 | BCGDan_sapMKO_2865 | BCGDan_sapMKO_2865 | 1128 |
| 3127364 | 3128272 | Rv2820c | Rv2820c | 909 | 3081341 | 3082249 | BCG_2839c | BCG_2839c | 909 | 3061288 | 3062196 | BCGDan_2865 | BCGDan_2865 | 909 | 3061353 | 3062261 | BCGDan_sapMKO_2866 | BCGDan_sapMKO_2866 | 909 |
| 3128253 | 3128963 | Rv2821c | Rv2821c | 711 | 3082230 | 3082940 | BCG_2840c | BCG_2840c | 711 | 3062177 | 3062887 | BCGDan_2866 | BCGDan_2866 | 711 | 3062242 | 3062952 | BCGDan_sapMKO_2867 | BCGDan_sapMKO_2867 | 711 |
| 3128973 | 3129347 | Rv2822c | Rv2822c | 375 | 3082950 | 3083324 | BCG_2841c | BCG_2841c | 375 | 3062897 | 3063271 | BCGDan_2867 | BCGDan_2867 | 375 | 3062962 | 3063336 | BCGDan_sapMKO_2868 | BCGDan_sapMKO_2868 | 375 |
| 3129344 | 3131773 | Rv2823c | Rv2823c | 2430 | 3083321 | 3085759 | BCG_2842c | BCG_2842c | 2439 | 3063268 | 3065706 | BCGDan_2868 | BCGDan_2868 | 2439 | 3063333 | 3065771 | BCGDan_sapMKO_2869 | BCGDan_sapMKO_2869 | 2439 |
| 3131770 | 3132714 | Rv2824c | Rv2824c | 945 | 3085756 | 3086700 | BCG_2843c | BCG_2843c | 945 | 3065703 | 3066647 | BCGDan_2869 | BCGDan_2869 | 945 | 3065768 | 3066712 | BCGDan_sapMKO_2870 | BCGDan_sapMKO_2870 | 945 |
| 3132892 | 3133539 | Rv2825c | Rv2825c | 648 | 3086878 | 3087525 | BCG_2844c | BCG_2844c | 648 | 3066825 | 3067472 | BCGDan_2870 | BCGDan_2870 | 648 | 3066890 | 3067537 | BCGDan_sapMKO_2871 | BCGDan_sapMKO_2871 | 648 |
| 3133709 | 3134593 | Rv2826c | Rv2826c | 885 | 3087695 | 3088579 | BCG_2845c | BCG_2845c | 885 | 3067642 | 3068526 | BCGDan_2871 | BCGDan_2871 | 885 | 3067707 | 3068591 | BCGDan_sapMKO_2872 | BCGDan_sapMKO_2872 | 885 |

| | | | | | | | | | | | | | | | | | | | |
|---|---|---|---|---|---|---|---|---|---|---|---|---|---|---|---|---|---|---|---|
| 3134596 | 3135483 | Rv2827c | Rv2827c | 888 | 3088582 | 3089469 | BCG_2846c | BCG_2846c | 888 | 3068529 | 3069416 | BCGDan_2872 | BCGDan_2872 | 888 | 3068594 | 3069496 | BCGDan_sapMKO_2873 | BCGDan_sapMKO_2873 | 903 |
| 3135788 | 3136333 | Rv2828c | Rv2828c | 546 | 3089774 | 3090319 | BCG_2847c | BCG_2847c | 546 | 3069721 | 3070266 | BCGDan_2873 | BCGDan_2873 | 546 | 3069786 | 3070331 | BCGDan_sapMKO_2874 | BCGDan_sapMKO_2874 | 546 |
| 3136330 | 3136599 | Rv2828A | Rv2828A | 270 | 3090316 | 3090585 | BCG_2848c | BCG_2848c | 270 | 3070263 | 3070532 | BCGDan_2874 | BCGDan_2874 | 270 | 3070328 | 3070597 | BCGDan_sapMKO_2875 | BCGDan_sapMKO_2875 | 270 |
| 3136620 | 3137012 | vapC22 | Rv2829c | 393 | 3090606 | 3090998 | BCG_2849c | BCG_2849c | 393 | 3070553 | 3070945 | BCGDan_2875 | BCGDan_2875 | 393 | 3070618 | 3071010 | BCGDan_sapMKO_2876 | BCGDan_sapMKO_2876 | 393 |
| 3137009 | 3137224 | vapB22 | Rv2830c | 216 | 3090995 | 3091210 | BCG_2850c | BCG_2850c | 216 | 3070942 | 3071157 | BCGDan_2876 | BCGDan_2876 | 216 | 3071007 | 3071234 | BCGDan_sapMKO_2877 | BCGDan_sapMKO_2877 | 228 |
| 3137271 | 3138020 | echA16 | Rv2831 | 750 | 3091257 | 3092006 | echA16 | BCG_2851 | 750 | 3071204 | 3071953 | echA16 | BCGDan_2877 | 750 | 3071269 | 3072018 | echA16 | BCGDan_sapMKO_2878 | 750 |
| 3138099 | 3139181 | ugpC | Rv2832c | 1083 | 3092085 | 3093167 | ugpC | BCG_2852c | 1083 | 3072032 | 3073114 | ugpC | BCGDan_2878 | 1083 | 3072097 | 3073179 | ugpC | BCGDan_sapMKO_2879 | 1083 |
| NA | NA | NA | NA | NA | 3093160 | 3093837 | ugpBb | BCG_2853A | 678 | 3073107 | 3073784 | ugpBb | BCGDan_2879 | 678 | 3073175 | 3073849 | BCGDan_sapMKO_2880 | BCGDan_sapMKO_2880 | 675 |
| 3139174 | 3140484 | ugpB | Rv2833c | 1311 | 3093838 | 3094470 | ugpBa | BCG_2853c | 633 | 3073785 | 3074417 | ugpBa | BCGDan_2880 | 633 | 3073850 | 3074482 | ugpBa | BCGDan_sapMKO_2881 | 633 |
| 3140487 | 3141314 | ugpE | Rv2834c | 828 | 3094473 | 3095300 | ugpE | BCG_2854c | 828 | 3074420 | 3075247 | ugpE | BCGDan_2881 | 828 | 3074485 | 3075270 | ugpE | BCGDan_sapMKO_2882 | 786 |
| 3141311 | 3142222 | ugpA | Rv2835c | 912 | 3095297 | 3096208 | ugpA | BCG_2855c | 912 | 3075244 | 3076155 | ugpA | BCGDan_2882 | 912 | 3075309 | 3076352 | ugpA | BCGDan_sapMKO_2883 | 1044 |
| 3142309 | 3143628 | dinF | Rv2836c | 1320 | 3096295 | 3097614 | dinF | BCG_2856c | 1320 | 3076242 | 3077561 | dinF | BCGDan_2883 | 1320 | 3076307 | 3077617 | dinF | BCGDan_sapMKO_2884 | 1311 |
| 3143635 | 3144645 | Rv2837c | Rv2837c | 1011 | 3097621 | 3098631 | BCG_2857c | BCG_2857c | 1011 | 3077568 | 3078578 | BCGDan_2884 | BCGDan_2884 | 1011 | 3077633 | 3078583 | BCGDan_sapMKO_2885 | BCGDan_sapMKO_2885 | 951 |
| 3144620 | 3145171 | rbfA | Rv2838c | 552 | 3098606 | 3099157 | rbfA | BCG_2858c | 552 | 3078553 | 3079104 | rbfA | BCGDan_2885 | 552 | 3078618 | 3079169 | rbfA | BCGDan_sapMKO_2886 | 552 |
| 3145171 | 3147873 | infB | Rv2839c | 2703 | 3099157 | 3101859 | infB | BCG_2859c | 2703 | 3079104 | 3081806 | infB | BCGDan_2886 | 2703 | 3079169 | 3081871 | infB | BCGDan_sapMKO_2887 | 2703 |
| 3147959 | 3148258 | Rv2840c | Rv2840c | 300 | 3101945 | 3102244 | BCG_2860c | BCG_2860c | 300 | 3081892 | 3082191 | BCGDan_2887 | BCGDan_2887 | 300 | 3081957 | 3082223 | BCGDan_sapMKO_2888 | BCGDan_sapMKO_2888 | 267 |
| 3148385 | 3149428 | nusA | Rv2841c | 1044 | 3102371 | 3103414 | nusA | BCG_2861c | 1044 | 3082318 | 3083361 | nusA | BCGDan_2888 | 1044 | 3082383 | 3083426 | nusA | BCGDan_sapMKO_2889 | 1044 |
| 3149425 | 3149976 | Rv2842c | Rv2842c | 552 | 3103411 | 3103962 | BCG_2862c | BCG_2862c | 552 | 3083358 | 3083909 | BCGDan_2889 | BCGDan_2889 | 552 | 3083423 | 3083974 | BCGDan_sapMKO_2890 | BCGDan_sapMKO_2890 | 552 |
| 3150171 | 3150716 | Rv2843 | Rv2843 | 546 | 3104157 | 3104702 | BCG_2863 | BCG_2863 | 546 | 3084104 | 3084649 | BCGDan_2890 | BCGDan_2890 | 546 | 3084229 | 3084714 | BCGDan_sapMKO_2891 | BCGDan_sapMKO_2891 | 486 |
| 3150713 | 3151201 | Rv2844 | Rv2844 | 489 | 3104699 | 3105187 | BCG_2864 | BCG_2864 | 489 | 3084646 | 3085134 | BCGDan_2891 | BCGDan_2891 | 489 | 3084711 | 3085199 | BCGDan_sapMKO_2892 | BCGDan_sapMKO_2892 | 489 |
| 3151202 | 3152950 | proS | Rv2845c | 1749 | 3105188 | 3106936 | proS | BCG_2865c | 1749 | 3085135 | 3086883 | proS | BCGDan_2892 | 1749 | 3085200 | 3086948 | proS | BCGDan_sapMKO_2893 | 1749 |
| 3153039 | 3154631 | efpA | Rv2846c | 1593 | 3107025 | 3108617 | efpA | BCG_2866c | 1593 | 3086972 | 3088564 | efpA | BCGDan_2893 | 1593 | 3087037 | 3088449 | efpA | BCGDan_sapMKO_2894 | 1413 |
| 3154654 | 3155871 | cysG | Rv2847c | 1218 | 3108640 | 3109857 | cysG | BCG_2867c | 1218 | 3088587 | 3089804 | cysG | BCGDan_2894 | 1218 | 3088652 | 3089869 | cysG | BCGDan_sapMKO_2895 | 1218 |
| 3156148 | 3157521 | cobB | Rv2848c | 1374 | 3110134 | 3111507 | cobB | BCG_2868c | 1374 | 3090027 | 3091400 | cobB | BCGDan_2895 | 1374 | 3090092 | 3091471 | cobB | BCGDan_sapMKO_2896 | 1380 |
| 3157521 | 3158144 | cobO | Rv2849c | 624 | 3111507 | 3112130 | cobO | BCG_2869c | 624 | 3091400 | 3092023 | cobO | BCGDan_2896 | 624 | 3091465 | 3092088 | cobO | BCGDan_sapMKO_2897 | 624 |
| 3158165 | 3160054 | Rv2850c | Rv2850c | 1890 | 3112151 | 3114040 | BCG_2870c | BCG_2870c | 1890 | 3092044 | 3093933 | BCGDan_2897 | BCGDan_2897 | 1890 | 3092109 | 3093998 | BCGDan_sapMKO_2898 | BCGDan_sapMKO_2898 | 1890 |
| 3160051 | 3160521 | Rv2851c | Rv2851c | 471 | 3114037 | 3114507 | BCG_2871c | BCG_2871c | 471 | 3093930 | 3094400 | BCGDan_2898 | BCGDan_2898 | 471 | 3093995 | 3094465 | BCGDan_sapMKO_2899 | BCGDan_sapMKO_2899 | 471 |
| 3160580 | 3162061 | mqo | Rv2852c | 1482 | 3114566 | 3116047 | mqo | BCG_2872c | 1482 | 3094459 | 3095940 | mqo | BCGDan_2899 | 1482 | 3094524 | 3095978 | mqo | BCGDan_sapMKO_2900 | 1455 |
| 3162268 | 3164115 | PE_PGRS48 | Rv2853 | 1848 | 3116254 | 3118047 | PE_PGRS48 | BCG_2873 | 1794 | 3096147 | 3097940 | PE_PGRS48 | BCGDan_2900 | 1794 | 3096212 | 3098005 | PE_PGRS48 | BCGDan_sapMKO_2901 | 1794 |
| 3164152 | 3165192 | Rv2854 | Rv2854 | 1041 | 3118084 | 3119124 | BCG_2874 | BCG_2874 | 1041 | 3097977 | 3099017 | BCGDan_2901 | BCGDan_2901 | 1041 | 3098042 | 3099082 | BCGDan_sapMKO_2902 | BCGDan_sapMKO_2902 | 1041 |
| 3165205 | 3166584 | mtr | Rv2855 | 1380 | 3119137 | 3120516 | mtr | BCG_2875 | 1380 | 3099030 | 3100409 | mtr | BCGDan_2902 | 1380 | 3099095 | 3100474 | mtr | BCGDan_sapMKO_2903 | 1380 |
| 3166684 | 3167802 | nicT | Rv2856 | 1119 | 3120616 | 3121734 | nicT | BCG_2876 | 1119 | 3100509 | 3101627 | nicT | BCGDan_2903 | 1119 | 3100574 | 3101692 | nicT | BCGDan_sapMKO_2904 | 1119 |
| NA | NA | NA | NA | NA | 3121800 | 3122132 | BCG_2877 | BCG_2877 | 333 | 3101693 | 3102025 | BCGDan_2904 | BCGDan_2904 | 333 | 3101758 | 3102090 | BCGDan_sapMKO_2905 | BCGDan_sapMKO_2905 | 333 |
| NA | NA | NA | NA | NA | 3122144 | 3122359 | BCG_2878 | BCG_2878 | 216 | 3102037 | 3102252 | BCGDan_2905 | BCGDan_2905 | 216 | 3102102 | 3102317 | BCGDan_sapMKO_2906 | BCGDan_sapMKO_2906 | 216 |
| 3168583 | 3169359 | Rv2857c | Rv2857c | 777 | 3122515 | 3123291 | BCG_2879c | BCG_2879c | 777 | 3102408 | 3103184 | BCGDan_2906 | BCGDan_2906 | 777 | 3102473 | 3103246 | BCGDan_sapMKO_2907 | BCGDan_sapMKO_2907 | 774 |
| 3169356 | 3170723 | aldC | Rv2858c | 1368 | 3123288 | 3124655 | aldC | BCG_2880c | 1368 | 3103181 | 3104548 | aldC | BCGDan_2907 | 1368 | 3103246 | 3104613 | aldC | BCGDan_sapMKO_2908 | 1368 |
| 3170720 | 3171646 | Rv2859c | Rv2859c | 927 | 3124652 | 3125578 | BCG_2881c | BCG_2881c | 927 | 3104545 | 3105471 | BCGDan_2908 | BCGDan_2908 | 927 | 3104610 | 3105464 | BCGDan_sapMKO_2909 | BCGDan_sapMKO_2909 | 855 |
| 3171627 | 3173000 | glnA4 | Rv2860c | 1374 | 3125559 | 3126932 | glnA4 | BCG_2882c | 1374 | 3105452 | 3106825 | glnA4 | BCGDan_2909 | 1374 | 3105517 | 3106890 | glnA4 | BCGDan_sapMKO_2910 | 1374 |
| 3173160 | 3174017 | mapB | Rv2861c | 858 | 3127092 | 3127949 | mapB | BCG_2883c | 858 | 3106985 | 3107842 | mapB | BCGDan_2910 | 858 | 3107050 | 3107856 | mapB | BCGDan_sapMKO_2911 | 807 |
| 3174059 | 3174643 | Rv2862c | Rv2862c | 585 | 3127991 | 3128575 | BCG_2884c | BCG_2884c | 585 | 3107884 | 3108468 | BCGDan_2911 | BCGDan_2911 | 585 | 3107949 | 3108533 | BCGDan_sapMKO_2912 | BCGDan_sapMKO_2912 | 585 |
| 3174992 | 3175372 | vapC23 | Rv2863 | 381 | 3128924 | 3129304 | BCG_2885 | BCG_2885 | 381 | 3108817 | 3109197 | BCGDan_2912 | BCGDan_2912 | 381 | 3108882 | 3109262 | BCGDan_sapMKO_2913 | BCGDan_sapMKO_2913 | 381 |
| 3175454 | 3177265 | Rv2864c | Rv2864c | 1812 | 3129386 | 3131197 | BCG_2886c | BCG_2886c | 1812 | 3109279 | 3111090 | BCGDan_2913 | BCGDan_2913 | 1812 | 3109344 | 3111116 | BCGDan_sapMKO_2914 | BCGDan_sapMKO_2914 | 1773 |
| 3177537 | 3177818 | relF | Rv2865 | 282 | 3131469 | 3131750 | BCG_2887 | BCG_2887 | 282 | 3111362 | 3111643 | BCGDan_2914 | BCGDan_2914 | 282 | 3111427 | 3111708 | BCGDan_sapMKO_2915 | BCGDan_sapMKO_2915 | 282 |
| 3177822 | 3178085 | relG | Rv2866 | 264 | 3131754 | 3132017 | BCG_2888 | BCG_2888 | 264 | 3111647 | 3111910 | BCGDan_2915 | BCGDan_2915 | 264 | 3111712 | 3111975 | BCGDan_sapMKO_2916 | BCGDan_sapMKO_2916 | 264 |
| 3178458 | 3179312 | Rv2867c | Rv2867c | 855 | 3132390 | 3133244 | BCG_2889c | BCG_2889c | 855 | 3112283 | 3113137 | BCGDan_2916 | BCGDan_2916 | 855 | 3112348 | 3113202 | BCGDan_sapMKO_2917 | BCGDan_sapMKO_2917 | 855 |
| 3179368 | 3180531 | gcpE | Rv2868c | 1164 | 3133300 | 3134463 | gcpE | BCG_2890c | 1164 | 3113193 | 3114356 | gcpE | BCGDan_2917 | 1164 | 3113258 | 3114349 | gcpE | BCGDan_sapMKO_2918 | 1092 |
| 3180548 | 3181762 | rip | Rv2869c | 1215 | 3134480 | 3135694 | BCG_2891c | BCG_2891c | 1215 | 3114373 | 3115587 | BCGDan_2918 | BCGDan_2918 | 1215 | 3114438 | 3115652 | BCGDan_sapMKO_2919 | BCGDan_sapMKO_2919 | 1215 |
| 3181770 | 3183011 | dxr | Rv2870c | 1242 | 3135702 | 3136943 | dxr | BCG_2892c | 1242 | 3115595 | 3116836 | dxr | BCGDan_2919 | 1242 | 3115660 | 3116970 | dxr | BCGDan_sapMKO_2920 | 1311 |
| 3183138 | 3183395 | vapB43 | Rv2871 | 258 | 3137070 | 3137327 | BCG_2893 | BCG_2893 | 258 | 3116963 | 3117220 | BCGDan_2920 | BCGDan_2920 | 258 | 3117028 | 3117285 | BCGDan_sapMKO_2921 | BCGDan_sapMKO_2921 | 258 |
| 3183382 | 3183825 | vapC43 | Rv2872 | 444 | 3137314 | 3137757 | BCG_2894 | BCG_2894 | 444 | 3117207 | 3117650 | BCGDan_2921 | BCGDan_2921 | 444 | 3117272 | 3117715 | BCGDan_sapMKO_2922 | BCGDan_sapMKO_2922 | 444 |
| 3183905 | 3184567 | mpt83 | Rv2873 | 663 | 3137837 | 3138499 | mpb83 | BCG_2895 | 663 | 3117730 | 3118392 | mpb83 | BCGDan_2922 | 663 | 3117795 | 3118457 | mpb83 | BCGDan_sapMKO_2923 | 663 |
| 3184847 | 3186934 | dipZ | Rv2874 | 2088 | 3138779 | 3140866 | dipZ | BCG_2896 | 2088 | 3118672 | 3120759 | dipZ | BCGDan_2923 | 2088 | 3118737 | 3120824 | dipZ | BCGDan_sapMKO_2924 | 2088 |
| 3187030 | 3187611 | mpt70 | Rv2875 | 582 | 3140962 | 3141543 | mpb70 | BCG_2897 | 582 | 3120855 | 3121436 | mpb70 | BCGDan_2924 | 582 | 3120911 | 3121501 | mpb70 | BCGDan_sapMKO_2925 | 591 |
| 3187663 | 3187977 | Rv2876 | Rv2876 | 315 | 3141595 | 3141909 | BCG_2898 | BCG_2898 | 315 | 3121488 | 3121802 | BCGDan_2925 | BCGDan_2925 | 315 | 3121553 | 3121867 | BCGDan_sapMKO_2926 | BCGDan_sapMKO_2926 | 315 |
| 3188008 | 3188871 | Rv2877c | Rv2877c | 864 | 3141940 | 3142803 | BCG_2899c | BCG_2899c | 864 | 3121833 | 3122696 | BCGDan_2926 | BCGDan_2926 | 864 | 3121898 | 3122761 | BCGDan_sapMKO_2927 | BCGDan_sapMKO_2927 | 864 |
| 3188876 | 3189397 | mpt53 | Rv2878c | 522 | 3142808 | 3143329 | mpb53 | BCG_2900c | 522 | 3122701 | 3123222 | mpb53 | BCGDan_2927 | 522 | 3122766 | 3123287 | mpb53 | BCGDan_sapMKO_2928 | 522 |
| 3189583 | 3190152 | Rv2879c | Rv2879c | 570 | 3143515 | 3144609 | BCG_2901c | BCG_2901c | 1095 | 3123408 | 3124502 | BCGDan_2928 | BCGDan_2928 | 1095 | 3123473 | 3124567 | BCGDan_sapMKO_2929 | BCGDan_sapMKO_2929 | 1095 |
| 3190701 | 3191621 | cdsA | Rv2881c | 921 | 3144632 | 3145552 | cdsA | BCG_2902c | 921 | 3124525 | 3125445 | cdsA | BCGDan_2929 | 921 | 3124590 | 3125510 | cdsA | BCGDan_sapMKO_2930 | 921 |
| 3191644 | 3192201 | frr | Rv2882c | 558 | 3145575 | 3146132 | frr | BCG_2903c | 558 | 3125468 | 3126025 | frr | BCGDan_2930 | 558 | 3125533 | 3126090 | frr | BCGDan_sapMKO_2931 | 558 |
| 3192373 | 3193158 | pyrH | Rv2883c | 786 | 3146304 | 3147089 | pyrH | BCG_2904c | 786 | 3126197 | 3126982 | pyrH | BCGDan_2931 | 786 | 3126262 | 3127047 | pyrH | BCGDan_sapMKO_2932 | 786 |
| 3193393 | 3194151 | Rv2884 | Rv2884 | 759 | 3147324 | 3148082 | BCG_2905 | BCG_2905 | 759 | 3127217 | 3127975 | BCGDan_2932 | BCGDan_2932 | 759 | 3127339 | 3128040 | BCGDan_sapMKO_2933 | BCGDan_sapMKO_2933 | 702 |
| 3194166 | 3195548 | Rv2885c | Rv2885c | 1383 | 3148161 | 3149480 | BCG_2906c | BCG_2906c | 1320 | 3128054 | 3129373 | BCGDan_2933 | BCGDan_2933 | 1320 | 3128119 | 3129402 | BCGDan_sapMKO_2934 | BCGDan_sapMKO_2934 | 1284 |
| 3195545 | 3196432 | Rv2886c | Rv2886c | 888 | 3149477 | 3150364 | BCG_2907c | BCG_2907c | 888 | 3129370 | 3130257 | BCGDan_2934 | BCGDan_2934 | 888 | 3129435 | 3130322 | BCGDan_sapMKO_2935 | BCGDan_sapMKO_2935 | 888 |
| 3196431 | 3196850 | Rv2887 | Rv2887 | 420 | 3150363 | 3150782 | BCG_2908 | BCG_2908 | 420 | 3130256 | 3130675 | BCGDan_2935 | BCGDan_2935 | 420 | 3130321 | 3130740 | BCGDan_sapMKO_2936 | BCGDan_sapMKO_2936 | 420 |
| 3196864 | 3198285 | amiC | Rv2888c | 1422 | 3150796 | 3152217 | amiC | BCG_2909c | 1422 | 3130689 | 3132110 | amiC | BCGDan_2936 | 1422 | 3130754 | 3132175 | amiC | BCGDan_sapMKO_2937 | 1422 |
| 3198292 | 3199107 | tsf | Rv2889c | 816 | 3152224 | 3153039 | tsf | BCG_2910c | 816 | 3132117 | 3132932 | tsf | BCGDan_2937 | 816 | 3132182 | 3132997 | tsf | BCGDan_sapMKO_2938 | 816 |
| 3199119 | 3199982 | rpsB | Rv2890c | 864 | 3153051 | 3153914 | rpsB | BCG_2911c | 864 | 3132944 | 3133807 | rpsB | BCGDan_2938 | 864 | 3133009 | 3133872 | rpsB | BCGDan_sapMKO_2939 | 864 |
| 3200266 | 3201015 | Rv2891 | Rv2891 | 750 | 3154198 | 3154947 | BCG_2912 | BCG_2912 | 750 | 3134091 | 3134840 | BCGDan_2939 | BCGDan_2939 | 750 | 3134156 | 3134905 | BCGDan_sapMKO_2940 | BCGDan_sapMKO_2940 | 750 |
| 3200794 | 3202020 | PPE45 | Rv2892c | 1227 | 3154726 | 3155952 | PPE45 | BCG_2913c | 1227 | 3134619 | 3135845 | PPE45 | BCGDan_2940 | 1227 | 3134684 | 3135841 | PPE45 | BCGDan_sapMKO_2941 | 1158 |
| 3202420 | 3203397 | Rv2893 | Rv2893 | 978 | 3156352 | 3157329 | BCG_2914 | BCG_2914 | 978 | 3136245 | 3137222 | BCGDan_2941 | BCGDan_2941 | 978 | 3136310 | 3137287 | BCGDan_sapMKO_2942 | BCGDan_sapMKO_2942 | 978 |
| 3203394 | 3204290 | xerC | Rv2894c | 897 | 3157326 | 3158222 | xerC | BCG_2915c | 897 | 3137219 | 3138115 | xerC | BCGDan_2942 | 897 | 3137284 | 3138180 | xerC | BCGDan_sapMKO_2943 | 897 |
| 3204381 | 3205232 | viuB | Rv2895c | 852 | 3158313 | 3159164 | viuB | BCG_2916c | 852 | 3138206 | 3139057 | viuB | BCGDan_2943 | 852 | 3138271 | 3139062 | viuB | BCGDan_sapMKO_2944 | 792 |
| 3205265 | 3206434 | Rv2896c | Rv2896c | 1170 | 3159197 | 3160366 | BCG_2917c | BCG_2917c | 1170 | 3139090 | 3140259 | BCGDan_2944 | BCGDan_2944 | 1170 | 3139155 | 3140324 | BCGDan_sapMKO_2945 | BCGDan_sapMKO_2945 | 1170 |
| 3206431 | 3207942 | Rv2897c | Rv2897c | 1512 | 3160363 | 3161874 | BCG_2918c | BCG_2918c | 1512 | 3140256 | 3141767 | BCGDan_2945 | BCGDan_2945 | 1512 | 3140321 | 3141832 | BCGDan_sapMKO_2946 | BCGDan_sapMKO_2946 | 1512 |
| 3207942 | 3208328 | Rv2898c | Rv2898c | 387 | 3161874 | 3162260 | BCG_2919c | BCG_2919c | 387 | 3141767 | 3142153 | BCGDan_2946 | BCGDan_2946 | 387 | 3141832 | 3142257 | BCGDan_sapMKO_2947 | BCGDan_sapMKO_2947 | 426 |
| 3208576 | 3209406 | fdhD | Rv2899c | 831 | 3162508 | 3163338 | fdhD | BCG_2920c | 831 | 3142401 | 3143231 | fdhD | BCGDan_2947 | 831 | 3142466 | 3143296 | fdhD | BCGDan_sapMKO_2948 | 831 |
| 3209406 | 3211745 | fdhF | Rv2900c | 2340 | 3163338 | 3165677 | fdhF | BCG_2921c | 2340 | 3143231 | 3145570 | fdhF | BCGDan_2948 | 2340 | 3143296 | 3145635 | fdhF | BCGDan_sapMKO_2949 | 2340 |
| 3211803 | 3212108 | Rv2901c | Rv2901c | 306 | 3165735 | 3166040 | BCG_2922c | BCG_2922c | 306 | 3145628 | 3145933 | BCGDan_2949 | BCGDan_2949 | 306 | 3145693 | 3145998 | BCGDan_sapMKO_2950 | BCGDan_sapMKO_2950 | 306 |
| 3212162 | 3212956 | rnhB | Rv2902c | 795 | 3166094 | 3166888 | rnhB | BCG_2923c | 795 | 3145987 | 3146781 | rnhB | BCGDan_2950 | 795 | 3146052 | 3146846 | rnhB | BCGDan_sapMKO_2951 | 795 |
| 3212970 | 3213854 | lepB | Rv2903c | 885 | 3166902 | 3167786 | lepB | BCG_2924c | 885 | 3146795 | 3147679 | lepB | BCGDan_2951 | 885 | 3146860 | 3147744 | lepB | BCGDan_sapMKO_2952 | 885 |
| 3213912 | 3214253 | rplS | Rv2904c | 342 | 3167844 | 3168185 | rplS | BCG_2925c | 342 | 3147737 | 3148078 | rplS | BCGDan_2952 | 342 | 3147802 | 3148143 | rplS | BCGDan_sapMKO_2953 | 342 |
| 3214628 | 3215572 | lppW | Rv2905 | 945 | 3168560 | 3169504 | lppW | BCG_2926 | 945 | 3148453 | 3149397 | lppW | BCGDan_2953 | 945 | 3148518 | 3149462 | lppW | BCGDan_sapMKO_2954 | 945 |
| 3215665 | 3216357 | trmD | Rv2906c | 693 | 3169597 | 3170289 | trmD | BCG_2927c | 693 | 3149490 | 3150182 | trmD | BCGDan_2954 | 693 | 3149555 | 3150247 | trmD | BCGDan_sapMKO_2955 | 693 |
| 3216361 | 3216891 | rimM | Rv2907c | 531 | 3170293 | 3170823 | rimM | BCG_2928c | 531 | 3150186 | 3150716 | rimM | BCGDan_2955 | 531 | 3150251 | 3150781 | rimM | BCGDan_sapMKO_2956 | 531 |
| 3216905 | 3217147 | Rv2908c | Rv2908c | 243 | 3170837 | 3171079 | BCG_2929c | BCG_2929c | 243 | 3150730 | 3150972 | BCGDan_2956 | BCGDan_2956 | 243 | 3150795 | 3151037 | BCGDan_sapMKO_2957 | BCGDan_sapMKO_2957 | 243 |
| 3217155 | 3217643 | rpsP | Rv2909c | 489 | 3171087 | 3171575 | rpsP | BCG_2930c | 489 | 3150980 | 3151468 | rpsP | BCGDan_2957 | 489 | 3151045 | 3151533 | rpsP | BCGDan_sapMKO_2958 | 489 |

| | | | | | | | | | | | | | | | | | | | |
|---|---|---|---|---|---|---|---|---|---|---|---|---|---|---|---|---|---|---|---|
| 3217827 | 3218270 | Rv2910c | Rv2910c | 444 | 3171759 | 3172202 | BCG_2931c | BCG_2931c | 444 | 3151652 | 3152095 | BCGDan_2958 | BCGDan_2958 | 444 | 3151717 | 3152160 | BCGDan_sapMKO_2959 | BCGDan_sapMKO_2959 | 444 |
| 3218339 | 3219214 | dacB2 | Rv2911 | 876 | 3172271 | 3173146 | dacB2 | BCG_2932 | 876 | 3152164 | 3153039 | dacB2 | BCGDan_2959 | 876 | 3152229 | 3153104 | dacB2 | BCGDan_sapMKO_2960 | 876 |
| 3219274 | 3219861 | Rv2912c | Rv2912c | 588 | 3173206 | 3173793 | BCG_2933c | BCG_2933c | 588 | 3153099 | 3153686 | BCGDan_2960 | BCGDan_2960 | 588 | 3153164 | 3153751 | BCGDan_sapMKO_2961 | BCGDan_sapMKO_2961 | 588 |
| 3219863 | 3221698 | Rv2913c | Rv2913c | 1836 | 3173795 | 3175630 | BCG_2934c | BCG_2934c | 1836 | 3153688 | 3155523 | BCGDan_2961 | BCGDan_2961 | 1836 | 3153753 | 3155588 | BCGDan_sapMKO_2962 | BCGDan_sapMKO_2962 | 1836 |
| 3221767 | 3223524 | pknI | Rv2914c | 1758 | 3175699 | 3177456 | pknI | BCG_2935c | 1758 | 3155592 | 3157349 | pknI | BCGDan_2962 | 1758 | 3155657 | 3157414 | pknI | BCGDan_sapMKO_2963 | 1758 |
| 3223568 | 3224680 | Rv2915c | Rv2915c | 1113 | 3177500 | 3178612 | BCG_2936c | BCG_2936c | 1113 | 3157393 | 3158505 | BCGDan_2963 | BCGDan_2963 | 1113 | 3157458 | 3158531 | BCGDan_sapMKO_2964 | BCGDan_sapMKO_2964 | 1074 |
| 3224708 | 3226285 | ffh | Rv2916c | 1578 | 3178640 | 3180217 | ffh | BCG_2937c | 1578 | 3158533 | 3160110 | ffh | BCGDan_2964 | 1578 | 3158598 | 3160175 | ffh | BCGDan_sapMKO_2965 | 1578 |
| 3226363 | 3228243 | Rv2917 | Rv2917 | 1881 | 3180295 | 3182175 | BCG_2938 | BCG_2938 | 1881 | 3160188 | 3162068 | BCGDan_2965 | BCGDan_2965 | 1881 | 3160412 | 3162133 | BCGDan_sapMKO_2966 | BCGDan_sapMKO_2966 | 1722 |
| 3228254 | 3230680 | glnD | Rv2918c | 2427 | 3182186 | 3184612 | glnD | BCG_2939c | 2427 | 3162079 | 3164505 | glnD | BCGDan_2966 | 2427 | 3162144 | 3164570 | glnD | BCGDan_sapMKO_2967 | 2427 |
| 3230738 | 3231076 | glnB | Rv2919c | 339 | 3184670 | 3185008 | glnB | BCG_2940c | 339 | 3164563 | 3164901 | glnB | BCGDan_2967 | 339 | 3164628 | 3164966 | glnB | BCGDan_sapMKO_2968 | 339 |
| 3231073 | 3232506 | amt | Rv2920c | 1434 | 3185005 | 3186438 | amt | BCG_2941c | 1434 | 3164898 | 3166331 | amt | BCGDan_2968 | 1434 | 3164963 | 3166396 | amt | BCGDan_sapMKO_2969 | 1434 |
| 3232871 | 3234139 | ftsY | Rv2921c | 1269 | 3186915 | 3188183 | ftsY | BCG_2942c | 1269 | 3166752 | 3168020 | ftsY | BCGDan_2969 | 1269 | 3166817 | 3168085 | ftsY | BCGDan_sapMKO_2970 | 1269 |
| 3234189 | 3237806 | smc | Rv2922c | 3618 | 3188233 | 3191850 | smc | BCG_2943c | 3618 | 3168070 | 3171687 | smc | BCGDan_2970 | 3618 | 3168135 | 3171752 | smc | BCGDan_sapMKO_2971 | 3618 |
| 3237818 | 3238099 | acyP | Rv2922A | 282 | 3191862 | 3192143 | acyP | BCG_2944c | 282 | 3171699 | 3171980 | acyP | BCGDan_2971 | 282 | 3171764 | 3172045 | acyP | BCGDan_sapMKO_2972 | 282 |
| 3238086 | 3238499 | Rv2923c | Rv2923c | 414 | 3192130 | 3192543 | BCG_2945c | BCG_2945c | 414 | 3171967 | 3172380 | BCGDan_2972 | BCGDan_2972 | 414 | 3172032 | 3172445 | BCGDan_sapMKO_2973 | BCGDan_sapMKO_2973 | 414 |
| 3238601 | 3239470 | fpg | Rv2924c | 870 | 3192645 | 3193514 | fpg | BCG_2946c | 870 | 3172482 | 3173351 | fpg | BCGDan_2973 | 870 | 3172547 | 3173518 | fpg | BCGDan_sapMKO_2974 | 972 |
| 3239829 | 3240551 | rnc | Rv2925c | 723 | 3193818 | 3194540 | rnc | BCG_2947c | 723 | 3173576 | 3174298 | rnc | BCGDan_2974 | 723 | 3173641 | 3174363 | rnc | BCGDan_sapMKO_2975 | 723 |
| 3240548 | 3241171 | Rv2926c | Rv2926c | 624 | 3194537 | 3195160 | BCG_2948c | BCG_2948c | 624 | 3174295 | 3174918 | BCGDan_2975 | BCGDan_2975 | 624 | 3174360 | 3174899 | BCGDan_sapMKO_2976 | BCGDan_sapMKO_2976 | 540 |
| 3241222 | 3241959 | Rv2927c | Rv2927c | 738 | 3195211 | 3195948 | BCG_2949c | BCG_2949c | 738 | 3174969 | 3175706 | BCGDan_2976 | BCGDan_2976 | 738 | 3175034 | 3175777 | BCGDan_sapMKO_2977 | BCGDan_sapMKO_2977 | 744 |
| 3242198 | 3242983 | tesA | Rv2928 | 786 | 3196187 | 3196972 | tesA | BCG_2950 | 786 | 3175945 | 3176730 | tesA | BCGDan_2977 | 786 | 3176004 | 3176795 | tesA | BCGDan_sapMKO_2978 | 792 |
| 3242970 | 3243281 | Rv2929 | Rv2929 | 312 | 3196959 | 3197270 | BCG_2951 | BCG_2951 | 312 | 3176717 | 3177028 | BCGDan_2978 | BCGDan_2978 | 312 | 3176782 | 3177093 | BCGDan_sapMKO_2979 | BCGDan_sapMKO_2979 | 312 |
| 3243697 | 3245448 | fadD26 | Rv2930 | 1752 | 3197686 | 3199437 | fadD26 | BCG_2952 | 1752 | 3177444 | 3179195 | fadD26 | BCGDan_2979 | 1752 | 3177509 | 3179260 | fadD26 | BCGDan_sapMKO_2980 | 1752 |
| 3245445 | 3251075 | ppsA | Rv2931 | 5631 | 3199434 | 3205064 | ppsA | BCG_2953 | 5631 | 3179192 | 3184822 | ppsA | BCGDan_2980 | 5631 | 3179257 | 3184887 | ppsA | BCGDan_sapMKO_2981 | 5631 |
| 3251072 | 3255688 | ppsB | Rv2932 | 4617 | 3205061 | 3209677 | ppsB | BCG_2954 | 4617 | 3184819 | 3189435 | ppsB | BCGDan_2981 | 4617 | 3184887 | 3189500 | ppsB | BCGDan_sapMKO_2982 | 4614 |
| 3255685 | 3262251 | ppsC | Rv2933 | 6567 | 3209674 | 3216240 | ppsC | BCG_2955 | 6567 | 3189432 | 3195998 | ppsC | BCGDan_2982 | 6567 | 3189497 | 3196063 | ppsC | BCGDan_sapMKO_2983 | 6567 |
| 3262248 | 3267731 | ppsD | Rv2934 | 5484 | 3216237 | 3221720 | ppsD | BCG_2956 | 5484 | 3195995 | 3201478 | ppsD | BCGDan_2983 | 5484 | 3196060 | 3201543 | ppsD | BCGDan_sapMKO_2984 | 5484 |
| 3267737 | 3272203 | ppsE | Rv2935 | 4467 | 3221726 | 3226192 | ppsE | BCG_2957 | 4467 | 3201484 | 3205950 | ppsE | BCGDan_2984 | 4467 | 3201549 | 3206015 | ppsE | BCGDan_sapMKO_2985 | 4467 |
| 3272214 | 3273209 | drrA | Rv2936 | 996 | 3226203 | 3227198 | drrA | BCG_2958 | 996 | 3205961 | 3206956 | drrA | BCGDan_2985 | 996 | 3206005 | 3207021 | drrA | BCGDan_sapMKO_2986 | 1017 |
| 3273206 | 3274075 | drrB | Rv2937 | 870 | 3227195 | 3228064 | drrB | BCG_2959 | 870 | 3206953 | 3207822 | drrB | BCGDan_2986 | 870 | 3207111 | 3207887 | drrB | BCGDan_sapMKO_2987 | 777 |
| 3274072 | 3274902 | drrC | Rv2938 | 831 | 3228061 | 3228891 | drrC | BCG_2960 | 831 | 3207819 | 3208649 | drrC | BCGDan_2987 | 831 | 3207884 | 3208714 | drrC | BCGDan_sapMKO_2988 | 831 |
| 3274949 | 3276217 | papA5 | Rv2939 | 1269 | 3228938 | 3230206 | papA5 | BCG_2961 | 1269 | 3208696 | 3209964 | papA5 | BCGDan_2988 | 1269 | 3208761 | 3210029 | papA5 | BCGDan_sapMKO_2989 | 1269 |
| 3276380 | 3282715 | mas | Rv2940c | 6336 | 3230369 | 3236704 | mas | BCG_2962c | 6336 | 3210127 | 3216462 | mas | BCGDan_2989 | 6336 | 3210192 | 3216527 | mas | BCGDan_sapMKO_2990 | 6336 |
| 3283335 | 3285077 | fadD28 | Rv2941 | 1743 | 3237324 | 3239066 | fadD28 | BCG_2963 | 1743 | 3217082 | 3218824 | fadD28 | BCGDan_2990 | 1743 | 3217222 | 3218889 | fadD28 | BCGDan_sapMKO_2991 | 1668 |
| 3285070 | 3287832 | mmpL7 | Rv2942 | 2763 | 3239059 | 3241821 | mmpL7 | BCG_2964 | 2763 | 3218817 | 3221579 | mmpL7 | BCGDan_2991 | 2763 | 3218927 | 3221644 | mmpL7 | BCGDan_sapMKO_2992 | 2718 |
| 3288464 | 3289705 | Rv2943 | Rv2943 | 1242 | 3242453 | 3243694 | BCG_2965 | BCG_2965 | 1242 | 3222211 | 3223452 | BCGDan_2992 | BCGDan_2992 | 1242 | 3222276 | 3223517 | BCGDan_sapMKO_2993 | BCGDan_sapMKO_2993 | 1242 |
| 3289705 | 3290235 | Rv2943A | Rv2943A | 531 | 3243694 | 3244224 | BCG_2966 | BCG_2966 | 531 | 3223452 | 3223982 | BCGDan_2993 | BCGDan_2993 | 531 | 3223517 | 3224047 | BCGDan_sapMKO_2994 | BCGDan_sapMKO_2994 | 531 |
| 3290624 | 3291325 | lppX | Rv2945c | 702 | 3244613 | 3245314 | lppX | BCG_2967c | 702 | 3224371 | 3225072 | lppX | BCGDan_2994 | 702 | 3224436 | 3225137 | lppX | BCGDan_sapMKO_2995 | 702 |
| 3291503 | 3296353 | pks1 | Rv2946c | 4851 | 3245663 | 3252001 | pks1 | BCG_2968c | 6339 | 3225421 | 3231759 | pks1 | BCGDan_2995 | 6339 | 3225486 | 3231824 | pks1 | BCGDan_sapMKO_2996 | 6339 |
| 3297837 | 3299954 | fadD22 | Rv2948c | 2118 | 3251998 | 3254115 | fadD22 | BCG_2969c | 2118 | 3231756 | 3233873 | fadD22 | BCGDan_2996 | 2118 | 3231821 | 3233938 | fadD22 | BCGDan_sapMKO_2997 | 2118 |
| 3299971 | 3300570 | Rv2949c | Rv2949c | 600 | 3254132 | 3254731 | BCG_2970c | BCG_2970c | 600 | 3233890 | 3234489 | BCGDan_2997 | BCGDan_2997 | 600 | 3233955 | 3234569 | BCGDan_sapMKO_2998 | BCGDan_sapMKO_2998 | 615 |
| 3300596 | 3302455 | fadD29 | Rv2950c | 1860 | 3254757 | 3256616 | fadD29 | BCG_2971c | 1860 | 3234515 | 3236374 | fadD29 | BCGDan_2998 | 1860 | 3234580 | 3236439 | fadD29 | BCGDan_sapMKO_2999 | 1860 |
| 3303103 | 3304248 | Rv2951c | Rv2951c | 1146 | 3257264 | 3258409 | BCG_2972c | BCG_2972c | 1146 | 3237022 | 3238167 | BCGDan_2999 | BCGDan_2999 | 1146 | 3237087 | 3238232 | BCGDan_sapMKO_3000 | BCGDan_sapMKO_3000 | 1146 |
| 3304441 | 3305253 | Rv2952 | Rv2952 | 813 | 3258602 | 3259414 | BCG_2973 | BCG_2973 | 813 | 3238360 | 3239172 | BCGDan_3000 | BCGDan_3000 | 813 | 3238512 | 3239237 | BCGDan_sapMKO_3001 | BCGDan_sapMKO_3001 | 726 |
| 3305279 | 3306535 | Rv2953 | Rv2953 | 1257 | 3259440 | 3260696 | BCG_2974 | BCG_2974 | 1257 | 3239198 | 3240454 | BCGDan_3001 | BCGDan_3001 | 1257 | 3239263 | 3240519 | BCGDan_sapMKO_3002 | BCGDan_sapMKO_3002 | 1257 |
| 3306666 | 3307391 | Rv2954c | Rv2954c | 726 | 3260827 | 3261552 | BCG_2975c | BCG_2975c | 726 | 3240585 | 3241310 | BCGDan_3002 | BCGDan_3002 | 726 | 3240650 | 3241393 | BCGDan_sapMKO_3003 | BCGDan_sapMKO_3003 | 744 |
| 3307580 | 3308545 | Rv2955c | Rv2955c | 966 | 3261741 | 3262706 | BCG_2976c | BCG_2976c | 966 | 3241499 | 3242464 | BCGDan_3003 | BCGDan_3003 | 966 | 3241564 | 3242538 | BCGDan_sapMKO_3004 | BCGDan_sapMKO_3004 | 975 |
| 3308668 | 3309399 | Rv2956 | Rv2956 | 732 | 3262829 | 3263560 | BCG_2977 | BCG_2977 | 732 | 3242587 | 3243318 | BCGDan_3004 | BCGDan_3004 | 732 | 3242652 | 3243383 | BCGDan_sapMKO_3005 | BCGDan_sapMKO_3005 | 732 |
| 3309470 | 3310297 | Rv2957 | Rv2957 | 828 | 3263631 | 3264458 | BCG_2978 | BCG_2978 | 828 | 3243389 | 3244216 | BCGDan_3005 | BCGDan_3005 | 828 | 3243454 | 3244281 | BCGDan_sapMKO_3006 | BCGDan_sapMKO_3006 | 828 |
| NA | NA | NA | NA | NA | NA | NA | NA | NA | NA | 3244633 | 3244767 | BCGDan_3006 | BCGDan_3006 | 135 | 3244698 | 3244832 | BCGDan_sapMKO_3007 | BCGDan_sapMKO_3007 | 135 |
| 3310714 | 3312000 | Rv2958c | Rv2958c | 1287 | 3265062 | 3266162 | BCG_2979c | BCG_2979c | 1101 | 3244820 | 3245920 | BCGDan_3007 | BCGDan_3007 | 1101 | 3244885 | 3245985 | BCGDan_sapMKO_3008 | BCGDan_sapMKO_3008 | 1101 |
| 3312101 | 3312838 | Rv2959c | Rv2959c | 738 | 3266263 | 3267000 | BCG_2980c | BCG_2980c | 738 | 3246021 | 3246758 | BCGDan_3008 | BCGDan_3008 | 738 | 3246086 | 3246760 | BCGDan_sapMKO_3009 | BCGDan_sapMKO_3009 | 675 |
| 3312953 | 3313201 | Rv2960c | Rv2960c | 249 | 3267115 | 3267363 | BCG_2981c | BCG_2981c | 249 | 3246873 | 3247121 | BCGDan_3009 | BCGDan_3009 | 249 | 3246938 | 3247186 | BCGDan_sapMKO_3010 | BCGDan_sapMKO_3010 | 249 |
| 3313283 | 3313672 | Rv2961 | Rv2961 | 390 | 3267445 | 3267834 | BCG_2982 | BCG_2982 | 390 | 3247203 | 3247592 | BCGDan_3010 | BCGDan_3010 | 390 | 3247268 | 3247657 | BCGDan_sapMKO_3011 | BCGDan_sapMKO_3011 | 390 |
| 3313773 | 3315122 | Rv2962c | Rv2962c | 1350 | 3267935 | 3269284 | BCG_2983c | BCG_2983c | 1350 | 3247693 | 3249042 | BCGDan_3011 | BCGDan_3011 | 1350 | 3247758 | 3248954 | BCGDan_sapMKO_3012 | BCGDan_sapMKO_3012 | 1197 |
| 3315236 | 3316456 | Rv2963 | Rv2963 | 1221 | 3269398 | 3270618 | BCG_2984 | BCG_2984 | 1221 | 3249156 | 3250376 | BCGDan_3012 | BCGDan_3012 | 1221 | 3249260 | 3250441 | BCGDan_sapMKO_3013 | BCGDan_sapMKO_3013 | 1182 |
| 3316529 | 3317461 | purU | Rv2964 | 933 | 3270691 | 3271623 | purU | BCG_2985 | 933 | 3250449 | 3251381 | purU | BCGDan_3013 | 933 | 3250514 | 3251446 | purU | BCGDan_sapMKO_3014 | 933 |
| 3318330 | 3318815 | kdtB | Rv2965c | 486 | 3272492 | 3272977 | kdtB | BCG_2986c | 486 | 3252250 | 3252735 | kdtB | BCGDan_3014 | 486 | 3252315 | 3252764 | kdtB | BCGDan_sapMKO_3015 | 450 |
| 3318901 | 3319467 | Rv2966c | Rv2966c | 567 | 3273063 | 3273629 | BCG_2987c | BCG_2987c | 567 | 3252821 | 3253387 | BCGDan_3015 | BCGDan_3015 | 567 | 3252886 | 3253452 | BCGDan_sapMKO_3016 | BCGDan_sapMKO_3016 | 567 |
| 3319663 | 3323046 | pca | Rv2967c | 3384 | 3273825 | 3277208 | pca | BCG_2988c | 3384 | 3253583 | 3256966 | pca | BCGDan_3016 | 3384 | 3253648 | 3257031 | pca | BCGDan_sapMKO_3017 | 3384 |
| 3323071 | 3323703 | Rv2968c | Rv2968c | 633 | 3277233 | 3277865 | BCG_2989c | BCG_2989c | 633 | 3256991 | 3257623 | BCGDan_3017 | BCGDan_3017 | 633 | 3257056 | 3257688 | BCGDan_sapMKO_3018 | BCGDan_sapMKO_3018 | 633 |
| 3323709 | 3324476 | Rv2969c | Rv2969c | 768 | 3277871 | 3278638 | BCG_2990c | BCG_2990c | 768 | 3257629 | 3258396 | BCGDan_3018 | BCGDan_3018 | 768 | 3257694 | 3258461 | BCGDan_sapMKO_3019 | BCGDan_sapMKO_3019 | 768 |
| 3324573 | 3325703 | lipN | Rv2970c | 1131 | 3278735 | 3279865 | lipN | BCG_2991c | 1131 | 3258493 | 3259623 | lipN | BCGDan_3019 | 1131 | 3258558 | 3259631 | lipN | BCGDan_sapMKO_3020 | 1074 |
| 3325934 | 3326104 | Rv2970A | Rv2970A | 171 | 3280096 | 3280266 | BCG_2992 | BCG_2992 | 171 | 3259854 | 3260024 | BCGDan_3020 | BCGDan_3020 | 171 | 3259919 | 3260089 | BCGDan_sapMKO_3021 | BCGDan_sapMKO_3021 | 171 |
| 3326101 | 3326949 | Rv2971 | Rv2971 | 849 | 3280263 | 3281111 | BCG_2993 | BCG_2993 | 849 | 3260021 | 3260869 | BCGDan_3021 | BCGDan_3021 | 849 | 3260086 | 3260934 | BCGDan_sapMKO_3022 | BCGDan_sapMKO_3022 | 849 |
| 3327023 | 3327736 | Rv2972c | Rv2972c | 714 | 3281185 | 3281898 | BCG_2994c | BCG_2994c | 714 | 3260943 | 3261656 | BCGDan_3022 | BCGDan_3022 | 714 | 3261008 | 3261721 | BCGDan_sapMKO_3023 | BCGDan_sapMKO_3023 | 714 |
| 3327733 | 3329946 | recG | Rv2973c | 2214 | 3281895 | 3284108 | recG | BCG_2995c | 2214 | 3261653 | 3263866 | recG | BCGDan_3023 | 2214 | 3261718 | 3263931 | recG | BCGDan_sapMKO_3024 | 2214 |
| 3329949 | 3331361 | Rv2974c | Rv2974c | 1413 | 3284111 | 3285772 | BCG_2996c | BCG_2996c | 1662 | 3263869 | 3265530 | BCGDan_3024 | BCGDan_3024 | 1662 | 3263934 | 3265595 | BCGDan_sapMKO_3025 | BCGDan_sapMKO_3025 | 1662 |
| NA | NA | NA | NA | NA | NA | NA | NA | NA | NA | 3265530 | 3265736 | BCGDan_3025 | BCGDan_3025 | 207 | 3265598 | 3265801 | BCGDan_sapMKO_3026 | BCGDan_sapMKO_3026 | 204 |
| NA | NA | NA | NA | NA | NA | NA | NA | NA | NA | 3265772 | 3265966 | BCGDan_3026 | BCGDan_3026 | 195 | 3265837 | 3266031 | BCGDan_sapMKO_3027 | BCGDan_sapMKO_3027 | 195 |
| 3332071 | 3332754 | ung | Rv2976c | 684 | 3286231 | 3286914 | ung | BCG_2997c | 684 | 3265989 | 3266672 | ung | BCGDan_3027 | 684 | 3266054 | 3266737 | ung | BCGDan_sapMKO_3028 | 684 |
| 3332787 | 3333788 | thiL | Rv2977c | 1002 | 3286947 | 3287948 | thiL | BCG_2998c | 1002 | 3266705 | 3267706 | thiL | BCGDan_3028 | 1002 | 3266770 | 3267771 | thiL | BCGDan_sapMKO_3029 | 1002 |
| 3333785 | 3335164 | Rv2978c | Rv2978c | 1380 | 3287945 | 3289324 | BCG_2999c | BCG_2999c | 1380 | 3267703 | 3269082 | BCGDan_3029 | BCGDan_3029 | 1380 | 3267768 | 3269147 | BCGDan_sapMKO_3030 | BCGDan_sapMKO_3030 | 1380 |
| 3335164 | 3335748 | Rv2979c | Rv2979c | 585 | 3289324 | 3289908 | BCG_3000c | BCG_3000c | 585 | 3269082 | 3269666 | BCGDan_3030 | BCGDan_3030 | 585 | 3269147 | 3269731 | BCGDan_sapMKO_3031 | BCGDan_sapMKO_3031 | 585 |
| 3335960 | 3336505 | Rv2980 | Rv2980 | 546 | 3290120 | 3290665 | BCG_3001 | BCG_3001 | 546 | 3269878 | 3270423 | BCGDan_3031 | BCGDan_3031 | 546 | 3269913 | 3270488 | BCGDan_sapMKO_3032 | BCGDan_sapMKO_3032 | 576 |
| 3336796 | 3337917 | ddlA | Rv2981c | 1122 | 3291310 | 3292431 | ddlA | BCG_3002c | 1122 | 3271127 | 3272248 | ddlA | BCGDan_3032 | 1122 | 3271192 | 3272268 | ddlA | BCGDan_sapMKO_3033 | 1077 |
| 3337995 | 3338999 | gpdA2 | Rv2982c | 1005 | 3292509 | 3293513 | gpdA2 | BCG_3003c | 1005 | 3272326 | 3273330 | gpdA2 | BCGDan_3033 | 1005 | 3272391 | 3273395 | gpdA2 | BCGDan_sapMKO_3034 | 1005 |
| 3339118 | 3339762 | Rv2983 | Rv2983 | 645 | 3293632 | 3294276 | BCG_3004 | BCG_3004 | 645 | 3273449 | 3274093 | BCGDan_3034 | BCGDan_3034 | 645 | 3273514 | 3274158 | BCGDan_sapMKO_3035 | BCGDan_sapMKO_3035 | 645 |
| 3339854 | 3342082 | ppk1 | Rv2984 | 2229 | 3294368 | 3296596 | ppk | BCG_3005 | 2229 | 3274185 | 3276413 | ppk | BCGDan_3035 | 2229 | 3274250 | 3276478 | ppk | BCGDan_sapMKO_3036 | 2229 |
| 3342165 | 3343118 | mutT1 | Rv2985 | 954 | 3296679 | 3297632 | mutT1 | BCG_3006 | 954 | 3276496 | 3277449 | mutT1 | BCGDan_3036 | 954 | 3276561 | 3277514 | mutT1 | BCGDan_sapMKO_3037 | 954 |
| 3343176 | 3343820 | hupB | Rv2986c | 645 | 3297690 | 3298307 | hupB | BCG_3007c | 618 | 3277507 | 3278124 | hupB | BCGDan_3037 | 618 | 3277572 | 3278189 | hupB | BCGDan_sapMKO_3038 | 618 |
| 3344033 | 3344629 | leuD | Rv2987c | 597 | 3298520 | 3299116 | leuD | BCG_3008c | 597 | 3278337 | 3278933 | leuD | BCGDan_3038 | 597 | 3278402 | 3278998 | leuD | BCGDan_sapMKO_3039 | 597 |
| 3344654 | 3346075 | leuC | Rv2988c | 1422 | 3299141 | 3300562 | leuC | BCG_3009c | 1422 | 3278958 | 3280379 | leuC | BCGDan_3039 | 1422 | 3279023 | 3280444 | leuC | BCGDan_sapMKO_3040 | 1422 |
| 3346147 | 3346848 | Rv2989 | Rv2989 | 702 | 3300634 | 3301335 | BCG_3010 | BCG_3010 | 702 | 3280451 | 3281152 | BCGDan_3040 | BCGDan_3040 | 702 | 3280516 | 3281217 | BCGDan_sapMKO_3041 | BCGDan_sapMKO_3041 | 702 |
| 3346859 | 3347719 | Rv2990c | Rv2990c | 861 | 3301346 | 3302206 | BCG_3011c | BCG_3011c | 861 | 3281163 | 3282023 | BCGDan_3041 | BCGDan_3041 | 861 | 3281228 | 3282088 | BCGDan_sapMKO_3042 | BCGDan_sapMKO_3042 | 861 |
| 3347982 | 3348473 | Rv2991 | Rv2991 | 492 | 3302469 | 3302960 | BCG_3012 | BCG_3012 | 492 | 3282286 | 3282777 | BCGDan_3042 | BCGDan_3042 | 492 | 3282351 | 3282842 | BCGDan_sapMKO_3043 | BCGDan_sapMKO_3043 | 492 |
| 3348805 | 3350277 | gltS | Rv2992c | 1473 | 3303292 | 3304764 | gltS | BCG_3013c | 1473 | 3283109 | 3284581 | gltS | BCGDan_3045 | 1473 | 3283174 | 3284646 | gltS | BCGDan_sapMKO_3046 | 1473 |

| | | | | | | | | | | | | | | | | | | | |
|---|---|---|---|---|---|---|---|---|---|---|---|---|---|---|---|---|---|---|---|
| 3350274 | 3350993 | Rv2993c | Rv2993c | 720 | 3304761 | 3305480 | BCG_3014c | BCG_3014c | 720 | 3284578 | 3285297 | BCGDan_3046 | BCGDan_3046 | 720 | 3284643 | 3285440 | BCGDan_sapMKO_3047 | BCGDan_sapMKO_3047 | 798 |
| 3351269 | 3352606 | Rv2994 | Rv2994 | 1338 | 3305756 | 3307093 | BCG_3015 | BCG_3015 | 1338 | 3285573 | 3286910 | BCGDan_3047 | BCGDan_3047 | 1338 | 3285638 | 3286975 | BCGDan_sapMKO_3048 | BCGDan_sapMKO_3048 | 1338 |
| 3352458 | 3353468 | leuB | Rv2995c | 1011 | 3306945 | 3307955 | leuB | BCG_3016c | 1011 | 3286762 | 3287772 | leuB | BCGDan_3048 | 1011 | 3286827 | 3287837 | leuB | BCGDan_sapMKO_3049 | 1011 |
| 3353483 | 3355069 | serA1 | Rv2996c | 1587 | 3307970 | 3309556 | serA1 | BCG_3017c | 1587 | 3287787 | 3289373 | serA1 | BCGDan_3049 | 1587 | 3287852 | 3289423 | serA1 | BCGDan_sapMKO_3050 | 1572 |
| 3355099 | 3356541 | Rv2997 | Rv2997 | 1443 | 3309586 | 3311028 | BCG_3018 | BCG_3018 | 1443 | 3289403 | 3290845 | BCGDan_3050 | BCGDan_3050 | 1443 | 3289468 | 3290910 | BCGDan_sapMKO_3051 | BCGDan_sapMKO_3051 | 1443 |
| 3356815 | 3357276 | Rv2998 | Rv2998 | 462 | 3311302 | 3311763 | BCG_3019 | BCG_3019 | 462 | 3291119 | 3291580 | BCGDan_3051 | BCGDan_3051 | 462 | 3291184 | 3291645 | BCGDan_sapMKO_3052 | BCGDan_sapMKO_3052 | 462 |
| 3357225 | 3357428 | Rv2998A | Rv2998A | 204 | 3311712 | 3311915 | BCG_3020c | BCG_3020c | 204 | 3291529 | 3291732 | BCGDan_3052 | BCGDan_3052 | 204 | 3291594 | 3291797 | BCGDan_sapMKO_3053 | BCGDan_sapMKO_3053 | 204 |
| 3357602 | 3358567 | lppY | Rv2999 | 966 | 3312089 | 3313054 | lppY | BCG_3021 | 966 | 3291906 | 3292871 | lppY | BCGDan_3053 | 966 | 3291956 | 3292936 | lppY | BCGDan_sapMKO_3054 | 981 |
| 3358612 | 3359271 | Rv3000 | Rv3000 | 660 | 3313099 | 3313758 | BCG_3022 | BCG_3022 | 660 | 3292916 | 3293575 | BCGDan_3054 | BCGDan_3054 | 660 | 3292981 | 3293640 | BCGDan_sapMKO_3055 | BCGDan_sapMKO_3055 | 660 |
| 3359585 | 3360586 | ilvC | Rv3001c | 1002 | 3314072 | 3315073 | ilvC | BCG_3023c | 1002 | 3293889 | 3294890 | ilvC | BCGDan_3055 | 1002 | 3293954 | 3294955 | ilvC | BCGDan_sapMKO_3056 | 1002 |
| 3360624 | 3361130 | ilvN | Rv3002c | 507 | 3315111 | 3315617 | ilvN | BCG_3024c | 507 | 3294928 | 3295434 | ilvN | BCGDan_3056 | 507 | 3294993 | 3295499 | ilvN | BCGDan_sapMKO_3057 | 507 |
| 3361130 | 3362986 | ilvB1 | Rv3003c | 1857 | 3315617 | 3317473 | ilvB1 | BCG_3025c | 1857 | 3295434 | 3297290 | ilvB1 | BCGDan_3057 | 1857 | 3295499 | 3297355 | ilvB1 | BCGDan_sapMKO_3058 | 1857 |
| 3363348 | 3363686 | cfp6 | Rv3004 | 339 | 3317835 | 3318173 | cfp6 | BCG_3026 | 339 | 3297652 | 3297990 | cfp6 | BCGDan_3058 | 339 | 3297678 | 3298055 | cfp6 | BCGDan_sapMKO_3059 | 378 |
| 3363693 | 3364532 | Rv3005c | Rv3005c | 840 | 3318180 | 3319019 | BCG_3027c | BCG_3027c | 840 | 3297997 | 3298836 | BCGDan_3059 | BCGDan_3059 | 840 | 3298062 | 3298847 | BCGDan_sapMKO_3060 | BCGDan_sapMKO_3060 | 786 |
| 3364709 | 3365830 | lppZ | Rv3006 | 1122 | 3319196 | 3320317 | lppZ | BCG_3028 | 1122 | 3299013 | 3300134 | lppZ | BCGDan_3060 | 1122 | 3299015 | 3300199 | lppZ | BCGDan_sapMKO_3061 | 1185 |
| 3365836 | 3366450 | Rv3007c | Rv3007c | 615 | 3320323 | 3320937 | BCG_3029c | BCG_3029c | 615 | 3300140 | 3300754 | BCGDan_3061 | BCGDan_3061 | 615 | 3300205 | 3300783 | BCGDan_sapMKO_3062 | BCGDan_sapMKO_3062 | 579 |
| 3366644 | 3367267 | Rv3008 | Rv3008 | 624 | 3321131 | 3321754 | BCG_3030 | BCG_3030 | 624 | 3300948 | 3301571 | BCGDan_3062 | BCGDan_3062 | 624 | 3301013 | 3301636 | BCGDan_sapMKO_3063 | BCGDan_sapMKO_3063 | 624 |
| 3367264 | 3368793 | gatB | Rv3009c | 1530 | 3321751 | 3323280 | gatB | BCG_3031c | 1530 | 3301568 | 3303097 | gatB | BCGDan_3063 | 1530 | 3301633 | 3303162 | gatB | BCGDan_sapMKO_3064 | 1530 |
| 3368823 | 3369854 | pfkA | Rv3010c | 1032 | 3323310 | 3324341 | pfkA | BCG_3032c | 1032 | 3303127 | 3304158 | pfkA | BCGDan_3064 | 1032 | 3303192 | 3304223 | pfkA | BCGDan_sapMKO_3065 | 1032 |
| 3369950 | 3371434 | gatA | Rv3011c | 1485 | 3324437 | 3325921 | gatA | BCG_3033c | 1485 | 3304254 | 3305738 | gatA | BCGDan_3065 | 1485 | 3304319 | 3305803 | gatA | BCGDan_sapMKO_3066 | 1485 |
| 3371431 | 3371730 | gatC | Rv3012c | 300 | 3325918 | 3326217 | gatC | BCG_3034c | 300 | 3305735 | 3306034 | gatC | BCGDan_3066 | 300 | 3305800 | 3306099 | gatC | BCGDan_sapMKO_3067 | 300 |
| 3371815 | 3372471 | Rv3013 | Rv3013 | 657 | 3326302 | 3326958 | BCG_3035 | BCG_3035 | 657 | 3306119 | 3306775 | BCGDan_3067 | BCGDan_3067 | 657 | 3306175 | 3306840 | BCGDan_sapMKO_3068 | BCGDan_sapMKO_3068 | 666 |
| 3372545 | 3374620 | ligA | Rv3014c | 2076 | 3327032 | 3329107 | ligA | BCG_3036c | 2076 | 3306849 | 3308924 | ligA | BCGDan_3068 | 2076 | 3306914 | 3308989 | ligA | BCGDan_sapMKO_3069 | 2076 |
| 3374651 | 3375664 | Rv3015c | Rv3015c | 1014 | 3329138 | 3330151 | BCG_3037c | BCG_3037c | 1014 | 3308955 | 3309968 | BCGDan_3069 | BCGDan_3069 | 1014 | 3309020 | 3310033 | BCGDan_sapMKO_3070 | BCGDan_sapMKO_3070 | 1014 |
| 3375758 | 3376387 | lpqA | Rv3016 | 630 | 3330245 | 3330874 | lpqA | BCG_3038 | 630 | 3310062 | 3310691 | lpqA | BCGDan_3070 | 630 | 3310127 | 3310756 | lpqA | BCGDan_sapMKO_3071 | 630 |
| 3376490 | 3376852 | esxQ | Rv3017c | 363 | 3330977 | 3331339 | esxQ | BCG_3039c | 363 | 3310794 | 3311156 | esxQ | BCGDan_3071 | 363 | 3310859 | 3311224 | esxQ | BCGDan_sapMKO_3072 | 366 |
| 3376939 | 3378243 | PPE46 | Rv3018c | 1305 | 3331426 | 3332739 | PPE46 | BCG_3040c | 1314 | 3311243 | 3312556 | PPE46 | BCGDan_3072 | 1314 | 3311308 | 3312645 | PPE46 | BCGDan_sapMKO_3073 | 1338 |
| 3378711 | 3379001 | esxR | Rv3019c | 291 | 3333142 | 3333432 | esxR | BCG_3042c | 291 | 3312959 | 3313249 | esxR | BCGDan_3073 | 291 | 3313024 | 3313314 | esxR | BCGDan_sapMKO_3074 | 291 |
| 3379036 | 3379329 | esxS | Rv3020c | 294 | 3333467 | 3333760 | esxS | BCG_3043c | 294 | 3313284 | 3313577 | esxS | BCGDan_3074 | 294 | 3313349 | 3313642 | esxS | BCGDan_sapMKO_3075 | 294 |
| 3379376 | 3380452 | PPE47 | Rv3021c | 1077 | 3333807 | 3335114 | PPE47 | BCG_3044c | 1308 | 3313624 | 3314931 | PPE47 | BCGDan_3075 | 1308 | 3313689 | 3314996 | PPE47 | BCGDan_sapMKO_3076 | 1308 |
| 3380679 | 3380993 | PE29 | Rv3022A | 315 | 3335111 | 3335425 | PE29 | BCG_3045c | 315 | 3314928 | 3315242 | PE29 | BCGDan_3076 | 315 | 3314993 | 3315316 | PE29 | BCGDan_sapMKO_3077 | 324 |
| 3381375 | 3382622 | Rv3023c | Rv3023c | 1248 | 3335807 | 3337054 | BCG_3046c | BCG_3046c | 1248 | 3315624 | 3316871 | BCGDan_3077 | BCGDan_3077 | 1248 | 3315689 | 3316999 | BCGDan_sapMKO_3078 | BCGDan_sapMKO_3078 | 1311 |
| 3382785 | 3383888 | trmU | Rv3024c | 1104 | 3337217 | 3338320 | trmU | BCG_3047c | 1104 | 3317034 | 3318137 | trmU | BCGDan_3078 | 1104 | 3317099 | 3318145 | trmU | BCGDan_sapMKO_3079 | 1047 |
| 3383885 | 3385066 | iscS | Rv3025c | 1182 | 3338317 | 3339498 | iscS | BCG_3048c | 1182 | 3318134 | 3319315 | iscS | BCGDan_3079 | 1182 | 3318199 | 3319380 | iscS | BCGDan_sapMKO_3080 | 1182 |
| 3385163 | 3386077 | Rv3026c | Rv3026c | 915 | 3339595 | 3340509 | BCG_3049c | BCG_3049c | 915 | 3319412 | 3320326 | BCGDan_3080 | BCGDan_3080 | 915 | 3319477 | 3320328 | BCGDan_sapMKO_3081 | BCGDan_sapMKO_3081 | 852 |
| 3386074 | 3386919 | Rv3027c | Rv3027c | 846 | 3340506 | 3341246 | BCG_3050c | BCG_3050c | 741 | 3320323 | 3321168 | BCGDan_3081 | BCGDan_3081 | 846 | 3320388 | 3321233 | BCGDan_sapMKO_3082 | BCGDan_sapMKO_3082 | 846 |
| 3387075 | 3388031 | fixB | Rv3028c | 957 | 3341507 | 3342463 | fixB | BCG_3051c | 957 | 3321324 | 3322280 | fixB | BCGDan_3082 | 957 | 3321389 | 3322345 | fixB | BCGDan_sapMKO_3083 | 957 |
| 3388070 | 3388870 | fixA | Rv3029c | 801 | 3342502 | 3343302 | fixA | BCG_3052c | 801 | 3322319 | 3323119 | fixA | BCGDan_3083 | 801 | 3322384 | 3323184 | fixA | BCGDan_sapMKO_3084 | 801 |
| 3389101 | 3389925 | Rv3030 | Rv3030 | 825 | 3343533 | 3344357 | BCG_3053 | BCG_3053 | 825 | 3323350 | 3324174 | BCGDan_3084 | BCGDan_3084 | 825 | 3323493 | 3324239 | BCGDan_sapMKO_3085 | BCGDan_sapMKO_3085 | 747 |
| 3389922 | 3391502 | Rv3031 | Rv3031 | 1581 | 3344354 | 3345934 | BCG_3054 | BCG_3054 | 1581 | 3324171 | 3325751 | BCGDan_3085 | BCGDan_3085 | 1581 | 3324257 | 3325816 | BCGDan_sapMKO_3086 | BCGDan_sapMKO_3086 | 1560 |
| 3391534 | 3392778 | Rv3032 | Rv3032 | 1245 | 3345966 | 3347210 | BCG_3055 | BCG_3055 | 1245 | 3325783 | 3327027 | BCGDan_3086 | BCGDan_3086 | 1245 | 3325860 | 3327092 | BCGDan_sapMKO_3087 | BCGDan_sapMKO_3087 | 1233 |
| 3392812 | 3393201 | Rv3032A | Rv3032A | 390 | 3347244 | 3347633 | BCG_3056 | BCG_3056 | 390 | 3327061 | 3327450 | BCGDan_3087 | BCGDan_3087 | 390 | 3327126 | 3327515 | BCGDan_sapMKO_3088 | BCGDan_sapMKO_3088 | 390 |
| 3393380 | 3393928 | Rv3033 | Rv3033 | 549 | 3347812 | 3348360 | BCG_3057 | BCG_3057 | 549 | 3327629 | 3328177 | BCGDan_3088 | BCGDan_3088 | 549 | 3327694 | 3328242 | BCGDan_sapMKO_3089 | BCGDan_sapMKO_3089 | 549 |
| 3394019 | 3394921 | Rv3034c | Rv3034c | 903 | 3348451 | 3349353 | BCG_3058c | BCG_3058c | 903 | 3328268 | 3329170 | BCGDan_3089 | BCGDan_3089 | 903 | 3328333 | 3329235 | BCGDan_sapMKO_3090 | BCGDan_sapMKO_3090 | 903 |
| 3395379 | 3396461 | Rv3035 | Rv3035 | 1083 | 3349811 | 3350893 | BCG_3059 | BCG_3059 | 1083 | 3329628 | 3330710 | BCGDan_3090 | BCGDan_3090 | 1083 | 3329768 | 3330775 | BCGDan_sapMKO_3091 | BCGDan_sapMKO_3091 | 1008 |
| 3396458 | 3397141 | TB22.2 | Rv3036c | 684 | 3350890 | 3351573 | TB22.2 | BCG_3060c | 684 | 3330707 | 3331390 | TB22.2 | BCGDan_3091 | 684 | 3330772 | 3331455 | TB22.2 | BCGDan_sapMKO_3092 | 684 |
| 3397214 | 3398290 | Rv3037c | Rv3037c | 1077 | 3351646 | 3352722 | BCG_3061c | BCG_3061c | 1077 | 3331463 | 3332539 | BCGDan_3092 | BCGDan_3092 | 1077 | 3331528 | 3332604 | BCGDan_sapMKO_3093 | BCGDan_sapMKO_3093 | 1077 |
| 3398425 | 3399408 | Rv3038c | Rv3038c | 984 | 3352857 | 3353840 | BCG_3062c | BCG_3062c | 984 | 3332674 | 3333657 | BCGDan_3093 | BCGDan_3093 | 984 | 3332739 | 3333752 | BCGDan_sapMKO_3094 | BCGDan_sapMKO_3094 | 1014 |
| 3399419 | 3400183 | echA17 | Rv3039c | 765 | 3353851 | 3354615 | echA17 | BCG_3063c | 765 | 3333668 | 3334432 | echA17 | BCGDan_3094 | 765 | 3333733 | 3334518 | echA17 | BCGDan_sapMKO_3095 | 786 |
| 3400192 | 3401058 | Rv3040c | Rv3040c | 867 | 3354624 | 3355490 | BCG_3064c | BCG_3064c | 867 | 3334441 | 3335307 | BCGDan_3095 | BCGDan_3095 | 867 | 3334506 | 3335315 | BCGDan_sapMKO_3096 | BCGDan_sapMKO_3096 | 810 |
| 3401055 | 3401918 | Rv3041c | Rv3041c | 864 | 3355487 | 3356350 | BCG_3065c | BCG_3065c | 864 | 3335304 | 3336167 | BCGDan_3096 | BCGDan_3096 | 864 | 3335369 | 3336232 | BCGDan_sapMKO_3097 | BCGDan_sapMKO_3097 | 864 |
| 3401933 | 3403162 | serB2 | Rv3042c | 1230 | 3356365 | 3357594 | serB2 | BCG_3066c | 1230 | 3336182 | 3337411 | serB2 | BCGDan_3097 | 1230 | 3336247 | 3337476 | serB2 | BCGDan_sapMKO_3098 | 1230 |
| 3403200 | 3404921 | ctaD | Rv3043c | 1722 | 3357632 | 3359353 | ctaD | BCG_3067c | 1722 | 3337449 | 3339170 | ctaD | BCGDan_3098 | 1722 | 3337514 | 3339235 | ctaD | BCGDan_sapMKO_3099 | 1722 |
| 3405136 | 3406215 | fecB | Rv3044 | 1080 | 3359568 | 3360647 | fecB | BCG_3068 | 1080 | 3339385 | 3340464 | fecB | BCGDan_3099 | 1080 | 3339546 | 3340529 | fecB | BCGDan_sapMKO_3100 | 984 |
| 3406285 | 3407325 | adhC | Rv3045 | 1041 | 3360717 | 3361757 | adhC | BCG_3069 | 1041 | 3340534 | 3341574 | adhC | BCGDan_3100 | 1041 | 3340599 | 3341639 | adhC | BCGDan_sapMKO_3101 | 1041 |
| 3407314 | 3407688 | Rv3046c | Rv3046c | 375 | 3361746 | 3362120 | BCG_3070c | BCG_3070c | 375 | 3341563 | 3341937 | BCGDan_3101 | BCGDan_3101 | 375 | 3341628 | 3342002 | BCGDan_sapMKO_3102 | BCGDan_sapMKO_3102 | 375 |
| 3408022 | 3408306 | Rv3047c | Rv3047c | 285 | 3362454 | 3362738 | BCG_3071c | BCG_3071c | 285 | 3342271 | 3342555 | BCGDan_3102 | BCGDan_3102 | 285 | 3342336 | 3342620 | BCGDan_sapMKO_3103 | BCGDan_sapMKO_3103 | 285 |
| 3408404 | 3409378 | nrdF2 | Rv3048c | 975 | 3362836 | 3363810 | nrdF2 | BCG_3072c | 975 | 3342653 | 3343627 | nrdF2 | BCGDan_3103 | 975 | 3342718 | 3343692 | nrdF2 | BCGDan_sapMKO_3104 | 975 |
| 3409509 | 3411083 | Rv3049c | Rv3049c | 1575 | 3363941 | 3365515 | BCG_3073c | BCG_3073c | 1575 | 3343758 | 3345332 | BCGDan_3104 | BCGDan_3104 | 1575 | 3343823 | 3345397 | BCGDan_sapMKO_3105 | BCGDan_sapMKO_3105 | 1575 |
| 3411217 | 3411957 | Rv3050c | Rv3050c | 741 | 3365649 | 3366389 | BCG_3074c | BCG_3074c | 741 | 3345466 | 3346206 | BCGDan_3105 | BCGDan_3105 | 741 | 3345531 | 3346229 | BCGDan_sapMKO_3106 | BCGDan_sapMKO_3106 | 699 |
| 3412085 | 3414166 | nrdE | Rv3051c | 2082 | 3366517 | 3368598 | nrdE | BCG_3075c | 2082 | 3346334 | 3348415 | nrdE | BCGDan_3106 | 2082 | 3346399 | 3348480 | nrdE | BCGDan_sapMKO_3107 | 2082 |
| 3414232 | 3414684 | nrdI | Rv3052c | 453 | 3368664 | 3369116 | nrdI | BCG_3076c | 453 | 3348481 | 3348933 | nrdI | BCGDan_3107 | 453 | 3348546 | 3348998 | nrdI | BCGDan_sapMKO_3108 | 453 |
| 3414719 | 3414958 | nrdH | Rv3053c | 240 | 3369151 | 3369390 | nrdH | BCG_3077c | 240 | 3348968 | 3349207 | nrdH | BCGDan_3108 | 240 | 3349033 | 3349272 | nrdH | BCGDan_sapMKO_3109 | 240 |
| NA | NA | NA | NA | NA | 3369368 | 3369736 | BCG_3078c | BCG_3078c | 369 | 3349185 | 3349553 | BCGDan_3109 | BCGDan_3109 | 369 | 3349250 | 3349618 | BCGDan_sapMKO_3110 | BCGDan_sapMKO_3110 | 369 |
| 3415435 | 3415989 | Rv3054c | Rv3054c | 555 | 3369853 | 3370407 | BCG_3079c | BCG_3079c | 555 | 3349670 | 3350224 | BCGDan_3110 | BCGDan_3110 | 555 | 3349735 | 3350289 | BCGDan_sapMKO_3111 | BCGDan_sapMKO_3111 | 555 |
| 3416081 | 3416695 | Rv3055 | Rv3055 | 615 | 3370499 | 3371113 | BCG_3080 | BCG_3080 | 615 | 3350316 | 3350930 | BCGDan_3111 | BCGDan_3111 | 615 | 3350381 | 3350995 | BCGDan_sapMKO_3112 | BCGDan_sapMKO_3112 | 615 |
| 3416705 | 3417745 | dinP | Rv3056 | 1041 | 3371123 | 3372163 | dinP | BCG_3081 | 1041 | 3350940 | 3351980 | dinP | BCGDan_3112 | 1041 | 3351005 | 3352045 | dinP | BCGDan_sapMKO_3113 | 1041 |
| 3417799 | 3418662 | Rv3057c | Rv3057c | 864 | 3372217 | 3373080 | BCG_3082c | BCG_3082c | 864 | 3352034 | 3352897 | BCGDan_3113 | BCGDan_3113 | 864 | 3352099 | 3352962 | BCGDan_sapMKO_3114 | BCGDan_sapMKO_3114 | 864 |
| 3418726 | 3419376 | Rv3058c | Rv3058c | 651 | 3373144 | 3373794 | BCG_3083c | BCG_3083c | 651 | 3352961 | 3353611 | BCGDan_3114 | BCGDan_3114 | 651 | 3353026 | 3353706 | BCGDan_sapMKO_3115 | BCGDan_sapMKO_3115 | 681 |
| 3419492 | 3420970 | cyp136 | Rv3059 | 1479 | 3373910 | 3375388 | cyp136 | BCG_3084 | 1479 | 3353727 | 3355205 | cyp136 | BCGDan_3115 | 1479 | 3353792 | 3355270 | cyp136 | BCGDan_sapMKO_3116 | 1479 |
| 3421741 | 3423213 | Rv3060c | Rv3060c | 1473 | 3376159 | 3377631 | BCG_3085c | BCG_3085c | 1473 | 3355976 | 3357448 | BCGDan_3116 | BCGDan_3116 | 1473 | 3356041 | 3357513 | BCGDan_sapMKO_3117 | BCGDan_sapMKO_3117 | 1473 |
| 3423262 | 3425427 | fadE22 | Rv3061c | 2166 | 3377748 | 3379913 | fadE22 | BCG_3086c | 2166 | 3357565 | 3359730 | fadE22 | BCGDan_3117 | 2166 | 3357630 | 3359795 | fadE22 | BCGDan_sapMKO_3118 | 2166 |
| 3425584 | 3427107 | ligB | Rv3062 | 1524 | 3380070 | 3381593 | ligB | BCG_3087 | 1524 | 3359887 | 3361410 | ligB | BCGDan_3118 | 1524 | 3359952 | 3361475 | ligB | BCGDan_sapMKO_3119 | 1524 |
| 3427243 | 3429519 | cstA | Rv3063 | 2277 | 3381729 | 3384005 | cstA | BCG_3088 | 2277 | 3361546 | 3363822 | cstA | BCGDan_3119 | 2277 | 3361605 | 3363887 | cstA | BCGDan_sapMKO_3120 | 2283 |
| 3429825 | 3430250 | Rv3064c | Rv3064c | 426 | 3384311 | 3384736 | BCG_3089c | BCG_3089c | 426 | 3364128 | 3364553 | BCGDan_3120 | BCGDan_3120 | 426 | 3364193 | 3364618 | BCGDan_sapMKO_3121 | BCGDan_sapMKO_3121 | 426 |
| 3430387 | 3430710 | mmr | Rv3065 | 324 | 3384873 | 3385196 | mmr | BCG_3090 | 324 | 3364690 | 3365013 | mmr | BCGDan_3121 | 324 | 3364755 | 3365078 | mmr | BCGDan_sapMKO_3122 | 324 |
| 3430707 | 3431315 | Rv3066 | Rv3066 | 609 | 3385193 | 3385801 | BCG_3091 | BCG_3091 | 609 | 3365010 | 3365618 | BCGDan_3122 | BCGDan_3122 | 609 | 3365075 | 3365683 | BCGDan_sapMKO_3123 | BCGDan_sapMKO_3123 | 609 |
| 3431428 | 3431838 | Rv3067 | Rv3067 | 411 | 3385914 | 3386324 | BCG_3092 | BCG_3092 | 411 | 3365731 | 3366141 | BCGDan_3123 | BCGDan_3123 | 411 | 3365796 | 3366206 | BCGDan_sapMKO_3124 | BCGDan_sapMKO_3124 | 411 |
| 3431979 | 3433622 | pgmA | Rv3068c | 1644 | 3386465 | 3388108 | pgmA | BCG_3093c | 1644 | 3366282 | 3367925 | pgmA | BCGDan_3125 | 1644 | 3366347 | 3367990 | pgmA | BCGDan_sapMKO_3126 | 1644 |
| 3433692 | 3434090 | Rv3069 | Rv3069 | 399 | 3388178 | 3388576 | BCG_3094 | BCG_3094 | 399 | 3367995 | 3368393 | BCGDan_3126 | BCGDan_3126 | 399 | 3368060 | 3368458 | BCGDan_sapMKO_3127 | BCGDan_sapMKO_3127 | 399 |
| 3434087 | 3434467 | Rv3070 | Rv3070 | 381 | 3388573 | 3388953 | BCG_3095 | BCG_3095 | 381 | 3368390 | 3368770 | BCGDan_3127 | BCGDan_3127 | 381 | 3368455 | 3368835 | BCGDan_sapMKO_3128 | BCGDan_sapMKO_3128 | 381 |
| 3434464 | 3435573 | Rv3071 | Rv3071 | 1110 | 3388950 | 3390059 | BCG_3096 | BCG_3096 | 1110 | 3368767 | 3369876 | BCGDan_3128 | BCGDan_3128 | 1110 | 3368832 | 3369941 | BCGDan_sapMKO_3129 | BCGDan_sapMKO_3129 | 1110 |
| 3435798 | 3436322 | Rv3072c | Rv3072c | 525 | 3390284 | 3390808 | BCG_3097c | BCG_3097c | 525 | 3370101 | 3370625 | BCGDan_3129 | BCGDan_3129 | 525 | 3370166 | 3370663 | BCGDan_sapMKO_3130 | BCGDan_sapMKO_3130 | 498 |
| 3436329 | 3436685 | Rv3073c | Rv3073c | 357 | 3390815 | 3391171 | BCG_3098c | BCG_3098c | 357 | 3370632 | 3370988 | BCGDan_3130 | BCGDan_3130 | 357 | 3370697 | 3371053 | BCGDan_sapMKO_3131 | BCGDan_sapMKO_3131 | 357 |
| 3436779 | 3438053 | Rv3074 | Rv3074 | 1275 | 3391265 | 3392539 | BCG_3099 | BCG_3099 | 1275 | 3371082 | 3372356 | BCGDan_3131 | BCGDan_3131 | 1275 | 3371147 | 3372421 | BCGDan_sapMKO_3132 | BCGDan_sapMKO_3132 | 1275 |
| 3438050 | 3438973 | Rv3075c | Rv3075c | 924 | 3392536 | 3393459 | BCG_3100c | BCG_3100c | 924 | 3372353 | 3373276 | BCGDan_3132 | BCGDan_3132 | 924 | 3372418 | 3373332 | BCGDan_sapMKO_3133 | BCGDan_sapMKO_3133 | 915 |

| | | | | | | | | | | | | | | | | | | | |
|---|---|---|---|---|---|---|---|---|---|---|---|---|---|---|---|---|---|---|---|
| 3439072 | 3439548 | Rv3076 | Rv3076 | 477 | 3393558 | 3394034 | BCG_3101 | BCG_3101 | 477 | 3373375 | 3373851 | BCGDan_3133 | BCGDan_3133 | 477 | 3373440 | 3373916 | BCGDan_sapMKO_3134 | BCGDan_sapMKO_3134 | 477 |
| 3439541 | 3441352 | Rv3077 | Rv3077 | 1812 | 3394027 | 3395838 | BCG_3102 | BCG_3102 | 1812 | 3373844 | 3375655 | BCGDan_3134 | BCGDan_3134 | 1812 | 3373909 | 3375720 | BCGDan_sapMKO_3135 | BCGDan_sapMKO_3135 | 1812 |
| 3441353 | 3441754 | hab | Rv3078 | 402 | 3395839 | 3396240 | hab | BCG_3103 | 402 | 3375656 | 3376057 | hab | BCGDan_3135 | 402 | 3375721 | 3376122 | hab | BCGDan_sapMKO_3136 | 402 |
| 3441770 | 3442597 | Rv3079c | Rv3079c | 828 | 3396256 | 3397083 | BCG_3104c | BCG_3104c | 828 | 3376073 | 3376900 | BCGDan_3136 | BCGDan_3136 | 828 | 3376138 | 3376977 | BCGDan_sapMKO_3137 | BCGDan_sapMKO_3137 | 840 |
| 3442656 | 3445988 | pknK | Rv3080c | 3333 | 3397142 | 3400474 | pknK | BCG_3105c | 3333 | 3376959 | 3380291 | pknK | BCGDan_3137 | 3333 | 3377024 | 3380356 | pknK | BCGDan_sapMKO_3138 | 3333 |
| 3446040 | 3447278 | Rv3081 | Rv3081 | 1239 | 3400526 | 3401764 | BCG_3106 | BCG_3106 | 1239 | 3380343 | 3381581 | BCGDan_3138 | BCGDan_3138 | 1239 | 3380408 | 3381646 | BCGDan_sapMKO_3139 | BCGDan_sapMKO_3139 | 1239 |
| 3447404 | 3448426 | virS | Rv3082c | 1023 | 3401890 | 3402912 | virS | BCG_3107c | 1023 | 3381707 | 3382729 | virS | BCGDan_3139 | 1023 | 3381772 | 3382794 | virS | BCGDan_sapMKO_3140 | 1023 |
| 3448504 | 3449991 | Rv3083 | Rv3083 | 1488 | 3402990 | 3404477 | BCG_3108 | BCG_3108 | 1488 | 3382807 | 3384294 | BCGDan_3140 | BCGDan_3140 | 1488 | 3382872 | 3384359 | BCGDan_sapMKO_3141 | BCGDan_sapMKO_3141 | 1488 |
| 3449997 | 3450923 | lipR | Rv3084 | 927 | 3404483 | 3405409 | lipR | BCG_3109 | 927 | 3384300 | 3385226 | lipR | BCGDan_3141 | 927 | 3384365 | 3385291 | lipR | BCGDan_sapMKO_3142 | 927 |
| 3450920 | 3451750 | Rv3085 | Rv3085 | 831 | 3405406 | 3406236 | BCG_3110 | BCG_3110 | 831 | 3385223 | 3386053 | BCGDan_3142 | BCGDan_3142 | 831 | 3385288 | 3386118 | BCGDan_sapMKO_3143 | BCGDan_sapMKO_3143 | 831 |
| 3451781 | 3452887 | adhD | Rv3086 | 1107 | 3406267 | 3407373 | adhD | BCG_3111 | 1107 | 3386084 | 3387190 | adhD | BCGDan_3143 | 1107 | 3386149 | 3387255 | adhD | BCGDan_sapMKO_3144 | 1107 |
| 3452925 | 3454343 | Rv3087 | Rv3087 | 1419 | 3407411 | 3408829 | BCG_3112 | BCG_3112 | 1419 | 3387228 | 3388646 | BCGDan_3144 | BCGDan_3144 | 1419 | 3387290 | 3388711 | BCGDan_sapMKO_3145 | BCGDan_sapMKO_3145 | 1422 |
| 3454340 | 3455764 | tgs4 | Rv3088 | 1425 | 3408826 | 3410250 | BCG_3113 | BCG_3113 | 1425 | 3388643 | 3390067 | BCGDan_3145 | BCGDan_3145 | 1425 | 3388708 | 3390132 | BCGDan_sapMKO_3146 | BCGDan_sapMKO_3146 | 1425 |
| 3455761 | 3457272 | fadD13 | Rv3089 | 1512 | 3410247 | 3411758 | fadD13 | BCG_3114 | 1512 | 3390064 | 3391575 | fadD13 | BCGDan_3146 | 1512 | 3390129 | 3391640 | fadD13 | BCGDan_sapMKO_3147 | 1512 |
| 3458211 | 3459098 | Rv3090 | Rv3090 | 888 | 3412697 | 3413584 | BCG_3115 | BCG_3115 | 888 | 3392514 | 3393401 | BCGDan_3147 | BCGDan_3147 | 888 | 3392579 | 3393466 | BCGDan_sapMKO_3148 | BCGDan_sapMKO_3148 | 888 |
| 3459116 | 3460807 | Rv3091 | Rv3091 | 1692 | 3413602 | 3415293 | BCG_3116 | BCG_3116 | 1692 | 3393419 | 3395110 | BCGDan_3148 | BCGDan_3148 | 1692 | 3393484 | 3395175 | BCGDan_sapMKO_3149 | BCGDan_sapMKO_3149 | 1692 |
| 3460814 | 3461734 | Rv3092c | Rv3092c | 921 | 3415300 | 3416220 | BCG_3117c | BCG_3117c | 921 | 3395117 | 3396037 | BCGDan_3149 | BCGDan_3149 | 921 | 3395182 | 3396102 | BCGDan_sapMKO_3150 | BCGDan_sapMKO_3150 | 921 |
| 3461760 | 3462764 | Rv3093c | Rv3093c | 1005 | 3416246 | 3417250 | BCG_3118c | BCG_3118c | 1005 | 3396063 | 3397067 | BCGDan_3150 | BCGDan_3150 | 1005 | 3396128 | 3397132 | BCGDan_sapMKO_3151 | BCGDan_sapMKO_3151 | 1005 |
| 3462761 | 3463891 | Rv3094c | Rv3094c | 1131 | 3417247 | 3418377 | BCG_3119c | BCG_3119c | 1131 | 3397064 | 3398194 | BCGDan_3151 | BCGDan_3151 | 1131 | 3397129 | 3398259 | BCGDan_sapMKO_3152 | BCGDan_sapMKO_3152 | 1131 |
| 3463973 | 3464449 | Rv3095 | Rv3095 | 477 | 3418459 | 3418935 | BCG_3120 | BCG_3120 | 477 | 3398276 | 3398752 | BCGDan_3152 | BCGDan_3152 | 477 | 3398341 | 3398817 | BCGDan_sapMKO_3153 | BCGDan_sapMKO_3153 | 477 |
| 3464547 | 3465686 | Rv3096 | Rv3096 | 1140 | 3419033 | 3420172 | BCG_3121 | BCG_3121 | 1140 | 3398850 | 3399989 | BCGDan_3153 | BCGDan_3153 | 1140 | 3398915 | 3400054 | BCGDan_sapMKO_3154 | BCGDan_sapMKO_3154 | 1140 |
| 3465778 | 3467091 | lipY | Rv3097c | 1314 | 3420264 | 3421577 | PE_PGRS63 | BCG_3122c | 1314 | 3400081 | 3401394 | PE_PGRS63 | BCGDan_3154 | 1314 | 3400146 | 3401459 | PE_PGRS63 | BCGDan_sapMKO_3155 | 1314 |
| 3467210 | 3467662 | Rv3098c | Rv3098c | 453 | 3421696 | 3422148 | BCG_3123c | BCG_3123c | 453 | 3401513 | 3401965 | BCGDan_3155 | BCGDan_3155 | 453 | 3401578 | 3402030 | BCGDan_sapMKO_3156 | BCGDan_sapMKO_3156 | 453 |
| 3468413 | 3469264 | Rv3099c | Rv3099c | 852 | 3422899 | 3423750 | BCG_3124c | BCG_3124c | 852 | 3402716 | 3403567 | BCGDan_3157 | BCGDan_3157 | 852 | 3402781 | 3403632 | BCGDan_sapMKO_3158 | BCGDan_sapMKO_3158 | 852 |
| 3469301 | 3469783 | smpB | Rv3100c | 483 | 3423787 | 3424269 | smpB | BCG_3125c | 483 | 3403604 | 3404086 | smpB | BCGDan_3158 | 483 | 3403669 | 3404151 | smpB | BCGDan_sapMKO_3159 | 483 |
| 3469786 | 3470679 | ftsX | Rv3101c | 894 | 3424272 | 3425165 | ftsX | BCG_3126c | 894 | 3404089 | 3404982 | ftsX | BCGDan_3159 | 894 | 3404154 | 3405047 | ftsX | BCGDan_sapMKO_3160 | 894 |
| 3470680 | 3471369 | ftsE | Rv3102c | 690 | 3425166 | 3425855 | ftsE | BCG_3127c | 690 | 3404983 | 3405672 | ftsE | BCGDan_3160 | 690 | 3405048 | 3405740 | ftsE | BCGDan_sapMKO_3161 | 693 |
| 3471413 | 3471850 | Rv3103c | Rv3103c | 438 | 3425899 | 3426336 | BCG_3128c | BCG_3128c | 438 | 3405716 | 3406153 | BCGDan_3161 | BCGDan_3161 | 438 | 3405781 | 3406218 | BCGDan_sapMKO_3162 | BCGDan_sapMKO_3162 | 438 |
| 3471852 | 3472778 | Rv3104c | Rv3104c | 927 | 3426338 | 3427264 | BCG_3129c | BCG_3129c | 927 | 3406155 | 3407081 | BCGDan_3162 | BCGDan_3162 | 927 | 3406220 | 3407146 | BCGDan_sapMKO_3163 | BCGDan_sapMKO_3163 | 927 |
| 3472768 | 3473904 | prfB | Rv3105c | 1137 | 3427254 | 3428390 | prfB | BCG_3130c | 1137 | 3407071 | 3408207 | prfB | BCGDan_3163 | 1137 | 3407136 | 3408272 | prfB | BCGDan_sapMKO_3164 | 1137 |
| 3474007 | 3475377 | fprA | Rv3106 | 1371 | 3428494 | 3429864 | fprA | BCG_3131 | 1371 | 3408311 | 3409681 | fprA | BCGDan_3164 | 1371 | 3408376 | 3409746 | fprA | BCGDan_sapMKO_3165 | 1371 |
| 3475378 | 3476961 | agpS | Rv3107c | 1584 | 3429865 | 3431448 | agpS | BCG_3132c | 1584 | 3409682 | 3411265 | agpS | BCGDan_3165 | 1584 | 3409747 | 3411330 | agpS | BCGDan_sapMKO_3166 | 1584 |
| 3477060 | 3477500 | Rv3108 | Rv3108 | 441 | 3431547 | 3431987 | BCG_3133 | BCG_3133 | 441 | 3411364 | 3411804 | BCGDan_3166 | BCGDan_3166 | 441 | 3411429 | 3411869 | BCGDan_sapMKO_3167 | BCGDan_sapMKO_3167 | 441 |
| 3477649 | 3478728 | moaA1 | Rv3109 | 1080 | 3432136 | 3433215 | moaA1 | BCG_3134 | 1080 | 3411953 | 3413032 | moaA1 | BCGDan_3167 | 1080 | 3412150 | 3413097 | moaA1 | BCGDan_sapMKO_3168 | 948 |
| 3478779 | 3479174 | moaB1 | Rv3110 | 396 | 3433266 | 3433661 | moaB1 | BCG_3135 | 396 | 3413083 | 3413478 | moaB1 | BCGDan_3168 | 396 | 3413148 | 3413543 | moaB1 | BCGDan_sapMKO_3169 | 396 |
| 3479171 | 3479683 | moaC1 | Rv3111 | 513 | 3433658 | 3434170 | moaC1 | BCG_3136 | 513 | 3413475 | 3413987 | moaC1 | BCGDan_3169 | 513 | 3413594 | 3414052 | moaC1 | BCGDan_sapMKO_3170 | 459 |
| 3479700 | 3479951 | moaD1 | Rv3112 | 252 | 3434187 | 3434438 | moaD1 | BCG_3137 | 252 | 3414004 | 3414255 | moaD1 | BCGDan_3170 | 252 | 3414069 | 3414320 | moaD1 | BCGDan_sapMKO_3171 | 252 |
| 3480074 | 3480742 | Rv3113 | Rv3113 | 669 | 3434560 | 3435228 | BCG_3138 | BCG_3138 | 669 | 3414377 | 3415045 | BCGDan_3171 | BCGDan_3171 | 669 | 3414442 | 3415110 | BCGDan_sapMKO_3172 | BCGDan_sapMKO_3172 | 669 |
| 3480759 | 3481289 | Rv3114 | Rv3114 | 531 | 3435245 | 3435775 | BCG_3139 | BCG_3139 | 531 | 3415062 | 3415592 | BCGDan_3172 | BCGDan_3172 | 531 | 3415094 | 3415657 | BCGDan_sapMKO_3173 | BCGDan_sapMKO_3173 | 564 |
| 3481451 | 3482698 | Rv3115 | Rv3115 | 1248 | 3435937 | 3437184 | BCG_3140 | BCG_3140 | 1248 | 3415754 | 3417001 | BCGDan_3173 | BCGDan_3173 | 1248 | 3415756 | 3417066 | BCGDan_sapMKO_3174 | BCGDan_sapMKO_3174 | 1311 |
| 3482776 | 3483945 | moeB2 | Rv3116 | 1170 | 3437262 | 3438431 | moeB2 | BCG_3141 | 1170 | 3417079 | 3418248 | moeB2 | BCGDan_3174 | 1170 | 3417144 | 3418313 | moeB2 | BCGDan_sapMKO_3175 | 1170 |
| 3483974 | 3484807 | cysA3 | Rv3117 | 834 | 3438460 | 3439422 | cysA3 | BCG_3142 | 963 | 3418277 | 3419239 | cysA3 | BCGDan_3175 | 963 | 3418342 | 3419304 | cysA3 | BCGDan_sapMKO_3176 | 963 |
| 3488089 | 3488559 | Rv3122 | Rv3122 | 471 | 3439800 | 3440270 | BCG_3143 | BCG_3143 | 471 | 3419617 | 3420087 | BCGDan_3176 | BCGDan_3176 | 471 | 3419727 | 3420152 | BCGDan_sapMKO_3177 | BCGDan_sapMKO_3177 | 426 |
| 3488569 | 3489063 | Rv3123 | Rv3123 | 495 | 3440280 | 3440774 | BCG_3144 | BCG_3144 | 495 | 3420097 | 3420591 | BCGDan_3177 | BCGDan_3177 | 495 | 3420162 | 3420656 | BCGDan_sapMKO_3178 | BCGDan_sapMKO_3178 | 495 |
| 3489506 | 3490375 | moaR1 | Rv3124 | 870 | 3441217 | 3442086 | BCG_3145 | BCG_3145 | 870 | 3421034 | 3421903 | BCGDan_3178 | BCGDan_3178 | 870 | 3421099 | 3421968 | BCGDan_sapMKO_3179 | BCGDan_sapMKO_3179 | 870 |
| NA | NA | NA | NA | NA | 3442187 | 3442501 | PPE49b | BCG_3146c | 315 | 3422004 | 3422318 | PPE49b | BCGDan_3179 | 315 | 3422069 | 3422383 | PPE49b | BCGDan_sapMKO_3180 | 315 |
| 3490476 | 3491651 | PPE49 | Rv3125c | 1176 | 3442513 | 3443361 | PPE49a | BCG_3147c | 849 | 3422330 | 3423178 | PPE49a | BCGDan_3180 | 849 | 3422395 | 3423243 | PPE49a | BCGDan_sapMKO_3181 | 849 |
| 3491808 | 3492122 | Rv3126c | Rv3126c | 315 | 3443518 | 3443832 | BCG_3148c | BCG_3148c | 315 | 3423335 | 3423649 | BCGDan_3181 | BCGDan_3181 | 315 | 3423400 | 3423672 | BCGDan_sapMKO_3182 | BCGDan_sapMKO_3182 | 273 |
| 3492147 | 3493181 | Rv3127 | Rv3127 | 1035 | 3443857 | 3444891 | BCG_3149 | BCG_3149 | 1035 | 3423674 | 3424708 | BCGDan_3182 | BCGDan_3182 | 1035 | 3423694 | 3424773 | BCGDan_sapMKO_3183 | BCGDan_sapMKO_3183 | 1080 |
| 3493168 | 3494181 | Rv3128c | Rv3128c | 1014 | 3444878 | 3445228 | BCG_3150c | BCG_3150c | 351 | 3424695 | 3425045 | BCGDan_3183 | BCGDan_3183 | 351 | 3424760 | 3425110 | BCGDan_sapMKO_3184 | BCGDan_sapMKO_3184 | 351 |
| NA | NA | NA | NA | NA | 3445310 | 3445891 | BCG_3151c | BCG_3151c | 582 | 3425127 | 3425708 | BCGDan_3184 | BCGDan_3184 | 582 | 3425192 | 3425725 | BCGDan_sapMKO_3185 | BCGDan_sapMKO_3185 | 534 |
| 3494660 | 3494992 | Rv3129 | Rv3129 | 333 | 3446370 | 3446702 | BCG_3152 | BCG_3152 | 333 | 3426187 | 3426519 | BCGDan_3185 | BCGDan_3185 | 333 | 3426252 | 3426584 | BCGDan_sapMKO_3186 | BCGDan_sapMKO_3186 | 333 |
| 3494975 | 3496366 | tgs1 | Rv3130c | 1392 | 3446685 | 3448076 | BCG_3153c | BCG_3153c | 1392 | 3426502 | 3427893 | BCGDan_3186 | BCGDan_3186 | 1392 | 3426567 | 3427958 | BCGDan_sapMKO_3187 | BCGDan_sapMKO_3187 | 1392 |
| 3496551 | 3497549 | Rv3131 | Rv3131 | 999 | 3448261 | 3449259 | BCG_3154 | BCG_3154 | 999 | 3428078 | 3429076 | BCGDan_3187 | BCGDan_3187 | 999 | 3428143 | 3429141 | BCGDan_sapMKO_3188 | BCGDan_sapMKO_3188 | 999 |
| 3497529 | 3499265 | devS | Rv3132c | 1737 | 3449239 | 3450975 | devS | BCG_3155c | 1737 | 3429056 | 3430792 | devS | BCGDan_3188 | 1737 | 3429121 | 3430857 | devS | BCGDan_sapMKO_3189 | 1737 |
| 3499262 | 3499915 | devR | Rv3133c | 654 | 3450972 | 3451625 | devR | BCG_3156c | 654 | 3430789 | 3431442 | devR | BCGDan_3189 | 654 | 3430854 | 3431507 | devR | BCGDan_sapMKO_3190 | 654 |
| 3499943 | 3500749 | Rv3134c | Rv3134c | 807 | 3451653 | 3452459 | BCG_3157c | BCG_3157c | 807 | 3431470 | 3432276 | BCGDan_3190 | BCGDan_3190 | 807 | 3431535 | 3432314 | BCGDan_sapMKO_3191 | BCGDan_sapMKO_3191 | 780 |
| 3501334 | 3501732 | PPE50 | Rv3135 | 399 | 3453044 | 3454189 | PPE50 | BCG_3158 | 1146 | 3432861 | 3434006 | PPE50 | BCGDan_3191 | 1146 | 3432926 | 3434071 | PPE50 | BCGDan_sapMKO_3192 | 1146 |
| 3501794 | 3502936 | PPE51 | Rv3136 | 1143 | 3454841 | 3455983 | PPE51 | BCG_3159 | 1143 | 3434658 | 3435800 | PPE51 | BCGDan_3192 | 1143 | 3434765 | 3435865 | PPE51 | BCGDan_sapMKO_3193 | 1101 |
| 3503393 | 3504175 | Rv3137 | Rv3137 | 783 | 3456440 | 3457222 | BCG_3160 | BCG_3160 | 783 | 3436257 | 3437039 | BCGDan_3193 | BCGDan_3193 | 783 | 3436340 | 3437104 | BCGDan_sapMKO_3194 | BCGDan_sapMKO_3194 | 765 |
| 3504195 | 3505283 | pflA | Rv3138 | 1089 | 3457242 | 3458330 | pflA | BCG_3161 | 1089 | 3437059 | 3438147 | pflA | BCGDan_3194 | 1089 | 3437124 | 3438212 | pflA | BCGDan_sapMKO_3195 | 1089 |
| 3505363 | 3506769 | fadE24 | Rv3139 | 1407 | 3458410 | 3459816 | fadE24 | BCG_3162 | 1407 | 3438227 | 3439633 | fadE24 | BCGDan_3195 | 1407 | 3438292 | 3439698 | fadE24 | BCGDan_sapMKO_3196 | 1407 |
| 3506790 | 3507995 | fadE23 | Rv3140 | 1206 | 3459837 | 3461042 | fadE23 | BCG_3163 | 1206 | 3439654 | 3440859 | fadE23 | BCGDan_3196 | 1206 | 3439719 | 3440924 | fadE23 | BCGDan_sapMKO_3197 | 1206 |
| 3508095 | 3509066 | fadB4 | Rv3141 | 972 | 3461142 | 3462113 | fadB4 | BCG_3164 | 972 | 3440959 | 3441930 | fadB4 | BCGDan_3197 | 972 | 3441024 | 3441995 | fadB4 | BCGDan_sapMKO_3198 | 972 |
| 3509118 | 3509546 | Rv3142c | Rv3142c | 429 | 3462165 | 3462593 | BCG_3165c | BCG_3165c | 429 | 3441982 | 3442410 | BCGDan_3198 | BCGDan_3198 | 429 | 3442047 | 3442475 | BCGDan_sapMKO_3199 | BCGDan_sapMKO_3199 | 429 |
| 3509654 | 3510055 | Rv3143 | Rv3143 | 402 | 3462701 | 3463102 | BCG_3166 | BCG_3166 | 402 | 3442518 | 3442919 | BCGDan_3199 | BCGDan_3199 | 402 | 3442583 | 3442984 | BCGDan_sapMKO_3200 | BCGDan_sapMKO_3200 | 402 |
| 3510088 | 3511317 | PPE52 | Rv3144c | 1230 | 3463135 | 3464364 | PPE52 | BCG_3167c | 1230 | 3442952 | 3444181 | PPE52 | BCGDan_3200 | 1230 | 3443017 | 3444234 | PPE52 | BCGDan_sapMKO_3201 | 1218 |
| 3511682 | 3512068 | nuoA | Rv3145 | 387 | 3464729 | 3465115 | nuoA | BCG_3168 | 387 | 3444546 | 3444932 | nuoA | BCGDan_3201 | 387 | 3444611 | 3444997 | nuoA | BCGDan_sapMKO_3202 | 387 |
| 3512077 | 3512631 | nuoB | Rv3146 | 555 | 3465124 | 3465678 | nuoB | BCG_3169 | 555 | 3444941 | 3445495 | nuoB | BCGDan_3202 | 555 | 3445006 | 3445560 | nuoB | BCGDan_sapMKO_3203 | 555 |
| 3512628 | 3513338 | nuoC | Rv3147 | 711 | 3465675 | 3466385 | nuoC | BCG_3170 | 711 | 3445492 | 3446202 | nuoC | BCGDan_3203 | 711 | 3445557 | 3446267 | nuoC | BCGDan_sapMKO_3204 | 711 |
| 3513338 | 3514660 | nuoD | Rv3148 | 1323 | 3466385 | 3467707 | nuoD | BCG_3171 | 1323 | 3446202 | 3447524 | nuoD | BCGDan_3204 | 1323 | 3446267 | 3447589 | nuoD | BCGDan_sapMKO_3205 | 1323 |
| 3514657 | 3515415 | nuoE | Rv3149 | 759 | 3467704 | 3468462 | nuoE | BCG_3172 | 759 | 3447521 | 3448279 | nuoE | BCGDan_3205 | 759 | 3447586 | 3448344 | nuoE | BCGDan_sapMKO_3206 | 759 |
| 3515412 | 3516749 | nuoF | Rv3150 | 1338 | 3468459 | 3469796 | nuoF | BCG_3173 | 1338 | 3448276 | 3449613 | nuoF | BCGDan_3206 | 1338 | 3448341 | 3449678 | nuoF | BCGDan_sapMKO_3207 | 1338 |
| 3516746 | 3519166 | nuoG | Rv3151 | 2421 | 3469793 | 3472213 | nuoG | BCG_3174 | 2421 | 3449610 | 3452030 | nuoG | BCGDan_3207 | 2421 | 3449675 | 3452095 | nuoG | BCGDan_sapMKO_3208 | 2421 |
| 3519282 | 3520514 | nuoH | Rv3152 | 1233 | 3472329 | 3473561 | nuoH | BCG_3175 | 1233 | 3452146 | 3453378 | nuoH | BCGDan_3208 | 1233 | 3452202 | 3453443 | nuoH | BCGDan_sapMKO_3209 | 1242 |
| 3520507 | 3521142 | nuoI | Rv3153 | 636 | 3473554 | 3474189 | nuoI | BCG_3176 | 636 | 3453371 | 3454006 | nuoI | BCGDan_3209 | 636 | 3453436 | 3454071 | nuoI | BCGDan_sapMKO_3210 | 636 |
| 3521139 | 3521927 | nuoJ | Rv3154 | 789 | 3474186 | 3474974 | nuoJ | BCG_3177 | 789 | 3454003 | 3454791 | nuoJ | BCGDan_3210 | 789 | 3454068 | 3454856 | nuoJ | BCGDan_sapMKO_3211 | 789 |
| 3521924 | 3522223 | nuoK | Rv3155 | 300 | 3474971 | 3475270 | nuoK | BCG_3178 | 300 | 3454788 | 3455087 | nuoK | BCGDan_3211 | 300 | 3454853 | 3455152 | nuoK | BCGDan_sapMKO_3212 | 300 |
| 3522234 | 3524135 | nuoL | Rv3156 | 1902 | 3475281 | 3477182 | nuoL | BCG_3179 | 1902 | 3455098 | 3456999 | nuoL | BCGDan_3212 | 1902 | 3455163 | 3457064 | nuoL | BCGDan_sapMKO_3213 | 1902 |
| 3524132 | 3525793 | nuoM | Rv3157 | 1662 | 3477179 | 3478840 | nuoM | BCG_3180 | 1662 | 3456993 | 3458657 | nuoM | BCGDan_3213 | 1665 | 3457058 | 3458722 | nuoM | BCGDan_sapMKO_3214 | 1665 |
| 3525790 | 3527385 | nuoN | Rv3158 | 1596 | 3478837 | 3480432 | nuoN | BCG_3181 | 1596 | 3458654 | 3460249 | nuoN | BCGDan_3214 | 1596 | 3458719 | 3460314 | nuoN | BCGDan_sapMKO_3215 | 1596 |
| 3527391 | 3529163 | PPE53 | Rv3159c | 1773 | 3480438 | 3482207 | PPE53 | BCG_3182c | 1770 | 3460255 | 3462024 | PPE53 | BCGDan_3215 | 1770 | 3460320 | 3462089 | PPE53 | BCGDan_sapMKO_3216 | 1770 |
| NA | NA | NA | NA | NA | 3482455 | 3484356 | PPE70 | BCG_3183c | 1902 | 3462272 | 3464173 | PPE70 | BCGDan_3216 | 1902 | 3462337 | 3464238 | PPE70 | BCGDan_sapMKO_3217 | 1902 |
| 3529338 | 3529979 | Rv3160c | Rv3160c | 642 | 3484531 | 3485172 | BCG_3184c | BCG_3184c | 642 | 3464348 | 3464989 | BCGDan_3217 | BCGDan_3217 | 642 | 3464413 | 3465054 | BCGDan_sapMKO_3218 | BCGDan_sapMKO_3218 | 642 |
| 3529990 | 3531138 | Rv3161c | Rv3161c | 1149 | 3485183 | 3486331 | BCG_3185c | BCG_3185c | 1149 | 3465000 | 3466148 | BCGDan_3218 | BCGDan_3218 | 1149 | 3465065 | 3466222 | BCGDan_sapMKO_3219 | BCGDan_sapMKO_3219 | 1158 |
| 3531208 | 3531645 | Rv3162c | Rv3162c | 438 | 3486247 | 3486837 | BCG_3186c | BCG_3186c | 591 | 3466064 | 3466654 | BCGDan_3219 | BCGDan_3219 | 591 | 3466129 | 3466719 | BCGDan_sapMKO_3220 | BCGDan_sapMKO_3220 | 591 |

| | | | | | | | | | | | | | | | | | | | |
|---|---|---|---|---|---|---|---|---|---|---|---|---|---|---|---|---|---|---|---|
| 3531642 | 3532913 | Rv3163c | Rv3163c | 1272 | 3486834 | 3488105 | BCG_3187c | BCG_3187c | 1272 | 3466651 | 3467922 | BCGDan_3220 | BCGDan_3220 | 1272 | 3466716 | 3467987 | BCGDan_sapMKO_3221 | BCGDan_sapMKO_3221 | 1272 |
| 3532943 | 3533905 | moxR3 | Rv3164c | 963 | 3488135 | 3489097 | moxR3 | BCG_3188c | 963 | 3467952 | 3468914 | moxR3 | BCGDan_3221 | 963 | 3468017 | 3468973 | moxR3 | BCGDan_sapMKO_3222 | 957 |
| 3533913 | 3534395 | Rv3165c | Rv3165c | 483 | 3489105 | 3489587 | BCG_3189c | BCG_3189c | 483 | 3468922 | 3469404 | BCGDan_3222 | BCGDan_3222 | 483 | 3468987 | 3469469 | BCGDan_sapMKO_3223 | BCGDan_sapMKO_3223 | 483 |
| 3534392 | 3535351 | Rv3166c | Rv3166c | 960 | 3489584 | 3490543 | BCG_3190c | BCG_3190c | 960 | 3469401 | 3470360 | BCGDan_3223 | BCGDan_3223 | 960 | 3469466 | 3470374 | BCGDan_sapMKO_3224 | BCGDan_sapMKO_3224 | 909 |
| 3535431 | 3536057 | Rv3167c | Rv3167c | 627 | 3490623 | 3491249 | BCG_3191c | BCG_3191c | 627 | 3470440 | 3471066 | BCGDan_3224 | BCGDan_3224 | 627 | 3470505 | 3471131 | BCGDan_sapMKO_3225 | BCGDan_sapMKO_3225 | 627 |
| 3536102 | 3537238 | Rv3168 | Rv3168 | 1137 | 3491294 | 3492430 | BCG_3192 | BCG_3192 | 1137 | 3471111 | 3472247 | BCGDan_3225 | BCGDan_3225 | 1137 | 3471176 | 3472312 | BCGDan_sapMKO_3226 | BCGDan_sapMKO_3226 | 1137 |
| 3537238 | 3538362 | Rv3169 | Rv3169 | 1125 | 3492430 | 3493554 | BCG_3193 | BCG_3193 | 1125 | 3472247 | 3473371 | BCGDan_3226 | BCGDan_3226 | 1125 | 3472312 | 3473436 | BCGDan_sapMKO_3227 | BCGDan_sapMKO_3227 | 1125 |
| 3538505 | 3539851 | aofH | Rv3170 | 1347 | 3493697 | 3495043 | aofH | BCG_3194 | 1347 | 3473514 | 3474860 | aofH | BCGDan_3227 | 1347 | 3473579 | 3474925 | aofH | BCGDan_sapMKO_3228 | 1347 |
| 3539846 | 3540745 | hpx | Rv3171c | 900 | 3495038 | 3495937 | hpx | BCG_3195c | 900 | 3474855 | 3475754 | hpx | BCGDan_3228 | 900 | 3474920 | 3475819 | hpx | BCGDan_sapMKO_3229 | 900 |
| 3540882 | 3541364 | Rv3172c | Rv3172c | 483 | 3496074 | 3496556 | BCG_3196c | BCG_3196c | 483 | 3475891 | 3476373 | BCGDan_3229 | BCGDan_3229 | 483 | 3475956 | 3476438 | BCGDan_sapMKO_3230 | BCGDan_sapMKO_3230 | 483 |
| 3541443 | 3542045 | Rv3173c | Rv3173c | 603 | 3496635 | 3497237 | BCG_3197c | BCG_3197c | 603 | 3476452 | 3477054 | BCGDan_3230 | BCGDan_3230 | 603 | 3476517 | 3477053 | BCGDan_sapMKO_3231 | BCGDan_sapMKO_3231 | 537 |
| 3542138 | 3542845 | Rv3174 | Rv3174 | 708 | 3497330 | 3498037 | BCG_3198 | BCG_3198 | 708 | 3477147 | 3477854 | BCGDan_3231 | BCGDan_3231 | 708 | 3477212 | 3477919 | BCGDan_sapMKO_3232 | BCGDan_sapMKO_3232 | 708 |
| 3542860 | 3544347 | Rv3175 | Rv3175 | 1488 | 3498052 | 3499539 | BCG_3199 | BCG_3199 | 1488 | 3477869 | 3479356 | BCGDan_3232 | BCGDan_3232 | 1488 | 3477934 | 3479421 | BCGDan_sapMKO_3233 | BCGDan_sapMKO_3233 | 1488 |
| NA | NA | NA | NA | NA | 3499536 | 3499853 | mesTb | BCG_3200c | 318 | 3479353 | 3479670 | mesTb | BCGDan_3233 | 318 | 3479418 | 3479735 | mesTb | BCGDan_sapMKO_3234 | 318 |
| 3544344 | 3545300 | mesT | Rv3176c | 957 | 3499865 | 3500491 | mesTa | BCG_3201c | 627 | 3479682 | 3480308 | mesTa | BCGDan_3234 | 627 | 3479747 | 3480373 | mesTa | BCGDan_sapMKO_3235 | 627 |
| 3545447 | 3546307 | Rv3177 | Rv3177 | 861 | 3500638 | 3501498 | BCG_3202 | BCG_3202 | 861 | 3480455 | 3481315 | BCGDan_3235 | BCGDan_3235 | 861 | 3480520 | 3481380 | BCGDan_sapMKO_3236 | BCGDan_sapMKO_3236 | 861 |
| 3546438 | 3546797 | Rv3178 | Rv3178 | 360 | 3501629 | 3501988 | BCG_3203 | BCG_3203 | 360 | 3481446 | 3481805 | BCGDan_3236 | BCGDan_3236 | 360 | 3481511 | 3481870 | BCGDan_sapMKO_3237 | BCGDan_sapMKO_3237 | 360 |
| NA | NA | NA | NA | NA | 3502228 | 3502521 | BCG_3204 | BCG_3204 | 294 | 3482045 | 3482338 | BCGDan_3237 | BCGDan_3237 | 294 | 3482110 | 3482403 | BCGDan_sapMKO_3238 | BCGDan_sapMKO_3238 | 294 |
| 3547618 | 3548907 | Rv3179 | Rv3179 | 1290 | 3502809 | 3504098 | BCG_3205 | BCG_3205 | 1290 | 3482626 | 3483915 | BCGDan_3238 | BCGDan_3238 | 1290 | 3482691 | 3483980 | BCGDan_sapMKO_3239 | BCGDan_sapMKO_3239 | 1290 |
| 3549254 | 3549688 | Rv3180c | Rv3180c | 435 | 3504445 | 3504879 | BCG_3206c | BCG_3206c | 435 | 3484262 | 3484696 | BCGDan_3239 | BCGDan_3239 | 435 | 3484327 | 3484761 | BCGDan_sapMKO_3240 | BCGDan_sapMKO_3240 | 435 |
| 3549691 | 3550143 | Rv3181c | Rv3181c | 453 | 3504882 | 3505334 | BCG_3207c | BCG_3207c | 453 | 3484699 | 3485151 | BCGDan_3240 | BCGDan_3240 | 453 | 3484764 | 3485216 | BCGDan_sapMKO_3241 | BCGDan_sapMKO_3241 | 453 |
| 3550374 | 3550718 | Rv3182 | Rv3182 | 345 | 3505565 | 3505909 | BCG_3208 | BCG_3208 | 345 | 3485382 | 3485726 | BCGDan_3241 | BCGDan_3241 | 345 | 3485453 | 3485791 | BCGDan_sapMKO_3242 | BCGDan_sapMKO_3242 | 339 |
| 3550715 | 3551044 | Rv3183 | Rv3183 | 330 | 3505906 | 3506235 | BCG_3209 | BCG_3209 | 330 | 3485723 | 3486052 | BCGDan_3242 | BCGDan_3242 | 330 | 3485788 | 3486117 | BCGDan_sapMKO_3243 | BCGDan_sapMKO_3243 | 330 |
| 3554298 | 3554645 | Rv3188 | Rv3188 | 348 | 3506773 | 3507120 | BCG_3210 | BCG_3210 | 348 | 3486590 | 3486937 | BCGDan_3243 | BCGDan_3243 | 348 | 3486655 | 3487002 | BCGDan_sapMKO_3244 | BCGDan_sapMKO_3244 | 348 |
| 3554642 | 3555262 | Rv3189 | Rv3189 | 621 | 3507117 | 3507737 | BCG_3211 | BCG_3211 | 621 | 3486934 | 3487554 | BCGDan_3244 | BCGDan_3244 | 621 | 3486999 | 3487619 | BCGDan_sapMKO_3245 | BCGDan_sapMKO_3245 | 621 |
| 3555422 | 3556687 | Rv3190c | Rv3190c | 1266 | 3507897 | 3509162 | BCG_3212c | BCG_3212c | 1266 | 3487714 | 3488979 | BCGDan_3245 | BCGDan_3245 | 1266 | 3487779 | 3489044 | BCGDan_sapMKO_3246 | BCGDan_sapMKO_3246 | 1266 |
| 3557311 | 3558345 | Rv3191c | Rv3191c | 1035 | 3509786 | 3510820 | BCG_3213c | BCG_3213c | 1035 | 3489603 | 3490637 | BCGDan_3246 | BCGDan_3246 | 1035 | 3489668 | 3490852 | BCGDan_sapMKO_3247 | BCGDan_sapMKO_3247 | 1185 |
| 3559563 | 3560024 | Rv3192 | Rv3192 | 462 | 3512035 | 3512496 | BCG_3214 | BCG_3214 | 462 | 3491852 | 3492313 | BCGDan_3248 | BCGDan_3248 | 462 | 3491914 | 3492378 | BCGDan_sapMKO_3249 | BCGDan_sapMKO_3249 | 465 |
| 3560194 | 3563172 | Rv3193c | Rv3193c | 2979 | 3512666 | 3515644 | BCG_3215c | BCG_3215c | 2979 | 3492483 | 3495461 | BCGDan_3249 | BCGDan_3249 | 2979 | 3492548 | 3495520 | BCGDan_sapMKO_3250 | BCGDan_sapMKO_3250 | 2973 |
| 3563264 | 3564286 | Rv3194c | Rv3194c | 1023 | 3515736 | 3516758 | BCG_3216c | BCG_3216c | 1023 | 3495553 | 3496575 | BCGDan_3250 | BCGDan_3250 | 1023 | 3495618 | 3496616 | BCGDan_sapMKO_3251 | BCGDan_sapMKO_3251 | 999 |
| 3564364 | 3565782 | Rv3195 | Rv3195 | 1419 | 3516836 | 3518254 | BCG_3217 | BCG_3217 | 1419 | 3496653 | 3498071 | BCGDan_3251 | BCGDan_3251 | 1419 | 3496718 | 3498136 | BCGDan_sapMKO_3252 | BCGDan_sapMKO_3252 | 1419 |
| 3565788 | 3566687 | Rv3196 | Rv3196 | 900 | 3518260 | 3519159 | BCG_3218 | BCG_3218 | 900 | 3498077 | 3498976 | BCGDan_3252 | BCGDan_3252 | 900 | 3498217 | 3499041 | BCGDan_sapMKO_3253 | BCGDan_sapMKO_3253 | 825 |
| 3566696 | 3566896 | Rv3196A | Rv3196A | 201 | 3519168 | 3519368 | BCG_3219c | BCG_3219c | 201 | 3498985 | 3499185 | BCGDan_3253 | BCGDan_3253 | 201 | 3499050 | 3499250 | BCGDan_sapMKO_3254 | BCGDan_sapMKO_3254 | 201 |
| 3567024 | 3568367 | Rv3197 | Rv3197 | 1344 | 3519506 | 3520849 | BCG_3220 | BCG_3220 | 1344 | 3499323 | 3500666 | BCGDan_3254 | BCGDan_3254 | 1344 | 3499388 | 3500731 | BCGDan_sapMKO_3255 | BCGDan_sapMKO_3255 | 1344 |
| 3568401 | 3568679 | whiB7 | Rv3197A | 279 | 3520883 | 3521161 | whiB7 | BCG_3221c | 279 | 3500700 | 3500978 | whiB7_1 | BCGDan_3255 | 279 | 3500765 | 3501043 | whiB7 | BCGDan_sapMKO_3256 | 279 |
| 3569109 | 3571211 | uvrD2 | Rv3198c | 2103 | 3521591 | 3523693 | UvrD2 | BCG_3222c | 2103 | 3501408 | 3503510 | UvrD2_1 | BCGDan_3256 | 2103 | 3501473 | 3503641 | UvrD2 | BCGDan_sapMKO_3257 | 2169 |
| 3571335 | 3571589 | Rv3198A | Rv3198A | 255 | 3523817 | 3524071 | BCG_3223 | BCG_3223 | 255 | 3503634 | 3503888 | BCGDan_3257 | BCGDan_3257 | 255 | 3503699 | 3503953 | BCGDan_sapMKO_3258 | BCGDan_sapMKO_3258 | 255 |
| 3571602 | 3572543 | nudC | Rv3199c | 942 | 3524084 | 3525025 | nudC | BCG_3224c | 942 | 3503901 | 3504842 | nudC_1 | BCGDan_3258 | 942 | 3503966 | 3504889 | nudC | BCGDan_sapMKO_3259 | 924 |
| 3572602 | 3573669 | Rv3200c | Rv3200c | 1068 | 3525084 | 3526151 | BCG_3225c | BCG_3225c | 1068 | 3504901 | 3505968 | BCGDan_3259 | BCGDan_3259 | 1068 | 3504966 | 3506033 | BCGDan_sapMKO_3260 | BCGDan_sapMKO_3260 | 1068 |
| 3573731 | 3577036 | Rv3201c | Rv3201c | 3306 | 3526213 | 3529518 | BCG_3226c | BCG_3226c | 3306 | 3506030 | 3509335 | BCGDan_3260 | BCGDan_3260 | 3306 | 3506095 | 3509457 | BCGDan_sapMKO_3261 | BCGDan_sapMKO_3261 | 3363 |
| 3577033 | 3580200 | Rv3202c | Rv3202c | 3168 | 3529515 | 3532682 | BCG_3227c | BCG_3227c | 3168 | 3509332 | 3512499 | BCGDan_3261 | BCGDan_3261 | 3168 | 3509397 | 3512564 | BCGDan_sapMKO_3262 | BCGDan_sapMKO_3262 | 3168 |
| NA | NA | NA | NA | NA | 3532759 | 3532977 | BCG_3228 | BCG_3228 | 219 | 3512576 | 3512794 | BCGDan_3262 | BCGDan_3262 | 219 | 3512641 | 3512859 | BCGDan_sapMKO_3263 | BCGDan_sapMKO_3263 | 219 |
| 3580638 | 3581312 | lipV | Rv3203 | 675 | 3533119 | 3533793 | lipV | BCG_3229 | 675 | 3512936 | 3513610 | lipV_1 | BCGDan_3263 | 675 | 3512890 | 3513675 | lipV | BCGDan_sapMKO_3264 | 786 |
| 3581315 | 3581620 | Rv3204 | Rv3204 | 306 | 3533796 | 3534101 | BCG_3230 | BCG_3230 | 306 | 3513613 | 3513918 | BCGDan_3264 | BCGDan_3264 | 306 | 3513678 | 3513983 | BCGDan_sapMKO_3265 | BCGDan_sapMKO_3265 | 306 |
| 3581627 | 3582505 | Rv3205c | Rv3205c | 879 | 3534108 | 3534986 | BCG_3231c | BCG_3231c | 879 | 3513925 | 3514803 | BCGDan_3265 | BCGDan_3265 | 879 | 3513990 | 3514844 | BCGDan_sapMKO_3266 | BCGDan_sapMKO_3266 | 855 |
| 3582532 | 3583710 | moeB1 | Rv3206c | 1179 | 3535013 | 3536191 | moeB1 | BCG_3232c | 1179 | 3514830 | 3516008 | moeB1_1 | BCGDan_3266 | 1179 | 3514895 | 3515935 | moeB1 | BCGDan_sapMKO_3267 | 1041 |
| 3583801 | 3584658 | Rv3207c | Rv3207c | 858 | 3536282 | 3537139 | BCG_3233c | BCG_3233c | 858 | 3516099 | 3516956 | BCGDan_3267 | BCGDan_3267 | 858 | 3516164 | 3517150 | BCGDan_sapMKO_3268 | BCGDan_sapMKO_3268 | 987 |
| 3585004 | 3585690 | Rv3208 | Rv3208 | 687 | 3537485 | 3538171 | BCG_3234 | BCG_3234 | 687 | 3517302 | 3517988 | BCGDan_3268 | BCGDan_3268 | 687 | 3517367 | 3518053 | BCGDan_sapMKO_3269 | BCGDan_sapMKO_3269 | 687 |
| 3585677 | 3585949 | TB9.4 | Rv3208A | 273 | 3538158 | 3538430 | TB9.4 | BCG_3235c | 273 | 3517975 | 3518247 | TB9.4_1 | BCGDan_3269 | 273 | 3518040 | 3518327 | TB9.4 | BCGDan_sapMKO_3270 | 288 |
| 3586274 | 3586834 | Rv3209 | Rv3209 | 561 | 3538755 | 3539315 | BCG_3236 | BCG_3236 | 561 | 3518572 | 3519132 | BCGDan_3270 | BCGDan_3270 | 561 | 3518637 | 3519197 | BCGDan_sapMKO_3271 | BCGDan_sapMKO_3271 | 561 |
| 3586844 | 3587539 | Rv3210c | Rv3210c | 696 | 3539325 | 3540020 | BCG_3237c | BCG_3237c | 696 | 3519142 | 3519837 | BCGDan_3271 | BCGDan_3271 | 696 | 3519207 | 3519902 | BCGDan_sapMKO_3272 | BCGDan_sapMKO_3272 | 696 |
| 3587798 | 3589381 | rhlE | Rv3211 | 1584 | 3540279 | 3541862 | rhlE | BCG_3238 | 1584 | 3520096 | 3521679 | rhlE_1 | BCGDan_3272 | 1584 | 3520161 | 3521744 | rhlE | BCGDan_sapMKO_3273 | 1584 |
| 3589394 | 3590617 | Rv3212 | Rv3212 | 1224 | 3541875 | 3543098 | BCG_3239 | BCG_3239 | 1224 | 3521692 | 3522915 | BCGDan_3273 | BCGDan_3273 | 1224 | 3521757 | 3522980 | BCGDan_sapMKO_3274 | BCGDan_sapMKO_3274 | 1224 |
| 3590692 | 3591492 | Rv3213c | Rv3213c | 801 | 3543174 | 3543974 | BCG_3240c | BCG_3240c | 801 | 3522991 | 3523791 | BCGDan_3274 | BCGDan_3274 | 801 | 3523056 | 3523856 | BCGDan_sapMKO_3275 | BCGDan_sapMKO_3275 | 801 |
| 3591646 | 3592257 | gpm2 | Rv3214 | 612 | 3544128 | 3544739 | gpm2_1 | BCG_3241 | 612 | 3523945 | 3524556 | gpm2_1 | BCGDan_3275 | 612 | 3524010 | 3524621 | gpm2 | BCGDan_sapMKO_3276 | 612 |
| 3592254 | 3593372 | entC | Rv3215 | 1119 | 3544736 | 3545854 | entC_1 | BCG_3242 | 1119 | 3524553 | 3525671 | entC_1 | BCGDan_3276 | 1119 | 3524618 | 3525736 | entC | BCGDan_sapMKO_3277 | 1119 |
| 3593439 | 3593852 | Rv3216 | Rv3216 | 414 | 3546002 | 3546334 | BCG_3243 | BCG_3243 | 333 | 3525819 | 3526151 | BCGDan_3277 | BCGDan_3277 | 333 | 3525926 | 3526216 | BCGDan_sapMKO_3278 | BCGDan_sapMKO_3278 | 291 |
| 3593804 | 3594235 | Rv3217c | Rv3217c | 432 | 3546286 | 3546717 | BCG_3244c | BCG_3244c | 432 | 3526103 | 3526534 | BCGDan_3278 | BCGDan_3278 | 432 | 3526168 | 3526575 | BCGDan_sapMKO_3279 | BCGDan_sapMKO_3279 | 408 |
| 3594468 | 3595433 | Rv3218 | Rv3218 | 966 | 3546950 | 3547915 | BCG_3245 | BCG_3245 | 966 | 3526767 | 3527732 | BCGDan_3279 | BCGDan_3279 | 966 | 3526832 | 3527797 | BCGDan_sapMKO_3280 | BCGDan_sapMKO_3280 | 966 |
| 3595713 | 3595967 | whiB1 | Rv3219 | 255 | 3548195 | 3548449 | whiB1_1 | BCG_3246 | 255 | 3528012 | 3528266 | whiB1_1 | BCGDan_3280 | 255 | 3528077 | 3528331 | whiB1 | BCGDan_sapMKO_3281 | 255 |
| 3596029 | 3597534 | Rv3220c | Rv3220c | 1506 | 3548511 | 3550016 | BCG_3247c | BCG_3247c | 1506 | 3528328 | 3529833 | BCGDan_3281 | BCGDan_3281 | 1506 | 3528393 | 3529898 | BCGDan_sapMKO_3282 | BCGDan_sapMKO_3282 | 1506 |
| 3597551 | 3597766 | TB7.3 | Rv3221c | 216 | 3550033 | 3550248 | TB7.3_1 | BCG_3248c | 216 | 3529850 | 3530065 | TB7.3_1 | BCGDan_3282 | 216 | 3529915 | 3530130 | TB7.3 | BCGDan_sapMKO_3283 | 216 |
| 3598051 | 3598356 | rshA | Rv3221A | 306 | 3550533 | 3550838 | BCG_3249c | BCG_3249c | 306 | 3530350 | 3530655 | BCGDan_3283 | BCGDan_3283 | 306 | 3530415 | 3530720 | BCGDan_sapMKO_3284 | BCGDan_sapMKO_3284 | 306 |
| 3598353 | 3598904 | Rv3222c | Rv3222c | 552 | 3550835 | 3551386 | BCG_3250c | BCG_3250c | 552 | 3530652 | 3531203 | BCGDan_3284 | BCGDan_3284 | 552 | 3530717 | 3531268 | BCGDan_sapMKO_3285 | BCGDan_sapMKO_3285 | 552 |
| 3598901 | 3599551 | sigH | Rv3223c | 651 | 3551383 | 3552033 | sigH_1 | BCG_3251c | 651 | 3531200 | 3531850 | sigH_1 | BCGDan_3285 | 651 | 3531265 | 3531915 | sigH | BCGDan_sapMKO_3286 | 651 |
| 3599851 | 3600699 | Rv3224 | Rv3224 | 849 | 3552333 | 3553181 | BCG_3252 | BCG_3252 | 849 | 3532150 | 3532998 | BCGDan_3286 | BCGDan_3286 | 849 | 3532215 | 3533063 | BCGDan_sapMKO_3287 | BCGDan_sapMKO_3287 | 849 |
| 3600635 | 3600823 | Rv3224A | Rv3224A | 189 | 3553117 | 3553305 | BCG_3253 | BCG_3253 | 189 | 3532934 | 3533122 | BCGDan_3287 | BCGDan_3287 | 189 | 3532999 | 3533187 | BCGDan_sapMKO_3288 | BCGDan_sapMKO_3288 | 189 |
| 3600801 | 3601019 | Rv3224B | Rv3224B | 219 | 3553283 | 3553501 | BCG_3254 | BCG_3254 | 219 | 3533100 | 3533318 | BCGDan_3288 | BCGDan_3288 | 219 | 3533165 | 3533383 | BCGDan_sapMKO_3289 | BCGDan_sapMKO_3289 | 219 |
| 3601016 | 3602440 | Rv3225c | Rv3225c | 1425 | 3553498 | 3554922 | BCG_3255c | BCG_3255c | 1425 | 3533315 | 3534739 | BCGDan_3289 | BCGDan_3289 | 1425 | 3533380 | 3534804 | BCGDan_sapMKO_3290 | BCGDan_sapMKO_3290 | 1425 |
| 3602564 | 3603322 | Rv3226c | Rv3226c | 759 | 3555046 | 3555804 | BCG_3256c | BCG_3256c | 759 | 3534863 | 3535621 | BCGDan_3290 | BCGDan_3290 | 759 | 3534928 | 3535686 | BCGDan_sapMKO_3291 | BCGDan_sapMKO_3291 | 759 |
| 3603377 | 3604729 | aroA | Rv3227 | 1353 | 3555859 | 3557211 | aroA_1 | BCG_3257 | 1353 | 3535676 | 3537028 | aroA_1 | BCGDan_3291 | 1353 | 3535777 | 3537093 | aroA | BCGDan_sapMKO_3292 | 1317 |
| 3604726 | 3605718 | Rv3228 | Rv3228 | 993 | 3557208 | 3558200 | BCG_3258 | BCG_3258 | 993 | 3537025 | 3538017 | BCGDan_3292 | BCGDan_3292 | 993 | 3537024 | 3538082 | BCGDan_sapMKO_3293 | BCGDan_sapMKO_3293 | 1059 |
| 3605751 | 3607034 | desA3 | Rv3229c | 1284 | 3558233 | 3559516 | desA3_1 | BCG_3259c | 1284 | 3538050 | 3539333 | desA3_1 | BCGDan_3293 | 1284 | 3538115 | 3539398 | desA3 | BCGDan_sapMKO_3294 | 1284 |
| 3607112 | 3608254 | Rv3230c | Rv3230c | 1143 | 3559594 | 3560736 | BCG_3260c | BCG_3260c | 1143 | 3539411 | 3540553 | BCGDan_3294 | BCGDan_3294 | 1143 | 3539476 | 3540618 | BCGDan_sapMKO_3295 | BCGDan_sapMKO_3295 | 1143 |
| 3608364 | 3608873 | Rv3231c | Rv3231c | 510 | 3560846 | 3561355 | BCG_3261c | BCG_3261c | 510 | 3540663 | 3541172 | BCGDan_3295 | BCGDan_3295 | 510 | 3540728 | 3541237 | BCGDan_sapMKO_3296 | BCGDan_sapMKO_3296 | 510 |
| 3608870 | 3609757 | ppk2 | Rv3232c | 888 | 3561352 | 3562239 | pvdS | BCG_3262c | 888 | 3541169 | 3542056 | pvdS | BCGDan_3296 | 888 | 3541234 | 3542100 | pvdS | BCGDan_sapMKO_3297 | 867 |
| 3609781 | 3610371 | Rv3233c | Rv3233c | 591 | 3562263 | 3563672 | BCG_3263c | BCG_3263c | 1410 | 3542080 | 3543489 | BCGDan_3297 | BCGDan_3297 | 1410 | 3542145 | 3543554 | BCGDan_sapMKO_3298 | BCGDan_sapMKO_3298 | 1410 |
| 3611300 | 3611941 | Rv3235 | Rv3235 | 642 | 3563783 | 3564424 | BCG_3264 | BCG_3264 | 642 | 3543600 | 3544241 | BCGDan_3298 | BCGDan_3298 | 642 | 3543755 | 3544306 | BCGDan_sapMKO_3299 | BCGDan_sapMKO_3299 | 552 |
| 3611959 | 3613116 | Rv3236c | Rv3236c | 1158 | 3564442 | 3565599 | BCG_3265c | BCG_3265c | 1158 | 3544259 | 3545416 | BCGDan_3299 | BCGDan_3299 | 1158 | 3544324 | 3545481 | BCGDan_sapMKO_3300 | BCGDan_sapMKO_3300 | 1158 |
| 3613121 | 3613603 | Rv3237c | Rv3237c | 483 | 3565604 | 3566086 | BCG_3266c | BCG_3266c | 483 | 3545421 | 3545903 | BCGDan_3300 | BCGDan_3300 | 483 | 3545486 | 3545968 | BCGDan_sapMKO_3301 | BCGDan_sapMKO_3301 | 483 |
| 3613664 | 3614398 | Rv3238c | Rv3238c | 735 | 3566147 | 3566881 | BCG_3267c | BCG_3267c | 735 | 3545964 | 3546698 | BCGDan_3301 | BCGDan_3301 | 735 | 3546029 | 3546763 | BCGDan_sapMKO_3302 | BCGDan_sapMKO_3302 | 735 |
| 3614457 | 3617603 | Rv3239c | Rv3239c | 3147 | 3566940 | 3570086 | BCG_3268c | BCG_3268c | 3147 | 3546757 | 3549903 | BCGDan_3302 | BCGDan_3302 | 3147 | 3546822 | 3549968 | BCGDan_sapMKO_3303 | BCGDan_sapMKO_3303 | 3147 |
| 3617682 | 3620531 | secA1 | Rv3240c | 2850 | 3570165 | 3573014 | secA1 | BCG_3269c | 2850 | 3549982 | 3552831 | secA1 | BCGDan_3303 | 2850 | 3550047 | 3552896 | secA1 | BCGDan_sapMKO_3304 | 2850 |
| 3620610 | 3621254 | Rv3241c | Rv3241c | 645 | 3573093 | 3573737 | BCG_3270c | BCG_3270c | 645 | 3552910 | 3553554 | BCGDan_3304 | BCGDan_3304 | 645 | 3552975 | 3553619 | BCGDan_sapMKO_3305 | BCGDan_sapMKO_3305 | 645 |
| 3621570 | 3622211 | Rv3242c | Rv3242c | 642 | 3574053 | 3574694 | BCG_3271c | BCG_3271c | 642 | 3553870 | 3554511 | BCGDan_3305 | BCGDan_3305 | 642 | 3553935 | 3554576 | BCGDan_sapMKO_3306 | BCGDan_sapMKO_3306 | 642 |
| 3622249 | 3623091 | Rv3243c | Rv3243c | 843 | 3574732 | 3575574 | BCG_3272c | BCG_3272c | 843 | 3554549 | 3555391 | BCGDan_3306 | BCGDan_3306 | 843 | 3554614 | 3555402 | BCGDan_sapMKO_3307 | BCGDan_sapMKO_3307 | 789 |

| | | | | | | | | | | | | | | | | | | | |
|---|---|---|---|---|---|---|---|---|---|---|---|---|---|---|---|---|---|---|---|
| 3623159 | 3624910 | lpqB | Rv3244c | 1752 | 3575642 | 3577393 | lpqB | BCG_3273c | 1752 | 3555459 | 3557210 | lpqB | BCGDan_3307 | 1752 | 3555524 | 3557287 | lpqB | BCGDan_sapMKO_3308 | 1764 |
| 3624910 | 3626613 | mtrB | Rv3245c | 1704 | 3577393 | 3579096 | mtrB | BCG_3274c | 1704 | 3557210 | 3558913 | mtrB | BCGDan_3308 | 1704 | 3557275 | 3558924 | mtrB | BCGDan_sapMKO_3309 | 1650 |
| 3626663 | 3627349 | mtrA | Rv3246c | 687 | 3579146 | 3579832 | mtrA | BCG_3275c | 687 | 3558963 | 3559649 | mtrA | BCGDan_3309 | 687 | 3559028 | 3559714 | mtrA | BCGDan_sapMKO_3310 | 687 |
| 3627419 | 3628063 | tmk | Rv3247c | 645 | 3579902 | 3580546 | tmk | BCG_3276c | 645 | 3559719 | 3560363 | tmk | BCGDan_3310 | 645 | 3559784 | 3560428 | tmk | BCGDan_sapMKO_3311 | 645 |
| 3628160 | 3629647 | sahH | Rv3248c | 1488 | 3580643 | 3582130 | sahH | BCG_3277c | 1488 | 3560460 | 3561947 | sahH | BCGDan_3311 | 1488 | 3560525 | 3562024 | sahH | BCGDan_sapMKO_3312 | 1500 |
| 3629752 | 3630387 | Rv3249c | Rv3249c | 636 | 3582235 | 3582870 | BCG_3278c | BCG_3278c | 636 | 3562052 | 3562687 | BCGDan_3312 | BCGDan_3312 | 636 | 3562117 | 3562722 | BCGDan_sapMKO_3313 | BCGDan_sapMKO_3313 | 606 |
| 3630384 | 3630566 | rubB | Rv3250c | 183 | 3582867 | 3583049 | BCG_3279c | BCG_3279c | 183 | 3562684 | 3562866 | BCGDan_3313 | BCGDan_3313 | 183 | 3562749 | 3562931 | BCGDan_sapMKO_3314 | BCGDan_sapMKO_3314 | 183 |
| 3630571 | 3630738 | rubA | Rv3251c | 168 | 3583054 | 3583221 | BCG_3280c | BCG_3280c | 168 | 3562871 | 3563038 | BCGDan_3314 | BCGDan_3314 | 168 | 3562936 | 3563103 | BCGDan_sapMKO_3315 | BCGDan_sapMKO_3315 | 168 |
| 3630738 | 3631988 | alkB | Rv3252c | 1251 | 3583221 | 3584471 | alkB | BCG_3281c | 1251 | 3563038 | 3564288 | alkB | BCGDan_3315 | 1251 | 3563103 | 3564353 | alkB | BCGDan_sapMKO_3316 | 1251 |
| 3632097 | 3633584 | Rv3253c | Rv3253c | 1488 | 3584580 | 3586067 | BCG_3282c | BCG_3282c | 1488 | 3564397 | 3565884 | BCGDan_3316 | BCGDan_3316 | 1488 | 3564462 | 3565931 | BCGDan_sapMKO_3317 | BCGDan_sapMKO_3317 | 1470 |
| 3633675 | 3635063 | Rv3254 | Rv3254 | 1389 | 3586158 | 3587546 | BCG_3283 | BCG_3283 | 1389 | 3565975 | 3567363 | BCGDan_3317 | BCGDan_3317 | 1389 | 3566040 | 3567428 | BCGDan_sapMKO_3318 | BCGDan_sapMKO_3318 | 1389 |
| 3635041 | 3636267 | manA | Rv3255c | 1227 | 3587524 | 3588750 | manA | BCG_3284c | 1227 | 3567341 | 3568567 | manA | BCGDan_3318 | 1227 | 3567406 | 3568632 | manA | BCGDan_sapMKO_3319 | 1227 |
| 3636275 | 3637315 | Rv3256c | Rv3256c | 1041 | 3588758 | 3589798 | BCG_3285c | BCG_3285c | 1041 | 3568575 | 3569615 | BCGDan_3319 | BCGDan_3319 | 1041 | 3568640 | 3569680 | BCGDan_sapMKO_3320 | BCGDan_sapMKO_3320 | 1041 |
| 3637312 | 3638709 | pmmA | Rv3257c | 1398 | 3589795 | 3591192 | manB | BCG_3286c | 1398 | 3569612 | 3571009 | manB | BCGDan_3320 | 1398 | 3569677 | 3571053 | manB | BCGDan_sapMKO_3321 | 1377 |
| 3638811 | 3639302 | Rv3258c | Rv3258c | 492 | 3591294 | 3591785 | BCG_3287c | BCG_3287c | 492 | 3571111 | 3571602 | BCGDan_3321 | BCGDan_3321 | 492 | 3571176 | 3571667 | BCGDan_sapMKO_3322 | BCGDan_sapMKO_3322 | 492 |
| 3639425 | 3639844 | Rv3259 | Rv3259 | 420 | 3591908 | 3592327 | BCG_3288 | BCG_3288 | 420 | 3571725 | 3572144 | BCGDan_3322 | BCGDan_3322 | 420 | 3571790 | 3572209 | BCGDan_sapMKO_3323 | BCGDan_sapMKO_3323 | 420 |
| 3639872 | 3640141 | whiB2 | Rv3260c | 270 | 3592355 | 3592624 | whiB2 | BCG_3289c | 270 | 3572172 | 3572441 | whiB2 | BCGDan_3323 | 270 | 3572237 | 3572524 | whiB2 | BCGDan_sapMKO_3324 | 288 |
| 3640543 | 3641538 | fbiA | Rv3261 | 996 | 3593026 | 3594021 | fbiA | BCG_3290 | 996 | 3572843 | 3573838 | fbiA | BCGDan_3324 | 996 | 3572908 | 3573903 | fbiA | BCGDan_sapMKO_3325 | 996 |
| 3641535 | 3642881 | fbiB | Rv3262 | 1347 | 3594018 | 3595364 | fbiB | BCG_3291 | 1347 | 3573835 | 3575181 | fbiB | BCGDan_3325 | 1347 | 3573900 | 3575246 | fbiB | BCGDan_sapMKO_3326 | 1347 |
| 3643177 | 3644838 | Rv3263 | Rv3263 | 1662 | 3595660 | 3597321 | BCG_3292 | BCG_3292 | 1662 | 3575477 | 3577138 | BCGDan_3326 | BCGDan_3326 | 1662 | 3575614 | 3577203 | BCGDan_sapMKO_3327 | BCGDan_sapMKO_3327 | 1590 |
| 3644898 | 3645977 | manB | Rv3264c | 1080 | 3597381 | 3598460 | manC | BCG_3293c | 1080 | 3577198 | 3578277 | manC | BCGDan_3327 | 1080 | 3577263 | 3578318 | manC | BCGDan_sapMKO_3328 | 1056 |
| 3645979 | 3646884 | wbbL1 | Rv3265c | 906 | 3598462 | 3599355 | wbbL1 | BCG_3294c | 894 | 3578279 | 3579172 | wbbL1 | BCGDan_3328 | 894 | 3578344 | 3579255 | wbbL1 | BCGDan_sapMKO_3329 | 912 |
| 3646895 | 3647809 | rmlD | Rv3266c | 915 | 3599366 | 3600280 | rmld | BCG_3295c | 915 | 3579183 | 3580097 | rmld | BCGDan_3329 | 915 | 3579248 | 3580162 | rmld | BCGDan_sapMKO_3330 | 915 |
| 3647885 | 3649381 | Rv3267 | Rv3267 | 1497 | 3600356 | 3601852 | BCG_3296 | BCG_3296 | 1497 | 3580173 | 3581669 | BCGDan_3330 | BCGDan_3330 | 1497 | 3580241 | 3581734 | BCGDan_sapMKO_3331 | BCGDan_sapMKO_3331 | 1494 |
| 3649420 | 3650109 | Rv3268 | Rv3268 | 690 | 3601891 | 3602580 | BCG_3297 | BCG_3297 | 690 | 3581708 | 3582397 | BCGDan_3331 | BCGDan_3331 | 690 | 3581722 | 3582462 | BCGDan_sapMKO_3332 | BCGDan_sapMKO_3332 | 741 |
| 3650234 | 3650515 | Rv3269 | Rv3269 | 282 | 3602705 | 3602986 | BCG_3298 | BCG_3298 | 282 | 3582522 | 3582803 | BCGDan_3332 | BCGDan_3332 | 282 | 3582587 | 3582868 | BCGDan_sapMKO_3333 | BCGDan_sapMKO_3333 | 282 |
| 3650526 | 3652682 | ctpC | Rv3270 | 2157 | 3602997 | 3605153 | ctpC | BCG_3299 | 2157 | 3582814 | 3584970 | ctpC | BCGDan_3333 | 2157 | 3582879 | 3585035 | ctpC | BCGDan_sapMKO_3334 | 2157 |
| 3652679 | 3653347 | Rv3271c | Rv3271c | 669 | 3605150 | 3605818 | BCG_3300c | BCG_3300c | 669 | 3584967 | 3585635 | BCGDan_3334 | BCGDan_3334 | 669 | 3585032 | 3585700 | BCGDan_sapMKO_3335 | BCGDan_sapMKO_3335 | 669 |
| 3653448 | 3654632 | Rv3272 | Rv3272 | 1185 | 3605919 | 3607103 | BCG_3301 | BCG_3301 | 1185 | 3585736 | 3586920 | BCGDan_3335 | BCGDan_3335 | 1185 | 3585801 | 3586985 | BCGDan_sapMKO_3336 | BCGDan_sapMKO_3336 | 1185 |
| 3654637 | 3656931 | Rv3273 | Rv3273 | 2295 | 3607108 | 3609402 | BCG_3302 | BCG_3302 | 2295 | 3586925 | 3589219 | BCGDan_3336 | BCGDan_3336 | 2295 | 3587014 | 3589284 | BCGDan_sapMKO_3337 | BCGDan_sapMKO_3337 | 2271 |
| 3656920 | 3658089 | fadE25 | Rv3274c | 1170 | 3609391 | 3610560 | fadE25 | BCG_3303c | 1170 | 3589208 | 3590377 | fadE25 | BCGDan_3337 | 1170 | 3589273 | 3590442 | fadE25 | BCGDan_sapMKO_3338 | 1170 |
| 3658114 | 3658638 | purE | Rv3275c | 525 | 3610585 | 3611109 | purE | BCG_3304c | 525 | 3590402 | 3590926 | purE | BCGDan_3338 | 525 | 3590467 | 3590991 | purE | BCGDan_sapMKO_3339 | 525 |
| 3658635 | 3659924 | purK | Rv3276c | 1290 | 3611106 | 3612395 | purK | BCG_3305c | 1290 | 3590923 | 3592212 | purK | BCGDan_3339 | 1290 | 3590988 | 3592214 | purK | BCGDan_sapMKO_3340 | 1227 |
| 3659878 | 3660696 | Rv3277 | Rv3277 | 819 | 3612349 | 3613167 | BCG_3306 | BCG_3306 | 819 | 3592166 | 3592984 | BCGDan_3340 | BCGDan_3340 | 819 | 3592231 | 3593049 | BCGDan_sapMKO_3341 | BCGDan_sapMKO_3341 | 819 |
| 3660651 | 3661169 | Rv3278c | Rv3278c | 519 | 3613122 | 3613640 | BCG_3307c | BCG_3307c | 519 | 3592939 | 3593457 | BCGDan_3341 | BCGDan_3341 | 519 | 3593004 | 3593522 | BCGDan_sapMKO_3342 | BCGDan_sapMKO_3342 | 519 |
| 3661212 | 3662012 | birA | Rv3279c | 801 | 3613683 | 3614483 | birA | BCG_3308c | 801 | 3593500 | 3594300 | birA | BCGDan_3342 | 801 | 3593565 | 3594365 | birA | BCGDan_sapMKO_3343 | 801 |
| 3662062 | 3663708 | accD5 | Rv3280 | 1647 | 3614533 | 3616179 | accD5 | BCG_3309 | 1647 | 3594350 | 3595996 | accD5 | BCGDan_3343 | 1647 | 3594415 | 3596061 | accD5 | BCGDan_sapMKO_3344 | 1647 |
| 3663689 | 3664222 | accE5 | Rv3281 | 534 | 3616160 | 3616630 | BCG_3310 | BCG_3310 | 471 | 3595977 | 3596447 | BCGDan_3344 | BCGDan_3344 | 471 | 3596042 | 3596512 | BCGDan_sapMKO_3345 | BCGDan_sapMKO_3345 | 471 |
| 3664219 | 3664887 | Rv3282 | Rv3282 | 669 | 3616627 | 3617295 | BCG_3311 | BCG_3311 | 669 | 3596444 | 3597112 | BCGDan_3345 | BCGDan_3345 | 669 | 3596509 | 3597177 | BCGDan_sapMKO_3346 | BCGDan_sapMKO_3346 | 669 |
| 3664928 | 3665821 | sseA | Rv3283 | 894 | 3617336 | 3618229 | sseA | BCG_3312 | 894 | 3597153 | 3598046 | sseA | BCGDan_3346 | 894 | 3597146 | 3598111 | sseA | BCGDan_sapMKO_3347 | 966 |
| 3665818 | 3666249 | Rv3284 | Rv3284 | 432 | 3618226 | 3618657 | BCG_3313 | BCG_3313 | 432 | 3598043 | 3598474 | BCGDan_3347 | BCGDan_3347 | 432 | 3598108 | 3598539 | BCGDan_sapMKO_3348 | BCGDan_sapMKO_3348 | 432 |
| 3666357 | 3668159 | accA3 | Rv3285 | 1803 | 3618765 | 3620567 | accA3 | BCG_3314 | 1803 | 3598582 | 3600384 | accA3 | BCGDan_3348 | 1803 | 3598647 | 3600449 | accA3 | BCGDan_sapMKO_3349 | 1803 |
| 3668169 | 3668954 | sigF | Rv3286c | 786 | 3620577 | 3621362 | sigF | BCG_3315c | 786 | 3600394 | 3601179 | sigF | BCGDan_3349 | 786 | 3600459 | 3601244 | sigF | BCGDan_sapMKO_3350 | 786 |
| 3668951 | 3669388 | rsbW | Rv3287c | 438 | 3621359 | 3621796 | rsbW | BCG_3316c | 438 | 3601176 | 3601613 | rsbW | BCGDan_3350 | 438 | 3601241 | 3601678 | rsbW | BCGDan_sapMKO_3351 | 438 |
| 3669586 | 3669999 | usfY | Rv3288c | 414 | 3621994 | 3622407 | usfY | BCG_3317c | 414 | 3601811 | 3602224 | usfY | BCGDan_3351 | 414 | 3601876 | 3602289 | usfY | BCGDan_sapMKO_3352 | 414 |
| 3670034 | 3670411 | Rv3289c | Rv3289c | 378 | 3622442 | 3622819 | BCG_3318c | BCG_3318c | 378 | 3602259 | 3602636 | BCGDan_3352 | BCGDan_3352 | 378 | 3602324 | 3602701 | BCGDan_sapMKO_3353 | BCGDan_sapMKO_3353 | 378 |
| 3670445 | 3671794 | lat | Rv3290c | 1350 | 3622853 | 3624202 | lat | BCG_3319c | 1350 | 3602670 | 3604019 | lat | BCGDan_3353 | 1350 | 3602735 | 3604003 | lat | BCGDan_sapMKO_3354 | 1269 |
| 3671845 | 3672297 | lrpA | Rv3291c | 453 | 3624253 | 3624705 | BCG_3320c | BCG_3320c | 453 | 3604070 | 3604522 | BCGDan_3354 | BCGDan_3354 | 453 | 3604135 | 3604587 | BCGDan_sapMKO_3355 | BCGDan_sapMKO_3355 | 453 |
| 3672328 | 3673575 | Rv3292 | Rv3292 | 1248 | 3624736 | 3625983 | BCG_3321 | BCG_3321 | 1248 | 3604553 | 3605800 | BCGDan_3355 | BCGDan_3355 | 1248 | 3604618 | 3605865 | BCGDan_sapMKO_3356 | BCGDan_sapMKO_3356 | 1248 |
| 3673602 | 3675086 | pcd | Rv3293 | 1485 | 3626010 | 3627494 | pcd_1 | BCG_3322 | 1485 | 3605827 | 3607311 | pcd_1 | BCGDan_3356 | 1485 | 3605892 | 3607376 | pcd | BCGDan_sapMKO_3357 | 1485 |
| 3675186 | 3675995 | Rv3294c | Rv3294c | 810 | 3627594 | 3628403 | BCG_3323c | BCG_3323c | 810 | 3607411 | 3608220 | BCGDan_3357 | BCGDan_3357 | 810 | 3607476 | 3608285 | BCGDan_sapMKO_3358 | BCGDan_sapMKO_3358 | 810 |
| 3676066 | 3676731 | Rv3295 | Rv3295 | 666 | 3628474 | 3629139 | BCG_3324 | BCG_3324 | 666 | 3608291 | 3608956 | BCGDan_3358 | BCGDan_3358 | 666 | 3608356 | 3609021 | BCGDan_sapMKO_3359 | BCGDan_sapMKO_3359 | 666 |
| 3676775 | 3681316 | lhr | Rv3296 | 4542 | 3629183 | 3633724 | lhr_1 | BCG_3325 | 4542 | 3609000 | 3613541 | lhr_1 | BCGDan_3359 | 4542 | 3609065 | 3613606 | lhr | BCGDan_sapMKO_3360 | 4542 |
| 3681320 | 3682087 | nei | Rv3297 | 768 | 3633728 | 3634495 | nei_1 | BCG_3326 | 768 | 3613545 | 3614312 | nei_1 | BCGDan_3360 | 768 | 3613610 | 3614377 | nei | BCGDan_sapMKO_3361 | 768 |
| 3682110 | 3683024 | lpqC | Rv3298c | 915 | 3634518 | 3635432 | lpqC_1 | BCG_3327c | 915 | 3614335 | 3615249 | lpqC_1 | BCGDan_3361 | 915 | 3614400 | 3615299 | lpqC | BCGDan_sapMKO_3362 | 900 |
| 3683051 | 3685963 | atsB | Rv3299c | 2913 | 3635459 | 3638371 | atsB | BCG_3328c | 2913 | 3615276 | 3618188 | atsB_1 | BCGDan_3362 | 2913 | 3615341 | 3618250 | atsB | BCGDan_sapMKO_3363 | 2910 |
| 3685983 | 3686900 | Rv3300c | Rv3300c | 918 | 3638391 | 3639308 | BCG_3329c | BCG_3329c | 918 | 3618208 | 3619125 | BCGDan_3363 | BCGDan_3363 | 918 | 3618273 | 3619157 | BCGDan_sapMKO_3364 | BCGDan_sapMKO_3364 | 885 |
| 3686912 | 3687577 | phoY1 | Rv3301c | 666 | 3639320 | 3639985 | phoY1_1 | BCG_3330c | 666 | 3619137 | 3619802 | phoY1_1 | BCGDan_3364 | 666 | 3619202 | 3619867 | phoY1 | BCGDan_sapMKO_3365 | 666 |
| 3687685 | 3689442 | glpD2 | Rv3302c | 1758 | 3640093 | 3641850 | glpD2_1 | BCG_3331c | 1758 | 3619910 | 3621667 | glpD2_1 | BCGDan_3365 | 1758 | 3619975 | 3621732 | glpD2 | BCGDan_sapMKO_3366 | 1758 |
| 3689457 | 3690938 | lpdA | Rv3303c | 1482 | 3641931 | 3642536 | 'lpdA | BCG_3332c | 606 | 3621748 | 3623163 | lpdA_1 | BCGDan_3366 | 1416 | 3621813 | 3623228 | lpdA | BCGDan_sapMKO_3367 | 1416 |
| 3691141 | 3691620 | Rv3304 | Rv3304 | 480 | 3679712 | 3680191 | BCG_3369 | BCG_3369 | 480 | 3623366 | 3623845 | BCGDan_3367 | BCGDan_3367 | 480 | 3623431 | 3623910 | BCGDan_sapMKO_3368 | BCGDan_sapMKO_3368 | 480 |
| 3691639 | 3692808 | amiA1 | Rv3305c | 1170 | 3680210 | 3681379 | amiA1 | BCG_3370c | 1170 | 3623864 | 3625033 | amiA1_1 | BCGDan_3368 | 1170 | 3623929 | 3625098 | amiA1 | BCGDan_sapMKO_3369 | 1170 |
| 3692805 | 3693989 | amiB1 | Rv3306c | 1185 | 3681376 | 3682560 | amiB1 | BCG_3371c | 1185 | 3625030 | 3626214 | amiB1_1 | BCGDan_3369 | 1185 | 3625095 | 3626279 | amiB1 | BCGDan_sapMKO_3370 | 1185 |
| 3694054 | 3694860 | deoD | Rv3307 | 807 | 3682625 | 3683431 | deoD | BCG_3372 | 807 | 3626279 | 3627085 | deoD_1 | BCGDan_3370 | 807 | 3626344 | 3627150 | deoD | BCGDan_sapMKO_3371 | 807 |
| 3694864 | 3696468 | pmmB | Rv3308 | 1605 | 3683435 | 3685039 | pmmB | BCG_3373 | 1605 | 3627089 | 3628693 | pmmB_1 | BCGDan_3371 | 1605 | 3627154 | 3628758 | pmmB | BCGDan_sapMKO_3372 | 1605 |
| 3696470 | 3697093 | upp | Rv3309c | 624 | 3685041 | 3685664 | upp | BCG_3374c | 624 | 3628695 | 3629318 | upp_1 | BCGDan_3372 | 624 | 3628760 | 3629383 | upp | BCGDan_sapMKO_3373 | 624 |
| 3697198 | 3698097 | sapM | Rv3310 | 900 | 3685769 | 3686668 | sapM | BCG_3375 | 900 | 3629423 | 3630322 | sapM_1 | BCGDan_3373 | 900 | 3629488 | 3629775 | truncated sapM | BCGDan_sapMKO_3374 | 288 |
| 3698121 | 3699383 | Rv3311 | Rv3311 | 1263 | 3686692 | 3687954 | BCG_3376 | BCG_3376 | 1263 | 3630346 | 3631608 | BCGDan_3374 | BCGDan_3374 | 1263 | 3629803 | 3631065 | BCGDan_sapMKO_3375 | BCGDan_sapMKO_3375 | 1263 |
| 3699404 | 3700330 | Rv3312c | Rv3312c | 927 | 3687975 | 3688901 | BCG_3377c | BCG_3377c | 927 | 3631629 | 3632555 | BCGDan_3375 | BCGDan_3375 | 927 | 3631086 | 3632018 | BCGDan_sapMKO_3376 | BCGDan_sapMKO_3376 | 933 |
| 3700705 | 3701016 | Rv3312A | Rv3312A | 312 | 3689276 | 3689587 | BCG_3378c | BCG_3378c | 312 | 3632930 | 3633241 | BCGDan_3376 | BCGDan_3376 | 312 | 3632387 | 3632698 | BCGDan_sapMKO_3377 | BCGDan_sapMKO_3377 | 312 |
| 3701087 | 3702184 | add | Rv3313c | 1098 | 3689658 | 3690755 | add | BCG_3379c | 1098 | 3633312 | 3634409 | add_1 | BCGDan_3377 | 1098 | 3632769 | 3633866 | add | BCGDan_sapMKO_3378 | 1098 |
| 3702184 | 3703467 | deoA | Rv3314c | 1284 | 3690755 | 3692038 | deoA | BCG_3380c | 1284 | 3634409 | 3635692 | deoA_1 | BCGDan_3378 | 1284 | 3633866 | 3635149 | deoA | BCGDan_sapMKO_3379 | 1284 |
| 3703464 | 3703865 | cdd | Rv3315c | 402 | 3692035 | 3692436 | cdd | BCG_3381c | 402 | 3635689 | 3636090 | cdd_1 | BCGDan_3379 | 402 | 3635146 | 3635547 | cdd | BCGDan_sapMKO_3380 | 402 |
| 3704102 | 3704440 | sdhC | Rv3316 | 339 | 3692673 | 3693011 | sdhC | BCG_3382 | 339 | 3636327 | 3636665 | sdhC_1 | BCGDan_3380 | 339 | 3635784 | 3636122 | sdhC | BCGDan_sapMKO_3381 | 339 |
| 3704437 | 3704871 | sdhD | Rv3317 | 435 | 3693008 | 3693442 | sdhD | BCG_3383 | 435 | 3636662 | 3637096 | sdhD_1 | BCGDan_3381 | 435 | 3636119 | 3636553 | sdhD | BCGDan_sapMKO_3382 | 435 |
| 3705000 | 3706772 | sdhA | Rv3318 | 1773 | 3693571 | 3695343 | sdhA | BCG_3384 | 1773 | 3637225 | 3638997 | sdhA_1 | BCGDan_3382 | 1773 | 3636706 | 3638454 | sdhA | BCGDan_sapMKO_3383 | 1749 |
| 3706772 | 3707563 | sdhB | Rv3319 | 792 | 3695343 | 3696134 | sdhB | BCG_3385 | 792 | 3638997 | 3639788 | sdhB_1 | BCGDan_3383 | 792 | 3638454 | 3639245 | sdhB | BCGDan_sapMKO_3384 | 792 |
| 3707642 | 3708070 | vapC44 | Rv3320c | 429 | 3696213 | 3696641 | BCG_3386c | BCG_3386c | 429 | 3639867 | 3640295 | BCGDan_3384 | BCGDan_3384 | 429 | 3639324 | 3639752 | BCGDan_sapMKO_3385 | BCGDan_sapMKO_3385 | 429 |
| 3708074 | 3708316 | vapB44 | Rv3321c | 243 | 3696645 | 3696887 | BCG_3387c | BCG_3387c | 243 | 3640299 | 3640541 | BCGDan_3385 | BCGDan_3385 | 243 | 3639756 | 3639998 | BCGDan_sapMKO_3386 | BCGDan_sapMKO_3386 | 243 |
| 3708438 | 3709052 | Rv3322c | Rv3322c | 615 | 3697009 | 3697623 | BCG_3388c | BCG_3388c | 615 | 3640663 | 3641277 | BCGDan_3386 | BCGDan_3386 | 615 | 3640120 | 3640734 | BCGDan_sapMKO_3387 | BCGDan_sapMKO_3387 | 615 |
| NA | NA | NA | NA | NA | NA | NA | NA | NA | NA | 3642265 | 3642543 | whiB7_2 | BCGDan_3387 | 279 | NA | NA | NA | NA | NA |
| NA | NA | NA | NA | NA | NA | NA | NA | NA | NA | 3642973 | 3645075 | UvrD2_2 | BCGDan_3388 | 2103 | NA | NA | NA | NA | NA |
| NA | NA | NA | NA | NA | NA | NA | NA | NA | NA | 3645199 | 3645453 | BCGDan_3389 | BCGDan_3389 | 255 | NA | NA | NA | NA | NA |
| NA | NA | NA | NA | NA | NA | NA | NA | NA | NA | 3645466 | 3646407 | nudC_2 | BCGDan_3390 | 942 | NA | NA | NA | NA | NA |
| NA | NA | NA | NA | NA | NA | NA | NA | NA | NA | 3646466 | 3647533 | BCGDan_3391 | BCGDan_3391 | 1068 | NA | NA | NA | NA | NA |
| NA | NA | NA | NA | NA | NA | NA | NA | NA | NA | 3647595 | 3650900 | BCGDan_3392 | BCGDan_3392 | 3306 | NA | NA | NA | NA | NA |

| | | | | | | | | | | | | | | | | | | | |
|---|---|---|---|---|---|---|---|---|---|---|---|---|---|---|---|---|---|---|---|
| NA | NA | NA | NA | NA | NA | NA | NA | NA | NA | 3650897 | 3654064 | BCGDan_3393 | BCGDan_3393 | 3168 | NA | NA | NA | NA | NA |
| NA | NA | NA | NA | NA | NA | NA | NA | NA | NA | 3654141 | 3654359 | BCGDan_3394 | BCGDan_3394 | 219 | NA | NA | NA | NA | NA |
| NA | NA | NA | NA | NA | NA | NA | NA | NA | NA | 3654501 | 3655175 | lipV_2 | BCGDan_3395 | 675 | NA | NA | NA | NA | NA |
| NA | NA | NA | NA | NA | NA | NA | NA | NA | NA | 3655178 | 3655483 | BCGDan_3396 | BCGDan_3396 | 306 | NA | NA | NA | NA | NA |
| NA | NA | NA | NA | NA | NA | NA | NA | NA | NA | 3655490 | 3656368 | BCGDan_3397 | BCGDan_3397 | 879 | NA | NA | NA | NA | NA |
| NA | NA | NA | NA | NA | NA | NA | NA | NA | NA | 3656395 | 3657573 | moeB1_2 | BCGDan_3398 | 1179 | NA | NA | NA | NA | NA |
| NA | NA | NA | NA | NA | NA | NA | NA | NA | NA | 3657664 | 3658521 | BCGDan_3399 | BCGDan_3399 | 858 | NA | NA | NA | NA | NA |
| NA | NA | NA | NA | NA | NA | NA | NA | NA | NA | 3658867 | 3659553 | BCGDan_3400 | BCGDan_3400 | 687 | NA | NA | NA | NA | NA |
| NA | NA | NA | NA | NA | NA | NA | NA | NA | NA | 3659540 | 3659812 | TB9.4_2 | BCGDan_3401 | 273 | NA | NA | NA | NA | NA |
| NA | NA | NA | NA | NA | NA | NA | NA | NA | NA | 3660137 | 3660697 | BCGDan_3402 | BCGDan_3402 | 561 | NA | NA | NA | NA | NA |
| NA | NA | NA | NA | NA | NA | NA | NA | NA | NA | 3660707 | 3661402 | BCGDan_3403 | BCGDan_3403 | 696 | NA | NA | NA | NA | NA |
| NA | NA | NA | NA | NA | NA | NA | NA | NA | NA | 3661661 | 3663244 | rhlE_2 | BCGDan_3404 | 1584 | NA | NA | NA | NA | NA |
| NA | NA | NA | NA | NA | NA | NA | NA | NA | NA | 3663257 | 3664480 | BCGDan_3405 | BCGDan_3405 | 1224 | NA | NA | NA | NA | NA |
| NA | NA | NA | NA | NA | 3642530 | 3643126 | BCG_3333c | BCG_3333c | 597 | 3664556 | 3665356 | BCGDan_3406 | BCGDan_3406 | 801 | NA | NA | NA | NA | NA |
| NA | NA | NA | NA | NA | 3643280 | 3643891 | gpm2_2 | BCG_3334 | 612 | 3665510 | 3666121 | gpm2_2 | BCGDan_3407 | 612 | NA | NA | NA | NA | NA |
| NA | NA | NA | NA | NA | 3643888 | 3645006 | entC_2 | BCG_3335 | 1119 | 3666118 | 3667236 | entC_2 | BCGDan_3408 | 1119 | NA | NA | NA | NA | NA |
| NA | NA | NA | NA | NA | 3645154 | 3645486 | BCG_3336 | BCG_3336 | 333 | 3667384 | 3667716 | BCGDan_3409 | BCGDan_3409 | 333 | NA | NA | NA | NA | NA |
| NA | NA | NA | NA | NA | 3645438 | 3645869 | BCG_3337c | BCG_3337c | 432 | 3667668 | 3668099 | BCGDan_3410 | BCGDan_3410 | 432 | NA | NA | NA | NA | NA |
| NA | NA | NA | NA | NA | 3646102 | 3647067 | BCG_3338 | BCG_3338 | 966 | 3668332 | 3669297 | BCGDan_3411 | BCGDan_3411 | 966 | NA | NA | NA | NA | NA |
| NA | NA | NA | NA | NA | 3647347 | 3647601 | whiB1_2 | BCG_3339 | 255 | 3669577 | 3669831 | whiB1_2 | BCGDan_3412 | 255 | NA | NA | NA | NA | NA |
| NA | NA | NA | NA | NA | 3647663 | 3649168 | BCG_3340c | BCG_3340c | 1506 | 3669893 | 3671398 | BCGDan_3413 | BCGDan_3413 | 1506 | NA | NA | NA | NA | NA |
| NA | NA | NA | NA | NA | 3649185 | 3649400 | TB7.3_2 | BCG_3341c | 216 | 3671415 | 3671630 | TB7.3_2 | BCGDan_3414 | 216 | NA | NA | NA | NA | NA |
| NA | NA | NA | NA | NA | 3649685 | 3649990 | BCG_3342c | BCG_3342c | 306 | 3671915 | 3672220 | BCGDan_3415 | BCGDan_3415 | 306 | NA | NA | NA | NA | NA |
| NA | NA | NA | NA | NA | 3649987 | 3650538 | BCG_3343c | BCG_3343c | 552 | 3672217 | 3672768 | BCGDan_3416 | BCGDan_3416 | 552 | NA | NA | NA | NA | NA |
| NA | NA | NA | NA | NA | 3650535 | 3651185 | sigH_2 | BCG_3344c | 651 | 3672765 | 3673415 | sigH_2 | BCGDan_3417 | 651 | NA | NA | NA | NA | NA |
| NA | NA | NA | NA | NA | 3651485 | 3652333 | BCG_3345 | BCG_3345 | 849 | 3673715 | 3674563 | BCGDan_3418 | BCGDan_3418 | 849 | NA | NA | NA | NA | NA |
| NA | NA | NA | NA | NA | 3652269 | 3652457 | BCG_3346 | BCG_3346 | 189 | 3674499 | 3674687 | BCGDan_3419 | BCGDan_3419 | 189 | NA | NA | NA | NA | NA |
| NA | NA | NA | NA | NA | 3652435 | 3652653 | BCG_3347 | BCG_3347 | 219 | 3674665 | 3674883 | BCGDan_3420 | BCGDan_3420 | 219 | NA | NA | NA | NA | NA |
| NA | NA | NA | NA | NA | 3652650 | 3654074 | BCG_3348c | BCG_3348c | 1425 | 3674880 | 3676304 | BCGDan_3421 | BCGDan_3421 | 1425 | NA | NA | NA | NA | NA |
| NA | NA | NA | NA | NA | 3654198 | 3654956 | BCG_3349c | BCG_3349c | 759 | 3676428 | 3677186 | BCGDan_3422 | BCGDan_3422 | 759 | NA | NA | NA | NA | NA |
| NA | NA | NA | NA | NA | 3655011 | 3656363 | aroA_2 | BCG_3350 | 1353 | 3677241 | 3678593 | aroA_2 | BCGDan_3423 | 1353 | NA | NA | NA | NA | NA |
| NA | NA | NA | NA | NA | 3656360 | 3657352 | BCG_3351 | BCG_3351 | 993 | 3678590 | 3679582 | BCGDan_3424 | BCGDan_3424 | 993 | NA | NA | NA | NA | NA |
| NA | NA | NA | NA | NA | 3657385 | 3658668 | desA3_2 | BCG_3352c | 1284 | 3679615 | 3680898 | desA3_2 | BCGDan_3425 | 1284 | NA | NA | NA | NA | NA |
| NA | NA | NA | NA | NA | 3658746 | 3659888 | BCG_3353c | BCG_3353c | 1143 | 3680976 | 3682118 | BCGDan_3426 | BCGDan_3426 | 1143 | NA | NA | NA | NA | NA |
| NA | NA | NA | NA | NA | 3659943 | 3660365 | lat' | BCG_3354c | 423 | 3682173 | 3682595 | lat | BCGDan_3427 | 423 | NA | NA | NA | NA | NA |
| NA | NA | NA | NA | NA | 3659998 | 3660108 | BCG_3355c | BCG_3355c | 111 | 3682228 | 3682338 | BCGDan_3428 | BCGDan_3428 | 111 | NA | NA | NA | NA | NA |
| NA | NA | NA | NA | NA | 3660416 | 3660868 | BCG_3356c | BCG_3356c | 453 | 3682646 | 3683098 | BCGDan_3429 | BCGDan_3429 | 453 | NA | NA | NA | NA | NA |
| NA | NA | NA | NA | NA | 3660899 | 3662146 | BCG_3357 | BCG_3357 | 1248 | 3683129 | 3684376 | BCGDan_3430 | BCGDan_3430 | 1248 | NA | NA | NA | NA | NA |
| NA | NA | NA | NA | NA | 3662173 | 3663657 | pcd_2 | BCG_3358 | 1485 | 3684403 | 3685887 | pcd_2 | BCGDan_3431 | 1485 | NA | NA | NA | NA | NA |
| NA | NA | NA | NA | NA | 3663757 | 3664566 | BCG_3359c | BCG_3359c | 810 | 3685987 | 3686796 | BCGDan_3432 | BCGDan_3432 | 810 | NA | NA | NA | NA | NA |
| NA | NA | NA | NA | NA | 3664637 | 3665302 | BCG_3360 | BCG_3360 | 666 | 3686867 | 3687532 | BCGDan_3433 | BCGDan_3433 | 666 | NA | NA | NA | NA | NA |
| NA | NA | NA | NA | NA | 3665346 | 3669887 | lhr_2 | BCG_3361 | 4542 | 3687576 | 3692117 | lhr_2 | BCGDan_3434 | 4542 | NA | NA | NA | NA | NA |
| NA | NA | NA | NA | NA | 3669891 | 3670658 | nei_2 | BCG_3362 | 768 | 3692121 | 3692888 | nei_2 | BCGDan_3435 | 768 | NA | NA | NA | NA | NA |
| NA | NA | NA | NA | NA | 3670681 | 3671595 | lpqC_2 | BCG_3363c | 915 | 3692911 | 3693825 | lpqC_2 | BCGDan_3436 | 915 | NA | NA | NA | NA | NA |
| NA | NA | NA | NA | NA | 3671622 | 3674534 | atsB_2 | BCG_3364c | 2913 | 3693852 | 3696764 | atsB_2 | BCGDan_3437 | 2913 | NA | NA | NA | NA | NA |
| NA | NA | NA | NA | NA | 3674554 | 3675471 | BCG_3365c | BCG_3365c | 918 | 3696784 | 3697701 | BCGDan_3438 | BCGDan_3438 | 918 | NA | NA | NA | NA | NA |
| NA | NA | NA | NA | NA | 3675483 | 3676148 | phoY1_2 | BCG_3366c | 666 | 3697713 | 3698378 | phoY1_2 | BCGDan_3439 | 666 | NA | NA | NA | NA | NA |
| NA | NA | NA | NA | NA | 3676256 | 3678013 | glpD2_2 | BCG_3367c | 1758 | 3698486 | 3700243 | glpD2_2 | BCGDan_3440 | 1758 | NA | NA | NA | NA | NA |
| NA | NA | NA | NA | NA | 3678094 | 3679509 | lpdA | BCG_3368c | 1416 | 3700324 | 3701739 | lpdA_2 | BCGDan_3441 | 1416 | NA | NA | NA | NA | NA |
| NA | NA | NA | NA | NA | NA | NA | NA | NA | NA | 3701942 | 3702421 | BCGDan_3442 | BCGDan_3442 | 480 | NA | NA | NA | NA | NA |
| NA | NA | NA | NA | NA | NA | NA | NA | NA | NA | 3702440 | 3703609 | amiA1_2 | BCGDan_3443 | 1170 | NA | NA | NA | NA | NA |
| NA | NA | NA | NA | NA | NA | NA | NA | NA | NA | 3703606 | 3704790 | amiB1_2 | BCGDan_3444 | 1185 | NA | NA | NA | NA | NA |
| NA | NA | NA | NA | NA | NA | NA | NA | NA | NA | 3704855 | 3705661 | deoD_2 | BCGDan_3445 | 807 | NA | NA | NA | NA | NA |
| NA | NA | NA | NA | NA | NA | NA | NA | NA | NA | 3705665 | 3707269 | pmmB_2 | BCGDan_3446 | 1605 | NA | NA | NA | NA | NA |
| NA | NA | NA | NA | NA | NA | NA | NA | NA | NA | 3707271 | 3707894 | upp_2 | BCGDan_3447 | 624 | NA | NA | NA | NA | NA |
| NA | NA | NA | NA | NA | NA | NA | NA | NA | NA | 3707999 | 3708898 | sapM_2 | BCGDan_3448 | 900 | NA | NA | NA | NA | NA |
| NA | NA | NA | NA | NA | NA | NA | NA | NA | NA | 3708922 | 3710184 | BCGDan_3449 | BCGDan_3449 | 1263 | NA | NA | NA | NA | NA |
| NA | NA | NA | NA | NA | NA | NA | NA | NA | NA | 3710205 | 3711131 | BCGDan_3450 | BCGDan_3450 | 927 | NA | NA | NA | NA | NA |
| NA | NA | NA | NA | NA | NA | NA | NA | NA | NA | 3711506 | 3711817 | BCGDan_3451 | BCGDan_3451 | 312 | NA | NA | NA | NA | NA |
| NA | NA | NA | NA | NA | NA | NA | NA | NA | NA | 3711888 | 3712985 | add_2 | BCGDan_3452 | 1098 | NA | NA | NA | NA | NA |
| NA | NA | NA | NA | NA | NA | NA | NA | NA | NA | 3712985 | 3714268 | deoA_2 | BCGDan_3453 | 1284 | NA | NA | NA | NA | NA |
| NA | NA | NA | NA | NA | NA | NA | NA | NA | NA | 3714265 | 3714666 | cdd_2 | BCGDan_3454 | 402 | NA | NA | NA | NA | NA |
| NA | NA | NA | NA | NA | NA | NA | NA | NA | NA | 3714903 | 3715241 | sdhC_2 | BCGDan_3455 | 339 | NA | NA | NA | NA | NA |
| NA | NA | NA | NA | NA | NA | NA | NA | NA | NA | 3715238 | 3715672 | sdhD_2 | BCGDan_3456 | 435 | NA | NA | NA | NA | NA |
| NA | NA | NA | NA | NA | NA | NA | NA | NA | NA | 3715801 | 3717573 | sdhA_2 | BCGDan_3457 | 1773 | NA | NA | NA | NA | NA |
| NA | NA | NA | NA | NA | NA | NA | NA | NA | NA | 3717573 | 3718364 | sdhB_2 | BCGDan_3458 | 792 | NA | NA | NA | NA | NA |
| NA | NA | NA | NA | NA | NA | NA | NA | NA | NA | 3718443 | 3718871 | BCGDan_3459 | BCGDan_3459 | 429 | NA | NA | NA | NA | NA |
| NA | NA | NA | NA | NA | NA | NA | NA | NA | NA | 3718875 | 3719117 | BCGDan_3460 | BCGDan_3460 | 243 | NA | NA | NA | NA | NA |
| NA | NA | NA | NA | NA | NA | NA | NA | NA | NA | 3719239 | 3719853 | BCGDan_3461 | BCGDan_3461 | 615 | NA | NA | NA | NA | NA |
| 3709049 | 3709714 | moaX | Rv3323c | 666 | 3697620 | 3698285 | moaX | BCG_3389c | 666 | 3719850 | 3720515 | moaX | BCGDan_3462 | 666 | 3640731 | 3641396 | moaX | BCGDan_sapMKO_3388 | 666 |
| 3709715 | 3710248 | moaC3 | Rv3324c | 534 | 3698286 | 3698819 | moaC3 | BCG_3390c | 534 | 3720516 | 3721049 | moaC3 | BCGDan_3463 | 534 | 3641397 | 3641930 | moaC3 | BCGDan_sapMKO_3389 | 534 |
| 3710245 | 3710379 | Rv3324A | Rv3324A | 135 | 3698816 | 3699190 | moaB3 | BCG_3391c | 375 | 3721046 | 3721420 | moaB3 | BCGDan_3464 | 375 | 3641927 | 3642253 | moaB3 | BCGDan_sapMKO_3390 | 327 |
| NA | NA | NA | NA | NA | 3699287 | 3700351 | moaA3 | BCG_3392c | 1065 | 3721517 | 3722581 | moaA3 | BCGDan_3465 | 1065 | 3642398 | 3643534 | moaA3 | BCGDan_sapMKO_3391 | 1137 |
| NA | NA | NA | NA | NA | 3700406 | 3700756 | BCG_3393 | BCG_3393 | 351 | 3722636 | 3722986 | BCGDan_3466 | BCGDan_3466 | 351 | 3643517 | 3643867 | BCGDan_sapMKO_3392 | BCGDan_sapMKO_3392 | 351 |
| NA | NA | NA | NA | NA | 3700734 | 3701120 | BCG_3394 | BCG_3394 | 387 | 3722964 | 3723350 | BCGDan_3467 | BCGDan_3467 | 387 | 3643845 | 3644231 | BCGDan_sapMKO_3393 | BCGDan_sapMKO_3393 | 387 |
| NA | NA | NA | NA | NA | 3700996 | 3702141 | embR2 | BCG_3395 | 1146 | 3723226 | 3724371 | embR2 | BCGDan_3468 | 1146 | 3644107 | 3645252 | embR2 | BCGDan_sapMKO_3394 | 1146 |
| NA | NA | NA | NA | NA | 3702236 | 3702880 | BCG_3396c | BCG_3396c | 645 | 3724466 | 3725110 | BCGDan_3469 | BCGDan_3469 | 645 | 3645347 | 3645991 | BCGDan_sapMKO_3395 | BCGDan_sapMKO_3395 | 645 |
| 3711749 | 3713461 | Rv3327 | Rv3327 | 1713 | 3703047 | 3704759 | BCG_3397 | BCG_3397 | 1713 | 3725277 | 3726989 | BCGDan_3470 | BCGDan_3470 | 1713 | 3646140 | 3647870 | BCGDan_sapMKO_3396 | BCGDan_sapMKO_3396 | 1731 |
| 3713394 | 3714332 | sigJ | Rv3328c | 939 | 3704692 | 3705630 | sigJ | BCG_3398c | 939 | 3726922 | 3727860 | sigJ | BCGDan_3471 | 939 | 3647803 | 3648750 | sigJ | BCGDan_sapMKO_3397 | 948 |
| 3714392 | 3715708 | Rv3329 | Rv3329 | 1317 | 3705690 | 3707006 | BCG_3399 | BCG_3399 | 1317 | 3727920 | 3729236 | BCGDan_3472 | BCGDan_3472 | 1317 | 3648801 | 3650117 | BCGDan_sapMKO_3398 | BCGDan_sapMKO_3398 | 1317 |
| 3715777 | 3716994 | dacB1 | Rv3330 | 1218 | 3707075 | 3708292 | dacB1 | BCG_3400 | 1218 | 3729305 | 3730522 | dacB1 | BCGDan_3473 | 1218 | 3650225 | 3651403 | dacB1 | BCGDan_sapMKO_3399 | 1179 |
| 3717090 | 3718598 | sugI | Rv3331 | 1509 | 3708388 | 3709881 | sugI | BCG_3401 | 1494 | 3730618 | 3732126 | sugI | BCGDan_3474 | 1509 | 3651499 | 3653007 | sugI | BCGDan_sapMKO_3400 | 1509 |
| 3718595 | 3719746 | nagA | Rv3332 | 1152 | 3709878 | 3711029 | nagA | BCG_3402 | 1152 | 3732123 | 3733274 | nagA | BCGDan_3475 | 1152 | 3653004 | 3654155 | nagA | BCGDan_sapMKO_3401 | 1152 |
| 3719937 | 3720782 | Rv3333c | Rv3333c | 846 | 3711220 | 3712065 | BCG_3403c | BCG_3403c | 846 | 3733465 | 3734310 | BCGDan_3476 | BCGDan_3476 | 846 | 3654346 | 3655191 | BCGDan_sapMKO_3402 | BCGDan_sapMKO_3402 | 846 |
| NA | NA | NA | NA | NA | 3712210 | 3712521 | BCG_3404 | BCG_3404 | 312 | 3734455 | 3734766 | BCGDan_3477 | BCGDan_3477 | 312 | 3655336 | 3655647 | BCGDan_sapMKO_3403 | BCGDan_sapMKO_3403 | 312 |
| 3721257 | 3721697 | Rv3334 | Rv3334 | 441 | 3712540 | 3712980 | BCG_3405 | BCG_3405 | 441 | 3734785 | 3735225 | BCGDan_3478 | BCGDan_3478 | 441 | 3655666 | 3656106 | BCGDan_sapMKO_3404 | BCGDan_sapMKO_3404 | 441 |

| | | | | | | | | | | | | | | | | | | | |
|---|---|---|---|---|---|---|---|---|---|---|---|---|---|---|---|---|---|---|---|
| 3721731 | 3722600 | Rv3335c | Rv3335c | 870 | 3713014 | 3713883 | BCG_3406c | BCG_3406c | 870 | 3735259 | 3736128 | BCGDan_3479 | BCGDan_3479 | 870 | 3656140 | 3657009 | BCGDan_sapMKO_3405 | BCGDan_sapMKO_3405 | 870 |
| 3722621 | 3723631 | trpS | Rv3336c | 1011 | 3713904 | 3714914 | trpS | BCG_3407c | 1011 | 3736149 | 3737159 | trpS | BCGDan_3480 | 1011 | 3657030 | 3657962 | trpS | BCGDan_sapMKO_3406 | 933 |
| 3723656 | 3724042 | Rv3337 | Rv3337 | 387 | 3714939 | 3715832 | BCG_3408 | BCG_3408 | 894 | 3737184 | 3738077 | BCGDan_3481 | BCGDan_3481 | 894 | 3658155 | 3658958 | BCGDan_sapMKO_3407 | BCGDan_sapMKO_3407 | 804 |
| 3724615 | 3725844 | icd1 | Rv3339c | 1230 | 3715899 | 3717128 | icd1 | BCG_3409c | 1230 | 3738144 | 3739373 | icd1 | BCGDan_3482 | 1230 | 3659025 | 3660254 | icd1 | BCGDan_sapMKO_3408 | 1230 |
| 3726127 | 3727476 | metC | Rv3340 | 1350 | 3717411 | 3718760 | metC | BCG_3410 | 1350 | 3739656 | 3741005 | metC | BCGDan_3483 | 1350 | 3660537 | 3661886 | metC | BCGDan_sapMKO_3409 | 1350 |
| 3727488 | 3728627 | metA | Rv3341 | 1140 | 3718772 | 3719911 | metA | BCG_3411 | 1140 | 3741017 | 3742156 | metA | BCGDan_3484 | 1140 | 3661898 | 3663037 | metA | BCGDan_sapMKO_3410 | 1140 |
| 3728624 | 3729355 | Rv3342 | Rv3342 | 732 | 3719908 | 3720639 | BCG_3412 | BCG_3412 | 732 | 3742153 | 3742884 | BCGDan_3485 | BCGDan_3485 | 732 | 3663025 | 3663765 | BCGDan_sapMKO_3411 | BCGDan_sapMKO_3411 | 741 |
| 3729364 | 3736935 | PPE54 | Rv3343c | 7572 | 3720648 | 3724664 | PPE54 | BCG_3413c | 4017 | 3742893 | 3746909 | PPE54 | BCGDan_3486 | 4017 | 3663774 | 3667775 | PPE54 | BCGDan_sapMKO_3412 | 4002 |
| 3736984 | 3738438 | PE_PGRS49 | Rv3344c | 1455 | 3724713 | 3730343 | PE_PGRS50 | BCG_3414c | 5631 | 3746958 | 3752588 | PE_PGRS50 | BCGDan_3487 | 5631 | 3667839 | 3673469 | PE_PGRS50 | BCGDan_sapMKO_3413 | 5631 |
| 3743198 | 3743455 | Rv3346c | Rv3346c | 258 | 3730767 | 3731024 | BCG_3415c | BCG_3415c | 258 | 3753012 | 3753269 | BCGDan_3488 | BCGDan_3488 | 258 | 3673893 | 3674150 | BCGDan_sapMKO_3414 | BCGDan_sapMKO_3414 | 258 |
| NA | NA | NA | NA | NA | 3731280 | 3734465 | PPE55b | BCG_3416c | 3186 | 3753525 | 3756710 | PPE55b | BCGDan_3489 | 3186 | 3674409 | 3677591 | PPE55b | BCGDan_sapMKO_3415 | 3183 |
| 3743711 | 3753184 | PPE55 | Rv3347c | 9474 | 3734462 | 3740752 | PPE55a | BCG_3417c | 6291 | 3756707 | 3762997 | PPE55a | BCGDan_3490 | 6291 | 3677588 | 3683842 | PPE55a | BCGDan_sapMKO_3416 | 6255 |
| 3753765 | 3754256 | Rv3348 | Rv3348 | 492 | 3741332 | 3741823 | BCG_3418 | BCG_3418 | 492 | 3763577 | 3764068 | BCGDan_3491 | BCGDan_3491 | 492 | 3684458 | 3684949 | BCGDan_sapMKO_3417 | BCGDan_sapMKO_3417 | 492 |
| 3754293 | 3755033 | Rv3349c | Rv3349c | 741 | 3741860 | 3742600 | BCG_3419c | BCG_3419c | 741 | 3764105 | 3764845 | BCGDan_3492 | BCGDan_3492 | 741 | 3684986 | 3685804 | BCGDan_sapMKO_3418 | BCGDan_sapMKO_3418 | 819 |
| NA | NA | NA | NA | NA | 3743514 | 3747173 | PPE56d | BCG_3420c | 3660 | 3765759 | 3769418 | PPE56d | BCGDan_3493 | 3660 | 3686640 | 3690299 | PPE56d | BCGDan_sapMKO_3419 | 3660 |
| NA | NA | NA | NA | NA | 3747167 | 3753301 | PPE56b | BCG_3421c | 6135 | 3769412 | 3775546 | PPE56b | BCGDan_3494 | 6135 | 3690293 | 3696427 | PPE56b | BCGDan_sapMKO_3420 | 6135 |
| 3755952 | 3767102 | PPE56 | Rv3350c | 11151 | 3753359 | 3754657 | PPE56a | BCG_3422c | 1299 | 3775604 | 3776902 | PPE56a | BCGDan_3495 | 1299 | 3696485 | 3697783 | PPE56a | BCGDan_sapMKO_3421 | 1299 |
| 3767346 | 3768140 | Rv3351c | Rv3351c | 795 | 3754901 | 3755695 | BCG_3423c | BCG_3423c | 795 | 3777146 | 3777940 | BCGDan_3496 | BCGDan_3496 | 795 | 3698027 | 3698821 | BCGDan_sapMKO_3422 | BCGDan_sapMKO_3422 | 795 |
| 3768222 | 3768593 | Rv3352c | Rv3352c | 372 | 3755777 | 3756148 | BCG_3424c | BCG_3424c | 372 | 3778022 | 3778393 | BCGDan_3497 | BCGDan_3497 | 372 | 3698903 | 3699274 | BCGDan_sapMKO_3423 | BCGDan_sapMKO_3423 | 372 |
| 3768736 | 3768996 | Rv3353c | Rv3353c | 261 | 3756291 | 3756551 | BCG_3425c | BCG_3425c | 261 | 3778536 | 3778796 | BCGDan_3498 | BCGDan_3498 | 261 | 3699417 | 3699677 | BCGDan_sapMKO_3424 | BCGDan_sapMKO_3424 | 261 |
| 3769111 | 3769500 | Rv3354 | Rv3354 | 390 | 3756666 | 3757055 | BCG_3426 | BCG_3426 | 390 | 3778911 | 3779300 | BCGDan_3499 | BCGDan_3499 | 390 | 3699792 | 3700181 | BCGDan_sapMKO_3425 | BCGDan_sapMKO_3425 | 390 |
| 3769514 | 3769807 | Rv3355c | Rv3355c | 294 | 3757069 | 3757362 | BCG_3427c | BCG_3427c | 294 | 3779314 | 3779607 | BCGDan_3500 | BCGDan_3500 | 294 | 3700195 | 3700488 | BCGDan_sapMKO_3426 | BCGDan_sapMKO_3426 | 294 |
| 3769804 | 3770649 | folD | Rv3356c | 846 | 3757359 | 3758204 | folD | BCG_3428c | 846 | 3779604 | 3780449 | folD | BCGDan_3501 | 846 | 3700485 | 3701330 | folD | BCGDan_sapMKO_3427 | 846 |
| 3770773 | 3771048 | relJ | Rv3357 | 276 | 3758328 | 3758603 | BCG_3429 | BCG_3429 | 276 | 3780573 | 3780848 | BCGDan_3502 | BCGDan_3502 | 276 | 3701454 | 3701729 | BCGDan_sapMKO_3428 | BCGDan_sapMKO_3428 | 276 |
| 3771045 | 3771302 | relK | Rv3358 | 258 | 3758600 | 3758857 | BCG_3430 | BCG_3430 | 258 | 3780845 | 3781102 | BCGDan_3503 | BCGDan_3503 | 258 | 3701726 | 3701983 | BCGDan_sapMKO_3429 | BCGDan_sapMKO_3429 | 258 |
| 3771344 | 3772534 | Rv3359 | Rv3359 | 1191 | 3758899 | 3760089 | BCG_3431 | BCG_3431 | 1191 | 3781144 | 3782334 | BCGDan_3504 | BCGDan_3504 | 1191 | 3702025 | 3703215 | BCGDan_sapMKO_3430 | BCGDan_sapMKO_3430 | 1191 |
| 3772651 | 3773019 | Rv3360 | Rv3360 | 369 | 3760206 | 3760574 | BCG_3432 | BCG_3432 | 369 | 3782451 | 3782819 | BCGDan_3505 | BCGDan_3505 | 369 | 3703332 | 3703700 | BCGDan_sapMKO_3431 | BCGDan_sapMKO_3431 | 369 |
| 3773016 | 3773567 | Rv3361c | Rv3361c | 552 | 3760571 | 3761122 | BCG_3433c | BCG_3433c | 552 | 3782816 | 3783367 | BCGDan_3506 | BCGDan_3506 | 552 | 3703697 | 3704248 | BCGDan_sapMKO_3432 | BCGDan_sapMKO_3432 | 552 |
| 3773574 | 3774155 | Rv3362c | Rv3362c | 582 | 3761129 | 3761710 | BCG_3434c | BCG_3434c | 582 | 3783374 | 3783955 | BCGDan_3507 | BCGDan_3507 | 582 | 3704255 | 3704836 | BCGDan_sapMKO_3433 | BCGDan_sapMKO_3433 | 582 |
| 3774136 | 3774504 | Rv3363c | Rv3363c | 369 | 3761691 | 3762059 | BCG_3435c | BCG_3435c | 369 | 3783936 | 3784304 | BCGDan_3508 | BCGDan_3508 | 369 | 3704817 | 3705185 | BCGDan_sapMKO_3434 | BCGDan_sapMKO_3434 | 369 |
| 3774482 | 3774874 | Rv3364c | Rv3364c | 393 | 3762037 | 3762429 | BCG_3436c | BCG_3436c | 393 | 3784282 | 3784674 | BCGDan_3509 | BCGDan_3509 | 393 | 3705163 | 3705555 | BCGDan_sapMKO_3435 | BCGDan_sapMKO_3435 | 393 |
| 3774871 | 3777501 | Rv3365c | Rv3365c | 2631 | 3762426 | 3765056 | BCG_3437c | BCG_3437c | 2631 | 3784671 | 3787301 | BCGDan_3510 | BCGDan_3510 | 2631 | 3705552 | 3708182 | BCGDan_sapMKO_3436 | BCGDan_sapMKO_3436 | 2631 |
| 3777737 | 3778201 | spoU | Rv3366 | 465 | 3765292 | 3765756 | spoU | BCG_3438 | 465 | 3787537 | 3788001 | spoU | BCGDan_3511 | 465 | 3708418 | 3708882 | spoU | BCGDan_sapMKO_3437 | 465 |
| 3778568 | 3780334 | PE_PGRS51 | Rv3367 | 1767 | 3766123 | 3767898 | PE_PGRS51 | BCG_3439 | 1776 | 3788368 | 3790143 | PE_PGRS51 | BCGDan_3512 | 1776 | 3709249 | 3711024 | PE_PGRS51 | BCGDan_sapMKO_3438 | 1776 |
| 3780335 | 3780979 | Rv3368c | Rv3368c | 645 | 3767899 | 3768543 | BCG_3440c | BCG_3440c | 645 | 3790144 | 3790788 | BCGDan_3513 | BCGDan_3513 | 645 | 3711025 | 3711669 | BCGDan_sapMKO_3439 | BCGDan_sapMKO_3439 | 645 |
| 3780978 | 3781412 | Rv3369 | Rv3369 | 435 | 3768542 | 3768976 | BCG_3441 | BCG_3441 | 435 | 3790787 | 3791221 | BCGDan_3514 | BCGDan_3514 | 435 | 3711668 | 3712102 | BCGDan_sapMKO_3440 | BCGDan_sapMKO_3440 | 435 |
| 3781501 | 3784740 | dnaE2 | Rv3370c | 3240 | 3769065 | 3772304 | dnaE2 | BCG_3442c | 3240 | 3791310 | 3794549 | dnaE2 | BCGDan_3515 | 3240 | 3712191 | 3715466 | dnaE2 | BCGDan_sapMKO_3441 | 3276 |
| 3784932 | 3786272 | Rv3371 | Rv3371 | 1341 | 3772496 | 3773836 | BCG_3443 | BCG_3443 | 1341 | 3794741 | 3796081 | BCGDan_3516 | BCGDan_3516 | 1341 | 3715622 | 3716962 | BCGDan_sapMKO_3442 | BCGDan_sapMKO_3442 | 1341 |
| 3786314 | 3787489 | otsB2 | Rv3372 | 1176 | 3773878 | 3775053 | otsB2 | BCG_3444 | 1176 | 3796123 | 3797298 | otsB2 | BCGDan_3517 | 1176 | 3717004 | 3718179 | otsB2 | BCGDan_sapMKO_3443 | 1176 |
| 3787726 | 3788367 | echA18 | Rv3373 | 642 | 3775290 | 3776180 | echA18 | BCG_3445 | 891 | 3797535 | 3798425 | echA18 | BCGDan_3518 | 891 | 3718431 | 3719306 | echA18 | BCGDan_sapMKO_3444 | 876 |
| 3788621 | 3790048 | amiD | Rv3375 | 1428 | 3776185 | 3777612 | amiD | BCG_3446 | 1428 | 3798430 | 3799857 | amiD | BCGDan_3519 | 1428 | 3719311 | 3720738 | amiD | BCGDan_sapMKO_3445 | 1428 |
| 3790156 | 3790809 | Rv3376 | Rv3376 | 654 | 3777720 | 3778373 | BCG_3447 | BCG_3447 | 654 | 3799965 | 3800618 | BCGDan_3520 | BCGDan_3520 | 654 | 3720882 | 3721499 | BCGDan_sapMKO_3446 | BCGDan_sapMKO_3446 | 618 |
| 3790848 | 3792353 | Rv3377c | Rv3377c | 1506 | 3778465 | 3779916 | BCG_3448c | BCG_3448c | 1452 | 3800710 | 3802161 | BCGDan_3521 | BCGDan_3521 | 1452 | 3721591 | 3723042 | BCGDan_sapMKO_3447 | BCGDan_sapMKO_3447 | 1452 |
| 3792358 | 3793248 | Rv3378c | Rv3378c | 891 | 3779921 | 3780811 | BCG_3449c | BCG_3449c | 891 | 3802166 | 3803056 | BCGDan_3522 | BCGDan_3522 | 891 | 3723047 | 3723937 | BCGDan_sapMKO_3448 | BCGDan_sapMKO_3448 | 891 |
| 3793257 | 3794867 | dxs2 | Rv3379c | 1611 | 3780820 | 3782430 | dxs2 | BCG_3450c | 1611 | 3803065 | 3804903 | dxs2 | BCGDan_3523 | 1839 | 3723946 | 3725784 | dxs2 | BCGDan_sapMKO_3449 | 1839 |
| 3796448 | 3797437 | lytB1 | Rv3382c | 990 | 3782655 | 3783644 | lytB1 | BCG_3451c | 990 | 3804900 | 3805889 | lytB1 | BCGDan_3524 | 990 | 3725781 | 3726776 | lytB1 | BCGDan_sapMKO_3450 | 996 |
| 3797437 | 3798489 | idsB | Rv3383c | 1053 | 3783644 | 3784696 | idsB | BCG_3452c | 1053 | 3805889 | 3806941 | idsB | BCGDan_3525 | 1053 | 3726770 | 3727723 | idsB | BCGDan_sapMKO_3451 | 954 |
| 3799243 | 3799635 | vapC46 | Rv3384c | 393 | 3785450 | 3785842 | BCG_3453c | BCG_3453c | 393 | 3807695 | 3808087 | BCGDan_3526 | BCGDan_3526 | 393 | 3728576 | 3728968 | BCGDan_sapMKO_3452 | BCGDan_sapMKO_3452 | 393 |
| 3799635 | 3799943 | vapB46 | Rv3385c | 309 | 3785842 | 3786150 | BCG_3454c | BCG_3454c | 309 | 3808087 | 3808362 | BCGDan_3527 | BCGDan_3527 | 276 | 3728968 | 3729243 | BCGDan_sapMKO_3453 | BCGDan_sapMKO_3453 | 276 |
| 3800092 | 3800796 | Rv3386 | Rv3386 | 705 | 3786299 | 3787003 | BCG_3455 | BCG_3455 | 705 | 3808544 | 3809248 | BCGDan_3528 | BCGDan_3528 | 705 | 3729350 | 3730129 | BCGDan_sapMKO_3454 | BCGDan_sapMKO_3454 | 780 |
| 3800786 | 3801463 | Rv3387 | Rv3387 | 678 | 3786993 | 3787670 | BCG_3456 | BCG_3456 | 678 | 3809238 | 3809915 | BCGDan_3529 | BCGDan_3529 | 678 | 3730056 | 3730796 | BCGDan_sapMKO_3455 | BCGDan_sapMKO_3455 | 741 |
| 3801653 | 3803848 | PE_PGRS52 | Rv3388 | 2196 | 3787860 | 3790112 | PE_PGRS52 | BCG_3457 | 2253 | 3810105 | 3812357 | PE_PGRS52 | BCGDan_3530 | 2253 | 3730986 | 3733238 | PE_PGRS52 | BCGDan_sapMKO_3456 | 2253 |
| 3803919 | 3804791 | htdY | Rv3389c | 873 | 3790183 | 3791055 | BCG_3458c | BCG_3458c | 873 | 3812428 | 3813300 | BCGDan_3531 | BCGDan_3531 | 873 | 3733309 | 3734151 | BCGDan_sapMKO_3457 | BCGDan_sapMKO_3457 | 843 |
| 3804865 | 3805575 | lpqD | Rv3390 | 711 | 3791129 | 3791839 | lpqD | BCG_3459 | 711 | 3813374 | 3814084 | lpqD | BCGDan_3532 | 711 | 3734255 | 3734965 | lpqD | BCGDan_sapMKO_3458 | 711 |
| 3805621 | 3807573 | acrA1 | Rv3391 | 1953 | 3791885 | 3793837 | acrA1 | BCG_3460 | 1953 | 3814130 | 3816082 | acrA1 | BCGDan_3533 | 1953 | 3735011 | 3736963 | acrA1 | BCGDan_sapMKO_3459 | 1953 |
| 3807574 | 3808437 | cmaA1 | Rv3392c | 864 | 3793838 | 3794701 | cmaA1 | BCG_3461c | 864 | 3816083 | 3816946 | cmaA1 | BCGDan_3534 | 864 | 3736964 | 3737827 | cmaA1 | BCGDan_sapMKO_3460 | 864 |
| 3808461 | 3809387 | iunH | Rv3393 | 927 | 3794725 | 3795651 | iunH | BCG_3462 | 927 | 3816970 | 3817896 | iunH | BCGDan_3535 | 927 | 3737851 | 3738777 | iunH | BCGDan_sapMKO_3461 | 927 |
| 3809442 | 3811025 | Rv3394c | Rv3394c | 1584 | 3795706 | 3797289 | BCG_3463c | BCG_3463c | 1584 | 3817951 | 3819534 | BCGDan_3536 | BCGDan_3536 | 1584 | 3738832 | 3740412 | BCGDan_sapMKO_3462 | BCGDan_sapMKO_3462 | 1581 |
| 3811022 | 3811636 | Rv3395c | Rv3395c | 615 | 3797286 | 3797900 | BCG_3464c | BCG_3464c | 615 | 3819531 | 3820145 | BCGDan_3537 | BCGDan_3537 | 615 | 3740412 | 3741026 | BCGDan_sapMKO_3463 | BCGDan_sapMKO_3463 | 615 |
| 3811719 | 3812345 | Rv3395A | Rv3395A | 627 | 3797983 | 3798609 | BCG_3465 | BCG_3465 | 627 | 3820228 | 3820854 | BCGDan_3538 | BCGDan_3538 | 627 | 3741025 | 3741735 | BCGDan_sapMKO_3464 | BCGDan_sapMKO_3464 | 711 |
| 3812501 | 3814078 | guaA | Rv3396c | 1578 | 3798765 | 3800342 | guaA | BCG_3466c | 1578 | 3821010 | 3822587 | guaA | BCGDan_3539 | 1578 | 3741891 | 3743420 | guaA | BCGDan_sapMKO_3465 | 1530 |
| 3814090 | 3814998 | phyA | Rv3397c | 909 | 3800354 | 3801262 | phyA | BCG_3467c | 909 | 3822599 | 3823507 | phyA | BCGDan_3540 | 909 | 3743480 | 3744400 | phyA | BCGDan_sapMKO_3466 | 921 |
| 3815027 | 3816106 | idsA1 | Rv3398c | 1080 | 3801291 | 3802370 | idsA1 | BCG_3468c | 1080 | 3823536 | 3824615 | idsA1 | BCGDan_3541 | 1080 | 3744417 | 3745496 | idsA1 | BCGDan_sapMKO_3467 | 1080 |
| 3816129 | 3817175 | Rv3399 | Rv3399 | 1047 | 3802393 | 3803439 | BCG_3469 | BCG_3469 | 1047 | 3824638 | 3825684 | BCGDan_3542 | BCGDan_3542 | 1047 | 3745645 | 3746565 | BCGDan_sapMKO_3468 | BCGDan_sapMKO_3468 | 921 |
| 3817239 | 3818027 | Rv3400 | Rv3400 | 789 | 3803503 | 3804291 | BCG_3470 | BCG_3470 | 789 | 3825748 | 3826536 | BCGDan_3543 | BCGDan_3543 | 789 | 3746674 | 3747417 | BCGDan_sapMKO_3469 | BCGDan_sapMKO_3469 | 744 |
| 3818042 | 3820402 | Rv3401 | Rv3401 | 2361 | 3804306 | 3806666 | BCG_3471 | BCG_3471 | 2361 | 3826551 | 3828911 | BCGDan_3544 | BCGDan_3544 | 2361 | 3747432 | 3749792 | BCGDan_sapMKO_3470 | BCGDan_sapMKO_3470 | 2361 |
| 3820653 | 3821891 | Rv3402c | Rv3402c | 1239 | NA | NA | NA | NA | NA | 3829400 | 3829555 | BCGDan_3545 | BCGDan_3545 | 156 | NA | NA | NA | NA | NA |
| NA | NA | NA | NA | NA | 3807341 | 3808393 | BCG_3472c | BCG_3472c | 1053 | 3829586 | 3830638 | BCGDan_3546 | BCGDan_3546 | 1053 | 3750467 | 3751519 | BCGDan_sapMKO_3471 | BCGDan_sapMKO_3471 | 1053 |
| 3822262 | 3823863 | Rv3403c | Rv3403c | 1602 | 3808764 | 3810365 | BCG_3473c | BCG_3473c | 1602 | 3831009 | 3832610 | BCGDan_3547 | BCGDan_3547 | 1602 | 3751890 | 3753491 | BCGDan_sapMKO_3472 | BCGDan_sapMKO_3472 | 1602 |
| 3823880 | 3824584 | Rv3404c | Rv3404c | 705 | 3810382 | 3811086 | BCG_3474c | BCG_3474c | 705 | 3832627 | 3833331 | BCGDan_3548 | BCGDan_3548 | 705 | 3753508 | 3754212 | BCGDan_sapMKO_3473 | BCGDan_sapMKO_3473 | 705 |
| 3824702 | 3825268 | Rv3405c | Rv3405c | 567 | 3811204 | 3811770 | BCG_3475c | BCG_3475c | 567 | 3833449 | 3834015 | BCGDan_3549 | BCGDan_3549 | 567 | 3754330 | 3754896 | BCGDan_sapMKO_3474 | BCGDan_sapMKO_3474 | 567 |
| 3825330 | 3826217 | Rv3406 | Rv3406 | 888 | 3811832 | 3812719 | BCG_3476 | BCG_3476 | 888 | 3834077 | 3834964 | BCGDan_3550 | BCGDan_3550 | 888 | 3754958 | 3755845 | BCGDan_sapMKO_3475 | BCGDan_sapMKO_3475 | 888 |
| 3826252 | 3826551 | vapB47 | Rv3407 | 300 | 3812754 | 3813053 | BCG_3477 | BCG_3477 | 300 | 3834999 | 3835298 | BCGDan_3551 | BCGDan_3551 | 300 | 3755880 | 3756179 | BCGDan_sapMKO_3476 | BCGDan_sapMKO_3476 | 300 |
| 3826548 | 3826958 | vapC47 | Rv3408 | 411 | 3813050 | 3813460 | BCG_3478 | BCG_3478 | 411 | 3835295 | 3835705 | BCGDan_3552 | BCGDan_3552 | 411 | 3756176 | 3756586 | BCGDan_sapMKO_3477 | BCGDan_sapMKO_3477 | 411 |
| 3826991 | 3828727 | choD | Rv3409c | 1737 | 3813493 | 3815229 | choD | BCG_3479c | 1737 | 3835738 | 3837474 | choD | BCGDan_3553 | 1737 | 3756619 | 3758355 | choD | BCGDan_sapMKO_3478 | 1737 |
| 3828783 | 3829910 | guaB3 | Rv3410c | 1128 | 3815285 | 3816412 | guaB3 | BCG_3480c | 1128 | 3837530 | 3838657 | guaB3 | BCGDan_3554 | 1128 | 3758411 | 3759538 | guaB3 | BCGDan_sapMKO_3479 | 1128 |
| 3829930 | 3831519 | guaB2 | Rv3411c | 1590 | 3816432 | 3818021 | guaB2 | BCG_3481c | 1590 | 3838677 | 3840266 | guaB2 | BCGDan_3555 | 1590 | 3759558 | 3761135 | guaB2 | BCGDan_sapMKO_3480 | 1578 |
| 3831726 | 3832136 | Rv3412 | Rv3412 | 411 | 3818228 | 3818638 | BCG_3482 | BCG_3482 | 411 | 3840473 | 3840883 | BCGDan_3556 | BCGDan_3556 | 411 | 3761354 | 3761764 | BCGDan_sapMKO_3481 | BCGDan_sapMKO_3481 | 411 |
| 3832146 | 3833045 | Rv3413c | Rv3413c | 900 | 3818648 | 3819547 | BCG_3483c | BCG_3483c | 900 | 3840893 | 3841792 | BCGDan_3557 | BCGDan_3557 | 900 | 3761774 | 3762673 | BCGDan_sapMKO_3482 | BCGDan_sapMKO_3482 | 900 |
| 3833038 | 3833676 | sigD | Rv3414c | 639 | 3819540 | 3820178 | sigD | BCG_3484c | 639 | 3841785 | 3842423 | sigD | BCGDan_3558 | 639 | 3762666 | 3763235 | sigD | BCGDan_sapMKO_3483 | 570 |
| 3833694 | 3834521 | Rv3415c | Rv3415c | 828 | 3820196 | 3821023 | BCG_3485c | BCG_3485c | 828 | 3842441 | 3843268 | BCGDan_3559 | BCGDan_3559 | 828 | 3763322 | 3764149 | BCGDan_sapMKO_3484 | BCGDan_sapMKO_3484 | 828 |
| 3834892 | 3835200 | whiB3 | Rv3416 | 309 | 3821394 | 3821702 | whiB3 | BCG_3486 | 309 | 3843639 | 3843947 | whiB3 | BCGDan_3560 | 309 | 3764520 | 3764828 | whiB3 | BCGDan_sapMKO_3485 | 309 |
| 3835272 | 3836891 | groEL1 | Rv3417c | 1620 | 3821774 | 3823393 | groEL1 | BCG_3487c | 1620 | 3844019 | 3845638 | groEL1 | BCGDan_3561 | 1620 | 3764900 | 3766519 | groEL1 | BCGDan_sapMKO_3486 | 1620 |
| 3836986 | 3837288 | groES | Rv3418c | 303 | 3823488 | 3823790 | groES | BCG_3488c | 303 | 3845733 | 3846035 | groES | BCGDan_3562 | 303 | 3766614 | 3766916 | groES | BCGDan_sapMKO_3487 | 303 |
| 3837555 | 3838589 | gcp | Rv3419c | 1035 | 3824057 | 3825091 | gcp | BCG_3489c | 1035 | 3846302 | 3847336 | gcp | BCGDan_3563 | 1035 | 3767183 | 3768217 | gcp | BCGDan_sapMKO_3488 | 1035 |
| 3838586 | 3839062 | rimI | Rv3420c | 477 | 3825088 | 3825564 | rimI | BCG_3490c | 477 | 3847333 | 3847809 | rimI | BCGDan_3564 | 477 | 3768214 | 3768690 | rimI | BCGDan_sapMKO_3489 | 477 |

| | | | | | | | | | | | | | | | | | | | |
|---|---|---|---|---|---|---|---|---|---|---|---|---|---|---|---|---|---|---|---|
| 3839059 | 3839694 | Rv3421c | Rv3421c | 636 | 3825561 | 3826196 | BCG_3491c | BCG_3491c | 636 | 3847806 | 3848441 | BCGDan_3565 | BCGDan_3565 | 636 | 3768687 | 3769322 | BCGDan_sapMKO_3490 | BCGDan_sapMKO_3490 | 636 |
| 3839691 | 3840197 | Rv3422c | Rv3422c | 507 | 3826193 | 3826699 | BCG_3492c | BCG_3492c | 507 | 3848438 | 3848944 | BCGDan_3566 | BCGDan_3566 | 507 | 3769319 | 3769825 | BCGDan_sapMKO_3491 | BCGDan_sapMKO_3491 | 507 |
| 3840194 | 3841420 | alr | Rv3423c | 1227 | 3826696 | 3827922 | alr | BCG_3493c | 1227 | 3848941 | 3850167 | alr | BCGDan_3567 | 1227 | 3769822 | 3770934 | alr | BCGDan_sapMKO_3492 | 1113 |
| 3841714 | 3842076 | Rv3424c | Rv3424c | 363 | 3828216 | 3828578 | BCG_3494c | BCG_3494c | 363 | 3850461 | 3850823 | BCGDan_3568 | BCGDan_3568 | 363 | 3771342 | 3771704 | BCGDan_sapMKO_3493 | BCGDan_sapMKO_3493 | 363 |
| 3842239 | 3842769 | PPE57 | Rv3425 | 531 | 3828741 | 3829277 | PPE57 | BCG_3495 | 537 | 3850986 | 3851522 | PPE57 | BCGDan_3569 | 537 | 3771876 | 3772403 | PPE57 | BCGDan_sapMKO_3494 | 528 |
| 3847642 | 3848805 | Rv3430c | Rv3430c | 1164 | 3829218 | 3830381 | BCG_3496c | BCG_3496c | 1164 | 3851463 | 3852626 | BCGDan_3570 | BCGDan_3570 | 1164 | 3772344 | 3773507 | BCGDan_sapMKO_3495 | BCGDan_sapMKO_3495 | 1164 |
| 3849294 | 3850139 | Rv3431c | Rv3431c | 846 | 3830870 | 3831715 | BCG_3497c | BCG_3497c | 846 | 3853115 | 3853960 | BCGDan_3571 | BCGDan_3571 | 846 | 3773996 | 3774841 | BCGDan_sapMKO_3496 | BCGDan_sapMKO_3496 | 846 |
| 3850372 | 3851754 | gadB | Rv3432c | 1383 | 3831948 | 3833330 | gadB | BCG_3498c | 1383 | 3854193 | 3855575 | gadB | BCGDan_3572 | 1383 | 3775074 | 3776396 | gadB | BCGDan_sapMKO_3497 | 1323 |
| 3851792 | 3853213 | Rv3433c | Rv3433c | 1422 | 3833368 | 3834786 | BCG_3499c | BCG_3499c | 1419 | 3855613 | 3857034 | BCGDan_3573 | BCGDan_3573 | 1422 | 3776494 | 3777915 | BCGDan_sapMKO_3498 | BCGDan_sapMKO_3498 | 1422 |
| 3853215 | 3853928 | Rv3434c | Rv3434c | 714 | 3834788 | 3835501 | BCG_3500c | BCG_3500c | 714 | 3857036 | 3857749 | BCGDan_3574 | BCGDan_3574 | 714 | 3777917 | 3778630 | BCGDan_sapMKO_3499 | BCGDan_sapMKO_3499 | 714 |
| 3853939 | 3854793 | Rv3435c | Rv3435c | 855 | 3835512 | 3836366 | BCG_3501c | BCG_3501c | 855 | 3857760 | 3858614 | BCGDan_3575 | BCGDan_3575 | 855 | 3778641 | 3779579 | BCGDan_sapMKO_3500 | BCGDan_sapMKO_3500 | 939 |
| 3855015 | 3856889 | glmS | Rv3436c | 1875 | 3836588 | 3838462 | glmS | BCG_3502c | 1875 | 3858836 | 3860710 | glmS | BCGDan_3576 | 1875 | 3779717 | 3781540 | glmS | BCGDan_sapMKO_3501 | 1824 |
| 3856911 | 3857387 | Rv3437 | Rv3437 | 477 | 3838484 | 3838960 | BCG_3503 | BCG_3503 | 477 | 3860732 | 3861208 | BCGDan_3577 | BCGDan_3577 | 477 | 3781613 | 3782089 | BCGDan_sapMKO_3502 | BCGDan_sapMKO_3502 | 477 |
| 3857397 | 3858239 | Rv3438 | Rv3438 | 843 | 3838970 | 3839812 | BCG_3504 | BCG_3504 | 843 | 3861218 | 3862060 | BCGDan_3578 | BCGDan_3578 | 843 | 3782099 | 3782941 | BCGDan_sapMKO_3503 | BCGDan_sapMKO_3503 | 843 |
| 3858259 | 3859662 | Rv3439c | Rv3439c | 1404 | 3839832 | 3841235 | BCG_3505c | BCG_3505c | 1404 | 3862080 | 3863483 | BCGDan_3579 | BCGDan_3579 | 1404 | 3782961 | 3784391 | BCGDan_sapMKO_3504 | BCGDan_sapMKO_3504 | 1431 |
| 3859665 | 3859976 | Rv3440c | Rv3440c | 312 | 3841238 | 3841549 | BCG_3506c | BCG_3506c | 312 | 3863486 | 3863797 | BCGDan_3580 | BCGDan_3580 | 312 | 3784367 | 3784678 | BCGDan_sapMKO_3505 | BCGDan_sapMKO_3505 | 312 |
| 3860024 | 3861370 | mrsA | Rv3441c | 1347 | 3841597 | 3842943 | mrsA | BCG_3507c | 1347 | 3863845 | 3865191 | mrsA | BCGDan_3581 | 1347 | 3784726 | 3786072 | mrsA | BCGDan_sapMKO_3506 | 1347 |
| 3861495 | 3861950 | rpsI | Rv3442c | 456 | 3843068 | 3843523 | rpsI | BCG_3508c | 456 | 3865316 | 3865771 | rpsI | BCGDan_3582 | 456 | 3786197 | 3786652 | rpsI | BCGDan_sapMKO_3507 | 456 |
| 3861947 | 3862390 | rplM | Rv3443c | 444 | 3843520 | 3843963 | rplM | BCG_3509c | 444 | 3865768 | 3866211 | rplM | BCGDan_3583 | 444 | 3786649 | 3787092 | rplM | BCGDan_sapMKO_3508 | 444 |
| 3862624 | 3862926 | esxT | Rv3444c | 303 | 3844196 | 3844498 | esxT | BCG_3510c | 303 | 3866444 | 3866746 | esxT | BCGDan_3584 | 303 | 3787325 | 3787627 | esxT | BCGDan_sapMKO_3509 | 303 |
| 3862947 | 3863264 | esxU | Rv3445c | 318 | 3844519 | 3844896 | esxU | BCG_3511c | 378 | 3866767 | 3867144 | esxU | BCGDan_3585 | 378 | 3787648 | 3788025 | esxU | BCGDan_sapMKO_3510 | 378 |
| 3863317 | 3864531 | Rv3446c | Rv3446c | 1215 | 3844889 | 3846103 | BCG_3512c | BCG_3512c | 1215 | 3867137 | 3868351 | BCGDan_3586 | BCGDan_3586 | 1215 | 3788018 | 3789232 | BCGDan_sapMKO_3511 | BCGDan_sapMKO_3511 | 1215 |
| 3864528 | 3868238 | eccC4 | Rv3447c | 3711 | 3846100 | 3849810 | BCG_3513c | BCG_3513c | 3711 | 3868348 | 3872058 | BCGDan_3587 | BCGDan_3587 | 3711 | 3789229 | 3792939 | BCGDan_sapMKO_3512 | BCGDan_sapMKO_3512 | 3711 |
| 3868352 | 3869755 | eccD4 | Rv3448 | 1404 | 3849924 | 3851327 | BCG_3514 | BCG_3514 | 1404 | 3872172 | 3873575 | BCGDan_3588 | BCGDan_3588 | 1404 | 3793053 | 3794456 | BCGDan_sapMKO_3513 | BCGDan_sapMKO_3513 | 1404 |
| 3869752 | 3871119 | mycP4 | Rv3449 | 1368 | 3851324 | 3852691 | BCG_3515 | BCG_3515 | 1368 | 3873572 | 3874939 | BCGDan_3589 | BCGDan_3589 | 1368 | 3794447 | 3795820 | BCGDan_sapMKO_3514 | BCGDan_sapMKO_3514 | 1374 |
| 3871084 | 3872496 | eccB4 | Rv3450c | 1413 | 3852656 | 3854068 | BCG_3516c | BCG_3516c | 1413 | 3874904 | 3876316 | BCGDan_3590 | BCGDan_3590 | 1413 | 3795785 | 3797197 | BCGDan_sapMKO_3515 | BCGDan_sapMKO_3515 | 1413 |
| 3872617 | 3873405 | cut3 | Rv3451 | 789 | 3854189 | 3854974 | cut3 | BCG_3517 | 786 | 3876437 | 3877225 | cut3 | BCGDan_3591 | 789 | 3797318 | 3798106 | cut3 | BCGDan_sapMKO_3516 | 789 |
| 3873452 | 3874132 | cut4 | Rv3452 | 681 | 3855021 | 3855701 | cut4 | BCG_3518 | 681 | 3877272 | 3877952 | cut4 | BCGDan_3592 | 681 | 3798138 | 3798833 | cut4 | BCGDan_sapMKO_3517 | 696 |
| 3874404 | 3874736 | Rv3453 | Rv3453 | 333 | 3855973 | 3857658 | BCG_3519 | BCG_3519 | 1686 | 3878224 | 3879909 | BCGDan_3593 | BCGDan_3593 | 1686 | 3799105 | 3800790 | BCGDan_sapMKO_3518 | BCGDan_sapMKO_3518 | 1686 |
| 3876052 | 3876822 | truA | Rv3455c | 771 | 3857620 | 3858390 | truA | BCG_3520c | 771 | 3879871 | 3880641 | truA | BCGDan_3594 | 771 | 3800752 | 3801522 | truA | BCGDan_sapMKO_3519 | 771 |
| 3876890 | 3877432 | rplQ | Rv3456c | 543 | 3858458 | 3859000 | rplQ | BCG_3521c | 543 | 3880709 | 3881251 | rplQ | BCGDan_3595 | 543 | 3801590 | 3802132 | rplQ | BCGDan_sapMKO_3520 | 543 |
| 3877464 | 3878507 | rpoA | Rv3457c | 1044 | 3859032 | 3860075 | rpoA | BCG_3522c | 1044 | 3881283 | 3882326 | rpoA | BCGDan_3596 | 1044 | 3802164 | 3803207 | rpoA | BCGDan_sapMKO_3521 | 1044 |
| 3878659 | 3879264 | rpsD | Rv3458c | 606 | 3860226 | 3860831 | rpsD | BCG_3523c | 606 | 3882477 | 3883082 | rpsD | BCGDan_3597 | 606 | 3803358 | 3803963 | rpsD | BCGDan_sapMKO_3522 | 606 |
| 3879273 | 3879692 | rpsK | Rv3459c | 420 | 3860840 | 3861259 | rpsK | BCG_3524c | 420 | 3883091 | 3883510 | rpsK | BCGDan_3598 | 420 | 3803972 | 3804391 | rpsK | BCGDan_sapMKO_3523 | 420 |
| 3879696 | 3880070 | rpsM | Rv3460c | 375 | 3861263 | 3861637 | rpsM | BCG_3525c | 375 | 3883514 | 3883888 | rpsM | BCGDan_3599 | 375 | 3804395 | 3804727 | rpsM | BCGDan_sapMKO_3524 | 333 |
| 3880286 | 3880399 | rpmJ | Rv3461c | 114 | 3861853 | 3861966 | rpmJ | BCG_3526c | 114 | 3884104 | 3884217 | rpmJ | BCGDan_3600 | 114 | 3804985 | 3805098 | rpmJ | BCGDan_sapMKO_3525 | 114 |
| 3880432 | 3880653 | infA | Rv3462c | 222 | 3861999 | 3862220 | infA | BCG_3527c | 222 | 3884250 | 3884471 | infA | BCGDan_3601 | 222 | 3805131 | 3805352 | infA | BCGDan_sapMKO_3526 | 222 |
| 3880907 | 3881764 | Rv3463 | Rv3463 | 858 | 3862474 | 3863331 | BCG_3528 | BCG_3528 | 858 | 3884725 | 3885582 | BCGDan_3602 | BCGDan_3602 | 858 | 3805606 | 3806463 | BCGDan_sapMKO_3527 | BCGDan_sapMKO_3527 | 858 |
| 3881837 | 3882832 | rmlB | Rv3464 | 996 | 3863404 | 3864399 | rmlB1 | BCG_3529 | 996 | 3885655 | 3886650 | rmlB1 | BCGDan_3603 | 996 | 3806536 | 3807531 | rmlB1 | BCGDan_sapMKO_3528 | 996 |
| 3882834 | 3883442 | rmlC | Rv3465 | 609 | 3864401 | 3865009 | rmlC | BCG_3530 | 609 | 3886652 | 3887260 | rmlC | BCGDan_3604 | 609 | 3807533 | 3808141 | rmlC | BCGDan_sapMKO_3529 | 609 |
| 3883525 | 3884193 | Rv3466 | Rv3466 | 669 | 3865092 | 3865760 | BCG_3531 | BCG_3531 | 669 | 3887343 | 3888011 | BCGDan_3605 | BCGDan_3605 | 669 | 3808224 | 3808892 | BCGDan_sapMKO_3530 | BCGDan_sapMKO_3530 | 669 |
| 3883964 | 3884917 | Rv3467 | Rv3467 | 954 | 3865531 | 3866484 | BCG_3532 | BCG_3532 | 954 | 3887782 | 3888735 | BCGDan_3606 | BCGDan_3606 | 954 | 3808657 | 3809616 | BCGDan_sapMKO_3531 | BCGDan_sapMKO_3531 | 960 |
| 3884975 | 3886069 | Rv3468c | Rv3468c | 1095 | 3866542 | 3867636 | rmlB2 | BCG_3533c | 1095 | 3888793 | 3889887 | rmlB2 | BCGDan_3607 | 1095 | 3809674 | 3810768 | rmlB2 | BCGDan_sapMKO_3532 | 1095 |
| 3886073 | 3887083 | mhpE | Rv3469c | 1011 | 3867640 | 3868650 | mhpE | BCG_3534c | 1011 | 3889891 | 3890901 | mhpE | BCGDan_3608 | 1011 | 3810772 | 3811782 | mhpE | BCGDan_sapMKO_3533 | 1011 |
| 3887144 | 3888802 | ilvB2 | Rv3470c | 1659 | 3868711 | 3870369 | ilvB2 | BCG_3535c | 1659 | 3890962 | 3892620 | ilvB2 | BCGDan_3609 | 1659 | 3811843 | 3813516 | ilvB2 | BCGDan_sapMKO_3534 | 1674 |
| 3888808 | 3889341 | Rv3471c | Rv3471c | 534 | 3870375 | 3870908 | BCG_3536c | BCG_3536c | 534 | 3892626 | 3893159 | BCGDan_3610 | BCGDan_3610 | 534 | 3813507 | 3813992 | BCGDan_sapMKO_3535 | BCGDan_sapMKO_3535 | 486 |
| 3889362 | 3889868 | Rv3472 | Rv3472 | 507 | 3870929 | 3871435 | BCG_3537 | BCG_3537 | 507 | 3893180 | 3893686 | BCGDan_3611 | BCGDan_3611 | 507 | 3814061 | 3814567 | BCGDan_sapMKO_3536 | BCGDan_sapMKO_3536 | 507 |
| 3889948 | 3890733 | bpoA | Rv3473c | 786 | 3871515 | 3872300 | bpoA | BCG_3538c | 786 | 3893766 | 3894551 | bpoA | BCGDan_3612 | 786 | 3814647 | 3815549 | bpoA | BCGDan_sapMKO_3537 | 903 |
| 3892371 | 3893720 | kgtP | Rv3476c | 1350 | 3872580 | 3873929 | kgtP | BCG_3539c | 1350 | 3894831 | 3896180 | kgtP | BCGDan_3613 | 1350 | 3815712 | 3817127 | kgtP | BCGDan_sapMKO_3538 | 1416 |
| 3894093 | 3894389 | PE31 | Rv3477 | 297 | 3874302 | 3874598 | PE31 | BCG_3540 | 297 | 3896553 | 3896849 | PE31 | BCGDan_3614 | 297 | 3817434 | 3817730 | PE31 | BCGDan_sapMKO_3539 | 297 |
| 3894426 | 3895607 | PPE60 | Rv3478 | 1182 | 3874635 | 3875816 | PPE60 | BCG_3541 | 1182 | 3896886 | 3898067 | PPE60 | BCGDan_3615 | 1182 | 3817767 | 3818948 | PPE60 | BCGDan_sapMKO_3540 | 1182 |
| 3895820 | 3898885 | Rv3479 | Rv3479 | 3066 | 3876029 | 3877330 | BCG_3542 | BCG_3542 | 1302 | 3898280 | 3899581 | BCGDan_3616 | BCGDan_3616 | 1302 | 3819068 | 3820462 | BCGDan_sapMKO_3541 | BCGDan_sapMKO_3541 | 1395 |
| NA | NA | NA | NA | NA | 3877335 | 3878381 | BCG_3543 | BCG_3543 | 1047 | 3899586 | 3900632 | BCGDan_3617 | BCGDan_3617 | 1047 | 3820467 | 3821513 | BCGDan_sapMKO_3542 | BCGDan_sapMKO_3542 | 1047 |
| 3898909 | 3900402 | Rv3480c | Rv3480c | 1494 | 3878405 | 3879898 | BCG_3544c | BCG_3544c | 1494 | 3900656 | 3902149 | BCGDan_3618 | BCGDan_3618 | 1494 | 3821537 | 3823030 | BCGDan_sapMKO_3543 | BCGDan_sapMKO_3543 | 1494 |
| 3900493 | 3901182 | Rv3481c | Rv3481c | 690 | 3879989 | 3880678 | BCG_3545c | BCG_3545c | 690 | 3902240 | 3902929 | BCGDan_3619 | BCGDan_3619 | 690 | 3823121 | 3823867 | BCGDan_sapMKO_3544 | BCGDan_sapMKO_3544 | 747 |
| 3901324 | 3902106 | Rv3482c | Rv3482c | 783 | 3880113 | 3881603 | BCG_3546c | BCG_3546c | 1491 | 3902364 | 3903854 | BCGDan_3620 | BCGDan_3620 | 1491 | 3823245 | 3824735 | BCGDan_sapMKO_3545 | BCGDan_sapMKO_3545 | 1491 |
| 3902150 | 3902812 | Rv3483c | Rv3483c | 663 | 3881647 | 3882309 | BCG_3547c | BCG_3547c | 663 | 3903898 | 3904560 | BCGDan_3621 | BCGDan_3621 | 663 | 3824779 | 3825441 | BCGDan_sapMKO_3546 | BCGDan_sapMKO_3546 | 663 |
| 3903078 | 3904616 | cpsA | Rv3484 | 1539 | 3882575 | 3884113 | cpsA | BCG_3548 | 1539 | 3904826 | 3906364 | cpsA | BCGDan_3622 | 1539 | 3825791 | 3827245 | cpsA | BCGDan_sapMKO_3547 | 1455 |
| 3904622 | 3905566 | Rv3485c | Rv3485c | 945 | 3884119 | 3885063 | BCG_3549c | BCG_3549c | 945 | 3906370 | 3907314 | BCGDan_3623 | BCGDan_3623 | 945 | 3827251 | 3828132 | BCGDan_sapMKO_3548 | BCGDan_sapMKO_3548 | 882 |
| 3905772 | 3906221 | Rv3486 | Rv3486 | 450 | 3885269 | 3885718 | BCG_3550 | BCG_3550 | 450 | 3907520 | 3907969 | BCGDan_3624 | BCGDan_3624 | 450 | 3828401 | 3828850 | BCGDan_sapMKO_3549 | BCGDan_sapMKO_3549 | 450 |
| 3906174 | 3907007 | lipF | Rv3487c | 834 | 3885671 | 3886504 | lipF | BCG_3551c | 834 | 3907922 | 3908755 | lipF | BCGDan_3625 | 834 | 3828803 | 3829621 | lipF | BCGDan_sapMKO_3550 | 819 |
| 3907667 | 3907990 | Rv3488 | Rv3488 | 324 | 3887164 | 3887487 | BCG_3552 | BCG_3552 | 324 | 3909415 | 3909738 | BCGDan_3626 | BCGDan_3626 | 324 | 3830296 | 3830619 | BCGDan_sapMKO_3551 | BCGDan_sapMKO_3551 | 324 |
| 3908072 | 3908236 | Rv3489 | Rv3489 | 165 | 3887569 | 3887733 | BCG_3553 | BCG_3553 | 165 | 3909820 | 3909984 | BCGDan_3627 | BCGDan_3627 | 165 | 3830701 | 3830865 | BCGDan_sapMKO_3552 | BCGDan_sapMKO_3552 | 165 |
| 3908236 | 3909738 | otsA | Rv3490 | 1503 | 3887733 | 3889235 | otsA | BCG_3554 | 1503 | 3909984 | 3911486 | otsA | BCGDan_3628 | 1503 | 3830865 | 3832367 | otsA | BCGDan_sapMKO_3553 | 1503 |
| 3909890 | 3910468 | Rv3491 | Rv3491 | 579 | 3889387 | 3889965 | BCG_3555 | BCG_3555 | 579 | 3911638 | 3912216 | BCGDan_3629 | BCGDan_3629 | 579 | 3832519 | 3833097 | BCGDan_sapMKO_3554 | BCGDan_sapMKO_3554 | 579 |
| 3910465 | 3910947 | Rv3492c | Rv3492c | 483 | 3889962 | 3890444 | BCG_3556c | BCG_3556c | 483 | 3912213 | 3912695 | BCGDan_3630 | BCGDan_3630 | 483 | 3833094 | 3833576 | BCGDan_sapMKO_3555 | BCGDan_sapMKO_3555 | 483 |
| 3910947 | 3911675 | Rv3493c | Rv3493c | 729 | 3890444 | 3891172 | BCG_3557c | BCG_3557c | 729 | 3912695 | 3913423 | BCGDan_3631 | BCGDan_3631 | 729 | 3833576 | 3834304 | BCGDan_sapMKO_3556 | BCGDan_sapMKO_3556 | 729 |
| 3911675 | 3913369 | mce4F | Rv3494c | 1695 | 3891172 | 3892866 | mce4F | BCG_3558c | 1695 | 3913423 | 3915117 | mce4F | BCGDan_3632 | 1695 | 3834304 | 3835998 | mce4F | BCGDan_sapMKO_3557 | 1695 |
| 3913380 | 3914534 | lprN | Rv3495c | 1155 | 3892877 | 3894031 | lprN | BCG_3559c | 1155 | 3915128 | 3916282 | lprN | BCGDan_3633 | 1155 | 3836009 | 3837163 | lprN | BCGDan_sapMKO_3558 | 1155 |
| 3914531 | 3915886 | mce4D | Rv3496c | 1356 | 3894028 | 3895383 | mce4D | BCG_3560c | 1356 | 3916279 | 3917634 | mce4D | BCGDan_3634 | 1356 | 3837160 | 3838512 | mce4D | BCGDan_sapMKO_3559 | 1353 |
| 3915883 | 3916956 | mce4C | Rv3497c | 1074 | 3895380 | 3896453 | mce4C | BCG_3561c | 1074 | 3917631 | 3918704 | mce4C | BCGDan_3635 | 1074 | 3838512 | 3839519 | mce4C | BCGDan_sapMKO_3560 | 1008 |
| 3916946 | 3917998 | mce4B | Rv3498c | 1053 | 3896443 | 3897495 | mce4B | BCG_3562c | 1053 | 3918694 | 3919746 | mce4B | BCGDan_3636 | 1053 | 3839575 | 3840627 | mce4B | BCGDan_sapMKO_3561 | 1053 |
| 3917998 | 3919200 | mce4A | Rv3499c | 1203 | 3897495 | 3898697 | mce4A | BCG_3563c | 1203 | 3919746 | 3920948 | mce4A | BCGDan_3637 | 1203 | 3840627 | 3841829 | mce4A | BCGDan_sapMKO_3562 | 1203 |
| 3919220 | 3920062 | yrbE4B | Rv3500c | 843 | 3898717 | 3899559 | yrbE4B | BCG_3564c | 843 | 3920968 | 3921810 | yrbE4B | BCGDan_3638 | 843 | 3841849 | 3842691 | yrbE4B | BCGDan_sapMKO_3563 | 843 |
| 3920097 | 3920861 | yrbE4A | Rv3501c | 765 | 3899594 | 3900358 | yrbE4A | BCG_3565c | 765 | 3921845 | 3922609 | yrbE4A | BCGDan_3639 | 765 | 3842726 | 3843490 | yrbE4A | BCGDan_sapMKO_3564 | 765 |
| 3921087 | 3922040 | Rv3502c | Rv3502c | 954 | 3900584 | 3901537 | BCG_3566c | BCG_3566c | 954 | 3922835 | 3923788 | BCGDan_3640 | BCGDan_3640 | 954 | 3843716 | 3844606 | BCGDan_sapMKO_3565 | BCGDan_sapMKO_3565 | 891 |
| 3922065 | 3922256 | fdxD | Rv3503c | 192 | 3901562 | 3901753 | fdxD | BCG_3567c | 192 | 3923813 | 3924004 | fdxD | BCGDan_3641 | 192 | 3844694 | 3844885 | fdxD | BCGDan_sapMKO_3566 | 192 |
| 3922471 | 3923673 | fadE26 | Rv3504 | 1203 | 3901968 | 3903170 | fadE26 | BCG_3568 | 1203 | 3924219 | 3925421 | fadE26 | BCGDan_3642 | 1203 | 3845100 | 3846302 | fadE26 | BCGDan_sapMKO_3567 | 1203 |
| 3923698 | 3924819 | fadE27 | Rv3505 | 1122 | 3903195 | 3904316 | fadE27 | BCG_3569 | 1122 | 3925446 | 3926567 | fadE27 | BCGDan_3643 | 1122 | 3846327 | 3847448 | fadE27 | BCGDan_sapMKO_3568 | 1122 |
| 3924890 | 3926398 | fadD17 | Rv3506 | 1509 | 3904387 | 3905895 | fadD17 | BCG_3570 | 1509 | 3926638 | 3928146 | fadD17 | BCGDan_3644 | 1509 | 3847519 | 3849027 | fadD17 | BCGDan_sapMKO_3569 | 1509 |
| 3926569 | 3930714 | PE_PGRS53 | Rv3507 | 4146 | 3906066 | 3910184 | PE_PGRS53 | BCG_3571 | 4119 | 3928317 | 3932435 | PE_PGRS53 | BCGDan_3645 | 4119 | 3849198 | 3853316 | PE_PGRS53 | BCGDan_sapMKO_3570 | 4119 |
| 3931005 | 3936710 | PE_PGRS54 | Rv3508 | 5706 | 3910475 | 3916759 | PE_PGRS54 | BCG_3572 | 6285 | 3932726 | 3939475 | PE_PGRS54 | BCGDan_3646 | 6750 | 3853607 | 3860356 | PE_PGRS54 | BCGDan_sapMKO_3571 | 6750 |
| 3936877 | 3938424 | ilvX | Rv3509c | 1548 | 3916926 | 3918473 | ilvX | BCG_3573c | 1548 | 3939642 | 3941189 | ilvX | BCGDan_3647 | 1548 | 3860523 | 3862070 | ilvX | BCGDan_sapMKO_3572 | 1548 |
| 3938421 | 3939257 | Rv3510c | Rv3510c | 837 | 3918470 | 3919306 | BCG_3574c | BCG_3574c | 837 | 3941186 | 3942022 | BCGDan_3648 | BCGDan_3648 | 837 | 3862067 | 3862903 | BCGDan_sapMKO_3573 | BCGDan_sapMKO_3573 | 837 |
| 3939617 | 3941761 | PE_PGRS55 | Rv3511 | 2145 | 3919666 | 3925098 | PE_PGRS55 | BCG_3575 | 5433 | 3942382 | 3947814 | PE_PGRS55 | BCGDan_3649 | 5433 | 3863263 | 3868695 | PE_PGRS55 | BCGDan_sapMKO_3574 | 5433 |
| 3945092 | 3945748 | fadD18 | Rv3513c | 657 | 3925227 | 3925883 | fadD18 | BCG_3576c | 657 | 3947943 | 3948599 | fadD18 | BCGDan_3650 | 657 | 3868824 | 3869480 | fadD18 | BCGDan_sapMKO_3575 | 657 |

| | | | | | | | | | | | | | | | | | | | |
|---|---|---|---|---|---|---|---|---|---|---|---|---|---|---|---|---|---|---|---|
| 3945794 | 3950263 | PE_PGRS57 | Rv3514 | 4470 | 3925929 | 3929156 | PE_PGRS57 | BCG_3577 | 3228 | 3948645 | 3951872 | PE_PGRS57 | BCGDan_3651 | 3228 | 3869526 | 3872753 | PE_PGRS57 | BCGDan_sapMKO_3576 | 3228 |
| 3950824 | 3952470 | fadD19 | Rv3515c | 1647 | 3929717 | 3931363 | fadD19 | BCG_3578c | 1647 | 3952433 | 3954079 | fadD19 | BCGDan_3652 | 1647 | 3873314 | 3874960 | fadD19 | BCGDan_sapMKO_3577 | 1647 |
| 3952544 | 3953335 | echA19 | Rv3516 | 792 | 3931437 | 3932228 | echA19 | BCG_3579 | 792 | 3954153 | 3954944 | echA19 | BCGDan_3653 | 792 | 3875016 | 3875825 | echA19 | BCGDan_sapMKO_3578 | 810 |
| 3953431 | 3954270 | Rv3517 | Rv3517 | 840 | 3932324 | 3933163 | BCG_3580 | BCG_3580 | 840 | 3955040 | 3955879 | BCGDan_3654 | BCGDan_3654 | 840 | 3875882 | 3876760 | BCGDan_sapMKO_3579 | BCGDan_sapMKO_3579 | 879 |
| NA | NA | NA | NA | NA | 3933218 | 3933799 | cyp142b | BCG_3581c | 582 | 3955934 | 3956515 | cyp142b | BCGDan_3655 | 582 | 3876815 | 3877402 | cyp142b | BCGDan_sapMKO_3580 | 588 |
| 3954325 | 3955521 | cyp142 | Rv3518c | 1197 | 3933796 | 3934413 | cyp142a | BCG_3582c | 618 | 3956512 | 3957129 | cyp142a | BCGDan_3656 | 618 | 3877393 | 3878010 | cyp142a | BCGDan_sapMKO_3581 | 618 |
| 3955550 | 3956260 | Rv3519 | Rv3519 | 711 | 3934442 | 3935152 | BCG_3583 | BCG_3583 | 711 | 3957158 | 3957868 | BCGDan_3657 | BCGDan_3657 | 711 | 3878039 | 3878749 | BCGDan_sapMKO_3582 | BCGDan_sapMKO_3582 | 711 |
| 3956325 | 3957368 | Rv3520c | Rv3520c | 1044 | 3935217 | 3936260 | BCG_3584c | BCG_3584c | 1044 | 3957933 | 3958976 | BCGDan_3658 | BCGDan_3658 | 1044 | 3878814 | 3879845 | BCGDan_sapMKO_3583 | BCGDan_sapMKO_3583 | 1032 |
| 3957521 | 3958432 | Rv3521 | Rv3521 | 912 | 3936413 | 3937324 | BCG_3585 | BCG_3585 | 912 | 3959129 | 3960040 | BCGDan_3659 | BCGDan_3659 | 912 | 3879917 | 3880921 | BCGDan_sapMKO_3584 | BCGDan_sapMKO_3584 | 1005 |
| 3958448 | 3959512 | ltp4 | Rv3522 | 1065 | 3937340 | 3938404 | BCG_3586 | BCG_3586 | 1065 | 3960056 | 3961120 | BCGDan_3660 | BCGDan_3660 | 1065 | 3880937 | 3882001 | BCGDan_sapMKO_3585 | BCGDan_sapMKO_3585 | 1065 |
| 3959529 | 3960713 | ltp3 | Rv3523 | 1185 | 3938421 | 3939605 | BCG_3587 | BCG_3587 | 1185 | 3961137 | 3962321 | BCGDan_3661 | BCGDan_3661 | 1185 | 3882018 | 3883202 | BCGDan_sapMKO_3586 | BCGDan_sapMKO_3586 | 1185 |
| 3960755 | 3961786 | Rv3524 | Rv3524 | 1032 | 3939647 | 3940678 | BCG_3588 | BCG_3588 | 1032 | 3962363 | 3963394 | BCGDan_3662 | BCGDan_3662 | 1032 | 3883244 | 3884275 | BCGDan_sapMKO_3587 | BCGDan_sapMKO_3587 | 1032 |
| 3961800 | 3962324 | Rv3525c | Rv3525c | 525 | 3940692 | 3941216 | BCG_3589c | BCG_3589c | 525 | 3963408 | 3963932 | BCGDan_3663 | BCGDan_3663 | 525 | 3884289 | 3884813 | BCGDan_sapMKO_3588 | BCGDan_sapMKO_3588 | 525 |
| 3962439 | 3963599 | kshA | Rv3526 | 1161 | 3941331 | 3942491 | BCG_3590 | BCG_3590 | 1161 | 3964047 | 3965207 | BCGDan_3664 | BCGDan_3664 | 1161 | 3884928 | 3886088 | BCGDan_sapMKO_3589 | BCGDan_sapMKO_3589 | 1161 |
| 3963605 | 3964054 | Rv3527 | Rv3527 | 450 | 3942497 | 3942946 | BCG_3591 | BCG_3591 | 450 | 3965213 | 3965662 | BCGDan_3665 | BCGDan_3665 | 450 | 3886094 | 3886543 | BCGDan_sapMKO_3590 | BCGDan_sapMKO_3590 | 450 |
| 3964479 | 3965192 | Rv3528c | Rv3528c | 714 | 3943371 | 3944084 | BCG_3592c | BCG_3592c | 714 | 3966087 | 3966800 | BCGDan_3666 | BCGDan_3666 | 714 | 3886968 | 3887681 | BCGDan_sapMKO_3591 | BCGDan_sapMKO_3591 | 714 |
| 3965884 | 3967038 | Rv3529c | Rv3529c | 1155 | 3944776 | 3945930 | BCG_3593c | BCG_3593c | 1155 | 3967492 | 3968646 | BCGDan_3667 | BCGDan_3667 | 1155 | 3888373 | 3889527 | BCGDan_sapMKO_3592 | BCGDan_sapMKO_3592 | 1155 |
| 3967038 | 3967820 | Rv3530c | Rv3530c | 783 | 3945930 | 3946712 | BCG_3594c | BCG_3594c | 783 | 3968646 | 3969428 | BCGDan_3668 | BCGDan_3668 | 783 | 3889527 | 3890309 | BCGDan_sapMKO_3593 | BCGDan_sapMKO_3593 | 783 |
| 3967817 | 3968944 | Rv3531c | Rv3531c | 1128 | 3946709 | 3947836 | BCG_3595c | BCG_3595c | 1128 | 3969425 | 3970552 | BCGDan_3669 | BCGDan_3669 | 1128 | 3890306 | 3891433 | BCGDan_sapMKO_3594 | BCGDan_sapMKO_3594 | 1128 |
| 3969343 | 3970563 | PPE61 | Rv3532 | 1221 | 3948235 | 3949455 | PPE61 | BCG_3596 | 1221 | 3970951 | 3972171 | PPE61 | BCGDan_3670 | 1221 | 3891832 | 3893052 | PPE61 | BCGDan_sapMKO_3595 | 1221 |
| 3970705 | 3972453 | PPE62 | Rv3533c | 1749 | 3949597 | 3951345 | PPE62 | BCG_3597c | 1749 | 3972313 | 3974061 | PPE62 | BCGDan_3671 | 1749 | 3893194 | 3894942 | PPE62 | BCGDan_sapMKO_3596 | 1749 |
| 3972552 | 3973592 | hsaF | Rv3534c | 1041 | 3951444 | 3952484 | BCG_3598c | BCG_3598c | 1041 | 3974160 | 3975200 | BCGDan_3672 | BCGDan_3672 | 1041 | 3895041 | 3896081 | BCGDan_sapMKO_3597 | BCGDan_sapMKO_3597 | 1041 |
| 3973589 | 3974500 | hsaG | Rv3535c | 912 | 3952481 | 3953392 | BCG_3599c | BCG_3599c | 912 | 3975197 | 3976108 | BCGDan_3673 | BCGDan_3673 | 912 | 3896078 | 3896989 | BCGDan_sapMKO_3598 | BCGDan_sapMKO_3598 | 912 |
| 3974511 | 3975296 | hsaE | Rv3536c | 786 | 3953403 | 3954188 | BCG_3600c | BCG_3600c | 786 | 3976119 | 3976904 | BCGDan_3674 | BCGDan_3674 | 786 | 3897000 | 3897785 | BCGDan_sapMKO_3599 | BCGDan_sapMKO_3599 | 786 |
| 3975369 | 3977060 | kstD | Rv3537 | 1692 | 3954261 | 3955952 | BCG_3601 | BCG_3601 | 1692 | 3976977 | 3978668 | BCGDan_3675 | BCGDan_3675 | 1692 | 3897849 | 3899549 | BCGDan_sapMKO_3600 | BCGDan_sapMKO_3600 | 1701 |
| 3977062 | 3977922 | Rv3538 | Rv3538 | 861 | 3955954 | 3956814 | BCG_3602 | BCG_3602 | 861 | 3978670 | 3979530 | BCGDan_3676 | BCGDan_3676 | 861 | 3899551 | 3900411 | BCGDan_sapMKO_3601 | BCGDan_sapMKO_3601 | 861 |
| 3978059 | 3979498 | PPE63 | Rv3539 | 1440 | 3956951 | 3958390 | PPE63 | BCG_3603 | 1440 | 3979667 | 3981106 | PPE63 | BCGDan_3677 | 1440 | 3900548 | 3901987 | PPE63 | BCGDan_sapMKO_3602 | 1440 |
| 3979499 | 3980659 | ltp2 | Rv3540c | 1161 | 3958391 | 3959551 | ltp2 | BCG_3604c | 1161 | 3981107 | 3982267 | ltp2 | BCGDan_3678 | 1161 | 3901988 | 3903148 | ltp2 | BCGDan_sapMKO_3603 | 1161 |
| 3980659 | 3981048 | Rv3541c | Rv3541c | 390 | 3959551 | 3959940 | BCG_3605c | BCG_3605c | 390 | 3982267 | 3982656 | BCGDan_3679 | BCGDan_3679 | 390 | 3903148 | 3903537 | BCGDan_sapMKO_3604 | BCGDan_sapMKO_3604 | 390 |
| 3981045 | 3981980 | Rv3542c | Rv3542c | 936 | 3959937 | 3960872 | BCG_3606c | BCG_3606c | 936 | 3982653 | 3983588 | BCGDan_3680 | BCGDan_3680 | 936 | 3903534 | 3904469 | BCGDan_sapMKO_3605 | BCGDan_sapMKO_3605 | 936 |
| 3981977 | 3983140 | fadE29 | Rv3543c | 1164 | 3960869 | 3962032 | fadE29 | BCG_3607c | 1164 | 3983585 | 3984748 | fadE29 | BCGDan_3681 | 1164 | 3904466 | 3905629 | fadE29 | BCGDan_sapMKO_3606 | 1164 |
| 3983125 | 3984144 | fadE28 | Rv3544c | 1020 | 3962017 | 3963036 | fadE28 | BCG_3608c | 1020 | 3984733 | 3985752 | fadE28 | BCGDan_3682 | 1020 | 3905614 | 3906633 | fadE28 | BCGDan_sapMKO_3607 | 1020 |
| 3984144 | 3985445 | cyp125 | Rv3545c | 1302 | 3963036 | 3964337 | cyp125 | BCG_3609c | 1302 | 3985752 | 3987053 | cyp125 | BCGDan_3683 | 1302 | 3906633 | 3907886 | cyp125 | BCGDan_sapMKO_3608 | 1254 |
| 3985557 | 3986732 | fadA5 | Rv3546 | 1176 | 3964449 | 3965624 | fadA5 | BCG_3610 | 1176 | 3987165 | 3988340 | fadA5 | BCGDan_3684 | 1176 | 3908046 | 3909221 | fadA5 | BCGDan_sapMKO_3609 | 1176 |
| 3986844 | 3987299 | ddn | Rv3547 | 456 | 3965736 | 3966191 | BCG_3611 | BCG_3611 | 456 | 3988452 | 3988907 | BCGDan_3685 | BCGDan_3685 | 456 | 3909333 | 3909788 | BCGDan_sapMKO_3610 | BCGDan_sapMKO_3610 | 456 |
| 3987382 | 3988296 | Rv3548c | Rv3548c | 915 | 3966274 | 3967188 | BCG_3612c | BCG_3612c | 915 | 3988990 | 3989904 | BCGDan_3686 | BCGDan_3686 | 915 | 3909871 | 3910785 | BCGDan_sapMKO_3611 | BCGDan_sapMKO_3611 | 915 |
| 3988319 | 3989098 | Rv3549c | Rv3549c | 780 | 3967211 | 3967990 | BCG_3613c | BCG_3613c | 780 | 3989927 | 3990706 | BCGDan_3687 | BCGDan_3687 | 780 | 3910808 | 3911587 | BCGDan_sapMKO_3612 | BCGDan_sapMKO_3612 | 780 |
| 3989153 | 3989896 | echA20 | Rv3550 | 744 | 3968045 | 3968788 | echA20 | BCG_3614 | 744 | 3990761 | 3991504 | echA20 | BCGDan_3688 | 744 | 3911642 | 3912385 | echA20 | BCGDan_sapMKO_3613 | 744 |
| 3989896 | 3990774 | Rv3551 | Rv3551 | 879 | 3968788 | 3969666 | BCG_3615 | BCG_3615 | 879 | 3991504 | 3992382 | BCGDan_3689 | BCGDan_3689 | 879 | 3912385 | 3913263 | BCGDan_sapMKO_3614 | BCGDan_sapMKO_3614 | 879 |
| 3990771 | 3991523 | Rv3552 | Rv3552 | 753 | 3969663 | 3970415 | BCG_3616 | BCG_3616 | 753 | 3992379 | 3993131 | BCGDan_3690 | BCGDan_3690 | 753 | 3913260 | 3914012 | BCGDan_sapMKO_3615 | BCGDan_sapMKO_3615 | 753 |
| 3991621 | 3992688 | Rv3553 | Rv3553 | 1068 | 3970513 | 3971580 | BCG_3617 | BCG_3617 | 1068 | 3993229 | 3994296 | BCGDan_3691 | BCGDan_3691 | 1068 | 3914110 | 3915177 | BCGDan_sapMKO_3616 | BCGDan_sapMKO_3616 | 1068 |
| 3992685 | 3994742 | fdxB | Rv3554 | 2058 | 3971577 | 3973634 | fdxB | BCG_3618 | 2058 | 3994293 | 3996350 | fdxB | BCGDan_3692 | 2058 | 3915213 | 3917231 | fdxB | BCGDan_sapMKO_3617 | 2019 |
| 3994830 | 3995699 | Rv3555c | Rv3555c | 870 | 3973722 | 3974591 | BCG_3619c | BCG_3619c | 870 | 3996438 | 3997307 | BCGDan_3693 | BCGDan_3693 | 870 | 3917319 | 3918188 | BCGDan_sapMKO_3618 | BCGDan_sapMKO_3618 | 870 |
| 3995804 | 3996964 | fadA6 | Rv3556c | 1161 | 3974696 | 3975856 | fadA6 | BCG_3620c | 1161 | 3997412 | 3998572 | fadA6 | BCGDan_3694 | 1161 | 3918293 | 3919453 | fadA6 | BCGDan_sapMKO_3619 | 1161 |
| 3997029 | 3997631 | Rv3557c | Rv3557c | 603 | 3975921 | 3976523 | BCG_3621c | BCG_3621c | 603 | 3998637 | 3999239 | BCGDan_3695 | BCGDan_3695 | 603 | 3919518 | 3920120 | BCGDan_sapMKO_3620 | BCGDan_sapMKO_3620 | 603 |
| 3997980 | 3999638 | PPE64 | Rv3558 | 1659 | 3976872 | 3978530 | PPE64 | BCG_3622 | 1659 | 3999588 | 4001246 | PPE64 | BCGDan_3696 | 1659 | 3920469 | 3922127 | PPE64 | BCGDan_sapMKO_3621 | 1659 |
| 3999647 | 4000435 | Rv3559c | Rv3559c | 789 | 3978539 | 3979327 | BCG_3623c | BCG_3623c | 789 | 4001255 | 4002043 | BCGDan_3697 | BCGDan_3697 | 789 | 3922136 | 3922924 | BCGDan_sapMKO_3622 | BCGDan_sapMKO_3622 | 789 |
| 4000432 | 4001589 | fadE30 | Rv3560c | 1158 | 3979324 | 3980481 | fadE30 | BCG_3624c | 1158 | 4002040 | 4003197 | fadE30 | BCGDan_3698 | 1158 | 3922921 | 3924078 | fadE30 | BCGDan_sapMKO_3623 | 1158 |
| 4001637 | 4003160 | fadD3 | Rv3561 | 1524 | 3980529 | 3982052 | fadD3 | BCG_3625 | 1524 | 4003245 | 4004768 | fadD3 | BCGDan_3699 | 1524 | 3924126 | 3925649 | fadD3 | BCGDan_sapMKO_3624 | 1524 |
| 4003161 | 4004294 | fadE31 | Rv3562 | 1134 | 3982053 | 3983186 | fadE31 | BCG_3626 | 1134 | 4004769 | 4005902 | fadE31 | BCGDan_3700 | 1134 | 3925650 | 3926783 | fadE31 | BCGDan_sapMKO_3625 | 1134 |
| 4004291 | 4005250 | fadE32 | Rv3563 | 960 | 3983183 | 3984142 | fadE32 | BCG_3627 | 960 | 4005899 | 4006858 | fadE32 | BCGDan_3701 | 960 | 3926780 | 3927739 | fadE32 | BCGDan_sapMKO_3626 | 960 |
| 4005247 | 4006203 | fadE33 | Rv3564 | 957 | 3984139 | 3985095 | fadE33 | BCG_3628 | 957 | 4006855 | 4007811 | fadE33 | BCGDan_3702 | 957 | 3927736 | 3928692 | fadE33 | BCGDan_sapMKO_3627 | 957 |
| 4006200 | 4007366 | aspB | Rv3565 | 1167 | 3985092 | 3986258 | aspB | BCG_3629 | 1167 | 4007808 | 4008974 | aspB | BCGDan_3703 | 1167 | 3928677 | 3929855 | aspB | BCGDan_sapMKO_3628 | 1179 |
| 4007331 | 4008182 | nat | Rv3566c | 852 | 3986223 | 3987074 | nat | BCG_3630c | 852 | 4008939 | 4009790 | nat | BCGDan_3704 | 852 | 3929820 | 3930671 | nat | BCGDan_sapMKO_3629 | 852 |
| 4008167 | 4008433 | Rv3566A | Rv3566A | 267 | 3987059 | 3987325 | BCG_3631c | BCG_3631c | 267 | 4009775 | 4010041 | BCGDan_3705 | BCGDan_3705 | 267 | 3930656 | 3930922 | BCGDan_sapMKO_3630 | BCGDan_sapMKO_3630 | 267 |
| 4008719 | 4009282 | hsaB | Rv3567c | 564 | 3987611 | 3988174 | BCG_3632c | BCG_3632c | 564 | 4010327 | 4010890 | BCGDan_3706 | BCGDan_3706 | 564 | 3931208 | 3931771 | BCGDan_sapMKO_3631 | BCGDan_sapMKO_3631 | 564 |
| 4009297 | 4010199 | hsaC | Rv3568c | 903 | 3988189 | 3989091 | bphC | BCG_3633c | 903 | 4010905 | 4011807 | bphC | BCGDan_3707 | 903 | 3931786 | 3932688 | bphC | BCGDan_sapMKO_3632 | 903 |
| 4010196 | 4011071 | hsaD | Rv3569c | 876 | 3989088 | 3989963 | bphD | BCG_3634c | 876 | 4011804 | 4012679 | bphD | BCGDan_3708 | 876 | 3932685 | 3933560 | bphD | BCGDan_sapMKO_3633 | 876 |
| 4011086 | 4012270 | hsaA | Rv3570c | 1185 | 3989978 | 3991162 | BCG_3635c | BCG_3635c | 1185 | 4012694 | 4013878 | BCGDan_3709 | BCGDan_3709 | 1185 | 3933575 | 3934723 | BCGDan_sapMKO_3634 | BCGDan_sapMKO_3634 | 1149 |
| 4012417 | 4013493 | kshB | Rv3571 | 1077 | 3991309 | 3992385 | hmp | BCG_3636 | 1077 | 4014025 | 4015101 | hmp | BCGDan_3710 | 1077 | 3934906 | 3935982 | hmp | BCGDan_sapMKO_3635 | 1077 |
| 4013511 | 4014041 | Rv3572 | Rv3572 | 531 | 3992403 | 3992933 | BCG_3637 | BCG_3637 | 531 | 4015119 | 4015649 | BCGDan_3711 | BCGDan_3711 | 531 | 3936000 | 3936530 | BCGDan_sapMKO_3636 | BCGDan_sapMKO_3636 | 531 |
| 4014077 | 4016212 | fadE34 | Rv3573c | 2136 | 3992969 | 3995104 | fadE34 | BCG_3638c | 2136 | 4015685 | 4017820 | fadE34 | BCGDan_3712 | 2136 | 3936566 | 3938701 | fadE34 | BCGDan_sapMKO_3637 | 2136 |
| 4016484 | 4017083 | kstR | Rv3574 | 600 | 3995376 | 3995975 | BCG_3639 | BCG_3639 | 600 | 4018092 | 4018691 | BCGDan_3713 | BCGDan_3713 | 600 | 3938967 | 3939572 | BCGDan_sapMKO_3638 | BCGDan_sapMKO_3638 | 606 |
| 4017089 | 4018168 | Rv3575c | Rv3575c | 1080 | 3995981 | 3997060 | BCG_3640c | BCG_3640c | 1080 | 4018697 | 4019776 | BCGDan_3714 | BCGDan_3714 | 1080 | 3939578 | 3940531 | BCGDan_sapMKO_3639 | BCGDan_sapMKO_3639 | 954 |
| 4018358 | 4019071 | lppH | Rv3576 | 714 | 3997250 | 3997963 | lppH | BCG_3641 | 714 | 4019966 | 4020679 | lppH | BCGDan_3715 | 714 | 3940847 | 3941560 | lppH | BCGDan_sapMKO_3640 | 714 |
| 4019262 | 4020128 | Rv3577 | Rv3577 | 867 | 3998154 | 3999020 | BCG_3642 | BCG_3642 | 867 | 4020870 | 4021736 | BCGDan_3716 | BCGDan_3716 | 867 | 3941793 | 3942617 | BCGDan_sapMKO_3641 | BCGDan_sapMKO_3641 | 825 |
| 4020142 | 4021383 | arsB2 | Rv3578 | 1242 | 3999034 | 4000275 | arsB2 | BCG_3643 | 1242 | 4021750 | 4022991 | arsB2 | BCGDan_3717 | 1242 | 3942664 | 3943872 | arsB2 | BCGDan_sapMKO_3642 | 1209 |
| 4021425 | 4022393 | Rv3579c | Rv3579c | 969 | 4000317 | 4001285 | BCG_3644c | BCG_3644c | 969 | 4023033 | 4024001 | BCGDan_3718 | BCGDan_3718 | 969 | 3943914 | 3944882 | BCGDan_sapMKO_3643 | BCGDan_sapMKO_3643 | 969 |
| 4022394 | 4023803 | cysS1 | Rv3580c | 1410 | 4001286 | 4002695 | cysS1 | BCG_3645c | 1410 | 4024002 | 4025411 | cysS1 | BCGDan_3719 | 1410 | 3944883 | 3946292 | cysS1 | BCGDan_sapMKO_3644 | 1410 |
| 4023868 | 4024347 | ispF | Rv3581c | 480 | 4002760 | 4003239 | ispF | BCG_3646c | 480 | 4025476 | 4025955 | ispF | BCGDan_3720 | 480 | 3946357 | 3946836 | ispF | BCGDan_sapMKO_3645 | 480 |
| 4024344 | 4025039 | ispD | Rv3582c | 696 | 4003236 | 4003931 | ispD | BCG_3647c | 696 | 4025952 | 4026647 | ispD | BCGDan_3721 | 696 | 3946833 | 3947528 | ispD | BCGDan_sapMKO_3646 | 696 |
| 4025056 | 4025544 | Rv3583c | Rv3583c | 489 | 4003948 | 4004436 | BCG_3648c | BCG_3648c | 489 | 4026664 | 4027152 | BCGDan_3722 | BCGDan_3722 | 489 | 3947545 | 3948033 | BCGDan_sapMKO_3647 | BCGDan_sapMKO_3647 | 489 |
| 4025830 | 4026378 | lpqE | Rv3584 | 549 | 4004721 | 4005269 | lpqE | BCG_3649 | 549 | 4027437 | 4027985 | lpqE | BCGDan_3723 | 549 | 3948354 | 3948866 | lpqE | BCGDan_sapMKO_3648 | 513 |
| 4026444 | 4027886 | radA | Rv3585 | 1443 | 4005335 | 4006777 | radA | BCG_3650 | 1443 | 4028051 | 4029493 | radA | BCGDan_3724 | 1443 | 3948932 | 3950374 | radA | BCGDan_sapMKO_3649 | 1443 |
| 4027891 | 4028967 | Rv3586 | Rv3586 | 1077 | 4006782 | 4007858 | BCG_3651 | BCG_3651 | 1077 | 4029498 | 4030574 | BCGDan_3725 | BCGDan_3725 | 1077 | 3950379 | 3951455 | BCGDan_sapMKO_3650 | BCGDan_sapMKO_3650 | 1077 |
| 4028968 | 4029762 | Rv3587c | Rv3587c | 795 | 4007859 | 4008650 | BCG_3652c | BCG_3652c | 792 | 4030575 | 4031390 | BCGDan_3726 | BCGDan_3726 | 816 | 3951456 | 3952271 | BCGDan_sapMKO_3651 | BCGDan_sapMKO_3651 | 816 |
| 4029871 | 4030494 | canB | Rv3588c | 624 | 4008759 | 4009382 | canB | BCG_3653c | 624 | 4031475 | 4032098 | canB | BCGDan_3727 | 624 | 3952356 | 3952979 | canB | BCGDan_sapMKO_3652 | 624 |
| 4030493 | 4031407 | mutY | Rv3589 | 915 | 4009381 | 4010295 | mutY | BCG_3654 | 915 | 4032097 | 4033011 | mutY | BCGDan_3728 | 915 | 3952978 | 3953892 | mutY | BCGDan_sapMKO_3653 | 915 |
| 4031404 | 4033158 | PE_PGRS58 | Rv3590c | 1755 | 4010292 | 4012046 | PE_PGRS58 | BCG_3655c | 1755 | 4033008 | 4034762 | PE_PGRS58 | BCGDan_3729 | 1755 | 3953889 | 3955643 | PE_PGRS58 | BCGDan_sapMKO_3654 | 1755 |
| 4033269 | 4034042 | Rv3591c | Rv3591c | 774 | 4012157 | 4012930 | BCG_3656c | BCG_3656c | 774 | 4034873 | 4035646 | BCGDan_3730 | BCGDan_3730 | 774 | 3955754 | 3956518 | BCGDan_sapMKO_3655 | BCGDan_sapMKO_3655 | 765 |
| 4034057 | 4034374 | mhuD | Rv3592 | 318 | 4012945 | 4013262 | TB11.2 | BCG_3657 | 318 | 4035661 | 4035978 | TB11.2 | BCGDan_3731 | 318 | 3956542 | 3956859 | TB11.2 | BCGDan_sapMKO_3656 | 318 |
| 4034352 | 4035710 | lpqF | Rv3593 | 1359 | 4013240 | 4014598 | lpqF | BCG_3658 | 1359 | 4035956 | 4037314 | lpqF | BCGDan_3732 | 1359 | 3956879 | 3958195 | lpqF | BCGDan_sapMKO_3657 | 1317 |
| 4035857 | 4036684 | Rv3594 | Rv3594 | 828 | 4014745 | 4015572 | BCG_3659 | BCG_3659 | 828 | 4037461 | 4038288 | BCGDan_3733 | BCGDan_3733 | 828 | 3958342 | 3959169 | BCGDan_sapMKO_3658 | BCGDan_sapMKO_3658 | 828 |
| 4036731 | 4038050 | PE_PGRS59 | Rv3595c | 1320 | 4015486 | 4016964 | PE_PGRS59 | BCG_3660c | 1479 | 4038202 | 4039680 | PE_PGRS59 | BCGDan_3734 | 1479 | 3959083 | 3960561 | PE_PGRS59 | BCGDan_sapMKO_3659 | 1479 |
| 4038158 | 4040704 | clpC1 | Rv3596c | 2547 | 4017072 | 4019618 | clpC | BCG_3661c | 2547 | 4039788 | 4042334 | clpC | BCGDan_3735 | 2547 | 3960669 | 3963215 | clpC | BCGDan_sapMKO_3660 | 2547 |
| 4040981 | 4041319 | lsr2 | Rv3597c | 339 | 4019895 | 4020233 | lsr2 | BCG_3662c | 339 | 4042611 | 4042949 | lsr2 | BCGDan_3736 | 339 | 3963492 | 3963830 | lsr2 | BCGDan_sapMKO_3661 | 339 |

| | | | | | | | | | | | | | | | | | | | |
|---|---|---|---|---|---|---|---|---|---|---|---|---|---|---|---|---|---|---|---|
| 4041423 | 4042940 | lysS | Rv3598c | 1518 | 4020337 | 4021854 | lysS | BCG_3663c | 1518 | 4043053 | 4044570 | lysS | BCGDan_3737 | 1518 | 3963934 | 3965451 | lysS | BCGDan_sapMKO_3662 | 1518 |
| 4043041 | 4043859 | Rv3600c | Rv3600c | 819 | 4021955 | 4022773 | BCG_3664c | BCG_3664c | 819 | 4044671 | 4045489 | BCGDan_3738 | BCGDan_3738 | 819 | 3965552 | 3966370 | BCGDan_sapMKO_3663 | BCGDan_sapMKO_3663 | 819 |
| 4043862 | 4044281 | panD | Rv3601c | 420 | 4022776 | 4023195 | panD | BCG_3665c | 420 | 4045492 | 4045911 | panD | BCGDan_3739 | 420 | 3966373 | 3966780 | panD | BCGDan_sapMKO_3664 | 408 |
| 4044281 | 4045210 | panC | Rv3602c | 930 | 4023195 | 4024124 | panC | BCG_3666c | 930 | 4045911 | 4046840 | panC | BCGDan_3740 | 930 | 3966792 | 3967721 | panC | BCGDan_sapMKO_3665 | 930 |
| 4045207 | 4046118 | Rv3603c | Rv3603c | 912 | 4024121 | 4025032 | BCG_3667c | BCG_3667c | 912 | 4046837 | 4047748 | BCGDan_3741 | BCGDan_3741 | 912 | 3967718 | 3968629 | BCGDan_sapMKO_3666 | BCGDan_sapMKO_3666 | 912 |
| 4046303 | 4047496 | Rv3604c | Rv3604c | 1194 | 4025217 | 4026410 | BCG_3668c | BCG_3668c | 1194 | 4047933 | 4049126 | BCGDan_3742 | BCGDan_3742 | 1194 | 3968814 | 3970007 | BCGDan_sapMKO_3667 | BCGDan_sapMKO_3667 | 1194 |
| 4047705 | 4048181 | Rv3605c | Rv3605c | 477 | 4026619 | 4027095 | BCG_3669c | BCG_3669c | 477 | 4049335 | 4049811 | BCGDan_3743 | BCGDan_3743 | 477 | 3970216 | 3970692 | BCGDan_sapMKO_3668 | BCGDan_sapMKO_3668 | 477 |
| 4048181 | 4048747 | folK | Rv3606c | 567 | 4027095 | 4027661 | folK | BCG_3670c | 567 | 4049811 | 4050377 | folK | BCGDan_3744 | 567 | 3970692 | 3971258 | folK | BCGDan_sapMKO_3669 | 567 |
| 4048744 | 4049145 | folB | Rv3607c | 402 | 4027658 | 4028059 | folB | BCG_3671c | 402 | 4050374 | 4050775 | folB | BCGDan_3745 | 402 | 3971255 | 3971656 | folB | BCGDan_sapMKO_3670 | 402 |
| 4049138 | 4049980 | folP1 | Rv3608c | 843 | 4028052 | 4028894 | folP1 | BCG_3672c | 843 | 4050768 | 4051610 | folP1 | BCGDan_3746 | 843 | 3971649 | 3972491 | folP1 | BCGDan_sapMKO_3671 | 843 |
| 4049977 | 4050585 | folE | Rv3609c | 609 | 4028891 | 4029499 | folE | BCG_3673c | 609 | 4051607 | 4052215 | folE | BCGDan_3747 | 609 | 3972488 | 3973096 | folE | BCGDan_sapMKO_3672 | 609 |
| 4050601 | 4052883 | ftsH | Rv3610c | 2283 | 4029515 | 4031797 | ftsH | BCG_3674c | 2283 | 4052231 | 4054513 | ftsH | BCGDan_3748 | 2283 | 3973112 | 3975394 | ftsH | BCGDan_sapMKO_3673 | 2283 |
| 4052950 | 4053603 | Rv3611 | Rv3611 | 654 | 4031864 | 4032517 | BCG_3675 | BCG_3675 | 654 | 4054580 | 4055233 | BCGDan_3749 | BCGDan_3749 | 654 | 3975461 | 3976114 | BCGDan_sapMKO_3674 | BCGDan_sapMKO_3674 | 654 |
| 4053518 | 4053847 | Rv3612c | Rv3612c | 330 | 4032432 | 4032761 | BCG_3676c | BCG_3676c | 330 | 4055148 | 4055477 | BCGDan_3750 | BCGDan_3750 | 330 | 3976029 | 3976358 | BCGDan_sapMKO_3675 | BCGDan_sapMKO_3675 | 330 |
| 4053881 | 4054042 | Rv3613c | Rv3613c | 162 | 4032795 | 4032956 | BCG_3677c | BCG_3677c | 162 | 4055511 | 4055672 | BCGDan_3751 | BCGDan_3751 | 162 | 3976392 | 3976553 | BCGDan_sapMKO_3676 | BCGDan_sapMKO_3676 | 162 |
| 4054142 | 4054696 | espD | Rv3614c | 555 | 4033056 | 4033610 | BCG_3678c | BCG_3678c | 555 | 4055772 | 4056326 | BCGDan_3752 | BCGDan_3752 | 555 | 3976653 | 3977225 | BCGDan_sapMKO_3677 | BCGDan_sapMKO_3677 | 573 |
| 4054812 | 4055123 | espC | Rv3615c | 312 | 4033726 | 4034037 | BCG_3679c | BCG_3679c | 312 | 4056442 | 4056753 | BCGDan_3753 | BCGDan_3753 | 312 | 3977323 | 3977634 | BCGDan_sapMKO_3678 | BCGDan_sapMKO_3678 | 312 |
| 4055197 | 4056375 | espA | Rv3616c | 1179 | 4034111 | 4035289 | BCG_3680c | BCG_3680c | 1179 | 4056827 | 4058005 | BCGDan_3754 | BCGDan_3754 | 1179 | 3977708 | 3978877 | BCGDan_sapMKO_3679 | BCGDan_sapMKO_3679 | 1170 |
| 4062527 | 4063249 | lpqG | Rv3623 | 723 | 4035755 | 4036270 | lpqG | BCG_3681 | 516 | 4058471 | 4058986 | lpqG | BCGDan_3755 | 516 | 3979352 | 3979867 | lpqG | BCGDan_sapMKO_3680 | 516 |
| 4063254 | 4063904 | hpt | Rv3624c | 651 | 4036275 | 4036925 | hpt | BCG_3682c | 651 | 4058991 | 4059641 | hpt | BCGDan_3756 | 651 | 3979872 | 3980480 | hpt | BCGDan_sapMKO_3681 | 609 |
| 4063901 | 4064872 | mesJ | Rv3625c | 972 | 4036922 | 4037893 | mesJ | BCG_3683c | 972 | 4059638 | 4060609 | mesJ | BCGDan_3757 | 972 | 3980519 | 3981403 | mesJ | BCGDan_sapMKO_3682 | 885 |
| 4064851 | 4065903 | Rv3626c | Rv3626c | 1053 | 4037872 | 4038924 | BCG_3684c | BCG_3684c | 1053 | 4060588 | 4061640 | BCGDan_3758 | BCGDan_3758 | 1053 | 3981469 | 3982542 | BCGDan_sapMKO_3683 | BCGDan_sapMKO_3683 | 1074 |
| 4065900 | 4067285 | Rv3627c | Rv3627c | 1386 | 4038921 | 4040306 | BCG_3685c | BCG_3685c | 1386 | 4061637 | 4063022 | BCGDan_3759 | BCGDan_3759 | 1386 | 3982518 | 3983903 | BCGDan_sapMKO_3684 | BCGDan_sapMKO_3684 | 1386 |
| 4067423 | 4067911 | ppa | Rv3628 | 489 | 4040444 | 4040932 | ppa | BCG_3686 | 489 | 4063160 | 4063648 | ppa | BCGDan_3760 | 489 | 3984041 | 3984529 | ppa | BCGDan_sapMKO_3685 | 489 |
| 4067957 | 4069054 | Rv3629c | Rv3629c | 1098 | 4040978 | 4042075 | BCG_3687c | BCG_3687c | 1098 | 4063694 | 4064791 | BCGDan_3761 | BCGDan_3761 | 1098 | 3984575 | 3985672 | BCGDan_sapMKO_3686 | BCGDan_sapMKO_3686 | 1098 |
| 4069175 | 4070470 | Rv3630 | Rv3630 | 1296 | 4042196 | 4043491 | BCG_3688 | BCG_3688 | 1296 | 4064912 | 4066207 | BCGDan_3762 | BCGDan_3762 | 1296 | 3985793 | 3987088 | BCGDan_sapMKO_3687 | BCGDan_sapMKO_3687 | 1296 |
| 4070514 | 4071239 | Rv3631 | Rv3631 | 726 | 4043535 | 4044260 | BCG_3689 | BCG_3689 | 726 | 4066251 | 4066976 | BCGDan_3763 | BCGDan_3763 | 726 | 3987132 | 3987857 | BCGDan_sapMKO_3688 | BCGDan_sapMKO_3688 | 726 |
| 4071236 | 4071580 | Rv3632 | Rv3632 | 345 | 4044257 | 4044601 | BCG_3690 | BCG_3690 | 345 | 4066973 | 4067317 | BCGDan_3764 | BCGDan_3764 | 345 | 3987854 | 3988198 | BCGDan_sapMKO_3689 | BCGDan_sapMKO_3689 | 345 |
| 4071791 | 4072666 | Rv3633 | Rv3633 | 876 | 4044812 | 4045687 | BCG_3691 | BCG_3691 | 876 | 4067528 | 4068403 | BCGDan_3765 | BCGDan_3765 | 876 | 3988409 | 3989284 | BCGDan_sapMKO_3690 | BCGDan_sapMKO_3690 | 876 |
| 4072667 | 4073611 | galE1 | Rv3634c | 945 | 4045688 | 4046632 | galE1 | BCG_3692c | 945 | 4068404 | 4069348 | galE1 | BCGDan_3766 | 945 | 3989285 | 3990253 | galE1 | BCGDan_sapMKO_3691 | 969 |
| 4073634 | 4075409 | Rv3635 | Rv3635 | 1776 | 4046655 | 4048430 | BCG_3693 | BCG_3693 | 1776 | 4069386 | 4071146 | BCGDan_3767 | BCGDan_3767 | 1761 | 3990267 | 3992027 | BCGDan_sapMKO_3692 | BCGDan_sapMKO_3692 | 1761 |
| 4075752 | 4076099 | Rv3636 | Rv3636 | 348 | 4048773 | 4049120 | BCG_3694 | BCG_3694 | 348 | 4071489 | 4071836 | BCGDan_3768 | BCGDan_3768 | 348 | 3992370 | 3992717 | BCGDan_sapMKO_3693 | BCGDan_sapMKO_3693 | 348 |
| 4076484 | 4076984 | Rv3637 | Rv3637 | 501 | 4049505 | 4050005 | BCG_3695 | BCG_3695 | 501 | 4072221 | 4072721 | BCGDan_3769 | BCGDan_3769 | 501 | 3993102 | 3993602 | BCGDan_sapMKO_3694 | BCGDan_sapMKO_3694 | 501 |
| 4076984 | 4077730 | Rv3638 | Rv3638 | 747 | 4050005 | 4050751 | BCG_3696 | BCG_3696 | 747 | 4072721 | 4073467 | BCGDan_3770 | BCGDan_3770 | 747 | 3993602 | 3994348 | BCGDan_sapMKO_3695 | BCGDan_sapMKO_3695 | 747 |
| 4077884 | 4078450 | Rv3639c | Rv3639c | 567 | 4050905 | 4051471 | BCG_3697c | BCG_3697c | 567 | 4073621 | 4074187 | BCGDan_3771 | BCGDan_3771 | 567 | 3994502 | 3995068 | BCGDan_sapMKO_3696 | BCGDan_sapMKO_3696 | 567 |
| 4078520 | 4079749 | Rv3640c | Rv3640c | 1230 | 4051541 | 4052770 | BCG_3698c | BCG_3698c | 1230 | 4074257 | 4075486 | BCGDan_3772 | BCGDan_3772 | 1230 | 3995138 | 3996406 | BCGDan_sapMKO_3697 | BCGDan_sapMKO_3697 | 1269 |
| 4079925 | 4080560 | fic | Rv3641c | 636 | 4052946 | 4053581 | fic | BCG_3699c | 636 | 4075662 | 4076297 | fic | BCGDan_3773 | 636 | 3996543 | 3997178 | fic | BCGDan_sapMKO_3698 | 636 |
| 4080571 | 4080765 | Rv3642c | Rv3642c | 195 | 4053592 | 4053786 | BCG_3700c | BCG_3700c | 195 | 4076308 | 4076502 | BCGDan_3774 | BCGDan_3774 | 195 | 3997189 | 3997383 | BCGDan_sapMKO_3699 | BCGDan_sapMKO_3699 | 195 |
| 4081160 | 4081351 | Rv3643 | Rv3643 | 192 | 4054181 | 4054372 | BCG_3701 | BCG_3701 | 192 | 4076897 | 4077088 | BCGDan_3775 | BCGDan_3775 | 192 | 3997778 | 3997969 | BCGDan_sapMKO_3700 | BCGDan_sapMKO_3700 | 192 |
| 4081516 | 4082721 | Rv3644c | Rv3644c | 1206 | 4054537 | 4055742 | BCG_3702c | BCG_3702c | 1206 | 4077253 | 4078458 | BCGDan_3777 | BCGDan_3777 | 1206 | 3998134 | 3999342 | BCGDan_sapMKO_3702 | BCGDan_sapMKO_3702 | 1209 |
| 4082807 | 4084456 | Rv3645 | Rv3645 | 1650 | 4055828 | 4057477 | BCG_3703 | BCG_3703 | 1650 | 4078544 | 4080193 | BCGDan_3778 | BCGDan_3778 | 1650 | 3999425 | 4001074 | BCGDan_sapMKO_3703 | BCGDan_sapMKO_3703 | 1650 |
| 4084453 | 4087257 | topA | Rv3646c | 2805 | 4057474 | 4060278 | topA | BCG_3704c | 2805 | 4080190 | 4082994 | topA | BCGDan_3779 | 2805 | 4001071 | 4003875 | topA | BCGDan_sapMKO_3704 | 2805 |
| 4087610 | 4088188 | Rv3647c | Rv3647c | 579 | 4060631 | 4061209 | BCG_3705c | BCG_3705c | 579 | 4083347 | 4083925 | BCGDan_3780 | BCGDan_3780 | 579 | 4004228 | 4004806 | BCGDan_sapMKO_3705 | BCGDan_sapMKO_3705 | 579 |
| 4088328 | 4088531 | cspA | Rv3648c | 204 | 4061349 | 4061552 | cspA | BCG_3706c | 204 | 4084065 | 4084268 | cspA | BCGDan_3781 | 204 | 4004946 | 4005149 | cspA | BCGDan_sapMKO_3706 | 204 |
| 4088781 | 4091096 | Rv3649 | Rv3649 | 2316 | 4061802 | 4064117 | BCG_3707 | BCG_3707 | 2316 | 4084518 | 4086833 | BCGDan_3782 | BCGDan_3782 | 2316 | 4005372 | 4007714 | BCGDan_sapMKO_3707 | BCGDan_sapMKO_3707 | 2343 |
| 4091233 | 4091517 | PE33 | Rv3650 | 285 | 4064254 | 4064538 | PE33 | BCG_3708 | 285 | 4086970 | 4087254 | PE33 | BCGDan_3783 | 285 | 4007851 | 4008135 | PE33 | BCGDan_sapMKO_3708 | 285 |
| 4091841 | 4092878 | Rv3651 | Rv3651 | 1038 | 4064862 | 4065899 | BCG_3709 | BCG_3709 | 1038 | 4087578 | 4088615 | BCGDan_3784 | BCGDan_3784 | 1038 | 4008459 | 4009496 | BCGDan_sapMKO_3709 | BCGDan_sapMKO_3709 | 1038 |
| 4093632 | 4093946 | PE_PGRS60 | Rv3652 | 315 | 4066653 | 4066967 | PE_PGRS60 | BCG_3710 | 315 | 4089369 | 4089683 | PE_PGRS60 | BCGDan_3785 | 315 | 4010250 | 4010564 | PE_PGRS60 | BCGDan_sapMKO_3710 | 315 |
| 4093940 | 4094527 | PE_PGRS61 | Rv3653 | 588 | 4066961 | 4067656 | PE_PGRS61 | BCG_3711 | 696 | 4089677 | 4090372 | PE_PGRS61 | BCGDan_3786 | 696 | 4010558 | 4011253 | PE_PGRS61 | BCGDan_sapMKO_3711 | 696 |
| 4094660 | 4094914 | Rv3654c | Rv3654c | 255 | 4067773 | 4068027 | BCG_3712c | BCG_3712c | 255 | 4090489 | 4090743 | BCGDan_3787 | BCGDan_3787 | 255 | 4011370 | 4011624 | BCGDan_sapMKO_3712 | BCGDan_sapMKO_3712 | 255 |
| 4094923 | 4095300 | Rv3655c | Rv3655c | 378 | 4068113 | 4068412 | BCG_3713c | BCG_3713c | 300 | 4090829 | 4091128 | BCGDan_3788 | BCGDan_3788 | 300 | 4011710 | 4012009 | BCGDan_sapMKO_3713 | BCGDan_sapMKO_3713 | 300 |
| 4095324 | 4095530 | Rv3656c | Rv3656c | 207 | 4068436 | 4068642 | BCG_3714c | BCG_3714c | 207 | 4091152 | 4091358 | BCGDan_3789 | BCGDan_3789 | 207 | 4012033 | 4012239 | BCGDan_sapMKO_3714 | BCGDan_sapMKO_3714 | 207 |
| 4095540 | 4096115 | Rv3657c | Rv3657c | 576 | 4068652 | 4069227 | BCG_3715c | BCG_3715c | 576 | 4091368 | 4091943 | BCGDan_3790 | BCGDan_3790 | 576 | 4012249 | 4012851 | BCGDan_sapMKO_3715 | BCGDan_sapMKO_3715 | 603 |
| 4096139 | 4096939 | Rv3658c | Rv3658c | 801 | 4069251 | 4070051 | BCG_3716c | BCG_3716c | 801 | 4091967 | 4092767 | BCGDan_3791 | BCGDan_3791 | 801 | 4012848 | 4013648 | BCGDan_sapMKO_3716 | BCGDan_sapMKO_3716 | 801 |
| 4096936 | 4097994 | Rv3659c | Rv3659c | 1059 | 4070048 | 4071106 | BCG_3717c | BCG_3717c | 1059 | 4092764 | 4093822 | BCGDan_3792 | BCGDan_3792 | 1059 | 4013645 | 4014808 | BCGDan_sapMKO_3717 | BCGDan_sapMKO_3717 | 1164 |
| 4098096 | 4099148 | Rv3660c | Rv3660c | 1053 | 4071208 | 4072260 | BCG_3718c | BCG_3718c | 1053 | 4093924 | 4094976 | BCGDan_3793 | BCGDan_3793 | 1053 | 4014805 | 4015878 | BCGDan_sapMKO_3718 | BCGDan_sapMKO_3718 | 1074 |
| 4099647 | 4100510 | Rv3661 | Rv3661 | 864 | 4072760 | 4073623 | BCG_3719 | BCG_3719 | 864 | 4095476 | 4096339 | BCGDan_3794 | BCGDan_3794 | 864 | 4016357 | 4017220 | BCGDan_sapMKO_3719 | BCGDan_sapMKO_3719 | 864 |
| 4101265 | 4102035 | Rv3662c | Rv3662c | 771 | 4074378 | 4075148 | BCG_3720c | BCG_3720c | 771 | 4097094 | 4097864 | BCGDan_3795 | BCGDan_3795 | 771 | 4017975 | 4018745 | BCGDan_sapMKO_3720 | BCGDan_sapMKO_3720 | 771 |
| 4102032 | 4103678 | dppD | Rv3663c | 1647 | 4075145 | 4076791 | dppD | BCG_3721c | 1647 | 4097861 | 4099507 | dppD | BCGDan_3796 | 1647 | 4018742 | 4020397 | dppD | BCGDan_sapMKO_3721 | 1656 |
| 4103675 | 4104475 | dppC | Rv3664c | 801 | 4076788 | 4077588 | dppC | BCG_3722c | 801 | 4099504 | 4100304 | dppC | BCGDan_3797 | 801 | 4020385 | 4021248 | dppC | BCGDan_sapMKO_3722 | 864 |
| 4104531 | 4105457 | dppB | Rv3665c | 927 | 4077644 | 4078570 | dppB | BCG_3723c | 927 | 4100360 | 4101286 | dppB | BCGDan_3798 | 927 | 4021241 | 4022167 | dppB | BCGDan_sapMKO_3723 | 927 |
| 4105459 | 4107084 | dppA | Rv3666c | 1626 | 4078572 | 4080197 | dppA | BCG_3724c | 1626 | 4101288 | 4102913 | dppA | BCGDan_3799 | 1626 | 4022169 | 4023794 | dppA | BCGDan_sapMKO_3724 | 1626 |
| 4107792 | 4109747 | acs | Rv3667 | 1956 | 4080905 | 4082860 | acs | BCG_3725 | 1956 | 4103621 | 4105576 | acs | BCGDan_3800 | 1956 | 4024502 | 4026457 | acs | BCGDan_sapMKO_3725 | 1956 |
| 4109783 | 4110481 | Rv3668c | Rv3668c | 699 | 4082896 | 4083594 | BCG_3726c | BCG_3726c | 699 | 4105612 | 4106310 | BCGDan_3801 | BCGDan_3801 | 699 | 4026493 | 4027191 | BCGDan_sapMKO_3726 | BCGDan_sapMKO_3726 | 699 |
| 4110827 | 4111345 | Rv3669 | Rv3669 | 519 | 4083940 | 4084458 | BCG_3727 | BCG_3727 | 519 | 4106656 | 4107174 | BCGDan_3802 | BCGDan_3802 | 519 | 4027537 | 4028055 | BCGDan_sapMKO_3727 | BCGDan_sapMKO_3727 | 519 |
| 4111346 | 4112329 | ephE | Rv3670 | 984 | 4084459 | 4085442 | ephE | BCG_3728 | 984 | 4107175 | 4108158 | ephE | BCGDan_3803 | 984 | 4028056 | 4029039 | ephE | BCGDan_sapMKO_3728 | 984 |
| 4112322 | 4113515 | Rv3671c | Rv3671c | 1194 | 4085435 | 4086628 | BCG_3729c | BCG_3729c | 1194 | 4108151 | 4109344 | BCGDan_3804 | BCGDan_3804 | 1194 | 4029032 | 4030234 | BCGDan_sapMKO_3729 | BCGDan_sapMKO_3729 | 1203 |
| 4113521 | 4114342 | Rv3672c | Rv3672c | 822 | 4086634 | 4087455 | BCG_3730c | BCG_3730c | 822 | 4109350 | 4110171 | BCGDan_3805 | BCGDan_3805 | 822 | 4030231 | 4031052 | BCGDan_sapMKO_3730 | BCGDan_sapMKO_3730 | 822 |
| 4114474 | 4115157 | Rv3673c | Rv3673c | 684 | 4087587 | 4088270 | BCG_3731c | BCG_3731c | 684 | 4110303 | 4110986 | BCGDan_3806 | BCGDan_3806 | 684 | 4031184 | 4031831 | BCGDan_sapMKO_3731 | BCGDan_sapMKO_3731 | 648 |
| 4115157 | 4115894 | nth | Rv3674c | 738 | 4088270 | 4089007 | nth | BCG_3732c | 738 | 4110986 | 4111723 | nth | BCGDan_3807 | 738 | 4031867 | 4032547 | nth | BCGDan_sapMKO_3732 | 681 |
| 4116002 | 4116379 | Rv3675 | Rv3675 | 378 | 4089115 | 4089492 | BCG_3733 | BCG_3733 | 378 | 4111831 | 4112208 | BCGDan_3808 | BCGDan_3808 | 378 | 4032727 | 4033089 | BCGDan_sapMKO_3733 | BCGDan_sapMKO_3733 | 363 |
| 4116478 | 4117152 | crp | Rv3676 | 675 | 4089591 | 4090265 | BCG_3734 | BCG_3734 | 675 | 4112307 | 4112981 | BCGDan_3809 | BCGDan_3809 | 675 | 4033188 | 4033862 | BCGDan_sapMKO_3734 | BCGDan_sapMKO_3734 | 675 |
| 4117258 | 4118052 | Rv3677c | Rv3677c | 795 | 4090371 | 4091165 | BCG_3735c | BCG_3735c | 795 | 4113087 | 4113881 | BCGDan_3810 | BCGDan_3810 | 795 | 4033968 | 4034762 | BCGDan_sapMKO_3735 | BCGDan_sapMKO_3735 | 795 |
| 4118059 | 4118514 | Rv3678c | Rv3678c | 456 | 4091172 | 4091627 | BCG_3736c | BCG_3736c | 456 | 4113888 | 4114343 | BCGDan_3811 | BCGDan_3811 | 456 | 4034769 | 4035176 | BCGDan_sapMKO_3736 | BCGDan_sapMKO_3736 | 408 |
| 4118530 | 4118691 | Rv3678A | Rv3678A | 162 | 4091643 | 4091804 | BCG_3737c | BCG_3737c | 162 | 4114359 | 4114520 | BCGDan_3812 | BCGDan_3812 | 162 | 4035240 | 4035401 | BCGDan_sapMKO_3737 | BCGDan_sapMKO_3737 | 162 |
| 4118776 | 4119798 | Rv3679 | Rv3679 | 1023 | 4091889 | 4092911 | BCG_3738 | BCG_3738 | 1023 | 4114605 | 4115627 | BCGDan_3813 | BCGDan_3813 | 1023 | 4035486 | 4036508 | BCGDan_sapMKO_3738 | BCGDan_sapMKO_3738 | 1023 |
| 4119795 | 4120955 | Rv3680 | Rv3680 | 1161 | 4092908 | 4094068 | BCG_3739 | BCG_3739 | 1161 | 4115624 | 4116784 | BCGDan_3814 | BCGDan_3814 | 1161 | 4036532 | 4037665 | BCGDan_sapMKO_3739 | BCGDan_sapMKO_3739 | 1134 |
| 4121198 | 4121554 | whiB4 | Rv3681c | 357 | 4094311 | 4094667 | whiB4 | BCG_3740c | 357 | 4117027 | 4117383 | whiB4 | BCGDan_3815 | 357 | 4037908 | 4038264 | whiB4 | BCGDan_sapMKO_3740 | 357 |
| 4121916 | 4124348 | ponA2 | Rv3682 | 2433 | 4095029 | 4097461 | ponA2 | BCG_3741 | 2433 | 4117745 | 4120177 | ponA2 | BCGDan_3816 | 2433 | 4038626 | 4041058 | ponA2 | BCGDan_sapMKO_3741 | 2433 |
| 4124417 | 4125376 | Rv3683 | Rv3683 | 960 | 4097530 | 4098489 | BCG_3742 | BCG_3742 | 960 | 4120246 | 4121205 | BCGDan_3817 | BCGDan_3817 | 960 | 4041127 | 4042086 | BCGDan_sapMKO_3742 | BCGDan_sapMKO_3742 | 960 |
| 4125439 | 4126479 | Rv3684 | Rv3684 | 1041 | 4098552 | 4099592 | BCG_3743 | BCG_3743 | 1041 | 4121268 | 4122308 | BCGDan_3818 | BCGDan_3818 | 1041 | 4042083 | 4043189 | BCGDan_sapMKO_3743 | BCGDan_sapMKO_3743 | 1107 |
| 4127295 | 4128725 | cyp137 | Rv3685c | 1431 | 4100408 | 4101838 | cyp137 | BCG_3744c | 1431 | 4123124 | 4124554 | cyp137 | BCGDan_3820 | 1431 | 4044005 | 4045366 | cyp137 | BCGDan_sapMKO_3745 | 1362 |
| 4128751 | 4129083 | Rv3686c | Rv3686c | 333 | 4101864 | 4102196 | BCG_3745c | BCG_3745c | 333 | 4124580 | 4124912 | BCGDan_3821 | BCGDan_3821 | 333 | 4045461 | 4045793 | BCGDan_sapMKO_3746 | BCGDan_sapMKO_3746 | 333 |
| 4129323 | 4129691 | rsfB | Rv3687c | 369 | 4102436 | 4102681 | BCG_3746c | BCG_3746c | 246 | 4125152 | 4125397 | BCGDan_3822 | BCGDan_3822 | 246 | 4046033 | 4046278 | BCGDan_sapMKO_3747 | BCGDan_sapMKO_3747 | 246 |
| NA | NA | NA | NA | NA | NA | NA | NA | NA | NA | 4125416 | 4125520 | BCGDan_3823 | BCGDan_3823 | 105 | 4046297 | 4046401 | BCGDan_sapMKO_3748 | BCGDan_sapMKO_3748 | 105 |
| 4129893 | 4130357 | Rv3688c | Rv3688c | 465 | 4103006 | 4103470 | BCG_3747c | BCG_3747c | 465 | 4125722 | 4126186 | BCGDan_3824 | BCGDan_3824 | 465 | 4046603 | 4047022 | BCGDan_sapMKO_3749 | BCGDan_sapMKO_3749 | 420 |

| | | | | | | | | | | | | | | | | | | | |
|---|---|---|---|---|---|---|---|---|---|---|---|---|---|---|---|---|---|---|---|
| 4130357 | 4131712 | Rv3689 | Rv3689 | 1356 | 4103470 | 4104825 | BCG_3748 | BCG_3748 | 1356 | 4126186 | 4127541 | BCGDan_3825 | BCGDan_3825 | 1356 | 4047067 | 4048422 | BCGDan_sapMKO_3750 | BCGDan_sapMKO_3750 | 1356 |
| 4131739 | 4132392 | Rv3690 | Rv3690 | 654 | 4104852 | 4105505 | BCG_3749 | BCG_3749 | 654 | 4127568 | 4128221 | BCGDan_3826 | BCGDan_3826 | 654 | 4048449 | 4049102 | BCGDan_sapMKO_3751 | BCGDan_sapMKO_3751 | 654 |
| 4132518 | 4133519 | Rv3691 | Rv3691 | 1002 | 4105631 | 4106632 | BCG_3750 | BCG_3750 | 1002 | 4128347 | 4129348 | BCGDan_3827 | BCGDan_3827 | 1002 | 4049084 | 4050229 | BCGDan_sapMKO_3752 | BCGDan_sapMKO_3752 | 1146 |
| 4133516 | 4134592 | moxR2 | Rv3692 | 1077 | 4106629 | 4107705 | moxR2 | BCG_3751 | 1077 | 4129345 | 4130421 | moxR2 | BCGDan_3828 | 1077 | 4050379 | 4051302 | moxR2 | BCGDan_sapMKO_3753 | 924 |
| 4134726 | 4136048 | Rv3693 | Rv3693 | 1323 | 4107839 | 4109161 | BCG_3752 | BCG_3752 | 1323 | 4130555 | 4131877 | BCGDan_3829 | BCGDan_3829 | 1323 | 4051436 | 4052758 | BCGDan_sapMKO_3754 | BCGDan_sapMKO_3754 | 1323 |
| 4136122 | 4137114 | Rv3694c | Rv3694c | 993 | 4109235 | 4110227 | BCG_3753c | BCG_3753c | 993 | 4131951 | 4132943 | BCGDan_3830 | BCGDan_3830 | 993 | 4052832 | 4053824 | BCGDan_sapMKO_3755 | BCGDan_sapMKO_3755 | 993 |
| 4137206 | 4138138 | Rv3695 | Rv3695 | 933 | 4110319 | 4111251 | BCG_3754 | BCG_3754 | 933 | 4133035 | 4133967 | BCGDan_3831 | BCGDan_3831 | 933 | 4053916 | 4054848 | BCGDan_sapMKO_3756 | BCGDan_sapMKO_3756 | 933 |
| 4138202 | 4139755 | glpK | Rv3696c | 1554 | 4111315 | 4112871 | glpK | BCG_3755c | 1557 | 4134031 | 4135587 | glpK | BCGDan_3832 | 1557 | 4054912 | 4056471 | glpK | BCGDan_sapMKO_3757 | 1560 |
| 4139805 | 4140242 | vapC48 | Rv3697c | 438 | 4112921 | 4113358 | BCG_3756c | BCG_3756c | 438 | 4135637 | 4136074 | BCGDan_3833 | BCGDan_3833 | 438 | 4056521 | 4056958 | BCGDan_sapMKO_3758 | BCGDan_sapMKO_3758 | 438 |
| 4140239 | 4140463 | vapB48 | Rv3697A | 225 | 4113355 | 4113579 | BCG_3756A | BCG_3756A | 225 | 4136071 | 4136295 | BCGDan_3834 | BCGDan_3834 | 225 | 4056955 | 4057179 | BCGDan_sapMKO_3759 | BCGDan_sapMKO_3759 | 225 |
| 4140493 | 4142022 | Rv3698 | Rv3698 | 1530 | 4113609 | 4115138 | BCG_3757 | BCG_3757 | 1530 | 4136325 | 4137854 | BCGDan_3835 | BCGDan_3835 | 1530 | 4057200 | 4058738 | BCGDan_sapMKO_3760 | BCGDan_sapMKO_3760 | 1539 |
| 4142044 | 4142745 | Rv3699 | Rv3699 | 702 | 4115160 | 4115861 | BCG_3758 | BCG_3758 | 702 | 4137876 | 4138577 | BCGDan_3836 | BCGDan_3836 | 702 | 4058775 | 4059461 | BCGDan_sapMKO_3761 | BCGDan_sapMKO_3761 | 687 |
| 4142748 | 4143920 | Rv3700c | Rv3700c | 1173 | 4115864 | 4117036 | BCG_3759c | BCG_3759c | 1173 | 4138580 | 4139752 | BCGDan_3837 | BCGDan_3837 | 1173 | 4059464 | 4060651 | BCGDan_sapMKO_3762 | BCGDan_sapMKO_3762 | 1188 |
| 4143951 | 4144916 | Rv3701c | Rv3701c | 966 | 4117067 | 4118032 | BCG_3760c | BCG_3760c | 966 | 4139783 | 4140748 | BCGDan_3838 | BCGDan_3838 | 966 | 4060667 | 4061626 | BCGDan_sapMKO_3763 | BCGDan_sapMKO_3763 | 960 |
| 4144913 | 4145614 | Rv3702c | Rv3702c | 702 | 4118029 | 4118730 | BCG_3761c | BCG_3761c | 702 | 4140745 | 4141446 | BCGDan_3839 | BCGDan_3839 | 702 | 4061629 | 4062330 | BCGDan_sapMKO_3764 | BCGDan_sapMKO_3764 | 702 |
| 4145614 | 4146891 | Rv3703c | Rv3703c | 1278 | 4118730 | 4120007 | BCG_3762c | BCG_3762c | 1278 | 4141446 | 4142723 | BCGDan_3840 | BCGDan_3840 | 1278 | 4062330 | 4063607 | BCGDan_sapMKO_3765 | BCGDan_sapMKO_3765 | 1278 |
| 4146888 | 4148186 | gshA | Rv3704c | 1299 | 4120004 | 4121302 | gshA | BCG_3763c | 1299 | 4142720 | 4144018 | gshA | BCGDan_3841 | 1299 | 4063604 | 4064902 | gshA | BCGDan_sapMKO_3766 | 1299 |
| 4148318 | 4148962 | Rv3705c | Rv3705c | 645 | 4121434 | 4122078 | BCG_3764c | BCG_3764c | 645 | 4144150 | 4144794 | BCGDan_3842 | BCGDan_3842 | 645 | 4065034 | 4065678 | BCGDan_sapMKO_3767 | BCGDan_sapMKO_3767 | 645 |
| 4149091 | 4149480 | Rv3705A | Rv3705A | 390 | 4122207 | 4122596 | BCG_3765c | BCG_3765c | 390 | 4144923 | 4145312 | BCGDan_3843 | BCGDan_3843 | 390 | 4065807 | 4066196 | BCGDan_sapMKO_3768 | BCGDan_sapMKO_3768 | 390 |
| 4149591 | 4149911 | Rv3706c | Rv3706c | 321 | 4122707 | 4123027 | BCG_3766c | BCG_3766c | 321 | 4145423 | 4145743 | BCGDan_3844 | BCGDan_3844 | 321 | 4066307 | 4066627 | BCGDan_sapMKO_3769 | BCGDan_sapMKO_3769 | 321 |
| 4150030 | 4151040 | Rv3707c | Rv3707c | 1011 | 4123146 | 4124156 | BCG_3767c | BCG_3767c | 1011 | 4145862 | 4146872 | BCGDan_3845 | BCGDan_3845 | 1011 | 4066746 | 4067894 | BCGDan_sapMKO_3770 | BCGDan_sapMKO_3770 | 1149 |
| 4151180 | 4152217 | asd | Rv3708c | 1038 | 4124296 | 4125333 | asd | BCG_3768c | 1038 | 4147012 | 4148049 | asd | BCGDan_3846 | 1038 | 4067896 | 4068933 | asd | BCGDan_sapMKO_3771 | 1038 |
| 4152218 | 4153483 | ask | Rv3709c | 1266 | 4125334 | 4126599 | ask | BCG_3769c | 1266 | 4148050 | 4149315 | ask | BCGDan_3847 | 1266 | 4068934 | 4070199 | ask | BCGDan_sapMKO_3772 | 1266 |
| 4153740 | 4155674 | leuA | Rv3710 | 1935 | 4126856 | 4129360 | leuA | BCG_3770 | 2505 | 4149572 | 4152076 | leuA | BCGDan_3848 | 2505 | 4070456 | 4072960 | leuA | BCGDan_sapMKO_3773 | 2505 |
| 4155740 | 4156729 | dnaQ | Rv3711c | 990 | 4129426 | 4130415 | dnaQ | BCG_3771c | 990 | 4152142 | 4153131 | dnaQ | BCGDan_3849 | 990 | 4073026 | 4074015 | dnaQ | BCGDan_sapMKO_3774 | 990 |
| 4156981 | 4158222 | Rv3712 | Rv3712 | 1242 | 4130549 | 4131790 | BCG_3772 | BCG_3772 | 1242 | 4153265 | 4154506 | BCGDan_3850 | BCGDan_3850 | 1242 | 4074149 | 4075390 | BCGDan_sapMKO_3775 | BCGDan_sapMKO_3775 | 1242 |
| 4158227 | 4158922 | cobQ2 | Rv3713 | 696 | 4131795 | 4132490 | cobQ2 | BCG_3773 | 696 | 4154511 | 4155206 | cobQ2 | BCGDan_3851 | 696 | 4075383 | 4076090 | cobQ2 | BCGDan_sapMKO_3776 | 708 |
| 4158931 | 4159821 | Rv3714c | Rv3714c | 891 | 4132499 | 4133389 | BCG_3774c | BCG_3774c | 891 | 4155215 | 4156105 | BCGDan_3852 | BCGDan_3852 | 891 | 4076099 | 4076989 | BCGDan_sapMKO_3777 | BCGDan_sapMKO_3777 | 891 |
| 4159889 | 4160500 | recR | Rv3715c | 612 | 4133457 | 4134068 | recR | BCG_3775c | 612 | 4156173 | 4156784 | recR | BCGDan_3853 | 612 | 4077057 | 4077668 | recR | BCGDan_sapMKO_3778 | 612 |
| 4160512 | 4160913 | Rv3716c | Rv3716c | 402 | 4134080 | 4134481 | BCG_3776c | BCG_3776c | 402 | 4156796 | 4157197 | BCGDan_3854 | BCGDan_3854 | 402 | 4077680 | 4078081 | BCGDan_sapMKO_3779 | BCGDan_sapMKO_3779 | 402 |
| 4161048 | 4161773 | Rv3717 | Rv3717 | 726 | 4134616 | 4135341 | BCG_3777 | BCG_3777 | 726 | 4157332 | 4158057 | BCGDan_3855 | BCGDan_3855 | 726 | 4078216 | 4078941 | BCGDan_sapMKO_3780 | BCGDan_sapMKO_3780 | 726 |
| 4161815 | 4162258 | Rv3718c | Rv3718c | 444 | 4135383 | 4135826 | BCG_3778c | BCG_3778c | 444 | 4158099 | 4158542 | BCGDan_3856 | BCGDan_3856 | 444 | 4078983 | 4079426 | BCGDan_sapMKO_3781 | BCGDan_sapMKO_3781 | 444 |
| 4162306 | 4163718 | Rv3719 | Rv3719 | 1413 | 4135874 | 4137286 | BCG_3779 | BCG_3779 | 1413 | 4158590 | 4160002 | BCGDan_3857 | BCGDan_3857 | 1413 | 4079561 | 4080886 | BCGDan_sapMKO_3782 | BCGDan_sapMKO_3782 | 1326 |
| 4163736 | 4164998 | Rv3720 | Rv3720 | 1263 | 4137304 | 4138566 | BCG_3780 | BCG_3780 | 1263 | 4160020 | 4161282 | BCGDan_3858 | BCGDan_3858 | 1263 | 4080904 | 4082166 | BCGDan_sapMKO_3783 | BCGDan_sapMKO_3783 | 1263 |
| 4164995 | 4166731 | dnaZX | Rv3721c | 1737 | 4138563 | 4140299 | dnaZX | BCG_3781c | 1737 | 4161279 | 4163015 | dnaZX | BCGDan_3859 | 1737 | 4082163 | 4083899 | dnaZX | BCGDan_sapMKO_3784 | 1737 |
| 4166821 | 4168128 | Rv3722c | Rv3722c | 1308 | 4140389 | 4141696 | BCG_3782c | BCG_3782c | 1308 | 4163105 | 4164412 | BCGDan_3860 | BCGDan_3860 | 1308 | 4083989 | 4085215 | BCGDan_sapMKO_3785 | BCGDan_sapMKO_3785 | 1227 |
| 4168536 | 4169300 | Rv3723 | Rv3723 | 765 | 4142104 | 4142868 | BCG_3783 | BCG_3783 | 765 | 4164820 | 4165584 | BCGDan_3862 | BCGDan_3862 | 765 | 4085704 | 4086468 | BCGDan_sapMKO_3787 | BCGDan_sapMKO_3787 | 765 |
| 4169467 | 4169709 | cut5a | Rv3724A | 243 | 4143035 | 4143736 | cut5 | BCG_3784 | 702 | 4165751 | 4166452 | cut5 | BCGDan_3863 | 702 | 4086728 | 4087336 | cut5 | BCGDan_sapMKO_3788 | 609 |
| 4170214 | 4171143 | Rv3725 | Rv3725 | 930 | 4143781 | 4144782 | BCG_3785 | BCG_3785 | 1002 | 4166497 | 4167498 | BCGDan_3864 | BCGDan_3864 | 1002 | 4087381 | 4088382 | BCGDan_sapMKO_3789 | BCGDan_sapMKO_3789 | 1002 |
| 4171421 | 4172614 | Rv3726 | Rv3726 | 1194 | 4144989 | 4146182 | BCG_3786 | BCG_3786 | 1194 | 4167705 | 4168898 | BCGDan_3865 | BCGDan_3865 | 1194 | 4088589 | 4089782 | BCGDan_sapMKO_3790 | BCGDan_sapMKO_3790 | 1194 |
| 4172955 | 4174763 | Rv3727 | Rv3727 | 1809 | 4146523 | 4148331 | BCG_3787 | BCG_3787 | 1809 | 4169239 | 4171047 | BCGDan_3866 | BCGDan_3866 | 1809 | 4090123 | 4091931 | BCGDan_sapMKO_3791 | BCGDan_sapMKO_3791 | 1809 |
| 4174873 | 4178070 | Rv3728 | Rv3728 | 3198 | 4148441 | 4151638 | BCG_3788 | BCG_3788 | 3198 | 4171157 | 4174354 | BCGDan_3867 | BCGDan_3867 | 3198 | 4092164 | 4095238 | BCGDan_sapMKO_3792 | BCGDan_sapMKO_3792 | 3075 |
| 4178285 | 4180615 | Rv3729 | Rv3729 | 2331 | 4151853 | 4154183 | BCG_3789 | BCG_3789 | 2331 | 4174569 | 4176899 | BCGDan_3868 | BCGDan_3868 | 2331 | 4095417 | 4097783 | BCGDan_sapMKO_3793 | BCGDan_sapMKO_3793 | 2367 |
| 4180680 | 4181720 | Rv3730c | Rv3730c | 1041 | 4154248 | 4155288 | BCG_3790c | BCG_3790c | 1041 | 4176964 | 4178004 | BCGDan_3869 | BCGDan_3869 | 1041 | 4097848 | 4098888 | BCGDan_sapMKO_3794 | BCGDan_sapMKO_3794 | 1041 |
| 4181758 | 4182834 | ligC | Rv3731 | 1077 | 4155326 | 4156402 | ligC | BCG_3791 | 1077 | 4178042 | 4179118 | ligC | BCGDan_3870 | 1077 | 4098941 | 4100002 | ligC | BCGDan_sapMKO_3795 | 1062 |
| 4182934 | 4183992 | Rv3732 | Rv3732 | 1059 | 4156502 | 4157560 | BCG_3792 | BCG_3792 | 1059 | 4179218 | 4180276 | BCGDan_3871 | BCGDan_3871 | 1059 | 4100102 | 4101160 | BCGDan_sapMKO_3796 | BCGDan_sapMKO_3796 | 1059 |
| 4184012 | 4184512 | Rv3733c | Rv3733c | 501 | 4157580 | 4158080 | BCG_3793c | BCG_3793c | 501 | 4180296 | 4180796 | BCGDan_3872 | BCGDan_3872 | 501 | 4101180 | 4101680 | BCGDan_sapMKO_3797 | BCGDan_sapMKO_3797 | 501 |
| 4184526 | 4185890 | tgs2 | Rv3734c | 1365 | 4158094 | 4159458 | BCG_3794c | BCG_3794c | 1365 | 4180810 | 4182174 | BCGDan_3873 | BCGDan_3873 | 1365 | 4101694 | 4103058 | BCGDan_sapMKO_3798 | BCGDan_sapMKO_3798 | 1365 |
| 4186089 | 4186577 | Rv3735 | Rv3735 | 489 | 4159657 | 4160145 | BCG_3795 | BCG_3795 | 489 | 4182373 | 4182861 | BCGDan_3874 | BCGDan_3874 | 489 | 4103257 | 4103745 | BCGDan_sapMKO_3799 | BCGDan_sapMKO_3799 | 489 |
| 4186634 | 4187695 | Rv3736 | Rv3736 | 1062 | 4160202 | 4161263 | BCG_3796 | BCG_3796 | 1062 | 4182918 | 4183979 | BCGDan_3875 | BCGDan_3875 | 1062 | 4103802 | 4104863 | BCGDan_sapMKO_3800 | BCGDan_sapMKO_3800 | 1062 |
| 4187699 | 4189288 | Rv3737 | Rv3737 | 1590 | 4161267 | 4162856 | BCG_3797 | BCG_3797 | 1590 | 4183983 | 4185572 | BCGDan_3876 | BCGDan_3876 | 1590 | 4104867 | 4106456 | BCGDan_sapMKO_3801 | BCGDan_sapMKO_3801 | 1590 |
| 4189285 | 4190232 | PPE66 | Rv3738c | 948 | 4162853 | 4163209 | PPE66 | BCG_3798c | 357 | 4185569 | 4185925 | PPE66 | BCGDan_3877 | 357 | 4106453 | 4106752 | PPE66 | BCGDan_sapMKO_3802 | 300 |
| 4190833 | 4192179 | Rv3740c | Rv3740c | 1347 | 4163235 | 4164581 | BCG_3799c | BCG_3799c | 1347 | 4185951 | 4187297 | BCGDan_3878 | BCGDan_3878 | 1347 | 4106835 | 4108190 | BCGDan_sapMKO_3803 | BCGDan_sapMKO_3803 | 1356 |
| 4192179 | 4192853 | Rv3741c | Rv3741c | 675 | 4164581 | 4165255 | BCG_3800c | BCG_3800c | 675 | 4187297 | 4187971 | BCGDan_3879 | BCGDan_3879 | 675 | 4108181 | 4108855 | BCGDan_sapMKO_3804 | BCGDan_sapMKO_3804 | 675 |
| 4192850 | 4193245 | Rv3742c | Rv3742c | 396 | 4165252 | 4165647 | BCG_3801c | BCG_3801c | 396 | 4187968 | 4188363 | BCGDan_3880 | BCGDan_3880 | 396 | 4108852 | 4109247 | BCGDan_sapMKO_3805 | BCGDan_sapMKO_3805 | 396 |
| 4193391 | 4195373 | ctpJ | Rv3743c | 1983 | 4165793 | 4167775 | ctpJ | BCG_3802c | 1983 | 4188509 | 4190491 | ctpJ | BCGDan_3881 | 1983 | 4109393 | 4111327 | ctpJ | BCGDan_sapMKO_3806 | 1935 |
| 4195440 | 4195802 | nmtR | Rv3744 | 363 | 4167842 | 4168204 | BCG_3803 | BCG_3803 | 363 | 4190558 | 4190920 | BCGDan_3882 | BCGDan_3882 | 363 | 4111484 | 4111804 | BCGDan_sapMKO_3807 | BCGDan_sapMKO_3807 | 321 |
| 4195886 | 4196098 | Rv3745c | Rv3745c | 213 | 4168288 | 4168500 | BCG_3804c | BCG_3804c | 213 | 4191004 | 4191216 | BCGDan_3883 | BCGDan_3883 | 213 | 4111888 | 4112100 | BCGDan_sapMKO_3808 | BCGDan_sapMKO_3808 | 213 |
| 4196171 | 4196506 | PE34 | Rv3746c | 336 | 4168573 | 4168908 | PE34 | BCG_3805c | 336 | 4191289 | 4191624 | PE34 | BCGDan_3884 | 336 | 4112173 | 4112508 | PE34 | BCGDan_sapMKO_3809 | 336 |
| 4196724 | 4197107 | Rv3747 | Rv3747 | 384 | 4169126 | 4169509 | BCG_3806 | BCG_3806 | 384 | 4191842 | 4192225 | BCGDan_3885 | BCGDan_3885 | 384 | 4112726 | 4113109 | BCGDan_sapMKO_3810 | BCGDan_sapMKO_3810 | 384 |
| 4197236 | 4197595 | Rv3748 | Rv3748 | 360 | 4169639 | 4169998 | BCG_3807 | BCG_3807 | 360 | 4192355 | 4192714 | BCGDan_3886 | BCGDan_3886 | 360 | 4113239 | 4113598 | BCGDan_sapMKO_3811 | BCGDan_sapMKO_3811 | 360 |
| 4197628 | 4198137 | Rv3749c | Rv3749c | 510 | 4170031 | 4170540 | BCG_3808c | BCG_3808c | 510 | 4192747 | 4193256 | BCGDan_3887 | BCGDan_3887 | 510 | 4113631 | 4114140 | BCGDan_sapMKO_3812 | BCGDan_sapMKO_3812 | 510 |
| 4198205 | 4198597 | Rv3750c | Rv3750c | 393 | 4170608 | 4171000 | BCG_3809c | BCG_3809c | 393 | 4193324 | 4193770 | BCGDan_3888 | BCGDan_3888 | 447 | 4114208 | 4114654 | BCGDan_sapMKO_3813 | BCGDan_sapMKO_3813 | 447 |
| 4198874 | 4199089 | Rv3751 | Rv3751 | 216 | 4171276 | 4171491 | BCG_3810 | BCG_3810 | 216 | 4193992 | 4194207 | BCGDan_3889 | BCGDan_3889 | 216 | 4114876 | 4115091 | BCGDan_sapMKO_3814 | BCGDan_sapMKO_3814 | 216 |
| 4199247 | 4199705 | Rv3752c | Rv3752c | 459 | 4171649 | 4172107 | BCG_3811c | BCG_3811c | 459 | 4194365 | 4194823 | BCGDan_3891 | BCGDan_3891 | 459 | 4115249 | 4115707 | BCGDan_sapMKO_3816 | BCGDan_sapMKO_3816 | 459 |
| 4199721 | 4200221 | Rv3753c | Rv3753c | 501 | 4172123 | 4172644 | BCG_3812c | BCG_3812c | 522 | 4194839 | 4195360 | BCGDan_3892 | BCGDan_3892 | 522 | 4115723 | 4116244 | BCGDan_sapMKO_3817 | BCGDan_sapMKO_3817 | 522 |
| 4200421 | 4201326 | tyrA | Rv3754 | 906 | 4172823 | 4173728 | tyrA | BCG_3813 | 906 | 4195539 | 4196444 | tyrA | BCGDan_3893 | 906 | 4116390 | 4117328 | tyrA | BCGDan_sapMKO_3818 | 939 |
| 4201289 | 4201888 | Rv3755c | Rv3755c | 600 | 4173691 | 4174290 | BCG_3814c | BCG_3814c | 600 | 4196407 | 4197006 | BCGDan_3894 | BCGDan_3894 | 600 | 4117291 | 4117845 | BCGDan_sapMKO_3819 | BCGDan_sapMKO_3819 | 555 |
| 4201894 | 4202613 | proZ | Rv3756c | 720 | 4174296 | 4175015 | proZ | BCG_3815c | 720 | 4197012 | 4197731 | proZ | BCGDan_3895 | 720 | 4117896 | 4118684 | proZ | BCGDan_sapMKO_3820 | 789 |
| 4202610 | 4203299 | proW | Rv3757c | 690 | 4175012 | 4175701 | proW | BCG_3816c | 690 | 4197728 | 4198417 | proW | BCGDan_3896 | 690 | 4118612 | 4119301 | proW | BCGDan_sapMKO_3821 | 690 |
| 4203287 | 4204417 | proV | Rv3758c | 1131 | 4175689 | 4176819 | proV | BCG_3817c | 1131 | 4198405 | 4199535 | proV | BCGDan_3897 | 1131 | 4119289 | 4120419 | proV | BCGDan_sapMKO_3822 | 1131 |
| 4204426 | 4205373 | proX | Rv3759c | 948 | 4176828 | 4177775 | proX | BCG_3818c | 948 | 4199544 | 4200491 | proX | BCGDan_3898 | 948 | 4120428 | 4121459 | proX | BCGDan_sapMKO_3823 | 1032 |
| 4205538 | 4205840 | Rv3760 | Rv3760 | 303 | 4177940 | 4178242 | BCG_3819 | BCG_3819 | 303 | 4200656 | 4200958 | BCGDan_3899 | BCGDan_3899 | 303 | 4121489 | 4121842 | BCGDan_sapMKO_3824 | BCGDan_sapMKO_3824 | 354 |
| 4205862 | 4206917 | fadE36 | Rv3761c | 1056 | 4178264 | 4179319 | fadE36 | BCG_3820c | 1056 | 4200980 | 4202035 | fadE36 | BCGDan_3900 | 1056 | 4121864 | 4122919 | fadE36 | BCGDan_sapMKO_3825 | 1056 |
| 4206996 | 4208876 | Rv3762c | Rv3762c | 1881 | 4179398 | 4181278 | BCG_3821c | BCG_3821c | 1881 | 4202114 | 4203994 | BCGDan_3901 | BCGDan_3901 | 1881 | 4122998 | 4124872 | BCGDan_sapMKO_3826 | BCGDan_sapMKO_3826 | 1875 |
| 4209047 | 4209526 | lpqH | Rv3763 | 480 | 4181449 | 4181928 | lpqH | BCG_3822 | 480 | 4204165 | 4204644 | lpqH | BCGDan_3902 | 480 | 4125049 | 4125528 | lpqH | BCGDan_sapMKO_3827 | 480 |
| 4209582 | 4211009 | tcrY | Rv3764c | 1428 | 4181984 | 4183411 | BCG_3823c | BCG_3823c | 1428 | 4204700 | 4206127 | BCGDan_3903 | BCGDan_3903 | 1428 | 4125584 | 4127011 | BCGDan_sapMKO_3828 | BCGDan_sapMKO_3828 | 1428 |
| 4211080 | 4211784 | tcrX | Rv3765c | 705 | 4183482 | 4184186 | BCG_3824c | BCG_3824c | 705 | 4206198 | 4206902 | BCGDan_3904 | BCGDan_3904 | 705 | 4127082 | 4127822 | BCGDan_sapMKO_3829 | BCGDan_sapMKO_3829 | 741 |
| 4212293 | 4212982 | Rv3766 | Rv3766 | 690 | 4184695 | 4185384 | BCG_3825 | BCG_3825 | 690 | 4207411 | 4208100 | BCGDan_3905 | BCGDan_3905 | 690 | 4128205 | 4128984 | BCGDan_sapMKO_3830 | BCGDan_sapMKO_3830 | 780 |
| 4212996 | 4213940 | Rv3767c | Rv3767c | 945 | 4185398 | 4186342 | BCG_3826c | BCG_3826c | 945 | 4208114 | 4209058 | BCGDan_3906 | BCGDan_3906 | 945 | 4128998 | 4129942 | BCGDan_sapMKO_3831 | BCGDan_sapMKO_3831 | 945 |
| 4214070 | 4214429 | Rv3768 | Rv3768 | 360 | 4186472 | 4186831 | BCG_3827 | BCG_3827 | 360 | 4209188 | 4209547 | BCGDan_3907 | BCGDan_3907 | 360 | 4130024 | 4130431 | BCGDan_sapMKO_3832 | BCGDan_sapMKO_3832 | 408 |
| 4214615 | 4214887 | Rv3769 | Rv3769 | 273 | 4187017 | 4187289 | BCG_3828 | BCG_3828 | 273 | 4209733 | 4210005 | BCGDan_3908 | BCGDan_3908 | 273 | 4130617 | 4130889 | BCGDan_sapMKO_3833 | BCGDan_sapMKO_3833 | 273 |
| 4215200 | 4215775 | Rv3770c | Rv3770c | 576 | 4187602 | 4188177 | BCG_3829c | BCG_3829c | 576 | 4210318 | 4210893 | BCGDan_3909 | BCGDan_3909 | 576 | 4131202 | 4131747 | BCGDan_sapMKO_3834 | BCGDan_sapMKO_3834 | 546 |
| 4215881 | 4216063 | Rv3770A | Rv3770A | 183 | 4188283 | 4188465 | BCG_3830c | BCG_3830c | 183 | 4210999 | 4211181 | BCGDan_3910 | BCGDan_3910 | 183 | 4131883 | 4132065 | BCGDan_sapMKO_3835 | BCGDan_sapMKO_3835 | 183 |
| 4216078 | 4216269 | Rv3770B | Rv3770B | 192 | 4188480 | 4188671 | BCG_3831c | BCG_3831c | 192 | 4211196 | 4211387 | BCGDan_3911 | BCGDan_3911 | 192 | 4132080 | 4132271 | BCGDan_sapMKO_3836 | BCGDan_sapMKO_3836 | 192 |
| 4216404 | 4216730 | Rv3771c | Rv3771c | 327 | 4188806 | 4189132 | BCG_3832c | BCG_3832c | 327 | 4211522 | 4211848 | BCGDan_3912 | BCGDan_3912 | 327 | 4132406 | 4132732 | BCGDan_sapMKO_3837 | BCGDan_sapMKO_3837 | 327 |

| | | | | | | | | | | | | | | | | | | | |
|---|---|---|---|---|---|---|---|---|---|---|---|---|---|---|---|---|---|---|---|
| 4217134 | 4218195 | hisC2 | Rv3772 | 1062 | 4189536 | 4190597 | hisC2 | BCG_3833 | 1062 | 4212252 | 4213313 | hisC2 | BCGDan_3915 | 1062 | 4133136 | 4134197 | hisC2 | BCGDan_sapMKO_3840 | 1062 |
| 4218241 | 4218825 | Rv3773c | Rv3773c | 585 | 4190643 | 4190987 | BCG_3834c | BCG_3834c | 345 | 4213359 | 4213703 | BCGDan_3916 | BCGDan_3916 | 345 | 4134243 | 4134644 | BCGDan_sapMKO_3841 | BCGDan_sapMKO_3841 | 402 |
| NA | NA | NA | NA | NA | 4190992 | 4191228 | BCG_3835c | BCG_3835c | 237 | 4213708 | 4213944 | BCGDan_3917 | BCGDan_3917 | 237 | 4134592 | 4134828 | BCGDan_sapMKO_3842 | BCGDan_sapMKO_3842 | 237 |
| 4218849 | 4219673 | echA21 | Rv3774 | 825 | 4191252 | 4192076 | echA21 | BCG_3836 | 825 | 4213968 | 4214792 | echA21 | BCGDan_3918 | 825 | 4134792 | 4135676 | echA21 | BCGDan_sapMKO_3843 | 885 |
| 4219685 | 4220932 | lipE | Rv3775 | 1248 | 4192088 | 4193335 | lipE | BCG_3837 | 1248 | 4214804 | 4216051 | lipE | BCGDan_3919 | 1248 | 4135688 | 4136935 | lipE | BCGDan_sapMKO_3844 | 1248 |
| 4221089 | 4222648 | Rv3776 | Rv3776 | 1560 | 4193492 | 4195051 | BCG_3838 | BCG_3838 | 1560 | 4216208 | 4217767 | BCGDan_3920 | BCGDan_3920 | 1560 | 4137092 | 4138651 | BCGDan_sapMKO_3845 | BCGDan_sapMKO_3845 | 1560 |
| 4222694 | 4223680 | Rv3777 | Rv3777 | 987 | 4195097 | 4196083 | BCG_3839 | BCG_3839 | 987 | 4217813 | 4218799 | BCGDan_3922 | BCGDan_3922 | 987 | 4138697 | 4139683 | BCGDan_sapMKO_3847 | BCGDan_sapMKO_3847 | 987 |
| 4223699 | 4224895 | Rv3778c | Rv3778c | 1197 | 4196102 | 4197298 | BCG_3840c | BCG_3840c | 1197 | 4218818 | 4220014 | BCGDan_3923 | BCGDan_3923 | 1197 | 4139702 | 4140898 | BCGDan_sapMKO_3848 | BCGDan_sapMKO_3848 | 1197 |
| 4224985 | 4226985 | Rv3779 | Rv3779 | 2001 | 4197388 | 4199388 | BCG_3841 | BCG_3841 | 2001 | 4220104 | 4222104 | BCGDan_3924 | BCGDan_3924 | 2001 | 4140988 | 4142988 | BCGDan_sapMKO_3849 | BCGDan_sapMKO_3849 | 2001 |
| 4226989 | 4227525 | Rv3780 | Rv3780 | 537 | 4199392 | 4199928 | BCG_3842 | BCG_3842 | 537 | 4222108 | 4222644 | BCGDan_3925 | BCGDan_3925 | 537 | 4143004 | 4143528 | BCGDan_sapMKO_3850 | BCGDan_sapMKO_3850 | 525 |
| 4227529 | 4228350 | rfbE | Rv3781 | 822 | 4199932 | 4200753 | rbfE | BCG_3843 | 822 | 4222648 | 4223469 | rbfE | BCGDan_3926 | 822 | 4143532 | 4144353 | rbfE | BCGDan_sapMKO_3851 | 822 |
| 4228347 | 4229261 | glfT1 | Rv3782 | 915 | 4200750 | 4201664 | BCG_3844 | BCG_3844 | 915 | 4223466 | 4224380 | BCGDan_3927 | BCGDan_3927 | 915 | 4144350 | 4145264 | BCGDan_sapMKO_3852 | BCGDan_sapMKO_3852 | 915 |
| 4229258 | 4230100 | rfbD | Rv3783 | 843 | 4201661 | 4202503 | rfbD | BCG_3845 | 843 | 4224377 | 4225219 | rfbD | BCGDan_3928 | 843 | 4145261 | 4146103 | rfbD | BCGDan_sapMKO_3853 | 843 |
| 4230256 | 4231236 | Rv3784 | Rv3784 | 981 | 4202659 | 4203639 | rfbB | BCG_3846 | 981 | 4225375 | 4226355 | rfbB | BCGDan_3929 | 981 | 4146259 | 4147239 | rfbB | BCGDan_sapMKO_3854 | 981 |
| 4231320 | 4232393 | Rv3785 | Rv3785 | 1074 | 4203723 | 4204796 | BCG_3847 | BCG_3847 | 1074 | 4226439 | 4227512 | BCGDan_3930 | BCGDan_3930 | 1074 | 4147356 | 4148396 | BCGDan_sapMKO_3855 | BCGDan_sapMKO_3855 | 1041 |
| 4232374 | 4233597 | Rv3786c | Rv3786c | 1224 | 4204777 | 4206000 | BCG_3848c | BCG_3848c | 1224 | 4227493 | 4228716 | BCGDan_3931 | BCGDan_3931 | 1224 | 4148377 | 4149600 | BCGDan_sapMKO_3856 | BCGDan_sapMKO_3856 | 1224 |
| 4233610 | 4234536 | Rv3787c | Rv3787c | 927 | 4206013 | 4206939 | BCG_3849c | BCG_3849c | 927 | 4228729 | 4229655 | BCGDan_3932 | BCGDan_3932 | 927 | 4149613 | 4150593 | BCGDan_sapMKO_3857 | BCGDan_sapMKO_3857 | 981 |
| 4234780 | 4235265 | Rv3788 | Rv3788 | 486 | 4207183 | 4207668 | BCG_3850 | BCG_3850 | 486 | 4229899 | 4230384 | BCGDan_3933 | BCGDan_3933 | 486 | 4150783 | 4151268 | BCGDan_sapMKO_3858 | BCGDan_sapMKO_3858 | 486 |
| 4235374 | 4235739 | Rv3789 | Rv3789 | 366 | 4207777 | 4208142 | BCG_3851 | BCG_3851 | 366 | 4230493 | 4230858 | BCGDan_3934 | BCGDan_3934 | 366 | 4151314 | 4151742 | BCGDan_sapMKO_3859 | BCGDan_sapMKO_3859 | 429 |
| 4235779 | 4237164 | dprE1 | Rv3790 | 1386 | 4208182 | 4209567 | BCG_3852 | BCG_3852 | 1386 | 4230898 | 4232283 | BCGDan_3935 | BCGDan_3935 | 1386 | 4151782 | 4153167 | BCGDan_sapMKO_3860 | BCGDan_sapMKO_3860 | 1386 |
| 4237165 | 4237929 | dprE2 | Rv3791 | 765 | 4209568 | 4210332 | BCG_3853 | BCG_3853 | 765 | 4232284 | 4233048 | BCGDan_3936 | BCGDan_3936 | 765 | 4153168 | 4153932 | BCGDan_sapMKO_3861 | BCGDan_sapMKO_3861 | 765 |
| 4237932 | 4239863 | aftA | Rv3792 | 1932 | 4210335 | 4212266 | BCG_3854 | BCG_3854 | 1932 | 4233051 | 4234982 | BCGDan_3937 | BCGDan_3937 | 1932 | 4154007 | 4155866 | BCGDan_sapMKO_3862 | BCGDan_sapMKO_3862 | 1860 |
| 4239863 | 4243147 | embC | Rv3793 | 3285 | 4212266 | 4215550 | embC | BCG_3855 | 3285 | 4234982 | 4238266 | embC | BCGDan_3938 | 3285 | 4155920 | 4159150 | embC | BCGDan_sapMKO_3863 | 3231 |
| 4243233 | 4246517 | embA | Rv3794 | 3285 | 4215636 | 4218920 | embA | BCG_3856 | 3285 | 4238352 | 4241636 | embA | BCGDan_3939 | 3285 | 4159236 | 4162520 | embA | BCGDan_sapMKO_3864 | 3285 |
| 4246514 | 4249810 | embB | Rv3795 | 3297 | 4218917 | 4222213 | embB | BCG_3857 | 3297 | 4241633 | 4244929 | embB | BCGDan_3940 | 3297 | 4162601 | 4165813 | embB | BCGDan_sapMKO_3865 | 3213 |
| 4249878 | 4251005 | Rv3796 | Rv3796 | 1128 | 4222281 | 4223408 | BCG_3858 | BCG_3858 | 1128 | 4244997 | 4246124 | BCGDan_3941 | BCGDan_3941 | 1128 | 4166049 | 4167008 | BCGDan_sapMKO_3866 | BCGDan_sapMKO_3866 | 960 |
| 4251085 | 4252866 | fadE35 | Rv3797 | 1782 | 4223488 | 4225269 | fadE35 | BCG_3859 | 1782 | 4246204 | 4247985 | fadE35 | BCGDan_3942 | 1782 | 4167088 | 4168869 | fadE35 | BCGDan_sapMKO_3867 | 1782 |
| 4252993 | 4254327 | Rv3798 | Rv3798 | 1335 | 4225396 | 4226730 | BCG_3860 | BCG_3860 | 1335 | 4248112 | 4249446 | BCGDan_3943 | BCGDan_3943 | 1335 | 4168996 | 4170330 | BCGDan_sapMKO_3868 | BCGDan_sapMKO_3868 | 1335 |
| 4254380 | 4255948 | accD4 | Rv3799c | 1569 | 4226783 | 4228351 | accD4 | BCG_3861c | 1569 | 4249499 | 4251067 | accD4 | BCGDan_3944 | 1569 | 4170383 | 4171951 | accD4 | BCGDan_sapMKO_3869 | 1569 |
| 4255945 | 4261146 | pks13 | Rv3800c | 5202 | 4228348 | 4233549 | pks13 | BCG_3862c | 5202 | 4251064 | 4256265 | pks13 | BCGDan_3945 | 5202 | 4171948 | 4177149 | pks13 | BCGDan_sapMKO_3870 | 5202 |
| 4261153 | 4263066 | fadD32 | Rv3801c | 1914 | 4233556 | 4235469 | fadD32 | BCG_3863c | 1914 | 4256272 | 4258185 | fadD32 | BCGDan_3946 | 1914 | 4177156 | 4179045 | fadD32 | BCGDan_sapMKO_3871 | 1890 |
| 4263355 | 4264365 | Rv3802c | Rv3802c | 1011 | 4235758 | 4236768 | BCG_3864c | BCG_3864c | 1011 | 4258474 | 4259484 | BCGDan_3947 | BCGDan_3947 | 1011 | 4179358 | 4180368 | BCGDan_sapMKO_3872 | BCGDan_sapMKO_3872 | 1011 |
| 4264563 | 4265462 | fbpD | Rv3803c | 900 | 4236966 | 4237865 | fbpD | BCG_3865c | 900 | 4259682 | 4260581 | fbpD | BCGDan_3948 | 900 | 4180566 | 4181465 | fbpD | BCGDan_sapMKO_3873 | 900 |
| 4265642 | 4266658 | fbpA | Rv3804c | 1017 | 4238045 | 4239061 | fbpA | BCG_3866c | 1017 | 4260761 | 4261777 | fbpA | BCGDan_3949 | 1017 | 4181645 | 4182622 | fbpA | BCGDan_sapMKO_3874 | 978 |
| 4266953 | 4268836 | aftB | Rv3805c | 1884 | 4239356 | 4241239 | BCG_3867c | BCG_3867c | 1884 | 4262072 | 4263955 | BCGDan_3950 | BCGDan_3950 | 1884 | 4182956 | 4184899 | BCGDan_sapMKO_3875 | BCGDan_sapMKO_3875 | 1944 |
| 4268925 | 4269833 | ubiA | Rv3806c | 909 | 4241328 | 4242236 | BCG_3868c | BCG_3868c | 909 | 4264044 | 4264952 | BCGDan_3951 | BCGDan_3951 | 909 | 4184928 | 4185836 | BCGDan_sapMKO_3876 | BCGDan_sapMKO_3876 | 909 |
| 4269840 | 4270337 | Rv3807c | Rv3807c | 498 | 4242243 | 4242740 | BCG_3869c | BCG_3869c | 498 | 4264959 | 4265456 | BCGDan_3952 | BCGDan_3952 | 498 | 4185843 | 4186376 | BCGDan_sapMKO_3877 | BCGDan_sapMKO_3877 | 534 |
| 4270366 | 4272279 | glfT2 | Rv3808c | 1914 | 4242769 | 4244682 | BCG_3870c | BCG_3870c | 1914 | 4265485 | 4267398 | BCGDan_3953 | BCGDan_3953 | 1914 | 4186369 | 4188282 | BCGDan_sapMKO_3878 | BCGDan_sapMKO_3878 | 1914 |
| 4272276 | 4273475 | glf | Rv3809c | 1200 | 4244679 | 4245878 | glf | BCG_3871c | 1200 | 4267395 | 4268594 | glf | BCGDan_3954 | 1200 | 4188279 | 4189478 | glf | BCGDan_sapMKO_3879 | 1200 |
| 4273739 | 4274593 | pirG | Rv3810 | 855 | 4246142 | 4246996 | pirG | BCG_3872 | 855 | 4268858 | 4269712 | pirG | BCGDan_3955 | 855 | 4189778 | 4190596 | pirG | BCGDan_sapMKO_3880 | 819 |
| 4274798 | 4276417 | Rv3811 | Rv3811 | 1620 | 4247201 | 4248820 | BCG_3873 | BCG_3873 | 1620 | 4269917 | 4271536 | BCGDan_3956 | BCGDan_3956 | 1620 | 4190759 | 4192420 | BCGDan_sapMKO_3881 | BCGDan_sapMKO_3881 | 1662 |
| 4276571 | 4278085 | PE_PGRS62 | Rv3812 | 1515 | 4248974 | 4250488 | PE_PGRS62 | BCG_3874 | 1515 | 4271690 | 4273204 | PE_PGRS62 | BCGDan_3957 | 1515 | 4192574 | 4194088 | PE_PGRS62 | BCGDan_sapMKO_3882 | 1515 |
| 4278394 | 4279215 | Rv3813c | Rv3813c | 822 | 4250797 | 4251618 | BCG_3875c | BCG_3875c | 822 | 4273513 | 4274334 | BCGDan_3958 | BCGDan_3958 | 822 | 4194397 | 4195206 | BCGDan_sapMKO_3883 | BCGDan_sapMKO_3883 | 810 |
| 4279230 | 4280015 | Rv3814c | Rv3814c | 786 | 4251633 | 4252418 | BCG_3876c | BCG_3876c | 786 | 4274349 | 4275134 | BCGDan_3959 | BCGDan_3959 | 786 | 4195233 | 4196018 | BCGDan_sapMKO_3884 | BCGDan_sapMKO_3884 | 786 |
| 4280033 | 4280788 | Rv3815c | Rv3815c | 756 | 4252436 | 4253191 | BCG_3877c | BCG_3877c | 756 | 4275152 | 4275907 | BCGDan_3960 | BCGDan_3960 | 756 | 4196036 | 4196791 | BCGDan_sapMKO_3885 | BCGDan_sapMKO_3885 | 756 |
| 4280792 | 4281571 | Rv3816c | Rv3816c | 780 | 4253195 | 4253974 | BCG_3878c | BCG_3878c | 780 | 4275911 | 4276690 | BCGDan_3961 | BCGDan_3961 | 780 | 4196795 | 4197574 | BCGDan_sapMKO_3886 | BCGDan_sapMKO_3886 | 780 |
| 4281647 | 4282402 | Rv3817 | Rv3817 | 756 | 4254050 | 4254805 | BCG_3879 | BCG_3879 | 756 | 4276766 | 4277521 | BCGDan_3962 | BCGDan_3962 | 756 | 4197650 | 4198405 | BCGDan_sapMKO_3887 | BCGDan_sapMKO_3887 | 756 |
| 4282449 | 4283999 | Rv3818 | Rv3818 | 1551 | 4254852 | 4256402 | BCG_3880 | BCG_3880 | 1551 | 4277568 | 4279118 | BCGDan_3963 | BCGDan_3963 | 1551 | 4198452 | 4200002 | BCGDan_sapMKO_3888 | BCGDan_sapMKO_3888 | 1551 |
| 4283996 | 4284331 | Rv3819 | Rv3819 | 336 | 4256399 | 4256734 | BCG_3881 | BCG_3881 | 336 | 4279115 | 4279450 | BCGDan_3964 | BCGDan_3964 | 336 | 4199999 | 4200334 | BCGDan_sapMKO_3889 | BCGDan_sapMKO_3889 | 336 |
| 4284419 | 4285825 | papA2 | Rv3820c | 1407 | 4256822 | 4258228 | papA2 | BCG_3882c | 1407 | 4279538 | 4280944 | papA2 | BCGDan_3965 | 1407 | 4200422 | 4201648 | papA2 | BCGDan_sapMKO_3890 | 1227 |
| 4285973 | 4286686 | Rv3821 | Rv3821 | 714 | 4258376 | 4259089 | BCG_3883 | BCG_3883 | 714 | 4281092 | 4281805 | BCGDan_3966 | BCGDan_3966 | 714 | 4201892 | 4202689 | BCGDan_sapMKO_3891 | BCGDan_sapMKO_3891 | 798 |
| 4286721 | 4287935 | Rv3822 | Rv3822 | 1215 | 4259124 | 4259492 | BCG_3884 | BCG_3884 | 369 | 4281840 | 4282208 | BCGDan_3967 | BCGDan_3967 | 369 | 4202724 | 4203092 | BCGDan_sapMKO_3892 | BCGDan_sapMKO_3892 | 369 |
| NA | NA | NA | NA | NA | 4259489 | 4260337 | BCG_3885 | BCG_3885 | 849 | 4282205 | 4283053 | BCGDan_3968 | BCGDan_3968 | 849 | 4203089 | 4203937 | BCGDan_sapMKO_3893 | BCGDan_sapMKO_3893 | 849 |
| 4288260 | 4291529 | mmpL8 | Rv3823c | 3270 | 4260662 | 4263931 | mmpL8 | BCG_3886c | 3270 | 4283378 | 4286647 | mmpL8 | BCGDan_3969 | 3270 | 4204262 | 4207399 | mmpL8 | BCGDan_sapMKO_3894 | 3138 |
| 4291639 | 4293174 | papA1 | Rv3824c | 1536 | 4264041 | 4265576 | papA1 | BCG_3887c | 1536 | 4286757 | 4288292 | papA1 | BCGDan_3970 | 1536 | 4207641 | 4208954 | papA1 | BCGDan_sapMKO_3895 | 1314 |
| 4293225 | 4299605 | pks2 | Rv3825c | 6381 | 4265627 | 4272007 | pks2 | BCG_3888c | 6381 | 4288343 | 4294723 | pks2 | BCGDan_3971 | 6381 | 4209227 | 4215514 | pks2 | BCGDan_sapMKO_3896 | 6288 |
| 4299812 | 4301566 | fadD23 | Rv3826 | 1755 | 4272214 | 4273968 | fadD23 | BCG_3889 | 1755 | 4294930 | 4296684 | fadD23 | BCGDan_3972 | 1755 | 4215814 | 4217568 | fadD23 | BCGDan_sapMKO_3897 | 1755 |
| 4301563 | 4302789 | Rv3827c | Rv3827c | 1227 | 4273965 | 4275191 | BCG_3890c | BCG_3890c | 1227 | 4296681 | 4297907 | BCGDan_3973 | BCGDan_3973 | 1227 | 4217565 | 4218788 | BCGDan_sapMKO_3898 | BCGDan_sapMKO_3898 | 1224 |
| 4302786 | 4303397 | Rv3828c | Rv3828c | 612 | 4275188 | 4275799 | BCG_3891c | BCG_3891c | 612 | 4297904 | 4298515 | BCGDan_3974 | BCGDan_3974 | 612 | 4218788 | 4219369 | BCGDan_sapMKO_3899 | BCGDan_sapMKO_3899 | 582 |
| 4303398 | 4305008 | Rv3829c | Rv3829c | 1611 | 4275800 | 4277410 | BCG_3892c | BCG_3892c | 1611 | 4298516 | 4300126 | BCGDan_3975 | BCGDan_3975 | 1611 | 4219400 | 4221010 | BCGDan_sapMKO_3900 | BCGDan_sapMKO_3900 | 1611 |
| 4305056 | 4305685 | Rv3830c | Rv3830c | 630 | 4277458 | 4278087 | BCG_3893c | BCG_3893c | 630 | 4300174 | 4300803 | BCGDan_3976 | BCGDan_3976 | 630 | 4221058 | 4221687 | BCGDan_sapMKO_3901 | BCGDan_sapMKO_3901 | 630 |
| 4305757 | 4306239 | Rv3831 | Rv3831 | 483 | 4278159 | 4278641 | BCG_3894 | BCG_3894 | 483 | 4300875 | 4301357 | BCGDan_3977 | BCGDan_3977 | 483 | 4221759 | 4222241 | BCGDan_sapMKO_3902 | BCGDan_sapMKO_3902 | 483 |
| 4306236 | 4306811 | Rv3832c | Rv3832c | 576 | 4278638 | 4279213 | BCG_3895c | BCG_3895c | 576 | 4301354 | 4301929 | BCGDan_3978 | BCGDan_3978 | 576 | 4222238 | 4222756 | BCGDan_sapMKO_3903 | BCGDan_sapMKO_3903 | 519 |
| 4306867 | 4307658 | Rv3833 | Rv3833 | 792 | 4279269 | 4280060 | BCG_3896 | BCG_3896 | 792 | 4301985 | 4302776 | BCGDan_3979 | BCGDan_3979 | 792 | 4222869 | 4223660 | BCGDan_sapMKO_3904 | BCGDan_sapMKO_3904 | 792 |
| 4307655 | 4308914 | serS | Rv3834c | 1260 | 4280057 | 4281316 | serS | BCG_3897c | 1260 | 4302773 | 4304032 | serS | BCGDan_3980 | 1260 | 4223657 | 4224916 | serS | BCGDan_sapMKO_3905 | 1260 |
| 4309047 | 4310396 | Rv3835 | Rv3835 | 1350 | 4281449 | 4282807 | BCG_3898 | BCG_3898 | 1359 | 4304165 | 4305514 | BCGDan_3981 | BCGDan_3981 | 1350 | 4225049 | 4226398 | BCGDan_sapMKO_3906 | BCGDan_sapMKO_3906 | 1350 |
| 4310401 | 4310814 | Rv3836 | Rv3836 | 414 | 4282804 | 4283154 | BCG_3899 | BCG_3899 | 351 | 4305519 | 4305869 | BCGDan_3982 | BCGDan_3982 | 351 | 4226415 | 4226753 | BCGDan_sapMKO_3908 | BCGDan_sapMKO_3908 | 339 |
| 4311009 | 4311707 | Rv3837c | Rv3837c | 699 | 4283413 | 4284111 | BCG_3900c | BCG_3900c | 699 | 4306128 | 4306826 | BCGDan_3983 | BCGDan_3983 | 699 | 4227012 | 4227710 | BCGDan_sapMKO_3909 | BCGDan_sapMKO_3909 | 699 |
| 4311704 | 4312669 | pheA | Rv3838c | 966 | 4284108 | 4285073 | pheA | BCG_3901c | 966 | 4306823 | 4307788 | pheA | BCGDan_3984 | 966 | 4227707 | 4228678 | pheA | BCGDan_sapMKO_3910 | 972 |
| 4312765 | 4313541 | Rv3839 | Rv3839 | 777 | 4285169 | 4285945 | BCG_3902 | BCG_3902 | 777 | 4307884 | 4308660 | BCGDan_3985 | BCGDan_3985 | 777 | 4228768 | 4229544 | BCGDan_sapMKO_3911 | BCGDan_sapMKO_3911 | 777 |
| 4313567 | 4313980 | Rv3840 | Rv3840 | 414 | 4285971 | 4286384 | BCG_3903 | BCG_3903 | 414 | 4308686 | 4309099 | BCGDan_3986 | BCGDan_3986 | 414 | 4229621 | 4229983 | BCGDan_sapMKO_3912 | BCGDan_sapMKO_3912 | 363 |
| 4314178 | 4314723 | bfrB | Rv3841 | 546 | 4286582 | 4287127 | bfrB | BCG_3904 | 546 | 4309297 | 4309842 | bfrB | BCGDan_3987 | 546 | 4230223 | 4230726 | bfrB | BCGDan_sapMKO_3913 | 504 |
| 4314738 | 4315562 | glpQ1 | Rv3842c | 825 | 4287142 | 4287966 | glpQ1 | BCG_3905c | 825 | 4309857 | 4310681 | glpQ1 | BCGDan_3988 | 825 | 4230741 | 4231571 | glpQ1 | BCGDan_sapMKO_3914 | 831 |
| 4315568 | 4316596 | Rv3843c | Rv3843c | 1029 | 4287972 | 4289000 | BCG_3906c | BCG_3906c | 1029 | 4310687 | 4311715 | BCGDan_3989 | BCGDan_3989 | 1029 | 4231571 | 4232599 | BCGDan_sapMKO_3915 | BCGDan_sapMKO_3915 | 1029 |
| 4318775 | 4319266 | Rv3844 | Rv3844 | 492 | 4291178 | 4291669 | BCG_3907 | BCG_3907 | 492 | 4313893 | 4314384 | BCGDan_3990 | BCGDan_3990 | 492 | 4234777 | 4235268 | BCGDan_sapMKO_3916 | BCGDan_sapMKO_3916 | 492 |
| 4319281 | 4319640 | Rv3845 | Rv3845 | 360 | 4291684 | 4292043 | BCG_3908 | BCG_3908 | 360 | 4314399 | 4314758 | BCGDan_3991 | BCGDan_3991 | 360 | 4235283 | 4235642 | BCGDan_sapMKO_3917 | BCGDan_sapMKO_3917 | 360 |
| NA | NA | NA | NA | NA | NA | NA | NA | NA | NA | 4314943 | 4315158 | BCGDan_3992 | BCGDan_3992 | 216 | 4235827 | 4236042 | BCGDan_sapMKO_3918 | BCGDan_sapMKO_3918 | 216 |
| NA | NA | NA | NA | NA | NA | NA | NA | NA | NA | 4315283 | 4315375 | BCGDan_3993 | BCGDan_3993 | 93 | 4236167 | 4236259 | BCGDan_sapMKO_3919 | BCGDan_sapMKO_3919 | 93 |
| NA | NA | NA | NA | NA | NA | NA | NA | NA | NA | 4315481 | 4315789 | BCGDan_3994 | BCGDan_3994 | 309 | 4236365 | 4236673 | BCGDan_sapMKO_3920 | BCGDan_sapMKO_3920 | 309 |
| 4320704 | 4321327 | sodA | Rv3846 | 624 | 4293107 | 4293730 | sodA | BCG_3909 | 624 | 4315822 | 4316445 | sodA | BCGDan_3995 | 624 | 4236706 | 4237329 | sodA | BCGDan_sapMKO_3921 | 624 |
| 4321538 | 4322071 | Rv3847 | Rv3847 | 534 | 4293941 | 4294474 | BCG_3910 | BCG_3910 | 534 | 4316656 | 4317189 | BCGDan_3996 | BCGDan_3996 | 534 | 4237540 | 4238073 | BCGDan_sapMKO_3922 | BCGDan_sapMKO_3922 | 534 |
| 4322326 | 4323234 | Rv3848 | Rv3848 | 909 | 4294729 | 4295637 | BCG_3911 | BCG_3911 | 909 | 4317444 | 4318352 | BCGDan_3997 | BCGDan_3997 | 909 | 4238328 | 4239236 | BCGDan_sapMKO_3923 | BCGDan_sapMKO_3923 | 909 |
| 4323499 | 4323897 | espR | Rv3849 | 399 | 4295902 | 4296300 | BCG_3912 | BCG_3912 | 399 | 4318617 | 4319015 | BCGDan_3998 | BCGDan_3998 | 399 | 4239486 | 4239899 | BCGDan_sapMKO_3924 | BCGDan_sapMKO_3924 | 414 |
| 4324015 | 4324671 | Rv3850 | Rv3850 | 657 | 4296418 | 4297074 | BCG_3913 | BCG_3913 | 657 | 4319133 | 4319789 | BCGDan_3999 | BCGDan_3999 | 657 | 4240017 | 4240673 | BCGDan_sapMKO_3925 | BCGDan_sapMKO_3925 | 657 |
| 4324683 | 4324967 | Rv3851 | Rv3851 | 285 | 4297086 | 4297370 | BCG_3914 | BCG_3914 | 285 | 4319801 | 4320085 | BCGDan_4000 | BCGDan_4000 | 285 | 4240685 | 4240969 | BCGDan_sapMKO_3926 | BCGDan_sapMKO_3926 | 285 |
| 4325074 | 4325478 | hns | Rv3852 | 405 | 4297477 | 4297881 | hns | BCG_3915 | 405 | 4320192 | 4320596 | hns | BCGDan_4001 | 405 | 4241076 | 4241480 | hns | BCGDan_sapMKO_3927 | 405 |

| | | | | | | | | | | | | | | | | | | | |
|---|---|---|---|---|---|---|---|---|---|---|---|---|---|---|---|---|---|---|---|
| 4325495 | 4325968 | rraA | Rv3853 | 474 | 4297898 | 4298371 | menG | BCG_3916 | 474 | 4320613 | 4321086 | menG | BCGDan_4002 | 474 | 4241497 | 4241970 | menG | BCGDan_sapMKO_3928 | 474 |
| 4326004 | 4327473 | ethA | Rv3854c | 1470 | 4298407 | 4299876 | ethA | BCG_3917c | 1470 | 4321122 | 4322591 | ethA | BCGDan_4003 | 1470 | 4242006 | 4243475 | ethA | BCGDan_sapMKO_3929 | 1470 |
| 4327549 | 4328199 | ethR | Rv3855 | 651 | 4299952 | 4300602 | ethR | BCG_3918 | 651 | 4322667 | 4323317 | ethR | BCGDan_4004 | 651 | 4243551 | 4244201 | ethR | BCGDan_sapMKO_3930 | 651 |
| 4328401 | 4329408 | Rv3856c | Rv3856c | 1008 | 4300804 | 4301811 | BCG_3919c | BCG_3919c | 1008 | 4323519 | 4324526 | BCGDan_4005 | BCGDan_4005 | 1008 | 4244403 | 4245410 | BCGDan_sapMKO_3931 | BCGDan_sapMKO_3931 | 1008 |
| 4329417 | 4329614 | Rv3857c | Rv3857c | 198 | 4301820 | 4302017 | BCG_3920c | BCG_3920c | 198 | 4324535 | 4324732 | BCGDan_4006 | BCGDan_4006 | 198 | 4245419 | 4245616 | BCGDan_sapMKO_3932 | BCGDan_sapMKO_3932 | 198 |
| 4330039 | 4331505 | gltD | Rv3858c | 1467 | 4302442 | 4303908 | gltD | BCG_3921c | 1467 | 4325157 | 4326623 | gltD | BCGDan_4007 | 1467 | 4246041 | 4247507 | gltD | BCGDan_sapMKO_3933 | 1467 |
| 4331498 | 4336081 | gltB | Rv3859c | 4584 | 4303901 | 4308484 | gltB | BCG_3922c | 4584 | 4326616 | 4331199 | gltB | BCGDan_4008 | 4584 | 4247500 | 4252083 | gltB | BCGDan_sapMKO_3934 | 4584 |
| 4336777 | 4337949 | Rv3860 | Rv3860 | 1173 | 4309180 | 4310241 | BCG_3923 | BCG_3923 | 1062 | 4331895 | 4332956 | BCGDan_4009 | BCGDan_4009 | 1062 | 4252779 | 4253840 | BCGDan_sapMKO_3935 | BCGDan_sapMKO_3935 | 1062 |
| 4337946 | 4338272 | Rv3861 | Rv3861 | 327 | 4310348 | 4310674 | BCG_3924 | BCG_3924 | 327 | 4333063 | 4333389 | BCGDan_4010 | BCGDan_4010 | 327 | 4253947 | 4254273 | BCGDan_sapMKO_3936 | BCGDan_sapMKO_3936 | 327 |
| 4338171 | 4338521 | whiB6 | Rv3862c | 351 | 4310573 | 4310923 | whiB6 | BCG_3925c | 351 | 4333288 | 4333638 | whiB6 | BCGDan_4011 | 351 | 4254172 | 4254522 | whiB6 | BCGDan_sapMKO_3937 | 351 |
| 4338849 | 4340027 | Rv3863 | Rv3863 | 1179 | 4311250 | 4312428 | BCG_3926 | BCG_3926 | 1179 | 4333965 | 4335143 | BCGDan_4012 | BCGDan_4012 | 1179 | 4254915 | 4256027 | BCGDan_sapMKO_3938 | BCGDan_sapMKO_3938 | 1113 |
| 4340270 | 4341478 | espE | Rv3864 | 1209 | 4312671 | 4313927 | BCG_3927 | BCG_3927 | 1257 | 4335386 | 4336642 | BCGDan_4013 | BCGDan_4013 | 1257 | 4256330 | 4257526 | BCGDan_sapMKO_3939 | BCGDan_sapMKO_3939 | 1197 |
| 4341566 | 4341877 | espF | Rv3865 | 312 | 4314015 | 4314326 | BCG_3928 | BCG_3928 | 312 | 4336730 | 4337041 | BCGDan_4014 | BCGDan_4014 | 312 | 4257614 | 4257925 | BCGDan_sapMKO_3940 | BCGDan_sapMKO_3940 | 312 |
| 4341880 | 4342731 | espG1 | Rv3866 | 852 | 4314329 | 4315180 | BCG_3929 | BCG_3929 | 852 | 4337044 | 4337895 | BCGDan_4015 | BCGDan_4015 | 852 | 4257928 | 4258779 | BCGDan_sapMKO_3941 | BCGDan_sapMKO_3941 | 852 |
| 4342770 | 4343321 | espH | Rv3867 | 552 | 4315219 | 4315770 | BCG_3930 | BCG_3930 | 552 | 4337934 | 4338485 | BCGDan_4016 | BCGDan_4016 | 552 | 4258818 | 4259369 | BCGDan_sapMKO_3942 | BCGDan_sapMKO_3942 | 552 |
| 4343314 | 4345035 | eccA1 | Rv3868 | 1722 | 4315763 | 4317484 | BCG_3931 | BCG_3931 | 1722 | 4338478 | 4340199 | BCGDan_4017 | BCGDan_4017 | 1722 | 4259362 | 4261083 | BCGDan_sapMKO_3943 | BCGDan_sapMKO_3943 | 1722 |
| 4345039 | 4346481 | eccB1 | Rv3869 | 1443 | 4317488 | 4318930 | BCG_3932 | BCG_3932 | 1443 | 4340203 | 4341645 | BCGDan_4018 | BCGDan_4018 | 1443 | 4261087 | 4262529 | BCGDan_sapMKO_3944 | BCGDan_sapMKO_3944 | 1443 |
| 4346481 | 4348724 | eccCa1 | Rv3870 | 2244 | 4318930 | 4321173 | BCG_3933 | BCG_3933 | 2244 | 4341645 | 4343888 | BCGDan_4019 | BCGDan_4019 | 2244 | 4262529 | 4264772 | BCGDan_sapMKO_3945 | BCGDan_sapMKO_3945 | 2244 |
| 4348827 | 4350602 | eccCb1 | Rv3871 | 1776 | 4321276 | 4322748 | BCG_3934 | BCG_3934 | 1473 | 4343991 | 4345463 | BCGDan_4020 | BCGDan_4020 | 1473 | 4264875 | 4266347 | BCGDan_sapMKO_3946 | BCGDan_sapMKO_3946 | 1473 |
| 4360199 | 4360546 | espL | Rv3880c | 348 | 4323191 | 4323538 | BCG_3935c | BCG_3935c | 348 | 4345906 | 4346253 | BCGDan_4021 | BCGDan_4021 | 348 | 4266790 | 4267131 | BCGDan_sapMKO_3947 | BCGDan_sapMKO_3947 | 342 |
| 4360543 | 4361925 | espB | Rv3881c | 1383 | 4323535 | 4324917 | BCG_3936c | BCG_3936c | 1383 | 4346250 | 4347632 | BCGDan_4022 | BCGDan_4022 | 1383 | 4267134 | 4268516 | BCGDan_sapMKO_3948 | BCGDan_sapMKO_3948 | 1383 |
| 4362032 | 4363420 | eccE1 | Rv3882c | 1389 | 4325024 | 4326412 | BCG_3937c | BCG_3937c | 1389 | 4347739 | 4349127 | BCGDan_4023 | BCGDan_4023 | 1389 | 4268623 | 4270011 | BCGDan_sapMKO_3949 | BCGDan_sapMKO_3949 | 1389 |
| 4363417 | 4364757 | mycP1 | Rv3883c | 1341 | 4326409 | 4327749 | BCG_3938c | BCG_3938c | 1341 | 4349124 | 4350464 | BCGDan_4024 | BCGDan_4024 | 1341 | 4270008 | 4271378 | BCGDan_sapMKO_3950 | BCGDan_sapMKO_3950 | 1371 |
| 4364979 | 4366838 | eccA2 | Rv3884c | 1860 | 4327971 | 4329830 | BCG_3939c | BCG_3939c | 1860 | 4350686 | 4352545 | BCGDan_4025 | BCGDan_4025 | 1860 | 4271570 | 4273420 | BCGDan_sapMKO_3951 | BCGDan_sapMKO_3951 | 1851 |
| 4366908 | 4368521 | eccE2 | Rv3885c | 1614 | 4329900 | 4331513 | BCG_3940c | BCG_3940c | 1614 | 4352615 | 4354228 | BCGDan_4026 | BCGDan_4026 | 1614 | 4273499 | 4275112 | BCGDan_sapMKO_3952 | BCGDan_sapMKO_3952 | 1614 |
| 4368518 | 4370170 | mycP2 | Rv3886c | 1653 | 4331510 | 4333162 | BCG_3941c | BCG_3941c | 1653 | 4354225 | 4355877 | BCGDan_4027 | BCGDan_4027 | 1653 | 4275109 | 4276545 | BCGDan_sapMKO_3953 | BCGDan_sapMKO_3953 | 1437 |
| 4370155 | 4371684 | eccD2 | Rv3887c | 1530 | 4333147 | 4334676 | BCG_3942c | BCG_3942c | 1530 | 4355862 | 4357391 | BCGDan_4028 | BCGDan_4028 | 1530 | 4276746 | 4278155 | BCGDan_sapMKO_3954 | BCGDan_sapMKO_3954 | 1410 |
| 4371681 | 4372706 | Rv3888c | Rv3888c | 1026 | 4334673 | 4335335 | BCG_3943c | BCG_3943c | 663 | 4357388 | 4358050 | BCGDan_4029 | BCGDan_4029 | 663 | 4278272 | 4279015 | BCGDan_sapMKO_3955 | BCGDan_sapMKO_3955 | 744 |
| NA | NA | NA | NA | NA | 4335332 | 4335697 | BCG_3944c | BCG_3944c | 366 | 4358047 | 4358412 | BCGDan_4030 | BCGDan_4030 | 366 | 4278931 | 4279296 | BCGDan_sapMKO_3956 | BCGDan_sapMKO_3956 | 366 |
| 4372800 | 4373630 | espG2 | Rv3889c | 831 | 4335791 | 4336621 | BCG_3945c | BCG_3945c | 831 | 4358506 | 4359336 | BCGDan_4031 | BCGDan_4031 | 831 | 4279390 | 4280220 | BCGDan_sapMKO_3957 | BCGDan_sapMKO_3957 | 831 |
| 4373726 | 4374013 | esxC | Rv3890c | 288 | 4336629 | 4337003 | esxC | BCG_3946c | 375 | 4359344 | 4359718 | esxC | BCGDan_4032 | 375 | 4280228 | 4280602 | esxC | BCGDan_sapMKO_3958 | 375 |
| 4374049 | 4374372 | esxD | Rv3891c | 324 | 4337039 | 4337362 | esxD | BCG_3947c | 324 | 4359754 | 4360077 | esxD | BCGDan_4033 | 324 | 4280638 | 4280961 | esxD | BCGDan_sapMKO_3959 | 324 |
| 4374484 | 4375683 | PPE69 | Rv3892c | 1200 | 4337473 | 4338672 | PPE69 | BCG_3948c | 1200 | 4360188 | 4361387 | PPE69 | BCGDan_4034 | 1200 | 4281072 | 4282271 | PPE69 | BCGDan_sapMKO_3960 | 1200 |
| 4375762 | 4375995 | PE36 | Rv3893c | 234 | 4338751 | 4338984 | PE36 | BCG_3949c | 234 | 4361466 | 4361699 | PE36 | BCGDan_4035 | 234 | 4282350 | 4282556 | PE36 | BCGDan_sapMKO_3961 | 207 |
| 4376262 | 4380452 | eccC2 | Rv3894c | 4191 | 4339251 | 4340936 | BCG_3950c | BCG_3950c | 1686 | 4361966 | 4363651 | BCGDan_4036 | BCGDan_4036 | 1686 | 4282850 | 4284733 | BCGDan_sapMKO_3962 | BCGDan_sapMKO_3962 | 1884 |
| NA | NA | NA | NA | NA | 4340941 | 4343442 | BCG_3951c | BCG_3951c | 2502 | 4363656 | 4366157 | BCGDan_4037 | BCGDan_4037 | 2502 | 4284540 | 4287041 | BCGDan_sapMKO_3963 | BCGDan_sapMKO_3963 | 2502 |
| 4380453 | 4381940 | eccB2 | Rv3895c | 1488 | 4343443 | 4344930 | BCG_3952c | BCG_3952c | 1488 | 4366158 | 4367645 | BCGDan_4038 | BCGDan_4038 | 1488 | 4287042 | 4288529 | BCGDan_sapMKO_3964 | BCGDan_sapMKO_3964 | 1488 |
| 4381943 | 4382851 | Rv3896c | Rv3896c | 909 | 4344974 | 4345840 | BCG_3953c | BCG_3953c | 867 | 4367689 | 4368555 | BCGDan_4039 | BCGDan_4039 | 867 | 4288573 | 4289439 | BCGDan_sapMKO_3965 | BCGDan_sapMKO_3965 | 867 |
| 4383008 | 4383640 | Rv3897c | Rv3897c | 633 | 4345990 | 4346979 | BCG_3954c | BCG_3954c | 990 | 4368705 | 4369694 | BCGDan_4040 | BCGDan_4040 | 990 | 4289589 | 4290584 | BCGDan_sapMKO_3966 | BCGDan_sapMKO_3966 | 996 |
| 4384147 | 4385379 | Rv3899c | Rv3899c | 1233 | 4347141 | 4348022 | BCG_3955c | BCG_3955c | 882 | 4369856 | 4370737 | BCGDan_4041 | BCGDan_4041 | 882 | 4290740 | 4291621 | BCGDan_sapMKO_3967 | BCGDan_sapMKO_3967 | 882 |
| NA | NA | NA | NA | NA | 4347629 | 4348372 | BCG_3956c | BCG_3956c | 744 | 4370344 | 4371087 | BCGDan_4042 | BCGDan_4042 | 744 | 4291231 | 4291971 | BCGDan_sapMKO_3968 | BCGDan_sapMKO_3968 | 741 |
| 4385373 | 4386308 | Rv3900c | Rv3900c | 936 | 4348366 | 4349301 | BCG_3957c | BCG_3957c | 936 | 4371081 | 4372016 | BCGDan_4043 | BCGDan_4043 | 936 | 4291965 | 4292861 | BCGDan_sapMKO_3969 | BCGDan_sapMKO_3969 | 897 |
| 4386365 | 4386814 | Rv3901c | Rv3901c | 450 | 4349410 | 4349805 | BCG_3958c | BCG_3958c | 396 | 4372125 | 4372520 | BCGDan_4044 | BCGDan_4044 | 396 | 4293009 | 4293404 | BCGDan_sapMKO_3970 | BCGDan_sapMKO_3970 | 396 |
| 4387365 | 4387895 | Rv3902c | Rv3902c | 531 | 4350356 | 4350886 | BCG_3959c | BCG_3959c | 531 | 4373071 | 4373601 | BCGDan_4045 | BCGDan_4045 | 531 | 4293955 | 4294485 | BCGDan_sapMKO_3971 | BCGDan_sapMKO_3971 | 531 |
| 4387892 | 4390432 | Rv3903c | Rv3903c | 2541 | 4350883 | 4353423 | BCG_3960c | BCG_3960c | 2541 | 4373598 | 4376138 | BCGDan_4046 | BCGDan_4046 | 2541 | 4294482 | 4297022 | BCGDan_sapMKO_3972 | BCGDan_sapMKO_3972 | 2541 |
| 4390437 | 4390709 | esxE | Rv3904c | 273 | 4353428 | 4353700 | esxE | BCG_3961c | 273 | 4376143 | 4376415 | esxE | BCGDan_4047 | 273 | 4297027 | 4297317 | esxE | BCGDan_sapMKO_3973 | 291 |
| 4390720 | 4391031 | esxF | Rv3905c | 312 | 4353849 | 4354022 | esxF | BCG_3962c | 174 | 4376426 | 4376536 | BCGDan_4048 | BCGDan_4048 | 111 | 4297310 | 4297420 | BCGDan_sapMKO_3974 | BCGDan_sapMKO_3974 | 111 |
| NA | NA | NA | NA | NA | NA | NA | NA | NA | NA | 4376564 | 4376737 | esxF | BCGDan_4049 | 174 | 4297448 | 4297621 | esxF | BCGDan_sapMKO_3975 | 174 |
| 4391097 | 4391606 | Rv3906c | Rv3906c | 510 | 4354088 | 4354597 | BCG_3963c | BCG_3963c | 510 | 4376803 | 4377312 | BCGDan_4050 | BCGDan_4050 | 510 | 4297687 | 4298196 | BCGDan_sapMKO_3976 | BCGDan_sapMKO_3976 | 510 |
| 4391631 | 4393073 | pcnA | Rv3907c | 1443 | 4354622 | 4356064 | pcnA | BCG_3964c | 1443 | 4377337 | 4378779 | pcnA | BCGDan_4051 | 1443 | 4298221 | 4299663 | pcnA | BCGDan_sapMKO_3977 | 1443 |
| 4393449 | 4394195 | mutT4 | Rv3908 | 747 | 4356440 | 4357186 | BCG_3965 | BCG_3965 | 747 | 4379155 | 4379901 | BCGDan_4052 | BCGDan_4052 | 747 | 4300039 | 4300785 | BCGDan_sapMKO_3978 | BCGDan_sapMKO_3978 | 747 |
| 4394192 | 4396600 | Rv3909 | Rv3909 | 2409 | 4357183 | 4359591 | BCG_3966 | BCG_3966 | 2409 | 4379898 | 4382306 | BCGDan_4053 | BCGDan_4053 | 2409 | 4300782 | 4303190 | BCGDan_sapMKO_3979 | BCGDan_sapMKO_3979 | 2409 |
| 4396597 | 4400151 | Rv3910 | Rv3910 | 3555 | 4359588 | 4363142 | BCG_3967 | BCG_3967 | 3555 | 4382303 | 4385857 | BCGDan_4054 | BCGDan_4054 | 3555 | 4303274 | 4306741 | BCGDan_sapMKO_3980 | BCGDan_sapMKO_3980 | 3468 |
| 4400186 | 4400854 | sigM | Rv3911 | 669 | 4363177 | 4363767 | sigMa_1 | BCG_3968 | 591 | 4385892 | 4386482 | sigMa | BCGDan_4055 | 591 | 4306806 | 4307366 | sigMa | BCGDan_sapMKO_3981 | 561 |
| NA | NA | NA | NA | NA | 4363656 | 4363844 | sigMb_1 | BCG_3969 | 189 | 4386371 | 4386559 | sigMb | BCGDan_4056 | 189 | 4307255 | 4307440 | sigMb | BCGDan_sapMKO_3982 | 186 |
| 4400870 | 4401634 | Rv3912 | Rv3912 | 765 | 4363860 | 4364624 | BCG_3970 | BCG_3970 | 765 | 4386575 | 4387339 | BCGDan_4057 | BCGDan_4057 | 765 | 4307459 | 4308223 | BCGDan_sapMKO_3983 | BCGDan_sapMKO_3983 | 765 |
| 4401728 | 4402735 | trxB2 | Rv3913 | 1008 | 4364718 | 4365725 | trxB2_1 | BCG_3971 | 1008 | 4387433 | 4388440 | trxB2 | BCGDan_4058 | 1008 | 4308317 | 4309324 | trxB2 | BCGDan_sapMKO_3984 | 1008 |
| 4402732 | 4403082 | trxC | Rv3914 | 351 | 4365722 | 4366072 | trxC_1 | BCG_3972 | 351 | 4388437 | 4388787 | trxC | BCGDan_4059 | 351 | 4309321 | 4309671 | trxC | BCGDan_sapMKO_3985 | 351 |
| 4403192 | 4404412 | Rv3915 | Rv3915 | 1221 | 4366182 | 4367402 | BCG_3973 | BCG_3973 | 1221 | 4388897 | 4390117 | BCGDan_4060 | BCGDan_4060 | 1221 | 4309781 | 4311001 | BCGDan_sapMKO_3986 | BCGDan_sapMKO_3986 | 1221 |
| 4404433 | 4405167 | Rv3916c | Rv3916c | 735 | 4367423 | 4368157 | BCG_3974c | BCG_3974c | 735 | 4390138 | 4390872 | BCGDan_4061 | BCGDan_4061 | 735 | 4311022 | 4311756 | BCGDan_sapMKO_3987 | BCGDan_sapMKO_3987 | 735 |
| 4405457 | 4406491 | parB | Rv3917c | 1035 | 4368447 | 4369481 | parB_1 | BCG_3975c | 1035 | 4391162 | 4392196 | parB_1 | BCGDan_4062 | 1035 | 4312046 | 4313080 | parB_1 | BCGDan_sapMKO_3988 | 1035 |
| 4406488 | 4407531 | parA | Rv3918c | 1044 | 4369478 | 4370521 | parA_1 | BCG_3976c | 1044 | 4392193 | 4393236 | parA_1 | BCGDan_4063 | 1044 | 4313077 | 4313985 | parA_1 | BCGDan_sapMKO_3989 | 909 |
| 4407528 | 4408202 | gid | Rv3919c | 675 | 4370518 | 4371192 | gid_1 | BCG_3977c | 675 | 4393233 | 4393907 | gid_1 | BCGDan_4064 | 675 | 4314117 | 4314791 | gid_1 | BCGDan_sapMKO_3990 | 675 |
| 4408334 | 4408897 | Rv3920c | Rv3920c | 564 | 4371324 | 4371887 | BCG_3978c | BCG_3978c | 564 | 4394039 | 4394602 | BCGDan_4065 | BCGDan_4065 | 564 | 4314923 | 4315486 | BCGDan_sapMKO_3991 | BCGDan_sapMKO_3991 | 564 |
| 4408969 | 4410069 | Rv3921c | Rv3921c | 1101 | 4371959 | 4373059 | BCG_3979c | BCG_3979c | 1101 | 4394674 | 4395774 | BCGDan_4066 | BCGDan_4066 | 1101 | 4315558 | 4316658 | BCGDan_sapMKO_3992 | BCGDan_sapMKO_3992 | 1101 |
| 4410053 | 4410415 | Rv3922c | Rv3922c | 363 | 4373043 | 4373405 | BCG_3980c | BCG_3980c | 363 | 4395758 | 4396120 | BCGDan_4067 | BCGDan_4067 | 363 | 4316642 | 4317004 | BCGDan_sapMKO_3993 | BCGDan_sapMKO_3993 | 363 |
| 4410412 | 4410789 | rnpA | Rv3923c | 378 | 4373402 | 4373779 | rnpA_1 | BCG_3981c | 378 | 4396117 | 4396494 | rnpA_1 | BCGDan_4068 | 378 | 4317001 | 4317267 | rnpA_1 | BCGDan_sapMKO_3994 | 267 |
| 4410786 | 4410929 | rpmH | Rv3924c | 144 | 4373776 | 4373919 | rpmH_1 | BCG_3982c | 144 | 4396491 | 4396634 | rpmH_1 | BCGDan_4069 | 144 | 4317375 | 4317518 | rpmH_1 | BCGDan_sapMKO_3995 | 144 |
| NA | NA | NA | NA | NA | 29668 | 31191 | dnaA_2 | BCG_0031 | 1524 | 4397238 | 4398761 | dnaA_2 | BCGDan_4070 | 1524 | 4318155 | 4319645 | dnaA_2 | BCGDan_sapMKO_3996 | 1491 |
| NA | NA | NA | NA | NA | 31719 | 32927 | dnaN_2 | BCG_0032 | 1209 | 4399289 | 4400497 | dnaN_2 | BCGDan_4071 | 1209 | 4320248 | 4321381 | dnaN_2 | BCGDan_sapMKO_3997 | 1134 |
| NA | NA | NA | NA | NA | 32947 | 34104 | recF_2 | BCG_0033 | 1158 | 4400517 | 4401674 | recF_2 | BCGDan_4072 | 1158 | 4321572 | 4322558 | recF_2 | BCGDan_sapMKO_3998 | 987 |
| NA | NA | NA | NA | NA | 34101 | 34664 | BCG_0034 | BCG_0034 | 564 | 4401671 | 4402234 | BCGDan_4073 | BCGDan_4073 | 564 | 4322555 | 4323118 | BCGDan_sapMKO_3999 | BCGDan_sapMKO_3999 | 564 |
| NA | NA | NA | NA | NA | 34790 | 36934 | gyrB_2 | BCG_0035 | 2145 | 4402360 | 4403364 | gyrB | BCGDan_4074 | 1005 | 4323244 | 4324248 | gyrB_2 | BCGDan_sapMKO_4000 | 1005 |
| NA | NA | NA | NA | NA | 21327 | 22547 | BCG_0021 | BCG_0021 | 1221 | 4403474 | 4404694 | BCGDan_4075 | BCGDan_4075 | 1221 | 4324358 | 4325578 | BCGDan_sapMKO_4001 | BCGDan_sapMKO_4001 | 1221 |
| NA | NA | NA | NA | NA | 22568 | 23302 | BCG_0022c | BCG_0022c | 735 | 4404715 | 4405449 | BCGDan_4076 | BCGDan_4076 | 735 | 4325599 | 4326333 | BCGDan_sapMKO_4002 | BCGDan_sapMKO_4002 | 735 |
| NA | NA | NA | NA | NA | 23592 | 24626 | parB_2 | BCG_0023c | 1035 | 4405739 | 4406773 | parB_2 | BCGDan_4077 | 1035 | 4326623 | 4327657 | parB_2 | BCGDan_sapMKO_4003 | 1035 |
| NA | NA | NA | NA | NA | 24623 | 25666 | parA_2 | BCG_0024c | 1044 | 4406770 | 4407813 | parA_2 | BCGDan_4078 | 1044 | 4327654 | 4328562 | parA_2 | BCGDan_sapMKO_4004 | 909 |
| NA | NA | NA | NA | NA | 25663 | 26337 | gid_2 | BCG_0025c | 675 | 4407810 | 4408484 | gid_2 | BCGDan_4079 | 675 | 4328694 | 4329368 | gid_2 | BCGDan_sapMKO_4005 | 675 |
| NA | NA | NA | NA | NA | 26469 | 27032 | BCG_0026c | BCG_0026c | 564 | 4408616 | 4409179 | BCGDan_4080 | BCGDan_4080 | 564 | 4329500 | 4330063 | BCGDan_sapMKO_4006 | BCGDan_sapMKO_4006 | 564 |
| NA | NA | NA | NA | NA | 27104 | 28204 | BCG_0027c | BCG_0027c | 1101 | 4409251 | 4410351 | BCGDan_4081 | BCGDan_4081 | 1101 | 4330135 | 4331235 | BCGDan_sapMKO_4007 | BCGDan_sapMKO_4007 | 1101 |
| NA | NA | NA | NA | NA | 28188 | 28550 | BCG_0028c | BCG_0028c | 363 | 4410335 | 4410697 | BCGDan_4082 | BCGDan_4082 | 363 | 4331219 | 4331581 | BCGDan_sapMKO_4008 | BCGDan_sapMKO_4008 | 363 |
| NA | NA | NA | NA | NA | 28547 | 28924 | rnpA_2 | BCG_0029c | 378 | 4410694 | 4411071 | rnpA_2 | BCGDan_4083 | 378 | 4331578 | 4331955 | rnpA_2 | BCGDan_sapMKO_4009 | 378 |
| NA | NA | NA | NA | NA | 28921 | 29064 | rpmH_2 | BCG_0030c | 144 | 4411068 | 4411211 | rpmH_2 | BCGDan_4084 | 144 | 4331952 | 4332095 | rpmH_2 | BCGDan_sapMKO_4010 | 144 |
| NA | NA | NA | NA | NA | 16731 | 18287 | BCG_0015 | BCG_0015 | 1557 | NA | NA | NA | NA | NA | NA | NA | NA | NA | NA |
| NA | NA | NA | NA | NA | 18322 | 18912 | sigMa_2 | BCG_0016 | 591 | NA | NA | NA | NA | NA | NA | NA | NA | NA | NA |
| NA | NA | NA | NA | NA | 18801 | 18989 | sigMb_2 | BCG_0017 | 189 | NA | NA | NA | NA | NA | NA | NA | NA | NA | NA |

| | | | | | | | | | | | | | | | | | | | |
|---|---|---|---|---|---|---|---|---|---|---|---|---|---|---|---|---|---|---|---|
| NA | NA | NA | NA | NA | 19005 | 19769 | BCG_0018 | BCG_0018 | 765 | NA | NA | NA | NA | NA | NA | NA | NA | NA | NA |
| NA | NA | NA | NA | NA | 19863 | 20870 | trxB2_2 | BCG_0019 | 1008 | NA | NA | NA | NA | NA | NA | NA | NA | NA | NA |
| NA | NA | NA | NA | NA | 20867 | 21217 | trxC_2 | BCG_0020 | 351 | NA | NA | NA | NA | NA | NA | NA | NA | NA | NA |
| NA | NA | NA | NA | NA | 36969 | 39485 | gyrA_2 | BCG_0036 | 2517 | NA | NA | NA | NA | NA | NA | NA | NA | NA | NA |
| NA | NA | NA | NA | NA | 39581 | 40495 | BCG_0037 | BCG_0037 | 915 | NA | NA | NA | NA | NA | NA | NA | NA | NA | NA |
| NA | NA | NA | NA | NA | 41541 | 41978 | BCG_0038c | BCG_0038c | 438 | NA | NA | NA | NA | NA | NA | NA | NA | NA | NA |
| NA | NA | NA | NA | NA | 42135 | 42683 | ppiA_2 | BCG_0039 | 549 | NA | NA | NA | NA | NA | NA | NA | NA | NA | NA |
| NA | NA | NA | NA | NA | 42889 | 43224 | BCG_0040c | BCG_0040c | 336 | NA | NA | NA | NA | NA | NA | NA | NA | NA | NA |
| NA | NA | NA | NA | NA | 43380 | 43661 | BCG_0041c | BCG_0041c | 282 | NA | NA | NA | NA | NA | NA | NA | NA | NA | NA |
| NA | NA | NA | NA | NA | 43755 | 44543 | BCG_0042 | BCG_0042 | 789 | NA | NA | NA | NA | NA | NA | NA | NA | NA | NA |
| NA | NA | NA | NA | NA | 44580 | 45278 | trpG_2 | BCG_0043 | 699 | NA | NA | NA | NA | NA | NA | NA | NA | NA | NA |
| NA | NA | NA | NA | NA | 45256 | 47136 | pknB | BCG_0044c | 1881 | NA | NA | NA | NA | NA | NA | NA | NA | NA | NA |
| NA | NA | NA | NA | NA | 722772 | 722912 | mce2Db | BCG_0638 | 141 | NA | NA | NA | NA | NA | NA | NA | NA | NA | NA |
| NA | NA | NA | NA | NA | 830619 | 830852 | BCG_0748 | BCG_0748 | 234 | NA | NA | NA | NA | NA | 801077 | 801310 | BCGDan_sapMKO_0726 | BCGDan_sapMKO_0726 | 234 |
| NA | NA | NA | NA | NA | 1758533 | 1759228 | BCG_1597 | BCG_1597 | 696 | NA | NA | NA | NA | NA | NA | NA | NA | NA | NA |
| NA | NA | NA | NA | NA | 2052848 | 2053696 | PPE33b | BCG_1843 | 849 | NA | NA | NA | NA | NA | 2033843 | 2034577 | BCGDan_sapMKO_1850 | BCGDan_sapMKO_1850 | 735 |
| 2052933 | 2053289 | Rv1810 | Rv1810 | 357 | 2053935 | 2054291 | BCG_1844 | BCG_1844 | 357 | NA | NA | NA | NA | NA | NA | NA | NA | NA | NA |
| NA | NA | NA | NA | NA | 2060520 | 2060600 | BCG_1851 | BCG_1851 | 81 | NA | NA | NA | NA | NA | 2040772 | 2040852 | BCGDan_sapMKO_1858 | BCGDan_sapMKO_1858 | 81 |
| NA | NA | NA | NA | NA | 3007599 | 3007631 | PE_PGRS47a | BCG_2755 | 33 | NA | NA | NA | NA | NA | NA | NA | NA | NA | NA |
| 3378329 | 3378415 | PE27A | Rv3018A | 87 | 3332760 | 3332846 | PE27A | BCG_3041c | 87 | NA | NA | NA | NA | NA | NA | NA | NA | NA | NA |
| 65012 | 65350 | Rv0061c | Rv0061c | 339 | NA | NA | NA | NA | NA | NA | NA | NA | NA | NA | NA | NA | NA | NA | NA |
| 186495 | 186623 | Rv0157A | Rv0157A | 129 | NA | NA | NA | NA | NA | NA | NA | NA | NA | NA | NA | NA | NA | NA | NA |
| 218390 | 218551 | mymT | Rv0186A | 162 | NA | NA | NA | NA | NA | NA | NA | NA | NA | NA | NA | NA | NA | NA | NA |
| 223607 | 223909 | Rv0192A | Rv0192A | 303 | NA | NA | NA | NA | NA | NA | NA | NA | NA | NA | NA | NA | NA | NA | NA |
| 265507 | 266295 | echA1 | Rv0222 | 789 | NA | NA | NA | NA | NA | NA | NA | NA | NA | NA | NA | NA | NA | NA | NA |
| 392273 | 392728 | Rv0326 | Rv0326 | 456 | NA | NA | NA | NA | NA | NA | NA | NA | NA | NA | NA | NA | NA | NA | NA |
| 424777 | 434679 | PPE8 | Rv0355c | 9903 | NA | NA | NA | NA | NA | NA | NA | NA | NA | NA | NA | NA | NA | NA | NA |
| 467459 | 468001 | PPE9 | Rv0388c | 543 | NA | NA | NA | NA | NA | NA | NA | NA | NA | NA | NA | NA | NA | NA | NA |
| 476394 | 476642 | Rv0397A | Rv0397A | 249 | NA | NA | NA | NA | NA | NA | NA | NA | NA | NA | NA | NA | NA | NA | NA |
| 547344 | 547517 | mazE1 | Rv0456B | 174 | NA | NA | NA | NA | NA | NA | NA | NA | NA | NA | NA | NA | NA | NA | NA |
| 591475 | 591576 | Rv0500B | Rv0500B | 102 | NA | NA | NA | NA | NA | NA | NA | NA | NA | NA | NA | NA | NA | NA | NA |
| 622121 | 622282 | Rv0530A | Rv0530A | 162 | NA | NA | NA | NA | NA | NA | NA | NA | NA | NA | NA | NA | NA | NA | NA |
| 688808 | 689062 | Rv0590A | Rv0590A | 255 | NA | NA | NA | NA | NA | NA | NA | NA | NA | NA | NA | NA | NA | NA | NA |
| 712174 | 712719 | galTb | Rv0619 | 546 | NA | NA | NA | NA | NA | NA | NA | NA | NA | NA | NA | NA | NA | NA | NA |
| 817531 | 817866 | Rv0724A | Rv0724A | 336 | NA | NA | NA | NA | NA | NA | NA | NA | NA | NA | NA | NA | NA | NA | NA |
| 874732 | 876390 | ptrBb | Rv0782 | 1659 | NA | NA | NA | NA | NA | NA | NA | NA | NA | NA | NA | NA | NA | NA | NA |
| 889072 | 889398 | Rv0795 | Rv0795 | 327 | NA | NA | NA | NA | NA | NA | NA | NA | NA | NA | NA | NA | NA | NA | NA |
| 889347 | 890333 | Rv0796 | Rv0796 | 987 | NA | NA | NA | NA | NA | NA | NA | NA | NA | NA | NA | NA | NA | NA | NA |
| 925361 | 927610 | PE_PGRS13 | Rv0833 | 2250 | NA | NA | NA | NA | NA | NA | NA | NA | NA | NA | NA | NA | NA | NA | NA |
| 1273355 | 1274767 | mmpL13b | Rv1146 | 1413 | NA | NA | NA | NA | NA | NA | NA | NA | NA | NA | NA | NA | NA | NA | NA |
| 1315234 | 1319982 | pks4 | Rv1181 | 4749 | NA | NA | NA | NA | NA | NA | NA | NA | NA | NA | NA | NA | NA | NA | NA |
| 1333931 | 1334845 | Rv1191 | Rv1191 | 915 | NA | NA | NA | NA | NA | NA | NA | NA | NA | NA | NA | NA | NA | NA | NA |
| 1403386 | 1404603 | cyp130 | Rv1256c | 1218 | NA | NA | NA | NA | NA | NA | NA | NA | NA | NA | NA | NA | NA | NA | NA |
| 1404717 | 1406084 | Rv1257c | Rv1257c | 1368 | NA | NA | NA | NA | NA | NA | NA | NA | NA | NA | NA | NA | NA | NA | NA |
| 1539180 | 1539440 | Rv1366A | Rv1366A | 261 | NA | NA | NA | NA | NA | NA | NA | NA | NA | NA | NA | NA | NA | NA | NA |
| 1541994 | 1542980 | Rv1369c | Rv1369c | 987 | NA | NA | NA | NA | NA | NA | NA | NA | NA | NA | NA | NA | NA | NA | NA |
| 1542929 | 1543255 | Rv1370c | Rv1370c | 327 | NA | NA | NA | NA | NA | NA | NA | NA | NA | NA | NA | NA | NA | NA | NA |
| 1694545 | 1695144 | Rv1504c | Rv1504c | 600 | NA | NA | NA | NA | NA | NA | NA | NA | NA | NA | NA | NA | NA | NA | NA |
| 1695943 | 1696443 | Rv1506c | Rv1506c | 501 | NA | NA | NA | NA | NA | NA | NA | NA | NA | NA | NA | NA | NA | NA | NA |
| 1696727 | 1697422 | Rv1507c | Rv1507c | 696 | NA | NA | NA | NA | NA | NA | NA | NA | NA | NA | NA | NA | NA | NA | NA |
| 1697356 | 1697859 | Rv1507A | Rv1507A | 504 | NA | NA | NA | NA | NA | NA | NA | NA | NA | NA | NA | NA | NA | NA | NA |
| 1698095 | 1699894 | Rv1508c | Rv1508c | 1800 | NA | NA | NA | NA | NA | NA | NA | NA | NA | NA | NA | NA | NA | NA | NA |
| 1699866 | 1700228 | Rv1508A | Rv1508A | 363 | NA | NA | NA | NA | NA | NA | NA | NA | NA | NA | NA | NA | NA | NA | NA |
| 1700212 | 1701093 | Rv1509 | Rv1509 | 882 | NA | NA | NA | NA | NA | NA | NA | NA | NA | NA | NA | NA | NA | NA | NA |
| 1701295 | 1702593 | Rv1510 | Rv1510 | 1299 | NA | NA | NA | NA | NA | NA | NA | NA | NA | NA | NA | NA | NA | NA | NA |
| 1703074 | 1704096 | gmdA | Rv1511 | 1023 | NA | NA | NA | NA | NA | NA | NA | NA | NA | NA | NA | NA | NA | NA | NA |
| 1704093 | 1705061 | epiA | Rv1512 | 969 | NA | NA | NA | NA | NA | NA | NA | NA | NA | NA | NA | NA | NA | NA | NA |
| 1705058 | 1705789 | Rv1513 | Rv1513 | 732 | NA | NA | NA | NA | NA | NA | NA | NA | NA | NA | NA | NA | NA | NA | NA |
| 1705807 | 1706595 | Rv1514c | Rv1514c | 789 | NA | NA | NA | NA | NA | NA | NA | NA | NA | NA | NA | NA | NA | NA | NA |
| 1706630 | 1707526 | Rv1515c | Rv1515c | 897 | NA | NA | NA | NA | NA | NA | NA | NA | NA | NA | NA | NA | NA | NA | NA |
| 1707529 | 1708539 | Rv1516c | Rv1516c | 1011 | NA | NA | NA | NA | NA | NA | NA | NA | NA | NA | NA | NA | NA | NA | NA |
| 1753716 | 1755431 | fadD11 | Rv1550 | 1716 | NA | NA | NA | NA | NA | NA | NA | NA | NA | NA | NA | NA | NA | NA | NA |
| 1760175 | 1760555 | frdC | Rv1554 | 381 | NA | NA | NA | NA | NA | NA | NA | NA | NA | NA | NA | NA | NA | NA | NA |
| 1779314 | 1779724 | Rv1573 | Rv1573 | 411 | NA | NA | NA | NA | NA | NA | NA | NA | NA | NA | NA | NA | NA | NA | NA |
| 1779930 | 1780241 | Rv1574 | Rv1574 | 312 | NA | NA | NA | NA | NA | NA | NA | NA | NA | NA | NA | NA | NA | NA | NA |
| 1780199 | 1780699 | Rv1575 | Rv1575 | 501 | NA | NA | NA | NA | NA | NA | NA | NA | NA | NA | NA | NA | NA | NA | NA |
| 1780643 | 1782064 | Rv1576c | Rv1576c | 1422 | NA | NA | NA | NA | NA | NA | NA | NA | NA | NA | NA | NA | NA | NA | NA |
| 1782072 | 1782584 | Rv1577c | Rv1577c | 513 | NA | NA | NA | NA | NA | NA | NA | NA | NA | NA | NA | NA | NA | NA | NA |
| 1782758 | 1783228 | Rv1578c | Rv1578c | 471 | NA | NA | NA | NA | NA | NA | NA | NA | NA | NA | NA | NA | NA | NA | NA |
| 1783309 | 1783623 | Rv1579c | Rv1579c | 315 | NA | NA | NA | NA | NA | NA | NA | NA | NA | NA | NA | NA | NA | NA | NA |
| 1783620 | 1783892 | Rv1580c | Rv1580c | 273 | NA | NA | NA | NA | NA | NA | NA | NA | NA | NA | NA | NA | NA | NA | NA |
| 1783906 | 1784301 | Rv1581c | Rv1581c | 396 | NA | NA | NA | NA | NA | NA | NA | NA | NA | NA | NA | NA | NA | NA | NA |
| 1784497 | 1785912 | Rv1582c | Rv1582c | 1416 | NA | NA | NA | NA | NA | NA | NA | NA | NA | NA | NA | NA | NA | NA | NA |
| 1785912 | 1786310 | Rv1583c | Rv1583c | 399 | NA | NA | NA | NA | NA | NA | NA | NA | NA | NA | NA | NA | NA | NA | NA |
| 1786307 | 1786528 | Rv1584c | Rv1584c | 222 | NA | NA | NA | NA | NA | NA | NA | NA | NA | NA | NA | NA | NA | NA | NA |
| 1786584 | 1787099 | Rv1585c | Rv1585c | 516 | NA | NA | NA | NA | NA | NA | NA | NA | NA | NA | NA | NA | NA | NA | NA |
| 1787096 | 1788505 | Rv1586c | Rv1586c | 1410 | NA | NA | NA | NA | NA | NA | NA | NA | NA | NA | NA | NA | NA | NA | NA |
| 1788162 | 1789163 | Rv1587c | Rv1587c | 1002 | NA | NA | NA | NA | NA | NA | NA | NA | NA | NA | NA | NA | NA | NA | NA |
| 1894224 | 1895342 | Rv1668c | Rv1668c | 1119 | NA | NA | NA | NA | NA | NA | NA | NA | NA | NA | NA | NA | NA | NA | NA |
| 1987745 | 1988731 | Rv1756c | Rv1756c | 987 | NA | NA | NA | NA | NA | NA | NA | NA | NA | NA | NA | NA | NA | NA | NA |
| 1988680 | 1989006 | Rv1757c | Rv1757c | 327 | NA | NA | NA | NA | NA | NA | NA | NA | NA | NA | NA | NA | NA | NA | NA |
| 1996152 | 1996478 | Rv1763 | Rv1763 | 327 | NA | NA | NA | NA | NA | NA | NA | NA | NA | NA | NA | NA | NA | NA | NA |
| 1996427 | 1997413 | Rv1764 | Rv1764 | 987 | NA | NA | NA | NA | NA | NA | NA | NA | NA | NA | NA | NA | NA | NA | NA |
| 2161566 | 2162762 | aceAb | Rv1916 | 1197 | NA | NA | NA | NA | NA | NA | NA | NA | NA | NA | NA | NA | NA | NA | NA |
| 2201277 | 2201579 | Rv1954A | Rv1954A | 303 | NA | NA | NA | NA | NA | NA | NA | NA | NA | NA | NA | NA | NA | NA | NA |
| 2208507 | 2209322 | yrbE3B | Rv1965 | 816 | NA | NA | NA | NA | NA | NA | NA | NA | NA | NA | NA | NA | NA | NA | NA |

| | | | | | | | | | | | | | | | | | | | |
|---|---|---|---|---|---|---|---|---|---|---|---|---|---|---|---|---|---|---|---|
| 2209327 | 2210604 | mce3A | Rv1966 | 1278 | NA | NA | NA | NA | NA | NA | NA | NA | NA | NA | NA | NA | NA | NA | NA |
| 2210601 | 2211629 | mce3B | Rv1967 | 1029 | NA | NA | NA | NA | NA | NA | NA | NA | NA | NA | NA | NA | NA | NA | NA |
| 2211626 | 2212858 | mce3C | Rv1968 | 1233 | NA | NA | NA | NA | NA | NA | NA | NA | NA | NA | NA | NA | NA | NA | NA |
| 2212855 | 2214126 | mce3D | Rv1969 | 1272 | NA | NA | NA | NA | NA | NA | NA | NA | NA | NA | NA | NA | NA | NA | NA |
| 2214123 | 2215256 | lprM | Rv1970 | 1134 | NA | NA | NA | NA | NA | NA | NA | NA | NA | NA | NA | NA | NA | NA | NA |
| 2215257 | 2216570 | mce3F | Rv1971 | 1314 | NA | NA | NA | NA | NA | NA | NA | NA | NA | NA | NA | NA | NA | NA | NA |
| 2216592 | 2217167 | Rv1972 | Rv1972 | 576 | NA | NA | NA | NA | NA | NA | NA | NA | NA | NA | NA | NA | NA | NA | NA |
| 2217164 | 2217646 | Rv1973 | Rv1973 | 483 | NA | NA | NA | NA | NA | NA | NA | NA | NA | NA | NA | NA | NA | NA | NA |
| 2217659 | 2218036 | Rv1974 | Rv1974 | 378 | NA | NA | NA | NA | NA | NA | NA | NA | NA | NA | NA | NA | NA | NA | NA |
| 2218052 | 2218717 | Rv1975 | Rv1975 | 666 | NA | NA | NA | NA | NA | NA | NA | NA | NA | NA | NA | NA | NA | NA | NA |
| 2218844 | 2219251 | Rv1976c | Rv1976c | 408 | NA | NA | NA | NA | NA | NA | NA | NA | NA | NA | NA | NA | NA | NA | NA |
| 2219754 | 2220800 | Rv1977 | Rv1977 | 1047 | NA | NA | NA | NA | NA | NA | NA | NA | NA | NA | NA | NA | NA | NA | NA |
| 2220908 | 2221756 | Rv1978 | Rv1978 | 849 | NA | NA | NA | NA | NA | NA | NA | NA | NA | NA | NA | NA | NA | NA | NA |
| 2221719 | 2223164 | Rv1979c | Rv1979c | 1446 | NA | NA | NA | NA | NA | NA | NA | NA | NA | NA | NA | NA | NA | NA | NA |
| 2223343 | 2224029 | mpt64 | Rv1980c | 687 | NA | NA | NA | NA | NA | NA | NA | NA | NA | NA | NA | NA | NA | NA | NA |
| 2224220 | 2225188 | nrdF1 | Rv1981c | 969 | NA | NA | NA | NA | NA | NA | NA | NA | NA | NA | NA | NA | NA | NA | NA |
| 2225413 | 2225832 | vapC36 | Rv1982c | 420 | NA | NA | NA | NA | NA | NA | NA | NA | NA | NA | NA | NA | NA | NA | NA |
| 2225841 | 2226101 | vapB36 | Rv1982A | 261 | NA | NA | NA | NA | NA | NA | NA | NA | NA | NA | NA | NA | NA | NA | NA |
| 2226244 | 2227920 | PE_PGRS35 | Rv1983 | 1677 | NA | NA | NA | NA | NA | NA | NA | NA | NA | NA | NA | NA | NA | NA | NA |
| 2227908 | 2228561 | cfp21 | Rv1984c | 654 | NA | NA | NA | NA | NA | NA | NA | NA | NA | NA | NA | NA | NA | NA | NA |
| 2228991 | 2229902 | Rv1985c | Rv1985c | 912 | NA | NA | NA | NA | NA | NA | NA | NA | NA | NA | NA | NA | NA | NA | NA |
| 2230011 | 2230610 | Rv1986 | Rv1986 | 600 | NA | NA | NA | NA | NA | NA | NA | NA | NA | NA | NA | NA | NA | NA | NA |
| 2231026 | 2231454 | Rv1987 | Rv1987 | 429 | NA | NA | NA | NA | NA | NA | NA | NA | NA | NA | NA | NA | NA | NA | NA |
| 2321057 | 2321467 | mazF7 | Rv2063A | 411 | NA | NA | NA | NA | NA | NA | NA | NA | NA | NA | NA | NA | NA | NA | NA |
| 2330214 | 2330963 | Rv2073c | Rv2073c | 750 | NA | NA | NA | NA | NA | NA | NA | NA | NA | NA | NA | NA | NA | NA | NA |
| 2330993 | 2331406 | Rv2074 | Rv2074 | 414 | NA | NA | NA | NA | NA | NA | NA | NA | NA | NA | NA | NA | NA | NA | NA |
| 2358033 | 2358206 | PE21 | Rv2099c | 174 | NA | NA | NA | NA | NA | NA | NA | NA | NA | NA | NA | NA | NA | NA | NA |
| 2365465 | 2365791 | Rv2105 | Rv2105 | 327 | NA | NA | NA | NA | NA | NA | NA | NA | NA | NA | NA | NA | NA | NA | NA |
| 2365740 | 2366726 | Rv2106 | Rv2106 | 987 | NA | NA | NA | NA | NA | NA | NA | NA | NA | NA | NA | NA | NA | NA | NA |
| 2421662 | 2422003 | Rv2160c | Rv2160c | 342 | NA | NA | NA | NA | NA | NA | NA | NA | NA | NA | NA | NA | NA | NA | NA |
| 2430159 | 2431145 | Rv2167c | Rv2167c | 987 | NA | NA | NA | NA | NA | NA | NA | NA | NA | NA | NA | NA | NA | NA | NA |
| 2431094 | 2431420 | Rv2168c | Rv2168c | 327 | NA | NA | NA | NA | NA | NA | NA | NA | NA | NA | NA | NA | NA | NA | NA |
| 2510351 | 2510587 | Rv2237A | Rv2237A | 237 | NA | NA | NA | NA | NA | NA | NA | NA | NA | NA | NA | NA | NA | NA | NA |
| 2525565 | 2526992 | Rv2251 | Rv2251 | 1428 | NA | NA | NA | NA | NA | NA | NA | NA | NA | NA | NA | NA | NA | NA | NA |
| 2534470 | 2535552 | Rv2262c | Rv2262c | 1083 | NA | NA | NA | NA | NA | NA | NA | NA | NA | NA | NA | NA | NA | NA | NA |
| 2546839 | 2547087 | mazE8 | Rv2274A | 249 | NA | NA | NA | NA | NA | NA | NA | NA | NA | NA | NA | NA | NA | NA | NA |
| 2550065 | 2550391 | Rv2278 | Rv2278 | 327 | NA | NA | NA | NA | NA | NA | NA | NA | NA | NA | NA | NA | NA | NA | NA |
| 2550340 | 2551326 | Rv2279 | Rv2279 | 987 | NA | NA | NA | NA | NA | NA | NA | NA | NA | NA | NA | NA | NA | NA | NA |
| 2564292 | 2565032 | Rv2293c | Rv2293c | 741 | NA | NA | NA | NA | NA | NA | NA | NA | NA | NA | NA | NA | NA | NA | NA |
| 2626223 | 2626519 | esxP | Rv2347c | 297 | NA | NA | NA | NA | NA | NA | NA | NA | NA | NA | NA | NA | NA | NA | NA |
| 2626654 | 2626980 | Rv2348c | Rv2348c | 327 | NA | NA | NA | NA | NA | NA | NA | NA | NA | NA | NA | NA | NA | NA | NA |
| 2627172 | 2628698 | plcC | Rv2349c | 1527 | NA | NA | NA | NA | NA | NA | NA | NA | NA | NA | NA | NA | NA | NA | NA |
| 2628781 | 2630319 | plcB | Rv2350c | 1539 | NA | NA | NA | NA | NA | NA | NA | NA | NA | NA | NA | NA | NA | NA | NA |
| 2630537 | 2632075 | plcA | Rv2351c | 1539 | NA | NA | NA | NA | NA | NA | NA | NA | NA | NA | NA | NA | NA | NA | NA |
| 2632923 | 2634098 | PPE38 | Rv2352c | 1176 | NA | NA | NA | NA | NA | NA | NA | NA | NA | NA | NA | NA | NA | NA | NA |
| 2634528 | 2635592 | PPE39 | Rv2353c | 1065 | NA | NA | NA | NA | NA | NA | NA | NA | NA | NA | NA | NA | NA | NA | NA |
| 2635628 | 2635954 | Rv2354 | Rv2354 | 327 | NA | NA | NA | NA | NA | NA | NA | NA | NA | NA | NA | NA | NA | NA | NA |
| 2635903 | 2636889 | Rv2355 | Rv2355 | 987 | NA | NA | NA | NA | NA | NA | NA | NA | NA | NA | NA | NA | NA | NA | NA |
| 2637688 | 2639535 | PPE40 | Rv2356c | 1848 | NA | NA | NA | NA | NA | NA | NA | NA | NA | NA | NA | NA | NA | NA | NA |
| 2692224 | 2692439 | aprA | Rv2395A | 216 | NA | NA | NA | NA | NA | NA | NA | NA | NA | NA | NA | NA | NA | NA | NA |
| 2692551 | 2692715 | aprB | Rv2395B | 165 | NA | NA | NA | NA | NA | NA | NA | NA | NA | NA | NA | NA | NA | NA | NA |
| 2784657 | 2785643 | Rv2479c | Rv2479c | 987 | NA | NA | NA | NA | NA | NA | NA | NA | NA | NA | NA | NA | NA | NA | NA |
| 2785592 | 2785918 | Rv2480c | Rv2480c | 327 | NA | NA | NA | NA | NA | NA | NA | NA | NA | NA | NA | NA | NA | NA | NA |
| 2869253 | 2869627 | Rv2548A | Rv2548A | 375 | NA | NA | NA | NA | NA | NA | NA | NA | NA | NA | NA | NA | NA | NA | NA |
| 2881758 | 2882147 | Rv2562 | Rv2562 | 390 | NA | NA | NA | NA | NA | NA | NA | NA | NA | NA | NA | NA | NA | NA | NA |
| 2970123 | 2970554 | Rv2645 | Rv2645 | 432 | NA | NA | NA | NA | NA | NA | NA | NA | NA | NA | NA | NA | NA | NA | NA |
| 2970551 | 2971549 | Rv2646 | Rv2646 | 999 | NA | NA | NA | NA | NA | NA | NA | NA | NA | NA | NA | NA | NA | NA | NA |
| 2971659 | 2972027 | Rv2647 | Rv2647 | 369 | NA | NA | NA | NA | NA | NA | NA | NA | NA | NA | NA | NA | NA | NA | NA |
| 2972160 | 2972486 | Rv2648 | Rv2648 | 327 | NA | NA | NA | NA | NA | NA | NA | NA | NA | NA | NA | NA | NA | NA | NA |
| 2972435 | 2973421 | Rv2649 | Rv2649 | 987 | NA | NA | NA | NA | NA | NA | NA | NA | NA | NA | NA | NA | NA | NA | NA |
| 2973795 | 2975234 | Rv2650c | Rv2650c | 1440 | NA | NA | NA | NA | NA | NA | NA | NA | NA | NA | NA | NA | NA | NA | NA |
| 2975242 | 2975775 | Rv2651c | Rv2651c | 534 | NA | NA | NA | NA | NA | NA | NA | NA | NA | NA | NA | NA | NA | NA | NA |
| 2975928 | 2976554 | Rv2652c | Rv2652c | 627 | NA | NA | NA | NA | NA | NA | NA | NA | NA | NA | NA | NA | NA | NA | NA |
| 2976586 | 2976909 | Rv2653c | Rv2653c | 324 | NA | NA | NA | NA | NA | NA | NA | NA | NA | NA | NA | NA | NA | NA | NA |
| 2976989 | 2977234 | Rv2654c | Rv2654c | 246 | NA | NA | NA | NA | NA | NA | NA | NA | NA | NA | NA | NA | NA | NA | NA |
| 2977231 | 2978658 | Rv2655c | Rv2655c | 1428 | NA | NA | NA | NA | NA | NA | NA | NA | NA | NA | NA | NA | NA | NA | NA |
| 2978660 | 2979052 | Rv2656c | Rv2656c | 393 | NA | NA | NA | NA | NA | NA | NA | NA | NA | NA | NA | NA | NA | NA | NA |
| 2979049 | 2979309 | Rv2657c | Rv2657c | 261 | NA | NA | NA | NA | NA | NA | NA | NA | NA | NA | NA | NA | NA | NA | NA |
| 2979326 | 2979688 | Rv2658c | Rv2658c | 363 | NA | NA | NA | NA | NA | NA | NA | NA | NA | NA | NA | NA | NA | NA | NA |
| 2979691 | 2980818 | Rv2659c | Rv2659c | 1128 | NA | NA | NA | NA | NA | NA | NA | NA | NA | NA | NA | NA | NA | NA | NA |
| 3174747 | 3174995 | vapB23 | Rv2862A | 249 | NA | NA | NA | NA | NA | NA | NA | NA | NA | NA | NA | NA | NA | NA | NA |
| 3189851 | 3190678 | Rv2880c | Rv2880c | 828 | NA | NA | NA | NA | NA | NA | NA | NA | NA | NA | NA | NA | NA | NA | NA |
| 3289790 | 3290506 | Rv2944 | Rv2944 | 717 | NA | NA | NA | NA | NA | NA | NA | NA | NA | NA | NA | NA | NA | NA | NA |
| 3296350 | 3297840 | pks15 | Rv2947c | 1491 | NA | NA | NA | NA | NA | NA | NA | NA | NA | NA | NA | NA | NA | NA | NA |
| 3331358 | 3331612 | Rv2975c | Rv2975c | 255 | NA | NA | NA | NA | NA | NA | NA | NA | NA | NA | NA | NA | NA | NA | NA |
| 3380440 | 3380682 | PPE48 | Rv3022c | 243 | NA | NA | NA | NA | NA | NA | NA | NA | NA | NA | NA | NA | NA | NA | NA |
| 3467606 | 3467926 | Rv3098A | Rv3098A | 321 | NA | NA | NA | NA | NA | NA | NA | NA | NA | NA | NA | NA | NA | NA | NA |
| 3484809 | 3485111 | sseC1 | Rv3118 | 303 | NA | NA | NA | NA | NA | NA | NA | NA | NA | NA | NA | NA | NA | NA | NA |
| 3485132 | 3485575 | moaE1 | Rv3119 | 444 | NA | NA | NA | NA | NA | NA | NA | NA | NA | NA | NA | NA | NA | NA | NA |
| 3485572 | 3486174 | Rv3120 | Rv3120 | 603 | NA | NA | NA | NA | NA | NA | NA | NA | NA | NA | NA | NA | NA | NA | NA |
| 3486509 | 3487711 | cyp141 | Rv3121 | 1203 | NA | NA | NA | NA | NA | NA | NA | NA | NA | NA | NA | NA | NA | NA | NA |
| 3502945 | 3503277 | Rv3136A | Rv3136A | 333 | NA | NA | NA | NA | NA | NA | NA | NA | NA | NA | NA | NA | NA | NA | NA |
| 3551281 | 3551607 | Rv3184 | Rv3184 | 327 | NA | NA | NA | NA | NA | NA | NA | NA | NA | NA | NA | NA | NA | NA | NA |
| 3551556 | 3552542 | Rv3185 | Rv3185 | 987 | NA | NA | NA | NA | NA | NA | NA | NA | NA | NA | NA | NA | NA | NA | NA |
| 3552764 | 3553090 | Rv3186 | Rv3186 | 327 | NA | NA | NA | NA | NA | NA | NA | NA | NA | NA | NA | NA | NA | NA | NA |
| 3553039 | 3554025 | Rv3187 | Rv3187 | 987 | NA | NA | NA | NA | NA | NA | NA | NA | NA | NA | NA | NA | NA | NA | NA |

| | | | | | | | | | | | | | | | | | | | |
|---|---|---|---|---|---|---|---|---|---|---|---|---|---|---|---|---|---|---|---|
| 3556855 | 3557064 | Rv3190A | Rv3190A | 210 | NA | NA | NA | NA | NA | NA | NA | NA | NA | NA | NA | NA | NA | NA | NA |
| 3593369 | 3593437 | Rv3216 | Rv3216 | 69 | NA | NA | NA | NA | NA | NA | NA | NA | NA | NA | NA | NA | NA | NA | NA |
| 3610374 | 3611189 | tgs3 | Rv3234c | 816 | NA | NA | NA | NA | NA | NA | NA | NA | NA | NA | NA | NA | NA | NA | NA |
| 3710433 | 3710759 | Rv3325 | Rv3325 | 327 | NA | NA | NA | NA | NA | NA | NA | NA | NA | NA | NA | NA | NA | NA | NA |
| 3710708 | 3711694 | Rv3326 | Rv3326 | 987 | NA | NA | NA | NA | NA | NA | NA | NA | NA | NA | NA | NA | NA | NA | NA |
| 3723904 | 3724548 | Rv3338 | Rv3338 | 645 | NA | NA | NA | NA | NA | NA | NA | NA | NA | NA | NA | NA | NA | NA | NA |
| 3738158 | 3742774 | PE_PGRS50 | Rv3345c | 4617 | NA | NA | NA | NA | NA | NA | NA | NA | NA | NA | NA | NA | NA | NA | NA |
| 3788368 | 3788616 | echA18.1 | Rv3374 | 249 | NA | NA | NA | NA | NA | NA | NA | NA | NA | NA | NA | NA | NA | NA | NA |
| 3795100 | 3796086 | Rv3380c | Rv3380c | 987 | NA | NA | NA | NA | NA | NA | NA | NA | NA | NA | NA | NA | NA | NA | NA |
| 3796035 | 3796361 | Rv3381c | Rv3381c | 327 | NA | NA | NA | NA | NA | NA | NA | NA | NA | NA | NA | NA | NA | NA | NA |
| 3843036 | 3843734 | PPE58 | Rv3426 | 699 | NA | NA | NA | NA | NA | NA | NA | NA | NA | NA | NA | NA | NA | NA | NA |
| 3843885 | 3844640 | Rv3427c | Rv3427c | 756 | NA | NA | NA | NA | NA | NA | NA | NA | NA | NA | NA | NA | NA | NA | NA |
| 3844738 | 3845970 | Rv3428c | Rv3428c | 1233 | NA | NA | NA | NA | NA | NA | NA | NA | NA | NA | NA | NA | NA | NA | NA |
| 3847165 | 3847701 | PPE59 | Rv3429 | 537 | NA | NA | NA | NA | NA | NA | NA | NA | NA | NA | NA | NA | NA | NA | NA |
| 3874822 | 3876090 | Rv3454 | Rv3454 | 1269 | NA | NA | NA | NA | NA | NA | NA | NA | NA | NA | NA | NA | NA | NA | NA |
| 3890830 | 3891156 | Rv3474 | Rv3474 | 327 | NA | NA | NA | NA | NA | NA | NA | NA | NA | NA | NA | NA | NA | NA | NA |
| 3891105 | 3892091 | Rv3475 | Rv3475 | 987 | NA | NA | NA | NA | NA | NA | NA | NA | NA | NA | NA | NA | NA | NA | NA |
| 3941724 | 3944963 | PE_PGRS56 | Rv3512 | 3240 | NA | NA | NA | NA | NA | NA | NA | NA | NA | NA | NA | NA | NA | NA | NA |
| 4042952 | 4043035 | Rv3599c | Rv3599c | 84 | NA | NA | NA | NA | NA | NA | NA | NA | NA | NA | NA | NA | NA | NA | NA |
| 4057733 | 4058701 | ephA | Rv3617 | 969 | NA | NA | NA | NA | NA | NA | NA | NA | NA | NA | NA | NA | NA | NA | NA |
| 4058698 | 4059885 | Rv3618 | Rv3618 | 1188 | NA | NA | NA | NA | NA | NA | NA | NA | NA | NA | NA | NA | NA | NA | NA |
| 4059984 | 4060268 | esxV | Rv3619c | 285 | NA | NA | NA | NA | NA | NA | NA | NA | NA | NA | NA | NA | NA | NA | NA |
| 4060295 | 4060591 | esxW | Rv3620c | 297 | NA | NA | NA | NA | NA | NA | NA | NA | NA | NA | NA | NA | NA | NA | NA |
| 4060648 | 4061889 | PPE65 | Rv3621c | 1242 | NA | NA | NA | NA | NA | NA | NA | NA | NA | NA | NA | NA | NA | NA | NA |
| 4061899 | 4062198 | PE32 | Rv3622c | 300 | NA | NA | NA | NA | NA | NA | NA | NA | NA | NA | NA | NA | NA | NA | NA |
| 4169606 | 4170169 | cut5b | Rv3724B | 564 | NA | NA | NA | NA | NA | NA | NA | NA | NA | NA | NA | NA | NA | NA | NA |
| 4190284 | 4190517 | PPE67 | Rv3739c | 234 | NA | NA | NA | NA | NA | NA | NA | NA | NA | NA | NA | NA | NA | NA | NA |
| 4350745 | 4351044 | PE35 | Rv3872 | 300 | NA | NA | NA | NA | NA | NA | NA | NA | NA | NA | NA | NA | NA | NA | NA |
| 4351075 | 4352181 | PPE68 | Rv3873 | 1107 | NA | NA | NA | NA | NA | NA | NA | NA | NA | NA | NA | NA | NA | NA | NA |
| 4352274 | 4352576 | esxB | Rv3874 | 303 | NA | NA | NA | NA | NA | NA | NA | NA | NA | NA | NA | NA | NA | NA | NA |
| 4352609 | 4352896 | esxA | Rv3875 | 288 | NA | NA | NA | NA | NA | NA | NA | NA | NA | NA | NA | NA | NA | NA | NA |
| 4353010 | 4355010 | espI | Rv3876 | 2001 | NA | NA | NA | NA | NA | NA | NA | NA | NA | NA | NA | NA | NA | NA | NA |
| 4355007 | 4356542 | eccD1 | Rv3877 | 1536 | NA | NA | NA | NA | NA | NA | NA | NA | NA | NA | NA | NA | NA | NA | NA |
| 4356693 | 4357535 | espJ | Rv3878 | 843 | NA | NA | NA | NA | NA | NA | NA | NA | NA | NA | NA | NA | NA | NA | NA |
| 4357593 | 4359782 | espK | Rv3879c | 2190 | NA | NA | NA | NA | NA | NA | NA | NA | NA | NA | NA | NA | NA | NA | NA |
| 4383653 | 4383985 | Rv3898c | Rv3898c | 333 | NA | NA | NA | NA | NA | NA | NA | NA | NA | NA | NA | NA | NA | NA | NA |
| NA | NA | NA | NA | NA | NA | NA | NA | NA | NA | NA | NA | NA | NA | NA | 2040695 | 2040775 | BCGDan_sapMKO_1857 | BCGDan_sapMKO_1857 | 81 |
