## Supplemental Table 7 Worksheet 2 for "Reference genome for the WHO reference strain for *Mycobacterium bovis* BCG Danish, the present tuberculosis vaccine"

| *M. bovis* BCG Pasteur 1173P2 (AM408590.1, duplicated CDS) | | | | | | | | | | *M. bovis* BCG Danish 1331 (NC..... (in process), duplicated CDS) | | | | | | | | | | *M. bovis* BCG Danish 1331 *sapM* KO (NC..... (in process), duplicated CDS) | | | | | | | | | |
|---|---|---|---|---|---|---|---|---|---|---|---|---|---|---|---|---|---|---|---|---|---|---|---|---|---|---|---|---|---|
| **start Past** | **end Past** | **CDS nam** | **locus_tag** | **length P** | **start Past** | **end Past** | **CDS nam** | **locus_tag** | **length P** | **start Dan** | **end Dani** | **CDS name/locus_ta** | **locus_tag Danish** | **length D** | **start Dan** | **end Dani** | **CDS name/locus_tag Da** | **locus_tag Danish duplicated** | **length D** | **start Dan** | **end Dani** | **CDS nam** | **locus_tag** | **length Da** | **start Dan** | **end Dani** | **CDS nam** | **locus_tag** | **length Danish sapM KO duplicated** |
| 1 | 1524 | dnaA_1 | BCG_000 | 1524 | 29668 | 31191 | dnaA_2 | BCG_003 | 1524 | 1 | 1524 | dnaA_1 | BCGDan_0001 | 1524 | 4397238 | 4398761 | dnaA_2 | BCGDan_4070 | 1524 | 1 | 1524 | dnaA_1 | BCGDan_ | 1524 | 4318155 | 4319645 | dnaA_2 | BCGDan_ | 1491 |
| 2052 | 3260 | dnaN_1 | BCG_000 | 1209 | 31719 | 32927 | dnaN_2 | BCG_003 | 1209 | 2052 | 3260 | dnaN_1 | BCGDan_0002 | 1209 | 4399289 | 4400497 | dnaN_2 | BCGDan_4071 | 1209 | 2127 | 3260 | dnaN_1 | BCGDan_ | 1134 | 4320248 | 4321381 | dnaN_2 | BCGDan_ | 1134 |
| 3280 | 4437 | recF_1 | BCG_000 | 1158 | 32947 | 34104 | recF_2 | BCG_003 | 1158 | 3280 | 4437 | recF_1 | BCGDan_0003 | 1158 | 4400517 | 4401674 | recF_2 | BCGDan_4072 | 1158 | 3451 | 4437 | recF_1 | BCGDan_ | 987 | 4321572 | 4322558 | recF_2 | BCGDan_ | 987 |
| 4434 | 4997 | BCG_000 | BCG_000 | 564 | 34101 | 34664 | BCG_003 | BCG_003 | 564 | 4434 | 4997 | BCGDan_0004 | BCGDan_0004 | 564 | 4401671 | 4402234 | BCGDan_4073 | BCGDan_4073 | 564 | 4434 | 4997 | BCGDan_ | BCGDan_ | 564 | 4322555 | 4323118 | BCGDan_ | BCGDan_ | 564 |
| 5123 | 7267 | gyrB_1 | BCG_000 | 2145 | 34790 | 36934 | gyrB_2 | BCG_003 | 2145 | 5123 | 7267 | gyrB_1 | BCGDan_0005 | 2145 | 4402360 | 4403364 | gyrB | BCGDan_4074 | 1005 | 5123 | 7267 | gyrB_1 | BCGDan_ | 2145 | 4323244 | 4324248 | gyrB_2 | BCGDan_ | 1005 |
| ND | ND | ND | ND | ND | ND | ND | ND | ND | ND | 3500700 | 3500978 | whiB7_1 | BCGDan_3255 | 279 | 3642265 | 3642543 | whiB7_2 | BCGDan_3387 | 279 | ND | ND | ND | ND | ND | ND | ND | ND | ND | ND |
| ND | ND | ND | ND | ND | ND | ND | ND | ND | ND | 3501408 | 3503510 | UvrD2_1 | BCGDan_3256 | 2103 | 3642973 | 3645075 | UvrD2_2 | BCGDan_3388 | 2103 | ND | ND | ND | ND | ND | ND | ND | ND | ND | ND |
| ND | ND | ND | ND | ND | ND | ND | ND | ND | ND | 3503634 | 3503888 | BCGDan_3257 | BCGDan_3257 | 255 | 3645199 | 3645453 | BCGDan_3389 | BCGDan_3389 | 255 | ND | ND | ND | ND | ND | ND | ND | ND | ND | ND |
| ND | ND | ND | ND | ND | ND | ND | ND | ND | ND | 3503901 | 3504842 | nudC_1 | BCGDan_3258 | 942 | 3645466 | 3646407 | nudC_2 | BCGDan_3390 | 942 | ND | ND | ND | ND | ND | ND | ND | ND | ND | ND |
| ND | ND | ND | ND | ND | ND | ND | ND | ND | ND | 3504901 | 3505968 | BCGDan_3259 | BCGDan_3259 | 1068 | 3646466 | 3647533 | BCGDan_3391 | BCGDan_3391 | 1068 | ND | ND | ND | ND | ND | ND | ND | ND | ND | ND |
| ND | ND | ND | ND | ND | ND | ND | ND | ND | ND | 3506030 | 3509335 | BCGDan_3260 | BCGDan_3260 | 3306 | 3647595 | 3650900 | BCGDan_3392 | BCGDan_3392 | 3306 | ND | ND | ND | ND | ND | ND | ND | ND | ND | ND |
| ND | ND | ND | ND | ND | ND | ND | ND | ND | ND | 3509332 | 3512499 | BCGDan_3261 | BCGDan_3261 | 3168 | 3650897 | 3654064 | BCGDan_3393 | BCGDan_3393 | 3168 | ND | ND | ND | ND | ND | ND | ND | ND | ND | ND |
| ND | ND | ND | ND | ND | ND | ND | ND | ND | ND | 3512576 | 3512794 | BCGDan_3262 | BCGDan_3262 | 219 | 3654141 | 3654359 | BCGDan_3394 | BCGDan_3394 | 219 | ND | ND | ND | ND | ND | ND | ND | ND | ND | ND |
| ND | ND | ND | ND | ND | ND | ND | ND | ND | ND | 3512936 | 3513610 | lipV_1 | BCGDan_3263 | 675 | 3654501 | 3655175 | lipV_2 | BCGDan_3395 | 675 | ND | ND | ND | ND | ND | ND | ND | ND | ND | ND |
| ND | ND | ND | ND | ND | ND | ND | ND | ND | ND | 3513613 | 3513918 | BCGDan_3264 | BCGDan_3264 | 306 | 3655178 | 3655483 | BCGDan_3396 | BCGDan_3396 | 306 | ND | ND | ND | ND | ND | ND | ND | ND | ND | ND |
| ND | ND | ND | ND | ND | ND | ND | ND | ND | ND | 3513925 | 3514803 | BCGDan_3265 | BCGDan_3265 | 879 | 3655490 | 3656368 | BCGDan_3397 | BCGDan_3397 | 879 | ND | ND | ND | ND | ND | ND | ND | ND | ND | ND |
| ND | ND | ND | ND | ND | ND | ND | ND | ND | ND | 3514830 | 3516008 | moeB1_1 | BCGDan_3266 | 1179 | 3656395 | 3657573 | moeB1_2 | BCGDan_3398 | 1179 | ND | ND | ND | ND | ND | ND | ND | ND | ND | ND |
| ND | ND | ND | ND | ND | ND | ND | ND | ND | ND | 3516099 | 3516956 | BCGDan_3267 | BCGDan_3267 | 858 | 3657664 | 3658521 | BCGDan_3399 | BCGDan_3399 | 858 | ND | ND | ND | ND | ND | ND | ND | ND | ND | ND |
| ND | ND | ND | ND | ND | ND | ND | ND | ND | ND | 3517302 | 3517988 | BCGDan_3268 | BCGDan_3268 | 687 | 3658867 | 3659553 | BCGDan_3400 | BCGDan_3400 | 687 | ND | ND | ND | ND | ND | ND | ND | ND | ND | ND |
| ND | ND | ND | ND | ND | ND | ND | ND | ND | ND | 3517975 | 3518247 | TB9.4_1 | BCGDan_3269 | 273 | 3659540 | 3659812 | TB9.4_2 | BCGDan_3401 | 273 | ND | ND | ND | ND | ND | ND | ND | ND | ND | ND |
| ND | ND | ND | ND | ND | ND | ND | ND | ND | ND | 3518572 | 3519132 | BCGDan_3270 | BCGDan_3270 | 561 | 3660137 | 3660697 | BCGDan_3402 | BCGDan_3402 | 561 | ND | ND | ND | ND | ND | ND | ND | ND | ND | ND |
| ND | ND | ND | ND | ND | ND | ND | ND | ND | ND | 3519142 | 3519837 | BCGDan_3271 | BCGDan_3271 | 696 | 3660707 | 3661402 | BCGDan_3403 | BCGDan_3403 | 696 | ND | ND | ND | ND | ND | ND | ND | ND | ND | ND |
| ND | ND | ND | ND | ND | ND | ND | ND | ND | ND | 3520096 | 3521679 | rhlE_1 | BCGDan_3272 | 1584 | 3661661 | 3663244 | rhlE_2 | BCGDan_3404 | 1584 | ND | ND | ND | ND | ND | ND | ND | ND | ND | ND |
| ND | ND | ND | ND | ND | ND | ND | ND | ND | ND | 3521692 | 3522915 | BCGDan_3273 | BCGDan_3273 | 1224 | 3663257 | 3664480 | BCGDan_3405 | BCGDan_3405 | 1224 | ND | ND | ND | ND | ND | ND | ND | ND | ND | ND |
| 3543174 | 3543974 | BCG_324 | BCG_324 | 801 | 3642530 | 3643126 | BCG_333 | BCG_333 | 597 | 3522991 | 3523791 | BCGDan_3274 | BCGDan_3274 | 801 | 3664556 | 3665356 | BCGDan_3406 | BCGDan_3406 | 801 | ND | ND | ND | ND | ND | ND | ND | ND | ND | ND |
| 3544128 | 3544739 | gpm2_1 | BCG_324 | 612 | 3643280 | 3643891 | gpm2_2 | BCG_333 | 612 | 3523945 | 3524556 | gpm2_1 | BCGDan_3275 | 612 | 3665510 | 3666121 | gpm2_2 | BCGDan_3407 | 612 | ND | ND | ND | ND | ND | ND | ND | ND | ND | ND |
| 3544736 | 3545854 | entC_1 | BCG_324 | 1119 | 3643888 | 3645006 | entC_2 | BCG_333 | 1119 | 3524553 | 3525671 | entC_1 | BCGDan_3276 | 1119 | 3666118 | 3667236 | entC_2 | BCGDan_3408 | 1119 | ND | ND | ND | ND | ND | ND | ND | ND | ND | ND |
| 3546002 | 3546334 | BCG_324 | BCG_324 | 333 | 3645154 | 3645486 | BCG_333 | BCG_333 | 333 | 3525819 | 3526151 | BCGDan_3277 | BCGDan_3277 | 333 | 3667384 | 3667716 | BCGDan_3409 | BCGDan_3409 | 333 | ND | ND | ND | ND | ND | ND | ND | ND | ND | ND |
| 3546286 | 3546717 | BCG_324 | BCG_324 | 432 | 3645438 | 3645869 | BCG_333 | BCG_333 | 432 | 3526103 | 3526534 | BCGDan_3278 | BCGDan_3278 | 432 | 3667668 | 3668099 | BCGDan_3410 | BCGDan_3410 | 432 | ND | ND | ND | ND | ND | ND | ND | ND | ND | ND |
| 3546950 | 3547915 | BCG_324 | BCG_324 | 966 | 3646102 | 3647067 | BCG_333 | BCG_333 | 966 | 3526767 | 3527732 | BCGDan_3279 | BCGDan_3279 | 966 | 3668332 | 3669297 | BCGDan_3411 | BCGDan_3411 | 966 | ND | ND | ND | ND | ND | ND | ND | ND | ND | ND |
| 3548195 | 3548449 | whiB1_1 | BCG_324 | 255 | 3647347 | 3647601 | whiB1_2 | BCG_333 | 255 | 3528012 | 3528266 | whiB1_1 | BCGDan_3280 | 255 | 3669577 | 3669831 | whiB1_2 | BCGDan_3412 | 255 | ND | ND | ND | ND | ND | ND | ND | ND | ND | ND |
| 3548511 | 3550016 | BCG_324 | BCG_324 | 1506 | 3647663 | 3649168 | BCG_334 | BCG_334 | 1506 | 3528328 | 3529833 | BCGDan_3281 | BCGDan_3281 | 1506 | 3669893 | 3671398 | BCGDan_3413 | BCGDan_3413 | 1506 | ND | ND | ND | ND | ND | ND | ND | ND | ND | ND |
| 3550033 | 3550248 | TB7.3_1 | BCG_324 | 216 | 3649185 | 3649400 | TB7.3_2 | BCG_334 | 216 | 3529850 | 3530065 | TB7.3_1 | BCGDan_3282 | 216 | 3671415 | 3671630 | TB7.3_2 | BCGDan_3414 | 216 | ND | ND | ND | ND | ND | ND | ND | ND | ND | ND |
| 3550533 | 3550838 | BCG_324 | BCG_324 | 306 | 3649685 | 3649990 | BCG_334 | BCG_334 | 306 | 3530350 | 3530655 | BCGDan_3283 | BCGDan_3283 | 306 | 3671915 | 3672220 | BCGDan_3415 | BCGDan_3415 | 306 | ND | ND | ND | ND | ND | ND | ND | ND | ND | ND |
| 3550835 | 3551386 | BCG_325 | BCG_325 | 552 | 3649987 | 3650538 | BCG_334 | BCG_334 | 552 | 3530652 | 3531203 | BCGDan_3284 | BCGDan_3284 | 552 | 3672217 | 3672768 | BCGDan_3416 | BCGDan_3416 | 552 | ND | ND | ND | ND | ND | ND | ND | ND | ND | ND |
| 3551383 | 3552033 | sigH_1 | BCG_325 | 651 | 3650535 | 3651185 | sigH_2 | BCG_334 | 651 | 3531200 | 3531850 | sigH_1 | BCGDan_3285 | 651 | 3672765 | 3673415 | sigH_2 | BCGDan_3417 | 651 | ND | ND | ND | ND | ND | ND | ND | ND | ND | ND |
| 3552333 | 3553181 | BCG_325 | BCG_325 | 849 | 3651485 | 3652333 | BCG_334 | BCG_334 | 849 | 3532150 | 3532998 | BCGDan_3286 | BCGDan_3286 | 849 | 3673715 | 3674563 | BCGDan_3418 | BCGDan_3418 | 849 | ND | ND | ND | ND | ND | ND | ND | ND | ND | ND |
| 3553117 | 3553305 | BCG_325 | BCG_325 | 189 | 3652269 | 3652457 | BCG_334 | BCG_334 | 189 | 3532934 | 3533122 | BCGDan_3287 | BCGDan_3287 | 189 | 3674499 | 3674687 | BCGDan_3419 | BCGDan_3419 | 189 | ND | ND | ND | ND | ND | ND | ND | ND | ND | ND |
| 3553283 | 3553501 | BCG_325 | BCG_325 | 219 | 3652435 | 3652653 | BCG_334 | BCG_334 | 219 | 3533100 | 3533318 | BCGDan_3288 | BCGDan_3288 | 219 | 3674665 | 3674883 | BCGDan_3420 | BCGDan_3420 | 219 | ND | ND | ND | ND | ND | ND | ND | ND | ND | ND |
| 3553498 | 3554922 | BCG_325 | BCG_325 | 1425 | 3652650 | 3654074 | BCG_334 | BCG_334 | 1425 | 3533315 | 3534739 | BCGDan_3289 | BCGDan_3289 | 1425 | 3674880 | 3676304 | BCGDan_3421 | BCGDan_3421 | 1425 | ND | ND | ND | ND | ND | ND | ND | ND | ND | ND |
| 3555046 | 3555804 | BCG_325 | BCG_325 | 759 | 3654198 | 3654956 | BCG_334 | BCG_334 | 759 | 3534863 | 3535621 | BCGDan_3290 | BCGDan_3290 | 759 | 3676428 | 3677186 | BCGDan_3422 | BCGDan_3422 | 759 | ND | ND | ND | ND | ND | ND | ND | ND | ND | ND |
| 3555859 | 3557211 | aroA_1 | BCG_325 | 1353 | 3655011 | 3656363 | aroA_2 | BCG_335 | 1353 | 3535676 | 3537028 | aroA_1 | BCGDan_3291 | 1353 | 3677241 | 3678593 | aroA_2 | BCGDan_3423 | 1353 | ND | ND | ND | ND | ND | ND | ND | ND | ND | ND |
| 3557208 | 3558200 | BCG_325 | BCG_325 | 993 | 3656360 | 3657352 | BCG_335 | BCG_335 | 993 | 3537025 | 3538017 | BCGDan_3292 | BCGDan_3292 | 993 | 3678590 | 3679582 | BCGDan_3424 | BCGDan_3424 | 993 | ND | ND | ND | ND | ND | ND | ND | ND | ND | ND |
| 3558233 | 3559516 | desA3_1 | BCG_325 | 1284 | 3657385 | 3658668 | desA3_2 | BCG_335 | 1284 | 3538050 | 3539333 | desA3_1 | BCGDan_3293 | 1284 | 3679615 | 3680898 | desA3_2 | BCGDan_3425 | 1284 | ND | ND | ND | ND | ND | ND | ND | ND | ND | ND |
| 3559594 | 3560736 | BCG_326 | BCG_326 | 1143 | 3658746 | 3659888 | BCG_335 | BCG_335 | 1143 | 3539411 | 3540553 | BCGDan_3294 | BCGDan_3294 | 1143 | 3680976 | 3682118 | BCGDan_3426 | BCGDan_3426 | 1143 | ND | ND | ND | ND | ND | ND | ND | ND | ND | ND |
| 3560846 | 3561355 | BCG_326 | BCG_326 | 510 | 3659998 | 3660108 | BCG_335 | BCG_335 | 111 | 3540663 | 3541172 | BCGDan_3295 | BCGDan_3295 | 510 | 3682228 | 3682338 | BCGDan_3428 | BCGDan_3428 | 111 | ND | ND | ND | ND | ND | ND | ND | ND | ND | ND |
| 3622853 | 3624202 | lat | BCG_331 | 1350 | 3659943 | 3660365 | lat' | BCG_335 | 423 | 3602670 | 3604019 | lat_1 | BCGDan_3353 | 1350 | 3682173 | 3682595 | lat_2 | BCGDan_3427 | 423 | ND | ND | ND | ND | ND | ND | ND | ND | ND | ND |
| 3624253 | 3624705 | BCG_332 | BCG_332 | 453 | 3660416 | 3660868 | BCG_335 | BCG_335 | 453 | 3604070 | 3604522 | BCGDan_3354 | BCGDan_3354 | 453 | 3682646 | 3683098 | BCGDan_3429 | BCGDan_3429 | 453 | ND | ND | ND | ND | ND | ND | ND | ND | ND | ND |
| 3624736 | 3625983 | BCG_332 | BCG_332 | 1248 | 3660899 | 3662146 | BCG_335 | BCG_335 | 1248 | 3604553 | 3605800 | BCGDan_3355 | BCGDan_3355 | 1248 | 3683129 | 3684376 | BCGDan_3430 | BCGDan_3430 | 1248 | ND | ND | ND | ND | ND | ND | ND | ND | ND | ND |
| 3626010 | 3627494 | pcd_1 | BCG_332 | 1485 | 3662173 | 3663657 | pcd_2 | BCG_335 | 1485 | 3605827 | 3607311 | pcd_1 | BCGDan_3356 | 1485 | 3684403 | 3685887 | pcd_2 | BCGDan_3431 | 1485 | ND | ND | ND | ND | ND | ND | ND | ND | ND | ND |
| 3627594 | 3628403 | BCG_332 | BCG_332 | 810 | 3663757 | 3664566 | BCG_335 | BCG_335 | 810 | 3607411 | 3608220 | BCGDan_3357 | BCGDan_3357 | 810 | 3685987 | 3686796 | BCGDan_3432 | BCGDan_3432 | 810 | ND | ND | ND | ND | ND | ND | ND | ND | ND | ND |
| 3628474 | 3629139 | BCG_332 | BCG_332 | 666 | 3664637 | 3665302 | BCG_336 | BCG_336 | 666 | 3608291 | 3608956 | BCGDan_3358 | BCGDan_3358 | 666 | 3686867 | 3687532 | BCGDan_3433 | BCGDan_3433 | 666 | ND | ND | ND | ND | ND | ND | ND | ND | ND | ND |
| 3629183 | 3633724 | lhr_1 | BCG_332 | 4542 | 3665346 | 3669887 | lhr_2 | BCG_336 | 4542 | 3609000 | 3613541 | lhr_1 | BCGDan_3359 | 4542 | 3687576 | 3692117 | lhr_2 | BCGDan_3434 | 4542 | ND | ND | ND | ND | ND | ND | ND | ND | ND | ND |
| 3633728 | 3634495 | nei_1 | BCG_332 | 768 | 3669891 | 3670658 | nei_2 | BCG_336 | 768 | 3613545 | 3614312 | nei_1 | BCGDan_3360 | 768 | 3692121 | 3692888 | nei_2 | BCGDan_3435 | 768 | ND | ND | ND | ND | ND | ND | ND | ND | ND | ND |
| 3634518 | 3635432 | lpqC_1 | BCG_332 | 915 | 3670681 | 3671595 | lpqC_2 | BCG_336 | 915 | 3614335 | 3615249 | lpqC_1 | BCGDan_3361 | 915 | 3692911 | 3693825 | lpqC_2 | BCGDan_3436 | 915 | ND | ND | ND | ND | ND | ND | ND | ND | ND | ND |
| 3635459 | 3638371 | atsB | BCG_332 | 2913 | 3671622 | 3674534 | atsB_2 | BCG_336 | 2913 | 3615276 | 3618188 | atsB_1 | BCGDan_3362 | 2913 | 3693852 | 3696764 | atsB_2 | BCGDan_3437 | 2913 | ND | ND | ND | ND | ND | ND | ND | ND | ND | ND |
| 3638391 | 3639308 | BCG_332 | BCG_332 | 918 | 3674554 | 3675471 | BCG_336 | BCG_336 | 918 | 3618208 | 3619125 | BCGDan_3363 | BCGDan_3363 | 918 | 3696784 | 3697701 | BCGDan_3438 | BCGDan_3438 | 918 | ND | ND | ND | ND | ND | ND | ND | ND | ND | ND |
| 3639320 | 3639985 | phoY1_1 | BCG_333 | 666 | 3675483 | 3676148 | phoY1_2 | BCG_336 | 666 | 3619137 | 3619802 | phoY1_1 | BCGDan_3364 | 666 | 3697713 | 3698378 | phoY1_2 | BCGDan_3439 | 666 | ND | ND | ND | ND | ND | ND | ND | ND | ND | ND |

**Legend**

| | |
|---|---|
| (light blue) | DU1-Paste |
| (light orange) | DU2-IV |
| (blue) | DU1-Danis |
| (orange) | DU2-III |
| ND | not duplica |

| | | | | | | | | | | | | | | | | | | | | | | | | | | | | | |
|---|---|---|---|---|---|---|---|---|---|---|---|---|---|---|---|---|---|---|---|---|---|---|---|---|---|---|---|---|---|
| 3640093 | 3641850 | glpD2_1 | BCG_333 | 1758 | 3676256 | 3678013 | glpD2_2 | BCG_336 | 1758 | 3619910 | 3621667 | glpD2_1 | BCGDan_3365 | 1758 | 3698486 | 3700243 | glpD2_2 | BCGDan_3440 | 1758 | ND | ND | ND | ND | ND | ND | ND | ND | ND | ND |
| 3678094 | 3679509 | lpdA | BCG_336 | 1416 | 3641931 | 3642536 | 'lpdA | BCG_333 | 606 | 3621748 | 3623163 | lpdA_1 | BCGDan_3366 | 1416 | 3700324 | 3701739 | lpdA_2 | BCGDan_3441 | 1416 | ND | ND | ND | ND | ND | ND | ND | ND | ND | ND |
| ND | ND | ND | ND | ND | ND | ND | ND | ND | ND | 3623366 | 3623845 | BCGDan_3367 | BCGDan_3367 | 480 | 3701942 | 3702421 | BCGDan_3442 | BCGDan_3442 | 480 | ND | ND | ND | ND | ND | ND | ND | ND | ND | ND |
| ND | ND | ND | ND | ND | ND | ND | ND | ND | ND | 3623864 | 3625033 | amiA1_1 | BCGDan_3368 | 1170 | 3702440 | 3703609 | amiA1_2 | BCGDan_3443 | 1170 | ND | ND | ND | ND | ND | ND | ND | ND | ND | ND |
| ND | ND | ND | ND | ND | ND | ND | ND | ND | ND | 3625030 | 3626214 | amiB1_1 | BCGDan_3369 | 1185 | 3703606 | 3704790 | amiB1_2 | BCGDan_3444 | 1185 | ND | ND | ND | ND | ND | ND | ND | ND | ND | ND |
| ND | ND | ND | ND | ND | ND | ND | ND | ND | ND | 3626279 | 3627085 | deoD_1 | BCGDan_3370 | 807 | 3704855 | 3705661 | deoD_2 | BCGDan_3445 | 807 | ND | ND | ND | ND | ND | ND | ND | ND | ND | ND |
| ND | ND | ND | ND | ND | ND | ND | ND | ND | ND | 3627089 | 3628693 | pmmB_1 | BCGDan_3371 | 1605 | 3705665 | 3707269 | pmmB_2 | BCGDan_3446 | 1605 | ND | ND | ND | ND | ND | ND | ND | ND | ND | ND |
| ND | ND | ND | ND | ND | ND | ND | ND | ND | ND | 3628695 | 3629318 | upp_1 | BCGDan_3372 | 624 | 3707271 | 3707894 | upp_2 | BCGDan_3447 | 624 | ND | ND | ND | ND | ND | ND | ND | ND | ND | ND |
| ND | ND | ND | ND | ND | ND | ND | ND | ND | ND | 3629423 | 3630322 | sapM_1 | BCGDan_3373 | 900 | 3707999 | 3708898 | sapM_2 | BCGDan_3448 | 900 | ND | ND | ND | ND | ND | ND | ND | ND | ND | ND |
| ND | ND | ND | ND | ND | ND | ND | ND | ND | ND | 3630346 | 3631608 | BCGDan_3374 | BCGDan_3374 | 1263 | 3708922 | 3710184 | BCGDan_3449 | BCGDan_3449 | 1263 | ND | ND | ND | ND | ND | ND | ND | ND | ND | ND |
| ND | ND | ND | ND | ND | ND | ND | ND | ND | ND | 3631629 | 3632555 | BCGDan_3375 | BCGDan_3375 | 927 | 3710205 | 3711131 | BCGDan_3450 | BCGDan_3450 | 927 | ND | ND | ND | ND | ND | ND | ND | ND | ND | ND |
| ND | ND | ND | ND | ND | ND | ND | ND | ND | ND | 3632930 | 3633241 | BCGDan_3376 | BCGDan_3376 | 312 | 3711506 | 3711817 | BCGDan_3451 | BCGDan_3451 | 312 | ND | ND | ND | ND | ND | ND | ND | ND | ND | ND |
| ND | ND | ND | ND | ND | ND | ND | ND | ND | ND | 3633312 | 3634409 | add_1 | BCGDan_3377 | 1098 | 3711888 | 3712985 | add_2 | BCGDan_3452 | 1098 | ND | ND | ND | ND | ND | ND | ND | ND | ND | ND |
| ND | ND | ND | ND | ND | ND | ND | ND | ND | ND | 3634409 | 3635692 | deoA_1 | BCGDan_3378 | 1284 | 3712985 | 3714268 | deoA_2 | BCGDan_3453 | 1284 | ND | ND | ND | ND | ND | ND | ND | ND | ND | ND |
| ND | ND | ND | ND | ND | ND | ND | ND | ND | ND | 3635689 | 3636090 | cdd_1 | BCGDan_3379 | 402 | 3714265 | 3714666 | cdd_2 | BCGDan_3454 | 402 | ND | ND | ND | ND | ND | ND | ND | ND | ND | ND |
| ND | ND | ND | ND | ND | ND | ND | ND | ND | ND | 3636327 | 3636665 | sdhC_1 | BCGDan_3380 | 339 | 3714903 | 3715241 | sdhC_2 | BCGDan_3455 | 339 | ND | ND | ND | ND | ND | ND | ND | ND | ND | ND |
| ND | ND | ND | ND | ND | ND | ND | ND | ND | ND | 3636662 | 3637096 | sdhD_1 | BCGDan_3381 | 435 | 3715238 | 3715672 | sdhD_2 | BCGDan_3456 | 435 | ND | ND | ND | ND | ND | ND | ND | ND | ND | ND |
| ND | ND | ND | ND | ND | ND | ND | ND | ND | ND | 3637225 | 3638997 | sdhA_1 | BCGDan_3382 | 1773 | 3715801 | 3717573 | sdhA_2 | BCGDan_3457 | 1773 | ND | ND | ND | ND | ND | ND | ND | ND | ND | ND |
| ND | ND | ND | ND | ND | ND | ND | ND | ND | ND | 3638997 | 3639788 | sdhB_1 | BCGDan_3383 | 792 | 3717573 | 3718364 | sdhB_2 | BCGDan_3458 | 792 | ND | ND | ND | ND | ND | ND | ND | ND | ND | ND |
| ND | ND | ND | ND | ND | ND | ND | ND | ND | ND | 3639867 | 3640295 | BCGDan_3384 | BCGDan_3384 | 429 | 3718443 | 3718871 | BCGDan_3459 | BCGDan_3459 | 429 | ND | ND | ND | ND | ND | ND | ND | ND | ND | ND |
| ND | ND | ND | ND | ND | ND | ND | ND | ND | ND | 3640299 | 3640541 | BCGDan_3385 | BCGDan_3385 | 243 | 3718875 | 3719117 | BCGDan_3460 | BCGDan_3460 | 243 | ND | ND | ND | ND | ND | ND | ND | ND | ND | ND |
| ND | ND | ND | ND | ND | ND | ND | ND | ND | ND | 3640663 | 3641277 | BCGDan_3386 | BCGDan_3386 | 615 | 3719239 | 3719853 | BCGDan_3461 | BCGDan_3461 | 615 | ND | ND | ND | ND | ND | ND | ND | ND | ND | ND |
| 21327 | 22547 | BCG_002 | BCG_002 | 1221 | 4366182 | 4367402 | BCG_397 | BCG_397 | 1221 | 4388897 | 4390117 | BCGDan_4060 | BCGDan_4060 | 1221 | 4403474 | 4404694 | BCGDan_4075 | BCGDan_4075 | 1221 | 4309781 | 4311001 | BCGDan_ | BCGDan_ | 1221 | 4324358 | 4325578 | BCGDan_ | BCGDan_ | 1221 |
| 22568 | 23302 | BCG_002 | BCG_002 | 735 | 4367423 | 4368157 | BCG_397 | BCG_397 | 735 | 4390138 | 4390872 | BCGDan_4061 | BCGDan_4061 | 735 | 4404715 | 4405449 | BCGDan_4076 | BCGDan_4076 | 735 | 4311022 | 4311756 | BCGDan_ | BCGDan_ | 735 | 4325599 | 4326333 | BCGDan_ | BCGDan_ | 735 |
| 23592 | 24626 | parB_2 | BCG_002 | 1035 | 4368447 | 4369481 | parB_1 | BCG_397 | 1035 | 4391162 | 4392196 | parB_1 | BCGDan_4062 | 1035 | 4405739 | 4406773 | parB_2 | BCGDan_4077 | 1035 | 4312046 | 4313080 | parB_1 | BCGDan_ | 1035 | 4326623 | 4327657 | parB_2 | BCGDan_ | 1035 |
| 24623 | 25666 | parA_2 | BCG_002 | 1044 | 4369478 | 4370521 | parA_1 | BCG_397 | 1044 | 4392193 | 4393236 | parA_1 | BCGDan_4063 | 1044 | 4406770 | 4407813 | parA_2 | BCGDan_4078 | 1044 | 4313077 | 4313985 | parA_1 | BCGDan_ | 909 | 4327654 | 4328562 | parA_2 | BCGDan_ | 909 |
| 25663 | 26337 | gid_2 | BCG_002 | 675 | 4370518 | 4371192 | gid_1 | BCG_397 | 675 | 4393233 | 4393907 | gid_1 | BCGDan_4064 | 675 | 4407810 | 4408484 | gid_2 | BCGDan_4079 | 675 | 4314117 | 4314791 | gid_1 | BCGDan_ | 675 | 4328694 | 4329368 | gid_2 | BCGDan_ | 675 |
| 26469 | 27032 | BCG_002 | BCG_002 | 564 | 4371324 | 4371887 | BCG_397 | BCG_397 | 564 | 4394039 | 4394602 | BCGDan_4065 | BCGDan_4065 | 564 | 4408616 | 4409179 | BCGDan_4080 | BCGDan_4080 | 564 | 4314923 | 4315486 | BCGDan_ | BCGDan_ | 564 | 4329500 | 4330063 | BCGDan_ | BCGDan_ | 564 |
| 27104 | 28204 | BCG_002 | BCG_002 | 1101 | 4371959 | 4373059 | BCG_397 | BCG_397 | 1101 | 4394674 | 4395774 | BCGDan_4066 | BCGDan_4066 | 1101 | 4409251 | 4410351 | BCGDan_4081 | BCGDan_4081 | 1101 | 4315558 | 4316658 | BCGDan_ | BCGDan_ | 1101 | 4330135 | 4331235 | BCGDan_ | BCGDan_ | 1101 |
| 28188 | 28550 | BCG_002 | BCG_002 | 363 | 4373043 | 4373405 | BCG_398 | BCG_398 | 363 | 4395758 | 4396120 | BCGDan_4067 | BCGDan_4067 | 363 | 4410335 | 4410697 | BCGDan_4082 | BCGDan_4082 | 363 | 4316642 | 4317004 | BCGDan_ | BCGDan_ | 363 | 4331219 | 4331581 | BCGDan_ | BCGDan_ | 363 |
| 28547 | 28924 | rnpA_2 | BCG_002 | 378 | 4373402 | 4373779 | rnpA_1 | BCG_398 | 378 | 4396117 | 4396494 | rnpA_1 | BCGDan_4068 | 378 | 4410694 | 4411071 | rnpA_2 | BCGDan_4083 | 378 | 4317001 | 4317267 | rnpA_1 | BCGDan_ | 267 | 4331578 | 4331955 | rnpA_2 | BCGDan_ | 378 |
| 28921 | 29064 | rpmH_2 | BCG_003 | 144 | 4373776 | 4373919 | rpmH_1 | BCG_398 | 144 | 4396491 | 4396634 | rpmH_1 | BCGDan_4069 | 144 | 4411068 | 4411211 | rpmH_2 | BCGDan_4084 | 144 | 4317375 | 4317518 | rpmH_1 | BCGDan_ | 144 | 4331952 | 4332095 | rpmH_2 | BCGDan_ | 144 |
| 7302 | 9818 | gyrA_1 | BCG_000 | 2517 | 36969 | 39485 | gyrA_2 | BCG_003 | 2517 | ND | ND | ND | ND | ND | ND | ND | ND | ND | ND | ND | ND | ND | ND | ND | ND | ND | ND | ND | ND |
| 9914 | 10828 | BCG_000 | BCG_000 | 915 | 39581 | 40495 | BCG_003 | BCG_003 | 915 | ND | ND | ND | ND | ND | ND | ND | ND | ND | ND | ND | ND | ND | ND | ND | ND | ND | ND | ND | ND |
| 11874 | 12311 | BCG_000 | BCG_000 | 438 | 41541 | 41978 | BCG_003 | BCG_003 | 438 | ND | ND | ND | ND | ND | ND | ND | ND | ND | ND | ND | ND | ND | ND | ND | ND | ND | ND | ND | ND |
| 12468 | 13016 | ppiA_1 | BCG_000 | 549 | 42135 | 42683 | ppiA_2 | BCG_003 | 549 | ND | ND | ND | ND | ND | ND | ND | ND | ND | ND | ND | ND | ND | ND | ND | ND | ND | ND | ND | ND |
| 13222 | 13557 | BCG_001 | BCG_001 | 336 | 42889 | 43224 | BCG_004 | BCG_004 | 336 | ND | ND | ND | ND | ND | ND | ND | ND | ND | ND | ND | ND | ND | ND | ND | ND | ND | ND | ND | ND |
| 13713 | 13994 | BCG_001 | BCG_001 | 282 | 43380 | 43661 | BCG_004 | BCG_004 | 282 | ND | ND | ND | ND | ND | ND | ND | ND | ND | ND | ND | ND | ND | ND | ND | ND | ND | ND | ND | ND |
| 14088 | 14876 | BCG_001 | BCG_001 | 789 | 43755 | 44543 | BCG_004 | BCG_004 | 789 | ND | ND | ND | ND | ND | ND | ND | ND | ND | ND | ND | ND | ND | ND | ND | ND | ND | ND | ND | ND |
| 14913 | 15611 | trpG_1 | BCG_001 | 699 | 44580 | 45278 | trpG_2 | BCG_004 | 699 | ND | ND | ND | ND | ND | ND | ND | ND | ND | ND | ND | ND | ND | ND | ND | ND | ND | ND | ND | ND |
| 15589 | 16734 | 'pknB | BCG_001 | 1146 | 45256 | 47136 | pknB | BCG_004 | 1881 | ND | ND | ND | ND | ND | ND | ND | ND | ND | ND | ND | ND | ND | ND | ND | ND | ND | ND | ND | ND |
| 16731 | 18287 | BCG_001 | BCG_001 | 1557 | 4359588 | 4363142 | BCG_396 | BCG_396 | 3555 | ND | ND | ND | ND | ND | ND | ND | ND | ND | ND | ND | ND | ND | ND | ND | ND | ND | ND | ND | ND |
| 18322 | 18912 | sigMa_2 | BCG_001 | 591 | 4363177 | 4363767 | sigMa_1 | BCG_396 | 591 | ND | ND | ND | ND | ND | ND | ND | ND | ND | ND | ND | ND | ND | ND | ND | ND | ND | ND | ND | ND |
| 18801 | 18989 | sigMb_2 | BCG_001 | 189 | 4363656 | 4363844 | sigMb_1 | BCG_396 | 189 | ND | ND | ND | ND | ND | ND | ND | ND | ND | ND | ND | ND | ND | ND | ND | ND | ND | ND | ND | ND |
| 19005 | 19769 | BCG_001 | BCG_001 | 765 | 4363860 | 4364624 | BCG_397 | BCG_397 | 765 | ND | ND | ND | ND | ND | ND | ND | ND | ND | ND | ND | ND | ND | ND | ND | ND | ND | ND | ND | ND |
| 19863 | 20870 | trxB2_2 | BCG_001 | 1008 | 4364718 | 4365725 | trxB2_1 | BCG_397 | 1008 | ND | ND | ND | ND | ND | ND | ND | ND | ND | ND | ND | ND | ND | ND | ND | ND | ND | ND | ND | ND |
| 20867 | 21217 | trxC_2 | BCG_002 | 351 | 4365722 | 4366072 | trxC_1 | BCG_397 | 351 | ND | ND | ND | ND | ND | ND | ND | ND | ND | ND | ND | ND | ND | ND | ND | ND | ND | ND | ND | ND |
